# Lonafarnib Partially Reverses Cardiac Senescence in Human and Mouse Progeria Models via Autophagy Activation

**DOI:** 10.1101/2025.06.27.661680

**Authors:** Luís M. Monteiro, Patrícia R. Pitrez, Simão Correia-Santos, Nicola Dark, Deolinda Santinha, Célia Aveleira, Megan Townsend, James Taylor, Xavier Nissan, Vilma Sardão, Luísa Cortes, Richard Mitter, Antonio Marino, Carlos Sousa-Soares, Rui Ribeiro, James C. Smith, Elsa Logarinho, Alessandro Ori, Andreia S. Bernardo, Lino Ferreira

## Abstract

Hutchinson-Gilford Progeria Syndrome (HGPS), characterised by accelerated ageing, causes cardiovascular defects resembling aspects of cardiovascular ageing. We used human left ventricle cardiomyocytes (CMs) derived from HGPS-induced pluripotent stem cells (iPSCs), and their isogenic-corrected controls, to investigate HGPS-CM dysfunction and identify potential therapies. Our results revealed that HGPS-iPSC-CMs exhibit greater maturity and associated elevated oxidative stress compared to controls, which they could not contend with, leading to cellular senescence. Increased senescence was also observed in cardiac tissue from mouse and human physiologically-aged and HGPS individuals. Functionally, HGPS-iPSC-CMs showed dysregulated mitochondrial respiration and calcium handling. Amongst the six drugs tested, rapamycin and lonafarnib were the most effective against HGPS-cardiac phenotypes. Although lonafarnib raised safety concerns, it partially reverted the cardiac senescent phenotype by inducing cellular autophagy and decreasing progerin expression in progeroid mice. Our study supports the use of HGPS-iPSC-CMs to identify novel biomarkers and therapies for HGPS, and potentially cardiac physiological-ageing.

## INTRODUCTION

Hutchinson-Gilford progeria syndrome (HGPS) is a rare incurable and fatal genetic disorder commonly caused by a heterozygous *de novo* point mutation (c.1824C>T; p.G608G) in the *LMNA* gene, which encodes for lamin A and C [1–3]. This mutation results in the formation of an abnormal form of the lamin A protein called progerin. Alterations in the vasculature and heart compartments are known phenotypes of the disease [3–6]. Most children affected die prematurely at an average age of 14.6 years due to myocardial infarction, heart failure, or stroke [3]. Lonafarnib, the gold-standard treatment and the only FDA-approved drug for HGPS, can extend the life expectancy of patients by a few years but remains far from being a cure [7, 8]. Other drugs currently in pre-clinical and clinical trials include: rapamycin and analogs [9]; MG-132 [10]; and NPY [11]). Yet, the mechanism of action and efficacy of these drugs in heart cells, particularly in CMs, remains elusive.

Heart function is used as a primary endpoint in HGPS clinical trials for the efficacy of new therapeutic drugs [8]. HGPS patients share many features of cardiac physiological ageing including left ventricular (LV) diastolic dysfunction, repolarization abnormalities, and prominent mislocalization of connexin43 (Cx43) in the LV [5, 6, 12]. Unfortunately, the molecular and cellular mechanisms by which progerin expression causes these alterations remain largely unknown. In addition, the effect of pharmacological interventions directly on human HGPS-CMs remains to be investigated.

Human iPSCs (hiPSCs) offer an unlimited source of HGPS cells and the opportunity to model HGPS in a human background [13–20]. HGPS-iPSCs have been used to model smooth muscle cells and endothelial cells, demonstrating that HGPS promotes a dysfunctional and senescent phenotype, similar to what has been observed in progeroid animals [14, 15]. However, it is still unclear if a similar phenotype is present in HGPS-CMs.

Cellular senescence is defined as an irreversible loss of division potential on mitotic cells, in many cases with the appearance of a senescence-associated secretory phenotype (SASP) [21]. Cell senescence has been described in rarely dividing/post-mitotic tissues, such as the heart, and can be induced by mitochondrial dysfunction [22].

In this study, we investigated whether cardiac senescence is present in CMs from HGPS patients and studied pharmacological interventions to revert their phenotype. We used a protocol to differentiate HGPS-iPSCs into left ventricle CMs and showed these cells accumulate progerin, in contrast with control CMs generated from a wild type isogenic cell line. Next, we investigated the molecular biology and cellular function of HGPS-iPSC-CMs, and compared their protein expression to that of mouse hearts (progeroid mice and aged WT mice) and human heart samples (HGPS or aged individuals). Finally, we studied the effect of chronic low-dose treatment with drugs relevant to physiological ageing, senescence or which are being tested in HGPS preclinical/clinical studies. The results herein presented elucidate molecular mechanisms underlying HGPS and physiological cardiac ageing and are a proof-of-concept that HGPS-iPSC-CMs can serve as a model for testing and identifying therapeutic targets.

## RESULTS

### HGPS-iPSCs were successfully differentiated into LV-CMs expressing progerin

CM immaturity has been highlighted as a limitation in a previous HGPS-iPSCs-CM study [13]. To circumvent this limitation, here we used a protocol leading to the generation of near-homogenous LV-CM populations with increased maturity [23]. HGPS-iPSCs, previously described in [20], and a newly generated wild type isogenic control (ISO-iPSCs) (**Fig. 1a**), were differentiated into LV-CMs (HGPS-iPSC-CMs and ISO-iPSCs-CMs, respectively) using the aforementioned protocol [23] (**Fig. 1b**). To generate the ISO-iPSCs controls we used CRISPR/Cas9 technology in combination with a repair template containing the WT sequence for exon 11 of *LMNA* (**Extended Data Fig. 1**). Both ISO- and HGPS-iPSCs expressed pluripotent markers prior to differentiation and were karyotypically normal.

**Figure 1-.**
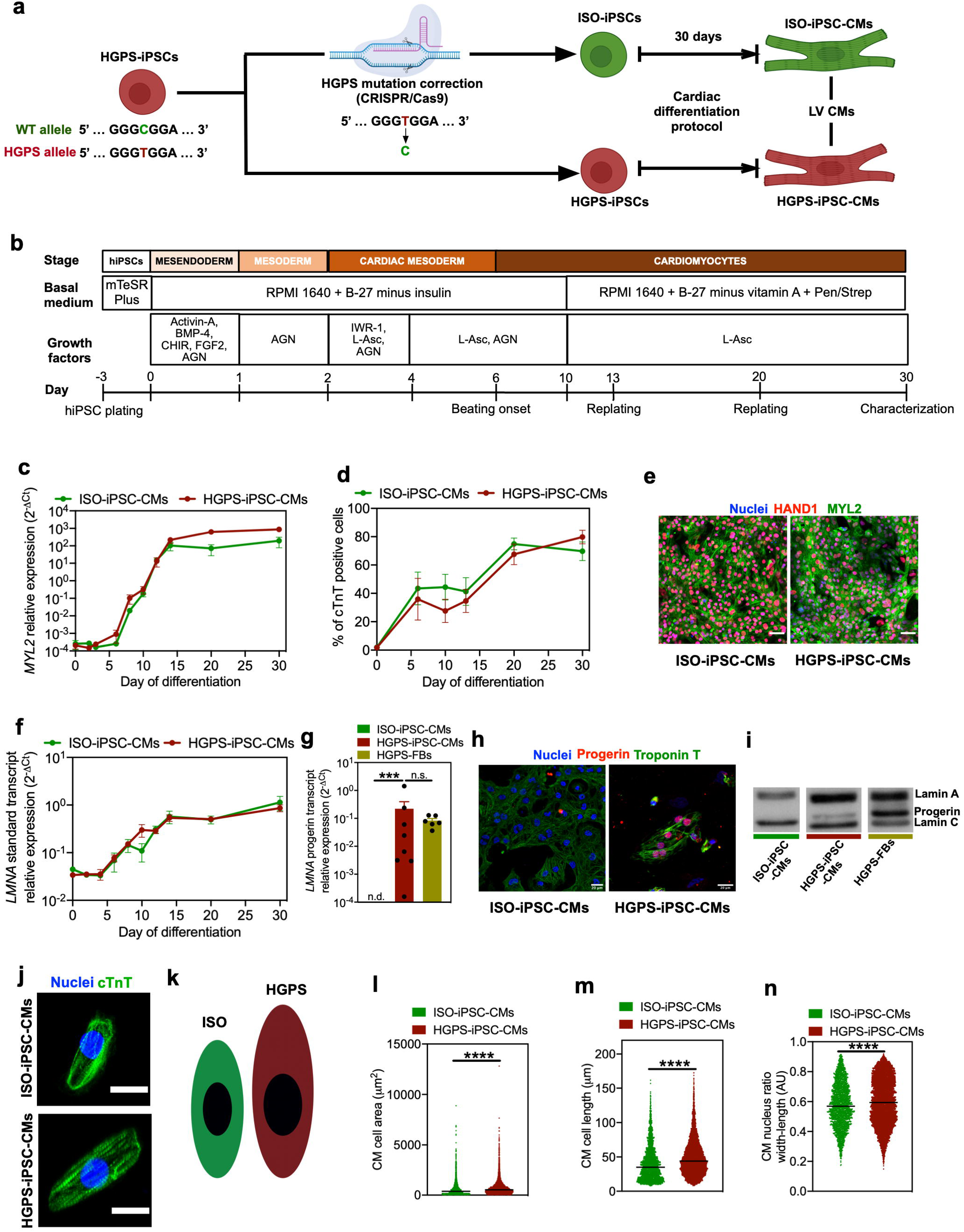
Differentiation of HGPS-iPSCs into LV CMs expressing progerin. **(a)** Scheme summarizing the process of correction of the HGPS heterozygous mutation in HGPS-iPSCs, obtaining a corrected isogenic control line (ISO-iPSCs), and differentiation of both iPSC lines into LV CMs, resorting to a 30-day protocol. **(b)** Cardiac differentiation protocol used. **(c)** Expression of *MYL2* mRNA transcripts quantified by RT-qPCR and normalized by *GAPDH*, throughout the differentiation period (n=3 independent differentiation batches). **(d)** Percentage of cardiac troponin T (cTnT) positive cells quantified by flow cytometry (for ISO-iPSC-CMs: n=3-13 independent differentiation batches per each day; for HGPS-iPSC-CMs: n= 3-11 independent differentiation batches per each day). **(e)** Co-expression of HAND1 and MYL2 in ISO- and HGPS-iPSC-CMs as evaluated by immunofluorescence. Scale bar: 50 μm. **(f)** Expression of *LMNA* standard mRNA transcript quantified by RT-qPCR and normalized by *GAPDH*, throughout the differentiation period. (n=3 independent differentiation batches). **(g)** *LMNA* progerin mRNA transcript expression quantified by TaqMan RT-qPCR, and normalized by *GAPDH*, in ISO-, HGPS-iPSC-CMs, and HGPS-FBs (n=6-8 independent experiments). n.d. - not detected. **(h)** Representative progerin immunostaining in ISO- and HGPS-iPSC-CMs. Scale bar: 20 μm. **(i)** Expression of lamin C (65 kDa), progerin (69 kDa) and lamin A (75 kDa) proteins from the nuclear fraction of ISO-iPSC-CMs, HGPS-iPSC-CMs and HGPS-FBs by western blot analyses. **(j)** Representative confocal fluorescence images of individual CMs. cTnT: cardiac troponin T. Scale bar: 20 μm. **(k)** Scheme representative of the size and shape of ISO-iPSC-CMs and HGPS-iPSC-CMs, in scale, taking into account the measurement of several parameters via the Opera Phenix system. More than 2500 cells were analyzed per experimental group. **(l)** Cell area quantification of individual CMs, resorting to MYL2 staining. **(m)** Cell length quantification of individual CMs, resorting to MYL2 staining. **(n)** Determination of the ratio between the length and width of the nuclei of individual CMs, resorting to MYL2 and DAPI stainings. In m, n and o, values are represented as the median. In all the other figures, the values presented are mean ± SEM. n.s. - not significant, *** - *p <* 0.001; **** - *p <* 0.0001.

Throughout the differentiation, there were no significant differences between ISO-iPSC-CMs and HGPS-iPSC-CMs regarding their gene expression pattern or differentiation efficiency, showing that the HGPS genotype did not perturb the cardiac differentiation process. Specifically, both cell lines exhibited an early transient expression of *TBXT* (Brachyury), a sharp increase and plateau in *GATA4* and *TBX5* expression, and a gradual and late increase in *TNNT2* and *MYL2* expression (**Extended Data Figs. 2a-2d, Fig. 1c**), suggesting that cells followed a typical ventricular differentiation path [23, 24]. Importantly, day 30 ISO- and HGPS-iPSC-CMs co-express, at the protein level, HAND1 and MYL2, confirming their LV phenotype [23, 25] (**Fig. 1e**). No statistical difference was observed in cardiac troponin T (cTnT) expression in ISO-iPSC-CMs and HGPS-iPSC-CMs, at day 30 (**Fig. 1d**). A small subset of fibroblasts and endothelial cells were also present in these cultures (**Extended Data Figs. 2e-2f**).

HGPS-iPSC-CMs show an accumulation of progerin relative to ISO-iPSC-CMs. The *LMNA* standard transcript increased gradually over time in both cell lines (**Fig. 1f**), in agreement with previous reports [26]. On day 30 of differentiation, progerin expression was detected at transcript (**Fig. 1g**) and protein levels (**Figs. 1h and i**) in HGPS-iPSC-CMs. Western blot analysis (**Fig. 1i**) further confirmed that only HGPS-iPSC-CMs express progerin, albeit at lower levels than HGPS fibroblasts (HGPS-FBs). Moreover, ISO-iPSC-CMs and HGPS-iPSC-CMs exhibited significant differences in cellular and nuclear morphology. HGPS-iPSC-CMs showed increased cellular area and anisotropy (**Fig.1j-n, and Extended Data Figs. 2g-j**) as well as vast nuclear area and geometry changes (**Extended Data Figs. 2k-2o**) relative to the isogenic counterparts. Of note, we observed two subpopulations of HGPS-iPSC-CMs, one with a wider nucleus than the other, suggesting some of the cells had nuclear deformities (**Extended Data Figs. 2n**).

Taken together, HGPS-iPSCs differentiate seemingly normally towards CMs, express progerin and hereafter, all presented results are from day 30 ISO- and HGPS-iPSC-CMs.

### HGPS-iPSCs-CMs show increased maturation, hypertrophy and senescence

To investigate in more depth the differences between HGPS-iPSC-CMs and ISO-iPSC-CMs, we performed single-cell RNASeq (**Fig. 2a**). We identified 12 distinct gene clusters (numbered from 0 to 11) in our dataset (**Fig. 2b**) some of which appeared to be over-represented in ISO- or HGPS-iPSC-CMs (**Figs. 2c-d** and **Extended Data Fig. 3a**). We identified ISO- or HGPS-iPSC-CMs enriched clusters by determining the percentage of cells in each cluster per cell line and calculating the relative cell number per cluster (**Fig. 2e**). Clusters 2 and 10 were enriched in ISO-iPSC-CMs while clusters 6, 7 and 11 were enriched in HGPS-iPSC-CMs (**Figs. 2e**).

**Figure 2-.**
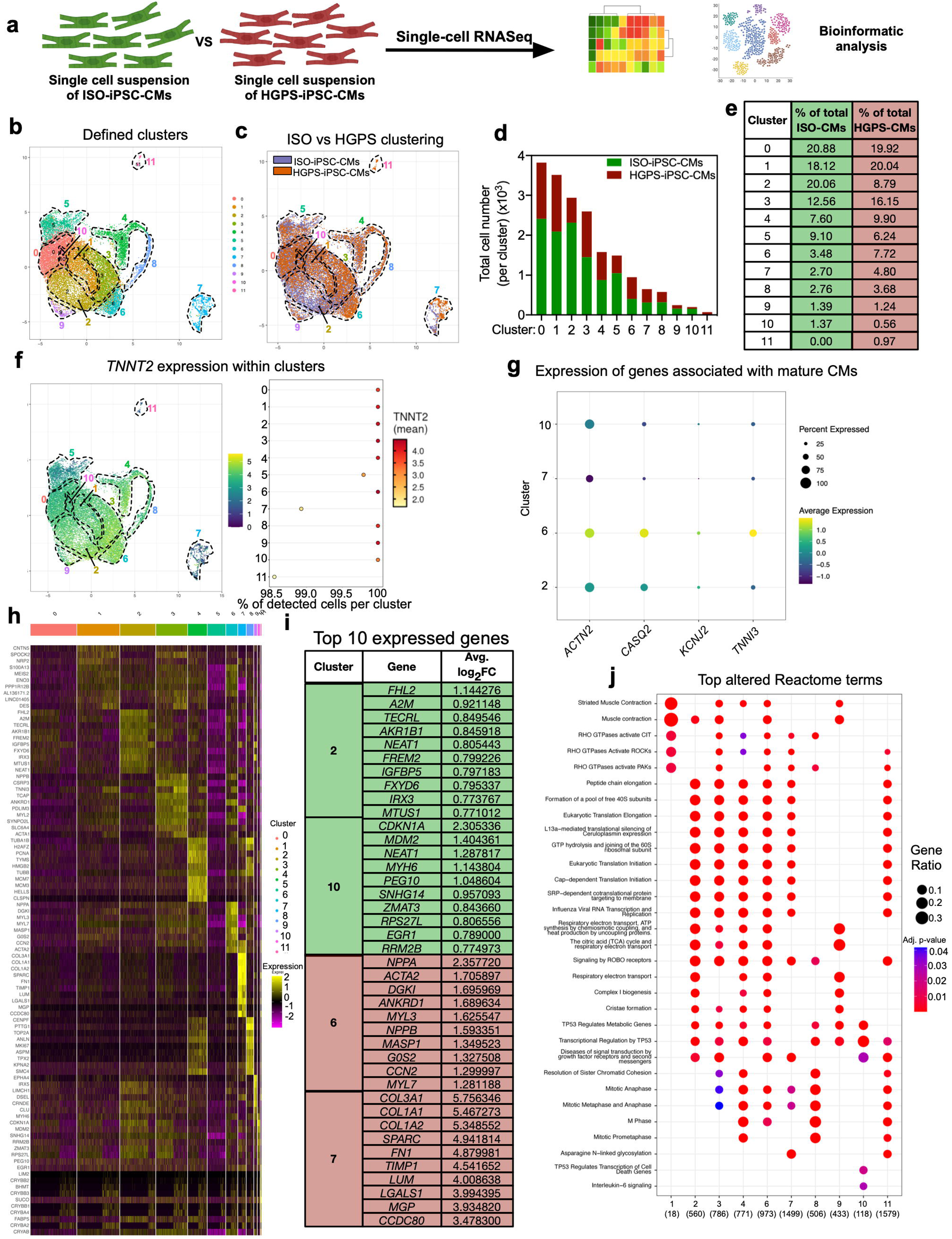
Genes and signaling pathways altered in HGPS-iPSC-CMs as evaluated by single-cell transcriptomics. **(a)** Single-cell RNASeq was applied in populations of day 30 ISO- and HGPS-iPSC-CMs. **(b)** Single-cell gene expression patterns revealed a subdivision in 12 clusters (from 0 to 11), as represented in a UMAP. **(c)** UMAP represents the separate distribution of ISO- and HGPS-iPSC-CMs through the 12 defined clusters. **(d)** Quantification of the total cell number and enrichment of each cluster in terms of ISO- and HGPS-iPSC-CMs. **(e)** Table showing the representativity of the number of ISO- or HGPS-iPSC-CMs present in each cluster, when compared with the total cell number of the respective group. **(f)** Expression of *TNNT2* (CM marker) by cluster, represented by UMAP (left) and quantified in terms of the percentage of expressing cells and level of expression (right). **(g)** Bubble plot representing, by cluster, the percentage of expressing cells and the level of expression of genes associated with increased CM maturity, for clusters 2, 6, 7 and 10. **(h)** Heatmap showing the top 10 expressed genes in each cluster, and the respective level of expression. **(i)** Top 10 genes and respective expression average log_2_ [fold-change] (FC). **(j)** Bubble plot representing, by cluster, the top altered Reactome terms, relative to their Gene Ratio and adjusted p-value.

On every cluster, the majority of cells (98.5% to 100%) expressed *TNNT2* at high levels (**Fig. 2f**), suggesting that differences in the gene expression profiles across clusters were not due to distinct cell type identities. Thus, we asked if the differences could be reflective of accelerated ageing in the HGPS-iPSC-CM cell model. Our results revealed that the HGPS-enriched cluster 6 had increased expression of genes associated with CM maturation (*e.g. ACTN2*, *CASQ2*, *KCNJ2*, *TNNI3*) compared with the ISO-enriched clusters 2 and 10 (**Fig. 2g** and **Extended Data Fig. 3b-e**). The HGPS-enriched cluster 7 did not follow this expression trend being, instead, enriched in senescence-related (CellAge dataset [27]) genes (**Fig. 2h** and **Extended Data Fig. 4a**).

To understand what distinguishes the different clusters, we identified the top 10 expressed genes per cluster and their respective expression levels (**Fig. 2i**). We particularly focused on the ISO-rich (2 and 10) and HGPS-rich (6 and 7) clusters (**Fig. 2i**). The highest expressed gene in cluster 2 was *FHL2* (**Fig. 2h, i; Extended Data Fig. 4b**), a gene encoding for the four-and-a-half LIM domain protein 2 (FHL2), which has been shown to exhibit cardioprotective and anti-hypertrophic roles [28]. Interestingly, we observed that FHL2 expression was downregulated in the LV heart tissue of HGPS mice using immunohistochemistry and Western blot analyses (**Extended Data Fig. 4c-d**). Conversely, the HGPS-enriched clusters 6 and 7 exhibited overexpression of hypertrophy-related genes (*NPPA*, *ANKRD1* and *NPPB*) [29] (**Fig. 2h, i**) and genes associated with cardiac ECM remodelling/fibrosis (*COL3A1*, *COL1A1*, *COL1A2*, *SPARC*, *FN1*, *TIMP1*, *LUM*) [30] (**Fig. 2h, i**).

Next, we performed pathway analysis, which revealed that HGPS-enriched clusters 6 and 7 exhibit alterations in RHO-related terms (**Fig. 2j**). It is known that RHO GTPases have crucial roles in cardiac hypertrophy and calcium handling [31].

Since both GO-term and pathway analyses suggest that ISO-iPSC-CMs are more protected from cardiac stressors than HGPS-iPSC-CMs, next we evaluated the heterogeneity of the HGPS-iPSC-CM cell population to determine if cluster phenotypes could be attributed to progerin expression. To this end, we selected reads whose sequences distinguish between the transcripts of the *LMNA* gene isoforms lamin A, lamin C and progerin, and eliminated the remaining reads. Given the similarity in sequence and sizes (3178 bp, 2461 bp, 2253 bp for lamin A, C and progerin, respectively), this approach has limitations and only allowed us to obtain relative proportions of cells expressing these isoforms throughout the clusters (**Extended Data Fig. 4f-h**). As expected, the proportion of detected progerin-expressing cells was greater in the HGPS-iPSC-CMs than in ISO-iPSC-CMs (3.41% vs. 0.02%, respectively). Moreover, the HGPS-iPSC-CM cluster most enriched in progerin-expressing cells was cluster 7, which also represents the cellular subpopulation that primarily accounts for the expression of senescence markers (CellAge dataset [27]) (**Fig. 2h** and **Extended Data Fig. 4a-4b)**.

Together these results identified populations of cells overrepresented in the HGPS-iPSC-CMs, which express higher levels of genes and pathways linked to cardiomyocyte stress and disease, and which includes a cluster enriched in progerin and senescence-related genes.

### HGPS-iPSCs-CMs show dysregulated mitochondrial respiration and calcium handling

The identified transcriptional differences prompted us to analyse the cell’s function including their metabolism and calcium handling (**Fig. 3; Supplementary Fig. 1a**). The SeaHorse XF Cell Mito Stress assay was used to assess mitochondrial respiration (**Fig. 3b-f**). Compared to ISO-iPSC-CMs, HGPS-iPSC-CMs showed an overall increased oxygen consumption rate (OCR) (**Fig. 3b**). The higher basal OCR suggests HGPS-iPSC-CMs have an increased mitochondrial activity relative to ISO-iPSCs-CMs (**Fig. 3b, c**) and/or that the cells are mostly relying on oxidative phosphorylation for ATP production (**Fig. 3b, d**) [32]. Furthermore, HGPS-iPSC-CMs exhibited significantly higher maximal respiration (**Fig. 3b, e**) and spare respiratory capacity (**Fig. 3b, f**), indicating that HGPS-iPSC-CMs increase their oxidative phosphorylation upon demanding conditions [32]. In contrast, ISO-iPSC-CMs were near their metabolic activity limit under normal culture conditions, as indicated by lower spare respiratory capacity (**Figs. 3b, f**). No differences were observed in the proton leak (**Supplementary Fig. 1b**).

**Figure 3-.**
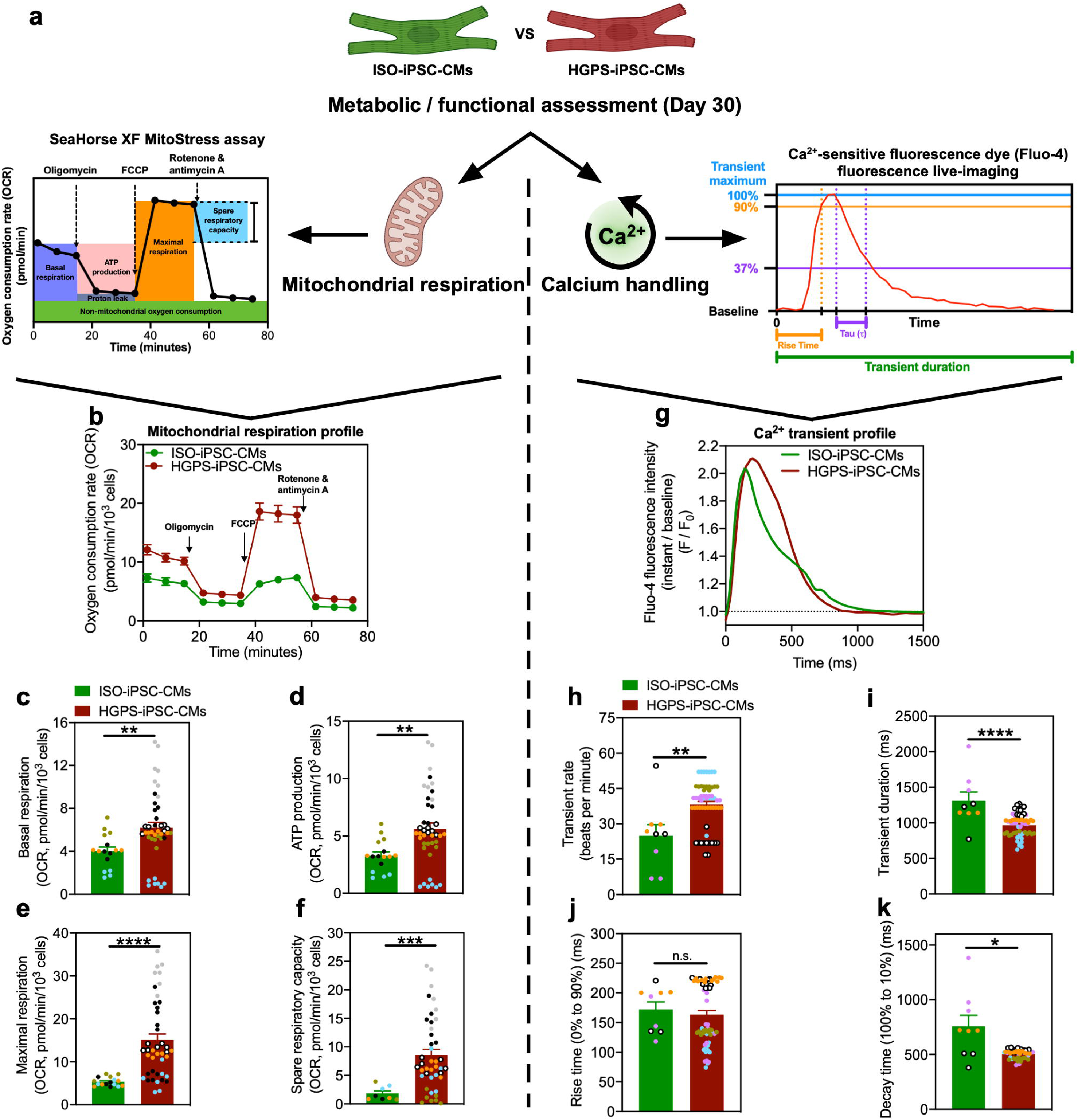
Mitochondrial respiration and calcium handling properties of HGPS-iPSC-CMs. **(a)** Overview of the experimental setup. **(b)** Average mitochondrial respiration profile curves represented as the oxygen consumption rate (OCR) normalized by cell number, over the assay time, and upon the sequential addition of the compounds oligomycin, FCCP, rotenone and antimycin A (n= 4-6 independent experiments, represented by color-coded symbols; n= 2-7 technical replicates per independent experiment). **(c-f)** Mitochondrial respiration parameters, all represented as OCR normalized by cell number for both ISO- and HGPS-iPSCs: **(c)** basal respiration, **(d)** ATP production, **(e)** maximal respiration and **(f)** spare respiratory capacity. **(g)** Average calcium transient curves of one individual cycle and represented by the Fluo-4 mean fluorescence intensity at a given acquisition moment, “F”, normalized to the respective signal baseline intensity, “F_0_” (n= 3-5 independent experiments, represented by color-coded symbols; n=3-12 technical replicates per independent experiment). **(h-k)** Determined calcium handling parameters: **(h)** transient rate, **(i)** transient duration, **(j)** rise time (time required for the calcium transient to increase from the saline, 0%, to 90% of its maximum), and **(k)** decay time (time required for the calcium transient to decrease from its maximum/peak to 10% of its maximum). In all figures, the values presented are mean ± SEM. n.s. - not significant, * - *p <* 0.05; ** - *p <* 0.01; *** - *p <* 0.001; **** - *p <* 0.0001.

Calcium handling was evaluated using high-speed fluorescence live imaging (**Figs. 3g-k**). HGPS-iPSC-CMs presented a higher rate of spontaneous calcium transients (**Fig. 3g, h**) and, consequently, their transient duration was shorter (**Figs. 3g, 3i**). Interestingly, HGPS-iPSC-CMs displayed a Ca^2+^ rise time similar to ISO-iPSC-CMs (**Fig. 3g, j**) but a significant decrease in the decay time (**Fig. 3g, k**), suggesting that HGPS-iPSC-CMs were more efficient than ISO-iPSC-CMs at removing the cytosolic Ca^2+^ (**Figs. 3i-k; Supplementary Fig. 1c**). No alterations were detected across groups in terms of calcium transient amplitude (**Supplementary Fig. 1c**).

Together our results indicated that HGPS-iPSC-CMs have increased mitochondrial respiration capacity and faster cytosolic calcium reuptake speed, which partially resembles the phenotype of a more mature CM.

### HGPS-iPSC-CMs express a senescent phenotype

To assess molecular alterations in HGPS-iPSC-CMs at the protein level, we performed proteomic analyses by mass spectrometry (**Fig. 4a-d**). 1014 proteins were significantly altered in HGPS-iPSC-CMs vs. ISO-iPSC-CMs (**Fig. 4b, c**). 129 out of 1014 proteins were specifically associated with cellular senescence as determined by the intersection with the CellAge database (composed of 1259 genes) [27] (**Fig. 4c** and **Supporting Table 1**). Proteome Ingenuity Pathway Analysis (IPA) further revealed that HGPS-iPSC-CMs have alterations in canonical pathways related to Oxidative Phosphorylation and the Sirtuin Signalling Pathway (**Fig. 4d**), which may reflect a compensatory mechanism to counteract ageing [33].

**Figure 4-.**
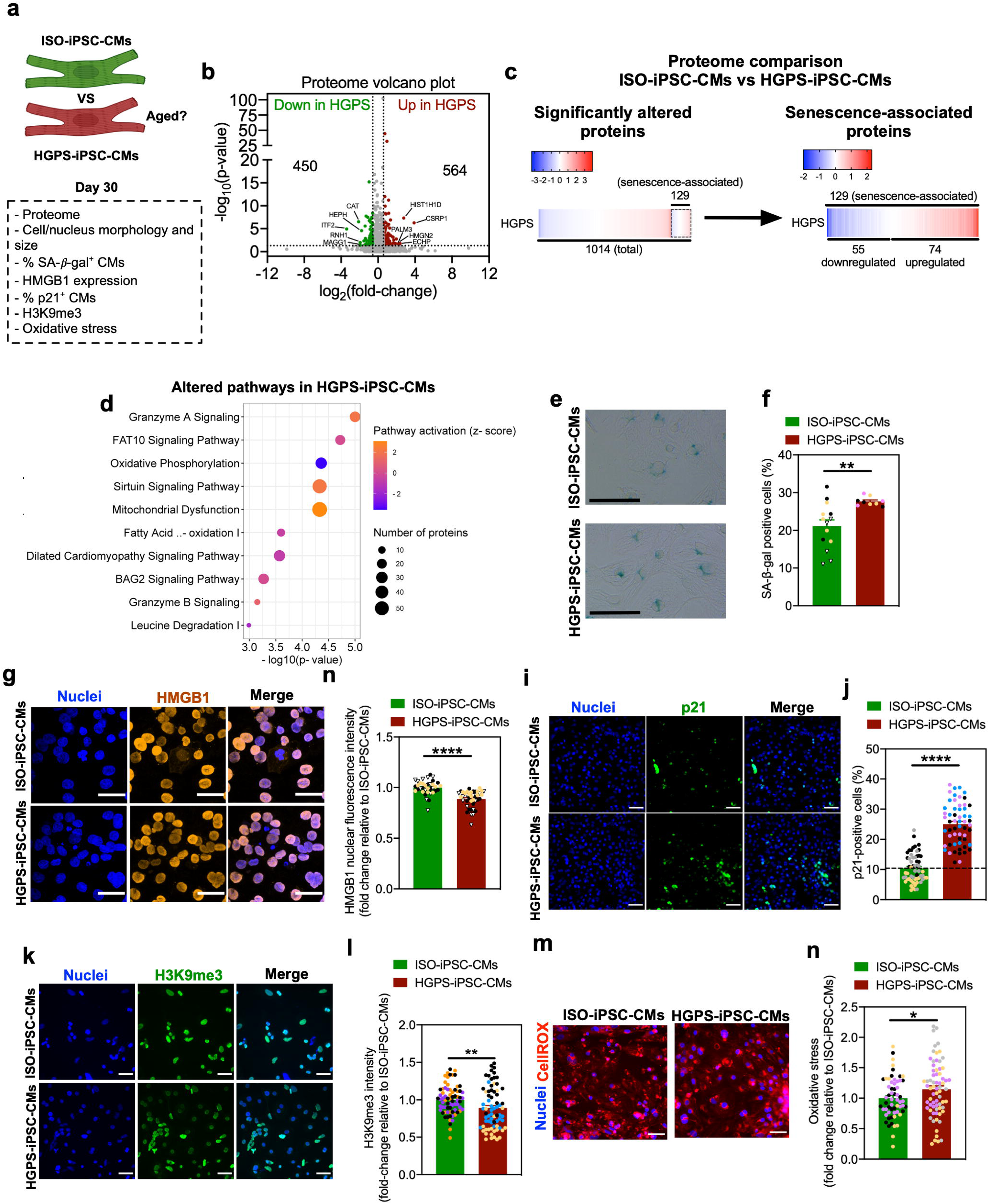
HGPS-iPSC-CMs present a senescent phenotype. **(a)** Summary of the experimental readouts. **(b)** Volcano plot comparing the proteome of ISO- and HGPS-iPSC-CMs (p-value<0.05; |log_2_(fold-change)|>0.58. **(c)** Heatmap for statistically altered proteins in HGPS-iPSC-CMs when compared with ISO-iPSC-CMs and, in particular, regarding senescence-associated proteins. **(d)** Bubble plot representing the canonical pathways that are altered in HGPS-iPSC-CMs, relative to ISO-iPSC-CMs, using the proteome data. Circle size represents the number of proteins associated with each pathway, −log(p-value) represents the likelihood of those proteins to be associated with that pathway, and color represents the predicted activation state of the pathway (z-score). **(e)** Representative images of the SA-β-gal colorimetric assay. Scale bars: 75 μm. **(f)** Percentage of SA-β-gal positive cells (n=3 independent experiments, represented by color-coded symbols; n=3-5 technical replicates per independent experiment). **(g)** Representative immunostaining for HMGB1 in ISO- and HGPS-iPSC-CMs. Scale bar: 30 μm. **(h)** Quantification of the nuclear HMGB1 protein content by image mean fluorescence intensity normalized to cell number and represented as a fold change of the ISO-iPSC-CMs (n=3 independent experiments, represented by color-coded symbols; n= 14-23 images per independent experiment). **(i)** Representative immunostaining for p21 in ISO- and HGPS-iPSC-CMs. Scale bar: 50 μm. **(j)** Percentage of p21-positive cells (n=3 independent experiments, represented by color-coded symbols; n=20 images per independent experiment). **(k)** Representative immunostaining for H3K9me3 in ISO- and HGPS-iPSC-CMs. Scale bar: 50 μm. **(l)** Quantification of the H3K9me3 content as image mean fluorescence intensity normalized to cell number and represented as a fold change of the ISO-iPSC-CMs (n=3 independent experiments, represented by color-coded symbols; n=24 images per independent experiment). **(m)** Representative fluorescence images with cells stained with CellROX (ROS-sensitive dye). Scale bar: 30 μm. **(n)** Quantification of the oxidative stress as the CellROX dye image mean fluorescence intensity normalized to cell number, and represented as a fold change of the ISO-iPSC-CMs (n=3 independent experiments, represented by color-coded symbols; n=23-24 images per independent experiment). All values are presented as mean ± SEM. * - *p <* 0.05, ** - *p <* 0.01, *** - *p <* 0.001, **** - *p <* 0.0001.

Next, we investigated in more detail the presence of senescence markers [21] in these cells (**Fig. 4e-n, Extended Data Figs. 5a-f**). HGPS-iPSC-CMs showed an increase of senescence markers relative to ISO-iPSC-CMs including higher levels of SA-/3-gal activity (**Fig. 4e, f**), lower levels of the high mobility group box 1 (HMGB1) protein in the nucleus (**Fig. 4g, h**), higher percentage of p21-expressing cells (**Fig. 4i, j**) and lower levels of the modification H3K9me3 (methylation of lysine 9 of histone H3) (**Fig. 4k, l**). Another common characteristic of senescent cells is the senescence-associated secretory phenotype (SASP) [21]. Although several canonical SASP-related factors released by HGPS-iPSC-CMs were seemingly increased relative to ISO-iPSC-CMs (**Extended Data Fig. 5a**), HGPS-iPSC-CMs only showed statistically increased expression of three non-canonical SASP genes (*EDN3*, *GDF15*, and *TGFB2*) **(Extended Data Fig. 5b-5d)** that have previously been shown to be overexpressed in aged/senescent mouse CMs [34].

To determine potential inducers of senescence in HGPS-iPSC-CMs we evaluated DNA damage and oxidative stress. No significant differences were found in the percentage of cells expressing 4 or more yH2AX foci (**Extended Data Figs. 5e-5f**); however, HGPS-iPSC-CMs presented an increased mean fluorescence intensity of CellROX, a ROS-sensitive dye (**Fig. 4m,n**), showing that these cells are under greater oxidative stress. This result is also supported by a downregulation of anti-oxidative proteins in HGPS-iPSC-CMs, namely SOD2, CAT, GPX3 and GPX7, as analysed by proteomics (**Supporting Table 2**).

Altogether, our results showed that HGPS-iPSC-CMs have a senescent phenotype possibly triggered by an increase in oxidative stress.

### A senescence phenotype is observed in the hearts of HGPS mice and human patients

Having demonstrated that HGPS-iPSC-CMs activated a senescent program, we questioned if such a senescent phenotype was also present in the cardiac tissue of mice and humans with HGPS and how it compared to the one observed in mice and humans during chronological ageing. For that purpose, we took advantage of a published proteomic dataset (<u>PXD039548</u>) [35] on LV tissue collected from chronological and HGPS aged hearts, from both mice and humans, and compared it with their young and non-progeroid counterparts, respectively (**Extended Data Figs. 6**). We focused on the LV as it is the most critical regarding cardiovascular function and the main chamber affected by heart failure in HGPS [6]. Proteome analysis showed that physiologically aged mice had 512 dysregulated proteins (**Supporting Table 3**), while HGPS mice presented 518 dysregulated proteins in comparison to their young/wild-type counterparts (**Supporting Table 4**), respectively (**Extended Data Figs. 6b, c**). Approximately 6.8% of the significantly altered proteins are shared by both mouse models and 30 proteins increase or decrease in the same direction in both models (**Extended Data Figs. 7a-d**). By intersecting the statistically altered proteins in both mouse models with the CellAge database [27], it was observed that physiologically aged and HGPS mice expressed 33 and 32 proteins, respectively, associated with senescence (**Extended Data Figs. 6b, c** and **Supporting Tables 5 and 6**). Moreover, gene set enrichment analysis showed that several pathways related to senescence were enriched in HGPS mouse heart LV tissue while metabolism and respiration pathways were downregulated (**Extended Data Figs. 6d**).

Next, we extended the previous analyses to human samples, also taking advantage of a published proteomic data set (<u>PXD039548</u>) [35], which was here re-analysed. We performed a similar proteomic analysis from formalin-fixed paraffin-embedded heart LV tissue from physiologically aged (65 to 70 years old) and HGPS individuals (10 to 14 years old), in comparison with their respective young/healthy controls (21 to 31 years old) (**Extended Data Figs. 6e-g**). 586 and 951 proteins were statistically altered in physiologically aged and HGPS individuals, respectively (**Extended Data Figs. 6f and Supporting Tables 7 and 8**). Approximately 13.6% of the significantly altered proteins were shared by both aged or HGPS individuals, with 117 proteins increasing or decreasing in the same direction (**Extended Data Figs. 6e-g**). By intersecting the statistically altered proteins in both patients with the CellAge database [27], 27 and 49 were related to senescence in physiologically aged and HGPS, respectively (**Extended Data Figs. 6f, g** and **Supporting Tables 9 and 10**).

Overall, our results confirmed that also *in vivo*, in mouse and human HGPS hearts, a senescence program is in place with an offset in metabolism. Moreover, we demonstrated that HGPS hearts show some similarities with chronological ageing, with both having increased expression of senescence-related genes, but the impact of ageing on the heart is broadly distinct from that of progerin accumulation in HGPS patients.

### Chronic drug effects in HGPS-iPSCs-CMs *vs* ISO-iPSCs-CMs: proteome

We next questioned whether drugs currently used and/or envisioned for HGPS treatment could revert the HGPS-iPSC-CMs’ phenotype. To this end, we investigated the impact of low-dose chronic (8 to 10 days) drug treatments on HGPS-iPSC-CMs with respect to their proteome (**Fig. 5**). Initially, we selected six compounds previously used in the context of HGPS, ageing and/or as a senotherapy, including: rapamycin (Rapa) [9, 36] and lonafarnib (Lona) [7, 8] (both tested in clinical trials in the context of HGPS); MG-132 [10] and neuropeptide Y (NPY) [11] (with promising results in HGPS pre-clinical studies); dasatinib + quercetin combination (D+Q) [37, 38] and fisetin [39, 40] (exploratory drugs in the context of HGPS but with a relevant history in the clinic as anti-senescent drugs). To define the drug concentrations to be used in the chronic treatment, we used concentrations reported in the literature [10, 11, 14, 40, 41]. Then, we evaluated their effect in ISO-and HGPS-iPSC-CMs after a 2-day or 10-day treatment with either a low dose (10% of the reported single-administration dose) or high dose (40% of the reported single-administration dose) (**Supplementary Fig. 2a-d**). In some cases, we had to decrease the concentration to avoid long-term cytotoxicity (**Supporting Table 11**). It is important to note that the proteomic methodology followed is unable to distinguish peptides from lamins and progerin, and thus, to evaluate the effect of the drugs in progerin. Initially, we assessed which of 1014 differentially expressed proteins in HGPS-iPSC CMs (**Fig. 4b**) had their expression altered by each drug (**Fig. 5b**). Rapa, Lona and D+Q reverted the expression tendency of 41, 37 and 51 downregulated proteins in untreated HGPS-iPSC-CMs, respectively. Rapa and Lona also reverted the expression tendency of 38 and 55 up-regulated proteins in untreated HGPS-iPSC-CMs, respectively, while D+Q had no effect. Of note, neither MG-132, fisetin nor NPY had a corrective effect on proteins altered in HGPS-iPSC-CMs (**Supplementary Fig. 3**).

**Figure 5-.**
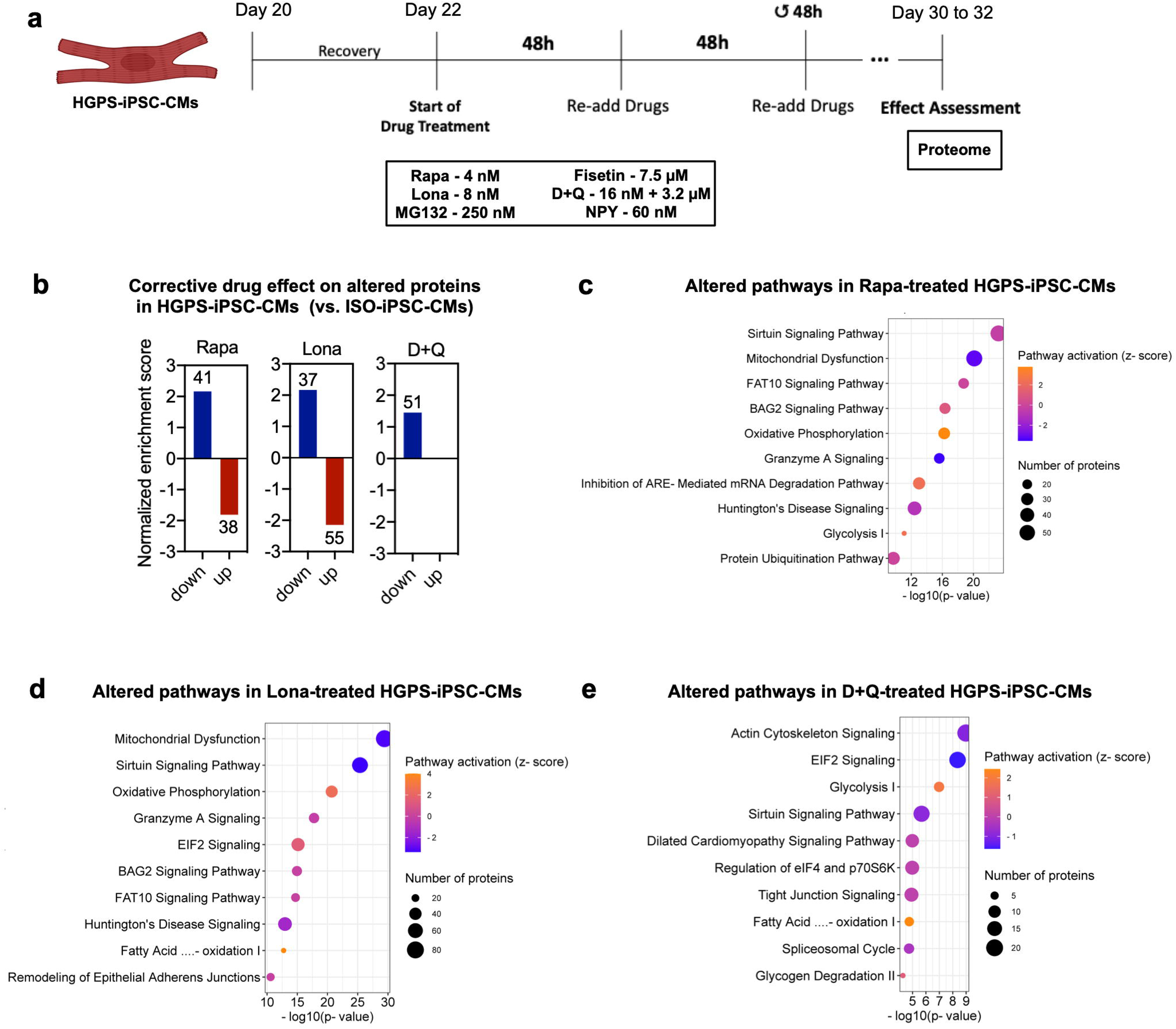
*In vitro* drug treatment effects on the proteome of HGPS-iPSCs-CMs. **(a)** Overview of the experimental setup. **(b-e)** Evaluation of the effect of drugs in the proteome of HGPS-iPSC-CMs. **(b)** Corrective effect of the drugs Rapa, Lona, and D+Q on the proteins that were originally downregulated or upregulated (regardless of statistical significance) in HGPS-iPSC-CMs relative to ISO-iPSC-CMs, plotted with the normalized enrichment score and with the number of proteins altered by the drugs presented over the bars. **(c-e)** Bubble plots representing the canonical pathways that are altered in HGPS-iPSC-CMs treated with the candidate drugs Rapa **(c)**, Lona **(d)**, and D+Q **(e)**, relative to untreated HGPS-iPSC-CMs, using the proteome data (cut off for pathway enrichment: p-value < 0-05). Circle size represents the number of proteins associated with each pathway, −log(p-value) represents the likelihood of those proteins to be associated with that pathway, and color represents the predicted activation state of the pathway (z-score).

Considering that Rapa, Lona and D+Q were the only drugs that partly reverted the HGPS-iPSC-CMs’ proteome, we explored in which pathways these three drugs acted (**Fig. 5c-e**). Rapa (**Fig. 5c**) and Lona (**Fig. 5d**) reverted the predicted activation state of relevant pathways that were altered in HGPS-iPSC-CMs (**Fig. 4d**), namely Oxidative Phosphorylation and Sirtuin Signalling canonical pathways (**Fig. 5c, d**), showing a potential corrective effect regarding mitochondrial function and mechanisms associated with ageing/senescence. On the other hand, D+Q only reverted the activation state of the Sirtuin Signalling Pathway (**Fig. 5e**).

Overall, our proteome results indicate that from the 6 drugs tested, the most promising in reverting the disease phenotype were Lona and Rapa, followed by D+Q drugs.

### Chronic drug effects in mitochondrial respiration and calcium handling of HGPS-iPSCs-CMs

Encouraged by the drug-induced proteomic results, next we evaluated drug effects in mitochondrial respiration (**Fig. 6a-e**) and calcium handling (**Fig. 6a, f-i**). MG-132, fisetin and NPY had little or no functional effect on HGPS-iPSC-CMs (**Extended Data Figs. 8a-k**). On the other hand, Rapa, Lona and D+Q altered HGPS-iPSC-CM function. Regarding mitochondrial respiration (**Fig. 6b-e; Extended Data Fig. 8a-8e; Supplementary Fig. 4**), Rapa was the best drug at reverting the metabolic profile of HGPS-iPSC-CMs, improving most parameters, namely basal respiration (**Fig. 6b**), ATP production (**Fig. 6c**), maximal respiration (**Fig. 6d**) and spare respiratory capacity (**Fig. 6e**). On the other hand, Lona corrected the basal respiration (**Fig. 6b**) and ATP production (**Fig. 6c**), and D+Q had practically no effect, despite showing a tendency to partially reverse the maximal respiration (**Fig. 6d**) and spare respiratory capacity (**Fig. 6e**).

**Figure 6-.**
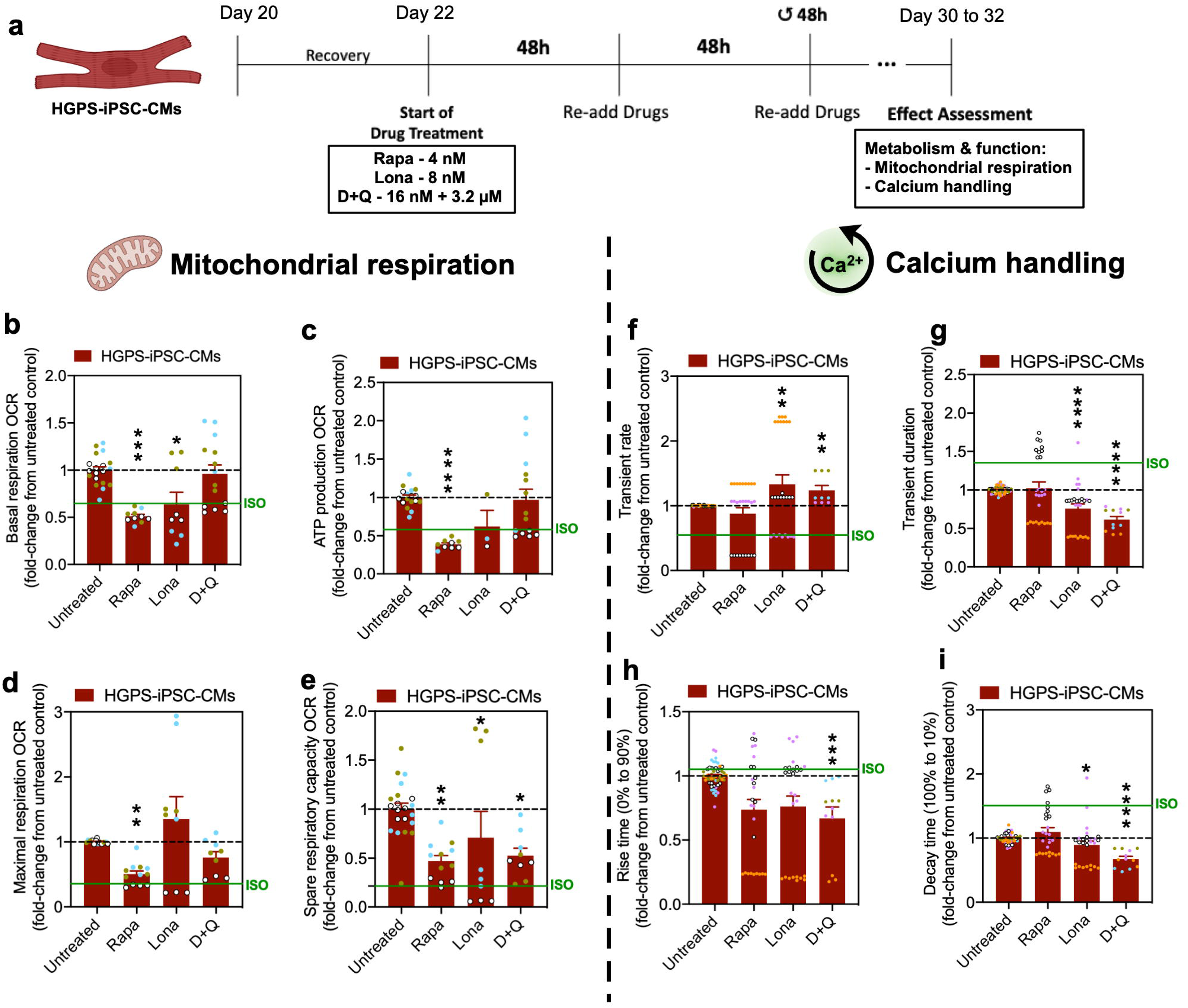
*In vitro* drug treatment effects on the metabolism and function of HGPS-iPSCs-CMs. **(a)** Overview of the experimental setup. **(b-e)** Mitochondrial respiration of HGPS-iPSC-CMs after drug exposure (the results are presented as fold-change of the OCR normalized to the number of cells and relative to the control, i.e., untreated cells; n=3 independent experiments, represented by color-coded symbols; n=1-7 technical replicates per independent experiment): **(b)** basal respiration; **(c)** ATP production; **(d)** maximal respiration and **(e)** spare respiratory capacity. **(f-i)** Evaluation of the effect of candidate drugs in the calcium handling of HGPS-iPSC-CMs (the results represent fold-change relative to control, i.e., untreated cells; n= 3-5 independent experiments, represented by color-coded symbols; n=3-13 technical replicates per independent experiment): **(f)** transient rate; **(g)** transient duration; **(h)** rise time (0% to 90%) and **(i)** decay time (100% to 10%). The dashed black line corresponds to the Untreated level. The continuous green line corresponds to the levels of ISO-iPSC-CMs on standard culture conditions. All values are presented as mean ± SEM. n.s. - not significant,* - *p <* 0.05, ** - *p <* 0.01, *** - *p <* 0.001, **** - *p <* 0.0001.

Regarding calcium handling (**Fig. 6f-i** and **Extended Data Fig. 8f-8k**), neither drug reverted the phenotype. However, Rapa showed a tendency to partially revert the HGPS phenotype, namely the transient rate (**Fig. 6f**) and the decay time (**Fig. 6i**) without disrupting the calcium transient amplitude (**Extended Data Fig. 8j**). It should also be noted that all drugs tested strongly decreased the rise time (0% to 90%) compared to the untreated control, *i.e.* further disrupting the calcium handling machinery in HGPS-CMs (**Fig. 6h**). Moreover, Lona and D+Q predominantly caused a detrimental effect in the HGPS-iPSC-CMs calcium handling, as not only did the calcium transients seem arrhythmic (increased rate and decreased durations) (**Fig. 6f, g**), but also because they led to a significant reduction in the calcium transient amplitude (Lona only, **Extended Data Fig. 8j**), and/or to generalized further disruption in calcium handling parameters (**Fig. 6f-i; Extended Data Fig. 8f-8k**), which suggests these drugs promoted further HGPS cardiomyocyte dysfunction.

Altogether, our results show that Rapa and Lona were better at reverting the metabolic profile of HGPS-iPSC-CMs than D+Q; however, only Rapa was able to revert partially some calcium handling features on HGPS-iPSC-CMs.

### Effect of chronic drug treatments in the senescence phenotype of HGPS-iPSCs-CMs

Having shown that the candidate drugs caused functional alterations on HGPS-iPSC-CMs, we then questioned if the drugs were able to revert the senescent phenotype in these cells (**Fig. 7a**). First, we evaluated their effect on the expression of nuclear HMGB1, a senescence marker (**Fig. 7b**). We complemented this analysis by evaluating the oxidative stress (**Fig. 7c**). Furthermore, we assessed the expression of senescence-related proteins that were significantly altered in HGPS-iPSC-CMs (**Fig. 7d, e**). Rapa and D+Q caused a conspicuous increase in HMGB1 in the nucleus (**Fig. 7b**) and a decrease in oxidative stress (**Fig. 7c**). Lona caused no relevant alterations in HMGB1 expression (**Fig. 7b**) nor in oxidative stress by CellROX staining (**Fig. 7c**), but induced the upregulation of anti-oxidative proteins, namely SOD2, GPX3, GPX4 and GPX7 (**Supporting Table 12)**. Importantly, all three compounds successfully reverted the expression pattern of several SASP proteins that were upregulated (**Fig. 7d**) or downregulated (**Fig. 7e**) in HGPS-iPSC-CMs. The other drugs, i.e. MG-132, fisetin and NPY, had little or no measurable effect (**Extended Data Fig. 9a-d**).

**Figure 7-.**
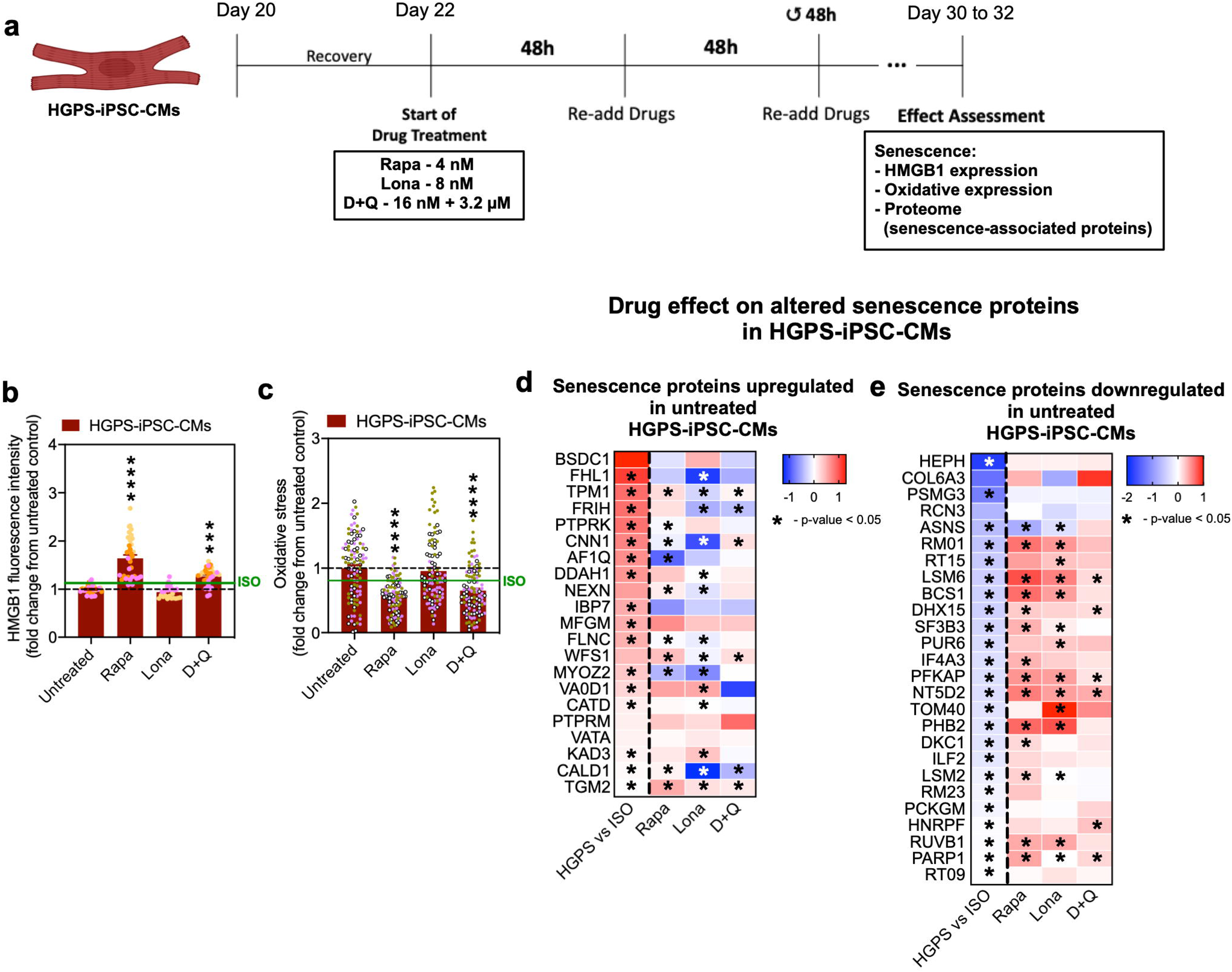
*In vitro* drug treatment effects on the senescence phenotype of HGPS-iPSCs-CMs. **(a)** Overview of the experimental setup. **(b)** Effect of the drugs on the HMGB1 nuclear protein content in HGPS-iPSC-CMs, represented as the fold-change of HMGB1 mean fluorescence intensity of each drug relative to control, i.e., untreated cells (n=3 independent experiments, represented by color-coded symbols; n=12 images pe independent experiment). **(c)** Effect of the drugs on the oxidative stress in HGPS iPSC-CMs, represented as the fold-change of the CellROX dye image mean fluorescence intensity of each drug relative to control (n=3 independent experiments, represented by color-coded symbols; n=36 images per independent experiment). **(d-e)** Heatmaps representing the effect of the candidate drugs on senescence-associated proteins that were originally significantly upregulated **(d)** or downregulated **(e)** in HGPS-iPSC-CMs, relative to ISO-iPSC-CMs. Heatmap colors represent the log_2_(fold-change) of protein expression. In the first, left-most column of each heatmap represents the change in protein expression of HGPS-iPSC-CMs relative to ISO-iPSC-CMs (as a reference), while the remaining columns represent the change of protein expression of drug-treated HGPS-iPSC-CMs in comparison with their Untreated counterparts. In figures b and c, the horizontal dashed black line corresponds to the Untreated level. The continuous green line corresponds to the levels of ISO-iPSC-CMs on standard culture conditions. Heatmaps tiles labelled with “*” correspond to comparisons where p-value < 0.05. All values are presented as mean ± SEM. *** - *p <* 0.001, **** - *p* 0.0001.

Altogether, Rapa, Lona and D+Q reverted the expression of SASP proteins in HGPS-iPSC-CMs, but only Rapa and D+Q improved HMGB1 accumulation in the nucleus and decreased oxidative stress.

### Chronic effect of Lona in the proteome and senescence of LV from progeria mice

Next, we asked how the results obtained *in vitro*, using our human cellular model, compared to the *in vivo* effect of the drug. Although Lona treatment did not stand out as more efficient than Rapa, we chose to perform this comparison using Lona, since it is the gold standard therapy for HGPS [7, 8]. Before conducting the *in vivo* test, we evaluated the senotherapeutic potential of Lona against HGPS-iPSC-CMs (**Extended Data Fig. 10**). As expected, Lona increased accumulation of prelamin A in HGPS-iPSC-CMs following 24 h of treatment. Our results further show that Lona had no senolytic effect (i.e. it did not eliminate selectively senescent cells under the concentrations tested) but it did show a senomorphic effect since Lona reverted the expression of non-canonical SASP genes in HGPS-iPSC-CMs, as evidenced by the downregulation of *EDN3* and *TGFB2* genes.

To evaluate the chronic effect of Lona treatment in the LV of *Lmna^G609G/G609G^* mice (abbreviated as LAKI mice)[42], we administered Lona daily (**Fig. 8a**). Lona treatment did not alter the body weight of either LAKI or control mice, with LAKI mice reaching, as previously reported, a body weight plateau at 6 weeks (**Supplementary Fig. 5a**). Irrespectively, Lona-treated LAKI mice exhibited statistically significant alterations in the expression of 186 proteins (48 upregulated and 138 downregulated proteins) in comparison to untreated LAKI mice (**Supporting Table 13**). Proteins involved in endoplasmic reticulum-Golgi transport (ERGIC1), sarcomere structure (MYL4 and MYL7), and intracellular trafficking of β1-adrenergic receptors (GOPC) were within those with the highest log_2_ fold-change (**Fig. 8b**). Amongst the top 30 proteins altered in untreated HGPS *vs.* WT animals (**Extended Data Fig. 7a-d**), only some (e.g. SERPINA1E, HMGA1) were improved in Lona-treated LAKI mice (**Fig. 8c**). Therefore, we then evaluated the effect of Lona treatment on senescence-related proteins (**Fig. 8d-f**). From the 186 proteins with significantly altered expression (**Supporting Table 13**), 21 were related to senescence/ageing including 3 upregulated (e.g. TAGLN) and 18 downregulated (e.g. EEF1E1, TGFB1, ZNF207, LBR) (**Supporting Table 14 and Fig. 8d**). Interestingly, amongst the top 20 senescence-related proteins altered in HGPS *vs.* LAKI mice (**Fig. 8e**) (regardless of statistical significance), 4 out of the 9 downregulated senescence proteins had their expression increased and, 17 out of the 19 upregulated proteins had their expression decreased (**Fig. 8e**). Noteworthy, down/up-regulation of these proteins upon Lona treatment was specific for the HGPS mice, i.e. Lona treatment had no effect on these proteins in WT mice (**Fig. 8e**). Moreover, Lona corrected the expression pattern of the senescence-related protein SFPQ [43] (**Fig. 8f**).

**Figure 8-.**
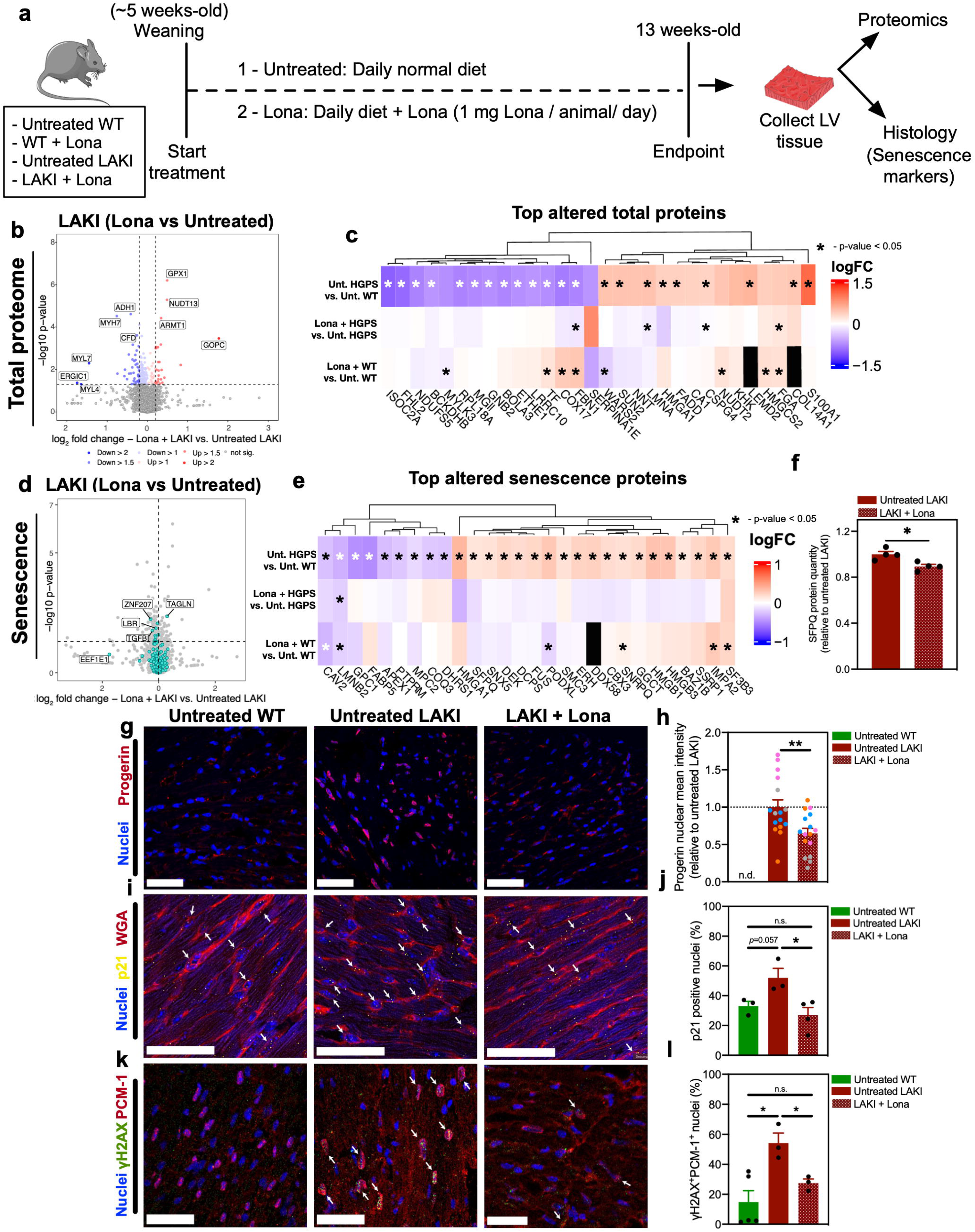
Effect of Lona treatment on *Lmna^G606G/G609G^* (LAKI) mouse heart. **(a)** Overview of the experimental setup. **(b-c)** Protein content comparison analysis within the LV tissue, in terms of the total proteome: **(b)** Volcano plot of proteins that have their expression altered in LAKI animals treated with Lona *vs* Untreated; **(c)** heatmap with the representation of top 30 total proteins with altered expression in LAKI *vs* WT animals, and their respective expression changes in Lona-treated LAKI and WT animals. **(d-f)** Expression of senescence-related proteins within the LV tissue of Lona-treated and untreated LAKI mice: **(d)** Volcano plot of senescence-related proteins that have their expression altered in LAKI animals treated with Lona *vs* untreated; **(e)** heatmap with the representation of senescence-related proteins with altered expression in LAKI *vs* WT animals, and their respective expression changes in Lona-treated LAKI and WT animals. Heatmaps tiles labelled with “*” correspond to comparisons where p-value < 0.05.; **(f)** protein quantity of the senescence-related protein SFPQ in Untreated and Lona-treated LAKI animals (normalized to Untreated LAKI). **(g-h)** Progerin expression analysis by immunofluorescence: **(g)** representative images of progerin staining in LV of the experimental groups (scale bar: 25 μm); **(h)** quantification of progerin nuclear fluorescence intensity (values normalized to Untreated LAKI; n=4 animals, colour-coded; n=3-5 images per animal). n.d. - not detected. **(i-j)** p21 expression analysis by RNAish: **(i)** representative images of p21 staining in LV of the experimental groups. Arrows point to examples of p21-positive nuclei (scale bar: 25 μm); **(j)** quantification of p21-positive nuclei in the LV tissue (n=3-4 animals, colour-coded; n=5 images per animal). **(k-l)** Expression of γΗ2ΑΧ in CMs (nuclear marker for CMs: PCM-1): **(k)** representative images of γH2AX foci nuclear staining specifically in CMs, in the LV tissue (arrows point to nuclei of CMs that are positive for γH2AX; scale bar: 25 μm); **(l)** quantification of γΗ2ΑΧ-positive nuclei of CMs (PCM-1 positive) in the LV tissue (n=3-5 animals; n=5 images per animal). All values are presented as mean ± SEM. n.s. - not significant, * - *p <* 0.05, ** - *p <*0.01.

Progerin (**Fig. 8g, h**) and senescence marker expression (**Fig. 8i-l**) were further evaluated by immunofluorescence and RNAish analyses. Progerin expression in LV tissue from Lona-treated LAKI mice was lower than in untreated ones (**Fig. 8g, h**). On the other hand, the percentage of p21-positive cells in LV tissue from Lona-treated LAKI mice was lower in untreated ones (**Fig. 8i, j**), but Lona did not alter p21 expression in the control animals (**Supplementary Fig. 5b**). Moreover, we evaluated the nuclear γH2AX (a marker of DNA damage) expression in CMs, which were identified by PCM-1 staining (**Fig. 8k, l**). In keeping with an improved senescent phenotype, the percentage of CMs (PCM-1-positive cells) accumulating γH2AX foci was lower in Lona-treated LAKI mice than in untreated LAKI mice (**Fig. 8k, l**).

Overall, our results underline that Lona treatment can prevent or revert senescence in LV-CMs of HGPS hearts, similarly to what happens *in vitro* in Lona-treated HGPS-iPSC-CMs.

### Lona triggers autophagy activity on progeria heart mice and HGPS-iPSC-CMs

To elucidate the mechanism by which Lona reverses senescence in HGPS mice hearts, we initially conducted proteomic analyses. Notably, Lona treatment elicited a statistically significant change in the expression of several autophagy-related proteins in both WT (**Fig. 9a**) and HGPS (**Fig. 9b**) hearts. The net effect on autophagy-related proteins in HGPS was higher than in WT hearts (**Fig. 9c**). To further validate autophagy induction, we evaluated the impact of prolonged Lona exposure on HGPS-iPSC-CMs via proteomics. Lona treatment resulted in the upregulation of several autophagy-related proteins such as ATG9A, VSP11 and VSP18 (**Fig. 9d-f**) compared to untreated HGPS-iPSC-CMs. Next, we assessed the cellular effects of this autophagy induction in HGPS-iPSC-CMs. Cells were treated with Lona for 24 h, after which they were exposed or not to chloroquine, an autophagy inhibitor, in the presence of Lona. The protein levels of the LC3B-II and LC3B-I, widely used autophagy markers, were assessed by Western blot (**Fig. 9g–k**). Results demonstrated that Lona enhanced autophagic flux (**Fig. 9j**) and increased autophagosome formation (**Fig. 9k**).

**Figure 9-.**
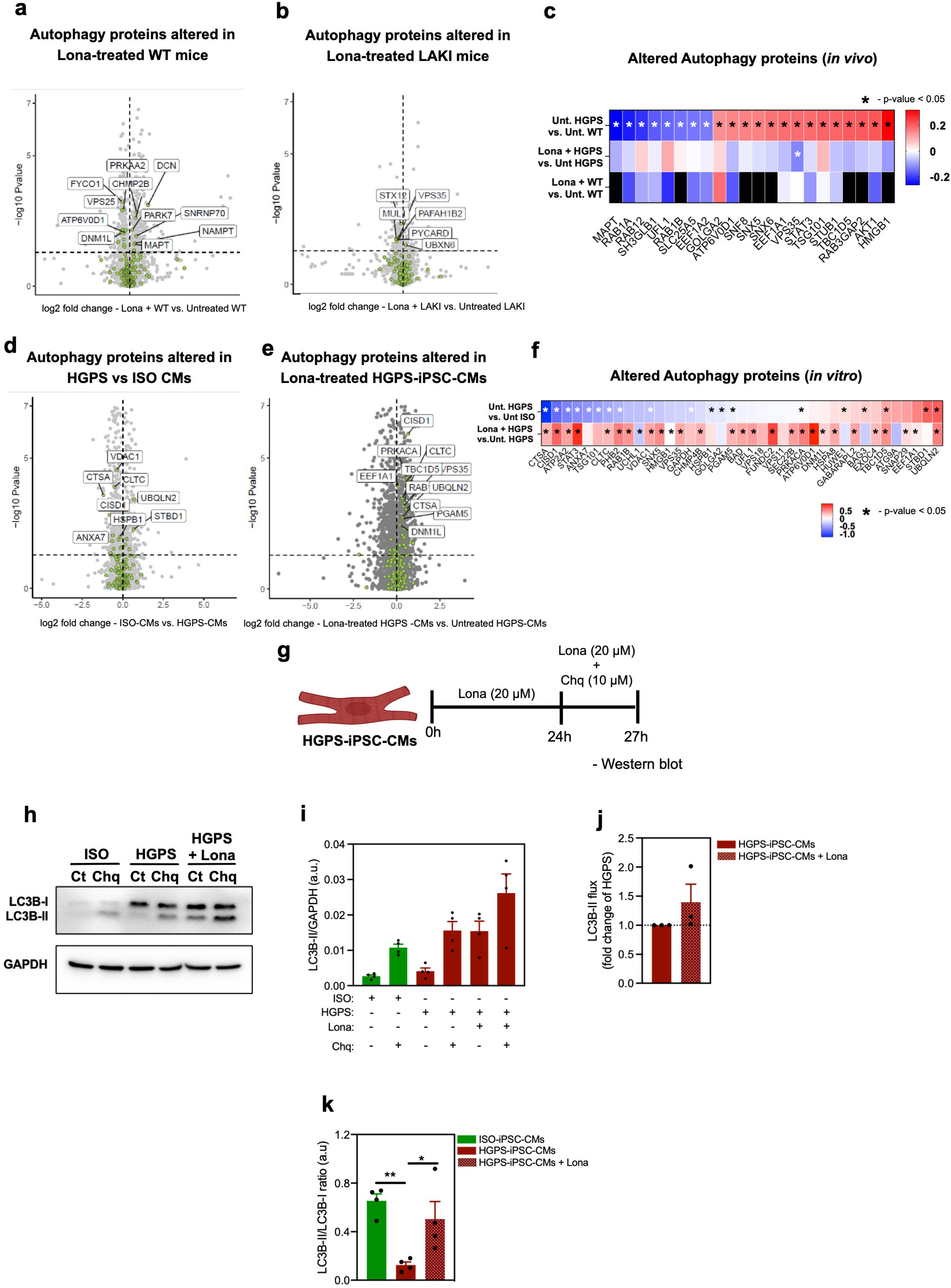
Effect of *in vivo* and *in vitro* Lona treatment on autophagy regulation. **(a-c)** *In vivo* proteomic analysis of autophagy-related proteins in WT and LAKI mice with or without Lona treatment. **(a)** Volcano plot of autophagy-related proteins that have their expression altered in WT animals treated with Lona *vs* Untreated. **(b)** Volcano plot of autophagy-related proteins that have their expression altered in LAKI animals treated with Lona *vs* Untreated. **(c)** Heatmap with the representation of top 22 autophagy-related proteins with altered expression in LAKI *vs* WT animals, and their respective expression changes in Lona-treated LAKI and WT animals. Heatmap tiles labelled with “*” correspond to comparisons where p-value < 0.05. **(d-f)** *In vitro* proteomic analysis of autophagy-related proteins in HGPS-iPSC-CMs with or without Lona treatment. **(d)** Volcano plot of autophagy-related proteins that have their expression altered in untreated ISO- vs. HGPS-iPSC-CMs. **(e)** Volcano plot of autophagy-related proteins that have their expression altered in Lona-treated HGPS-iPSC-CMs vs. Untreated. **(f)** Heatmap with the representation of top 36 autophagy-related proteins with altered expression in ISO- vs. HGPS-iPSC-CM lines, and their respective expression changes in Lona-treated ISO- and HGPS-iPSC-CM lines. Heatmap tiles labelled with “*” correspond to comparisons where p-value < 0.05. **(g-k)** Effects of *in vitro* acute Lona treatment in HGPS-iPSC-CMs on autophagy. **(g)** Experimental design of the acute *in vitro* treatment of HGPS-iPSC-CMs with Lona and chloroquine (Chq). **(h)** Representative Western blot of autophagy-related proteins (LC3B-I and LC3B-II) and the housekeeping protein (GAPDH) on protein extracts obtained from ISO-iPSC-CMs treated (Chq) or not (Ct) with chloroquine, and from HGPS-iPSC-CMs, treated or not with chloroquine and/or Lona. **(i)** LC3B-II protein levels normalized by the housekeeping protein GAPDH. **(j)** Autophagic flux on non-treated and Lona-treated HGPS-iPSC-CMs. **(k)** LC3B-II/LC3B-I protein ratio on both ISO-iPSC-CMs and HGPS-iPSC-CMs (non-treated or Lona-treated). All values are presented as mean ± SEM. *-*p <* 0.05, **-*p <*0.01.

To further validate the effect of Lona as autophagy inducer we have used the human osteosarcoma U2OS cell line expressing the tandem reporter mCherry–green fluorescent protein (GFP)–LC3B (**Extended Data Figs. 9e-h**). This cell line assesses both induction of autophagy and autophagic flux, with autophagosomes being identifiable as puncta labelled with both mRFP and GFP. Upon fusion of autophagosomes with lysosomes to generate degradative autolysosomes the GFP signal is quenched, allowing for the flux through the autophagic pathway to be monitored by the ratio of GFP-RFP puncta to RPF only puncta (autophagic flux). Cells exposed for 24 h to Lona showed an increased autophagic flux in a dose-dependent manner with a minimum effect concentration (MEC) of 10 μM (**Extended Data Figs. 9e and f**). Importantly, Lona’s induction of autophagy was not attributable to non-specific lysosomal stress induced by lysosomotropism, as confirmed by additional assays (data not shown). Next, we examined lysosomal proteolytic activity as an orthogonal measure of autophagic degradation. U20S cells were treated with different concentrations of Lona for 18 h, then fed the dye-protein conjugate DQ-BSA. Uptake by cells and degradation of DQ-BSA with lysosomes/autolysomes results in the formation of fluorescent puncta. We observed that Lona treatment enhanced lysosomal activity in U2OS cells at concentrations above 5 μM, consistent with activation of the autophagic pathway by the drug (**Extended Data Figs. 9g and h**).

Overall, our results indicate that Lona treatment resulted in an overall improvement in lysosomal function which might prevent or attenuate the effects of CM senescence.

## DISCUSSION

Our study provides mechanistic insights into cardiac dysfunction associated with HGPS by highlighting the role of cardiac senescence. Additionally, it investigates potential pharmacological interventions for cardiac dysfunction and elucidates Lona’s mode of action within the cardiac context. We found that HGPS-CMs undergo premature maturation, resulting in elevated ROS production. The inability to adequately regulate ROS levels leads to pathological features such as increased calcium sparks and cellular senescence. Notably, HGPS-CMs exhibit senescent characteristics similar to those observed in physiological ageing in both mice and humans. Chronic treatment with rapamycin and Lona partially reversed senescence markers and improved mitochondrial function in HGPS-iPSC-CMs. Furthermore, we showed for the first time Lona’s senotherapeutic effects in the hearts of progeroid mouse models. Our data suggest that Lona mitigates cardiomyocyte senescence primarily by inducing autophagy. Despite its progression into clinical use, Lona’s safety profile warrants careful consideration. We observed that Lona treatment reduced calcium amplitude and increased transient rates in CMs, indicating potential impacts on cardiac contractility. These findings highlight the importance of monitoring cardiovascular effects during Lona therapy. Overall, our results highlight the power of our cell model to explore new biomarkers and therapies for HGPS and, possibly, aspects of cardiac physiological-ageing such as senescence.

HGPS, characterised by accelerated ageing, presents pathological cardiovascular defects that resemble aspects of natural cardiovascular ageing [3, 5, 6, 16], and has been used as a model to study ageing not only in animal models but also in human cells [14, 15, 17–20, 44]. By performing high-content proteomic analysis comparing human and mouse chronological-aged and HGPS heart LV tissue with their young and non-progeroid counterparts, respectively, we were able to demonstrate that while there is a high concordance between HGPS and aged CMs expressed proteins (97.8% of proteins expressed, 2843 of 2906), only a subset of significantly altered proteins in HGPS were also altered in physiologically aged hearts (13% and 19% in mouse and human, respectively). The main similarity this analysis flagged was the presence of a senescent phenotype in both physiologically-aged and HGPS, suggesting senotherapeutic therapies may help improve both HGPS and aged individuals.

Heart function, including LV diastolic dysfunction assessment, is used as a primary endpoint in HGPS clinical trials [7]. Therefore, any therapeutical drugs under development should be evaluated for their effect on LV function also in a pre-clinical setting. Our HGPS-iPSC-derived LV-CMs, generated based on our previously published protocol [23], offer such a platform for disease modelling and drug screening, as exemplified by their ability to accumulate progerin. These cultures differ from others that could be obtained using standard CM differentiation protocols [45, 46], due to their LV-specificity and higher maturity, which increases the robustness of its readouts in disease modelling and drug screening studies. Interestingly, our single-cell transcriptomic results indicate that only a small percentage of cells accumulate progerin (enriched within senescent cluster), suggesting this model represents the early stages of disease and unexpectedly revealing that, prior to developing a stereotypical senescent phenotype, HGPS-CMs undergo accelerated maturation as evidenced by the overrepresentation of the maturity/hypertrophy-enriched gene cluster in HGPS cultures.

During normal development, maturation is driven by metabolic changes whereby cells become more and more reliant on oxidative phosphorylation (OXPHOS) [47, 48]. Increased ROS production is seen during this glycolysis–OXPHOS transition and is usually offset by an increased ability the more mature cardiomyocytes have to deal with oxidative stress. HGPS-iPSC-CMs appear to prematurely transition to OXPHOS, resulting in an increase in ROS levels but, unlike healthy matured cardiomyocytes, these cells cannot keep ROS levels in check, leading to an increase in calcium sparks and senescence, known pathological consequences of increased ROS levels in other disease contexts [49, 50]. Our data is therefore in line with a model where HGPS-iPSC-CMs undergo premature-maturity, as would be expected of cells that are ageing faster, but which fail to contend with the increased metabolic output, possibly a response to the deleterious effects of progerin, leading to secondary calcium imbalance and cellular senescence. This is in keeping with the increased basal respiration and ROS levels seen in CMs isolated from old mice [51], senescent (late-passage) human cardiac fibroblasts, and HGPS human dermal fibroblasts [52].

The senescent phenotype observed in LV-CMs *in vitro* and *in vivo* led us to explore six drugs, some of them in clinical use (Lona, Rapa, D+Q, Fisetin). Rapa and Lona gave the best results in terms of reversion of the metabolic and senescent phenotypes. Rapa is an FDA-approved drug known to slow down ageing [9], but its main use in the clinic is as an immunosuppressant drug. The beneficial aspects of Rapa might be ascribed to its effectiveness in: (i) reverting the mitochondrial respiration phenotype and (ii) inducing autophagy, which is beneficial to clear progerin and other damaged molecules [9]. Rapa was the only drug that improved both metabolism and ROS levels and may be for that reason the only drug which showed a beneficial trend in calcium handling modulation, appearing to help revert the increase of calcium decay seen in HGPS-iPSC-CMs as well as to reduce their transient rate. Overall, our data are indicative that Rapa may be a hopeful treatment for HGPS patients, improving their CM function and reducing their senescent phenotype. It remains however to be addressed if Rapa has the same global positive impact in other cell types affected by progeria. Moreover, given the immunosuppressor role of Rapa, its dose may need to be carefully titrated for treating HGPS patients.

Lona, the only FDA-approved drug used to treat HGPS patients, was effective at: (i) reverting the expression of proteins that were differentially expressed in HGPS *vs* WT, including senescence-related and autophagy-related proteins and biomarkers; (ii) reducing the expression of genes linked with non-typical SASP; and (iii) reducing progerin levels. Both *in vivo* and *in vitro* results demonstrate that Lona induces autophagy in cardiac tissue, particularly in CMs. While autophagy induction helps mitigate the senescence phenotype in cardiac cells, it was insufficient to counteract the increased production of ROS by the premature maturation observed in HGPS-iPSC-CMs. This partial response may explain why Lona treatment failed to improve calcium handling and only partially rectified mitochondrial respiration back to control levels. Nonetheless, in this work, we also showed that chronic treatment (8 weeks) of LAKI mice with a Lona-supplemented diet caused a conspicuous reversion of the expression of senescence proteins and biomarkers in the LV tissue of these animals confirming the results we had unveiled with our *in vitro* cell model.

In conclusion, our data show that CM senescence is present in HGPS as well as in aged individuals, both in mice and humans, and the accumulation of senescence in CMs correlates with cellular dysfunction. We further show that Lona, a farnesyl transferase inhibitor approved by FDA for the HGPS patients, can prevent or attenuate the senescence phenotype on CMs but is unable to resolve the senescence trigger, i.e., to reduce the ROS levels in HGPS-CMs. Overall, our work opens new avenues in the research of molecular mechanisms, biomarkers, and therapeutic targets associated with cardiac ageing in the context of HGPS and possibly also in physiological ageing.

## ACKNOWLEDGMENTS

The authors would like to thank the input of Armindo Salvador, Francisco Santos, Rodrigo Santos, Jing Cai and Rebecca Hulbert for support with parts of the current work. The authors gratefully acknowledge support from the FLI Core Facility Proteomics. We thank the following scientific platforms/units at the Francis Crick Institute: advanced sequencing, light microscopy, and HESCU. We thank Lyn Healy for advice and support with stem cells. This work was funded by FEDER through the Program COMPETE and by Portuguese fund through FCT in the context of the projects 2022.07615.PTDC and PTDC/BTM-SAL/5174/2020, as well as the European projects RESETaging (ref. 952266), “CHAngeing – Connected Hubs in Ageing: Healthy Living to Protect Cerebrovascular Function” (Ref. 101087071), “DREAMs - Drug Repurposing with Artificial Intelligence for Muscular Disorders” (Ref: 101080229) and PRR project “HfPT”. The Bernardo lab was funded by the Wellcome Trust (210987/Z/18/Z). We were also funded by the Francis Crick Institute, which receives its core funding from Cancer Research UK (FC001157), the UK Medical Research Council (FC001157), and the Wellcome Trust (FC001157). The FLI is a member of the Leibniz Association and is financially supported by the Federal Government of Germany and the State of Thuringia.

## DATA AVAILABILITY

All data generated and analyzed are included in the main manuscript, extended data and supplementary information. For reviewing purposes, all associated raw data files related to the *in vitro* and *in vivo* proteomics (relative to Figures 2 and 8) are currently deposited in the public repository MassIVE under the following accession number: MSV000096058_reviewer (password: FLIReviewer_2024). Regarding the *in vivo* proteomics shown in Figure 5, it originates from a previously published dataset [35], and is available in ProteomeXchange Consortium via the PRIDE [53] partner repository, and can be accessed with the dataset identifier PXD039548. Relatively to the scRNA-seq data (Figure 2), the GEO ID number is yet to be provided.

## ONLINE METHODS

For more detailed materials and methods please see *Supporting Information*.

### iPSC differentiation into CMs

The 30-day protocol used for the differentiation of iPSCs into left ventricle CMs was based on the one used by Dark *et al.* [23]. Before initiating a differentiation, iPSCs were dissociated to a single-cell suspension using TrypLE™ Express Enzyme (ThermoFisher Scientific) (5 min, 37°C) and then centrifuged (300*g*, 5 min, RT). After being resuspended in mTeSR™ Plus medium (STEMCELL Technologies) supplemented with Y-27632 dihydrochloride (5 µM; 1254; Tocris), the cells were plated at the densities of 8.4×10^4^ and 7.6×10^4^ cells/cm^2^ for ISO- and HGPS-iPSCs, respectively. The medium was replaced the following day with mTeSR™ Plus medium (STEMCELL Technologies). The differentiation was initiated (day 0) when the iPSCs reached approximately 80% of confluency, 2-3 days after plating. Basal medium comprised of RPMI 1640 Medium, GlutaMAX™ (Thermo Fisher Scientific) either: supplemented with B-27™ Supplement, minus insulin (Thermo Fisher Scientific) for days 0 to 9; or supplemented with B-27™ Supplement, minus vitamin A (Thermo Fisher Scientific) and Penicillin-Streptomycin (Pen/Strep) (25 U/mL:25 µg/mL; Corning) for days 10 to 30. Regarding the added factors, AGN 193109 Sodium Salt (100 nM; sc-210768; Santa Cruz) was used for days 0 to 9, and L-Ascorbic acid (65 µg/mL; A92902; Sigma) was first added on day 2 and remained throughout all of the differentiation. Moreover, Activin A (5 ng/mL; 338-AC; R&D Systems), BMP-4 (3 ng/mL; 314-BP; R&D Systems), CHIR-99021 HCl (3 nM; S2924; Selleckchem) and FGF2 (2 ng/mL; 233-FB; R&D Systems) were only added on day 0 and removed 24 h later. Finally, IWR-1 (1 nM; I0161; Sigma) was added on day 2 and removed on day 4. During the differentiations, the medium was replaced (and fresh factors added) when the medium composition changed, the day after replating (*i.e.* days 14 and 21) or, otherwise, every two days. To maintain, expand and increase the purity of the CM population, cells were dissociated and replated on days 13 and 20.

### CM dissociation and replating

Not only as part of the differentiation protocol (days 13 and 20) but also to perform some of the characterizations on day 30, the CMs had to be dissociated and replated. For dissociation, CMs were incubated with STEMdiff™ CM Dissociation Medium (STEM Cell Technologies) (12 min, 37°C). Then, 2 parts (in volume) of STEMdiff™ CM Support Medium (STEM Cell Technologies) were added and mixed with 1 part of the dissociation medium, previously added to the CMs. Dissociation was finalized by pipetting the CMs to a single-cell suspension. Only at day 13, the CMs were strained (Falcon® 40 µm Cell Strainer; 352340; Life Sciences) before being counted and replated. The cell suspension resulting from the dissociation step was centrifuged (300*g*, 5 min, RT) and resuspended in a differentiation medium (see iPSC differentiation into CMs section) supplemented with Y-27632 dihydrochloride (5 µM; 1254; Tocris) and replated on Matrigel-coated culture plates at a density of 2.5 x 10^5^ cells/cm^2^, except on assays in which low-density plating was needed. On the following day, the medium was replaced with a standard differentiation medium.

### Genome editing: plasmids, sgRNA and repair template

The sgRNA used (5’-AGGAGATGGGTCCACCCACC-3’) was designed to target the HGPS mutation site, specifically on the mutated HGPS allele. This sgRNA was cloned into a pSpCas9(BB)-2A-GFP (px458) plasmid (Addgene), containing an insert for Cas9 from *Streptococcus pyogenes*. For the repair template, required to correct the HGPS point mutation by homology directed repair (HDR), the following sequence inserted into pUC57 plasmid, was used.

5’- TCCTTGGGCACAGAACCACACCTTCCTGCCTGGCGGCTGGGAGCCTGCAGGAGCCTGGAGCCTGGTTGGGCCTGAGTGGTCAGTCCCAGACTCGCCGTCCCGCCTGAGCCTTGTCTCCCTTCCCAGGGCTCCCACTGCAGCAGCTCGGGGGACCCCGCTGAGTACAACCTGCGCTCGC GCACCGTGCTGTGCGGGACCTGCGGGCAGCCTGCCGACAAGGCATCTGCCAGCGGCTCAGGAGCGCAGGTGGGCGGACCCATCTCCTCTGGCTCTTCTGCCTCCAGTGTCACGGTCACTCGCAGCTACCGCAGTGTGGGGGGCAGTGGGGGTGGCAGCTTCGGGGACAATCTGGTCACCCGCTCCTACCTCCTGGGCAACTCCAGCCCCCGAACCCAGGTGAGTTGTCTCTGCTTTGTCTCCAAATCCTGCAGGCGGGTCCCTGGTCATCGAGGGGTAGGACGAGGTGGCCTTGCAGGGGGGAGAGCCTGCCTTCTCTTCCGCA-3’

This 502bp sequence matches the wild-type sequence of the *LMNA* gene *locus* where the base that gets mutated in HGPS (*i.e.* C) is in the center, except for the PAM site that was altered by site-directed mutagenesis from CCA to GCA. This alteration was performed to prevent the sgRNA would cutting the repair template during the editing, and also to allow one to distinguish by Sanger sequencing, upon genotyping, the corrected alleles (by the repair template) from wild-type alleles.

### High-content transcriptome analysis (single-cell RNASeq): methanol-fixation

Prior to processing and transcript sequencing, 8.9×10^5^ and 1.1×10^6^ ISO- and HGPS-iPSC-CMs, respectively, were collected and then methanol-fixed. Briefly, first cells were centrifuged (300*g*, 5 min, RT) and washed with PBS twice, and the supernatants were discarded. Thereafter, chilled PBS (ca. 4°C) was added to the pellets (200 μL per 1×10^6^ cells) followed by drop by drop addition of 100% methanol (ca. −20°C) (800 μL per 1×10^6^ cells). Cells were then incubated at −20°C for 30 min and stored at −80°C until further processing. For rehydration, methanol fixed cells were first equilibrated to 4°C, centrifuged (1000*g*, 5 min, 4°C), the supernatant discarded and the resuspended in a Wash-Resuspension buffer from 10x Genomics. Cells were then loaded into a 10x Genomics system for processing and sequencing.

### High-content transcriptome analysis (single-cell RNASeq): 10x processing and sequencing

Libraries were prepared by using a 10x 3’ mRNA v3.1 kit. They were sequenced on a NovaSeq Illumina sequencer to a depth of at least 50K reads/ cell, following manufacturer’s instructions. Low pass sequencing was used to check how many cells were captured and to confirm the appropriate number of cells and reads per cell were obtained. These samples were sequenced 3 times to obtain 50K reads/cell.

### High-content transcriptome analysis (single-cell RNASeq): bioinformatic analysis

Sample pre-processing (Cell Ranger): sample-level count data were generated using 10x Genomics Cell Ranger 6.0.1 0.0 with bcl2fastq-2.20.0 against 10X human reference dataset refdata-gex-GRCh38-2020-A. The resulting filtered feature-barcode matrices were used as input into R for further analysis using the Seurat package with default settings unless otherwise stated [54].

Sample QC: simple QC metrics were derived based on the total counts per cell, the number of expressed genes per cell and each gene’s percentage contribution to a total cell’s counts. The percentage of cell counts from mitochondrial genes and separately from ribosomal genes was also calculated. Metrics were plotted as distributions to manually identify outlier cells that failed QC. Cells with a mitochondrial percentage >20%, where a single gene contributed >20%, or where the total number of detected genes was < 1000 or > 6000 were flagged as outliers and discarded from further analysis.

Sample Processing (Seurat): count matrices were then normalised per cell using a centered log ratio (CLR) transformation [Seurat::NormalizeData(normalization.method = “CLR”, margin = 2)]. The top 2000 most variable genes that exhibited high cell-to-cell variation in the dataset were identified [Seurat::FindVariableFeatures(selection.method = “vst”, nfeatures = 2000)]. Data were scaled and centred using all available features [Seurat::ScaleData] such that the mean expression across cells was 0 and the variance was 1 in order to ensure that highly-expressed genes did not dominate downstream analysis. PCA dimensionality reduction was run using the 2000 most variable genes to assess the dimensionality of the dataset. The non-linear dimensional reduction technique UMAP was then applied using the first 50 PC so that similar cells were placed together in low-dimensional space. A k-nearest neighbours (KNN) graph was constructed from the Euclidean distance in PCA space, refining edge-weights between cells based on shared overlap of local neighborhoods (Jaccard similarity) [Seurat::FindNeighbors(dims = 50, k.param = 20, compute.SNN = TRUE)]. The KNN-graph was used to group cells into clusters showing similar expression profiles via the Louvain algorithm [Seurat::FindClusters(resolution=0.5)]. The effects of cell cycle heterogeneity were assessed by scoring each cell based on the expression of the default canonical G2/M and S phase marker genes from Tirosh et al. [55] loaded with Seurat [Seurat::CellCycleScoring]. Potential doublet cells were flagged, though not removed, using the R package DoubletFinder’s doubletFinder_v3 function [56], assuming a doublet rate of 7.5%.

Sample integration (Seurat): after each sample dataset had been separately processed, a set of integration features was selected that represented genes repeatedly variable across them all [Seurat::SelectIntegrationFeatures]. Integration anchors, i.e. cross-dataset pairs of cells matched in biological state, were identified using reciprocal PCA (rPCA) focussing on just the integration features. The anchors were used to integrate the sample datasets into a single object using the Seurat::IntegrateData function with default settings. The integrated data were then centred and scaled, before PCA and UMAP dimensionality reduction, KNN-graph construction and Louvain clustering using the same parameters as for the single samples (see above). The number of cells from each cell contributing to each cluster was reported.

Cluster marker gene identification: positive marker genes differentially expressed in each cluster relative to all others were identified using a Wilcoxon rank sum test, thresholding on a log-scale fold-change of 0.25 and a p-value < 0.01. Only genes expressed in 0.25% of cells were included in testing. [Seurat::FindAllMarkers(assay=“RNA”,only.pos = TRUE, min.pct = 0.25, logfc.threshold = 0.25, random.seed = 1, return.thresh = 0.01)]. The top 10 genes based on average logFC were highlighted.

Cluster enrichment analysis: enrichment analysis of biological terms mapped to genes differentially expressed in each cluster was assessed against a genomic background using i) Reactome and ii) Gene Ontology (GO) Biological Process (BP) classifications via the ClusterProfiler package’s compareClusters function [57]. Genes were mapped to Entrez gene ids. Only gene-sets between 10-500 in size were considered. A Benjamini-Hochberg (BH) multiple testing correction was applied. Results were thresholded at a p-value < 0.05 and a q-value < 0.1. The top 5 most enriched terms based on the overlap of cluster/term membership (“GeneRatio”) were highlighted.

Feature plots: the normalised abundance of selected genes was overlaid on top of the integrated sample UMAP dimension reduction using Seurat’s FeaturePlot function (slot=”data”).

Dotplots: dotplots were used to visualise how selected genes’ normalised expression altered across clusters using Seurat’s DotPlot function (assay=”RNA”). Normalised expression values were exponentiated prior to averageing so that averageing was done in non-log space. The size of the dots encodes the percentage of cells within a cluster, while the colour encodes the average expression level across all cells within a cluster.

Senescence markers: a table of senescence-associated genes was downloaded from the CellAge website [27] to assess changes in expression across cells. Genes were stratified into 2 groups based on their “Senescence Effect” being classed as either “Induces” or “Inhibits” by CellAge. https://genomics.senescence.info/cells/

Expression of LMNA isoforms: this analysis was an attempt to get isoform specific counts for 3 specific transcript isoforms of the LMNA gene which differ at their 3’ ends. It was conducted by running a second iteration of cellranger count, replacing the original LMNA gene in the standard gtf file with three new “genes” (details below), representing the 3 isoforms of interest. Because the isoforms overlap in genomic space and share multiple common exons this analysis was somewhat compromised. Reads mapping equally well to multiple isoforms were discarded, hence the counts for the individual isoforms were not comparable to the parent gene. This type of analysis was not well suited to 3’ sequencing and so the results should only be used in an observational sense.

LMNA, the original LMNA gene and all its many isoforms.

LMNA-A, LaminA, LMNA-205, ENST00000368300.9

LMNA-C, LaminC, LMNA-206, ENST00000368301.6

LMNA-P, Progerin, LMNA-204, ENST00000368299.7

### Mitochondrial respiration assessment

Day 20 CMs were plated in Seahorse XF 96 Cell Culture microplates (Agilent), previously coated with Matrigel, at a density of 2.85×10^4^ cells/well. On day 30, the CMs had their oxygen consumption rates (OCR) measured and determined resorting to a Seahorse XFe96 analyzer (Agilent). For the XF Cell Mito Stress kit (Agilent) assay, the plate was washed three times with a basal medium composed of Seahorse XF RPMI medium, pH 7.4 (103576-100; Agilent), D-(+)-Glucose (11 mM; G7021; Sigma), L-Glutamine (3 mM; G3126; Sigma) and L-Ascorbic acid (65 µg/mL; A92902; Sigma). Then, B-27™ Supplement, minus vitamin A (Thermo Fisher Scientific) was added to the wells and the cells were incubated for 1 h at 37°C. OCR measurements were obtained in different respiratory states by the sequential injection of oligomycin (1.76 µM; Sigma), FCCP (1.77 µM; Sigma), and rotenone/antimycin (0.88 µM; Sigma). Following the assay, CMs were fixed with 4 % PFA (10 min; RT), stained with DAPI (2 µg/mL; D9542; Sigma) (5 min; RT), and the nuclei were counted. OCR values were normalized for 1×10^3^ nuclei and results were analyzed using the Wave Desktop Version 2.6.3 software (Agilent).

### Calcium imaging: calcium-sensitive dye loading procedure

Prior to calcium imaging, day 20 ISO- and/or HGPS-iPSC-CMs were plated in a 96-well µ-Plate (ibidi) and then drug-treated for 8 to 10 days. iPSC-CMs were incubated (30 min, 37°C) with Fluo-4, AM, cell permeant (4 µM; F14201; Invitrogen^TM^) in a mixture of conditioned medium and an equivalent medium without Phenol Red (1:1). After incubation, the medium was replaced with a mixture of conditioned medium and fresh Phenol Red-free medium (1:1). Cells were then imaged.

### Calcium imaging: live transient imaging and acquisition

Acquisitions were performed in a Zeiss Cell Observer SD spinning disk confocal microscope (Carl Zeiss), in controlled conditions of temperature (37°C) and humidity, using a 488 nm laser and 20x objective. Three 90 second-long recordings were obtained per well of each tested condition.

### Calcium imaging: transient data extraction and analysis

Fluorescence calcium transient data extraction from the recordings and further processing were performed resorting to the software MATLAB® (version R2021a; MathWorks®) in combination with an adjusted script of CalTrack, an open-source tool that allows for the high-throughput automated calcium transient analysis in CMs [60]. Regarding the analysis, in brief, by means of fluorescence-based segmentation, CalTrack allowed for the detection of several CMs per recording, whose data was further used. Recordings where less than three CMs were detected, were discarded from the analysis. The fluorescence background was automatically determined and subtracted from the signals. Furthermore, CalTrack performed photobleaching corrections and excluded noisy signals. From the full signals, the calcium transient rate was determined, and the average calcium transient cycle was obtained that, in turn, was used for plotting and calculating the following parameters: transient maximum (peak of the transient relative to the background), transient duration, rise time (*i.e.* the time that it takes for the calcium intracellular levels to increase from the initial baseline level to 90% of the transient maximum, corresponding to the upstroke) and the tau (r) decay constant (*i.e.* the time that it takes for the calcium intracellular levels to decrease from the transient maximum to 37% of the transient maximum, corresponding to the initial downstroke).

### *In vitro* drug treatments

Day 20 HGPS-iPSC-CMs were plated in the desired plate format and on day 21, the medium was replaced with one without Y-27632 dihydrochloride and left without manipulation for that day. On day 22 of differentiation, the chronic low-dose treatment was initiated. The medium was supplemented with the DMSO (control) or the tested drug compounds, and the medium + growth factors + drugs mixture was refreshed every other day until the CMs were characterized on day 30 of differentiation. The compounds tested were the following: i) DMSO (0.4%) as the untreated control; ii) rapamycin (4 nM; Sigma); iii) lonafarnib (Lona) (8 µM; Selleckchem); iv) MG-132 (250 nM; Calbiochem); v) fisetin (7.5 µM; Hangzhou Yuechen Chemical); vi) the combination of dasatinib (16 nM; Hangzhou Yuechen Chemical) and quercetin (3.2 µM; Hangzhou Yuechen Chemical) (D+Q); and vii) neuropeptide Y (NPY) (60 nM; Phoenix Pharmaceuticals). The aforementioned concentrations were selected based on: i) concentrations from previous reports where these compounds were used with single-administration and in a short-term timeframe [10, 11, 14, 40, 41]; and by performing two cell viability assays: a short-term, 48-h treatment test and then a long-term, 10-day treatment test, performed as with the aforementioned final methodology. The 48 h concentration test was performed by treating ISO- and HGPS-iPSC-CMs (day 22) once with DMSO (0.4%), Rapa (2 and 4 nM), Lona (4 and 8 µM), MG-132 (500 nM and 2 µM), fisetin (2 and 20 µM), D+Q (8 nM of D + 1.6 µM of Q; and 32 nM of D + 6.4 µM of Q), and NPY (10 and 40 nM). These concentrations correspond to 10% and 40% of the concentrations found in the literature for single-dose treatments. Then, 48 h after treatment, we resorted to the CellTiter-Glo® Luminescence Cell Viability Assay (Promega), according to the manufacturer’s instructions. Briefly, cells and culture medium contents were equilibrated for 30 min at RT, and then one part of the CellTiter-Glo® was added to induce cell lysis and incubated for 2 min with agitation. Then, the lysate mix was added to an opaque 96-well multiwell plate and left at RT for 10 min to stabilize the luminescence signal. Luminescence was acquired in a plate reader (Biotek). Cell viability data were normalized for DMSO. For the 10-day drug concentration test, ISO- and HGPS-iPSC-CMs (day 22) were chronically treated with drug refreshing every other day, as previously explained, with the following concentrations: DMSO (0.4%), Rapa (2 and 4 nM), Lona (4 and 8 µM), MG-132 (125 and 250 nM, fisetin (2 and 10 µM), D+Q (8 nM of D + 1.6 µM of Q; and 16 nM of D + 3.2 µM of Q), and NPY (10 and 40 nM). After the 8 to 10 days, cells were fixed as previously described, their nuclei stained with DAPI and counted by means of an IN Cell Analyzer 2200 (GE Healthcare) equipment. For data analysis of the parameters evaluated in drug-treated HGPS-iPSC-CMs, values were normalized with the untreated (DMSO) control within each experiment/differentiation batch.

### Sample preparation for proteomics analysis of drug-treated cells

Pellets from untreated ISO-iPSC-CMs, untreated HGPS-iPSC-CMs and HGPS-iPSC-CMs treated with Rapa, Lona, MG-132, fisetin, D+Q and NPY were processed for quantitative proteome analysis. For proteomics analysis, 1 Mio cells (equals approximately 100 µg protein) was harvested for each sample and was resuspended in 100 µl PBS and lysis buffer (fc 4% SDS, 100 mM HEPES, pH 8.5, 50 mM DTT). Samples were then sonicated in a Bioruptor (Diagenode, Belgium) (10 cycles with 1 minute on and 30s off with high intensity @ 20°C). Samples were then heated at 95 °C for 10 minutes, before being subjected to another round of sonication in the Bioruptor. The lysates were clarified and debris precipitated by centrifugation at 14000 rpm for 10 minutes, then incubated with iodacetamide (room temperature, in the dark, 20 minutes, 15 mM). 5% of the sample was removed to check lysis on a coomassie gel. Based on the gel, an estimated 30 µg of each sample was treated with 4 volumes ice-cold acetone and left overnight at −20 °C to precipitate the proteins. The samples were then centrifuged at 14000 rpm for 30 minutes, 4 °C. After removal of the supernatant, the precipitates were washed twice with 300 µL of a solution of ice-cold 80 % acetone. After the addition of each wash solution, the samples were vortexed and centrifuged again for 10 minutes at 4°C. The pellets were then allowed to air-dry before being dissolved in a digestion buffer at 1 µg/µL (1M guanidine HCl in 0.1M HEPES, pH 8). To facilitate the resuspension of the protein pellet, the samples were subjected to 5 cycles of sonication in the Bioruptor, as described above. Afterwards, LysC (Wako) was added at 1:100 (w/w) enzyme : protein ratio and digestion proceeded for 4 h at 37 °C under shaking (1000 rpm for 1 h, then 650 rpm). The samples were diluted 1:1 with milliQ water (to reach 1.5M urea) and were incubated with a 1:100 w/w amount of trypsin (Promega sequencing grade) overnight at 37 °C, 650 rpm. The digests were then acidified with 10% trifluoroacetic acid and then desalted with Waters Oasis® HLB µElution Plate 30µm in the presence of a slow vacuum. In this process, the columns were conditioned with 3×100 µL solvent B (80% acetonitrile; 0.05% formic acid) and equilibrated with 3×100 µL solvent A (0.05% formic acid in milliQ water). The samples were loaded, washed 3 times with 100 µL solvent A, and then eluted into PCR tubes with 50 µL solvent B. The eluates were dried down and before analysis samples were reconstituted in in MS Buffer (5% acetonitrile, 95% Milli-Q water, with 0.1% formic acid) and spiked with iRT peptides (Biognosys, Switzerland).

### LC-MS Data independent analysis (DIA)

Peptides were separated in trap/elute mode using the nanoAcquity MClass Ultra-High Performance Liquid Chromatography system (Waters, Waters Corporation, Milford, MA, USA) equipped with a trapping (nanoAcquity Symmetry C18, 5 μm, 180 μm × 20 mm) and an analytical column (nanoAcquity BEH C18, 1.7 μm, 75 μm × 250 mm). Solvent A was water and 0.1% formic acid, and solvent B was acetonitrile and 0.1% formic acid. 1 µl of the sample (∼1 μg on column) was loaded with a constant flow of solvent A at 5 μl/min onto the trapping column. Trapping time was 6 min. Peptides were eluted via the analytical column with a constant flow of 0.3 μl/min. During the elution, the percentage of solvent B increased in a nonlinear fashion from 0–40% in 120 min. The total run time was 145 min, including equilibration and conditioning. The LC was coupled to an Orbitrap Exploris 480 (Thermo Fisher Scientific, Bremen, Germany) using the Proxeon nanospray source. The peptides were introduced into the mass spectrometer via a Pico-Tip Emitter 360-μm outer diameter × 20-μm inner diameter, 10-μm tip (New Objective) heated at 300 °C, and a spray voltage of 2.2 kV was applied. The capillary temperature was set at 300°C. The radio frequency ion funnel was set to 30%. For DIA data acquisition, full scan mass spectrometry (MS) spectra with a mass range 350–1650 m/z were acquired in profile mode in the Orbitrap with a resolution of 120,000 FWHM. The default charge state was set to 3+. The filling time was set at a maximum of 60 ms with limitation of 3 × 10^6^ ions. DIA scans were acquired with 40 mass window segments of differing widths across the MS1 mass range. Higher collisional dissociation fragmentation (stepped normalized collision energy; 25, 27.5, and 30%) was applied and MS/MS spectra were acquired with a resolution of 30,000 FWHM with a fixed first mass of 200 m/z after accumulation of 3 × 10^6^ ions or after filling time of 35 ms (whichever occurred first). Data were acquired in profile mode. For data acquisition and processing of the raw data Xcalibur 4.3 (Thermo) and Tune version 2.0 were used.

### Proteomic data processing

DIA raw data were analyzed using the directDIA pipeline in Spectronaut v.16 (Biognosys, Switzerland) with BGS settings besides the following parameters: Protein LFQ method= QUANT 2.0, Proteotypicity Filter = Only protein group specific, Major Group Quantity = Median peptide quantity, Minor Group Quantity = Median precursor quantity, Data Filtering = Qvalue, Normalizing strategy = Local Normalization. The data were searched against a UniProt (Homo Sapiens, 20,375 entries, v. 160126) and a contaminants (247 entries) database. The identifications were filtered to satisfy FDR of 1 % on peptide and protein level. Relative protein quantification was performed in Spectronaut using a pairwise t-test performed at the precursor level followed by multiple testing correction. For further proteome data analysis, we resorted to the IPA software (Qiagen) to obtain the altered canonical pathways and respective activation z-score. Cross-referencing with the senescence gene database CellAge [27] was performed. Bubble plot representation was obtained using SRplot (http://www.bioinformatics.com.cn/srplot). Cross-referencing with the GO database, using GO:0006914 autophagy term was also performed.

### *In vivo* analyses: mouse strains

The HGPS mouse model carrying a p.Gly609Gly mutation in the Lmna gene (LAKI - Lmna^G609G/G609G^) was kindly provided by Carlos López-Otín [42]. Procedures involving animals and their care were conducted in compliance with institutional ethical guidelines (i3S Animal Welfare and Ethics Review Body, ORBEA) and with the National and European Union rules (2010/63/EU), under DGAV license (DGAV 0421/000/000/2017). Mice were kept on pathogen-free barrier areas, under a 12 h:12 h light–dark cycle, at the i3S animal facility. Experiments were performed using mice of both sexes, randomly assigned to control and experimental groups.

### *In vivo* analyses: lonafarnib treatment

The farnesyltransferase inhibitor lonafarnib (kindly provided by The Progeria Research Foundation) was mixed thoroughly in DietGel® Boost (Clear H2O) to a final concentration of 450 mg per kg of soft diet [61]. Mice were fed daily a fresh 3g dose of diet placed on floor level, starting at 5 weeks of age. The vehicle control consisted of a soft diet without lonafarnib. Previous work showed that mixing the drug in the mouse diet allows the maintenance of steady-state drug levels. Mice were weighted weekly from the start of the treatment until euthanasia by decapitation at 13 weeks of age. Hearts were snap-frozen in liquid nitrogen and plasma was obtained by centrifugation of whole blood at 2,000 g for 10 min.

### *In vivo* analyses: sample preparation for total proteome

The methodology used for the protein sample isolation, preparation and analysis is similar to previously published [35]. Snap-frozen hearts were thawed and transferred into Precellys® lysing kit tubes (Keramik-kit 1.4/2.8□mm, 2□mL (CKM)) containing PBS supplemented with cOmplete™, Mini, EDTA-free Protease Inhibitor (Roche,11836170001). The volume of PBS added was calculated based on an estimated protein content (5% of fresh tissue weight) to reach a 10 µg/µL concentration. Tissues were homogenized twice at 6000□rpm for 30□s using Precellys® 24 Dual (Bertin Instruments, Montigny-le-Bretonneux, France), and the homogenates were transferred to new 2□mL Eppendorf tubes. Samples corresponding to ∼150 µg of protein were used as starting material for each biological replicate. Volumes were adjusted using PBS and samples were lysed by the addition of 4× lysis buffer (8% SDS, 100 mM HEPES, pH8). Samples were sonicated twice in a Bioruptor Plus for 10 cycles with 1□min ON and 30□s OFF with high intensity at 20□°C. The lysates were centrifuged at 18,407 xg for 1□min and transferred to new 1.5□mL Eppendorf tubes. Subsequently, samples were reduced using 20□mM DTT (Carl Roth, 6908) for 15□min at 45□°C and alkylated using freshly made 200□mM iodoacetamide (IAA) (Sigma-Aldrich, I1149) for 30□min at room temperature in the dark. An aliquot of each lysate was used for estimating the precise protein quantity using Qubit protein assay, and volumes were adjusted accordingly to have a total protein amount of 100 µg (Thermo Scientific, Q33211). Subsequently, proteins were precipitated using cold acetone and resuspended in digestion buffer (3 M urea, 100 mM HEPES pH 8.0). Proteins were digested using LysC 1:100 enzyme:proteins ratio for 4 h (Wako sequencing grade, 125-05061) and trypsin 1:100 enzyme:proteins ratio for 16 h (Promega sequencing grade, V5111). The digested proteins were then acidified with 10% (v/v) trifluoroacetic acid. Peptides were desalted using Waters Oasis® HLB µElution Plate 30□µm following manufacturer instructions. The eluates were dried down using a vacuum concentrator and reconstituted in MS buffer A (5% (v/v) acetonitrile, 0.1% (v/v) formic acid).

### *In vivo* analyses: TMT labelling for proteomic analyses

The solution containing the resuspended peptides was brought to a pH of 8.5 and a final concentration of 100 mM HEPES (Sigma H3375) prior to labeling. 20 µg of peptides were used for each label reaction. TMT-10plex reagents (Thermo Fisher #90111) were reconstituted in 41 µL of acetonitrile (Biosolve #0001204102BS). TMT labeling was performed in two steps by the addition of 2x of the TMT reagent per mg of peptide (e.g., 40 µg of TMT reagent for 20 µg of peptides). TMT reagents were added to samples at room temperature, followed by incubation in a thermomixer (Eppendorf) under constant shaking at 600 rpm for 30 min. After incubation, a second portion of TMT reagent was added and followed by incubation for another 30 min. After checking the labeling efficiency by MS, equal amounts of samples were pooled (200 µg total), desalted using two wells of a Waters Oasis HLB mElution Plate 30 mm (Waters #186001828BA) and subjected to high pH fractionation before MS analysis.

### *In vivo* analyses: high pH peptide fractionation for proteomic analyses

Offline high pH reverse phase fractionation was performed using an Agilent 1260 Infinity HPLC System equipped with a binary pump, degasser, variable wavelength UV detector (set to 220 and 254 nm), peltier-cooled autosampler (set at 10C) and a fraction collector. The column used was a Waters XBridge C18 column (3.5 mm, 100 3 1.0 mm, Waters) with a Gemini C18, 4 3 2.0 mm SecurityGuard (Phenomenex) cartridge as a guard column. The solvent system consisted of 20 mM ammonium formate (20 mM formic acid (Biosolve #00069141A8BS), 20 mM (Fluka #9857) pH 10.0) as mobile phase (A) and 100% acetonitrile (Biosolve #0001204102BS) as mobile phase (B). The separation was performed at a mobile phase flow rate of 0.1 mL/min using a non-linear gradient from 95% A to 40% B for 91 min. Forty-eight fractions were collected along with the LC separation and subsequently pooled into 24 fractions. Pooled fractions were dried in a speed vacuum centrifuge and then stored at −80 °C until MS analysis.

### *In vivo* analyses: data acquisition for TMT-labelled samples (proteomic analyses)

For TMT experiments, fractions were resuspended in 20 µL reconstitution buffer (5% (v/v) acetonitrile (Biosolve #0001204102BS), 0.1% (v/v) TFA in water) and 5 µL were injected into the mass spectrometer. Peptides were separated using the nanoAcquity UPLC system (Waters) fitted with a trapping (nanoAcquity Symmetry C18, 5 µm, 180 µm x 20 mm) and an analytical column (nanoAcquity BEH C18, 2.5 µm, 75 µm x 250 mm). The analytical column outlet was coupled directly to an Orbitrap Fusion Lumos (Thermo Fisher Scientific) using the Proxeon nanospray source. Solvent A was water with 0.1% (v/v) formic acid and solvent B was acetonitrile, 0.1% (v/v) formic acid. The samples were loaded with a constant flow of solvent A at 5 µL/min, onto the trapping column. Trapping time was 6 min. Peptides were eluted via the analytical column at a constant flow rate of 0.3 µL/ min, at 40 °C. During the elution step, the percentage of solvent B increased in a linear fashion from 5% to 7% in the first 10 min, then from 7% B to 30% B in the following 105 min and to 45% B by 130 min. The peptides were introduced into the mass spectrometer via a Pico-Tip Emitter 360 µm OD x 20 µm ID; 10 mm tip (New Objective) and a spray voltage of 2.2kV was applied. The capillary temperature was set at 300 °C. Full scan MS spectra with a mass range of 375-1500 m/z were acquired in profile mode in the Orbitrap with a resolution of 60000 FWHM using the quad isolation. The RF on the ion funnel was set to 40%. The filling time was set to a maximum of 100 ms with an AGC target of 4 x10^5^ ions and 1 microscan. The peptide monoisotopic precursor selection was enabled along with relaxed restrictions if too few precursors were found. The most intense ions (instrument operated for a 3 s cycle time) from the full scan MS were selected for MS2, using quadrupole isolation and a window of 1 Da. HCD was performed with a collision energy of 35%. A maximum fill time of 50 ms for each precursor ion was set. MS2 data were acquired with a fixed first mass of 120 m/z and acquired in the ion trap in Rapid scan mode. The dynamic exclusion list was set with a maximum retention period of 60 s and a relative mass window of 10 ppm. For the MS3, the precursor selection window was set to the range 400-2000 m/z, with an exclusion width of 18 m/z (high) and 5 m/z (low). The most intense fragments from the MS2 experiment were co-isolated (using Synchronus Precursor Selection = 8) and fragmented using HCD (65%). MS3 spectra were acquired in the Orbitrap over the mass range of 100-1000 m/z and the resolution was set to 30000 FWHM. The maximum injection time was set to 105 ms and the instrument was set not to inject ions for all available parallelizable times. The Xcalibur v4.0 and Tune v2.1 were used to acquire and process raw data.

### *In vivo* analyses: data processing for TMT labelled samples

TMT-10plex data were processed using Spectromine v4.4 (Biognosys). Data were searched against the relevant species-specific fasta database (Uniprot database, Swissprot entry only, release 2016_01 for Mus musculus, 16748 entries) with an appended list of common contaminants. The data were searched with the following modifications: carbamidomethyl (C) as fixed modification, and oxidation (M), acetyl (protein N-term), as variable modifications, and TMT_Lys and TMT_Nter for TMT label reagents. A maximum of 2 missed cleavages were allowed. MS3 quantitation was allowed. Global normalization was applied using the median and single hits proteins were excluded. Fraction aggregation was performed as sum. Relative quantification was performed in Spectromine for each pairwise comparison using the replicate samples from each condition using default settings. Candidates and report tables were exported from Spectromine for downstream analysis.

### *In vivo* analyses: RNA *in situ* hybridization (RNAish) in LV tissue

RNA in situ hybridization was performed on sections from FFPE fragments of LV muscle of Untreated WT mice, and Untreated and Lona-treated LAKI mice, resorting to a RNAscope Multiplex Fluorescent Assay v2 (Advanced Cell Diagnostics) kit. First, the paraffin sections were baked in a dry oven (1 h, 60°C). Afterwards, the sections were deparaffinized by immersion in xylene (2×5 min), followed by absolute ethanol (2×2 min). Then, sections were left to dry at RT. After drying, the sections were incubated with RNAscope Hydrogen Peroxide (Advanced Cell Diagnostics) (10 min, RT). For target retrieval, sections were incubated in Target Retrieval Solution (Advanced Cell Diagnostics) (30 min, 100°C), washed with distilled water (15 sec), incubated with ethanol (3 min, RT), and left to dry (5 min, RT). Following the drawing of a hydrophobic barrier around the sections - using an ImmEdge® Hydrophobic Barrier PAP pen (Vector Laboratories) – the sections were then incubated (30 min, 40°C) in a HybEZ™ II Hybridization System (Advanced Cell Diagnostics) with RNAscope Protease Plus (Advanced Cell Diagnostics) and washed with distilled water. For hybridization, sections were incubated (2 h, 40°C) with the RNAscope probe that targets the gene encoding p21 (i.e. *Cdkn1a*) (Mm-Cdkn1a, 408551, Advanced Cell Diagnostics). Sections were then rinsed twice with agitation in the Wash buffer (Advanced Cell Diagnostics) (2 min, RT). To amplify the signal, sections were incubated sequentially in RNAscope Multiplex FL v2 Amp1 (30 min, 40°C), RNAscope Multiplex FL v2 Amp2 (30 min, 40°C), RNAscope Multiplex FL v2 Amp3 (15 min, 40°C), and RNAscope Multiplex FL v2 HRP-C1 (15 min, 40°C), with 2×2 min (RT) washes in Wash buffer in between those incubations. For developing the HRP-C1 signal, sections were incubated with Opal 570 (FP1488001KT, Akoya Biosciences) (1:1000, 30 min, 40°C) diluted in RNAscope Multiplex TSA buffer (Advanced Cell Diagnostics). After washing with Wash buffer (2×2 min, RT), the sections were blocked with RNAscope Multiplex FL v2 HRP blocker (Advanced Cell Diagnostics) (15 min, 40°C) and washed with Wash buffer (2×2min, RT). Nuclei were stained with DAPI (2 μg/mL) (5 min, RT) and the cell membrane with Wheat Germ Agglutinin (WGA) (ThermoFisher) (10 μg/mL, 30 min, RT). Finally, sections were mounted with Fluorescent mounting medium (Dako). Capture was carried out in LSM 710 confocal microscope (ZEISS) with the Plan-Apochromat 40×/1,4 OIL DIC M27 objective.

P21 *foci* were counted manually in the Qupath Software resorting the “add points” functionality.

### *In vivo* analyses: γH2AX staining in PCM-1 positive cells in LV tissue

Cryosections were permeabilized with 1% Triton for 15 min, washed 3× PBS 1×, and blocked for 1 h RT with blocking solution (5% BSA + 1:60 NGS + 0,01% Triton), and incubated with anti-Phospho-Histone H2Ax ab (1:100 cell signaling, ref 9718). On the following day, the samples were then washed 3× in PBS 1× for 5 min RT, and incubated with secondary Goat anti-Rabbit IgG, Alexa Fluor™ 488 (1:500, Invitrogen, cat no. A11034). The tissue was then fixated for 20 min with 4% PFA and incubated with PCM1 (1:100, Santa Cruz Biotechnology, ref sc-398365), washed and incubated with secondary Goat anti Mouse IgG, Alexa Fluor 633 (1:500, Invitrogen, ref. A21052). The samples were stained with DAPI (2 µg/mL, 20 min) and the slides mounted with Fluorescent Mounting Media (Dako, ref 2S302380-2) and let dry for 24 h. All the antibodies were diluted in blocking solution and incubated 16-24 h at 4 °C, and the sample washed between every step in 1× PBS for 5 min RT. Capture was carried out in LSM 710 confocal microscope (ZEISS) with the Plan-Apochromat 40×/1,4 OIL DIC M27 objective. Capture was carried out in LSM 710 confocal microscope (ZEISS) with the Plan-Apochromat 40×/1,4 OIL DIC M27 objective.

γH2AX *foci* were counted manually in the Qupath Software resorting the “add points” functionality.

### Acute treatment of iPSC-CMs with Lona for viability, senolytic activity, prelamin A accumulation and autophagy assessments

For assessment of the acute effect of Lona in viability and senolytic potential, ISO- and HGPS-iPSC-CMs were treated for 24h with Lona (20 µM) while an Untreated condition (0 µM) was used as a control and to normalize all the date. Viability/senolytic potential by performing automatic nuclei count, following cell fixation (4% PFA, 10 min., RT) and nuclei staining (DAPI, 2 µg/mL, 5 min.). Nuclei count was then normalized to the Untreated condition of each cell line.

For the validation of the expected activity of Lona only in HGPS-iPSC-CMs, these cells were also acutely treated with Lona (20 µM, 24h) and then cells were fixed (4% PFA, 10 min., RT), immunostained for prelamin A (see section *Immunocytochemistry* from *Materials and Methods*). Finally, nuclear fluorescence intensity corresponding to prelamin A was automatically quantified.

For the assessment of autophagic activitiy in ISO- and HGPS-iPSC-CMs, cell were treated continuously for 27 h with Lona (20 μM) and for the last 3 h with chloroquine (Chq) (an autophagy inhibitor) between the 24 h and 27 h timepoints, i.e. in combination with the Lona. Then, cells were collected and Western blot was performed (see section *11.2 Western blot for autophapgy marker* from *Materials and Methods). Autophagic flux was assessed by* the LC3B turnover assay measures the amount of LC3B-II delivered to the lysosomes by comparing the LC3B-II amount in the presence and absence of the lysosomal inhibitor chloroquine by Western blotting. For each experimental condition, untreated HGPS-iPSC-CMs and HGPS-iPSC-CMs treated with Lona, “LC3B-II flux,” was determined by subtracting the densitometric value of the LC3B-II band of the chloroquine-untreated sample (ChQ-LC3B-II) from the densitometric value of the LC3B-II band of the corresponding chloroquine-treated sample (ChQ + LC3B-II). If autophagic flux is stimulated, the amount of LC3B-II will be higher in the presence of lysosomal inhibitor (ChQ). Conversely, if the LC3B-II protein does not increase in the presence of ChQ, it indicates that autophagic flux is impaired, and a defect or delay earlier in the process, prior to degradation at the autolysosome, has occurred. For each independent experiment, the values obtained upon subtraction for each condition were normalized to the control condition (untreated HGPS-iPSC-CMs cells). The results are represented as mean values for each experimental condition.

### General statistical analysis

Values shown in text and figures are mean ± standard error of the mean (SEM). Data statistical analysis was executed by resorting to GraphPad Prism version 8 for Mac (Graphpad Software, www.graphpad.com). Outliers were detected using the ROUT method (using Q = 10%) and then excluded from the statistical analysis. Shapiro-Wilk’s test was used to evaluate if the data displayed a normal distribution. If so, the homoscedasticity of the data was tested by an F test. These results defined the statistical test(s) used further. Normal distributed and homoscedastic data were tested with parametric tests (independent samples Student’s t-test or ordinary one-way ANOVA for two or more conditions, respectively). Non-normal distributed and/or heteroscedastic data were tested with non-parametric tests (Mann-Whitney Test or Kruskal-Wallis test for two or more conditions, respectively). For more than two conditions, post-hoc tests for the correction of multiple comparisons were performed (Dunnett’s multiple comparison tests for normal distributed and homoscedastic data and Dunn’s multiple comparison tests for non-normal distributed and/or heteroscedastic data. “N” represents the number of independent experiments and/or differentiation batches, while “n” represents the number of quantified images, wells or technical replicates. The statistical significance level chosen for all statistical tests was *p* < 0.05.

---

**Extended Data Fig. 1.**
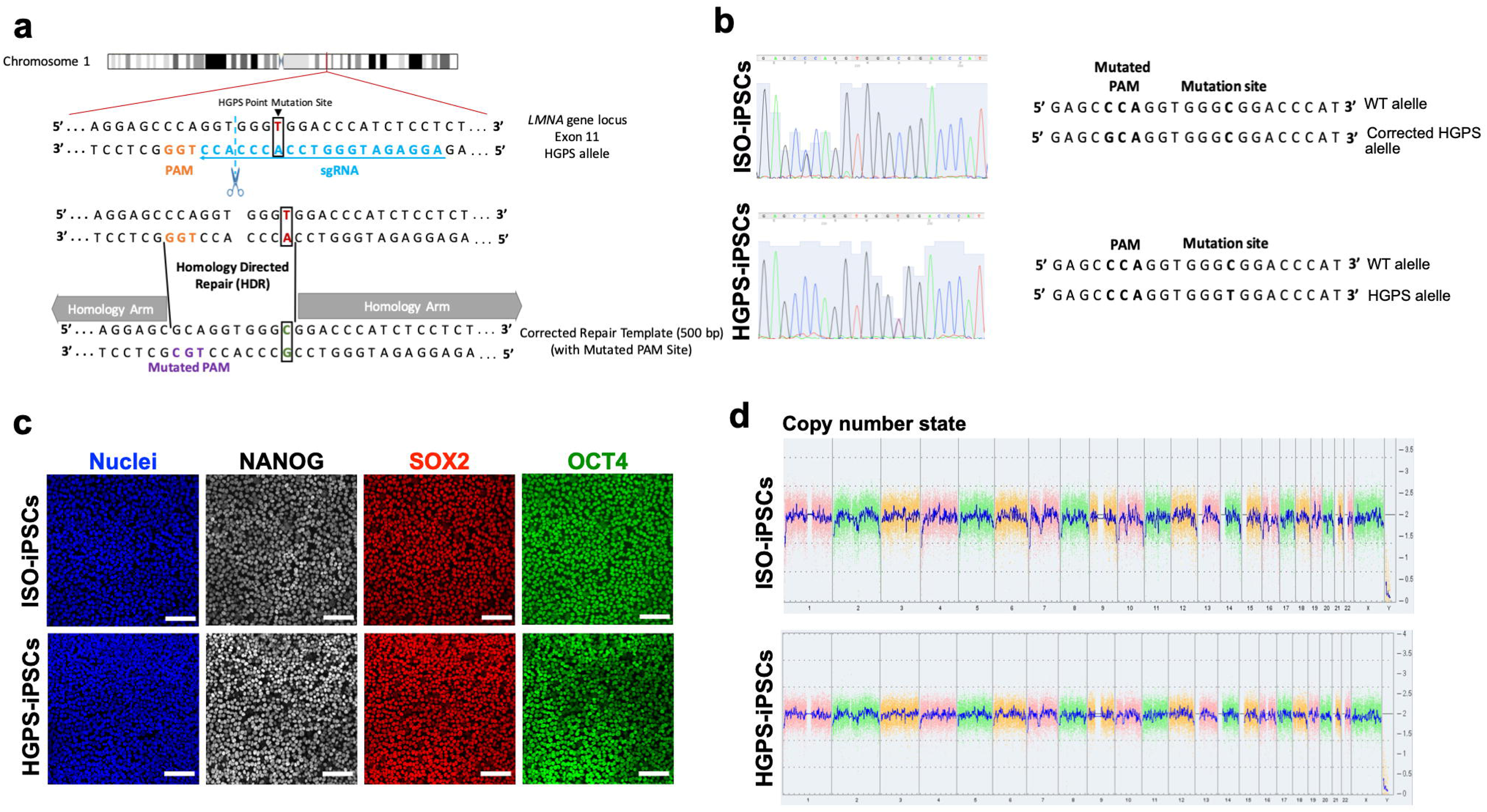

**Extended Data Fig. 2.**
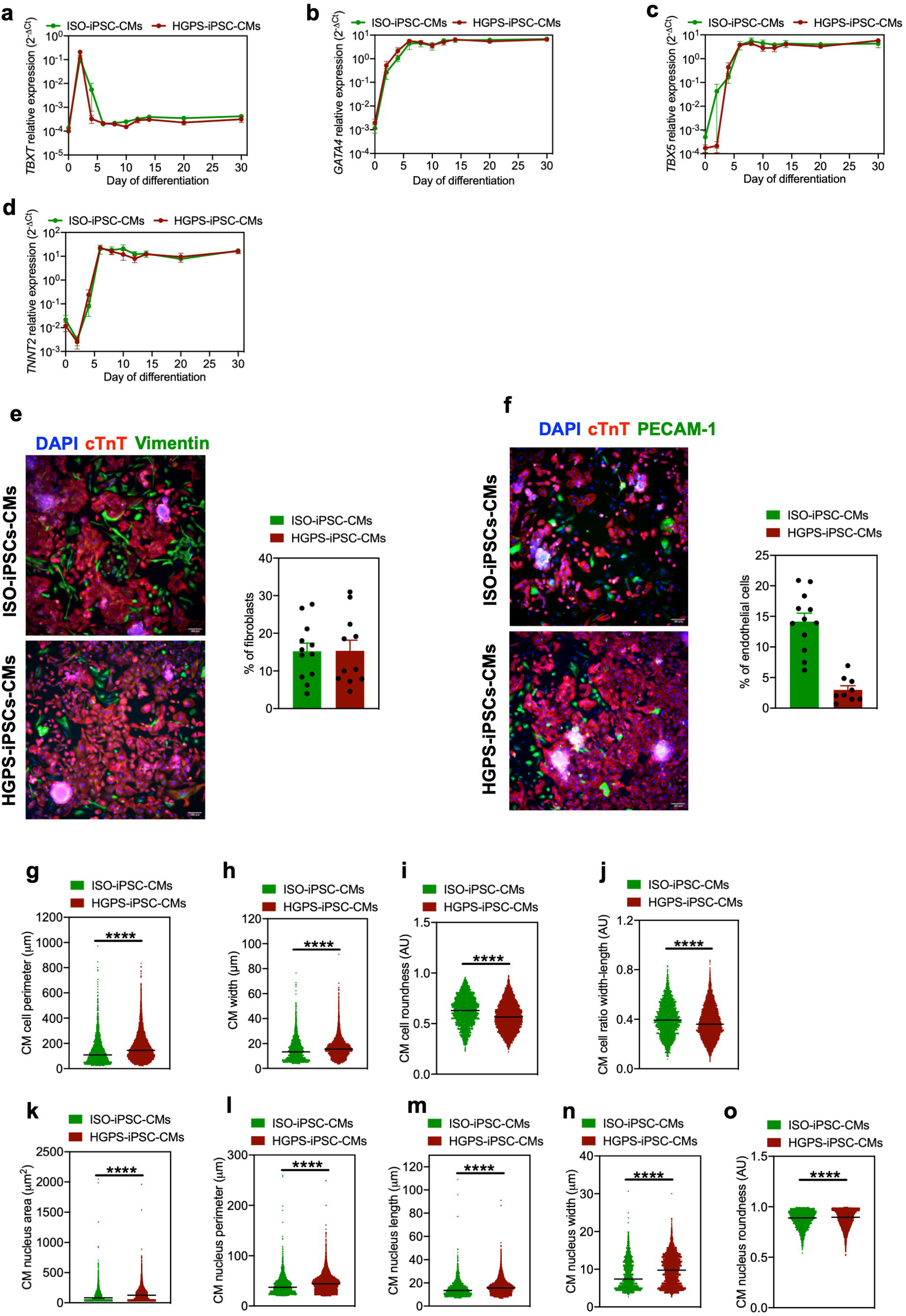

**Extended Data Fig. 3.**
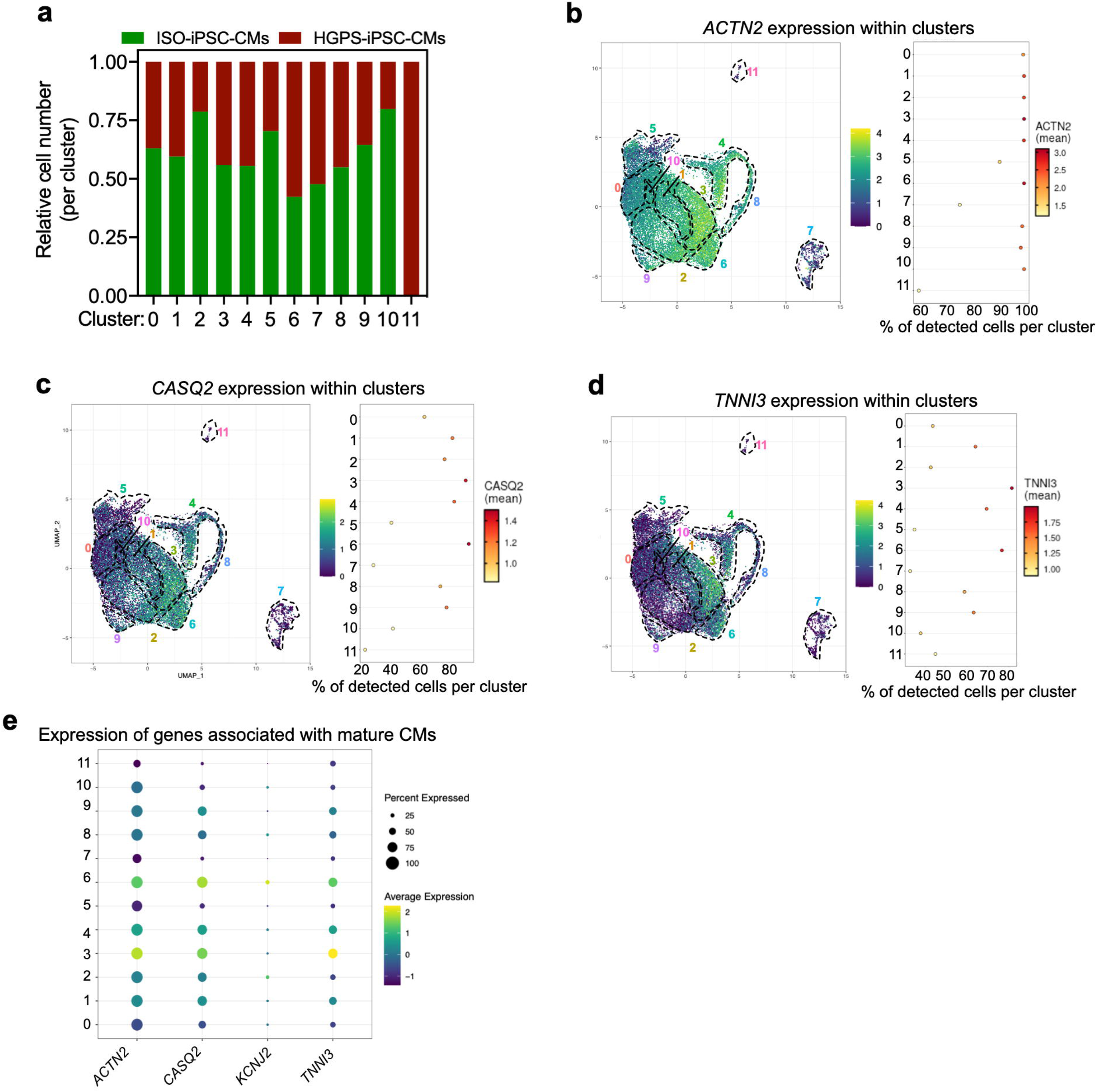

**Extended Data Fig. 4.**
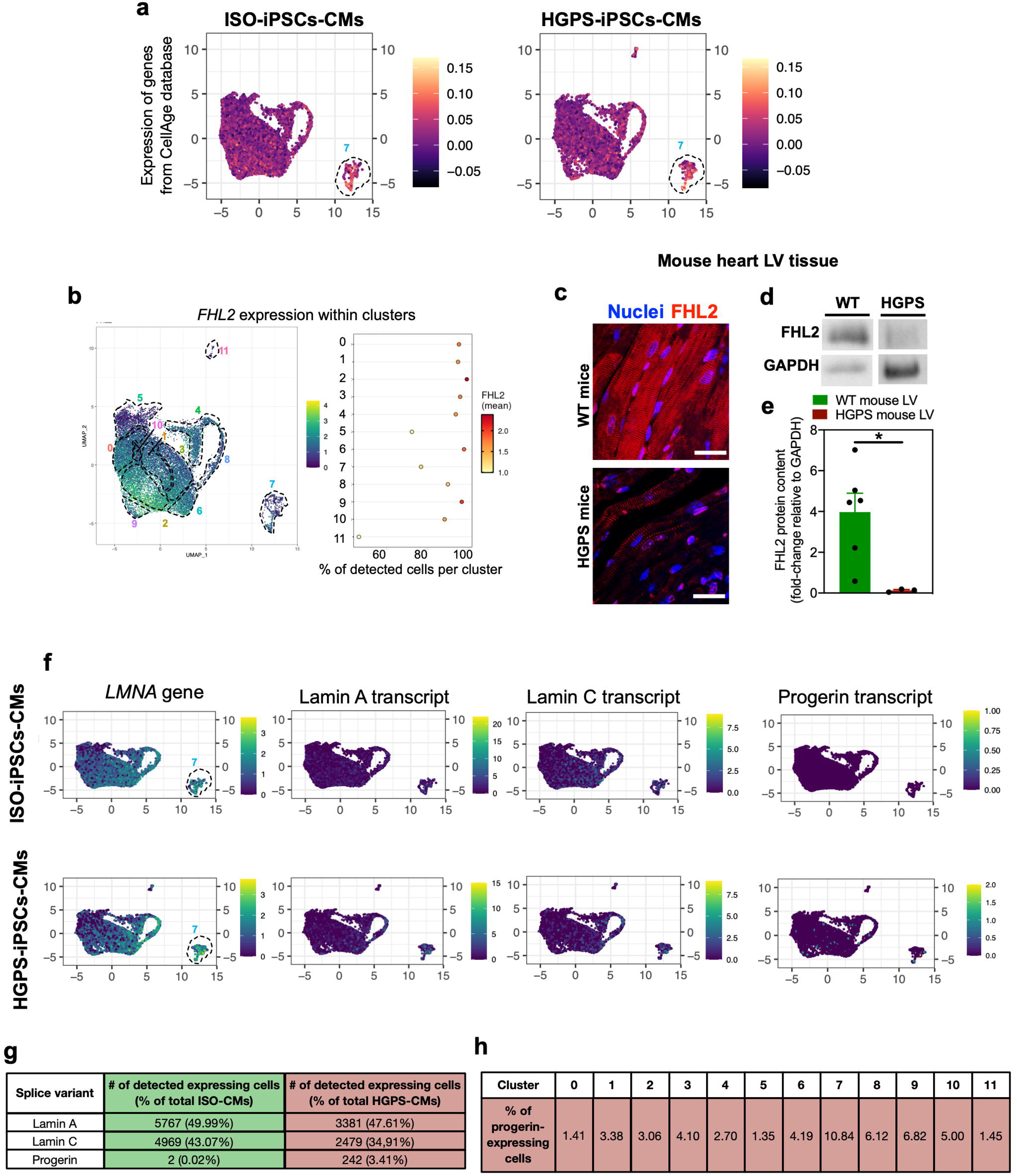

**Extended Data Fig. 5.**
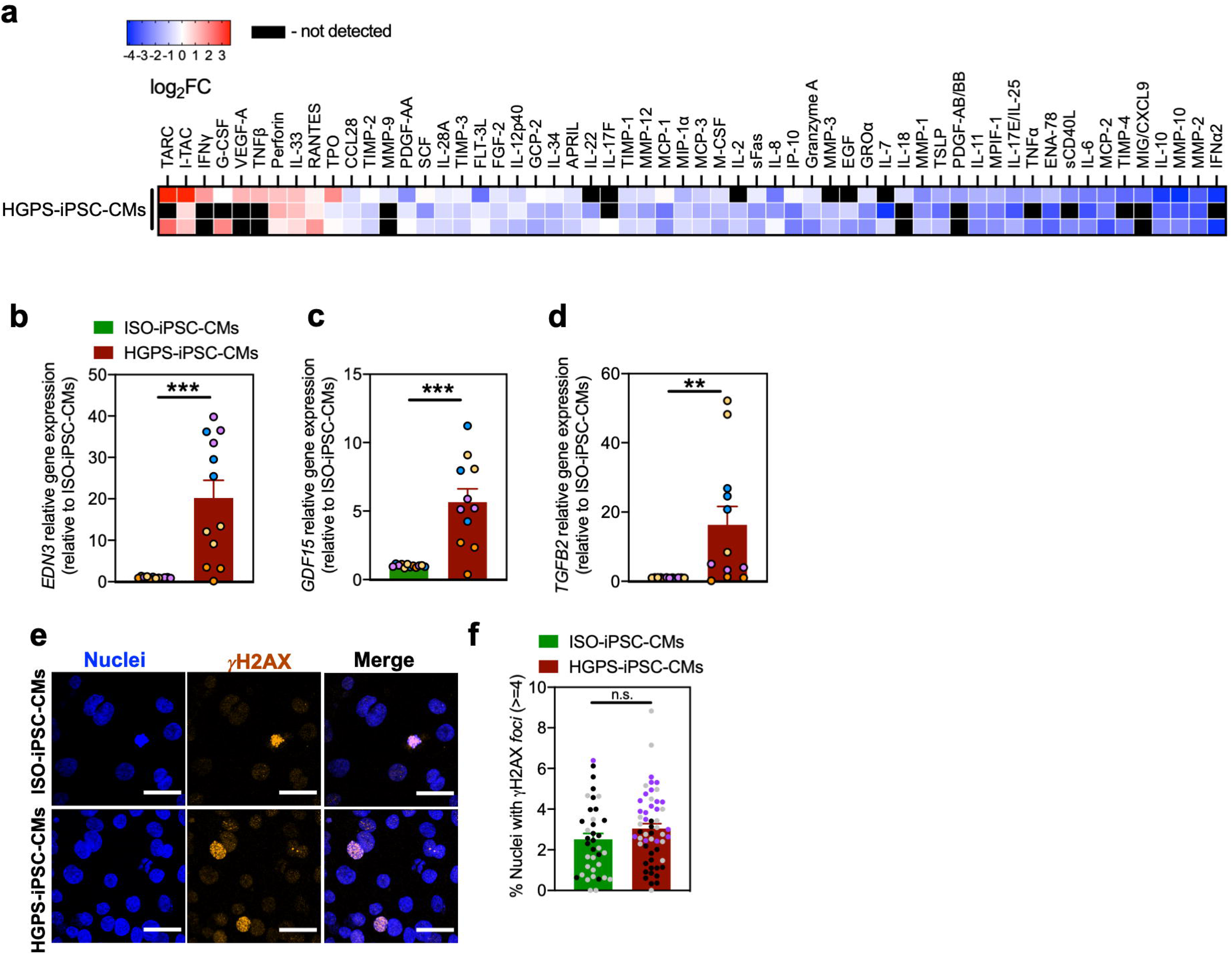

**Extended Data Fig. 6.**
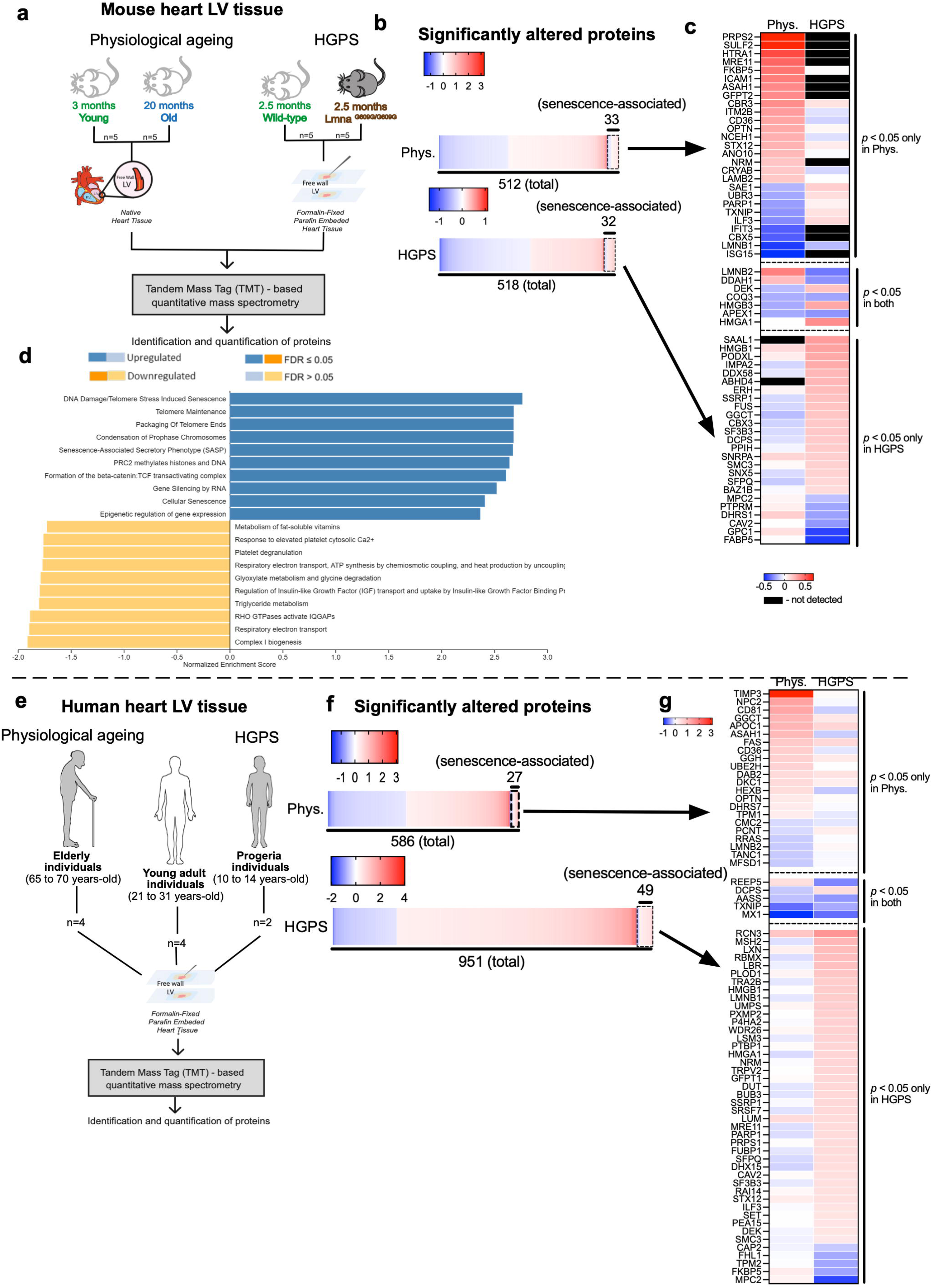

**Extended Data Fig. 7.**
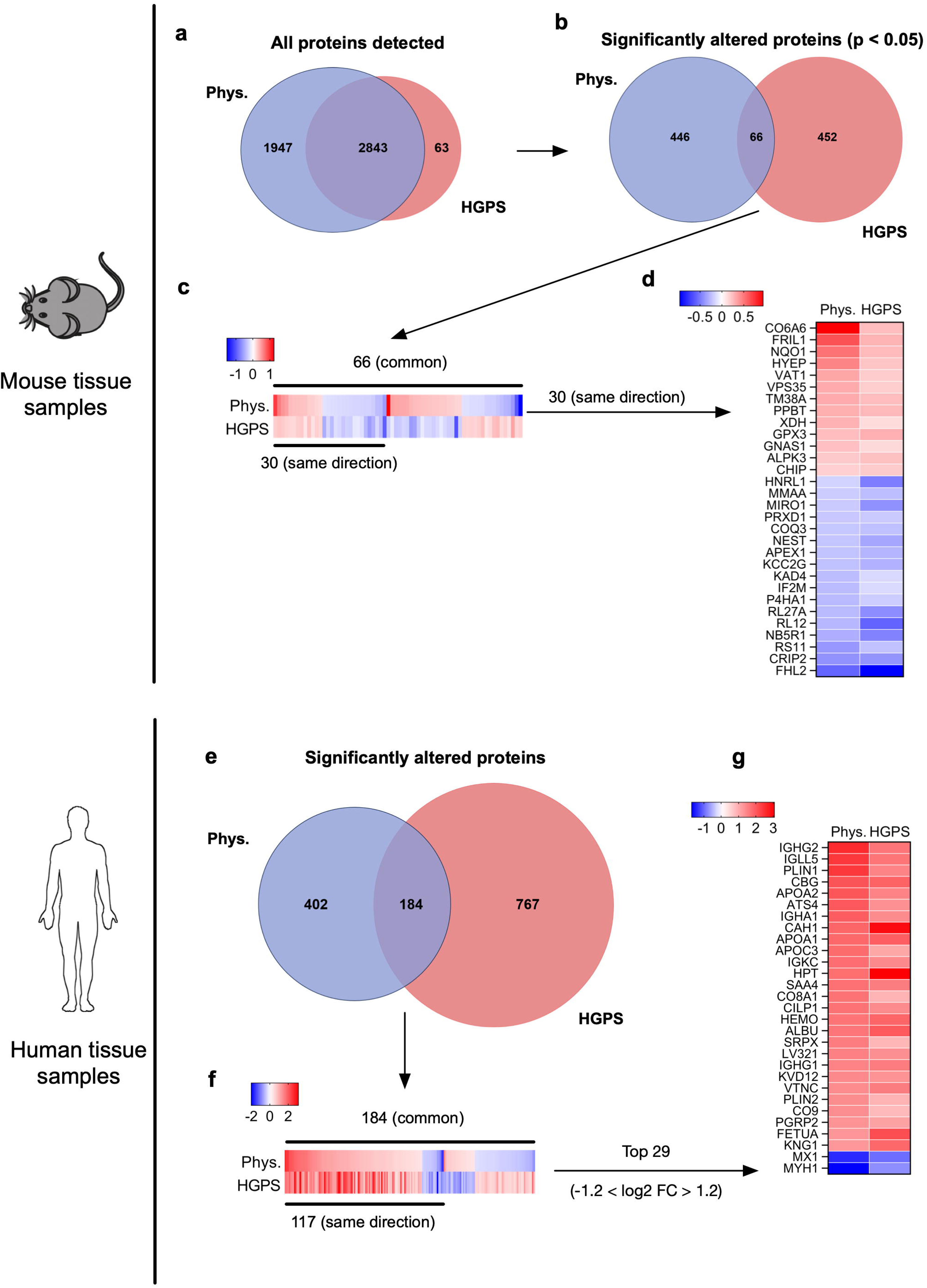

**Extended Data Fig. 8.**
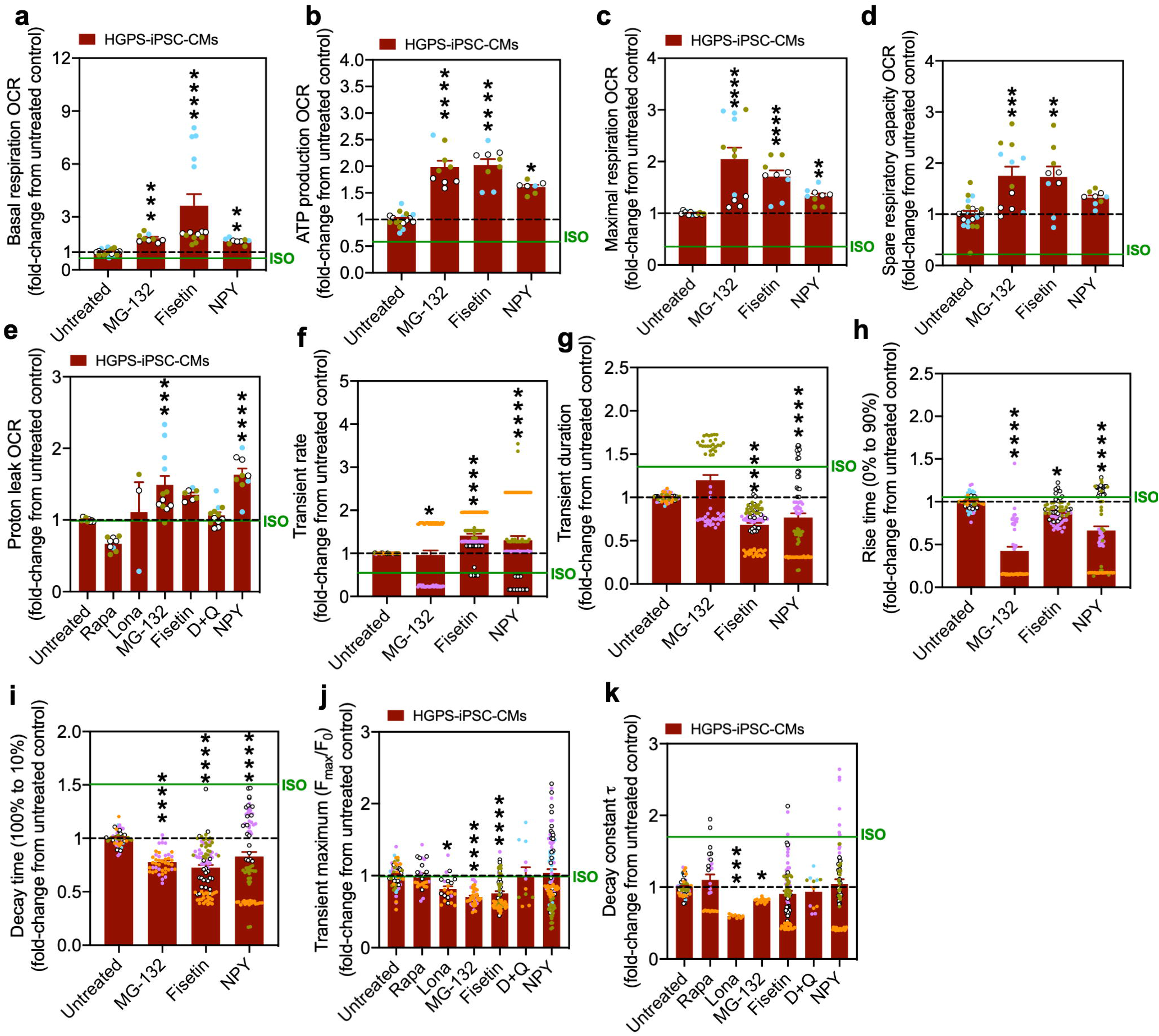

**Extended Data Fig. 9.**
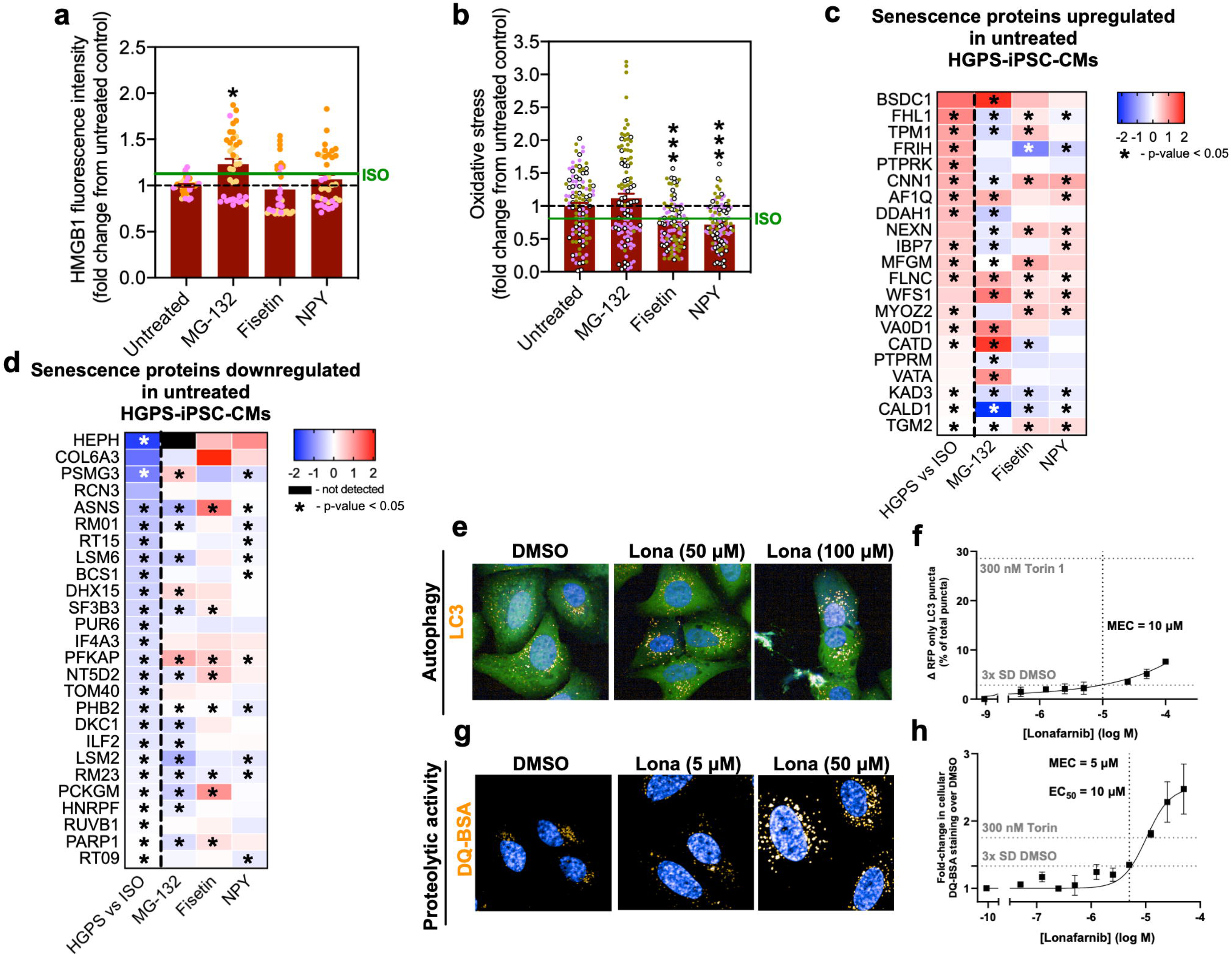

**Extended Data Fig. 10.**
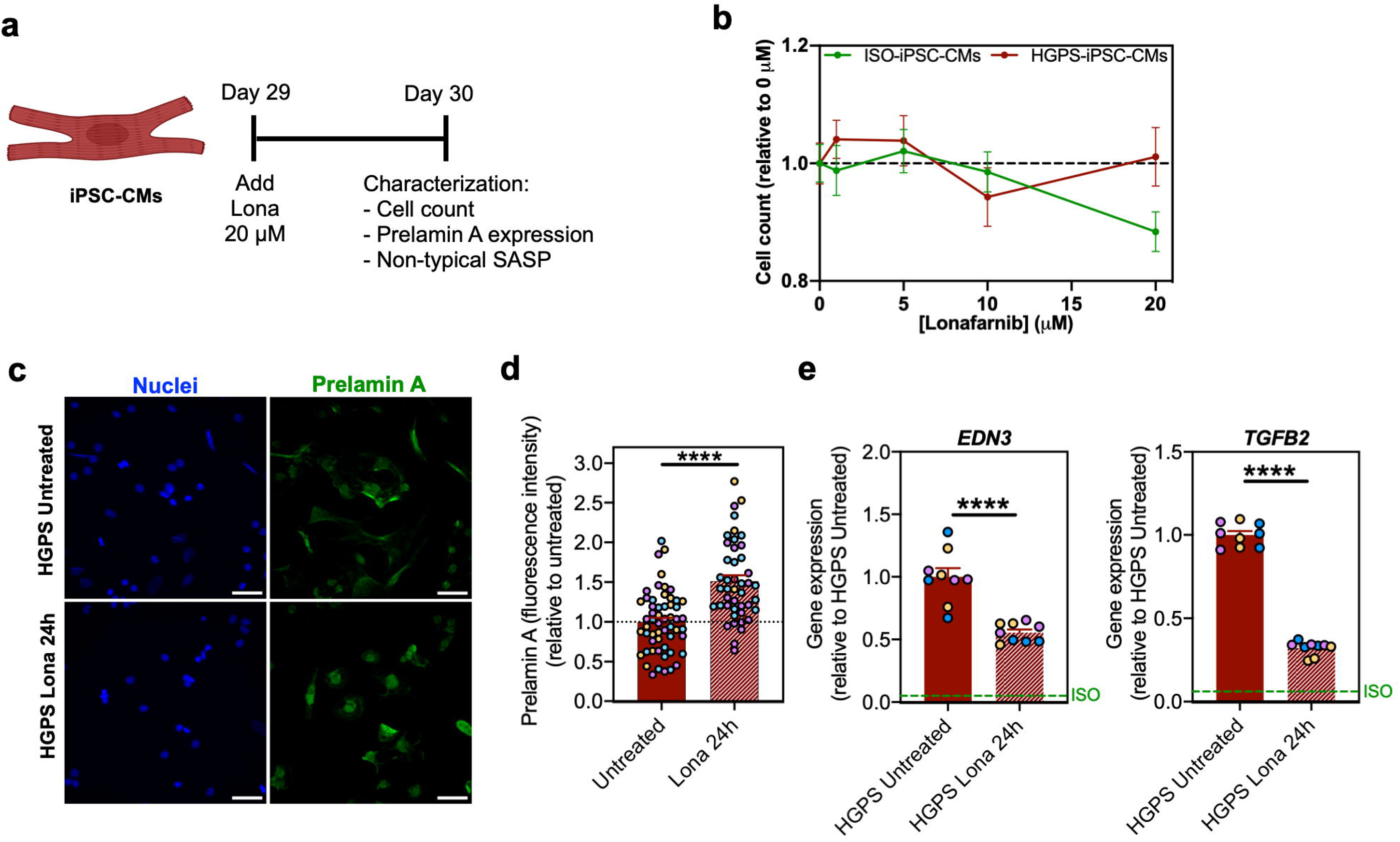

**Supplementary Fig. 1.**
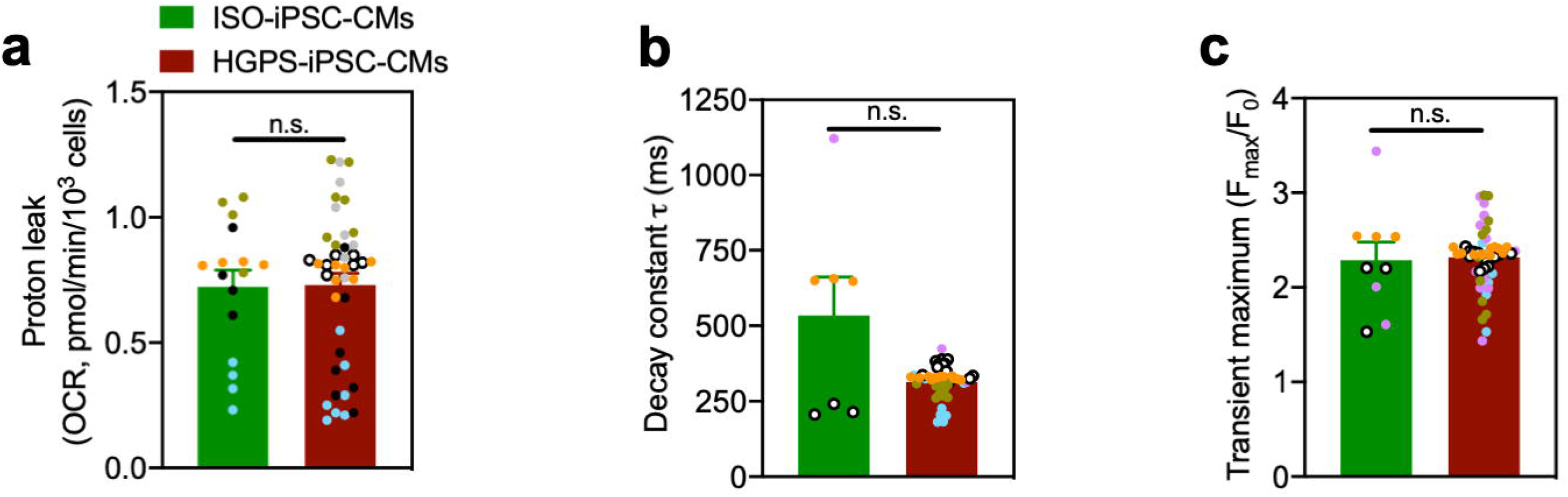

**Supplementary Fig. 2.**
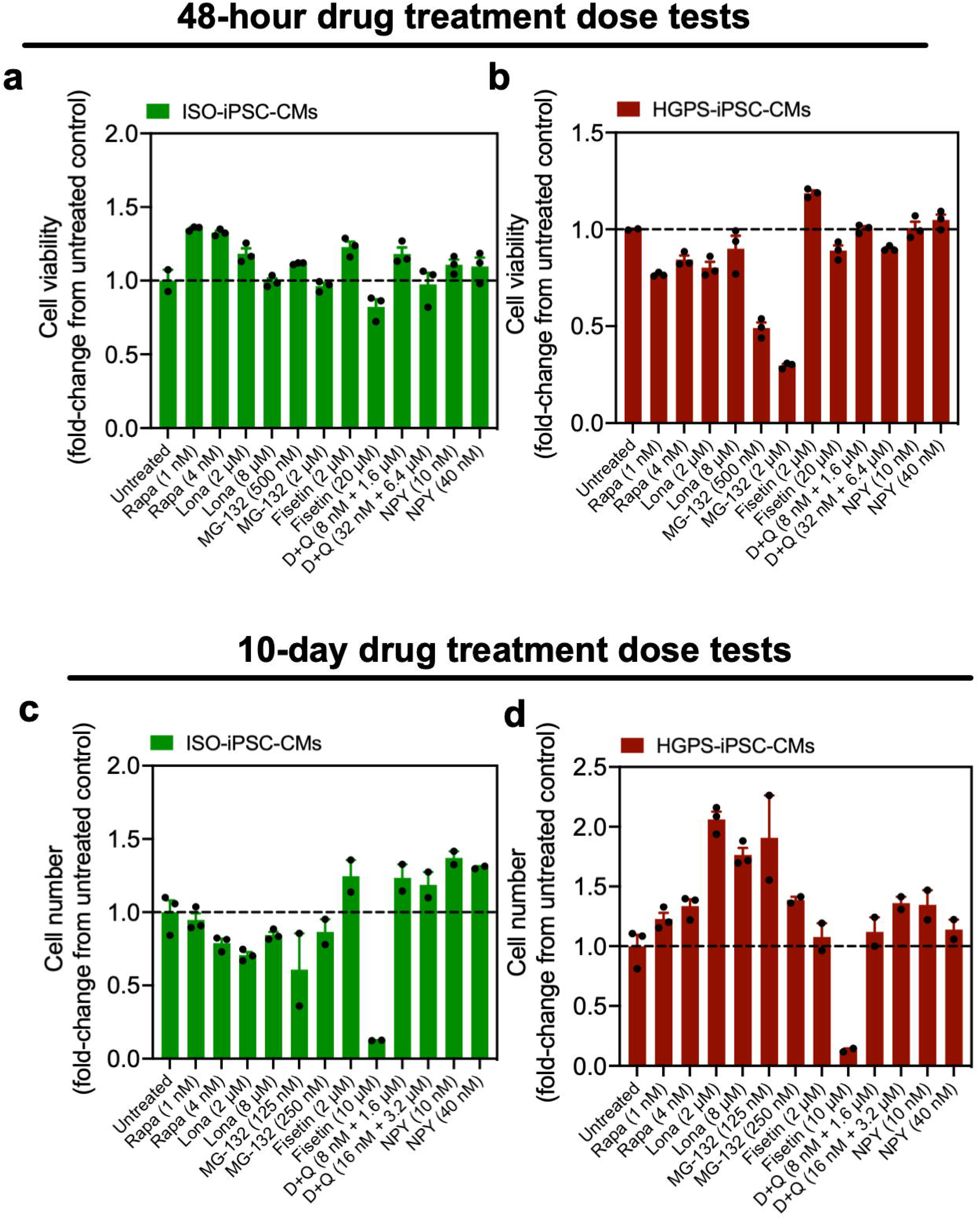

**Supplementary Fig. 3.**
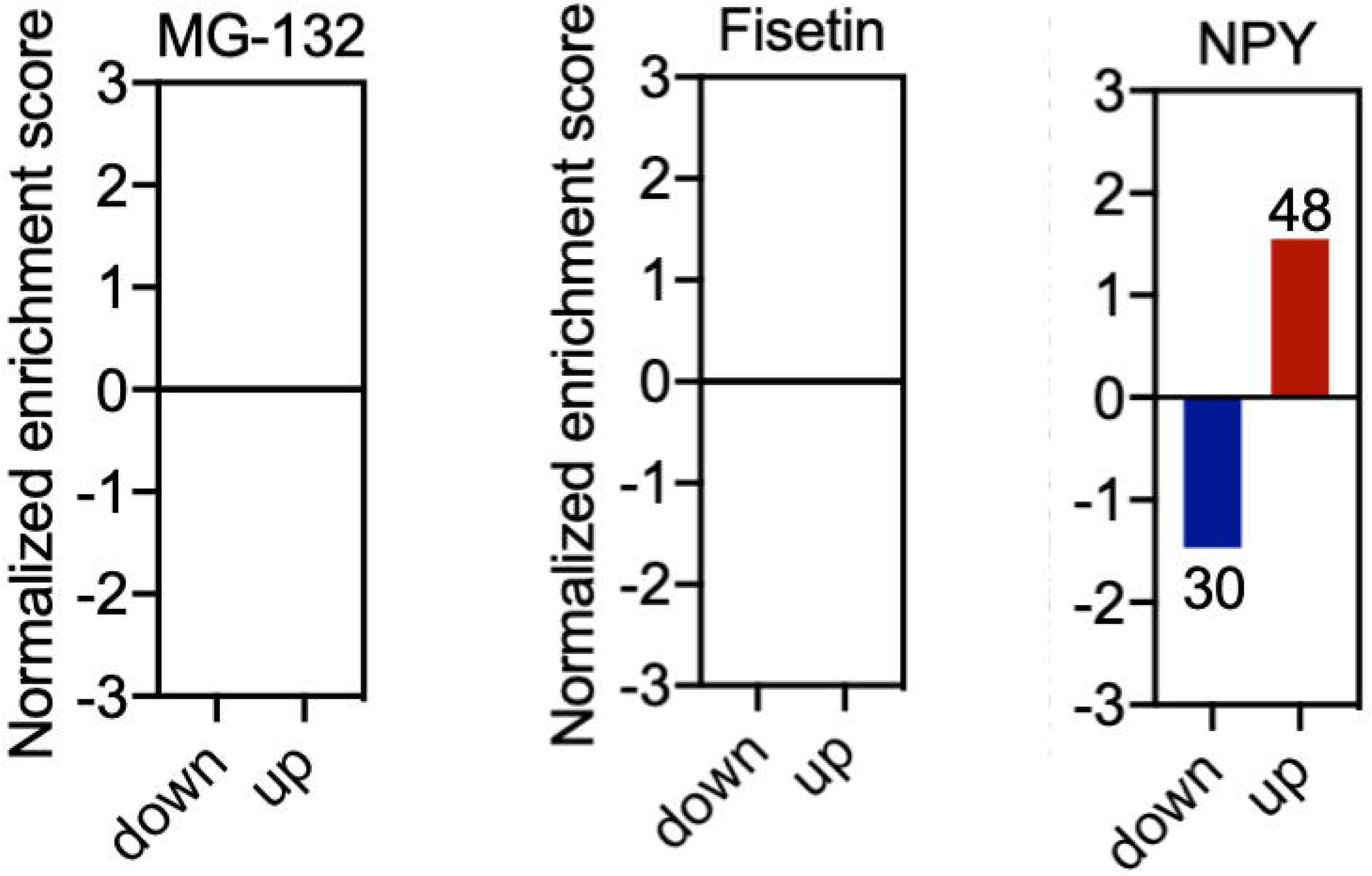

**Supplementary Fig. 4.**
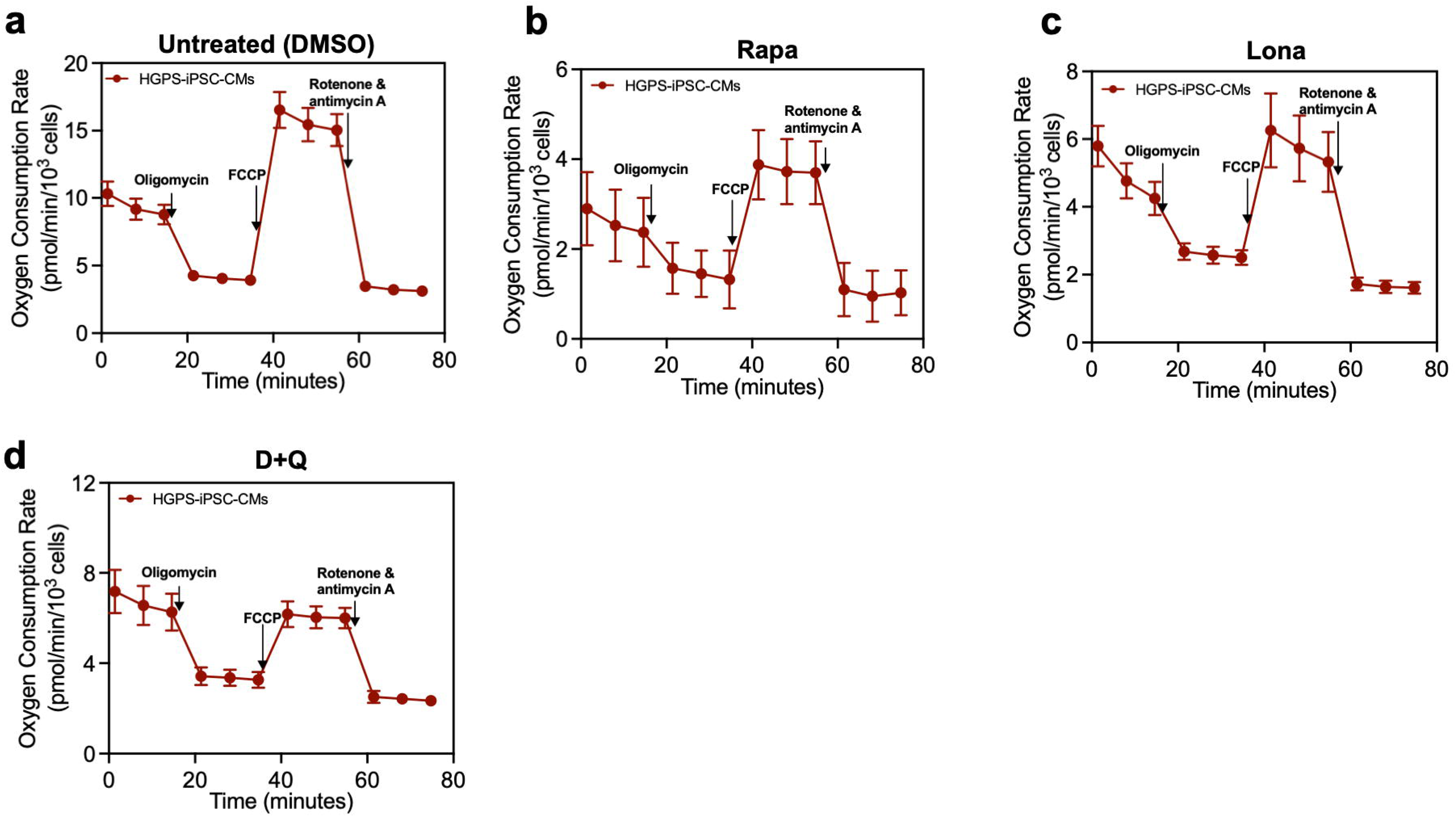

**Supplementary Fig. 5.**
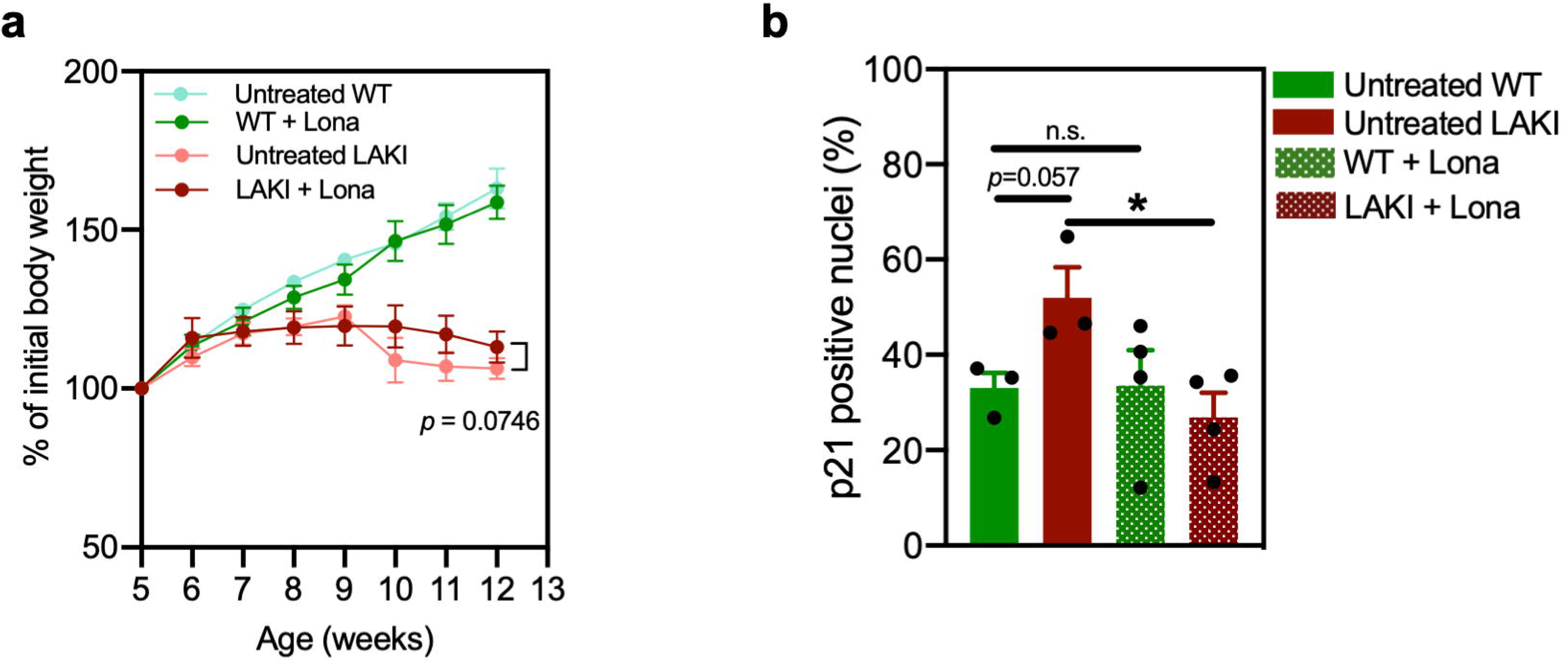

## REFERENCES

1. De Sandre-Giovannoli, A., et al., Lamin a truncation in Hutchinson-Gilford progeria. Science, 2003. 300(5628): p. 2055.

2. Eriksson, M., et al., Recurrent de novo point mutations in lamin A cause Hutchinson-Gilford progeria syndrome. Nature, 2003. 423(6937): p. 293–8.

3. Hamczyk, M.R., L. del Campo, and V. Andres, Aging in the Cardiovascular System: Lessons from Hutchinson-Gilford Progeria Syndrome. Annu Rev Physiol, 2018. 80: p. 27–48.

4. Olive, M., et al., Cardiovascular pathology in Hutchinson-Gilford progeria: correlation with the vascular pathology of aging. Arterioscler Thromb Vasc Biol, 2010. 30(11): p. 2301–9.

5. Rivera-Torres, J., et al., Cardiac electrical defects in progeroid mice and Hutchinson-Gilford progeria syndrome patients with nuclear lamina alterations. Proc Natl Acad Sci U S A, 2016. 113(46): p. E7250–E7259.

6. Prakash, A., Cardiac Abnormalities in Patients With Hutchinson-Gilford Progeria Syndrome. JAMA Cardiology, 2018. 3(4): p. 8.

7. Gordon, L.B., et al., Association of Lonafarnib Treatment vs No Treatment With Mortality Rate in Patients With Hutchinson-Gilford Progeria Syndrome. JAMA, 2018. 319(16): p. 1687–1695.

8. Gordon, L.B., et al., Clinical trial of a farnesyltransferase inhibitor in children with Hutchinson-Gilford progeria syndrome. Proc Natl Acad Sci U S A, 2012. 109(41): p. 16666–71.

9. Cao, K., et al., Rapamycin reverses cellular phenotypes and enhances mutant protein clearance in Hutchinson-Gilford progeria syndrome cells. Sci Transl Med, 2011. 3(89): p. 89ra58.

10. Harhouri, K., et al., MG132-induced progerin clearance is mediated by autophagy activation and splicing regulation. EMBO Mol Med, 2017. 9(9): p. 1294–1313.

11. Aveleira, C.A., et al., Neuropeptide Y Enhances Progerin Clearance and Ameliorates the Senescent Phenotype of Human Hutchinson-Gilford Progeria Syndrome Cells. J Gerontol A Biol Sci Med Sci, 2020. 75(6): p. 1073–1078.

12. Macias, A., et al., Paclitaxel mitigates structural alterations and cardiac conduction system defects in a mouse model of Hutchinson-Gilford progeria syndrome. Cardiovasc Res, 2022. 118(2): p. 503–516.

13. Monnerat, G., et al., Modelling premature cardiac aging with induced pluripotent stem cells from a hutchinson-gilford Progeria Syndrome patient. Front Physiol, 2022. 13: p. 1007418.

14. Pitrez, P.R., et al., Vulnerability of progeroid smooth muscle cells to biomechanical forces is mediated by MMP13. Nat Commun, 2020. 11(1): p. 4110.

15. Xu, Q., et al., Vascular senescence in progeria: role of endothelial dysfunction. Eur Heart J Open, 2022. 2(4): p. oeac047.

16. Pitrez, P.R., et al., Cellular reprogramming as a tool to model human aging in a dish. Nat Commun, 2024. 15(1): p. 1816.

17. Liu, G.H., et al., Recapitulation of premature ageing with iPSCs from Hutchinson-Gilford progeria syndrome. Nature, 2011. 472(7342): p. 221–5.

18. Zhang, J., et al., A human iPSC model of Hutchinson Gilford Progeria reveals vascular smooth muscle and mesenchymal stem cell defects. Cell Stem Cell, 2011. 8(1): p. 31–45.

19. Miller, J.D., et al., Human iPSC-based modeling of late-onset disease via progerin-induced aging. Cell Stem Cell, 2013. 13(6): p. 691–705.

20. Nissan, X., et al., Unique preservation of neural cells in Hutchinson-Gilford progeria syndrome is due to the expression of the neural-specific miR-9 microRNA. Cell Rep, 2012. 2(1): p. 1–9.

21. Gorgoulis, V., et al., Cellular Senescence: Defining a Path Forward. Cell, 2019. 179(4): p. 813–827.

22. Anderson, R., et al., Length-independent telomere damage drives post-mitotic cardiomyocyte senescence. EMBO J, 2019. 38(5).

23. Dark, N., et al., Generation of left ventricle-like cardiomyocytes with improved structural, functional, and metabolic maturity from human pluripotent stem cells. Cell Reports Methods, 2023. 3(4): p. 100456.

24. Churko, J.M., et al., Defining human cardiac transcription factor hierarchies using integrated single-cell heterogeneity analysis. Nat Commun, 2018. 9(1): p. 4906.

25. Cui, Y., et al., Single-Cell Transcriptome Analysis Maps the Developmental Track of the Human Heart. Cell Rep, 2019. 26(7): p. 1934–1950 e5.

26. Zuo, B., et al., Influences of lamin A levels on induction of pluripotent stem cells. Biol Open, 2012. 1(11): p. 1118–27.

27. Tacutu, R., et al., Human Ageing Genomic Resources: integrated databases and tools for the biology and genetics of ageing. Nucleic Acids Res, 2013. 41(Database issue): p. D1027–33.

28. Hojayev, B., et al., FHL2 binds calcineurin and represses pathological cardiac growth. Mol Cell Biol, 2012. 32(19): p. 4025–34.

29. Vigil-Garcia, M., et al., Gene expression profiling of hypertrophic cardiomyocytes identifies new players in pathological remodelling. Cardiovasc Res, 2021. 117(6): p. 1532–1545.

30. Frangogiannis, N.G., The Extracellular Matrix in Ischemic and Nonischemic Heart Failure. Circ Res, 2019. 125(1): p. 117–146.

31. Brown, J.H., D.P. Del Re, and M.A. Sussman, The Rac and Rho hall of fame: a decade of hypertrophic signaling hits. Circ Res, 2006. 98(6): p. 730–42.

32. Divakaruni, A.S., et al., Analysis and interpretation of microplate-based oxygen consumption and pH data. Methods Enzymol, 2014. 547: p. 309–54.

33. Bonkowski, M.S. and D.A. Sinclair, Slowing ageing by design: the rise of NAD(+) and sirtuin-activating compounds. Nat Rev Mol Cell Biol, 2016. 17(11): p. 679–690.

34. Anderson, R., et al., Length-independent telomere damage drives post-mitotic cardiomyocyte senescence. The EMBO Journal, 2019. 38(5): p. e100492.

35. Santinha, D., et al., Remodeling of the Cardiac Extracellular Matrix Proteome During Chronological and Pathological Aging. Mol Cell Proteomics, 2024. 23(1): p. 100706.

36. Hospital, B.C.s., Phase I/II Trial of Everolimus in Combination With Lonafarnib in Progeria. 2015, https://ClinicalTrials.gov/show/NCT02579044.

37. Hickson, L.J., et al., Senolytics decrease senescent cells in humans: Preliminary report from a clinical trial of Dasatinib plus Quercetin in individuals with diabetic kidney disease. EBioMedicine, 2019. 47: p. 446–456.

38. Salerno, N., et al., Pharmacological clearance of senescent cells improves cardiac remodeling and function after myocardial infarction in female aged mice. Mech Ageing Dev, 2022. 208: p. 111740.

39. Clinic, M., Alleviation by Fisetin of Frailty, Inflammation, and Related Measures in Older Adults. 2018, https://ClinicalTrials.gov/show/NCT03675724.

40. Yousefzadeh, M.J., et al., Fisetin is a senotherapeutic that extends health and lifespan. EBioMedicine, 2018. 36: p. 18–28.

41. Zhu, Y., et al., The Achilles’ heel of senescent cells: from transcriptome to senolytic drugs. Aging Cell, 2015. 14(4): p. 644–58.

42. Osorio, F.G., et al., Splicing-directed therapy in a new mouse model of human accelerated aging. Sci Transl Med, 2011. 3(106): p. 106ra107.

43. Petti, E., et al., SFPQ and NONO suppress RNA:DNA-hybrid-related telomere instability. Nature Communications, 2019. 10(1): p. 1001.

44. Lo Cicero, A., et al., Pathological modelling of pigmentation disorders associated with Hutchinson-Gilford Progeria Syndrome (HGPS) revealed an impaired melanogenesis pathway in iPS-derived melanocytes. Sci Rep, 2018. 8(1): p. 9112.

45. Cyganek, L., et al., Deep phenotyping of human induced pluripotent stem cell-derived atrial and ventricular cardiomyocytes. JCI Insight, 2018. 3(12).

46. Lian, X., et al., Directed cardiomyocyte differentiation from human pluripotent stem cells by modulating Wnt/beta-catenin signaling under fully defined conditions. Nat Protoc, 2013. 8(1): p. 162–75.

47. Guo, Y. and W.T. Pu, Cardiomyocyte Maturation: New Phase in Development. Circ Res, 2020. 126(8): p. 1086–1106.

48. Gouveia, P.J., et al., Flexible nanofilms coated with aligned piezoelectric microfibers preserve the contractility of cardiomyocytes. Biomaterials, 2017. 139: p. 213–228.

49. Harper, J.W., et al., The p21 Cdk-interacting protein Cip1 is a potent inhibitor of G1 cyclin-dependent kinases. Cell, 1993. 75(4): p. 805–16.

50. Wagner, K.D. and N. Wagner, The Senescence Markers p16INK4A, p14ARF/p19ARF, and p21 in Organ Development and Homeostasis. Cells, 2022. 11(12).

51. Zhang, H., et al., Reduction of elevated proton leak rejuvenates mitochondria in the aged cardiomyocyte. Elife, 2020. 9.

52. Vehns, E., R. Arnold, and K. Djabali, Impact of MnTBAP and Baricitinib Treatment on Hutchinson–Gilford Progeria Fibroblasts. 2022. 15(8): p. 945.

53. Perez-Riverol, Y., et al., The PRIDE database resources in 2022: a hub for mass spectrometry-based proteomics evidences. Nucleic Acids Res, 2022. 50(D1): p. D543–D552.

54. Hao, Y., et al., Integrated analysis of multimodal single-cell data. Cell, 2021. 184(13): p. 3573–3587.e29.

55. Tirosh, I., et al., Dissecting the multicellular ecosystem of metastatic melanoma by single-cell RNA-seq. Science, 2016. 352(6282): p. 189–96.

56. McGinnis, C.S., L.M. Murrow, and Z.J. Gartner, DoubletFinder: Doublet Detection in Single-Cell RNA Sequencing Data Using Artificial Nearest Neighbors. Cell Syst, 2019. 8(4): p. 329–337.e4.

57. Wu, T., et al., clusterProfiler 4.0: A universal enrichment tool for interpreting omics data. Innovation (Camb), 2021. 2(3): p. 100141.

58. Holden, P. and W.A. Horton, Crude subcellular fractionation of cultured mammalian cell lines. BMC Res Notes, 2009. 2: p. 243.

59. Ferreira, L.L., et al., Doxorubicin persistently rewires cardiac circadian homeostasis in mice. Arch Toxicol, 2020. 94(1): p. 257–271.

60. Psaras, Y., et al., CalTrack: High-Throughput Automated Calcium Transient Analysis in Cardiomyocytes. Circ Res, 2021. 129(2): p. 326–341.

61. Murtada, S.-I., et al., Lonafarnib improves cardiovascular function and survival in a mouse model of Hutchinson-Gilford progeria syndrome. eLife, 2023. 12: p. e82728.

