## Supplementary material for "Lonafarnib Partially Reverses Cardiac Senescence in Human and Mouse Progeria Models via Autophagy Activation": LEGENDS_Extended_Data_and_Suppl_Figs

**Extended Data Figures**

**Extended Data Fig 1 - iPSC genome editing and pluripotency assessment. (a)** CRISPR/Cas9-mediated genome editing methodology to obtain mutation-corrected ISO-iPSCs. **(b)** Sanger sequencing in the *LMNA* locus of the genome, confirming the correction of the HGPS point mutation. **(c)** Representative immunostainings in ISO- and HGPS-iPSCs for the pluripotency markers NANOG, SOX2 and OCT4. Scale bars: 100 𝜇m. **(d)** Karyotype profile of ISO- (top) and HGPS-iPSCs (bottom). No chromosomal aberrations were found. The whole genome view displays all somatic and sex chromosomes in one frame with a high-level copy number. The smooth signal plot (right y-axis) is the smoothing of the log2 ratios which depict the signal intensities of probes on the microarray. A value of 2 represents a normal copy number state (CN=2). A value of 3 represents chromosomal gain (CN=3). A value of 1 represents a chromosomal loss (CN=1). The pink, green and yellow colors indicate the raw signal for each chromosome probe, while the blue signal represents the normalized probe signal which is used to identify copy number and aberrations (if any).

**Extended Data Fig 2 – ISO-iPSC and HGPS-iPSC differentiation into CMs. (a-d)** RT-qPCR quantification of mRNA encoding *TBXT* **(a)**, *GATA4* **(b)**, *TBX5* **(c)**, and *TNNT2* **(d)**. Expression was normalized by *GAPDH* (n=3 independent differentiations). Values are presented as mean± SEM. **(e)** Representative immunostainings for CM and fibroblast markers (cTnT and vimentin, respectively), and quantification of the percentage of fibroblasts for each cell line. Values are presented as mean ± SEM (n=12 images for ISO; n=11 images for HGPS). **(f)** Representative immunostainings for CM and endothelial cell markers (cTnT and PECAM-1, respectively), and quantification of the percentage of endothelial cells for each cell line. Values are presented as mean ± SEM (n=12 images for ISO; n=9 images for HGPS). Scale bars: 100 𝜇m. cTnT - cardiac troponin T. **(g-o)** Measurement of cellular and nuclear morphological parameters of individual CMs (staining for MYL2 and DAPI): **(g)** cell perimeter; **(h)** cell width; **(i)** cell roundness; **(j)** cell width/length ratio; **(k)** nuclear area; **(l)** nuclear perimeter; **(m)** nuclear length; **(n)** nuclear width and **(o)** nuclear roundness. Values are represented as the median (ISO-iPSC-CMs: n=2503 cells analyzed; HGPS-iPSC-CMs: n=7509 individual CMs analyzed). AU: arbitrary units.

**Extended Data Fig 3** **- Single-cell transcriptomics analyses of ISO- or HGPS-iPSC-CMs: expression of CM maturation markers. (a)** Proportion/enrichment of the number of ISO- and HGPS-iPSC-CMs in each cluster, normalized by the total cell number in each cluster. **(b-d)** UMAPs representing the expression, by cluster, of *ACTN2* **(b)**, *CASQ2* **(c)**, and *TNNI3* **(d)**, markers of mature CMs. **(e)** Bubble plot representing, by cluster, the percentage of expressing cells and the level of expression of genes associated with increased CM maturity.

**Extended Data Fig 4 - Single-cell transcriptomics analyses of ISO- or HGPS-iPSC-CMs: expression of ageing and progeria markers. (a)** UMAPs representing the expression of CellAge database genes in ISO- or HGPS-iPSC-CMs. **(b)** Single-cell transcriptome expression of *FHL2* (CM marker) by cluster, represented by UMAP (left) and quantified in terms of the percentage of expressing cells and level of expression (right). **(c)** Representative FHL2 immunostainings in LV cardiac tissue from WT and HGPS mice. Scale bar: 25 μm. **(d)** Representative FHL2 (32 kDa) western blot. Loading control: GAPDH (36 kDa). **(e)** Relative FHL2 western blot quantification (normalized to GAPDH). Values are presented as mean + SEM (n=6 for WT; n=3 for HGPS).  **(f)** UMAP visual representation of the number of expressing cells and levels of expression of the *LMNA* gene and its individual isoform transcripts lamin A, lamin C and progerin. **(g)** Absolute number and relative percentage of *LMNA* isoform detections (splice variants: lamin A, lamin C, or progerin) in ISO-iPSC-CMs and HGPS-iPSC-CMs. **(h)** Estimated percentage of HGPS-iPSC-CM cells that express the progerin transcript per cluster. The RNA-seq assay platform used (3’-centric) was not designed to properly distinguish different isoforms of a gene. The isoforms transcripts are very similar in sequence and sizes (3178 bp, 2461 bp, 2253 bp for lamin A, C and progerin, respectively) and thus greatly limiting the number of usable reads for this particular analysis. For example, it was expected for all the cells to express lamin A and C transcripts, but the percentage of cells that were detected to express them was below 50% in either of the cell lines. Nonetheless, we were able to determine the amount of progeria-expressing cells in relative terms.

**Extended Data Fig 5 – Characterization of the senescence phenotype in HGPS-iPSC-CMs. (a)** Heatmap representing the log_2_(fold-change) of concentration of SASP-related molecules released to the culture medium, relative to ISO-iPSC-CMs. Each row represents an independent differentiation batch (n=3 independent differentiations analyzed per cell line). Only factors where average (|log_2_(fold-change) |) ≥ 0.58 are presented. **(b-d)** *EDN3* **(b)**, *GDF15* **(c)** and *TGFB2* **(d)** gene expression (relative to *GAPDH*) in ISO-iPSC-CMs and HGPS-iPSC-CMs, both normalised to the ISO-iPSC-CMs. All values are presented as mean± SEM (n=3 independent experiments for both ISO- and HGPS-iPSC-CMs, represented by color-coded symbols; n=9 technical replicates both groups). **(e)** Representative immunostaining for 𝛾H2AX in ISO- and HGPS-iPSC-CMs. Scale bar: 50 𝜇m. **(f)** Quantification of the percentage of cells having 4 or more 𝛾H2AX *foci* in the nucleus. Values are presented as mean ± SEM (n=3 independent experiments for both ISO- and HGPS-iPSC-CMs, represented by color-coded symbols; n=18 images for both groups). n.s. - not significant; ** - *p <* 0.01; *** - *p <* 0.001.

**Extended Data Fig 6 – Proteomic analyses on physiologically aged and HGPS heart LV tissue from mouse and human. (a)** Schematic workflow of the proteomics procedure with mouse heart LV tissue samples. **(b)** Heatmaps representing the log_2_(fold-change) of the significantly altered (*p* < 0.05) proteins in the LV heart tissue of physiologically/chronologically aged (Phys.) (top) and HGPS mice (bottom), when compared to their young/healthy counterparts. The number of senescence-related proteins (cross-referenced with CellAge database) is highlighted and their number is displayed. **(c)** Heatmap representing the log_2_(fold-change) of senescence-related proteins (cross-referenced with CellAge database) that are statistically altered (*p* < 0.05), and which direction of change aligns with a senescent phenotype, in physiologically aged mice (top), HGPS mice (bottom), or in both (middle). **(d)** Gene set enrichment analysis display of the twenty most relevant affected pathways in HGPS, in comparison with their WT counterparts. **(e)** Schematic workflow of the proteomics procedure with human heart LV tissue samples. **(f)** Heatmaps representing the log_2_(fold-change) of the significantly altered (*p* < 0.05) proteins in the LV heart tissue of physiologically/chronologically aged (Phys.) (top) and HGPS human individuals (bottom), when compared to their young/healthy counterparts. The number of senescence-related proteins (cross-referenced with CellAge database) is highlighted and their number is displayed. **(g)** Heatmap representing the log_2_(fold-change) of senescence-related proteins (cross-referenced with CellAge database) that are statistically altered (*p* < 0.05), and which direction of change aligns with a senescent phenotype, in physiologically aged human individuals (top), HGPS human individuals (bottom), or in both (middle).

**Extended Data Fig 7 – Proteomic analyses of physiologically aged and HGPS mice’s and human’s heart LV tissue. (a)** Venn diagram depicting the number of all proteins detected in mouse heart LV tissue in aged and HGPS mice. **(b)** Venn diagram depicting the significantly altered proteins (*p* < 0.05) in both aged and HGPS mice heart LV tissue. **(c)** Heatmap representing the log2(fold-change) of common significantly altered (*p* < 0.05) proteins between the LV heart tissue of physiologically aged (Phys.) (top) and HGPS mice (bottom). The number of proteins altered in the same direction is highlighted. **(d)** Heatmap depicting the 30 proteins altered in the same direction. **(e)** Venn diagram depicting the significantly altered proteins in both aged and HGPS human heart LV tissue. **(f)** Heatmap representing the log2(fold-change) of common significantly altered (*p* < 0.05) proteins between the LV heart tissue of physiologically/ aged (Phys.) (top) and HGPS (bottom). The number of proteins altered in the same direction is highlighted. **(g)** Heatmap of the most altered proteins ( |log_2_(fold-change)| ≥ 1.2 ).

**Extended Data Fig 8 - *In vitro* drug effects. (a-e)** Evaluation of the effect of drugs in the mitochondrial respiration of HGPS-iPSC-CMs. Values were normalized to control and number of cells. These parameters were represented as the fold-change of the OCR of each drug. **(a)** Basal respiration. Values presented as mean±SEM (n=3 independent experiments, represented by color-coded symbols; n=7, 4, 5, and 3 technical replicates for Untreated, MG-132, Fisetin, and NPY, respectively). **(b)** ATP production. Values presented as mean±SEM (n=3 independent experiments, represented by color-coded symbols; n=6, 4, 3 and 3 technical replicates for Untreated, MG-132, Fisetin and NPY, respectively). **(c)** Maximal respiration. Values presented as mean±SEM (n=3 independent experiments, represented by color-coded symbols; n=7, 4, 3 and 3 technical replicates for Untreated, MG-132, Fisetin and NPY, respectively). **(d)** Spare respiratory capacity. Values presented as mean±SEM (n=3 independent experiments, represented by color-coded symbols; n=7, 4, 3 and 3 technical replicates for Untreated, MG-132, Fisetin and NPY, respectively). **(e)** Proton leak, mitochondrial respiration parameter. Values presented as mean±SEM (n=3 independent experiments, represented by color-coded symbols; n=5, 4, 1, 4, 3, 5 and 3 technical replicates for Untreated, Rapa, Lona, MG-132, Fisetin, D+Q and NPY, respectively). **(f-k)** Evaluation of the effect of drugs in the calcium handling of HGPS-iPSC-CMs, represented as a fold-change relative to the untreated control. **(f)** Transient rate. Values presented as mean±SEM (n=5, 2, 4 and 4 independent experiments for Untreated, MG-132, Fisetin and NPY, respectively, represented by color-coded symbols; n=13, 26, 22 and 19 technical replicates for Untreated, MG-132, Fisetin and NPY, respectively). **(g)** Transient duration. Values presented as mean±SEM (n=5, 2, 4 and 4 independent experiments for Untreated, MG-132, Fisetin and NPY, respectively, represented by color-coded symbols; n=13, 26, 22 and 19 technical replicates for Untreated, MG-132, Fisetin and NPY, respectively). **(h)** Rise time (0% to 90%). Values presented as mean±SEM (n=5, 2, 4 and 4 independent experiments for Untreated, MG-132, Fisetin and NPY, respectively, represented by color-coded symbols; n=13, 26, 22 and 19 technical replicates for Untreated, MG-132, Fisetin and NPY, respectively). **(i)** Decay time (100% to 10%). Values presented as mean±SEM (n=5, 2, 4 and 4 independent experiments for Untreated, MG-132, Fisetin and NPY, respectively, represented by color-coded symbols; n=13, 26, 22 and 19 technical replicates for Untreated, MG-132, Fisetin and NPY, respectively. **(j)** Transient maximum. Values presented as mean±SEM (n=5, 3, 3, 2, 4, 4 and 4 independent experiments for Untreated, Rapa, Lona, MG-132, Fisetin, D+Q and NPY, respectively, represented by color-coded symbols; n=13, 9, 9, 26, 22, 3 and 19 technical replicates for Untreated, Rapa, Lona, MG-132, Fisetin, D+Q and NPY, respectively. **(k)** Decay constant 𝜏. Values presented as mean±SEM (n=5, 3, 3, 2, 4, 4 and 4 independent experiments for Untreated, Rapa, Lona, MG-132, Fisetin, D+Q and NPY, respectively, represented by color-coded symbols; n=13, 9, 9, 15, 22, 3 and 19 technical replicates for Untreated, Rapa, Lona, MG-132, Fisetin, D+Q and NPY, respectively).The dashed black line corresponds to the Untreated level. The continuous green line corresponds to the levels of ISO-iPSC-CMs on standard culture conditions. Rapa - rapamycin; Lona - lonafarnib; D+Q - dasatinib + quercetin; NPY: - neuropeptide Y. All values presented as mean ± SEM. n.s. - not significant,* - *p <* 0.05, ** - *p <* 0.01, *** - *p <* 0.001, **** - *p <* 0.0001.

**Extended Data Fig 9** **– *In vitro* effect of drugs on the senescent phenotype and autophagy.**

**(a-d)** Effect of tested drugs in the senescence phenotype of HGPS-iPSC-CMs. **(a)** Effect of the tested drugs on the HMGB1 nuclear protein content in HGPS-iPSC-CMs, represented as the fold-change of HMGB1 image mean fluorescence intensity of each drug relative to the untreated control. Values are presented as mean±SEM (n=3 independent experiments, represented by color-coded symbols; n=12 images per independent experiment). **(b)** Effect of the tested drugs on the oxidative stress in HGPS-iPSC-CMs, represented as the fold-change of the CellROX dye image mean fluorescence intensity of each drug relative to the untreated control. Values are presented as mean±SEM (n=3 independent experiments, represented by color-coded symbols; n=36 images per independent experiment). **(c and d)** Heatmaps representing the effect of the drugs on senescence-associated proteins that were originally significantly upregulated **(c)** or downregulated **(d)** in HGPS-iPSC-CMs, relative to ISO-iPSC-CMs. Heatmap colors represent the log_2_(fold-change) of protein expression. The first column of each heatmap represents the change in protein expression of HGPS-iPSC-CMs when compared with ISO-iPSC-CMs (as a reference), while the remaining columns represent the change of protein expression of drug-treated HGPS-iPSC-CMs in comparison with their Untreated counterparts. NPY: - neuropeptide Y. Heatmaps tiles labelled with “*” correspond to comparisons where p-value < 0.05. **(e-h)** Effect of the tested drugs in autophagy-reporter U2OS cell lines. **(e)** Representative images of U2OS LC3-eGFP-RFP cells treated with Lona. Blue: Hoechst 33342. Green: GFP. Orange: RFP. **(f)** Quantification of images. Autophagic flux was defined as the percentage increase in the proportion of autolysosomes (RFP only) out of all LC3 positive vesicles (yellow, GFP+RFP signal), over DMSO. Error bars represent the SEM of two replicates from different cell passages and source. Data were fitted to a variable slope dose-response curve, which was used to determine the minimum effect concentration (MEC) for Lona to be 10 µM by its intersection with a threshold of 3× the standard deviation of the DMSO controls. **(g)** Representative images of U2OS cells treated with Lona for 24 h before staining with DQ-BSA (lysosome tracker, orange fluorescence) for 6 h. Blue: Hoechst 33342. Orange: DQ-BSA. **(h)** Quantification of images. The total DQ-BSA intensity per cell was calculated after background subtraction and normalized to DMSO treatment. Error bars represent the SEM of two replicates from different cell passages and source. Data were fitted to a variable slope dose-response curve, which was used to determine the minimum effect concentration (MEC) for Lonafarnib to be 5 µM by its intersection with a threshold of 3× the standard deviation of the DMSO controls.

**Extended Data Fig 10** **– Effect of Lona on HGPS-iPSC-CMs. (a)** Experimental design of the acute *in vitro* treatment with Lona.  **(b)** Effect of the acute Lona treatment in the nuclei counts (n=3 independent experiments). **(c and d)** Validation of the Lona effect by assessment of the prelamin A expression changes. **(c)** Representative images of the prelamin A staining in HGPS-iPSC-CMs with or without the Lona 24 h treatment. Scale bar: 50 μm. **(d)** Quantification of the prelamin A fluorescence intensity levels in the nucleus in HGPS-iPSC-CMs with or without the Lona 24 h treatment. Values are presented as mean±SEM (n=3 independent experiments, represented by color-coded symbols; n=20 images per independent experiment). **(e)** Gene expression of the non-canonical SASP genes *EDN3* (left) and *TGFB2* (right) in HGPS-iPSC-CMs with or without the Lona 24 h treatment. Values are presented as mean±SEM (n=3 independent experiments, represented by color-coded symbols; n=3 technical replicates per experiment).

**Supplementary Figures**

**Supplementary Figure 1 - Mitochondrial respiration and calcium handling. (a)** Proton leak, mitochondrial respiration parameter represented as OCR and normalized by cell number for both ISO- and HGPS-iPSCs. Values are presented as mean±SEM (n=4 and 6 independent experiments for ISO- and HGPS-iPSC-CMs, respectively, represented by color-coded symbols; ISO-iPSC-CMs: n= 4 technical replicates per independent experiment; HGPS-iPSC-CMs: n=7 technical replicates per independent experiment). **(b and c)** Calcium handling parameters: decay constant 𝜏 (time required for the calcium transient to decrease from its maximum/peak to 37% of its maximum) and transient maximum. Values are presented as mean±SEM (n=3 and 5 independent experiments for ISO- and HGPS-iPSC-CMs, respectively, represented by color-coded symbols; ISO-iPSC-CMs: n=3 technical replicates per independent experiment for all parameters; HGPS-iPSC-CMs: n=12 technical replicates per independent experiment).

**Supplementary Figure 2** **- Short- and long-term *in vitro* drug dose tests in ISO- and HGPS-iPSC-CMs. (a and b)** Viability (normalized for the control) of ISO-iPSC-CMs **(a)** and HGPS-iPSC-CMs **(b)** after a 48 h (short-term) treatment with two test concentrations per compound. Values presented as mean±SEM (n=1 independent experiment for each cell line; n=2-3 technical replicates per independent experiment). **(c and d)** Cell number normalized for the Untreated control of ISO-iPSC-CMs **(c)** and HGPS-iPSC-CMs **(d)** after a 10-day (long-term) treatment with two test concentrations per compound. Values presented as mean±SEM (n=1 independent experiment for each cell line; n=3 technical replicates per independent experiment). Rapa - rapamycin; Lona - lonafarnib; D+Q - dasatinib + quercetin; NPY: - neuropeptide Y. Although Rapa (1 and 4 nM) caused a tendency to decrease cell viability in HGPS-iPSC-CMs, it increased it in ISO-iPSC-CMs, therefore we maintained these concentrations to be used in the long-term dose test. Considering that both concentrations of MG-132 (500 nM and 2 𝜇M) caused a great decrease in viability in HGPS-iPSC-CMs, we reduced them to 125 and 250 nM. As fisetin (20 𝜇M) decreased viability in both cell lines, we reduced it by half (10 𝜇M) for the long-term test, along with the low dose (2 𝜇M). The highest dose of D+Q (*i.e.* 32 nM + 6.4 𝜇M) reduced viability in both cell lines, therefore we halved this higher concentration (*i.e.* 16 nM + 3.2 𝜇M) and kept the lower one (*i.e.* 8 nM + 1.6 𝜇M). Finally, the remaining drug concentrations were maintained as they did not cause relevant alterations in viability **(a** **and** **b)**. For the long-term drug concentration test, we used the final cell number as a measurement of cell viability, after chronically treating both cell lines with two selected drug concentrations per compound **(c and d)**. As a compromise between cell viability and bioactivity, we selected the highest tested concentrations for most compounds, except for fisetin and NPY, as they only cause a modest decrease in cell number in both cell lines **(c and d)**. As the lowest fisetin concentration caused a cell number increase in both cell lines while the highest caused a major decrease **(c and d)**, we reduced the latter concentration by 25% (i.e. to 7.5 𝜇M) and considered it final. Finally, as the highest tested NPY concentration (40 nM) caused an increase in cell viability, we increased it by 50% (i.e. to 60 nM) for more bioactivity. Of note, the concentration selected for Lona (8 𝜇M), although higher, is within the same order of magnitude of the range of maximal concentrations of Lona detected in the plasma of HGPS patients in a clinical trial (from 2.8 to 4.1 𝜇M, depending on the given dose). Regarding the selected concentration of Rapa (4 nM) is similar, although lower, than the recommended maximal plasma concentration in patients when this compound is used in the context of organ transplantations, as an immunosuppressive drug (from 6.3 to 10.3 nM).

**Supplementary Figure 3** **- Corrective effects of MG132, Fisetin and NPY.** These drugs did not have a corrective effect on the proteins that were originally downregulated or upregulated in HGPS-iPSC-CMs relative to ISO-iPSC-CMs. Plot with the normalized enrichment score.

**Supplementary Figure 4** **- Average mitochondrial respiration profile curves of drug-treated HGPS-iPSC-CMs.** Oxygen consumption rate (OCR) curves normalized by cell number, over the assay time, and upon the sequential addition of the compounds oligomycin, FCCP, rotenone and antimycin A (n= 4-6 independent experiments; n= 2-7 technical replicates per independent experiment) of HGPS-iPSC-CMs subjected to a prolonged exposure to DMSO (Untreated control) **(a)**, Rapa **(b)**, Lona **(c)** and D+Q **(d)**.

**Supplementary Figure 5** **- *In vivo* drug treatment. (a)** Body weight alterations were monitored weekly during the treatment period in all animal groups: Untreated WT, Untreated LAKI, WT + Lona, LAKI + Lona. Values within each animal group were normalised by the initial body weight (at week 5). Values are presented as mean±SEM (Untreated WT: n=6 animals for week 13 and n=7 for the rest of timepoints; Untreated LAKI: n=9 animals for weeks 5 and 6, n=6 for weeks 7 and 11, n = 8 for weeks 8 and 12, n=4 for week 9, n=5 for week 10; WT + Lona: n=5 animals for all timepoints; LAKI + Lona: n=4 animals for all timepoints). **(b)** Quantification of p21-positive nuclei in the LV tissue for all animal groups. Values are presented as mean±SEM (n=3 animals for Untreated WT and Untreated LAKI; n=4 animals for WT+Lona and LAKI + Lona).
