## Supplementary material for "Lonafarnib Partially Reverses Cardiac Senescence in Human and Mouse Progeria Models via Autophagy Activation": Supp_Information

Supplementary Methods

**iPSC culture: origin and culture**

HGPS-iPSCs were generated from HGPS skin fibroblasts (AG01972 from Coriell Institute for Medical Research; 14 year-old female HGPS patient) provided by Coriell Institute and characterized according to Nissan et al. [1], resorting to the Yamanaka’s original method with OCT4, KLF4, SOX2, c-MYC, transferred by retroviral vectors. HGPS fibroblasts were obtained from Coriell Institute for Medical Research, which in turn were collected under Institutional Review Board approval and individual informed consent (https://[www.coriell.org/0/Sections/Support/NIA/Model.aspx?PgId=351](http://www.coriell.org/0/Sections/Support/NIA/Model.aspx?PgId=351)). Isogenic iPSCs (ISO-iPSCs) were obtained by CRISPR/Cas9-mediated correction of the HGPS point mutation (see *Genome editing*).

iPSC lines were splitted, as colonies, every 3-4 days (when confluency was 70-80%) using Versene Solution (ThermoFisher Scientific) (3-4 min at RT) with a 1:6 – 1:12 ratio. iPSC lines were maintained using mTeSR™ Plus medium (STEMCELL Technologies) and 6-well culture plates (Corning® Costar®) coated with hESC-qualified Matrigel® matrix (Corning). iPSCs were kept at 37°C, 5% (v/v) CO_2_ in a humidified atmosphere, and the medium was replaced every 2 days. For cryopreservation, iPSCs were dissociated as colonies (1:2 ratio) with Versene Solution (ThermoFisher Scientific) (3-4 min at RT), centrifuged (157*g*, 5 min, RT), and then resuspended and cryopreserved as clumps with CryoStor® CS10 (STEMCELL Technologies) supplemented with Y-27632 dihydrochloride (5 𝜇M; 1254; Tocris). Cryovials were stored in liquid nitrogen.

**Cardiomyocyte cryopreservation**

For cryopreservation, day 20 cardiomyocytes were dissociated as previously described (see *Cardiomyocyte dissociation and replating*) except that the mechanical dissociation was reduced so that the cardiomyocytes were kept as aggregates. From this point on, the process of cardiomyocytes cryopreservation was the same as the cryopreservation of iPSC colonies (see *Human iPSC lines origin and culture*).

**Multiplex analysis of cytokines for SASP evaluation**

Prior to running the multiplex analysis assay, conditioned medium from independent batches of ISO- and HGPS-iPSC-CMs was collected at day 30 of differentiation, after being in contact with the cells for 3 days. The conditioned medium was stored at -80°C until being processed.

For the multiplex analysis, it was used the Luminex xMAP technology for multiplexed quantification of 96 Human cytokines, chemokines, and growth factors; or 9 human matrix metalloproteinases and 4 tissue inhibitors of metalloproteinases. The multiplexing analysis was performed using the Luminex™ 200 system (Luminex, Austin, TX, USA) by Eve Technologies Corp. (Calgary, Alberta). Ninety-six markers were simultaneously measured in the samples using Eve Technologies' Human Cytokine 96-Plex Discovery Assay® which consists of two separate kits; the Panel A 48-plex and the Panel B 48-plex (MilliporeSigma, Burlington, Massachusetts, USA). Thirteen markers were simultaneously measured in the samples using Eve Technologies' Human MMP/TIMP 13-Plex Discovery Assay® which consists of two separate kits; one 9 plex and one 4 plex (R&D Systems, Inc., Minneapolis, MN, USA). The assays were run according to the manufacturer's protocol. The Panel A 48-plex consisted of sCD40L, EGF, Eotaxin, FGF-2, FLT-3 Ligand, Fractalkine, G-CSF, GM-CSF, GROα, IFN-α2, IFN-γ, IL-1α, IL-1β, IL-1RA, IL-2, IL-3, IL-4, IL-5, IL-6, IL-7, IL-8, IL-9, IL-10, IL-12(p40), IL-12(p70), IL-13, IL-15, IL-17A, IL-17E/IL-25, IL-17F, IL-18, IL-22, IL-27, IP-10, MCP-1, MCP-3, M-CSF, MDC, MIG/CXCL9, MIP-1α, MIP-1β, PDGF-AA, PDGF-AB/BB, RANTES, TGFα, TNF-α, TNF-β, and VEGF-A. The Panel B 48-plex consisted of 6CKine, APRIL, BAFF, BCA-1, CCL28, CTACK, CXCL16, ENA-78, Eotaxin-2, Eotaxin-3, GCP-2, Granzyme A, Granzyme B, HMGB1, I-309, I-TAC, IFNβ, IFNω, IL-11, IL-16, IL-20, IL-21, IL-23, IL-24, IL-28A, IL-29, IL-31, IL-33, IL-34, IL-35, LIF, Lymphotactin, MCP-2, MCP-4, MIP-1δ, MIP-3α, MIP-3β, MPIF-1, Perforin, sCD137, SCF, SDF-1, sFAS, sFASL, TARC, TPO, TRAIL, and TSLP. The 9-plex consisted of MMP-1, MMP-2, MMP-3, MMP-7, MMP-8, MMP-9, MMP-10, MMP-12 and MMP-13. The 4-Plex consisted of TIMP-1, TIMP-2, TIMP-3 and TIMP-4. Assay sensitivities of the markers range from 0.05 – 100 pg/mL for the 96-plex, and 0.28 – 253 pg/mL for the 13-Plex. For the 96-Plex, individual analyte sensitivity values are available in the MILLIPLEX® MAP protocol. Individual analyte sensitivity values are available in the product datasheet for the 4-Plex and by building the panel on the R&D Systems Magnetic Luminex Performance product page for the 9-Plex.

**Tissue RNA and protein extraction**

For protein extraction, 50 - 100 g of mice left ventricle tissue were homogenized in 1 mL of TRIzol™ (ThermoFisher Scientific) resorting to a Tissuelyser II (Qiagen) through 4 cycles of 30 Hz for 15 s followed by 30 s intervals on ice. Downstream processing for TRIzol™ was carried out according to the manufacturer protocol for RNA extraction while organic (protein) phase processing was adapted from [2]. Briefly, 200 µL phenol-ethanol phase were loaded into Spectra/Por® 6 MWCO 2000 regenerated cellulose dialysis membrane (REPLIGEN) and dialyzed against 3 changes of 50 mL 0.1 % SDS (VWR) for 22 h. The eluent was renewed at 16 h and 20 h of dialysis. During the dialysis, a globular mass and a supernatant formed, which was discarded. The globular mass was resuspended by 5 cycles of 15 s sonication in a Branson 2510MT (Emerson) with 30 s pause on ice in 100-200 µL of protein solubilizing solution (1:1 of 1 % SDS and 8 M Urea in 1 M Tris, pH 8). Finally, the protein solution was centrifuged at 3400*g*, at 4 °C for 10 min, the supernatant was collected and kept at -20 °C, and the pellet was discarded.

**Genome editing: procedure**

Prior to transfection, HGPS-iPSCs were incubated for 2 h with Y-27632 dihydrochloride (5 𝜇M; 1254; Tocris), then detached using TrypLE Express® (ThermoFischer) (5 min, 37°C) and then centrifuged for (300*g*, 5 min, RT). Thereafter, a total of 2×10^6^ live HGPS-iPSCs were distributed into two nucleofection reactions, where 1×10^6^ cells were resuspended in a 100 μL-reaction volume composed of: 82 μL of Nucleofector^TM^ solution and 18 μL of Supplement from the Amaxa^TM^ P3 Primary Cell 4D-Nucleofector^TM^ X Kit (Lonza), with 1 μg of sgRNA plasmid and 3.8 μg of repair template plasmid. The mixture was added into a Nucleocuvette^TM^ (Lonza) and nucleofected using a Nucleofector^TM^ system (Lonza). Nucleofected cells were plated on feeders (CF1 MEFs, generated in-house) in KSR medium supplemented with Y-27632 dihydrochloride (5 𝜇M; 1254; Tocris). Nucleofected cells were plated into two full 6-well plates to allow for single-cell colonies to emerge. After two weeks, 480 single colonies were manually picked and plated in individual 96-well plate wells pre-seeded with CF1 MEFs and in KSR Medium. Individual clones were maintained in culture until genotyping was completed.

**Genome editing: clone genotyping**

To isolate genomic DNA from the individual clones, cells from confluent 96-well plate wells were detached using TrypLE Express® (ThermoFisher), centrifuged (300*g*, 3 min, RT), and then resuspended and incubated (55 min, 37°C) in 20 μL of Lysis Buffer (lysis buffer composition: 670 mM Tris HCl at pH 9.1, 160 mM (NH_4_)_2_SO_4_, 35 mM MgCl_2_, 1.5 mg/mL BSA; 1 μL of 10% Igepal (Sigma); 0.8 μL of 20 mg/mL Proteinase K (New England Biolabs) and 16.2 μL of dH_2_O). The lysis reaction was stopped by incubating the samples for 15 min at 95°C. The resulting lysate was used directly in the PCR reaction. A 1098 bp region containing the HGPS mutation was amplified by PCR using a Q5® High-Fidelity DNA Polymerase (New England Biolabs) and pre-tested primers (Amplification Forward primer: 5 ́-TCTCTCCCCCATTCTTGTTGCA-3’; Amplification Reverse primer: 5 ́-CATGAGGTGAGGAGGACGCAG-3’). Sanger sequencing was then used to determine if the targeted site had been repaired. To this end, we designed primers for a 500 bp region contained within the 1098 bp amplicons; both forward and reverse primers (Sequencing Forward primer: 5 ́- CCTTGGGCACAGAACCACAC -3’; Sequencing Reverse primer: 5 ́- GCGGAAGAGAAGGCAGGCTC-3’) were used to confirm if the region had been repaired, which was that the clone had a “C” on both alleles of the HGPS mutation site and that one allele assessed by confirming had the altered PAM site.

**Karyotype analysis**

The karyotype of HGPS-iPSCs and HGPSΔ2-iPSC samples were analyzed using KaryoStatTM assay (ThermoFisher Scientific). Briefly, genomic DNA (gDNA) was extracted using PureLink Genomic DNA Mini Kit (catalog # K182000) and estimated by Qubit dsDNA BR Assay Kits. gDNA was processed according to the manufacturer’s instructions. Briefly, gDNA was digested with Nsp I restriction enzyme. Digested DNA was adapter ligated and PCR amplified. DNA was then fragmented and labeled with biotin. Fragmented-labeled DNA was hybridized onto GeneChip arrays in a GeneChip Hybridization Oven 645 overnight. Chips were washed and stained simultaneously using GeneChip Fluidics Station 450 and scanned using GeneChip Scanner 3000 7 G. Data were analyzed using ChAS 3.2.

**Quantitative reverse transcription-polymerase chain reaction (qRT-PCR) analysis**

Total RNA was extracted from samples resorting to the RNeasy Micro Kit (Qiagen), following the manufacturer’s instructions. cDNA was then synthesized using the qScript® cDNA SuperMix (Quantabio), according to the manufacturer’s instructions. Quantification of the gene expression was carried out using quantitative real-time PCR. All genes had their mRNA quantified using the SYBR® Green technology, except for Progerin where the Taqman technology was used. For the genes based on the SYBR® Green technology, KiCqStart® SYBR® Green primers (Sigma) diluted in Ambion® molecular biology grade Tris-EDTA (TE) solution, pH 8 (Invitrogen) were used, with a final qPCR reaction concentration of 400 nM. For the qPCR, the NZYSpeedy qPCR Green Master Mix (NZYTech) was used in a 10 𝜇L-reaction according to the manufacturer’s instructions and the following thermocycling conditions: 95°C for 2 min, followed by 40 cycles of 95°C for 5 s and 60°C for 30 s. cDNA samples were used at a final reaction concentration of 0.5 ng/𝜇L. The list of the SYBR® Green-based primers can be found in **Supporting Table 15**. For the quantification of the progerin transcript, customized progerin-specific Taqman primers (Applied Biosystems™; ThermoFisher) and probe (6-FAM™ / MGB probe; Applied Biosystems™; ThermoFisher) were used at a final reaction concentration of 900 nM and 200 nM, respectively. For normalization, Human GAPD (GAPDH) Endogenous Control (VIC™/MGB probe, primer limited) was used, according to the manufacturer’s instructions. For the TaqMan qPCR, the TaqMan™ Gene Expression Master Mix (Applied Biosystems™; ThermoFisher) was used in a 20 𝜇L-reaction according to the manufacturer’s instructions and the following thermocycling conditions: 50°C for 2 min, 95°C for 10 min, followed by 44 cycles of 95°C for 15 s and 60°C for 1 min. cDNA samples were used at a final reaction concentration of 0.625 ng/𝜇L. The sequences of the progerin-specific primers and probe can be found in **Supporting Table 16**. Both cDNA synthesis and qPCR were performed using a CFC Connect Real-Time PCR Detection System (Bio-Rad), and the respective software. The expression of the tested genes was determined relative to the housekeeping gene GAPDH gene, according to the equation: 2^[ - (Ct sample - Ct GADPH)]^. The mean minimal cycle threshold values (Ct) were calculated from triplicate reactions.

**Flow cytometry analysis**

To determine the percentage of cardiomyocytes within the differentiation cell populations over time, we measured the percentage of cells expressing cardiac troponin T (cTnT) through flow cytometry analysis.

Differentiated cells from the analyzed timepoints (days of differentiation 0, 6, 10, 13, 20, and 30) were dissociated: using TrypLE™ Express Enzyme (ThermoFisher Scientific) (5 min, 37°C) for day 0 (as described in *Cardiac differentiation protocol*); and using STEMdiff™ Cardiomyocyte Dissociation Medium (STEM Cell Technologies) (12 min, 37°C) and STEMdiff™ Cardiomyocyte Support Medium (STEM Cell Technologies) for the rest of the timepoints (as described in *Cardiomyocyte dissociation and replating*). Thereafter, cells were centrifuged (300*g*; 5 min) and the pellet resuspend and incubated in 4% PFA (15 min, RT) for fixation. Then, the pellet was washed twice with PBS (400*g*, 5 min).

To block, permeabilize and stain the fixed cells, they were resuspended and incubated (30 min, RT) in a solution of 1% BSA (Sigma) and 0.5% Tween 20 (Merck) in PBS, mixed with the conjugated antibody PE Mouse Anti-Cardiac Troponin T (1:100; 564767; BD Biosciences). Then samples were washed twice with PBS (400*g*, 5 min, 4°C). Unstained controls went through the same procedure, except for the addition of the antibody.

Samples were run and data was acquired in a BD Accuri™ C6 flow cytometer (BD Biosciences). Data were further analyzed using the FlowJo™ v10 software (BD Biosciences). Briefly, first gating was used to exclude events related to debris and then to exclude doublet events. At least, ten thousand singlet events were collected. From those events, the percentage of cTnT-positive cells was determined by defining the gate for unstained control samples at 1%, and considering cTnT-positive cells as the events in which fluorescence intensity was higher than in that unstained control gate.

**Immunocytochemistry analysis**

Cells were fixed with 4% paraformaldehyde (PFA) (10 min, RT), and washed with PBS. For permeabilization, cells were incubated with 1% Triton X-100 (Sigma) in PBS (15 min, RT) and then washed with PBS. Thereafter, cells with blocked with 5% BSA (Sigma) (1 h, RT). After blocking, the cells were incubated (1 h, RT) with anti-human primary antibodies specific for: CD31 (1:200 dilution; M0823; Agilent), H3K9me3 (1:500 dilution; ab8898; Abcam), HAND1 (1:100 dilution; AF3168; R&D Systems), MLC-2V (1:100; 10906-1-AP; Proteintech), progerin (1:50 dilution; sc-81611; Santa Cruz Biotechnology), cardiac Troponin T (cTnT) (1:200 dilution; MA5-12960), NANOG (1: 200 dilution; AF1997; R&D Systems), SOX2 (1: 200 dilution; AB5603; Merck Millipore), OCT4 (1:200 dilution; sc-5279; Santa Cruz Biotechnology), HMGB1 (1:250 dilution; ab18256; Abcam), p21 (1:100 dilution; sc-817: Santa Cruz Biotechnology), prelamin A (1:400; ANT0045; Diatheva), 𝛾H2A.X (1:1000 dilution; ab11174; Abcam), 𝛼-actinin (sarcomeric) (𝛼-SA) (1:200 dilution; A7811; Sigma), FHL2 (1:50 dilution; ab137477; Abcam) and vimentin (1:100 dilution; MA5-16409, Invitrogen). The binding of the primary antibodies was detected resorting to Alexa Fluor 488 (1:200 dilution; Thermo Fisher) and Alexa Fluor 555 (1:200 dilution; Thermo Fisher) (1 h, RT). As a negative control for unspecific binding of the secondary antibodies, the latter were added without their respective primary antibody incubation. Cell nuclei were stained with 4', 6'-diamidino-2-phenylindole (DAPI) (2 𝜇g/mL; D9542; Sigma) (5 min; RT). Then, images were acquired by the IN Cell Analyzer 2200 microscope (GE Healthcare) or using a Zeiss LSM 710 Confocal microscope (Carl Zeiss) equipment. Fiji or IN Cell Developer software’s were used to quantify the intensity or presence of fluorescence regarding the aforementioned markers, normalized by the nuclei number.

**Immunostaining for high-content imaging using Opera Phenix**

Cells were grown on black-rimed 96-well plates (Greiner Bio-One, 655090) and on day 30 were fixed with 4% paraformaldehyde Solution in PBS (4% PFA) (Boster, UK) for 15 minutes at room temperature. After washing cells with PBS twice, cells were kept in fresh PBS at 4°C in the dark until ready to be stained.

Immunocytochemistry was started by removing the PBS and permeabilizing cells in PBS supplemented with 1 mL/L Triton X-100 (ABCAM, UK) (PBST solution) for 10 minutes at room temperature. After washing, cells were stained with Cell Mask (Thermo Fisher Scientific, UK) in PBS solution (1:10,000) for 20 minutes at room temperature in the dark. Next, cells were incubated with Blocking Buffer (5% Donkey Serum in PBS) for 1 hour at room temperature. Cells were stained with 1:100 rabbit anti-MYL2, Proteintech (10906-1-AP) made in 5% Donkey Serum in Gold Buffer (pH = 7.15, 10 mM Tris, 155 mM NaCl, 2 mM EGTA, 2 mM MgCl2). Cells were incubated in antibody solutions overnight at 4°C. On the second day, antibody solutions were removed and cells were washed with PBS three times. Then, cells were incubated with Secondary Antibody Solution (5% Donkey Serum in Gold Buffer) (1:500 donkey Alexa-antibodies were used, Thermo Fisher Scientific) for 1 hour at room temperature in the dark. After, cells were washed with PBS three times and next incubated with DAPI (1:10,000) (Thermo Fisher Scientific, UK) in PBS for 10 minutes in the dark. Lastly, cells were washed with PBS twice and ready to be analyzed in the Opera Phenix High Content Screening System (PerkinElmer). Imaging was done within 7 days of staining. The cells’ phenotype was scanned using a ×40/NA 1.1 water lens. Z-stacks from -1 to 5 µm with a step size of 1 µm were acquired using excitation lasers at 405, 488, 568 and 640 nm and emission filters at 450, 540, 600 and 690 nm. The quantification and phenotype was analyzed using Harmony software 5.0. First, segmentation of nuclei and cytoplasm was performed using DAPI and cell mask after applying a smoothed filter. Determination of MYL2-positive cells was based on a cut-off baseline for MYL2 intensity per cell.

**Whole-cell protein extraction and nuclear fraction enrichment**

Whole-cell protein was isolated from 5×10^5^ iPSC-CMs. For such, cells were priorly detached and washed with DPBS and then lysed with 2× RIPA buffer (EMD Millipore Corporation) supplemented with protease/phosphatase inhibitor cocktail (Cell Signaling Technology) and let incubate for 15 min at 4 °C. Afterwards cell lysates were centrifuged at 10000*g* for 10 min at 4 °C, and the insoluble pellet was discarded. Protein was stored at -80 °C. For cellular fraction enrichment, approximately 0.5×10^6^ cells/condition were detached and washed in cold DPBS, centrifuged at 400*g* for 10 min, the supernatant discarded and the pellet stored at -80 °C. Subsequent steps were carried out accordingly to [3]. Briefly, cytosol fraction was obtained by incubating for 10 min with 120 µL of Digitonin Buffer (150 mM NaCl, 50 mM HEPES pH 7.4, 25 µg/mL digitonin (AppliChem, #A1905,0100) followed by centrifugation at 7000*g*, 10 min. The supernatant was then kept and the pellet and then homogenized and incubated with 40 µL of NP40 Buffer (150 mM NaCl, 50 mM HEPES pH 7.4, 1 % NP40) for 30 min. The cells were then centrifuged at 7000*g* for 10 min, 4 °C. The supernatant containing the organelle fraction was then kept and transferred to a clean 1.5 mL centrifuge tube while the pellet was homogenized with 90 µL RIPA Buffer (10 mM NaCl, 50 mM HEPES pH 7.4, 0.5 % Sodium Deoxycholate, 0.1 % SDS, DNase I (Qiagen) 1 U/mL) for 60 min. The solution was then centrifuged at 7800*g* for 10 min, the pellet was discarded, and the nuclear fraction (supernatant) was kept. All steps were carried out at 4 °C.

**Western blots**

Western blot

Western Blot was carried as in [4]. Briefly, tissue (50 µg) or whole (40 µg) and fractioned (12 µg) cell lysates were denaturated for 5 min in 95 °C with Laemmli Buffer, loaded into SDS-polyacrylamide gels and transferred into Amersham™ Hybond™ P 0.45 μm PVDF Membrane (Cytiva). After 1 h blocking in 5% non-fat milk on TBS-T at RT, the membranes were incubated overnight at 4 °C under continuous agitation with primary antibodies specific for: FHL2 (32 kDa; 1:500 dilution; ab137477; Abcam), mouse GAPDH (37 kDa; 1:10000 dilution; #2118S; Cell Signaling Technology), human GAPDH (37 kDa; 1:1000 dilution; sc-365062; Santa Cruz Biotechnology), lamin A/C (75/65 kDa; 1:1000 dilution; sc-7293; Santa Cruz Biotechnology) and progerin (69 kDa; 1:1000 dilution; sc-81611; Santa Cruz Biotechnology). Of note, the anti-FHL2 antibody was incubated for an increased period of 2 h at RT. Afterward, the membranes were washed in TBS-T at RT and incubated for 1 h with anti-rabbit or anti-mouse HRP-conjugated antibodies. Images were acquired in an Imager Chemi 5QE (VWR) using WesternBright™ Quantum (Advansta) substrate. Band intensity was quantified using TotalLab TL120 software (Nonlinear Dynamics Ltd).

Western blot for autophagy analyses

For the assessment of autophagic activity in ISO- and HGPS-iPSC-CMs, cells were treated continuously for 27 h with Lona (20 μM, in DMSO) and for 3 h with chloroquine (Chq or CQ) (10 μM, in water) (an autophagy inhibitor) between the 24 h and 27 h timepoints, i.e. in combination with the Lona. At the end, cells were lysed on ice in 2× RIPA buffer (EMD Millipore Corporation) supplemented with protease/phosphatase inhibitor cocktail (Cell Signaling Technology) and let incubate for 15 min at 4 °C, and the insoluble material was pelleted by centrifugation for 10 min at 16,000 g and 4 °C. The protein concentration of each sample was determined by the micro bicinchoninic acid (micro BCA) protein assay (Pierce Biotechnology). The samples were denaturated by adding 6× concentrated sample buffer (0.5 M Tris, 30% glycerol, 10% SDS, 0.6 M DTT, 0.012% bromophenol blue) and heating for 5 min at 95 °C. Samples were stored at -20 °C until use. Standard sodium dodecyl sulfate polyacrylamide gel electrophoresis (SDS-PAGE) was used to analyse protein levels. Protein samples underwent separation through electrophoresis on custom-made 4-12% polyacrylamide gel (BioRad) and transferred onto an Immobilon-P PVDF membrane (Amersham, Cytiva) using a BioRad transfer system. Membranes were blocked in 5% milk in TBST (Tris-buffered saline (TBS) with 0.1% Tween-20) at room temperature for 1 h and then incubated with the primary rabbit polyclonal anti-LC3B (#2775, Cell Signalling Technology) in a 1:1000 dilution in 1× TBST with 5% BSA overnight. Membranes were washed and then incubated for 1 h, at room temperature (RT), with a horseradish peroxidase (HRP)-linked secondary antibody, specific to rabbit immunoglobulin G (IgG) in a 1:5,000 dilution. After washing, protein immunoreactive bands were visualized by enhanced chemiluminescence with the ECL substrate (Amersham, Cytiva) on the ImageQuant 8000 imaging system (Amersham, cytiva). The membranes were reprobed with a mouse monoclonal anti-GAPDH (#MAB374, Millipore; 1:5000) antibody for equal protein loading control. The optical density of the bands was quantified using Image Lab Software (BioRad). The results were normalized to the amount of the loading control and are expressed as the relative amount compared with experiment control. Autophagy flux was calculated as in [5] or determined in the presence of the lysosomal inhibitor chloroquine, and expressed as “Autophagic flux” calculated by subtracting the densitometric value of LC3B-II – Chq from those corresponding LC3B-II + Chq values.

**Immunohistochemistry analysis**

Firstly, histological tissue sections (4 𝜇m-thick) from the heart LV of WT and LAKI mice were deparaffinized in xylene (two times, 5 min) and rehydrated through a decreasing ethanol solutions gradient: absolute ethanol (two times, 5 min), to 95% ethanol (two times, 5 min), to 70% ethanol (3 min), and then to deionized water (5 min).

In order to stain the tissues with the anti-FHL2 antibody, a heat-induced antigen retrieval procedure was first needed. Briefly, tissue sections were immersed and incubated in citrate buffer (10 mM citric acid, 0.05% Tween 20, pH 6.0) (20 min, 97 °C) followed by a cooldown (20 min, RT) and a PBS wash (5 min).

For permeabilization, sections were incubated with 1% Triton X-100 (Sigma) (15 min, RT), and then washed with PBS.

Tissue sections were blocked by resorting to 5% BSA in PBS (1 h, RT) and then incubated with the FHL2-specific antibody (1:100; ab137477; Abcam) (overnight, 4 °C). To detect the binding of the anti-FHL2 antibody, the Alexa Fluor 555 secondary antibody (1:200; Thermo Fisher) (1 h, RT) was used. Finally, nuclei were stained with DAPI (2 𝜇g/mL; D9542; Sigma) (5 min; RT), and the tissue slides were mounted using the Dako Fluorescence Mounting Medium (Agilent).

Images were acquired by the IN Cell Analyzer 2200 microscope equipment (GE Healthcare) or using a Zeiss LSM 710 Confocal microscope (Carl Zeiss).

**SA-𝛽-galactosidase assay**

Day 20 cardiomyocytes were plated at a density of 3.6 x 10^4^ cells/cm^2^ and the activity of the senescence-associated-𝛽-galactosidase (SA-𝛽-gal) was assessed at day 30 of differentiation. For that purpose, we resorted to the Senescence Detection Kit (ab65351, Abcam) using an adaptation of the manufacturer’s protocol to the 96-well microplates. Cells were fixed by incubating them with the Fixative Solution (12 min, RT), then washed three times with PBS. Thereafter, cells were stained by incubating them with the Staining Solution mixture (overnight, 37°C, protected from light, without CO_2_ injection), which comprised of a Staining Solution mixed with 1% of Staining Supplement and 5% of the X-Gal substrate (20 mg/mL in DMSO), and, finally, washed three times with PBS and the nuclei stained with DAPI (2 𝜇g/mL; D9542; Sigma) (5 min; RT).

Using IN Cell Analyzer 2200 (GE Healthcare), images were acquired in the DAPI and brightfield filters for further quantification. For colored representative images, the Zeiss Observer Z1 microscope (Carl Zeiss) with a 40x FLUAR objective (Carl Zeiss) was used.

For quantification, nuclei counting was performed automatically using the QuPath-0.2.3 software and SA-𝛽-gal positive cells were manually counted.

**Multiplex analysis of cytokines for SASP evaluation**

Prior to running the multiplex analysis assay, conditioned medium from independent batches of ISO- and HGPS-iPSC-CMs was collected at day 30 of differentiation, after being in contact with the cells for 3 days. The conditioned medium was stored at -80°C until being processed. For the multiplex analysis, it was used the Luminex xMAP technology for multiplexed quantification of 96 Human cytokines, chemokines, and growth factors; or 9 human matrix metalloproteinases and 4 tissue inhibitors of metalloproteinases. The multiplexing analysis was performed using the Luminex™ 200 system (Luminex, Austin, TX, USA) by Eve Technologies Corp. (Calgary, Alberta). Ninety-six markers were simultaneously measured in the samples using Eve Technologies' Human Cytokine 96-Plex Discovery Assay® which consists of two separate kits; the Panel A 48-plex and the Panel B 48-plex (MilliporeSigma, Burlington, Massachusetts, USA). Thirteen markers were simultaneously measured in the samples using Eve Technologies' Human MMP/TIMP 13-Plex Discovery Assay® which consists of two separate kits; one 9 plex and one 4 plex (R&D Systems, Inc., Minneapolis, MN, USA). The assays were run according to the manufacturer's protocol. Assay sensitivities of the markers range from 0.05 – 100 pg/mL for the 96-plex, and 0.28 – 253 pg/mL for the 13-Plex. For the 96-Plex, individual analyte sensitivity values are available in the MILLIPLEX® MAP protocol. Individual analyte sensitivity values are available in the product datasheet for the 4-Plex and by building the panel on the R&D Systems Magnetic Luminex Performance product page for the 9-Plex.

**Oxidative stress determination**

Day 20 cardiomyocytes were plated in 96 well microplates, previously coated with Matrigel, at a density of 2.5 x 10^5^ cells/cm^2^. On day 30, cardiomyocytes were stained with the ROS-sensitive fluorescent probe CellROX® Deep Red (5 𝜇M; C10422; Thermo Fisher Scientific), by incubating it with the cells (2 hrs, 37°C) and then fixing the cells with a 2.5% formaldehyde solution (30 min, RT), followed by two washes with PBS, and the nuclei stained with DAPI (2 𝜇g/mL; D9542; Sigma) (5 min; RT), for nuclei counting.

Images were acquired with a 40x objective from the IN Cell Analyzer 2200 (GE Healthcare) equipment and using the Cy5 channel. For oxidative stress assessment, images were analyzed through the IN Cell Developer software to calculate the CellROX® fluorescence intensity. Briefly, the CellROX-expressing area (A) was segmented by a minimum intensity threshold, then the mean fluorescence intensity within the area was calculated (D), and the multiplication of both parameters (DxA) was considered for the determination. Finally, DxA was normalized by nuclei number.

**mCherry-eGFP-LC3B tandem reporter assay**

U2OS cells stably expressing the mCherry-eGFP-LC3B tandem reporter and kept at below 10 passages were plated into 384-well black, optically-clear cell culture plates (PerkinElmer PhenoPlate) using a MultiDrop Combi liquid handling system (ThermoFisher) at 2,500 cells per well in 50 µl per well of cell culture media containing DMEM supplemented with 10% FBS (Invitrogen). Cells were then incubated overnight at 37 °C in 5% CO2 to allow cell adhesion and equilibrium. Compounds, initially prepared as 50 mM stocks, were arrayed in a 384-well compound plate (Costar V-bottom) and each plate design was dispensed using an Echo 650 Acoustic liquid handler (LabCyte), with the final DMSO levels across the wells being held constant relative to the total compound dispensed for that assay plate. On the day of the treatment, 25 µl of pre-warmed media was dispensed across the compound plate and the plate was incubated for 30 minutes to allow sufficient mixing. Using the CyBio Felix liquid dispenser, 40 µl of media was aspirated from the cell culture plates and replaced with 35 µl of fresh media and 5 µl of compound solutions, such that the final concentrations of the compound added to the cells ranged from 10 µM to 1 nM in a ten-point dose-response. The cell culture plates were then incubated for a further 24 h, before the media was aspirated using the CyBio Felix and replaced with 40 µl of fixing solution (4% formaldehyde, 1% glutaraldehyde in dPBS) using the Combi system and fixing was allowed to proceed in the dark at room temperature for 10 minutes. Afterwards, the fixing solution was removed by inverting the plate on a paper towel and tapping and replaced with 50 µl of PBS containing 5 µg/mL of Hoechst nuclear stain. The plates were then sealed and stored at 4 °C until imaging.

Cells were imaged using a Opera Phenix (Revvity) and a 40x water objective in confocal mode, 2 binning. 11 fields of view were captured in the middle of each well with two Z-stacks 0.5 µM apart. Data were collected in the DAPI, GFP and mCherry channels and analysed using the Harmony image analysis software (Revvity) as follows. Cells were segmented using a 3 pixel gaussian of the DAPI channel to segment their nuclei. Cytoplasmic areas were then segmented using a mask generated from the sum of the three channels.

To segment puncta, the GFP and mCherry channels were first background subtracted using a sliding parabola filter of 0.01. These background subtracted channels were filtered by a 12 pixel median blur. Enhanced GFP and RFP images were then produced by diving the background subtracted images by the median blur images. RFP puncta with cells were identified from the enhanced RFP channel using the Find Spots Harmony building block, then spots with a size less than 2 pixels were excluded. The ratio of RFP:GFP inside these puncta was calculated from the enhanced GFP and mCherry channels. Puncta were classified into autophagosomes or autolysosomes based on their RFP:GFP ratio, with a threshold set based on 0.2 µM Bafilomycin A1 and 0.2 µM Torin1 controls. Treatment with Bafilomycin A1 blocks autophagosome maturation into lysosomes, which results in the accumulation of autophagosomes, whereas treatment with the autophagy stimulator Torin1 results in autolysosome accumulation. The area of autolysosomes per cell as well as the percentage of puncta which were autolysosomes (the autophagic flux) were plotted against the concentrations of compound on a log scale.

Dose-response data were baseline subtracted by subtracting the values from the DMSO control. Dose-response curves were fitted to a log (agonist) vs response curve modified with a hill co-efficient to generate EC50s:

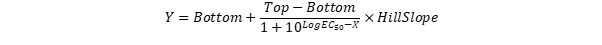

*Equation. 1 Dose-response curve modified with a hill-coefficient for determining autophagy activity EC_50_s*

Where Bottom = the lower limit of the curve, Top = upper limit of the curve, EC50 = the middle of curve inflection and HillSlope = the cooperativity coefficient.  The Bottom was constrained to 0 and the hill slope was constrained to <2.

**DQ-BSA assay**

Autophagic activity can also be measured by monitoring the uptake and degradation of DQ-BSA via confocal microscopy. DQ-BSA is a highly labelled conjugate of the BSA protein with the BODIPY® TR-X dye. In its native state the fluorescence of the conjugate is quenched. DQ-BSA fed to cells in endocytosed and degraded via the autophagic pathway. Upon cleavage by proteases the dye within the conjugate is released and able to fluoresce.

U2OS kept below 10 passages were plated into 384-well black, optically clear cell culture plates (PerkinElmer PhenoPlate) using a MultiDrop Combi liquid handling system (ThermoFisher) at 2,500 cells per well in 50 µL per well of cell culture media containing DMEM supplemented with 10% FBS (Invitrogen). Cells were then incubated overnight at 37 °C in 5% CO_2_ to allow cell adhesion and equilibrium. Compounds, initially prepared as 50 mM stocks, were arrayed in a 384-well compound plate (Costar V-bottom) and each plate design was dispensed using an Echo 650 Acoustic liquid handler (LabCyte), with the final DMSO levels across the wells being held constant relative to the total compound dispensed for that assay plate. On the day of the treatment, 25 µL of pre-warmed media was dispensed across the compound source plate and the plate was incubated for 30 min to allow sufficient mixing. Using the CyBio Felix liquid handler, 40 µL of media was aspirated from the cell culture plates and replaced with 35 µL of fresh media and 5 µL of compound solutions, such that the final concentrations of the compound added to the cells ranged from 10 µM to 1 nM in a ten-point dose-response.

After 18 h, a 5 µL of a solution containing DQ-BSA Red and Lysotracker in media stains was added to a final concentration of 5 µg/mL Hoechst 33342 and 25 µg /mL DQ-BSA Red (Invitrogen). The cell culture plates were then incubated for a further 6 h, before 40 µl media was aspirated using the CyBio Felix and replaced with fresh media.

Cells were imaged and analyzed as described above, except that the cells were maintained at 37 ºC and 5% CO_2_ throughout the imaging process. The mCherry channel was used separately to identify proteolytically active lysosomes/autolysosomes. The area of mCherry positive puncta a per cell was plotted to generate EC50s as described above.

**High-content proteomic analysis (LV human and mouse tissue)**

### The mice and human mass spectrometry proteomics data underlying this article was based on a previously published dataset [6], and is available in ProteomeXchange Consortium via the PRIDE [7] partner repository, and can be accessed with the dataset identifier PXD039548.

### Mouse LV tissue

This study was conducted in accordance with the guidelines of Directive 2010/63/EU of the European Parliament on the protection of animals used for scientific purposes and was approved by the local Ethical Committees. Aged (20 months-old) and young (3 months-old) WT C57BL/6 mice were obtained from in-house breeding at the Leibniz Institute on Ageing – Fritz Lipmann Institute (FLI) and at UC-Biotech (University of Coimbra, Coimbra), respectively. Animals were housed in a dedicated pathogen-free animal facility with a 12-h light/dark cycle and fed ad libitum with a standard diet. For these animals fresh LV tissue was used. Moreover, FFPE mouse heart samples from LmnaG609G/G609G mice (10 weeks-old) and age-matched WT mice were kindly provided by Annachiara De Sandre-Giovannoli.

### Human LV tissue

Biopsies of human LV were obtained from Coimbra Hospital University Centre (CHUC, Coimbra, Portugal) from four elderly donors (65 to 70 years old) and one young donor (31 years old), who had no cardiac pathology. These samples were not suitable for transplantation, in agreement with the regulations of the tissue bank and the approval of the Ethics Committee of the Faculty of Medicine of the University of Coimbra (CE-022/2017). Additional 3 human LV FFPE samples from young donors (20 and 21 years old) were kindly provided by the Centre for Heart Lung Innovation Providence Health Care / University of British Columbia, Vancouver, BC, Canada. Clinical details for both elderly and young donors are outlined in [6]. FFPE tissue sections from LV tissue of two HGPS patient donors (10 and 14 years old) were provided by Progeria Research Foundation (PRF). As per the Common Rule and National Law No. 12/93 of April 22, deceased individuals are not considered human subjects, so informed consent was not required for postmortem sample use. However, the study was approved by the ethics committee of the Faculty of Medicine at the University of Coimbra (CE -022/2017) and followed institutional guidelines and the Declaration of Helsinki principles.

**Progerin protein staining in LV tissue**

Formalin-fixed paraffin included slices were deparaffinized and rehydrated with subsequential submersion in xylene (2×5 min), ethanol 100% ( 2×5 min), ethanol 96% (2×3 min), ethanol 70% (3 min), ethanol 40% (3 min) and dH_2_O (5 min). Heat-induced antigen retrieval was carried in a water bath with Tris-EDTA buffer (pH 9, 10 mM Tris Base, 1 mM EDTA solution, 0.05% Tween-20) at 97 ºC for 20 min. The slides were let cool down for 20 min, and washed in PBS. Then the tissues were permeabilized with 1% Triton for 15 min, washed 3× PBS 1×, and blocked for 1 h RT with 5% 3× in PBS 1× for 5 min RT, and incubated with secondary Goat anti-Rabbit IgG, BSA, and incubated with anti-progerin (1:50, Santa Cruz Biotechnology, ref sc-81611) for 48 h in a humid chamber at 4 ºC. The samples were then washed and incubated with goat anti-mouse Alexa Fluor™ 555 (1:500, Invitrogen, cat no. A21422) for 24 h, 4 ºC. The samples were stained with 2 µg/mL DAPI (20 min) and the slides were mounted with Fluorescent Mounting Media (Dako, ref 2S302380-2) and let dry for 24 h. All the antibodies were diluted in a blocking solution and incubated 16-24 h at 4 ºC, and the sample was washed between every step in 1x PBS for 5 min RT. Capture was carried out in LSM 710 confocal microscope (ZEISS) with the Plan-Apochromat 40x/1,4 OIL DIC M27 objective.

Maximum intensity projections were quantified in Qupath Software. Nuclei were detected resorting to a nuclei detection script. False positive nuclei detection and nuclei in high autofluorescent areas were manually removed. Mean nuclei intensity was then exported directly from Qupath. To remove background variation, 3 different ROI were defined per image of areas solely with tissue autofluorescence, and the mean background value was discounted from the mean nuclei intensity per image.

Supplementary Tables

**Supplementary Data Table 1. List of significantly altered senescence-associated proteins (*p*-value<0.05) when compared with ISO-iPSC-CMs**

| **Uniprot ID** | **Protein Symbol** | **Protein Description** | **Average Log2 Ratio** | ***p*-value** | ***q*-value** | **% of change** | **# of Ratios** | **Ratio** | **# Unique Total Peptides** |
| --- | --- | --- | --- | --- | --- | --- | --- | --- | --- |
| P12235 | ADT1 | ADP/ATP translocase 1 | -0,214348444 | 6,58E-05 | 0,0004137 | -13,80646576 | 102 | 0,861935342 | 13 |
| P05141 | ADT2 | ADP/ATP translocase 2 | 0,121889727 | 0,00336 | 0,01121318 | 8,815926432 | 327 | 1,088159264 | 36 |
| Q13015 | AF1Q | Protein AF1q | 0,6804972 | 0,049398 | 0,08771464 | 60,26920006 | 37 | 1,602692001 | 5 |
| Q02952 | AKA12 | A-kinase anchor protein 12 | 0,434870588 | 1,72E-08 | 2,52E-07 | 35,17895698 | 348 | 1,35178957 | 48 |
| P15121 | ALDR | Aldose reductase | -0,791141837 | 3,89E-07 | 4,37E-06 | -42,21134637 | 223 | 0,577886536 | 22 |
| P39687 | AN32A | Acidic leucine-rich nuclear phosphoprotein 32 family member A | 0,606499746 | 0,009527 | 0,0257304 | 52,25607007 | 153 | 1,522560701 | 15 |
| Q9BTT0 | AN32E | Acidic leucine-rich nuclear phosphoprotein 32 family member E | 0,304877978 | 0,034048 | 0,06711549 | 23,53141511 | 60 | 1,235314151 | 8 |
| Q9NQW6 | ANLN | Actin-binding protein anillin | 1,560679491 | 0,000628 | 0,00281162 | 194,9927486 | 40 | 2,949927486 | 6 |
| Q9BQE5 | APOL2 | Apolipoprotein L2 | -0,250892403 | 0,000765 | 0,00330103 | -15,96235743 | 56 | 0,840376426 | 8 |
| P08243 | ASNS | Asparagine synthetase [glutamine-hydrolyzing] | -0,676592107 | 3,34E-08 | 4,63E-07 | -37,43585976 | 135 | 0,625641402 | 24 |
| Q9UIG0 | BAZ1B | Tyrosine-protein kinase BAZ1B | 0,007938428 | 0,03169 | 0,06369421 | 0,551766548 | 63 | 1,005517665 | 10 |
| Q9Y276 | BCS1 | Mitochondrial chaperone BCS1 | -0,476535696 | 0,04616 | 0,08349909 | -28,12986486 | 47 | 0,718701351 | 8 |
| P16278 | BGAL | Beta-galactosidase | -0,38946025 | 0,000648 | 0,00288508 | -23,65848355 | 103 | 0,763415165 | 15 |
| Q8TDN6 | BRX1 | Ribosome biogenesis protein BRX1 homolog | 0,205625552 | 0,039944 | 0,07523408 | 15,31862595 | 47 | 1,15318626 | 7 |
| Q9NW68 | BSDC1 | BSD domain-containing protein 1 | 1,29259651 | 0,017391 | 0,04064803 | 144,9685445 | 8 | 2,449685445 | 2 |
| Q05682 | CALD1 | Caldesmon | 0,034246015 | 1,50E-08 | 2,23E-07 | 2,402150673 | 401 | 1,024021507 | 45 |
| P07858 | CATB | Cathepsin B | -0,0977153 | 0,011929 | 0,0306096 | -6,548825561 | 89 | 0,934511744 | 10 |
| P07339 | CATD | Cathepsin D | 0,110463998 | 1,14E-06 | 1,15E-05 | 7,957539246 | 279 | 1,079575392 | 31 |
| P35520 | CBS | Cystathionine beta-synthase | 1,158794453 | 4,11E-09 | 6,76E-08 | 123,2707798 | 122 | 2,232707798 | 18 |
| Q13185 | CBX3 | Chromobox protein homolog 3 | 0,136656815 | 0,028167 | 0,05843899 | 9,935460591 | 113 | 1,099354606 | 14 |
| P06493 | CDK1 | Cyclin-dependent kinase 1 | 0,302795352 | 0,003319 | 0,01109075 | 23,35321796 | 67 | 1,23353218 | 9 |
| P24941 | CDK2 | Cyclin-dependent kinase 2 | 0,085495353 | 0,002061 | 0,00753295 | 6,105199409 | 74 | 1,061051994 | 9 |
| Q15021 | CND1 | Condensin complex subunit 1 | 0,538191222 | 0,026889 | 0,05637765 | 45,21507424 | 21 | 1,452150742 | 4 |
| P51911 | CNN1 | Calponin-1 | 0,745109632 | 3,73E-05 | 0,00025079 | 67,61016331 | 287 | 1,676101633 | 33 |
| P12111 | CO6A3 | Collagen alpha-3(VI) chain | -1,340829916 | 0,002307 | 0,00824144 | -60,52065158 | 22 | 0,394793484 | 6 |
| Q96F07 | CYFP2 | Cytoplasmic FMR1-interacting protein 2 | -0,168220875 | 0,009804 | 0,02630805 | -11,0060526 | 56 | 0,889939474 | 10 |
| O94760 | DDAH1 | N(G),N(G)-dimethylarginine dimethylaminohydrolase 1 | 0,643701878 | 0,032657 | 0,06507056 | 56,23328833 | 109 | 1,562332883 | 11 |
| O43143 | DHX15 | Pre-mRNA-splicing factor ATP-dependent RNA helicase DHX15 | -0,473618562 | 0,000275 | 0,00140827 | -27,98439621 | 270 | 0,720156038 | 36 |
| O60832 | DKC1 | H/ACA ribonucleoprotein complex subunit 4 | -0,28544921 | 0,00885 | 0,02427499 | -17,95139045 | 159 | 0,820486095 | 22 |
| P26358 | DNMT1 | DNA (cytosine-5)-methyltransferase 1 | 0,687125454 | 0,040335 | 0,07577319 | 61,00722778 | 59 | 1,610072278 | 8 |
| O14531 | DPYL4 | Dihydropyrimidinase-related protein 4 | -0,7143583 | 0,000202 | 0,00108285 | -39,05238381 | 33 | 0,609476162 | 5 |
| P06733 | ENOA | Alpha-enolase | 0,513747022 | 0,006697 | 0,01949831 | 42,77536001 | 608 | 1,4277536 | 53 |
| Q13642 | FHL1 | Four and a half LIM domains protein 1 | 1,109763806 | 0,012821 | 0,03232601 | 115,8103125 | 78 | 2,158103125 | 10 |
| Q14315 | FLNC | Filamin-C | 0,486166281 | 4,58E-19 | 2,85E-17 | 40,07177545 | 2601 | 1,400717755 | 247 |
| Q96AE4 | FUBP1 | Far upstream element-binding protein 1 | 0,094257133 | 0,00141 | 0,00550104 | 6,751558583 | 339 | 1,067515586 | 40 |
| Q92945 | FUBP2 | Far upstream element-binding protein 2 | 0,180384246 | 0,011615 | 0,02999955 | 13,31856568 | 362 | 1,133185657 | 39 |
| P07954 | FUMH | Fumarate hydratase, mitochondrial | 0,014229594 | 0,00033 | 0,0016372 | 0,991200474 | 373 | 1,009912005 | 37 |
| P35637 | FUS | RNA-binding protein FUS | 0,577613053 | 0,043772 | 0,08043782 | 49,23780569 | 164 | 1,492378057 | 16 |
| Q9NZC3 | GDE1 | Glycerophosphodiester phosphodiesterase 1 | -0,077712028 | 0,00547 | 0,01657991 | -5,244080904 | 22 | 0,947559191 | 3 |
| Q06210 | GFPT1 | Glutamine--fructose-6-phosphate aminotransferase [isomerizing] 1 | -0,283251797 | 0,006645 | 0,0193748 | -17,8263245 | 143 | 0,821736755 | 19 |
| O94925 | GLSK | Glutaminase kidney isoform, mitochondrial | -0,210474409 | 0,006808 | 0,01977074 | -13,57470124 | 187 | 0,864252988 | 28 |
| P34897 | GLYM | Serine hydroxymethyltransferase, mitochondrial | 0,117387778 | 0,033574 | 0,06641509 | 8,47689418 | 276 | 1,084768942 | 36 |
| P35052 | GPC1 | Glypican-1 | -0,108741663 | 0,037544 | 0,07199756 | -7,260340346 | 30 | 0,927396597 | 5 |
| P49915 | GUAA | GMP synthase [glutamine-hydrolyzing] | 0,13354546 | 0,036023 | 0,06985996 | 9,698626317 | 242 | 1,096986263 | 33 |
| P16401 | H15 | Histone H1,5 | 0,826229746 | 0,017243 | 0,04039446 | 77,30457296 | 210 | 1,77304573 | 15 |
| Q9BQS7 | HEPH | Hephaestin | -1,785142685 | 2,63E-05 | 0,00018481 | -70,98527188 | 14 | 0,290147281 | 3 |
| P07686 | HEXB | Beta-hexosaminidase subunit beta | -0,77345574 | 3,34E-06 | 3,00E-05 | -41,49855116 | 69 | 0,585014488 | 13 |
| P05204 | HMGN2 | Non-histone chromosomal protein HMG-17 | 2,299518665 | 0,016398 | 0,0389343 | 392,293491 | 98 | 4,92293491 | 10 |
| P52597 | HNRPF | Heterogeneous nuclear ribonucleoprotein F | -0,033184335 | 0,038124 | 0,07273773 | -2,273910777 | 151 | 0,977260892 | 15 |
| P52272 | HNRPM | Heterogeneous nuclear ribonucleoprotein M | 1,002605756 | 0,00234 | 0,00833856 | 100,3615609 | 647 | 2,003615609 | 67 |
| P54652 | HSP72 | Heat shock-related 70 kDa protein 2 | 0,954728825 | 0,00029 | 0,00146964 | 93,82152841 | 230 | 1,938215284 | 26 |
| Q92743 | HTRA1 | Serine protease HTRA1 | -1,126718116 | 0,033831 | 0,0667833 | -54,20436868 | 18 | 0,457956313 | 3 |
| Q16270 | IBP7 | Insulin-like growth factor-binding protein 7 | 0,511962787 | 0,017247 | 0,04039641 | 42,59889351 | 30 | 1,425988935 | 4 |
| P38919 | IF4A3 | Eukaryotic initiation factor 4A-III | -0,343913149 | 0,005442 | 0,01651522 | -21,20986836 | 165 | 0,787901316 | 22 |
| Q12905 | ILF2 | Interleukin enhancer-binding factor 2 | -0,264017132 | 0,001887 | 0,00700171 | -16,72341184 | 183 | 0,832765882 | 22 |
| P52292 | IMA1 | Importin subunit alpha-1 | 0,601038892 | 3,17E-06 | 2,87E-05 | 51,68084346 | 99 | 1,516808435 | 13 |
| Q9UIJ7 | KAD3 | GTP:AMP phosphotransferase AK3, mitochondrial | 0,070830081 | 0,000154 | 0,00085477 | 5,032083088 | 234 | 1,050320831 | 27 |
| Q07666 | KHDR1 | KH domain-containing, RNA-binding, signal transduction-associated protein 1 | 0,529006598 | 0,02012 | 0,0454762 | 44,2935286 | 215 | 1,442935286 | 24 |
| P42167 | LAP2B | Lamina-associated polypeptide 2, isoforms beta/gamma | 0,262058986 | 0,00641 | 0,01881611 | 19,91889427 | 266 | 1,199188943 | 31 |
| Q9NZU5 | LMCD1 | LIM and cysteine-rich domains protein 1 | -0,039751334 | 0,01048 | 0,02770526 | -2,717738952 | 85 | 0,97282261 | 11 |
| P20700 | LMNB1 | Lamin-B1 | 0,341776017 | 0,002759 | 0,00951467 | 26,73157519 | 714 | 1,267315752 | 70 |
| Q03252 | LMNB2 | Lamin-B2 | 0,2808116 | 0,015162 | 0,03670352 | 21,48781321 | 511 | 1,214878132 | 64 |
| Q07954 | LRP1 | Prolow-density lipoprotein receptor-related protein 1 | -0,158162795 | 0,002737 | 0,00945474 | -10,38344304 | 54 | 0,89616557 | 10 |
| Q9Y333 | LSM2 | U6 snRNA-associated Sm-like protein LSm2 | -0,160787483 | 0,012319 | 0,03135174 | -10,54633375 | 42 | 0,894536662 | 5 |
| P62312 | LSM6 | U6 snRNA-associated Sm-like protein LSm6 | -0,491962197 | 0,007546 | 0,02145594 | -28,8942663 | 54 | 0,711057337 | 8 |
| P28300 | LYOX | Protein-lysine 6-oxidase | -0,700387795 | 0,028263 | 0,05857743 | -38,45932363 | 13 | 0,615406764 | 2 |
| O43772 | MCAT | Mitochondrial carnitine/acylcarnitine carrier protein | -0,572847584 | 0,00135 | 0,00530266 | -32,77114817 | 42 | 0,672288518 | 7 |
| P33991 | MCM4 | DNA replication licensing factor MCM4 | 0,674463682 | 0,021447 | 0,0477528 | 59,60033526 | 208 | 1,596003353 | 29 |
| Q08431 | MFGM | Lactadherin | 0,508609169 | 0,038519 | 0,07325259 | 42,26780002 | 82 | 1,422678 | 12 |
| Q9NPC6 | MYOZ2 | Myozenin-2 | 0,34034488 | 0,000132 | 0,00075066 | 26,60592122 | 333 | 1,266059212 | 30 |
| P49321 | NASP | Nuclear autoantigenic sperm protein | 0,429098749 | 0,000142 | 0,00079637 | 34,63922237 | 389 | 1,346392224 | 45 |
| Q0ZGT2 | NEXN | Nexilin | 0,583897552 | 0,036754 | 0,0708663 | 49,88931593 | 612 | 1,498893159 | 62 |
| Q92542 | NICA | Nicastrin | -0,197224003 | 0,000536 | 0,00247092 | -12,7772733 | 41 | 0,872227267 | 6 |
| O00567 | NOP56 | Nucleolar protein 56 | 0,009553774 | 0,039235 | 0,07427007 | 0,664414666 | 295 | 1,006644147 | 32 |
| P06748 | NPM | Nucleophosmin | 1,086080869 | 0,006111 | 0,01811882 | 112,2965416 | 378 | 2,122965416 | 30 |
| Q9H857 | NT5D2 | 5'-nucleotidase domain-containing protein 2 | -0,338422026 | 0,026341 | 0,05552993 | -20,90940934 | 64 | 0,790905907 | 11 |
| Q8NFH5 | NUP53 | Nucleoporin NUP53 | 0,699391202 | 0,021492 | 0,04782775 | 62,38194176 | 63 | 1,623819418 | 8 |
| Q9UQ80 | PA2G4 | Proliferation-associated protein 2G4 | 0,073439481 | 0,015064 | 0,03653049 | 5,222226342 | 250 | 1,052222263 | 31 |
| O95340 | PAPS2 | Bifunctional 3'-phosphoadenosine 5'-phosphosulfate synthase 2 | -0,269169235 | 0,010697 | 0,02812279 | -17,02027591 | 32 | 0,829797241 | 6 |
| P09874 | PARP1 | Poly [ADP-ribose] polymerase 1 | -0,012195884 | 0,001777 | 0,00666217 | -0,841791203 | 525 | 0,991582088 | 56 |
| Q16822 | PCKGM | Phosphoenolpyruvate carboxykinase [GTP], mitochondrial | -0,122920658 | 2,83E-05 | 0,00019696 | -8,167333536 | 201 | 0,918326665 | 31 |
| Q9UHG3 | PCYOX | Prenylcysteine oxidase 1 | -0,422817117 | 0,014555 | 0,03560938 | -25,40334304 | 103 | 0,74596657 | 15 |
| Q15119 | PDK2 | [Pyruvate dehydrogenase (acetyl-transferring)] kinase isozyme 2, mitochondrial | -0,719116295 | 0,012271 | 0,03126263 | -39,25305737 | 44 | 0,607469426 | 8 |
| Q01813 | PFKAP | ATP-dependent 6-phosphofructokinase, platelet type | -0,342637314 | 1,75E-06 | 1,70E-05 | -21,14016012 | 590 | 0,788598399 | 58 |
| Q99623 | PHB2 | Prohibitin-2 | -0,288608048 | 1,79E-10 | 3,80E-09 | -18,13084259 | 264 | 0,818691574 | 33 |
| Q02809 | PLOD1 | Procollagen-lysine,2-oxoglutarate 5-dioxygenase 1 | -0,391588515 | 0,002448 | 0,00864673 | -23,7710196 | 107 | 0,762289804 | 18 |
| P36873 | PP1G | Serine/threonine-protein phosphatase PP1-gamma catalytic subunit | 0,032275442 | 0,001439 | 0,00559614 | 2,262375297 | 97 | 1,022623753 | 10 |
| P10619 | PPGB | Lysosomal protective protein | -0,140691275 | 1,76E-06 | 1,71E-05 | -9,291558315 | 62 | 0,907084417 | 7 |
| P60891 | PRPS1 | Ribose-phosphate pyrophosphokinase 1 | -0,039392952 | 0,013617 | 0,03387751 | -2,693569868 | 182 | 0,973064301 | 23 |
| Q9BRX5 | PSF3 | DNA replication complex GINS protein PSF3 | 0,870324009 | 0,029788 | 0,06090171 | 82,8073414 | 21 | 1,828073414 | 4 |
| Q9BT73 | PSMG3 | Proteasome assembly chaperone 3 | -1,215925023 | 0,030724 | 0,06234082 | -56,95030335 | 12 | 0,430496966 | 2 |
| Q15262 | PTPRK | Receptor-type tyrosine-protein phosphatase kappa | 0,840343308 | 0,00391 | 0,01263607 | 79,04761584 | 85 | 1,790476158 | 14 |
| P28827 | PTPRM | Receptor-type tyrosine-protein phosphatase mu | 0,104941625 | 0,048131 | 0,08607638 | 7,545087418 | 36 | 1,075450874 | 5 |
| O15067 | PUR4 | Phosphoribosylformylglycinamidine synthase | 0,223909363 | 0,002237 | 0,00804881 | 16,78940198 | 353 | 1,16789402 | 42 |
| P22234 | PUR6 | Multifunctional protein ADE2 | -0,372755754 | 0,021306 | 0,04751032 | -22,76941277 | 210 | 0,772305872 | 25 |
| Q9NRW1 | RAB6B | Ras-related protein Rab-6B | -0,720318363 | 0,019377 | 0,04411588 | -39,30365128 | 23 | 0,606963487 | 3 |
| P38159 | RBMX | RNA-binding motif protein, X chromosome | 0,50015237 | 0,001509 | 0,00582323 | 41,43629323 | 508 | 1,414362932 | 51 |
| Q96D15 | RCN3 | Reticulocalbin-3 | -0,739108828 | 0,004324 | 0,01374859 | -40,0890684 | 33 | 0,599109316 | 6 |
| P35244 | RFA3 | Replication protein A 14 kDa subunit | 0,38118738 | 0,011755 | 0,03024841 | 30,24133384 | 42 | 1,302413338 | 6 |
| Q7LG56 | RIR2B | Ribonucleoside-diphosphate reductase subunit M2 B | -0,479434171 | 0,047232 | 0,08491271 | -28,27411204 | 40 | 0,71725888 | 6 |
| Q9BYD6 | RM01 | 39S ribosomal protein L1, mitochondrial | -0,603806162 | 0,001348 | 0,00529666 | -34,19843335 | 66 | 0,658015666 | 10 |
| Q16540 | RM23 | 39S ribosomal protein L23, mitochondrial | -0,126676956 | 0,014833 | 0,03611453 | -8,406124248 | 52 | 0,915938758 | 7 |
| P09651 | ROA1 | Heterogeneous nuclear ribonucleoprotein A1 | 0,740113616 | 0,025628 | 0,05440725 | 67,03073745 | 412 | 1,670307375 | 35 |
| P22626 | ROA2 | Heterogeneous nuclear ribonucleoproteins A2/B1 | 0,707155407 | 0,036093 | 0,06995511 | 63,25819445 | 512 | 1,632581945 | 44 |
| P82933 | RT09 | 28S ribosomal protein S9, mitochondrial | -0,009492618 | 0,040122 | 0,07549893 | -0,655818219 | 147 | 0,993441818 | 20 |
| P82914 | RT15 | 28S ribosomal protein S15, mitochondrial | -0,545888746 | 0,000131 | 0,00074886 | -31,50306906 | 84 | 0,684969309 | 13 |
| Q9Y265 | RUVB1 | RuvB-like 1 | -0,032512865 | 0,009098 | 0,02480114 | -2,228415747 | 217 | 0,977715843 | 26 |
| Q15393 | SF3B3 | Splicing factor 3B subunit 3 | -0,431614235 | 0,000404 | 0,00194629 | -25,85682685 | 206 | 0,741431732 | 31 |
| P23246 | SFPQ | Splicing factor, proline- and glutamine-rich | 0,579418951 | 0,006725 | 0,01955864 | 49,42473156 | 600 | 1,494247316 | 63 |
| Q9NTJ3 | SMC4 | Structural maintenance of chromosomes protein 4 | 0,012602764 | 0,007281 | 0,02082652 | 0,877383652 | 90 | 1,008773837 | 14 |
| P51688 | SPHM | N-sulphoglucosamine sulphohydrolase | -0,692253572 | 0,002479 | 0,00873523 | -38,11136413 | 35 | 0,618886359 | 5 |
| P16949 | STMN1 | Stathmin | 0,976584828 | 0,019088 | 0,04359446 | 96,78016772 | 294 | 1,967801677 | 32 |
| P14868 | SYDC | Aspartate--tRNA ligase, cytoplasmic | 0,020905869 | 3,31E-06 | 2,98E-05 | 1,459634532 | 206 | 1,014596345 | 30 |
| Q6PI48 | SYDM | Aspartate--tRNA ligase, mitochondrial | 0,054584093 | 0,48114 | 0,37925428 | 3,85596595 | 121 | 1,03855966 | 17 |
| P68371 | TBB4B | Tubulin beta-4B chain | 0,2922703 | 0,019268 | 0,04391617 | 22,45658023 | 463 | 1,224565802 | 35 |
| Q13428 | TCOF | Treacle protein | 0,682056439 | 0,002202 | 0,00794414 | 60,44250972 | 297 | 1,604425097 | 39 |
| P21980 | TGM2 | Protein-glutamine gamma-glutamyltransferase 2 | 0,007570175 | 0,001261 | 0,00500848 | 0,526103655 | 487 | 1,005261037 | 48 |
| Q86V81 | THOC4 | THO complex subunit 4 | 0,833470915 | 0,009379 | 0,02540553 | 78,19673648 | 128 | 1,781967365 | 14 |
| Q13263 | TIF1B | Transcription intermediary factor 1-beta | 0,350329645 | 0,014609 | 0,03570728 | 27,48518885 | 412 | 1,274851889 | 47 |
| Q9NQ88 | TIGAR | Fructose-2,6-bisphosphatase TIGAR | -0,614753512 | 2,05E-05 | 0,00014822 | -34,69585424 | 56 | 0,653041458 | 8 |
| Q9NQ34 | TMM9B | Transmembrane protein 9B | -0,579162463 | 0,034141 | 0,06725127 | -33,0647752 | 13 | 0,669352248 | 3 |
| O96008 | TOM40 | Mitochondrial import receptor subunit TOM40 homolog | -0,314214133 | 0,005694 | 0,01712914 | -19,57110151 | 75 | 0,804288985 | 10 |
| P11388 | TOP2A | DNA topoisomerase 2-alpha | 1,695561909 | 0,002843 | 0,00974417 | 223,9030177 | 15 | 3,239030177 | 4 |
| P09493 | TPM1 | Tropomyosin alpha-1 chain | 0,866395115 | 0,000991 | 0,00410988 | 82,31017913 | 452 | 1,823101791 | 40 |
| Q9ULW0 | TPX2 | Targeting protein for Xklp2 | 1,513577737 | 0,036719 | 0,0708178 | 185,5172155 | 12 | 2,855172155 | 2 |
| Q12931 | TRAP1 | Heat shock protein 75 kDa, mitochondrial | 0,121306803 | 0,000409 | 0,00196432 | 8,771968015 | 219 | 1,08771968 | 34 |
| P61421 | VA0D1 | V-type proton ATPase subunit d 1 | 0,269257056 | 0,001654 | 0,00627283 | 20,51870329 | 42 | 1,205187033 | 8 |
| P38606 | VATA | V-type proton ATPase catalytic subunit A | 0,082648525 | 0,020234 | 0,04565977 | 5,896031486 | 197 | 1,058960315 | 25 |
| O76024 | WFS1 | Wolframin | 0,423911702 | 0,03849 | 0,07322444 | 34,15601137 | 117 | 1,341560114 | 16 |

**Supplementary Data Table 2. List of relevant antioxidant proteins with altered expression in HGPS-iPSC-CMs (vs ISO-iPSC-CMs)**

| **Uniprot ID** | **Protein Symbol** | **Protein Description** | **Average Log2 Ratio** | ***p*-value** | **Adjusted *p*-value** |
| --- | --- | --- | --- | --- | --- |
| P04040 | CATA | Catalase | -2.1174278 | 1.78E-05 | 0.0001145 |
| P22352 | GPX3 | Glutathione peroxidase 3 | -0.0985667 | 0.032515 | 0.0551774 |
| Q96SL4 | GPX7 | Glutathione peroxidase 7 | -0.2358686 | 0.000224 | 0.00101521 |
| P04179 | SOD2 | Superoxide dismutase [Mn], mitochondrial | -0.0046289 | 0.0346497 | 0.057764 |

**Supplementary Data Table 3. List of total significantly altered proteins (p-value<0.05) in Physiologically aged mice**

| **Uniprot ID** | **Protein Symbol** | **Protein Description** | **Average Log2 Ratio** | ***p*-value** | **Adjusted *p*-value** |
| --- | --- | --- | --- | --- | --- |
| Q99L13 | 3HIDH | 3-hydroxyisobutyrate dehydrogenase, mitochondrial (HIBADH) (EC 1.1.1.31) | 0,316678181 | 0,003623 | 0,133505209 |
| P56379 | 68MP | 6.8 kDa mitochondrial proteolipid | -0,606380684 | 0,003548 | 0,131737888 |
| Q3THG9 | AASD1 | Alanyl-tRNA editing protein Aarsd1 (Alanyl-tRNA deacylase alaX) (AlaX) (AlaXp-II) (Alanyl-tRNA synthetase domain-containing protein 1) | -0,4455358 | 0,001323 | 0,069422675 |
| Q91V24 | ABCA7 | ATP-binding cassette sub-family A member 7 | 0,216569472 | 0,037304 | 0,415857046 |
| Q9DC29 | ABCB6 | ATP-binding cassette sub-family B member 6, mitochondrial | 0,304760329 | 0,00606 | 0,169247437 |
| Q8R2Y0 | ABHD6 | Monoacylglycerol lipase ABHD6 (EC 3.1.1.23) (2-arachidonoylglycerol hydrolase) (Abhydrolase domain-containing protein 6) | 0,312538783 | 0,005084 | 0,157323553 |
| Q8BUZ1 | ABRA | Actin-binding Rho-activating protein (Striated muscle activator of Rho-dependent signaling) (STARS) | 0,270172678 | 0,002226 | 0,099736537 |
| P45952 | ACADM | Medium-chain specific acyl-CoA dehydrogenase, mitochondrial (MCAD) (EC 1.3.8.7) | 0,195298752 | 0,040956 | 0,434498617 |
| Q5XG73 | ACBD5 | Acyl-CoA-binding domain-containing protein 5 | 0,198907606 | 0,04078 | 0,433575702 |
| Q9EPL9 | ACOX3 | Peroxisomal acyl-coenzyme A oxidase 3 (EC 1.3.3.6) (Branched-chain acyl-CoA oxidase) (BRCACox) (Pristanoyl-CoA oxidase) | 0,386480301 | 0,001473 | 0,074079015 |
| Q8VCW8 | ACSF2 | Acyl-CoA synthetase family member 2, mitochondrial (EC 6.2.1.-) | 0,456903954 | 0,000522 | 0,038163706 |
| Q91WC3 | ACSL6 | Long-chain-fatty-acid--CoA ligase 6 (EC 6.2.1.3) (Long-chain acyl-CoA synthetase 6) (LACS 6) | 0,735648521 | 1,94E-06 | 0,000668811 |
| Q8BGA8 | ACSM5 | Acyl-coenzyme A synthetase ACSM5, mitochondrial (EC 6.2.1.2) | 1,559860898 | 3,05E-08 | 3,69E-05 |
| Q80V03 | ADCK5 | Uncharacterized aarF domain-containing protein kinase 5 (EC 2.7.11.-) | 0,22201472 | 0,015515 | 0,265570551 |
| P00329 | ADH1 | Alcohol dehydrogenase 1 (EC 1.1.1.1) (ADH-A2) (Alcohol dehydrogenase A subunit) | 0,291842313 | 0,002935 | 0,119059186 |
| Q920A7 | AFG31 | AFG3-like protein 1 (EC 3.4.24.-) | -0,225125066 | 0,021907 | 0,312487475 |
| Q9ESW4 | AGK | Acylglycerol kinase, mitochondrial (EC 2.7.1.107) (EC 2.7.1.94) (Multiple substrate lipid kinase) (MuLK) (Multi-substrate lipid kinase) | -0,223706194 | 0,019429 | 0,299568128 |
| P31230 | AIMP1 | Aminoacyl tRNA synthase complex-interacting multifunctional protein 1 (Multisynthase complex auxiliary component p43) [Cleaved into: Endothelial monocyte-activating polypeptide 2 (EMAP-2) (Endothelial monocyte-activating polypeptide II) (EMAP-II) (Small inducible cytokine subfamily E member 1)] | -0,201766589 | 0,049903 | 0,470472203 |
| Q9JII6 | AK1A1 | Alcohol dehydrogenase [NADP(+)] (EC 1.1.1.2) (Aldehyde reductase) (Aldo-keto reductase family 1 member A1) | 0,339438874 | 0,037918 | 0,415979268 |
| Q9CZS1 | AL1B1 | Aldehyde dehydrogenase X, mitochondrial (EC 1.2.1.3) (Aldehyde dehydrogenase family 1 member B1) | -0,432667362 | 0,003908 | 0,13550848 |
| Q8R0Y6 | AL1L1 | Cytosolic 10-formyltetrahydrofolate dehydrogenase (10-FTHFDH) (FDH) (EC 1.5.1.6) (Aldehyde dehydrogenase family 1 member L1) | 0,266906256 | 0,035848 | 0,411990545 |
| Q8K009 | AL1L2 | Mitochondrial 10-formyltetrahydrofolate dehydrogenase (Mitochondrial 10-FTHFDH) (mtFDH) (EC 1.5.1.6) (Aldehyde dehydrogenase family 1 member L2) | -0,548015937 | 0,000216 | 0,021766647 |
| Q8CHT0 | AL4A1 | Delta-1-pyrroline-5-carboxylate dehydrogenase, mitochondrial (P5C dehydrogenase) (EC 1.2.1.88) (Aldehyde dehydrogenase family 4 member A1) (L-glutamate gamma-semialdehyde dehydrogenase) | 0,320010021 | 0,043236 | 0,438864843 |
| Q9JLJ2 | AL9A1 | 4-trimethylaminobutyraldehyde dehydrogenase (TMABADH) (EC 1.2.1.47) (Aldehyde dehydrogenase family 9 member A1) (EC 1.2.1.3) | 0,392450567 | 0,007459 | 0,189372626 |
| Q8BGT5 | ALAT2 | Alanine aminotransferase 2 (ALT2) (EC 2.6.1.2) (Glutamate pyruvate transaminase 2) (GPT 2) (Glutamic--alanine transaminase 2) (Glutamic--pyruvic transaminase 2) | -0,283187236 | 0,008446 | 0,197909434 |
| Q91Y97 | ALDOB | Fructose-bisphosphate aldolase B (EC 4.1.2.13) (Aldolase 2) (Liver-type aldolase) | 1,687548145 | 9,31E-07 | 0,000374694 |
| Q3TSG4 | ALKB5 | RNA demethylase ALKBH5 (EC 1.14.11.-) (Alkylated DNA repair protein alkB homolog 5) (Alpha-ketoglutarate-dependent dioxygenase alkB homolog 5) | 0,202197759 | 0,016824 | 0,27716535 |
| Q924C5 | ALPK3 | Alpha-protein kinase 3 (EC 2.7.11.-) (Myocytic induction/differentiation originator) (Midori) | 0,19737432 | 0,024131 | 0,331853082 |
| Q9R045 | ANGL2 | Angiopoietin-related protein 2 (Angiopoietin-like protein 2) | 1,052387332 | 1,36E-07 | 9,27E-05 |
| Q8C8R3 | ANK2 | Ankyrin-2 (ANK-2) (Ankyrin-B) (Brain ankyrin) | -0,301699565 | 0,003846 | 0,13550848 |
| Q8BH79 | ANO10 | Anoctamin-10 (Transmembrane protein 16K) | 0,246628194 | 0,02444 | 0,335153023 |
| Q99NH0 | ANR17 | Ankyrin repeat domain-containing protein 17 (Ankyrin repeat domain-containing protein FOE) (Gene trap ankyrin repeat protein) | -0,155362617 | 0,041621 | 0,437527 |
| Q64133 | AOFA | Amine oxidase [flavin-containing] A (EC 1.4.3.4) (Monoamine oxidase type A) (MAO-A) | 0,304849824 | 0,003239 | 0,128155431 |
| Q8BW75 | AOFB | Amine oxidase [flavin-containing] B (EC 1.4.3.4) (Monoamine oxidase type B) (MAO-B) | 0,32314598 | 0,011472 | 0,229977318 |
| P61967 | AP1S1 | AP-1 complex subunit sigma-1A (Adaptor protein complex AP-1 subunit sigma-1A) (Adaptor-related protein complex 1 subunit sigma-1A) (Clathrin assembly protein complex 1 sigma-1A small chain) (Clathrin coat assembly protein AP19) (Golgi adaptor HA1/AP1 adaptin sigma-1A subunit) (HA1 19 kDa subunit) (Sigma 1a subunit of AP-1 clathrin) (Sigma-adaptin 1A) (Sigma1A-adaptin) | -0,357845379 | 0,001269 | 0,068533932 |
| Q9WV76 | AP4B1 | AP-4 complex subunit beta-1 (AP-4 adaptor complex subunit beta) (Adaptor-related protein complex 4 subunit beta-1) (Beta subunit of AP-4) (Beta4-adaptin) | -0,238645809 | 0,036499 | 0,41357165 |
| Q80V94 | AP4E1 | AP-4 complex subunit epsilon-1 (AP-4 adaptor complex subunit epsilon) (Adaptor-related protein complex 4 subunit epsilon-1) (Epsilon subunit of AP-4) (Epsilon-adaptin) | -0,317966199 | 0,030575 | 0,380052223 |
| Q9DBR4 | APBB2 | Amyloid-beta A4 precursor protein-binding family B member 2 | -0,267081943 | 0,004614 | 0,149483602 |
| P28352 | APEX1 | DNA-(apurinic or apyrimidinic site) lyase (EC 3.1.-.-) (EC 4.2.99.18) (APEX nuclease) (APEN) (Apurinic-apyrimidinic endonuclease 1) (AP endonuclease 1) (REF-1) (Redox factor-1) [Cleaved into: DNA-(apurinic or apyrimidinic site) lyase, mitochondrial] | -0,229350527 | 0,029078 | 0,365516943 |
| P06728 | APOA4 | Apolipoprotein A-IV (Apo-AIV) (ApoA-IV) (Apolipoprotein A4) | 0,261307933 | 0,021146 | 0,310172316 |
| P08030 | APT | Adenine phosphoribosyltransferase (APRT) (EC 2.4.2.7) | 0,408975756 | 0,005512 | 0,166310605 |
| Q02013 | AQP1 | Aquaporin-1 (AQP-1) (Aquaporin-CHIP) (Delayed early response protein 2) (DER2) (Water channel protein for red blood cells and kidney proximal tubule) | -0,202435224 | 0,044887 | 0,445196515 |
| Q9EPJ9 | ARFG1 | ADP-ribosylation factor GTPase-activating protein 1 (ARF GAP 1) (ADP-ribosylation factor 1 GTPase-activating protein) (ARF1 GAP) (ARF1-directed GTPase-activating protein) | -0,239194722 | 0,018259 | 0,289918189 |
| Q9Z1K5 | ARI1 | E3 ubiquitin-protein ligase ARIH1 (EC 2.3.2.31) (Protein ariadne-1 homolog) (ARI-1) (RING-type E3 ubiquitin transferase ARIH1) (UbcH7-binding protein) (UbcM4-interacting protein 77) (Ubiquitin-conjugating enzyme E2-binding protein 1) | -0,331077042 | 0,037431 | 0,415979268 |
| A2BH40 | ARI1A | AT-rich interactive domain-containing protein 1A (ARID domain-containing protein 1A) (BRG1-associated factor 250) (BAF250) (BRG1-associated factor 250a) (BAF250A) (Osa homolog 1) (SWI-like protein) (SWI/SNF complex protein p270) (SWI/SNF-related, matrix-associated, actin-dependent regulator of chromatin subfamily F member 1) | 0,795058152 | 0,002008 | 0,094087926 |
| Q9D0L7 | ARM10 | Armadillo repeat-containing protein 10 | -0,179061745 | 0,03669 | 0,414206096 |
| Q9WV54 | ASAH1 | Acid ceramidase (AC) (ACDase) (Acid CDase) (EC 3.5.1.23) (Acylsphingosine deacylase) (N-acylsphingosine amidohydrolase) [Cleaved into: Acid ceramidase subunit alpha; Acid ceramidase subunit beta] | 0,396926334 | 0,015452 | 0,265494888 |
| Q8K0L0 | ASB2 | Ankyrin repeat and SOCS box protein 2 (ASB-2) | 0,254707061 | 0,027361 | 0,35802201 |
| Q64191 | ASPG | N(4)-(beta-N-acetylglucosaminyl)-L-asparaginase (EC 3.5.1.26) (Aspartylglucosaminidase) (AGA) (Glycosylasparaginase) (N4-(N-acetyl-beta-glucosaminyl)-L-asparagine amidase) [Cleaved into: Glycosylasparaginase alpha chain; Glycosylasparaginase beta chain] | 0,293123612 | 0,012486 | 0,235257931 |
| Q99MQ4 | ASPN | Asporin (Periodontal ligament-associated protein 1) (PLAP-1) | 0,242300493 | 0,019103 | 0,297447243 |
| Q925I1 | ATAD3 | ATPase family AAA domain-containing protein 3 (AAA-ATPase TOB3) | -0,18000366 | 0,037194 | 0,415590823 |
| O35143 | ATIF1 | ATPase inhibitor, mitochondrial (Inhibitor of F(1)F(o)-ATPase) (IF(1)) (IF1) | -0,40066985 | 0,006018 | 0,169247437 |
| Q8BH66 | ATLA1 | Atlastin-1 (EC 3.6.5.-) (Spastic paraplegia 3A homolog) | 0,414260919 | 0,000436 | 0,035078895 |
| Q91YH5 | ATLA3 | Atlastin-3 (EC 3.6.5.-) | 0,226691763 | 0,043658 | 0,43995108 |
| P56135 | ATPK | ATP synthase subunit f, mitochondrial | -0,371379057 | 0,020283 | 0,306686651 |
| O70305 | ATX2 | Ataxin-2 (Spinocerebellar ataxia type 2 protein homolog) | -0,215665887 | 0,032908 | 0,395146624 |
| P58158 | B3GA3 | Galactosylgalactosylxylosylprotein 3-beta-glucuronosyltransferase 3 (EC 2.4.1.135) (Beta-1,3-glucuronyltransferase 3) (Glucuronosyltransferase I) (GlcAT-I) (UDP-GlcUA:Gal beta-1,3-Gal-R glucuronyltransferase) (GlcUAT-I) | -0,214269428 | 0,047906 | 0,460644448 |
| O08734 | BAK | Bcl-2 homologous antagonist/killer (Apoptosis regulator BAK) | -0,450930293 | 0,000703 | 0,04547563 |
| Q9CWI3 | BCCIP | BRCA2 and CDKN1A-interacting protein | 0,225357566 | 0,027369 | 0,35802201 |
| Q80XN0 | BDH | D-beta-hydroxybutyrate dehydrogenase, mitochondrial (EC 1.1.1.30) (3-hydroxybutyrate dehydrogenase) (BDH) | 1,237141821 | 7,38E-08 | 5,94E-05 |
| P82198 | BGH3 | Transforming growth factor-beta-induced protein ig-h3 (Beta ig-h3) | 0,450622911 | 0,000313 | 0,029574232 |
| O55003 | BNIP3 | BCL2/adenovirus E1B 19 kDa protein-interacting protein 3 | 0,221702 | 0,021205 | 0,310172316 |
| Q9D6W8 | BORC6 | BLOC-1-related complex subunit 6 | 0,179079867 | 0,045773 | 0,447706624 |
| Q8R2Q8 | BST2 | Bone marrow stromal antigen 2 (BST-2) (HM1.24 antigen) (CD antigen CD317) | -0,414656146 | 0,002294 | 0,099736537 |
| Q8BJE2 | BTNL9 | Butyrophilin-like protein 9 | 0,4128485 | 0,000678 | 0,04485405 |
| O89103 | C1QR1 | Complement component C1q receptor (C1q/MBL/SPA receptor) (C1qR(p)) (C1qRp) (Cell surface antigen AA4) (Complement component 1 q subcomponent receptor 1) (Lymphocyte antigen 68) (Ly-68) (CD antigen CD93) | -0,245371327 | 0,014406 | 0,25471402 |
| Q8K1A6 | C2D1A | Coiled-coil and C2 domain-containing protein 1A (Five prime repressor element under dual repression-binding protein 1) (FRE under dual repression-binding protein 1) (Freud-1) | 0,187955097 | 0,046196 | 0,449574379 |
| P08607 | C4BPA | C4b-binding protein (C4bp) | 0,492534058 | 0,012943 | 0,238461417 |
| Q61112 | CAB45 | 45 kDa calcium-binding protein (Cab45) (Stromal cell-derived factor 4) (SDF-4) | 0,262362998 | 0,004838 | 0,153653996 |
| Q8BI72 | CARF | CDKN2A-interacting protein (Collaborator of ARF) | -0,34030328 | 0,000666 | 0,04485405 |
| P10605 | CATB | Cathepsin B (EC 3.4.22.1) (Cathepsin B1) [Cleaved into: Cathepsin B light chain; Cathepsin B heavy chain] | 0,444052821 | 0,003489 | 0,131584414 |
| P18242 | CATD | Cathepsin D (EC 3.4.23.5) | 0,421445442 | 0,000479 | 0,036604522 |
| Q9R013 | CATF | Cathepsin F (EC 3.4.22.41) | 0,97779818 | 3,75E-05 | 0,006039781 |
| P06797 | CATL1 | Cathepsin L1 (EC 3.4.22.15) (Cathepsin L) (Major excreted protein) (MEP) (p39 cysteine proteinase) [Cleaved into: Cathepsin L1 heavy chain; Cathepsin L1 light chain] | 0,319935501 | 0,04242 | 0,437527 |
| O70370 | CATS | Cathepsin S (EC 3.4.22.27) | 0,555943229 | 0,000813 | 0,049705084 |
| P49817 | CAV1 | Caveolin-1 | 0,192613212 | 0,036727 | 0,414206096 |
| Q9JJN5 | CBPN | Carboxypeptidase N catalytic chain (CPN) (EC 3.4.17.3) (Carboxypeptidase N polypeptide 1) (Carboxypeptidase N small subunit) | 0,292252863 | 0,036229 | 0,412690946 |
| Q8K354 | CBR3 | Carbonyl reductase [NADPH] 3 (EC 1.1.1.184) (NADPH-dependent carbonyl reductase 3) | 0,383316699 | 0,008323 | 0,196936869 |
| P83917 | CBX1 | Chromobox protein homolog 1 (Heterochromatin protein 1 homolog beta) (HP1 beta) (Heterochromatin protein p25) (M31) (Modifier 1 protein) | -0,185764982 | 0,028367 | 0,362241093 |
| Q61686 | CBX5 | Chromobox protein homolog 5 (Heterochromatin protein 1 homolog alpha) (HP1 alpha) | -0,414164338 | 0,015445 | 0,265494888 |
| A3KGF9 | CCD9B | Coiled-coil domain-containing protein 9B | 0,256257224 | 0,043804 | 0,440507248 |
| Q9WU84 | CCS | Copper chaperone for superoxide dismutase (Superoxide dismutase copper chaperone) | 0,272952491 | 0,010164 | 0,214516338 |
| Q08857 | CD36 | Platelet glycoprotein 4 (Glycoprotein IIIb) (GPIIIB) (PAS IV) (PAS-4) (Platelet glycoprotein IV) (GPIV) (CD antigen CD36) | 0,308286015 | 0,002244 | 0,099736537 |
| Q9QWK4 | CD5L | CD5 antigen-like (Apoptosis inhibitor expressed by macrophages) (mAIM) (Apoptosis inhibitory 6) (SP-alpha) | 0,728536813 | 2,51E-05 | 0,00433148 |
| P41731 | CD63 | CD63 antigen (CD antigen CD63) | 0,337929618 | 0,031881 | 0,38958903 |
| Q9Z0H4 | CELF2 | CUGBP Elav-like family member 2 (CELF-2) (Bruno-like protein 3) (CUG triplet repeat RNA-binding protein 2) (CUG-BP2) (CUG-BP- and ETR-3-like factor 2) (ELAV-type RNA-binding protein 3) (ETR-3) (mETR-3) (Neuroblastoma apoptosis-related RNA-binding protein) (mNapor) (RNA-binding protein BRUNOL-3) | -0,248084061 | 0,041319 | 0,437388354 |
| Q8C0D9 | CEP68 | Centrosomal protein of 68 kDa (Cep68) | 0,309891038 | 0,008545 | 0,198767737 |
| Q61147 | CERU | Ceruloplasmin (EC 1.16.3.1) (Ferroxidase) | 0,318217737 | 0,006734 | 0,181594765 |
| P04186 | CFAB | Complement factor B (EC 3.4.21.47) (C3/C5 convertase) [Cleaved into: Complement factor B Ba fragment; Complement factor B Bb fragment] | 0,538294915 | 0,023846 | 0,328869424 |
| P06909 | CFAH | Complement factor H (Protein beta-1-H) | 0,443562119 | 0,007128 | 0,184972448 |
| P63038 | CH60 | 60 kDa heat shock protein, mitochondrial (EC 3.6.4.9) (60 kDa chaperonin) (Chaperonin 60) (CPN60) (HSP-65) (Heat shock protein 60) (HSP-60) (Hsp60) (Mitochondrial matrix protein P1) | -0,210422253 | 0,010499 | 0,219018194 |
| Q9WUD1 | CHIP | STIP1 homology and U box-containing protein 1 (EC 2.3.2.27) (Carboxy terminus of Hsp70-interacting protein) (E3 ubiquitin-protein ligase CHIP) (RING-type E3 ubiquitin transferase CHIP) | 0,172166322 | 0,037703 | 0,415979268 |
| P60824 | CIRBP | Cold-inducible RNA-binding protein (A18 hnRNP) (Glycine-rich RNA-binding protein CIRP) | 0,325889975 | 0,00099 | 0,0567535 |
| Q8BMK4 | CKAP4 | Cytoskeleton-associated protein 4 (63-kDa cytoskeleton-linking membrane protein) (Climp-63) (p63) | -0,409102193 | 0,003296 | 0,128509517 |
| P49300 | CLC10 | C-type lectin domain family 10 member A (MMGL) (Macrophage asialoglycoprotein-binding protein 1) (M-ASGP-BP-1) (Macrophage galactose/N-acetylgalactosamine-specific lectin) | 0,523975542 | 0,001147 | 0,063634264 |
| Q8BWY2 | CLC1A | C-type lectin domain family 1 member A | 0,214710117 | 0,027734 | 0,358905168 |
| Q8BXK9 | CLIC5 | Chloride intracellular channel protein 5 | 0,277838622 | 0,042194 | 0,437527 |
| Q922J3 | CLIP1 | CAP-Gly domain-containing linker protein 1 (Cytoplasmic linker protein 170) (CLIP-170) (Restin) | 0,252027996 | 0,005994 | 0,169247437 |
| Q8CI96 | CLIP4 | CAP-Gly domain-containing linker protein 4 (Restin-like protein 2) | -0,713689966 | 0,000488 | 0,036604522 |
| Q1ERP8 | CLM9 | CMRF35-like molecule 9 (CLM-9) (CD300 antigen-like family member G) (Nepmucin) (CD antigen CD300g) | 0,268099339 | 0,009246 | 0,206612763 |
| Q9JHS4 | CLPX | ATP-dependent Clp protease ATP-binding subunit clpX-like, mitochondrial | -0,177976786 | 0,038458 | 0,42034693 |
| Q06890 | CLUS | Clusterin (Apolipoprotein J) (Apo-J) (Clustrin) (Sulfated glycoprotein 2) (SGP-2) [Cleaved into: Clusterin beta chain; Clusterin alpha chain] | 0,542946681 | 0,000431 | 0,035078895 |
| P21844 | CMA1 | Chymase (EC 3.4.21.39) (Alpha-chymase) (Mast cell chymase 1) (Mast cell protease 5) (mMCP-5) (Mast cell protease I) | 0,301515272 | 0,043277 | 0,438864843 |
| Q3U5Q7 | CMPK2 | UMP-CMP kinase 2, mitochondrial (EC 2.7.4.14) (Nucleoside-diphosphate kinase) (EC 2.7.4.6) (Thymidylate kinase LPS-inducible member) (TYKi) | -0,299072389 | 0,027537 | 0,358456285 |
| P02463 | CO4A1 | Collagen alpha-1(IV) chain [Cleaved into: Arresten] | 0,35096717 | 0,003323 | 0,128509517 |
| P08122 | CO4A2 | Collagen alpha-2(IV) chain [Cleaved into: Canstatin] | 0,263753533 | 0,008392 | 0,197608361 |
| P06684 | CO5 | Complement C5 (Hemolytic complement) [Cleaved into: Complement C5 beta chain; Complement C5 alpha chain; C5a anaphylatoxin; Complement C5 alpha' chain] | 0,711344201 | 0,004365 | 0,144302072 |
| Q04857 | CO6A1 | Collagen alpha-1(VI) chain | 0,708350839 | 3,46E-05 | 0,005762629 |
| Q02788 | CO6A2 | Collagen alpha-2(VI) chain | 0,793854814 | 3,03E-06 | 0,000914694 |
| Q8C6K9 | CO6A6 | Collagen alpha-6(VI) chain | 0,941403008 | 5,29E-06 | 0,001503267 |
| P06683 | CO9 | Complement component C9 | 0,419676474 | 0,049706 | 0,469528405 |
| Q80X19 | COEA1 | Collagen alpha-1(XIV) chain | -0,613791956 | 0,006066 | 0,169247437 |
| P39061 | COIA1 | Collagen alpha-1(XVIII) chain [Cleaved into: Endostatin; Non-collagenous domain 1 (NC1)] | 0,347215459 | 0,011625 | 0,229977318 |
| Q9JHH9 | COPZ2 | Coatomer subunit zeta-2 (Zeta-2-coat protein) (Zeta-2 COP) | -0,191633247 | 0,04938 | 0,469204211 |
| Q8BMS4 | COQ3 | Ubiquinone biosynthesis O-methyltransferase, mitochondrial (3-demethylubiquinol 3-O-methyltransferase) (EC 2.1.1.64) (Polyprenyldihydroxybenzoate methyltransferase) (EC 2.1.1.114) | -0,21781604 | 0,015943 | 0,268323331 |
| Q60936 | COQ8A | Atypical kinase COQ8A, mitochondrial (EC 2.7.-.-) (Chaperone activity of bc1 complex-like) (Chaperone-ABC1-like) (Coenzyme Q protein 8A) (aarF domain-containing protein kinase 3) | -0,294697625 | 0,016453 | 0,272922086 |
| P17665 | COX7C | Cytochrome c oxidase subunit 7C, mitochondrial (Cytochrome c oxidase polypeptide VIIc) | -0,367127561 | 0,004935 | 0,154696257 |
| Q9DBG1 | CP27A | Sterol 26-hydroxylase, mitochondrial (EC 1.14.15.15) (5-beta-cholestane-3-alpha,7-alpha,12-alpha-triol 27-hydroxylase) (Cytochrome P-450C27/25) (Cytochrome P450 27) (Sterol 27-hydroxylase) (Vitamin D(3) 25-hydroxylase) | 0,281905521 | 0,015176 | 0,264460609 |
| P12791 | CP2BA | Cytochrome P450 2B10 (EC 1.14.14.1) (CYPIIB10) (CYPIIB20) (Cytochrome P450 2B20) (Cytochrome P450 clone PF3/46) (Cytochrome P450-16-alpha) (P24) (Testosterone 16-alpha hydroxylase) | 2,091727797 | 1,20E-09 | 5,77E-06 |
| Q8CIM7 | CP2DQ | Cytochrome P450 2D26 (EC 1.14.14.1) (CYPIID26) | 0,39834949 | 0,000171 | 0,019711171 |
| Q64462 | CP4B1 | Cytochrome P450 4B1 (EC 1.14.14.1) (CYPIVB1) | 0,169705518 | 0,037766 | 0,415979268 |
| Q8BT60 | CPNE3 | Copine-3 (Copine III) | -0,212646743 | 0,045201 | 0,445196515 |
| Q9DCT8 | CRIP2 | Cysteine-rich protein 2 (CRP-2) (Heart LIM protein) | -0,411703605 | 0,009127 | 0,205504763 |
| P23927 | CRYAB | Alpha-crystallin B chain (Alpha(B)-crystallin) (P23) | 0,193195144 | 0,037041 | 0,414842867 |
| P50462 | CSRP3 | Cysteine and glycine-rich protein 3 (Cysteine-rich protein 3) (CRP3) (LIM domain protein, cardiac) (Muscle LIM protein) | 0,195640161 | 0,030628 | 0,380052223 |
| P97792 | CXAR | Coxsackievirus and adenovirus receptor homolog (CAR) (mCAR) | -0,33731034 | 0,005634 | 0,166310605 |
| P56395 | CYB5 | Cytochrome b5 | 0,493473252 | 0,000204 | 0,021549195 |
| Q8BPM0 | DAAM1 | Disheveled-associated activator of morphogenesis 1 | -0,207803616 | 0,026042 | 0,346910452 |
| Q8BHC4 | DCAKD | Dephospho-CoA kinase domain-containing protein | -0,240482443 | 0,016584 | 0,274149053 |
| Q9CWS0 | DDAH1 | N(G),N(G)-dimethylarginine dimethylaminohydrolase 1 (DDAH-1) (Dimethylarginine dimethylaminohydrolase 1) (EC 3.5.3.18) (DDAHI) (Dimethylargininase-1) | 0,231120544 | 0,035294 | 0,410296115 |
| O88533 | DDC | Aromatic-L-amino-acid decarboxylase (AADC) (EC 4.1.1.28) (DOPA decarboxylase) (DDC) | -0,375487667 | 0,003122 | 0,124526285 |
| Q62095 | DDX3Y | ATP-dependent RNA helicase DDX3Y (EC 3.6.4.13) (D1Pas1-related sequence 1) (DEAD box protein 3, Y-chromosomal) (DEAD-box RNA helicase DEAD2) (mDEAD2) | -0,348663459 | 0,011579 | 0,229977318 |
| Q7TNV0 | DEK | Protein DEK | -0,183267001 | 0,025233 | 0,341171474 |
| Q9WV69 | DEMA | Dematin (Dematin actin-binding protein) (Erythrocyte membrane protein band 4.9) | -0,192916586 | 0,037735 | 0,415979268 |
| Q9R0P5 | DEST | Destrin (Actin-depolymerizing factor) (ADF) (Sid 23) | 0,255840878 | 0,036336 | 0,412690946 |
| O88736 | DHB7 | 3-keto-steroid reductase (EC 1.1.1.270) (17-beta-hydroxysteroid dehydrogenase 7) (17-beta-HSD 7) (Estradiol 17-beta-dehydrogenase 7) (EC 1.1.1.62) | 0,328909032 | 0,02907 | 0,365516943 |
| Q64442 | DHSO | Sorbitol dehydrogenase (EC 1.1.1.14) (L-iditol 2-dehydrogenase) | 0,294497539 | 0,025334 | 0,341580355 |
| Q9QZD8 | DIC | Mitochondrial dicarboxylate carrier (Solute carrier family 25 member 10) | 0,550621862 | 0,00191 | 0,090400426 |
| P63037 | DNJA1 | DnaJ homolog subfamily A member 1 (DnaJ protein homolog 2) (Heat shock 40 kDa protein 4) (Heat shock protein J2) (HSJ-2) | -0,463442687 | 0,001567 | 0,076415726 |
| P56542 | DNS2A | Deoxyribonuclease-2-alpha (EC 3.1.22.1) (Acid DNase) (Deoxyribonuclease II alpha) (DNase II alpha) (Lysosomal DNase II) | 0,301385668 | 0,015456 | 0,265494888 |
| Q99LN9 | DOHH | Deoxyhypusine hydroxylase (DOHH) (EC 1.14.99.29) (Deoxyhypusine dioxygenase) (Deoxyhypusine monooxygenase) | 0,179696975 | 0,032772 | 0,395082396 |
| P31428 | DPEP1 | Dipeptidase 1 (EC 3.4.13.19) (Membrane-bound dipeptidase 1) (MBD-1) (Microsomal dipeptidase) (Renal dipeptidase) | 0,290618045 | 0,004117 | 0,139940674 |
| Q570Y9 | DPTOR | DEP domain-containing mTOR-interacting protein (DEP domain-containing protein 6) | 0,263060697 | 0,007798 | 0,194879948 |
| Q8CHR6 | DPYD | Dihydropyrimidine dehydrogenase [NADP(+)] (DHPDHase) (DPD) (EC 1.3.1.2) (Dihydrothymine dehydrogenase) (Dihydrouracil dehydrogenase) | 0,324429783 | 0,00064 | 0,044140705 |
| Q62188 | DPYL3 | Dihydropyrimidinase-related protein 3 (DRP-3) (Unc-33-like phosphoprotein 1) (ULIP-1) | -0,465468834 | 8,94E-05 | 0,012326075 |
| Q8CHS7 | DRS7C | Dehydrogenase/reductase SDR family member 7C (EC 1.1.-.-) (Short-chain dehydrogenase/reductase family 32C member 2) | 0,31592131 | 0,009153 | 0,205504763 |
| Q8BHA3 | DTD2 | D-aminoacyl-tRNA deacylase 2 (EC 3.1.1.96) (Animalia-specifc tRNA deacylase) (ATD) (D-tyrosyl-tRNA deacylase 2) (L-alanyl-tRNA deacylase) (EC 3.1.1.-) | 0,391802542 | 0,007124 | 0,184972448 |
| Q148W8 | DUS27 | Inactive dual specificity phosphatase 27 | -0,257180143 | 0,012111 | 0,23226961 |
| Q8VDW0 | DX39A | ATP-dependent RNA helicase DDX39A (EC 3.6.4.13) (DEAD box protein 39) | 0,389662705 | 0,045004 | 0,445196515 |
| Q9ESD7 | DYSF | Dysferlin (Dystrophy-associated fer-1-like protein) (Fer-1-like protein 1) | -0,206160837 | 0,028142 | 0,361546694 |
| Q61508 | ECM1 | Extracellular matrix protein 1 (Secretory component p85) | 0,399368225 | 0,001178 | 0,064608076 |
| P10126 | EF1A1 | Elongation factor 1-alpha 1 (EF-1-alpha-1) (Elongation factor Tu) (EF-Tu) (Eukaryotic elongation factor 1 A-1) (eEF1A-1) | -0,385676618 | 0,034824 | 0,409166927 |
| P57776 | EF1D | Elongation factor 1-delta (EF-1-delta) | -0,45722027 | 0,000711 | 0,04547563 |
| Q9D4J1 | EFHD1 | EF-hand domain-containing protein D1 (EF-hand domain-containing protein 1) (Mitocalcin) (Swiprosin-2) | 1,152571189 | 7,77E-06 | 0,001804631 |
| O35393 | EFNB3 | Ephrin-B3 | 0,346763324 | 0,007076 | 0,184972448 |
| Q61070 | EI24 | Etoposide-induced protein 2.4 (p53-induced gene 8 protein) | -0,246341046 | 0,017559 | 0,283474302 |
| Q8C522 | ENDD1 | Endonuclease domain-containing 1 protein (EC 3.1.30.-) | 0,205114539 | 0,027956 | 0,360816734 |
| P17183 | ENOG | Gamma-enolase (EC 4.2.1.11) (2-phospho-D-glycerate hydro-lyase) (Enolase 2) (Neural enolase) (Neuron-specific enolase) (NSE) | 0,599347079 | 0,016042 | 0,268323331 |
| Q8BTJ4 | ENPP4 | Bis(5'-adenosyl)-triphosphatase enpp4 (EC 3.6.1.29) (AP3A hydrolase) (AP3Aase) (Ectonucleotide pyrophosphatase/phosphodiesterase family member 4) (E-NPP 4) (NPP-4) | 0,277573787 | 0,00619 | 0,171716228 |
| O55026 | ENTP2 | Ectonucleoside triphosphate diphosphohydrolase 2 (NTPDase 2) (EC 3.6.1.-) (CD39 antigen-like 1) (Ecto-ATP diphosphohydrolase 2) (Ecto-ATPDase 2) (Ecto-ATPase 2) | 0,352965449 | 0,009478 | 0,208898509 |
| Q8VEH5 | EPMIP | EPM2A-interacting protein 1 (Laforin-interacting protein) | 0,272442569 | 0,034036 | 0,403042317 |
| Q8R1L4 | ERD23 | ER lumen protein-retaining receptor 3 (KDEL endoplasmic reticulum protein retention receptor 3) (KDEL receptor 3) | -0,249164964 | 0,012326 | 0,234915904 |
| Q8BK48 | EST2E | Pyrethroid hydrolase Ces2e (EC 3.1.1.88) (carboxylesterase 2E) | 0,55790295 | 0,005737 | 0,166310605 |
| Q5XG69 | F169A | Soluble lamin-associated protein of 75 kDa (SLAP75) (Protein FAM169A) | 0,91497521 | 7,64E-06 | 0,001804631 |
| Q3U2I3 | F16A2 | FTS and Hook-interacting protein (FHIP) (Protein FAM160A2) | 0,301196897 | 0,042925 | 0,438864843 |
| Q501J2 | F173A | Protein N-lysine methyltransferase FAM173A (EC 2.1.1.-) | 0,21034226 | 0,0204 | 0,306686651 |
| Q9CYH2 | F213A | Redox-regulatory protein FAM213A (Peroxiredoxin-like 2 activated in M-CSF stimulated monocytes) (Protein PAMM) | 0,426393217 | 0,000535 | 0,038534665 |
| P70266 | F261 | 6-phosphofructo-2-kinase/fructose-2,6-bisphosphatase 1 (6PF-2-K/Fru-2,6-P2ase 1) (PFK/FBPase 1) (6PF-2-K/Fru-2,6-P2ase liver isozyme) [Includes: 6-phosphofructo-2-kinase (EC 2.7.1.105); Fructose-2,6-bisphosphatase (EC 3.1.3.46)] | 0,644223438 | 0,000493 | 0,036604522 |
| Q80YC5 | FA12 | Coagulation factor XII (EC 3.4.21.38) (Hageman factor) (HAF) [Cleaved into: Coagulation factor XIIa heavy chain; Coagulation factor XIIa light chain] | 0,24431253 | 0,043908 | 0,440627624 |
| P35505 | FAAA | Fumarylacetoacetase (FAA) (EC 3.7.1.2) (Beta-diketonase) (Fumarylacetoacetate hydrolase) | 0,59683981 | 0,001278 | 0,068533932 |
| Q3TC72 | FAHD2 | Fumarylacetoacetate hydrolase domain-containing protein 2A (EC 3.-.-.-) | 0,3251901 | 0,007261 | 0,186862261 |
| Q8BPB5 | FBLN3 | EGF-containing fibulin-like extracellular matrix protein 1 (Fibulin-3) (FIBL-3) | -0,360385864 | 0,013391 | 0,243565124 |
| Q9WVH9 | FBLN5 | Fibulin-5 (FIBL-5) (Developmental arteries and neural crest EGF-like protein) (Dance) | -0,56456816 | 0,004471 | 0,146801182 |
| Q9CPU7 | FBX32 | F-box only protein 32 (Atrogin-1) (Muscle atrophy F-box protein) (MAFbx) | -0,480737227 | 0,002483 | 0,107018139 |
| P23591 | FCL | GDP-L-fucose synthase (EC 1.1.1.271) (GDP-4-keto-6-deoxy-D-mannose-3,5-epimerase-4-reductase) (Protein FX) (Red cell NADP(H)-binding protein) (Transplantation antigen P35B) (Tum-P35B antigen) | 0,389962589 | 0,013596 | 0,245788922 |
| Q80UZ0 | FGD5 | FYVE, RhoGEF and PH domain-containing protein 5 | 0,30098153 | 0,009964 | 0,212776202 |
| O70433 | FHL2 | Four and a half LIM domains protein 2 (FHL-2) (Skeletal muscle LIM-protein 3) (SLIM-3) | -0,587857859 | 0,023406 | 0,324658096 |
| Q9R059 | FHL3 | Four and a half LIM domains protein 3 (FHL-3) (Skeletal muscle LIM-protein 2) (SLIM-2) | -0,242438593 | 0,006959 | 0,184557321 |
| Q64378 | FKBP5 | Peptidyl-prolyl cis-trans isomerase FKBP5 (PPIase FKBP5) (EC 5.2.1.8) (51 kDa FK506-binding protein) (51 kDa FKBP) (FKBP-51) (FK506-binding protein 5) (FKBP-5) (Rotamase) | 0,454663094 | 0,017554 | 0,283474302 |
| Q8K2I3 | FMO2 | Dimethylaniline monooxygenase [N-oxide-forming] 2 (EC 1.14.13.8) (Dimethylaniline oxidase 2) (Pulmonary flavin-containing monooxygenase 2) (FMO 2) | 0,4537906 | 0,002031 | 0,094283395 |
| P97872 | FMO5 | Dimethylaniline monooxygenase [N-oxide-forming] 5 (EC 1.14.13.8) (Dimethylaniline oxidase 5) (Hepatic flavin-containing monooxygenase 5) (FMO 5) | 0,314818366 | 0,00828 | 0,196936869 |
| P09528 | FRIH | Ferritin heavy chain (Ferritin H subunit) (EC 1.16.3.1) [Cleaved into: Ferritin heavy chain, N-terminally processed] | 0,444181597 | 0,001412 | 0,071739638 |
| P29391 | FRIL1 | Ferritin light chain 1 (Ferritin L subunit 1) | 0,637484358 | 7,51E-06 | 0,001804631 |
| Q8BGW1 | FTO | Alpha-ketoglutarate-dependent dioxygenase FTO (EC 1.14.11.-) (Fat mass and obesity-associated protein) (Protein fatso) | -0,235809857 | 0,031008 | 0,383771212 |
| Q00612 | G6PD1 | Glucose-6-phosphate 1-dehydrogenase X (G6PD) (EC 1.1.1.49) | 0,297522295 | 0,015375 | 0,265494888 |
| Q8CFX1 | G6PE | GDH/6PGL endoplasmic bifunctional protein [Includes: Glucose 1-dehydrogenase (EC 1.1.1.47) (Glucose-6-phosphate dehydrogenase) (EC 1.1.1.363); 6-phosphogluconolactonase (6PGL) (EC 3.1.1.31)] | 0,52405402 | 0,001533 | 0,076277639 |
| P61922 | GABT | 4-aminobutyrate aminotransferase, mitochondrial (EC 2.6.1.19) ((S)-3-amino-2-methylpropionate transaminase) (EC 2.6.1.22) (GABA aminotransferase) (GABA-AT) (Gamma-amino-N-butyrate transaminase) (GABA transaminase) (GABA-T) (L-AIBAT) | 1,625964461 | 0,000167 | 0,019711171 |
| Q8BVW0 | GANC | Neutral alpha-glucosidase C (EC 3.2.1.20) | 0,25629938 | 0,012126 | 0,23226961 |
| Q9Z0E6 | GBP2 | Guanylate-binding protein 2 (EC 3.6.5.-) (GTP-binding protein 2) (GBP-2) (mGBP-2) (mGBP2) (Guanine nucleotide-binding protein 2) (Interferon-induced guanylate-binding protein 2) | -0,32398126 | 0,007971 | 0,196317279 |
| Q61107 | GBP4 | Guanylate-binding protein 4 (EC 3.6.5.-) (GTP-binding protein 3) (GBP-3) (GTP-binding protein 4) (GBP-4) (Guanine nucleotide-binding protein 4) (Guanylate-binding protein 3) | -0,417116952 | 0,003539 | 0,131737888 |
| P01864 | GCAB | Ig gamma-2A chain C region secreted form (B allele) | 1,007981961 | 0,03248 | 0,393920662 |
| Q07235 | GDN | Glia-derived nexin (GDN) (Peptidase inhibitor 7) (PI-7) (Protease nexin 1) (PN-1) (Protease nexin I) (Serine protease-inhibitor 4) (Serpin E2) | 0,693664147 | 0,025206 | 0,341171474 |
| Q9Z2Z9 | GFPT2 | Glutamine--fructose-6-phosphate aminotransferase [isomerizing] 2 (EC 2.6.1.16) (D-fructose-6-phosphate amidotransferase 2) (Glutamine:fructose-6-phosphate amidotransferase 2) (GFAT 2) (GFAT2) (Hexosephosphate aminotransferase 2) | 0,389060902 | 0,033116 | 0,396657108 |
| Q99JY3 | GIMA4 | GTPase IMAP family member 4 (Immunity-associated nucleotide 1 protein) (IAN-1) (Immunity-associated protein 4) | -0,243458824 | 0,009831 | 0,211856445 |
| P15105 | GLNA | Glutamine synthetase (GS) (EC 6.3.1.2) (Glutamate decarboxylase) (EC 4.1.1.15) (Glutamate--ammonia ligase) | 0,504493492 | 0,001796 | 0,086707447 |
| D3Z7P3 | GLSK | Glutaminase kidney isoform, mitochondrial (GLS) (EC 3.5.1.2) | -0,359654744 | 0,018701 | 0,295005233 |
| Q9JL62 | GLTP | Glycolipid transfer protein (GLTP) | 0,363608393 | 0,000673 | 0,04485405 |
| P50431 | GLYC | Serine hydroxymethyltransferase, cytosolic (SHMT) (EC 2.1.2.1) (Glycine hydroxymethyltransferase) (Serine methylase) | 0,51485778 | 0,003955 | 0,13550848 |
| Q8K0C9 | GMDS | GDP-mannose 4,6 dehydratase (EC 4.2.1.47) (GDP-D-mannose dehydratase) (GMD) | 0,264975294 | 0,042609 | 0,438534548 |
| Q9CQI3 | GMFB | Glia maturation factor beta (GMF-beta) | 0,24184175 | 0,007832 | 0,194879948 |
| Q6R0H7 | GNAS1 | Guanine nucleotide-binding protein G(s) subunit alpha isoforms XLas (Adenylate cyclase-stimulating G alpha protein) (Extra large alphas protein) (XLalphas) | 0,231099867 | 0,020501 | 0,306686651 |
| Q8BFR4 | GNS | N-acetylglucosamine-6-sulfatase (EC 3.1.6.14) (Glucosamine-6-sulfatase) (G6S) | 0,189084101 | 0,033716 | 0,400854464 |
| Q8BH60 | GOPC | Golgi-associated PDZ and coiled-coil motif-containing protein (PDZ protein interacting specifically with TC10) (PIST) | -0,321062774 | 0,004605 | 0,149483602 |
| Q99JX3 | GORS2 | Golgi reassembly-stacking protein 2 (GRS2) (Golgi reassembly-stacking protein of 55 kDa) (GRASP55) | -0,206113715 | 0,010558 | 0,219018194 |
| Q6IR34 | GPSM1 | G-protein-signaling modulator 1 (Activator of G-protein signaling 3) | -0,306869106 | 0,013814 | 0,247108523 |
| P46412 | GPX3 | Glutathione peroxidase 3 (GPx-3) (GSHPx-3) (EC 1.11.1.9) (Plasma glutathione peroxidase) (GPx-P) (GSHPx-P) | 0,241163784 | 0,047027 | 0,456040272 |
| O88396 | GRPE2 | GrpE protein homolog 2, mitochondrial (Mt-GrpE#2) | 0,195320239 | 0,020863 | 0,30796633 |
| P13745 | GSTA1 | Glutathione S-transferase A1 (EC 2.5.1.18) (GST class-alpha member 1) (Glutathione S-transferase Ya) (Glutathione S-transferase Ya1) [Cleaved into: Glutathione S-transferase A1, N-terminally processed] | 1,443744494 | 9,71E-09 | 2,34E-05 |
| P15626 | GSTM2 | Glutathione S-transferase Mu 2 (EC 2.5.1.18) (GST 5-5) (GST class-mu 2) (Glutathione S-transferase pmGT2) | 0,487139518 | 0,022619 | 0,317733016 |
| Q64471 | GSTT1 | Glutathione S-transferase theta-1 (EC 2.5.1.18) (GST class-theta-1) | 0,244977812 | 0,015994 | 0,268323331 |
| Q61133 | GSTT2 | Glutathione S-transferase theta-2 (EC 2.5.1.18) (GST class-theta-2) | 0,32370149 | 0,011149 | 0,22611708 |
| Q99L20 | GSTT3 | Glutathione S-transferase theta-3 (EC 2.5.1.18) | 0,391747658 | 0,000401 | 0,033976051 |
| P43275 | H11 | Histone H1.1 (H1 VAR.3) (Histone H1a) (H1a) | -1,010592362 | 1,97E-05 | 0,003662492 |
| P43277 | H13 | Histone H1.3 (H1 VAR.4) (H1d) | -0,404813972 | 0,019686 | 0,301666323 |
| P43276 | H15 | Histone H1.5 (H1 VAR.5) (H1b) | -1,530684171 | 4,24E-07 | 0,000204628 |
| Q9QY80 | HACD1 | Very-long-chain (3R)-3-hydroxyacyl-CoA dehydratase 1 (EC 4.2.1.134) (3-hydroxyacyl-CoA dehydratase 1) (HACD1) (Protein-tyrosine phosphatase-like member A) | -0,459775931 | 0,006933 | 0,184557321 |
| Q9QXE0 | HACL1 | 2-hydroxyacyl-CoA lyase 1 (EC 4.1.-.-) (2-hydroxyphytanoyl-CoA lyase) (2-HPCL) (Phytanoyl-CoA 2-hydroxylase 2) | 0,257426787 | 0,013822 | 0,247108523 |
| Q6NZM9 | HDAC4 | Histone deacetylase 4 (HD4) (EC 3.5.1.98) | -0,205271299 | 0,026088 | 0,346910452 |
| Q9D1N2 | HDBP1 | Glycosylphosphatidylinositol-anchored high density lipoprotein-binding protein 1 (GPI-HBP1) (GPI-anchored HDL-binding protein 1) (High density lipoprotein-binding protein 1) | 0,333236101 | 0,007285 | 0,186862261 |
| Q9R257 | HEBP1 | Heme-binding protein 1 (p22HBP) | 0,845835706 | 1,06E-05 | 0,002094406 |
| Q6NVF4 | HELB | DNA helicase B (EC 3.6.4.12) | 0,380642934 | 0,019487 | 0,299568128 |
| Q9JJK5 | HERP1 | Homocysteine-responsive endoplasmic reticulum-resident ubiquitin-like domain member 1 protein | -0,409953616 | 0,0028 | 0,115515908 |
| P29416 | HEXA | Beta-hexosaminidase subunit alpha (EC 3.2.1.52) (Beta-N-acetylhexosaminidase subunit alpha) (Hexosaminidase subunit A) (N-acetyl-beta-glucosaminidase subunit alpha) | 0,246433953 | 0,0441 | 0,440648118 |
| Q9D1G3 | HHATL | Protein-cysteine N-palmitoyltransferase HHAT-like protein (Glycerol uptake/transporter homolog) (Hedgehog acyltransferase-like protein) | -0,273702374 | 0,013422 | 0,243565124 |
| P70349 | HINT1 | Histidine triad nucleotide-binding protein 1 (EC 3.-.-.-) (Adenosine 5'-monophosphoramidase) (Protein kinase C inhibitor 1) (Protein kinase C-interacting protein 1) (PKCI-1) | 0,334022723 | 0,002657 | 0,110545052 |
| P54869 | HMCS2 | Hydroxymethylglutaryl-CoA synthase, mitochondrial (HMG-CoA synthase) (EC 2.3.3.10) (3-hydroxy-3-methylglutaryl coenzyme A synthase) | 0,636408068 | 0,028177 | 0,361546694 |
| O54879 | HMGB3 | High mobility group protein B3 (High mobility group protein 2a) (HMG-2a) (High mobility group protein 4) (HMG-4) | -0,225512471 | 0,010177 | 0,214516338 |
| Q8VDM6 | HNRL1 | Heterogeneous nuclear ribonucleoprotein U-like protein 1 | -0,170494599 | 0,034839 | 0,409166927 |
| Q9D0E1 | HNRPM | Heterogeneous nuclear ribonucleoprotein M (hnRNP M) | -0,179526096 | 0,042994 | 0,438864843 |
| Q8R0N6 | HOT | Hydroxyacid-oxoacid transhydrogenase, mitochondrial (HOT) (EC 1.1.99.24) (Alcohol dehydrogenase iron-containing protein 1) (ADHFe1) | -0,235298255 | 0,038723 | 0,420979619 |
| Q8K248 | HPDL | 4-hydroxyphenylpyruvate dioxygenase-like protein (EC 1.13.-.-) (Glyoxalase domain-containing protein 1) | -0,33109134 | 0,0495 | 0,46942088 |
| Q61646 | HPT | Haptoglobin (Zonulin) [Cleaved into: Haptoglobin alpha chain; Haptoglobin beta chain] | 3,250534274 | 2,57E-06 | 0,000825595 |
| Q61699 | HS105 | Heat shock protein 105 kDa (42 degrees C-HSP) (Heat shock 110 kDa protein) (Heat shock-related 100 kDa protein E7I) (HSP-E7I) | -0,23786199 | 0,014923 | 0,260996195 |
| Q9CZJ2 | HS12B | Heat shock 70 kDa protein 12B | 0,171016934 | 0,037037 | 0,414842867 |
| P11499 | HS90B | Heat shock protein HSP 90-beta (Heat shock 84 kDa) (HSP 84) (HSP84) (Tumor-specific transplantation 84 kDa antigen) (TSTA) | -0,23574876 | 0,016445 | 0,272922086 |
| P35385 | HSPB7 | Heat shock protein beta-7 (HspB7) (Cardiovascular heat shock protein) (cvHsp) (Heat shock protein 25 kDa 2) (Protein p19/6.8) | 0,261399423 | 0,012113 | 0,23226961 |
| Q9Z2G9 | HTAI2 | Oxidoreductase HTATIP2 (EC 1.1.1.-) | 0,337322713 | 0,000545 | 0,038712216 |
| Q9R118 | HTRA1 | Serine protease HTRA1 (EC 3.4.21.-) (High-temperature requirement A serine peptidase 1) (Serine protease 11) | 0,58478275 | 0,003871 | 0,13550848 |
| P06330 | HVM51 | Ig heavy chain V region AC38 205.12 | 1,096734905 | 0,040744 | 0,433575702 |
| Q9D379 | HYEP | Epoxide hydrolase 1 (EC 3.3.2.9) (Epoxide hydratase) (Microsomal epoxide hydrolase) (mEH) | 0,454816788 | 0,001314 | 0,069422675 |
| Q8K3X4 | I2BPL | Interferon regulatory factor 2-binding protein-like (Enhanced at puberty protein 1) | -0,465752358 | 0,000163 | 0,019711171 |
| P13597 | ICAM1 | Intercellular adhesion molecule 1 (ICAM-1) (MALA-2) (MyD10) (CD antigen CD54) | 0,429103437 | 0,001011 | 0,0567535 |
| P70404 | IDHG1 | Isocitrate dehydrogenase [NAD] subunit gamma 1, mitochondrial (Isocitric dehydrogenase subunit gamma) (NAD(+)-specific ICDH subunit gamma) | 0,35336662 | 0,003444 | 0,130887168 |
| P54071 | IDHP | Isocitrate dehydrogenase [NADP], mitochondrial (IDH) (EC 1.1.1.42) (ICD-M) (IDP) (NADP(+)-specific ICDH) (Oxalosuccinate decarboxylase) | 0,344171538 | 0,005798 | 0,16660284 |
| Q91YJ5 | IF2M | Translation initiation factor IF-2, mitochondrial (IF-2(Mt)) (IF-2Mt) (IF2(mt)) | -0,258599572 | 0,009718 | 0,211297351 |
| Q64345 | IFIT3 | Interferon-induced protein with tetratricopeptide repeats 3 (IFIT-3) (Glucocorticoid-attenuated response gene 49 protein) (GARG-49) (P49) (IRG2) | -0,411450991 | 0,005534 | 0,166310605 |
| P01867 | IGG2B | Ig gamma-2B chain C region | 1,182289197 | 0,04234 | 0,437527 |
| P01872 | IGHM | Immunoglobulin heavy constant mu [Cleaved into: Mu' chain (55 kDa mu' chain)] | 1,913265958 | 2,26E-05 | 0,00404754 |
| P01837 | IGKC | Immunoglobulin kappa constant (Ig kappa chain C region MOPC 21) | 1,046024406 | 0,002282 | 0,099736537 |
| Q9Z1X4 | ILF3 | Interleukin enhancer-binding factor 3 | -0,30257862 | 0,009079 | 0,205504763 |
| P40936 | INMT | Indolethylamine N-methyltransferase (Indolamine N-methyltransferase) (EC 2.1.1.49) (EC 2.1.1.96) (Aromatic alkylamine N-methyltransferase) (Amine N-methyltransferase) (Arylamine N-methyltransferase) (Thioether S-methyltransferase) (TEMT) | 1,294460512 | 0,000373 | 0,033115228 |
| Q9EPW0 | INP4A | Type I inositol 3,4-bisphosphate 4-phosphatase (EC 3.1.3.66) (Inositol polyphosphate 4-phosphatase type I) | 0,376273565 | 6,32E-05 | 0,00924269 |
| O70139 | IPKG | cAMP-dependent protein kinase inhibitor gamma (PKI-gamma) | 0,607477717 | 0,006569 | 0,178150577 |
| Q62406 | IRAK1 | Interleukin-1 receptor-associated kinase 1 (IRAK) (IRAK-1) (EC 2.7.11.1) (Pelle-like protein kinase) (mPLK) | -0,256804033 | 0,009039 | 0,205504763 |
| P85094 | ISC2A | Isochorismatase domain-containing protein 2A | 0,201029131 | 0,03181 | 0,38958903 |
| Q64339 | ISG15 | Ubiquitin-like protein ISG15 (Interferon-induced 15 kDa protein) (Interferon-induced 17 kDa protein) (IP17) (Ubiquitin cross-reactive protein) | -0,549140921 | 0,000639 | 0,044140705 |
| P11835 | ITB2 | Integrin beta-2 (Cell surface adhesion glycoproteins LFA-1/CR3/p150,95 subunit beta) (Complement receptor C3 subunit beta) (CD antigen CD18) | 0,470816484 | 0,028631 | 0,363349811 |
| Q61703 | ITIH2 | Inter-alpha-trypsin inhibitor heavy chain H2 (ITI heavy chain H2) (ITI-HC2) (Inter-alpha-inhibitor heavy chain 2) | 0,424284863 | 0,011579 | 0,229977318 |
| Q61704 | ITIH3 | Inter-alpha-trypsin inhibitor heavy chain H3 (ITI heavy chain H3) (ITI-HC3) (Inter-alpha-inhibitor heavy chain 3) | 0,524003999 | 0,000319 | 0,029574232 |
| A6X935 | ITIH4 | Inter alpha-trypsin inhibitor, heavy chain 4 (ITI heavy chain H4) (ITI-HC4) (Inter-alpha-inhibitor heavy chain 4) | 0,645096281 | 0,000179 | 0,020083533 |
| O89051 | ITM2B | Integral membrane protein 2B (Immature BRI2) (imBRI2) (Protein E25B) (Transmembrane protein BRI) (Bri) [Cleaved into: BRI2, membrane form (Mature BRI2) (mBRI2); BRI2 intracellular domain (BRI2 ICD); BRI2C, soluble form; Bri23 peptide (Bri2-23) (ABri23) (C-terminal peptide) (P23 peptide)] | 0,332984063 | 0,03536 | 0,410296115 |
| P70227 | ITPR3 | Inositol 1,4,5-trisphosphate receptor type 3 (IP3 receptor isoform 3) (IP3R 3) (InsP3R3) (Type 3 inositol 1,4,5-trisphosphate receptor) (Type 3 InsP3 receptor) | 0,238005789 | 0,030145 | 0,377949026 |
| Q9WTP7 | KAD3 | GTP:AMP phosphotransferase AK3, mitochondrial (EC 2.7.4.10) (Adenylate kinase 3) (AK 3) (Adenylate kinase 3 alpha-like 1) | -0,237600857 | 0,023019 | 0,321133804 |
| Q9WUR9 | KAD4 | Adenylate kinase 4, mitochondrial (AK 4) (EC 2.7.4.10) (EC 2.7.4.6) (Adenylate kinase 3-like) (Adenylate kinase isoenzyme 4) (GTP:AMP phosphotransferase AK4) | -0,256206371 | 0,012664 | 0,235257931 |
| Q71RI9 | KAT3 | Kynurenine--oxoglutarate transaminase 3 (EC 2.6.1.7) (Cysteine-S-conjugate beta-lyase 2) (EC 4.4.1.13) (Kynurenine aminotransferase 3) (Kynurenine aminotransferase III) (KATIII) (Kynurenine--glyoxylate transaminase) (EC 2.6.1.63) (Kynurenine--oxoglutarate transaminase III) | 0,370785711 | 0,007493 | 0,189372626 |
| O88986 | KBL | 2-amino-3-ketobutyrate coenzyme A ligase, mitochondrial (AKB ligase) (EC 2.3.1.29) (Aminoacetone synthase) (Glycine acetyltransferase) | 0,319359994 | 0,022643 | 0,317733016 |
| Q91YS8 | KCC1A | Calcium/calmodulin-dependent protein kinase type 1 (EC 2.7.11.17) (CaM kinase I) (CaM-KI) (CaM kinase I alpha) (CaMKI-alpha) | -0,407339242 | 0,010663 | 0,219018194 |
| Q923T9 | KCC2G | Calcium/calmodulin-dependent protein kinase type II subunit gamma (CaM kinase II subunit gamma) (CaMK-II subunit gamma) (EC 2.7.11.17) | -0,23895278 | 0,008068 | 0,196676452 |
| Q6WVG3 | KCD12 | BTB/POZ domain-containing protein KCTD12 (Pfetin) (Predominantly fetal expressed T1 domain) | -0,220742826 | 0,03849 | 0,42034693 |
| O88448 | KLC2 | Kinesin light chain 2 (KLC 2) | 0,3764214 | 0,00603 | 0,169247437 |
| Q02111 | KPCT | Protein kinase C theta type (EC 2.7.11.13) (nPKC-theta) | 1,318398127 | 2,35E-07 | 0,000126026 |
| Q8C050 | KS6A5 | Ribosomal protein S6 kinase alpha-5 (S6K-alpha-5) (EC 2.7.11.1) (90 kDa ribosomal protein S6 kinase 5) (Nuclear mitogen- and stress-activated protein kinase 1) (RSK-like protein kinase) (RLSK) | -0,297038911 | 0,003801 | 0,13550848 |
| P01643 | KV5AA | Ig kappa chain V-V region MOPC 173 | 1,109643848 | 0,012838 | 0,23742082 |
| Q61001 | LAMA5 | Laminin subunit alpha-5 (Laminin-10 subunit alpha) (Laminin-11 subunit alpha) (Laminin-15 subunit alpha) | 0,219159377 | 0,044059 | 0,440648118 |
| Q61292 | LAMB2 | Laminin subunit beta-2 (Laminin-11 subunit beta) (Laminin-14 subunit beta) (Laminin-15 subunit beta) (Laminin-3 subunit beta) (Laminin-4 subunit beta) (Laminin-7 subunit beta) (Laminin-9 subunit beta) (S-laminin subunit beta) (S-LAM beta) | 0,192806774 | 0,042808 | 0,438864843 |
| Q61029 | LAP2B | Lamina-associated polypeptide 2, isoforms beta/delta/epsilon/gamma (Thymopoietin isoforms beta/delta/epsilon/gamma) (TP beta/delta/epsilon/gamma) | -0,301191794 | 0,017958 | 0,28798242 |
| Q5SUF2 | LC7L3 | Luc7-like protein 3 (Cisplatin resistance-associated-overexpressed protein) | -0,370691062 | 0,009823 | 0,211856445 |
| Q7TNG8 | LDHD | Probable D-lactate dehydrogenase, mitochondrial (DLD) (Lactate dehydrogenase D) (EC 1.1.2.4) | -0,307308632 | 0,025521 | 0,342191281 |
| P16110 | LEG3 | Galectin-3 (Gal-3) (35 kDa lectin) (Carbohydrate-binding protein 35) (CBP 35) (Galactose-specific lectin 3) (IgE-binding protein) (L-34 galactoside-binding lectin) (Laminin-binding protein) (Lectin L-29) (Mac-2 antigen) | 0,807522817 | 9,11E-06 | 0,001912032 |
| O08573 | LEG9 | Galectin-9 (Gal-9) | -0,355171423 | 0,020757 | 0,307344621 |
| P14733 | LMNB1 | Lamin-B1 | -0,501124499 | 0,000119 | 0,015548468 |
| P21619 | LMNB2 | Lamin-B2 | 0,425482916 | 0,000467 | 0,036604522 |
| Q3UJB3 | LR14B | Leucine-rich repeat-containing protein 14B | 0,367651702 | 0,023309 | 0,324237803 |
| Q9QXA5 | LSM4 | U6 snRNA-associated Sm-like protein LSm4 | -0,187921325 | 0,027551 | 0,358456285 |
| Q8CG19 | LTBP1 | Latent-transforming growth factor beta-binding protein 1 (LTBP-1) (Transforming growth factor beta-1-binding protein 1) (TGF-beta1-BP-1) | 0,35160524 | 0,021742 | 0,312487475 |
| Q8K4G1 | LTBP4 | Latent-transforming growth factor beta-binding protein 4 (LTBP-4) | -0,503544762 | 0,00038 | 0,033115228 |
| Q922B1 | MACD1 | O-acetyl-ADP-ribose deacetylase MACROD1 (EC 3.2.2.-) (EC 3.5.1.-) (MACRO domain-containing protein 1) (Protein LRP16) ([Protein ADP-ribosylglutamate] hydrolase) | 0,2783527 | 0,003763 | 0,13550848 |
| Q91WP0 | MASP2 | Mannan-binding lectin serine protease 2 (EC 3.4.21.104) (MBL-associated serine protease 2) (Mannose-binding protein-associated serine protease 2) (MASP-2) [Cleaved into: Mannan-binding lectin serine protease 2 A chain; Mannan-binding lectin serine protease 2 B chain] | 0,394434704 | 0,010006 | 0,212776202 |
| Q60592 | MAST2 | Microtubule-associated serine/threonine-protein kinase 2 (EC 2.7.11.1) | 0,291133951 | 0,008317 | 0,196936869 |
| P39039 | MBL1 | Mannose-binding protein A (MBP-A) (Mannan-binding protein) (Ra-reactive factor polysaccharide-binding component p28B) (RaRF p28B) | 0,556672526 | 0,023644 | 0,327012559 |
| A6H5Y3 | METH | Methionine synthase (EC 2.1.1.13) (5-methyltetrahydrofolate--homocysteine methyltransferase) (Vitamin-B12 dependent methionine synthase) (MS) | 0,199063468 | 0,039897 | 0,428097146 |
| Q6PCP5 | MFF | Mitochondrial fission factor | -0,20325989 | 0,043037 | 0,438864843 |
| P21956 | MFGM | Lactadherin (MFGM) (Milk fat globule-EGF factor 8) (MFG-E8) (SED1) (Sperm surface protein SP47) (MP47) | 0,637278779 | 1,08E-05 | 0,002094406 |
| Q5NCS9 | MID49 | Mitochondrial dynamics protein MID49 (Mitochondrial dynamics protein of 49 kDa homolog) (Mitochondrial elongation factor 2) (Smith-Magenis syndrome chromosomal region candidate gene 7 protein homolog) | -0,190941774 | 0,032821 | 0,395082396 |
| P34884 | MIF | Macrophage migration inhibitory factor (MIF) (EC 5.3.2.1) (Delayed early response protein 6) (DER6) (Glycosylation-inhibiting factor) (GIF) (L-dopachrome isomerase) (L-dopachrome tautomerase) (EC 5.3.3.12) (Phenylpyruvate tautomerase) | 0,326626367 | 0,042089 | 0,437527 |
| Q62000 | MIME | Mimecan (Osteoglycin) | 0,540235098 | 0,000945 | 0,056299993 |
| Q9JM52 | MINK1 | Misshapen-like kinase 1 (EC 2.7.11.1) (GCK family kinase MiNK) (MAPK/ERK kinase kinase kinase 6) (MEK kinase kinase 6) (MEKKK 6) (Misshapen/NIK-related kinase) (Mitogen-activated protein kinase kinase kinase kinase 6) | 0,197464789 | 0,043159 | 0,438864843 |
| Q8BG51 | MIRO1 | Mitochondrial Rho GTPase 1 (MIRO-1) (EC 3.6.5.-) (Ras homolog gene family member T1) | -0,203917274 | 0,021495 | 0,312487475 |
| Q9QWV4 | MLF1 | Myeloid leukemia factor 1 (Hematopoietic lineage switch 7) (Myelodysplasia-myeloid leukemia factor 1) | -0,419898461 | 0,004774 | 0,152623272 |
| Q9CQV6 | MLP3B | Microtubule-associated proteins 1A/1B light chain 3B (Autophagy-related protein LC3 B) (Autophagy-related ubiquitin-like modifier LC3 B) (MAP1 light chain 3-like protein 2) (MAP1A/MAP1B light chain 3 B) (MAP1A/MAP1B LC3 B) (Microtubule-associated protein 1 light chain 3 beta) | 0,428644867 | 0,002623 | 0,110079866 |
| Q8C7H1 | MMAA | Methylmalonic aciduria type A homolog, mitochondrial (EC 3.6.-.-) | -0,194917587 | 0,026032 | 0,346910452 |
| Q9D273 | MMAB | Cob(I)yrinic acid a,c-diamide adenosyltransferase, mitochondrial (EC 2.5.1.17) (Cob(I)alamin adenosyltransferase) (Methylmalonic aciduria type B homolog) | 0,295169643 | 0,005297 | 0,162863549 |
| P33434 | MMP2 | 72 kDa type IV collagenase (EC 3.4.24.24) (72 kDa gelatinase) (Gelatinase A) (Matrix metalloproteinase-2) (MMP-2) [Cleaved into: PEX] | 0,275876049 | 0,020559 | 0,306686651 |
| O09110 | MP2K3 | Dual specificity mitogen-activated protein kinase kinase 3 (MAP kinase kinase 3) (MAPKK 3) (EC 2.7.12.2) (MAPK/ERK kinase 3) (MEK 3) | -0,199142299 | 0,034643 | 0,40885667 |
| Q9WV34 | MPP2 | MAGUK p55 subfamily member 2 (Discs large homolog 2) (Protein MPP2) | 0,298754097 | 0,009708 | 0,211297351 |
| Q7TT50 | MRCKB | Serine/threonine-protein kinase MRCK beta (EC 2.7.11.1) (CDC42-binding protein kinase beta) (DMPK-like beta) (Myotonic dystrophy kinase-related CDC42-binding kinase beta) (MRCK beta) (Myotonic dystrophy protein kinase-like beta) | -0,319495579 | 0,006437 | 0,177049395 |
| Q61216 | MRE11 | Double-strand break repair protein MRE11 (Double-strand break repair protein MRE11A) (MmMRE11A) (Meiotic recombination 11 homolog 1) (MRE11 homolog 1) (Meiotic recombination 11 homolog A) (MRE11 homolog A) | 0,497888779 | 0,025152 | 0,341171474 |
| Q99J25 | MRM1 | rRNA methyltransferase 1, mitochondrial (EC 2.1.1.-) (16S rRNA (guanosine(1145)-2'-O)-methyltransferase) (16S rRNA [Gm1145] 2'-O-methyltransferase) | -0,331128021 | 0,003328 | 0,128509517 |
| Q9Z2C4 | MTMR1 | Myotubularin-related protein 1 (Phosphatidylinositol-3,5-bisphosphate 3-phosphatase) (EC 3.1.3.95) (Phosphatidylinositol-3-phosphate phosphatase) (EC 3.1.3.64) | 0,242422748 | 0,03343 | 0,399417743 |
| Q5HZI1 | MTUS1 | Microtubule-associated tumor suppressor 1 homolog (AT2 receptor-binding protein) (Angiotensin-II type 2 receptor-interacting protein) (Coiled-coiled tumor suppressor gene 1 protein) (Mitochondrial tumor suppressor 1 homolog) | -0,348255852 | 0,030479 | 0,380052223 |
| Q8C854 | MYEF2 | Myelin expression factor 2 (MEF-2) (MyEF-2) | -0,399424035 | 0,002093 | 0,095316291 |
| Q91Z83 | MYH7 | Myosin-7 (Myosin heavy chain 7) (Myosin heavy chain slow isoform) (MyHC-slow) (Myosin heavy chain, cardiac muscle beta isoform) (MyHC-beta) | 1,270868705 | 0,000172 | 0,019711171 |
| Q8CI43 | MYL6B | Myosin light chain 6B (Smooth muscle and nonmuscle myosin light chain alkali 6B) | 0,498780093 | 0,000428 | 0,035078895 |
| P21271 | MYO5B | Unconventional myosin-Vb | 0,314546384 | 0,017236 | 0,280125886 |
| Q80YT7 | MYOME | Myomegalin (Phosphodiesterase 4D-interacting protein) | -0,434020298 | 0,005687 | 0,166310605 |
| Q9JIF9 | MYOTI | Myotilin (Myofibrillar titin-like Ig domains protein) (Titin immunoglobulin domain protein) | 1,1592966 | 2,31E-08 | 3,69E-05 |
| Q9D7V9 | NAAA | N-acylethanolamine-hydrolyzing acid amidase (EC 3.5.1.-) (N-acylsphingosine amidohydrolase-like) (ASAH-like protein) [Cleaved into: N-acylethanolamine-hydrolyzing acid amidase subunit alpha; N-acylethanolamine-hydrolyzing acid amidase subunit beta] | 0,287599067 | 0,026408 | 0,349584253 |
| P70414 | NAC1 | Sodium/calcium exchanger 1 (Na(+)/Ca(2+)-exchange protein 1) (Solute carrier family 8 member 1) | 0,212047369 | 0,017178 | 0,280122219 |
| Q9QWR8 | NAGAB | Alpha-N-acetylgalactosaminidase (EC 3.2.1.49) (Alpha-galactosidase B) | 0,44026526 | 0,000216 | 0,021766647 |
| Q9QZ08 | NAGK | N-acetyl-D-glucosamine kinase (N-acetylglucosamine kinase) (EC 2.7.1.59) (GlcNAc kinase) | 0,416084272 | 0,010427 | 0,218832595 |
| Q8C5H8 | NAKD2 | NAD kinase 2, mitochondrial (EC 2.7.1.23) (Mitochondrial NAD kinase) (NAD kinase domain-containing protein 1, mitochondrial) | 0,291273961 | 0,032211 | 0,391646954 |
| Q9DB73 | NB5R1 | NADH-cytochrome b5 reductase 1 (b5R.1) (EC 1.6.2.2) (NAD(P)H:quinone oxidoreductase type 3 polypeptide A2) | -0,306043547 | 0,032024 | 0,390350735 |
| Q9DCN2 | NB5R3 | NADH-cytochrome b5 reductase 3 (B5R) (Cytochrome b5 reductase) (EC 1.6.2.2) (Diaphorase-1) [Cleaved into: NADH-cytochrome b5 reductase 3 membrane-bound form; NADH-cytochrome b5 reductase 3 soluble form] | 0,224676774 | 0,013399 | 0,243565124 |
| P97432 | NBR1 | Next to BRCA1 gene 1 protein (Membrane component chromosome 17 surface marker 2 homolog) (Neighbor of BRCA1 gene 1 protein) | 0,287474402 | 0,005026 | 0,156516703 |
| Q91WV0 | NC2B | Protein Dr1 (Down-regulator of transcription 1) (Negative cofactor 2-beta) (NC2-beta) (TATA-binding protein-associated phosphoprotein) | -0,22306762 | 0,023001 | 0,321133804 |
| Q8BLF1 | NCEH1 | Neutral cholesterol ester hydrolase 1 (NCEH) (EC 3.1.1.-) (Arylacetamide deacetylase-like 1) (Chlorpyrifos oxon-binding protein) (CPO-BP) | 0,273673806 | 0,049685 | 0,469528405 |
| Q8BTG7 | NDRG4 | Protein NDRG4 (N-myc downstream-regulated gene 4 protein) (Protein Ndr4) | -0,352936676 | 0,022425 | 0,316512815 |
| P0DN34 | NDUB1 | NADH dehydrogenase [ubiquinone] 1 beta subcomplex subunit 1 (Complex I-MNLL) (CI-MNLL) (NADH-ubiquinone oxidoreductase MNLL subunit) | -0,681201462 | 0,003695 | 0,134113685 |
| Q9CQC7 | NDUB4 | NADH dehydrogenase [ubiquinone] 1 beta subcomplex subunit 4 (Complex I-B15) (CI-B15) (NADH-ubiquinone oxidoreductase B15 subunit) | -0,247940723 | 0,048468 | 0,464199682 |
| Q3UIU2 | NDUB6 | NADH dehydrogenase [ubiquinone] 1 beta subcomplex subunit 6 (Complex I-B17) (CI-B17) (NADH-ubiquinone oxidoreductase B17 subunit) | -0,194640536 | 0,036263 | 0,412690946 |
| Q6R891 | NEB2 | Neurabin-2 (Neurabin-II) (Protein phosphatase 1 regulatory subunit 9B) (Spinophilin) | -0,220442 | 0,047248 | 0,456655537 |
| Q6P5H2 | NEST | Nestin | -0,223381612 | 0,021724 | 0,312487475 |
| P08551 | NFL | Neurofilament light polypeptide (NF-L) (68 kDa neurofilament protein) (Neurofilament triplet L protein) | 0,791080901 | 0,000473 | 0,036604522 |
| Q8K4Z3 | NNRE | NAD(P)H-hydrate epimerase (EC 5.1.99.6) (Apolipoprotein A-I-binding protein) (AI-BP) (NAD(P)HX epimerase) | 0,316506423 | 0,047302 | 0,456655537 |
| Q61941 | NNTM | NAD(P) transhydrogenase, mitochondrial (EC 1.6.1.2) (Nicotinamide nucleotide transhydrogenase) (Pyridine nucleotide transhydrogenase) | 1,513368414 | 0,007913 | 0,195884789 |
| Q9JJG9 | NOA1 | Nitric oxide-associated protein 1 | 0,456641424 | 0,002261 | 0,099736537 |
| Q6WKZ7 | NOSTN | Nostrin (Disabled homolog 2-interacting protein 2) (Dab2-interacting protein 2) (Nitric oxide synthase trafficker) (eNOS-trafficking inducer) | 0,316455986 | 0,000987 | 0,0567535 |
| Q9DCJ9 | NPL | N-acetylneuraminate lyase (NALase) (EC 4.1.3.3) (N-acetylneuraminate pyruvate-lyase) (N-acetylneuraminic acid aldolase) (Sialate lyase) (Sialate-pyruvate lyase) (Sialic acid aldolase) (Sialic acid lyase) | 0,377601452 | 0,035485 | 0,410752267 |
| Q64669 | NQO1 | NAD(P)H dehydrogenase [quinone] 1 (EC 1.6.5.2) (Azoreductase) (DT-diaphorase) (DTD) (Menadione reductase) (NAD(P)H:quinone oxidoreductase 1) (Phylloquinone reductase) (Quinone reductase 1) (QR1) | 0,517126399 | 0,022011 | 0,312487475 |
| Q8VC65 | NRM | Nurim (Nuclear envelope membrane protein) (Nuclear rim protein) | 0,202981208 | 0,03279 | 0,395082396 |
| Q9D720 | NSE1 | Non-structural maintenance of chromosomes element 1 homolog (Non-SMC element 1 homolog) (EC 2.3.2.27) | -0,2428602 | 0,043591 | 0,43995108 |
| Q8VCE6 | NT5M | 5'(3')-deoxyribonucleotidase, mitochondrial (5',3'-nucleotidase, mitochondrial) (EC 3.1.3.-) (Deoxy-5'-nucleotidase 2) (dNT-2) | 0,261431768 | 0,005532 | 0,166310605 |
| Q8JZU0 | NUD13 | Nucleoside diphosphate-linked moiety X motif 13 (Nudix motif 13) (EC 3.-.-.-) | 0,293152017 | 0,034947 | 0,409433891 |
| Q6P3D0 | NUD16 | U8 snoRNA-decapping enzyme (EC 3.6.1.62) (IDP phosphatase) (IDPase) (EC 3.6.1.64) (Inosine diphosphate phosphatase) (Nucleoside diphosphate-linked moiety X motif 16) (Nudix motif 16) (m7GpppN-mRNA hydrolase) | 0,713991557 | 7,85E-06 | 0,001804631 |
| D3YYU8 | OBSL1 | Obscurin-like protein 1 | -0,151609176 | 0,042127 | 0,437527 |
| Q9D8W7 | OCAD2 | OCIA domain-containing protein 2 | 0,382874724 | 0,002063 | 0,094860698 |
| P35487 | ODPAT | Pyruvate dehydrogenase E1 component subunit alpha, testis-specific form, mitochondrial (EC 1.2.4.1) (PDHE1-A type II) | 0,273216791 | 0,020649 | 0,306686651 |
| Q8K3K8 | OPTN | Optineurin | 0,291156233 | 0,010625 | 0,219018194 |
| Q9WVD5 | ORNT1 | Mitochondrial ornithine transporter 1 (Solute carrier family 25 member 15) | -0,185787993 | 0,02868 | 0,363349811 |
| Q91XL9 | OSBL1 | Oxysterol-binding protein-related protein 1 (ORP-1) (OSBP-related protein 1) | 0,324723446 | 0,015948 | 0,268323331 |
| Q8C165 | P20D1 | N-fatty-acyl-amino acid synthase/hydrolase PM20D1 (EC 3.5.1.-) (EC 4.3.-.-) (Peptidase M20 domain-containing protein 1) | 0,255341358 | 0,045845 | 0,447706624 |
| Q9JJX6 | P2RX4 | P2X purinoceptor 4 (P2X4) (ATP receptor) (Purinergic receptor) | 0,484919179 | 7,75E-05 | 0,010999664 |
| Q60715 | P4HA1 | Prolyl 4-hydroxylase subunit alpha-1 (4-PH alpha-1) (EC 1.14.11.2) (Procollagen-proline,2-oxoglutarate-4-dioxygenase subunit alpha-1) | -0,26351539 | 0,010755 | 0,219982975 |
| Q9DCC4 | P5CR3 | Pyrroline-5-carboxylate reductase 3 (P5C reductase 3) (P5CR 3) (EC 1.5.1.2) (Pyrroline-5-carboxylate reductase-like protein) | -0,238103197 | 0,040481 | 0,433265159 |
| Q8K212 | PACS1 | Phosphofurin acidic cluster sorting protein 1 (PACS-1) | -0,207096462 | 0,024742 | 0,338327834 |
| P11103 | PARP1 | Poly [ADP-ribose] polymerase 1 (PARP-1) (EC 2.4.2.30) (ADP-ribosyltransferase diphtheria toxin-like 1) (ARTD1) (NAD(+) ADP-ribosyltransferase 1) (ADPRT 1) (Poly[ADP-ribose] synthase 1) (msPARP) | -0,259012422 | 0,012584 | 0,235257931 |
| Q64338 | PDE1C | Calcium/calmodulin-dependent 3',5'-cyclic nucleotide phosphodiesterase 1C (Cam-PDE 1C) (EC 3.1.4.17) | -0,468200118 | 0,000384 | 0,033115228 |
| Q8BFP9 | PDK1 | [Pyruvate dehydrogenase (acetyl-transferring)] kinase isozyme 1, mitochondrial (EC 2.7.11.2) (Pyruvate dehydrogenase kinase isoform 1) (PDH kinase 1) | 0,174249873 | 0,04705 | 0,456040272 |
| P70271 | PDLI4 | PDZ and LIM domain protein 4 (LIM protein RIL) (Reversion-induced LIM protein) | 0,239549445 | 0,044261 | 0,440648118 |
| Q99MZ7 | PECR | Peroxisomal trans-2-enoyl-CoA reductase (TERP) (EC 1.3.1.38) | 0,279147359 | 0,007317 | 0,186862261 |
| Q9R269 | PEPL | Periplakin | 0,475425548 | 0,000318 | 0,029574232 |
| P15331 | PERI | Peripherin | 0,433558979 | 0,00253 | 0,108077501 |
| Q9DB60 | PGFS | Prostamide/prostaglandin F synthase (Prostamide/PG F synthase) (Prostamide/PGF synthase) (EC 1.11.1.20) | 0,242578459 | 0,025068 | 0,341171474 |
| P28654 | PGS2 | Decorin (Bone proteoglycan II) (PG-S2) (PG40) | 0,385127399 | 0,008292 | 0,196936869 |
| Q9DB26 | PHYD1 | Phytanoyl-CoA dioxygenase domain-containing protein 1 (EC 1.-.-.-) | 0,423387436 | 0,002597 | 0,109966344 |
| P42337 | PK3CA | Phosphatidylinositol 4,5-bisphosphate 3-kinase catalytic subunit alpha isoform (PI3-kinase subunit alpha) (PI3K-alpha) (PI3Kalpha) (PtdIns-3-kinase subunit alpha) (EC 2.7.1.153) (Phosphatidylinositol 4,5-bisphosphate 3-kinase 110 kDa catalytic subunit alpha) (PtdIns-3-kinase subunit p110-alpha) (p110alpha) (Phosphoinositide-3-kinase catalytic alpha polypeptide) (Serine/threonine protein kinase PIK3CA) (EC 2.7.11.1) | -0,461741717 | 0,037841 | 0,415979268 |
| P70268 | PKN1 | Serine/threonine-protein kinase N1 (EC 2.7.11.13) (Protein kinase C-like 1) (Protein kinase C-like PKN) (Protein-kinase C-related kinase 1) (Serine-threonine protein kinase N) | -0,215387263 | 0,019032 | 0,297300125 |
| Q68FH0 | PKP4 | Plakophilin-4 (Armadillo-related protein) | 0,301690205 | 0,004215 | 0,142207105 |
| Q8CJH3 | PLXB1 | Plexin-B1 | 0,274627047 | 0,012469 | 0,235257931 |
| B2RXS4 | PLXB2 | Plexin-B2 | -0,193126845 | 0,026857 | 0,353236889 |
| Q9D358 | PPAC | Low molecular weight phosphotyrosine protein phosphatase (LMW-PTP) (LMW-PTPase) (EC 3.1.3.48) (Low molecular weight cytosolic acid phosphatase) (EC 3.1.3.2) | 0,557651064 | 0,021992 | 0,312487475 |
| P09242 | PPBT | Alkaline phosphatase, tissue-nonspecific isozyme (AP-TNAP) (TNSALP) (EC 3.1.3.1) (Alkaline phosphatase 2) (Alkaline phosphatase liver/bone/kidney isozyme) | 0,293767901 | 0,013711 | 0,246948997 |
| P16675 | PPGB | Lysosomal protective protein (EC 3.4.16.5) (Carboxypeptidase C) (Carboxypeptidase L) (Cathepsin A) (Protective protein cathepsin A) (PPCA) (Protective protein for beta-galactosidase) [Cleaved into: Lysosomal protective protein 32 kDa chain; Lysosomal protective protein 20 kDa chain] | 0,225835291 | 0,043567 | 0,43995108 |
| Q9EPX2 | PPN | Papilin | 0,289111868 | 0,005745 | 0,166310605 |
| Q8BKR5 | PPR37 | Protein phosphatase 1 regulatory subunit 37 (Leucine-rich repeat-containing protein 68) | 0,28804902 | 0,015888 | 0,268323331 |
| O88531 | PPT1 | Palmitoyl-protein thioesterase 1 (PPT-1) (EC 3.1.2.22) (Palmitoyl-protein hydrolase 1) | 1,116476595 | 4,18E-08 | 4,04E-05 |
| Q0VBB0 | PRLD2 | PRELI domain-containing protein 2 | 0,591720889 | 0,004334 | 0,144271139 |
| Q9WU79 | PROD | Proline dehydrogenase 1, mitochondrial (EC 1.5.5.2) (Proline oxidase) | -0,283948217 | 0,011378 | 0,22979149 |
| Q9CS42 | PRPS2 | Ribose-phosphate pyrophosphokinase 2 (EC 2.7.6.1) (Phosphoribosyl pyrophosphate synthase II) (PRS-II) | 0,718657789 | 0,000259 | 0,02552931 |
| Q52KR3 | PRUN2 | Protein prune homolog 2 (BNIP2 motif-containing molecule at the C-terminal region 1) | 0,416549981 | 0,00134 | 0,069568739 |
| Q9D820 | PRXD1 | Prolyl-tRNA synthetase associated domain-containing protein 1 (PrdX deacylase domain-containing protein 1) | -0,210805282 | 0,031774 | 0,38958903 |
| Q99JF8 | PSIP1 | PC4 and SFRS1-interacting protein (Lens epithelium-derived growth factor) (mLEDGF) | -0,273625853 | 0,018118 | 0,288637127 |
| Q91Z31 | PTBP2 | Polypyrimidine tract-binding protein 2 (Brain-enriched polypyrimidine tract-binding protein) (Brain-enriched PTB) (Neural polypyrimidine tract-binding protein) (RRM-type RNA-binding protein brPTB) | -0,444856542 | 0,001564 | 0,076415726 |
| O09114 | PTGDS | Prostaglandin-H2 D-isomerase (EC 5.3.99.2) (Glutathione-independent PGD synthase) (Lipocalin-type prostaglandin-D synthase) (Prostaglandin-D2 synthase) (L-PGDS) (PGD2 synthase) (PGDS2) | 1,215756715 | 8,29E-07 | 0,000363608 |
| Q3UQ28 | PXDN | Peroxidasin homolog (EC 1.11.1.7) | -0,60571749 | 1,10E-06 | 0,000409345 |
| Q69ZS0 | PZRN3 | E3 ubiquitin-protein ligase PDZRN3 (EC 2.3.2.27) (PDZ domain-containing RING finger protein 3) (RING-type E3 ubiquitin transferase PDZRN3) (Semaphorin cytoplasmic domain-associated protein 3) (Protein SEMACAP3) | -0,321076108 | 0,011912 | 0,23226961 |
| Q8BU31 | RAP2C | Ras-related protein Rap-2c | 0,376700793 | 0,027654 | 0,3588391 |
| O88851 | RBBP9 | Putative hydrolase RBBP9 (EC 3.-.-.-) (B5T-overexpressed gene protein) (Protein BOG) (Retinoblastoma-binding protein 9) (RBBP-9) | 0,206617518 | 0,028497 | 0,362935654 |
| B2RY56 | RBM25 | RNA-binding protein 25 (RNA-binding motif protein 25) | -0,285527886 | 0,008197 | 0,196936869 |
| Q8VH51 | RBM39 | RNA-binding protein 39 (Coactivator of activating protein 1 and estrogen receptors) (Coactivator of AP-1 and ERs) (RNA-binding motif protein 39) (RNA-binding region-containing protein 2) (Transcription coactivator CAPER) | -0,234504953 | 0,034067 | 0,403042317 |
| Q91YE7 | RBM5 | RNA-binding protein 5 (Putative tumor suppressor LUCA15) (RNA-binding motif protein 5) | -0,275855858 | 0,006883 | 0,184557321 |
| Q8CEE7 | RDH13 | Retinol dehydrogenase 13 (EC 1.1.1.-) | -0,205253857 | 0,036853 | 0,414658859 |
| Q8C0D4 | RHG12 | Rho GTPase-activating protein 12 (Rho-type GTPase-activating protein 12) | -0,222778176 | 0,01205 | 0,23226961 |
| Q8K0Q5 | RHG18 | Rho GTPase-activating protein 18 (Rho-type GTPase-activating protein 18) | -0,312462119 | 0,030569 | 0,380052223 |
| P59729 | RIN3 | Ras and Rab interactor 3 (Ras interaction/interference protein 3) | -0,337681773 | 0,005754 | 0,166310605 |
| Q920A5 | RISC | Retinoid-inducible serine carboxypeptidase (EC 3.4.16.-) (Serine carboxypeptidase 1) | 0,237775431 | 0,035592 | 0,411015969 |
| P53026 | RL10A | 60S ribosomal protein L10a (CSA-19) (Neural precursor cell expressed developmentally down-regulated protein 6) (NEDD-6) | -0,227914847 | 0,016053 | 0,268323331 |
| P35979 | RL12 | 60S ribosomal protein L12 | -0,269973715 | 0,018818 | 0,295877771 |
| P14115 | RL27A | 60S ribosomal protein L27a (L29) | -0,265794628 | 0,008048 | 0,196676452 |
| P47915 | RL29 | 60S ribosomal protein L29 | -0,191495623 | 0,048983 | 0,466350571 |
| Q9D8E6 | RL4 | 60S ribosomal protein L4 | -0,22159747 | 0,019751 | 0,301695415 |
| Q8BSE0 | RMD2 | Regulator of microtubule dynamics protein 2 (RMD-2) (mRMD-2) (Protein FAM82A1) | -0,193554247 | 0,042314 | 0,437527 |
| Q8CFI7 | RPB2 | DNA-directed RNA polymerase II subunit RPB2 (EC 2.7.7.6) (DNA-directed RNA polymerase II 140 kDa polypeptide) (DNA-directed RNA polymerase II subunit B) (RNA polymerase II subunit 2) (RNA polymerase II subunit B2) | 0,219874224 | 0,020229 | 0,306686651 |
| P62281 | RS11 | 40S ribosomal protein S11 | -0,391944808 | 0,0014 | 0,071739638 |
| P14206 | RSSA | 40S ribosomal protein SA (37 kDa laminin receptor precursor) (37LRP) (37 kDa oncofetal antigen) (37/67 kDa laminin receptor) (LRP/LR) (67 kDa laminin receptor) (67LR) (Laminin receptor 1) (LamR) (Laminin-binding protein precursor p40) (LBP/p40) (OFA/iLRP) | -0,301721817 | 0,039257 | 0,422987119 |
| Q9CPX7 | RT16 | 28S ribosomal protein S16, mitochondrial (MRP-S16) (S16mt) | -0,185268183 | 0,045377 | 0,445196515 |
| Q99P72 | RTN4 | Reticulon-4 (Neurite outgrowth inhibitor) (Nogo protein) | 0,165868711 | 0,045288 | 0,445196515 |
| Q91W34 | RUS1 | RUS1 family protein C16orf58 homolog | 0,274308316 | 0,005585 | 0,166310605 |
| P14069 | S10A6 | Protein S100-A6 (5B10) (Calcyclin) (Prolactin receptor-associated protein) (S100 calcium-binding protein A6) | 0,458791852 | 5,45E-05 | 0,008215527 |
| Q5SWT3 | S2535 | Solute carrier family 25 member 35 | -0,292544471 | 0,011944 | 0,23226961 |
| Q8CIW6 | S26A6 | Solute carrier family 26 member 6 (Anion exchange transporter) (Chloride-formate exchanger) (Pendrin-L1) (Pendrin-like protein 1) (Putative anion transporter-1) (Pat-1) | -0,522459466 | 0,000783 | 0,049064782 |
| Q9JLC8 | SACS | Sacsin (DnaJ homolog subfamily C member 29) (DNAJC29) | -0,239978871 | 0,005619 | 0,166310605 |
| Q9R1T2 | SAE1 | SUMO-activating enzyme subunit 1 (Ubiquitin-like 1-activating enzyme E1A) [Cleaved into: SUMO-activating enzyme subunit 1, N-terminally processed] | -0,199151756 | 0,012361 | 0,234915904 |
| Q9Z1F9 | SAE2 | SUMO-activating enzyme subunit 2 (EC 2.3.2.-) (Anthracycline-associated resistance ARX) (Ubiquitin-like 1-activating enzyme E1B) (Ubiquitin-like modifier-activating enzyme 2) | -0,202328374 | 0,021855 | 0,312487475 |
| Q61207 | SAP | Prosaposin (Sulfated glycoprotein 1) (SGP-1) [Cleaved into: Saposin-A; Saposin-B-Val; Saposin-B; Saposin-C; Saposin-D] | 0,333377596 | 0,003693 | 0,134113685 |
| O35609 | SCAM3 | Secretory carrier-associated membrane protein 3 (Secretory carrier membrane protein 3) | -0,290357216 | 0,003898 | 0,13550848 |
| Q61009 | SCRB1 | Scavenger receptor class B member 1 (SRB1) (SR-BI) | -0,491285917 | 0,001822 | 0,087078883 |
| O35114 | SCRB2 | Lysosome membrane protein 2 (85 kDa lysosomal membrane sialoglycoprotein) (LGP85) (Lysosome membrane protein II) (LIMP II) (Scavenger receptor class B member 2) | 0,413032616 | 0,036151 | 0,412690946 |
| Q9CZC8 | SCRN1 | Secernin-1 | 0,256259623 | 0,048673 | 0,465238409 |
| O08992 | SDCB1 | Syntenin-1 (Scaffold protein Pbp1) (Syndecan-binding protein 1) | 0,7050996 | 0,000119 | 0,015548468 |
| Q8CHH9 | SEPT8 | Septin-8 | -0,453163045 | 0,000716 | 0,04547563 |
| Q9QZI9 | SERC3 | Serine incorporator 3 (Axotomy-induced glyco/Golgi protein 1) (AIGP-1) (Axotomy-induced glycoprotein 1) (Membrane protein TMS-1) (Tumor differentially expressed protein 1) | 0,184907624 | 0,045048 | 0,445196515 |
| Q9EPB5 | SERHL | Serine hydrolase-like protein (SHL) (EC 3.1.-.-) | 0,23939699 | 0,009942 | 0,212776202 |
| P19324 | SERPH | Serpin H1 (47 kDa heat shock protein) (Collagen-binding protein) (Colligin) (Serine protease inhibitor J6) | -0,251690386 | 0,021574 | 0,312487475 |
| Q8R0X7 | SGPL1 | Sphingosine-1-phosphate lyase 1 (S1PL) (SP-lyase 1) (SPL 1) (mSPL) (EC 4.1.2.27) (Sphingosine-1-phosphate aldolase) | -0,432002996 | 0,013091 | 0,240263216 |
| Q9ES52 | SHIP1 | Phosphatidylinositol 3,4,5-trisphosphate 5-phosphatase 1 (EC 3.1.3.86) (Inositol polyphosphate-5-phosphatase of 145 kDa) (SIP-145) (SH2 domain-containing inositol 5'-phosphatase 1) (SH2 domain-containing inositol phosphatase 1) (SHIP-1) (p150Ship) | -0,35442138 | 0,042127 | 0,437527 |
| Q6P549 | SHIP2 | Phosphatidylinositol 3,4,5-trisphosphate 5-phosphatase 2 (EC 3.1.3.86) (AblSH3-binding protein) (Inositol polyphosphate phosphatase-like protein 1) (INPPL-1) (SH2 domain-containing inositol 5'-phosphatase 2) (SH2 domain-containing inositol phosphatase 2) (SHIP-2) | -0,18967663 | 0,016065 | 0,268323331 |
| Q80TE4 | SI1L2 | Signal-induced proliferation-associated 1-like protein 2 (SIPA1-like protein 2) | -0,243094279 | 0,017106 | 0,279901394 |
| Q61165 | SL9A1 | Sodium/hydrogen exchanger 1 (Na(+)/H(+) exchanger 1) (NHE-1) (Solute carrier family 9 member 1) | -0,278854919 | 0,040575 | 0,43331119 |
| P97471 | SMAD4 | Mothers against decapentaplegic homolog 4 (MAD homolog 4) (Mothers against DPP homolog 4) (Deletion target in pancreatic carcinoma 4 homolog) (SMAD family member 4) (SMAD 4) (Smad4) | -0,200734923 | 0,012611 | 0,235257931 |
| Q6DIC0 | SMCA2 | Probable global transcription activator SNF2L2 (EC 3.6.4.-) (ATP-dependent helicase SMARCA2) (BRG1-associated factor 190B) (BAF190B) (Protein brahma homolog) (SNF2-alpha) (SWI/SNF-related matrix-associated actin-dependent regulator of chromatin subfamily A member 2) | -0,208199324 | 0,047521 | 0,457853906 |
| Q3TKT4 | SMCA4 | Transcription activator BRG1 (EC 3.6.4.-) (ATP-dependent helicase SMARCA4) (BRG1-associated factor 190A) (BAF190A) (Protein brahma homolog 1) (SNF2-beta) (SWI/SNF-related matrix-associated actin-dependent regulator of chromatin subfamily A member 4) | -0,343513614 | 0,004931 | 0,154696257 |
| Q3UMB5 | SMCR8 | Guanine nucleotide exchange protein SMCR8 (Smith-Magenis syndrome chromosomal region candidate gene 8 protein homolog) | 0,233690187 | 0,028976 | 0,365516943 |
| Q8C1Q6 | SMIM4 | Small integral membrane protein 4 (Small nucleolar RNA host gene 8) | 0,249309526 | 0,028238 | 0,361546694 |
| O55101 | SNG2 | Synaptogyrin-2 (Cellugyrin) | -0,499848427 | 0,003958 | 0,13550848 |
| Q99L88 | SNTB1 | Beta-1-syntrophin (59 kDa dystrophin-associated protein A1 basic component 1) (DAPA1B) (Syntrophin-2) | -0,677799576 | 8,67E-06 | 0,001903089 |
| O70492 | SNX3 | Sorting nexin-3 (SDP3 protein) | -0,145630251 | 0,045325 | 0,445196515 |
| O35892 | SP100 | Nuclear autoantigen Sp-100 (Nuclear dot-associated Sp100 protein) (Speckled 100 kDa) | -0,22821156 | 0,036213 | 0,412690946 |
| Q91WP6 | SPA3N | Serine protease inhibitor A3N (Serpin A3N) | 0,374562163 | 0,007059 | 0,184972448 |
| O08800 | SPB8 | Serpin B8 | 0,203219997 | 0,035166 | 0,410017256 |
| Q9CQC8 | SPG21 | Maspardin (Acid cluster protein 33) (Spastic paraplegia 21 autosomal recessive Mast syndrome protein homolog) | 0,287484212 | 0,009111 | 0,205504763 |
| Q9ESJ4 | SPN90 | NCK-interacting protein with SH3 domain (54 kDa VacA-interacting protein) (VIP54) (90 kDa N-WASP-interacting protein) (90 kDa SH3 protein interacting with Nck) (SH3 adapter protein SPIN90) (WASP-interacting SH3-domain protein) (WISH) (Wiskott-Aldrich syndrome protein-binding protein) (N-WASP-binding protein) | -0,176731012 | 0,042178 | 0,437527 |
| P07214 | SPRC | SPARC (Basement-membrane protein 40) (BM-40) (Osteonectin) (ON) (Secreted protein acidic and rich in cysteine) | -0,35865402 | 0,0218 | 0,312487475 |
| P52840 | ST1A1 | Sulfotransferase 1A1 (ST1A1) (EC 2.8.2.1) (Aryl sulfotransferase) (Phenol sulfotransferase) (Phenol/aryl sulfotransferase) (mSTp1) (ST1A4) (Sulfokinase) | 0,301375835 | 0,011113 | 0,22611708 |
| Q9WVL2 | STAT2 | Signal transducer and activator of transcription 2 | -0,20131577 | 0,020547 | 0,306686651 |
| Q8C7E7 | STBD1 | Starch-binding domain-containing protein 1 (Genethonin-1) (Glycophagy cargo receptor stbd1) | 0,30503437 | 0,01483 | 0,260315294 |
| Q3TBT3 | STING | Stimulator of interferon genes protein (mSTING) (Endoplasmic reticulum interferon stimulator) (ERIS) (Mediator of IRF3 activation) (MMITA) (Transmembrane protein 173) | -0,317242079 | 0,008781 | 0,201835689 |
| O55098 | STK10 | Serine/threonine-protein kinase 10 (EC 2.7.11.1) (Lymphocyte-oriented kinase) | -0,213874294 | 0,031087 | 0,383771212 |
| P54116 | STOM | Erythrocyte band 7 integral membrane protein (Protein 7.2b) (Stomatin) | 0,248416806 | 0,014315 | 0,254047082 |
| Q9ER00 | STX12 | Syntaxin-12 | 0,26079686 | 0,011708 | 0,23066239 |
| Q8K1E0 | STX5 | Syntaxin-5 | -0,257574399 | 0,01921 | 0,298155278 |
| Q9Z2I8 | SUCB2 | Succinate--CoA ligase [GDP-forming] subunit beta, mitochondrial (EC 6.2.1.4) (GTP-specific succinyl-CoA synthetase subunit beta) (G-SCS) (GTPSCS) (Succinyl-CoA synthetase beta-G chain) (SCS-betaG) | 0,416273018 | 0,000969 | 0,0567535 |
| Q8CFG0 | SULF2 | Extracellular sulfatase Sulf-2 (mSulf-2) (EC 3.1.6.-) | 0,700448362 | 0,000183 | 0,020125148 |
| Q9DBX3 | SUSD2 | Sushi domain-containing protein 2 | 0,295231071 | 0,016982 | 0,27881005 |
| Q9CXJ1 | SYEM | Probable glutamate--tRNA ligase, mitochondrial (EC 6.1.1.17) (Glutamyl-tRNA synthetase) (GluRS) | -0,274068781 | 0,011532 | 0,229977318 |
| Q4FZC9 | SYNE3 | Nesprin-3 (KASH domain-containing protein 3) (KASH3) (Nuclear envelope spectrin repeat protein 3) | 0,202370037 | 0,038961 | 0,421667798 |
| Q9Z0F7 | SYUG | Gamma-synuclein (Persyn) | 0,592226039 | 0,000357 | 0,032489453 |
| Q8K1H7 | T11L2 | T-complex protein 11-like protein 2 | -0,342169645 | 0,006523 | 0,177885125 |
| Q6P8M1 | TATD1 | Putative deoxyribonuclease TATDN1 (EC 3.1.21.-) | 0,361654723 | 0,025483 | 0,342191281 |
| Q9JJZ2 | TBA8 | Tubulin alpha-8 chain (Alpha-tubulin 8) | -0,231892577 | 0,0351 | 0,410017256 |
| P10711 | TCEA1 | Transcription elongation factor A protein 1 (Transcription elongation factor S-II protein 1) (Transcription elongation factor TFIIS.o) | 1,089308397 | 0,002865 | 0,117197441 |
| O08784 | TCOF | Treacle protein (Treacher Collins syndrome protein homolog) | -0,303574279 | 0,008165 | 0,196936869 |
| P11031 | TCP4 | Activated RNA polymerase II transcriptional coactivator p15 (Positive cofactor 4) (PC4) (SUB1 homolog) (Single-stranded DNA-binding protein p9) (p14) | -0,287027697 | 0,007683 | 0,193156495 |
| P80315 | TCPD | T-complex protein 1 subunit delta (TCP-1-delta) (A45) (CCT-delta) | -0,211026746 | 0,035767 | 0,411990545 |
| P70210 | TEAD3 | Transcriptional enhancer factor TEF-5 (DTEF-1) (ETF-related factor 1) (ETFR-1) (TEA domain family member 3) (TEAD-3) | -0,206347607 | 0,048022 | 0,460843788 |
| Q3TL26 | TFB2M | Dimethyladenosine transferase 2, mitochondrial (EC 2.1.1.-) (Mitochondrial 12S rRNA dimethylase 2) (Mitochondrial transcription factor B2) (mTFB2M) (mtTFB2) (S-adenosylmethionine-6-N', N'-adenosyl(rRNA) dimethyltransferase 2) | 0,236617345 | 0,008565 | 0,198767737 |
| Q9ERA6 | TFP11 | Tuftelin-interacting protein 11 (Septin and tuftelin-interacting protein 1) (STIP-1) (Tuftelin-interacting protein 39) | 0,501830395 | 0,012011 | 0,23226961 |
| Q62351 | TFR1 | Transferrin receptor protein 1 (TR) (TfR) (TfR1) (Trfr) (CD antigen CD71) | -0,562741551 | 0,000205 | 0,021549195 |
| O08583 | THOC4 | THO complex subunit 4 (Tho4) (Ally of AML-1 and LEF-1) (Aly/REF export factor) (REF1-I) (RNA and export factor-binding protein 1) (Transcriptional coactivator Aly/REF) | -0,213597114 | 0,026775 | 0,353124341 |
| Q9CR76 | TM186 | Transmembrane protein 186 | 0,397991399 | 0,018923 | 0,296557535 |
| Q9CXT7 | TM192 | Transmembrane protein 192 | 0,398023932 | 0,004242 | 0,142207105 |
| Q91XE8 | TM205 | Transmembrane protein 205 | 0,429087482 | 0,022372 | 0,316512815 |
| Q3TMP8 | TM38A | Trimeric intracellular cation channel type A (TRIC-A) (TRICA) (27 kDa sarcoplasmic reticulum protein) (Mitsugumin-33A) (SPR-27) (Transmembrane protein 38A) | 0,30654964 | 0,037512 | 0,415979268 |
| Q9JJG6 | TMM47 | Transmembrane protein 47 (Transmembrane 4 superfamily member 10) | 0,372790301 | 0,010656 | 0,219018194 |
| Q9D0Z3 | TMM53 | Transmembrane protein 53 | 0,199061975 | 0,01764 | 0,283829174 |
| Q8VBT0 | TMX1 | Thioredoxin-related transmembrane protein 1 (Thioredoxin domain-containing protein 1) | -0,283082765 | 0,003075 | 0,123698629 |
| Q99LG2 | TNPO2 | Transportin-2 (Karyopherin beta-2b) | 0,182543045 | 0,04224 | 0,437527 |
| B1AXP6 | TOM5 | Mitochondrial import receptor subunit TOM5 homolog | -0,299845874 | 0,02159 | 0,312487475 |
| Q04750 | TOP1 | DNA topoisomerase 1 (EC 5.99.1.2) (DNA topoisomerase I) | -0,244860999 | 0,014762 | 0,260056703 |
| Q9DAM5 | TPC | Mitochondrial thiamine pyrophosphate carrier (Solute carrier family 25 member 19) | -0,309985056 | 0,008694 | 0,200791768 |
| O54818 | TPD53 | Tumor protein D53 (mD53) (Tumor protein D52-like 1) | 0,270785671 | 0,018412 | 0,291394385 |
| O89023 | TPP1 | Tripeptidyl-peptidase 1 (TPP-1) (EC 3.4.14.9) (Lysosomal pepstatin-insensitive protease) (LPIC) (Tripeptidyl aminopeptidase) (Tripeptidyl-peptidase I) (TPP-I) | 0,835606957 | 1,54E-07 | 9,27E-05 |
| Q38HM4 | TRI63 | E3 ubiquitin-protein ligase TRIM63 (EC 2.3.2.27) (Muscle-specific RING finger protein 1) (MuRF-1) (MuRF1) (Muscle RING finger protein 1) (RING-type E3 ubiquitin transferase TRIM63) (Tripartite motif-containing protein 63) | -0,222675063 | 0,039258 | 0,422987119 |
| Q923T7 | TRIM7 | E3 ubiquitin-protein ligase TRIM7 (EC 2.3.2.27) (Glycogenin-interacting protein) (Tripartite motif-containing protein 7) | -0,204121908 | 0,020502 | 0,306686651 |
| Q80YV3 | TRRAP | Transformation/transcription domain-associated protein (Tra1 homolog) | -0,258282558 | 0,009461 | 0,208898509 |
| Q9D0B5 | TSTD3 | Thiosulfate sulfurtransferase/rhodanese-like domain-containing protein 3 (Rhodanese domain-containing protein 3) | 0,37030624 | 0,000853 | 0,051497071 |
| Q8CC21 | TTC19 | Tetratricopeptide repeat protein 19, mitochondrial (TPR repeat protein 19) | 0,216326066 | 0,042033 | 0,437527 |
| Q8BG60 | TXNIP | Thioredoxin-interacting protein (Vitamin D3 up-regulated protein 1) | -0,282808061 | 0,021176 | 0,310172316 |
| P24529 | TY3H | Tyrosine 3-monooxygenase (EC 1.14.16.2) (Tyrosine 3-hydroxylase) (TH) | 0,367548285 | 0,000166 | 0,019711171 |
| Q6ZWY8 | TYB10 | Thymosin beta-10 | -0,476195602 | 0,012672 | 0,235257931 |
| Q3TW96 | UAP1L | UDP-N-acetylhexosamine pyrophosphorylase-like protein 1 (EC 2.7.7.-) | 0,409786213 | 0,003412 | 0,130722391 |
| Q9QZU9 | UB2L6 | Ubiquitin/ISG15-conjugating enzyme E2 L6 (EC 2.3.2.23) (E2 ubiquitin-conjugating enzyme L6) (UbcM8) (Ubiquitin carrier protein L6) (Ubiquitin-protein ligase L6) | -0,29278856 | 0,020599 | 0,306686651 |
| Q6ZWZ2 | UB2R2 | Ubiquitin-conjugating enzyme E2 R2 (EC 2.3.2.23) (E2 ubiquitin-conjugating enzyme R2) (Ubiquitin carrier protein R2) (Ubiquitin-conjugating enzyme E2-CDC34B) (Ubiquitin-protein ligase R2) | -0,240082668 | 0,045911 | 0,447706624 |
| Q8VE47 | UBA5 | Ubiquitin-like modifier-activating enzyme 5 (Ubiquitin-activating enzyme 5) (UFM1-activating enzyme) (Ubiquitin-activating enzyme E1 domain-containing protein 1) | -0,473123591 | 5,06E-05 | 0,007879449 |
| Q9ES00 | UBE4B | Ubiquitin conjugation factor E4 B (EC 2.3.2.27) (RING-type E3 ubiquitin transferase E4 B) (Ubiquitin fusion degradation protein 2) | -0,228329458 | 0,033513 | 0,399431234 |
| Q5U430 | UBR3 | E3 ubiquitin-protein ligase UBR3 (EC 2.3.2.27) (N-recognin-3) (RING-type E3 ubiquitin transferase UBR3) (Ubiquitin-protein ligase E3-alpha-3) (Ubiquitin-protein ligase E3-alpha-III) (Zinc finger protein 650) | -0,233201416 | 0,019282 | 0,298307804 |
| Q62465 | VAT1 | Synaptic vesicle membrane protein VAT-1 homolog (EC 1.-.-.-) | 0,326385203 | 0,000803 | 0,049663338 |
| Q80TB8 | VAT1L | Synaptic vesicle membrane protein VAT-1 homolog-like (EC 1.-.-.-) | 0,35443352 | 0,00969 | 0,211297351 |
| P29533 | VCAM1 | Vascular cell adhesion protein 1 (V-CAM 1) (VCAM-1) (CD antigen CD106) | 0,353290448 | 0,001 | 0,0567535 |
| P35918 | VGFR2 | Vascular endothelial growth factor receptor 2 (VEGFR-2) (EC 2.7.10.1) (Fetal liver kinase 1) (FLK-1) (Kinase NYK) (Protein-tyrosine kinase receptor flk-1) (CD antigen CD309) | 0,174709095 | 0,048942 | 0,466350571 |
| Q8R307 | VPS18 | Vacuolar protein sorting-associated protein 18 homolog | -0,192062856 | 0,03991 | 0,428097146 |
| Q9EQH3 | VPS35 | Vacuolar protein sorting-associated protein 35 (Maternal-embryonic 3) (Vesicle protein sorting 35) | 0,30755528 | 0,006455 | 0,177049395 |
| Q91XD6 | VPS36 | Vacuolar protein-sorting-associated protein 36 (ESCRT-II complex subunit VPS36) | -0,179848392 | 0,038951 | 0,421667798 |
| Q9CR26 | VTA1 | Vacuolar protein sorting-associated protein VTA1 homolog (SKD1-binding protein 1) (SBP1) | 0,255963732 | 0,026434 | 0,349584253 |
| P29788 | VTNC | Vitronectin (VN) (S-protein) (Serum-spreading factor) | 0,519108621 | 0,014157 | 0,252154535 |
| Q8R2Z5 | VWA1 | von Willebrand factor A domain-containing protein 1 (von Willebrand factor A domain-related protein) | 0,345193194 | 0,004743 | 0,152620144 |
| E9Q4P1 | WDFY1 | WD repeat and FYVE domain-containing protein 1 (WD40- and FYVE domain-containing protein 1) | 0,753482216 | 0,03147 | 0,387513809 |
| Q00519 | XDH | Xanthine dehydrogenase/oxidase [Includes: Xanthine dehydrogenase (XD) (EC 1.17.1.4); Xanthine oxidase (XO) (EC 1.17.3.2) (Xanthine oxidoreductase) (XOR)] | 0,285016616 | 0,009371 | 0,208451208 |
| Q04736 | YES | Tyrosine-protein kinase Yes (EC 2.7.10.2) (Proto-oncogene c-Yes) (p61-Yes) | -0,240572727 | 0,044275 | 0,440648118 |
| Q6PIU9 | YJ005 | Uncharacterized protein FLJ45252 homolog | -0,211957827 | 0,041913 | 0,437527 |
| Q9DCH6 | ZFAN6 | AN1-type zinc finger protein 6 (Associated with PRK1 protein) (Zinc finger A20 domain-containing protein 3) | -0,209649309 | 0,03871 | 0,420979619 |
| Q8R121 | ZPI | Protein Z-dependent protease inhibitor (PZ-dependent protease inhibitor) (PZI) (Serpin A10) | 0,407104296 | 0,018107 | 0,288637127 |

**Supplementary Data Table 4. List of total significantly altered proteins (p-value<0.05) in HGPS mice**

| **Uniprot ID** | **Protein Symbol** | **Protein Description** | **Average Log2 Ratio** | ***p*-value** | **Adjusted *p*-value** |
| --- | --- | --- | --- | --- | --- |
| Q61151 | 2A5E | Serine/threonine-protein phosphatase 2A 56 kDa regulatory subunit epsilon isoform (PP2A B subunit isoform B'-epsilon) (PP2A B subunit isoform B56-epsilon) (PP2A B subunit isoform PR61-epsilon) (PP2A B subunit isoform R5-epsilon) | 0,232616488 | 0,003037 | 0,062377375 |
| Q3UFY7 | 5NT3B | 7-methylguanosine phosphate-specific 5'-nucleotidase (7-methylguanosine nucleotidase) (EC 3.1.3.91) (Cytosolic 5'-nucleotidase 3B) (Cytosolic 5'-nucleotidase III-like protein) (cN-III-like protein) (EC 3.1.3.5) (N(7)-methylguanylate 5'-phosphatase) | 0,145261305 | 0,015133 | 0,154883083 |
| Q3THG9 | AASD1 | Alanyl-tRNA editing protein Aarsd1 (Alanyl-tRNA deacylase alaX) (AlaX) (AlaXp-II) (Alanyl-tRNA synthetase domain-containing protein 1) | 0,233637427 | 0,025421 | 0,203142138 |
| P05202 | AATM | Aspartate aminotransferase, mitochondrial (mAspAT) (EC 2.6.1.1) (EC 2.6.1.7) (Fatty acid-binding protein) (FABP-1) (Glutamate oxaloacetate transaminase 2) (Kynurenine aminotransferase 4) (Kynurenine aminotransferase IV) (Kynurenine--oxoglutarate transaminase 4) (Kynurenine--oxoglutarate transaminase IV) (Plasma membrane-associated fatty acid-binding protein) (FABPpm) (Transaminase A) | -0,179785654 | 0,003456 | 0,065880674 |
| Q9DC29 | ABCB6 | ATP-binding cassette sub-family B member 6, mitochondrial | -0,131998144 | 0,040209 | 0,255035879 |
| Q9JI39 | ABCBA | ATP-binding cassette sub-family B member 10, mitochondrial (ABC-mitochondrial erythroid protein) (ABC-me protein) (ATP-binding cassette transporter 10) (ABC transporter 10 protein) | 0,179310778 | 0,009864 | 0,122276261 |
| P61222 | ABCE1 | ATP-binding cassette sub-family E member 1 (RNase L inhibitor) (Ribonuclease 4 inhibitor) (RNS4I) | -0,1590266 | 0,022819 | 0,190682325 |
| Q9WV35 | ABEC2 | C->U-editing enzyme APOBEC-2 (EC 3.5.4.36) (mRNA(cytosine(6666)) deaminase 2) | 0,18332446 | 0,027772 | 0,212119635 |
| Q8VD66 | ABHD4 | Protein ABHD4 (EC 3.-.-.-) (Alpha/beta hydrolase domain-containing protein 4) (Abhydrolase domain-containing protein 4) (Alpha/beta-hydrolase 4) (Lyso-N-acylphosphatidylethanolamine lipase) | 0,248931523 | 0,005881 | 0,091744717 |
| Q9D7B6 | ACAD8 | Isobutyryl-CoA dehydrogenase, mitochondrial (EC 1.3.99.-) (Acyl-CoA dehydrogenase family member 8) (ACAD-8) | -0,224242929 | 0,011237 | 0,129052993 |
| Q8K370 | ACD10 | Acyl-CoA dehydrogenase family member 10 (ACAD-10) (EC 1.3.99.-) | 0,243041717 | 0,043097 | 0,262537322 |
| Q8VHQ9 | ACO11 | Acyl-coenzyme A thioesterase 11 (Acyl-CoA thioesterase 11) (EC 3.1.2.-) (Acyl-CoA thioester hydrolase 11) (Adipose-associated thioesterase) (Brown fat-inducible thioesterase) (BFIT) | -0,12567838 | 0,048214 | 0,275765509 |
| P28271 | ACOC | Cytoplasmic aconitate hydratase (Aconitase) (EC 4.2.1.3) (Citrate hydro-lyase) (Iron regulatory protein 1) (IRP1) (Iron-responsive element-binding protein 1) (IRE-BP 1) | -0,269492658 | 0,000236 | 0,009612549 |
| Q9CR21 | ACPM | Acyl carrier protein, mitochondrial (ACP) (CI-SDAP) (NADH-ubiquinone oxidoreductase 9.6 kDa subunit) | -0,412136642 | 0,003182 | 0,063752636 |
| Q8VCW8 | ACSF2 | Acyl-CoA synthetase family member 2, mitochondrial (EC 6.2.1.-) | -0,137715089 | 0,029581 | 0,218451424 |
| P41216 | ACSL1 | Long-chain-fatty-acid--CoA ligase 1 (EC 6.2.1.3) (Long-chain acyl-CoA synthetase 1) (LACS 1) | -0,143319016 | 0,015061 | 0,154691136 |
| P62737 | ACTA | Actin, aortic smooth muscle (Alpha-actin-2) [Cleaved into: Actin, aortic smooth muscle, intermediate form] | -0,243945566 | 0,004122 | 0,074227832 |
| P60710 | ACTB | Actin, cytoplasmic 1 (Beta-actin) [Cleaved into: Actin, cytoplasmic 1, N-terminally processed] | -0,312448811 | 0,003298 | 0,064213044 |
| P51881 | ADT2 | ADP/ATP translocase 2 (ADP,ATP carrier protein 2) (Adenine nucleotide translocator 2) (ANT 2) (Solute carrier family 25 member 5) [Cleaved into: ADP/ATP translocase 2, N-terminally processed] | -0,143513138 | 0,030837 | 0,221038409 |
| Q9CYR6 | AGM1 | Phosphoacetylglucosamine mutase (PAGM) (EC 5.4.2.3) (Acetylglucosamine phosphomutase) (N-acetylglucosamine-phosphate mutase) (Phosphoglucomutase-3) (PGM 3) | 0,142742361 | 0,043433 | 0,262537322 |
| Q8CJG0 | AGO2 | Protein argonaute-2 (Argonaute2) (mAgo2) (EC 3.1.26.n2) (Argonaute RISC catalytic component 2) (Eukaryotic translation initiation factor 2C 2) (eIF-2C 2) (eIF2C 2) (Piwi/argonaute family protein meIF2C2) (Protein slicer) | 0,155414333 | 0,025407 | 0,203142138 |
| Q8BK64 | AHSA1 | Activator of 90 kDa heat shock protein ATPase homolog 1 (AHA1) | -0,23582661 | 0,00206 | 0,047975731 |
| Q8R010 | AIMP2 | Aminoacyl tRNA synthase complex-interacting multifunctional protein 2 (Multisynthase complex auxiliary component p38) (Protein JTV-1) | -0,430685839 | 4,08E-05 | 0,002837649 |
| P31750 | AKT1 | RAC-alpha serine/threonine-protein kinase (EC 2.7.11.1) (AKT1 kinase) (Protein kinase B) (PKB) (Protein kinase B alpha) (PKB alpha) (Proto-oncogene c-Akt) (RAC-PK-alpha) (Thymoma viral proto-oncogene) | 0,21033211 | 0,008957 | 0,117848645 |
| Q60823 | AKT2 | RAC-beta serine/threonine-protein kinase (EC 2.7.11.1) (Protein kinase Akt-2) (Protein kinase B beta) (PKB beta) (RAC-PK-beta) | 0,279184112 | 0,001268 | 0,033751711 |
| P47740 | AL3A2 | Fatty aldehyde dehydrogenase (EC 1.2.1.3) (Aldehyde dehydrogenase 3) (Aldehyde dehydrogenase family 3 member A2) | -0,129868431 | 0,036308 | 0,240254199 |
| Q924C5 | ALPK3 | Alpha-protein kinase 3 (EC 2.7.11.-) (Myocytic induction/differentiation originator) (Midori) | 0,229317489 | 0,027917 | 0,212119635 |
| P97449 | AMPN | Aminopeptidase N (AP-N) (mAPN) (EC 3.4.11.2) (Alanyl aminopeptidase) (Aminopeptidase M) (AP-M) (Membrane protein p161) (Microsomal aminopeptidase) (CD antigen CD13) | 0,313560392 | 0,010626 | 0,124983812 |
| O35381 | AN32A | Acidic leucine-rich nuclear phosphoprotein 32 family member A (Acidic nuclear phosphoprotein pp32) (Leucine-rich acidic nuclear protein) (LANP) (Potent heat-stable protein phosphatase 2A inhibitor I1PP2A) | 0,268512308 | 0,004588 | 0,07919585 |
| P59672 | ANS1A | Ankyrin repeat and SAM domain-containing protein 1A (Odin) | 0,15627993 | 0,038773 | 0,24966942 |
| P35585 | AP1M1 | AP-1 complex subunit mu-1 (AP-mu chain family member mu1A) (Adaptor protein complex AP-1 subunit mu-1) (Adaptor-related protein complex 1 subunit mu-1) (Clathrin assembly protein complex 1 mu-1 medium chain 1) (Clathrin coat assembly protein AP47) (Clathrin coat-associated protein AP47) (Golgi adaptor HA1/AP1 adaptin mu-1 subunit) (Mu-adaptin 1) (Mu1A-adaptin) | 0,187181015 | 0,021588 | 0,188260494 |
| P28352 | APEX1 | DNA-(apurinic or apyrimidinic site) lyase (EC 3.1.-.-) (EC 4.2.99.18) (APEX nuclease) (APEN) (Apurinic-apyrimidinic endonuclease 1) (AP endonuclease 1) (REF-1) (Redox factor-1) [Cleaved into: DNA-(apurinic or apyrimidinic site) lyase, mitochondrial] | -0,280376441 | 0,00127 | 0,033751711 |
| P09813 | APOA2 | Apolipoprotein A-II (Apo-AII) (ApoA-II) (Apolipoprotein A2) [Cleaved into: Proapolipoprotein A-II (ProapoA-II)] | -0,274211137 | 0,004053 | 0,073636148 |
| Q9WV32 | ARC1B | Actin-related protein 2/3 complex subunit 1B (Arp2/3 complex 41 kDa subunit) (p41-ARC) | -0,279683532 | 0,00021 | 0,009612549 |
| Q8CG76 | ARK72 | Aflatoxin B1 aldehyde reductase member 2 (EC 1.1.1.n11) (Succinic semialdehyde reductase) (SSA reductase) | -0,448811493 | 7,34E-06 | 0,001020165 |
| Q9QZB7 | ARP10 | Actin-related protein 10 (Actin-related protein 11) | -0,211930067 | 0,005335 | 0,086935286 |
| O54984 | ASNA | ATPase Asna1 (EC 3.6.-.-) (Arsenical pump-driving ATPase) (Arsenite-stimulated ATPase) | 0,156907713 | 0,040646 | 0,256765664 |
| Q6PIE5 | AT1A2 | Sodium/potassium-transporting ATPase subunit alpha-2 (Na(+)/K(+) ATPase alpha-2 subunit) (EC 3.6.3.9) (Na(+)/K(+) ATPase alpha(+) subunit) (Sodium pump subunit alpha-2) | 0,312534228 | 0,000766 | 0,023045423 |
| P14094 | AT1B1 | Sodium/potassium-transporting ATPase subunit beta-1 (Sodium/potassium-dependent ATPase subunit beta-1) | -0,29958819 | 0,00067 | 0,021753075 |
| G5E829 | AT2B1 | Plasma membrane calcium-transporting ATPase 1 (PMCA1) (EC 3.6.3.8) (Plasma membrane calcium ATPase isoform 1) (Plasma membrane calcium pump isoform 1) | -0,120725752 | 0,0363 | 0,240254199 |
| Q6Q477 | AT2B4 | Plasma membrane calcium-transporting ATPase 4 (PMCA4) (EC 3.6.3.8) | -0,296085844 | 0,000727 | 0,022554625 |
| P56382 | ATP5E | ATP synthase subunit epsilon, mitochondrial (ATPase subunit epsilon) (ATP synthase F1 subunit epsilon) | -0,482773284 | 4,56E-06 | 0,000700045 |
| Q9DCX2 | ATP5H | ATP synthase subunit d, mitochondrial (ATPase subunit d) | -0,278678598 | 0,004048 | 0,073636148 |
| Q9CPQ8 | ATP5L | ATP synthase subunit g, mitochondrial (ATPase subunit g) | 0,342405635 | 0,035339 | 0,239730685 |
| Q64373 | B2CL1 | Bcl-2-like protein 1 (Bcl2-L-1) (Apoptosis regulator Bcl-X) | -0,136965814 | 0,012662 | 0,139268033 |
| Q9Z1R2 | BAG6 | Large proline-rich protein BAG6 (BAG family molecular chaperone regulator 6) (BCL2-associated athanogene 6) (BAG-6) (HLA-B-associated transcript 3) (Protein Scythe) | -0,165867204 | 0,027997 | 0,212122278 |
| Q61334 | BAP29 | B-cell receptor-associated protein 29 (BCR-associated protein 29) (Bap29) | -0,16991852 | 0,022876 | 0,190682325 |
| Q9Z277 | BAZ1B | Tyrosine-protein kinase BAZ1B (EC 2.7.10.2) (Bromodomain adjacent to zinc finger domain protein 1B) (Williams syndrome transcription factor homolog) (Williams-Beuren syndrome chromosomal region 9 protein homolog) | 0,129076502 | 0,041749 | 0,259388001 |
| Q9R069 | BCAM | Basal cell adhesion molecule (B-CAM cell surface glycoprotein) (Lutheran antigen) (CD antigen CD239) | -0,325601169 | 0,000231 | 0,009612549 |
| O35855 | BCAT2 | Branched-chain-amino-acid aminotransferase, mitochondrial (BCAT(m)) (EC 2.6.1.42) | -0,330517829 | 0,000126 | 0,006445939 |
| Q9CZP5 | BCS1 | Mitochondrial chaperone BCS1 (BCS1-like protein) | -0,203561972 | 0,022842 | 0,190682325 |
| Q80XN0 | BDH | D-beta-hydroxybutyrate dehydrogenase, mitochondrial (EC 1.1.1.30) (3-hydroxybutyrate dehydrogenase) (BDH) | -0,317334991 | 0,003699 | 0,069605759 |
| Q8R016 | BLMH | Bleomycin hydrolase (BH) (BLM hydrolase) (BMH) (EC 3.4.22.40) | -0,504917218 | 0,012165 | 0,137006045 |
| Q8CEI1 | BOLA3 | BolA-like protein 3 | -0,790214948 | 2,06E-06 | 0,000400219 |
| Q8BXV2 | BRI3B | BRI3-binding protein (I3-binding protein) | -0,166013974 | 0,020328 | 0,182129082 |
| Q91VR8 | BRK1 | Protein BRICK1 (BRK1) | 0,117237386 | 0,041408 | 0,259388001 |
| Q8BHG2 | CA123 | UPF0587 protein C1orf123 homolog | -0,678867558 | 0,000168 | 0,007894432 |
| O08532 | CA2D1 | Voltage-dependent calcium channel subunit alpha-2/delta-1 (Voltage-gated calcium channel subunit alpha-2/delta-1) [Cleaved into: Voltage-dependent calcium channel subunit alpha-2-1; Voltage-dependent calcium channel subunit delta-1] | -0,21740855 | 0,012418 | 0,13879024 |
| O08529 | CAN2 | Calpain-2 catalytic subunit (EC 3.4.22.53) (80 kDa M-calpain subunit) (CALP80) (Calcium-activated neutral proteinase 2) (CANP 2) (Calpain M-type) (Calpain-2 large subunit) (Millimolar-calpain) (M-calpain) | -0,408369143 | 0,000226 | 0,009612549 |
| O09161 | CASQ2 | Calsequestrin-2 (Calsequestrin, cardiac muscle isoform) | -0,126675027 | 0,036157 | 0,240254199 |
| Q9WVC3 | CAV2 | Caveolin-2 | -0,273504107 | 0,002818 | 0,059147505 |
| A2AMM0 | CAVN4 | Caveolae-associated protein 4 (Muscle-related coiled-coil protein) (Muscle-restricted coiled-coil protein) | -0,313233247 | 0,000212 | 0,009612549 |
| P47754 | CAZA2 | F-actin-capping protein subunit alpha-2 (CapZ alpha-2) | 0,114566575 | 0,046496 | 0,270714162 |
| P23198 | CBX3 | Chromobox protein homolog 3 (Heterochromatin protein 1 homolog gamma) (HP1 gamma) (M32) (Modifier 2 protein) | 0,191189309 | 0,005808 | 0,091573162 |
| Q8CH18 | CCAR1 | Cell division cycle and apoptosis regulator protein 1 (Cell cycle and apoptosis regulatory protein 1) (CARP-1) | 0,168521099 | 0,019978 | 0,180820079 |
| Q3URS9 | CCD51 | Coiled-coil domain-containing protein 51 | -0,190679626 | 0,013116 | 0,141705536 |
| Q8R349 | CDC16 | Cell division cycle protein 16 homolog (Anaphase-promoting complex subunit 6) (APC6) (Cyclosome subunit 6) | 0,143498554 | 0,022818 | 0,190682325 |
| Q6A068 | CDC5L | Cell division cycle 5-like protein (Cdc5-like protein) | 0,195240738 | 0,013371 | 0,143391231 |
| P28659 | CELF1 | CUGBP Elav-like family member 1 (CELF-1) (50 kDa nuclear polyadenylated RNA-binding protein) (Brain protein F41) (Bruno-like protein 2) (CUG triplet repeat RNA-binding protein 1) (CUG-BP1) (CUG-BP- and ETR-3-like factor 1) (Deadenylation factor CUG-BP) (Deadenylation factor EDEN-BP) (Embryo deadenylation element-binding protein homolog) (EDEN-BP homolog) (RNA-binding protein BRUNOL-2) | 0,200986467 | 0,009722 | 0,121718662 |
| Q9CXS4 | CENPV | Centromere protein V (CENP-V) (Proline-rich protein 6) | 0,182395392 | 0,042171 | 0,259606922 |
| Q8VCT4 | CES1D | Carboxylesterase 1D (Carboxylesterase 3) (EC 3.1.1.1) (EC 3.1.1.67) (Fatty acid ethyl ester synthase) (FAEE synthase) (Triacylglycerol hydrolase) (TGH) | -0,286865023 | 0,007312 | 0,105149698 |
| Q64433 | CH10 | 10 kDa heat shock protein, mitochondrial (Hsp10) (10 kDa chaperonin) (Chaperonin 10) (CPN10) | -0,158373863 | 0,02175 | 0,188260494 |
| Q9D1L0 | CHCH2 | Coiled-coil-helix-coiled-coil-helix domain-containing protein 2 | -0,317393583 | 0,002955 | 0,061140497 |
| Q6PDQ2 | CHD4 | Chromodomain-helicase-DNA-binding protein 4 (CHD-4) (EC 3.6.4.12) | 0,190806214 | 0,025475 | 0,203142138 |
| Q9WUD1 | CHIP | STIP1 homology and U box-containing protein 1 (EC 2.3.2.27) (Carboxy terminus of Hsp70-interacting protein) (E3 ubiquitin-protein ligase CHIP) (RING-type E3 ubiquitin transferase CHIP) | 0,192227418 | 0,008086 | 0,110740283 |
| D3Z7H8 | CILP2 | Cartilage intermediate layer protein 2 [Cleaved into: Cartilage intermediate layer protein 2 C1; Cartilage intermediate layer protein 2 C2] | -0,773982662 | 0,014341 | 0,149626304 |
| Q99LM2 | CK5P3 | CDK5 regulatory subunit-associated protein 3 | 0,122083686 | 0,041763 | 0,259388001 |
| Q8BRT1 | CLAP2 | CLIP-associating protein 2 (Cytoplasmic linker-associated protein 2) | 0,182966148 | 0,008079 | 0,110740283 |
| Q9QYB1 | CLIC4 | Chloride intracellular channel protein 4 (mc3s5/mtCLIC) | -0,184929814 | 0,014033 | 0,148307529 |
| Q8BXK9 | CLIC5 | Chloride intracellular channel protein 5 | -0,224411807 | 0,007873 | 0,109408805 |
| Q8R1G2 | CMBL | Carboxymethylenebutenolidase homolog (EC 3.1.-.-) | -0,129210847 | 0,047633 | 0,275137781 |
| Q8BIG7 | CMTD1 | Catechol O-methyltransferase domain-containing protein 1 (EC 2.1.1.-) | -0,310085441 | 0,00013 | 0,006520296 |
| Q08093 | CNN2 | Calponin-2 (Calponin H2, smooth muscle) (Neutral calponin) | -0,337110591 | 0,002644 | 0,05588513 |
| P02463 | CO4A1 | Collagen alpha-1(IV) chain [Cleaved into: Arresten] | -0,591566207 | 2,61E-05 | 0,002164174 |
| Q3U962 | CO5A2 | Collagen alpha-2(V) chain | 0,217075973 | 0,014142 | 0,148930139 |
| Q8C6K9 | CO6A6 | Collagen alpha-6(VI) chain | 0,241516452 | 0,001423 | 0,036090038 |
| Q921H9 | COA7 | Cytochrome c oxidase assembly factor 7 (Beta-lactamase hcp-like protein) (Respiratory chain assembly factor 1) (Sel1 repeat-containing protein 1) | -0,216401079 | 0,039613 | 0,25340093 |
| Q80X19 | COEA1 | Collagen alpha-1(XIV) chain | 0,437209888 | 0,024835 | 0,202353408 |
| O35206 | COFA1 | Collagen alpha-1(XV) chain [Cleaved into: Restin (Endostatin-XV)] | 0,169809079 | 0,042973 | 0,262537322 |
| Q8BMS4 | COQ3 | Ubiquinone biosynthesis O-methyltransferase, mitochondrial (3-demethylubiquinol 3-O-methyltransferase) (EC 2.1.1.64) (Polyprenyldihydroxybenzoate methyltransferase) (EC 2.1.1.114) | -0,238851286 | 0,003997 | 0,073636148 |
| Q9CXI0 | COQ5 | 2-methoxy-6-polyprenyl-1,4-benzoquinol methylase, mitochondrial (EC 2.1.1.201) (Ubiquinone biosynthesis methyltransferase COQ5) | -0,339919867 | 7,56E-05 | 0,004081112 |
| P56394 | COX17 | Cytochrome c oxidase copper chaperone | -0,858105549 | 9,43E-07 | 0,000250148 |
| Q8K0C8 | COX19 | Cytochrome c oxidase assembly protein COX19 | -0,340998536 | 0,000234 | 0,009612549 |
| Q9D7J4 | COX20 | Cytochrome c oxidase assembly protein COX20, mitochondrial | -0,282961678 | 0,03117 | 0,222849593 |
| Q9CPZ8 | COXM1 | COX assembly mitochondrial protein homolog (Cmc1p) | -0,484442623 | 2,75E-05 | 0,002164174 |
| Q8K199 | COXM2 | COX assembly mitochondrial protein 2 homolog | -0,17395313 | 0,012898 | 0,140385658 |
| Q6NVF9 | CPSF6 | Cleavage and polyadenylation specificity factor subunit 6 | 0,146031669 | 0,038403 | 0,247834295 |
| Q3TYS2 | CQ062 | Uncharacterized protein C17orf62 homolog | 0,154465304 | 0,037211 | 0,243371494 |
| Q9DCT8 | CRIP2 | Cysteine-rich protein 2 (CRP-2) (Heart LIM protein) | -0,406468887 | 1,30E-05 | 0,001453288 |
| P47941 | CRKL | Crk-like protein | 0,21328301 | 0,018643 | 0,173938529 |
| O88545 | CSN6 | COP9 signalosome complex subunit 6 (SGN6) (Signalosome subunit 6) (JAB1-containing signalosome subunit 6) | 0,237087169 | 0,007127 | 0,104466739 |
| Q8VHY0 | CSPG4 | Chondroitin sulfate proteoglycan 4 (Chondroitin sulfate proteoglycan NG2) (Proteoglycan AN2) | 0,352130491 | 0,000569 | 0,019520396 |
| P50462 | CSRP3 | Cysteine and glycine-rich protein 3 (Cysteine-rich protein 3) (CRP3) (LIM domain protein, cardiac) (Muscle LIM protein) | -0,279788462 | 0,000144 | 0,006879383 |
| Q8BY89 | CTL2 | Choline transporter-like protein 2 (Solute carrier family 44 member 2) | -0,600492061 | 2,95E-05 | 0,00220392 |
| P26231 | CTNA1 | Catenin alpha-1 (102 kDa cadherin-associated protein) (Alpha E-catenin) (CAP102) | -0,213099823 | 0,010521 | 0,124983812 |
| P56391 | CX6B1 | Cytochrome c oxidase subunit 6B1 (Cytochrome c oxidase subunit VIb isoform 1) (COX VIb-1) | -0,488371102 | 7,19E-05 | 0,004034167 |
| P56392 | CX7A1 | Cytochrome c oxidase subunit 7A1, mitochondrial (Cytochrome c oxidase subunit VIIa-heart) (Cytochrome c oxidase subunit VIIa-H) (Cytochrome c oxidase subunit VIIa-muscle) (Cytochrome c oxidase subunit VIIa-M) | -0,153802947 | 0,016713 | 0,164700744 |
| P48771 | CX7A2 | Cytochrome c oxidase subunit 7A2, mitochondrial (Cytochrome c oxidase subunit VIIa-liver/heart) (Cytochrome c oxidase subunit VIIa-L) | 0,145181261 | 0,04338 | 0,262537322 |
| P21460 | CYTC | Cystatin-C (Cystatin-3) | 0,20387547 | 0,014376 | 0,149626304 |
| Q5M8N4 | D39U1 | Epimerase family protein SDR39U1 (EC 1.1.1.-) (Short-chain dehydrogenase/reductase family 39U member 1) | -0,148754374 | 0,036435 | 0,240254199 |
| Q9JII5 | DAZP1 | DAZ-associated protein 1 (Deleted in azoospermia-associated protein 1) | 0,135888219 | 0,026883 | 0,210416436 |
| Q62418 | DBNL | Drebrin-like protein (Actin-binding protein 1) (SH3 domain-containing protein 7) | -0,36941546 | 0,000823 | 0,024051674 |
| Q8R1Q8 | DC1L1 | Cytoplasmic dynein 1 light intermediate chain 1 (Dynein light chain A) (DLC-A) (Dynein light intermediate chain 1, cytosolic) | -0,198874348 | 0,013298 | 0,143138526 |
| Q9DAR7 | DCPS | m7GpppX diphosphatase (EC 3.6.1.59) (DCS-1) (Decapping scavenger enzyme) (Hint-related 7meGMP-directed hydrolase) (Histidine triad nucleotide-binding protein 5) (Histidine triad protein member 5) (HINT-5) (Scavenger mRNA-decapping enzyme DcpS) | 0,186058408 | 0,018207 | 0,17344108 |
| Q99KJ8 | DCTN2 | Dynactin subunit 2 (50 kDa dynein-associated polypeptide) (Dynactin complex 50 kDa subunit) (DCTN-50) (Growth cone membrane protein 23-48K) (GMP23-48K) (p50 dynamitin) | -0,136960925 | 0,030164 | 0,219420978 |
| Q9CWS0 | DDAH1 | N(G),N(G)-dimethylarginine dimethylaminohydrolase 1 (DDAH-1) (Dimethylarginine dimethylaminohydrolase 1) (EC 3.5.3.18) (DDAHI) (Dimethylargininase-1) | -0,286442274 | 0,001165 | 0,032359119 |
| Q99LD8 | DDAH2 | N(G),N(G)-dimethylarginine dimethylaminohydrolase 2 (DDAH-2) (Dimethylarginine dimethylaminohydrolase 2) (EC 3.5.3.18) (DDAHII) (Dimethylargininase-2) | -0,373825844 | 0,000825 | 0,024051674 |
| Q62167 | DDX3X | ATP-dependent RNA helicase DDX3X (EC 3.6.4.13) (D1Pas1-related sequence 2) (DEAD box RNA helicase DEAD3) (mDEAD3) (DEAD box protein 3, X-chromosomal) (Embryonic RNA helicase) | 0,248088241 | 0,049054 | 0,278258066 |
| Q569Z5 | DDX46 | Probable ATP-dependent RNA helicase DDX46 (EC 3.6.4.13) (DEAD box protein 46) | 0,114291066 | 0,043823 | 0,262985417 |
| Q6Q899 | DDX58 | Probable ATP-dependent RNA helicase DDX58 (EC 3.6.4.13) (DEAD box protein 58) (RIG-I-like receptor 1) (RLR-1) (Retinoic acid-inducible gene 1 protein) (RIG-1) (Retinoic acid-inducible gene I protein) (RIG-I) | 0,250038602 | 0,04445 | 0,264869442 |
| Q7TNV0 | DEK | Protein DEK | 0,212517783 | 0,001626 | 0,040203758 |
| A2RT67 | DEND3 | DENN domain-containing protein 3 | 0,189444014 | 0,01462 | 0,151227565 |
| P50171 | DHB8 | Estradiol 17-beta-dehydrogenase 8 (EC 1.1.1.62) (17-beta-hydroxysteroid dehydrogenase 8) (17-beta-HSD 8) (3-ketoacyl-[acyl-carrier-protein] reductase alpha subunit) (KAR alpha subunit) (3-oxoacyl-[acyl-carrier-protein] reductase) (EC 1.1.1.-) (Protein Ke6) (Ke-6) (Testosterone 17-beta-dehydrogenase 8) (EC 1.1.1.239) | -0,203722376 | 0,004783 | 0,081535452 |
| Q99L04 | DHRS1 | Dehydrogenase/reductase SDR family member 1 (EC 1.1.-.-) | -0,251148457 | 0,001751 | 0,042208398 |
| Q99LB2 | DHRS4 | Dehydrogenase/reductase SDR family member 4 (EC 1.1.1.184) (NADPH-dependent carbonyl reductase/NADP-retinol dehydrogenase) (CR) (PHCR) (NADPH-dependent retinol dehydrogenase/reductase) (NDRD) (mouNRDR) (Peroxisomal short-chain alcohol dehydrogenase) (PSCD) | -0,440339484 | 1,27E-05 | 0,001453288 |
| Q9QYI3 | DNJC7 | DnaJ homolog subfamily C member 7 (Cytoplasmic CAR retention protein) (CCRP) (MDj11) (Tetratricopeptide repeat protein 2) (TPR repeat protein 2) | 0,208162477 | 0,038379 | 0,247834295 |
| Q8VDR9 | DOCK6 | Dedicator of cytokinesis protein 6 | 0,154918367 | 0,049453 | 0,279020789 |
| O08553 | DPYL2 | Dihydropyrimidinase-related protein 2 (DRP-2) (Unc-33-like phosphoprotein 2) (ULIP-2) | 0,127813067 | 0,049817 | 0,280531462 |
| Q62188 | DPYL3 | Dihydropyrimidinase-related protein 3 (DRP-3) (Unc-33-like phosphoprotein 1) (ULIP-1) | 0,142309576 | 0,032803 | 0,230034777 |
| P32233 | DRG1 | Developmentally-regulated GTP-binding protein 1 (DRG-1) (Neural precursor cell expressed developmentally down-regulated protein 3) (NEDD-3) | 0,153939554 | 0,03262 | 0,229840195 |
| Q7TSF1 | DSG1B | Desmoglein-1-beta (Dsg1-beta) (Desmoglein-5) | -1,110112798 | 0,000783 | 0,023299563 |
| P10126 | EF1A1 | Elongation factor 1-alpha 1 (EF-1-alpha-1) (Elongation factor Tu) (EF-Tu) (Eukaryotic elongation factor 1 A-1) (eEF1A-1) | 0,181584352 | 0,036627 | 0,240254199 |
| P62631 | EF1A2 | Elongation factor 1-alpha 2 (EF-1-alpha-2) (Eukaryotic elongation factor 1 A-2) (eEF1A-2) (Statin-S1) | -0,137512591 | 0,017688 | 0,169724042 |
| Q9DCH4 | EIF3F | Eukaryotic translation initiation factor 3 subunit F (eIF3f) (Deubiquitinating enzyme eIF3f) (EC 3.4.19.12) (Eukaryotic translation initiation factor 3 subunit 5) (eIF-3-epsilon) (eIF3 p47) | 0,164198247 | 0,013993 | 0,148307529 |
| Q99K41 | EMIL1 | EMILIN-1 (Elastin microfibril interface-located protein 1) (Elastin microfibril interfacer 1) | 0,228609661 | 0,007317 | 0,105149698 |
| P48193 | EPB41 | Protein 4.1 (P4.1) (4.1R) (Band 4.1) | 0,141792697 | 0,036111 | 0,240254199 |
| P84089 | ERH | Enhancer of rudimentary homolog (Mer) | 0,232110926 | 0,002898 | 0,060389411 |
| Q9R0P3 | ESTD | S-formylglutathione hydrolase (FGH) (EC 3.1.2.12) (Esterase 10) (Esterase D) (Sid 478) | -0,459968081 | 1,40E-05 | 0,001476951 |
| Q9DCM0 | ETHE1 | Persulfide dioxygenase ETHE1, mitochondrial (EC 1.13.11.18) (Ethylmalonic encephalopathy protein 1 homolog) (Hepatoma subtracted clone one protein) (Sulfur dioxygenase ETHE1) | -0,72682235 | 4,49E-07 | 0,000160491 |
| O35250 | EXOC7 | Exocyst complex component 7 (Exocyst complex component Exo70) | 0,13847393 | 0,046252 | 0,270374134 |
| Q9CRA8 | EXOS5 | Exosome complex component RRP46 (Exosome component 5) (Ribosomal RNA-processing protein 46) | 0,229137488 | 0,00585 | 0,091744717 |
| Q80VD1 | FA98B | Protein FAM98B | 0,149703304 | 0,03333 | 0,232037843 |
| Q05816 | FABP5 | Fatty acid-binding protein 5 (Epidermal-type fatty acid-binding protein) (E-FABP) (Fatty acid-binding protein, epidermal) (Keratinocyte lipid-binding protein) (Psoriasis-associated fatty acid-binding protein homolog) (PA-FABP) | -0,498434268 | 0,000971 | 0,028033803 |
| P12710 | FABPL | Fatty acid-binding protein, liver (14 kDa selenium-binding protein) (Fatty acid-binding protein 1) (Liver-type fatty acid-binding protein) (L-FABP) | -1,440838663 | 0,026416 | 0,208254519 |
| Q61160 | FADD | FAS-associated death domain protein (FAS-associating death domain-containing protein) (Mediator of receptor induced toxicity) (Protein FADD) | 0,369795525 | 0,043929 | 0,262985417 |
| Q3TC72 | FAHD2 | Fumarylacetoacetate hydrolase domain-containing protein 2A (EC 3.-.-.-) | -0,556596499 | 9,66E-06 | 0,001237666 |
| Q71FD7 | FBLI1 | Filamin-binding LIM protein 1 (FBLP-1) (CSX-associated LIM) | -0,364190012 | 7,02E-05 | 0,004034167 |
| Q61554 | FBN1 | Fibrillin-1 [Cleaved into: Asprosin] | -0,794464847 | 4,95E-07 | 0,000160491 |
| Q9QZN4 | FBX6 | F-box only protein 6 (F-box only protein 6b) (F-box protein that recognizes sugar chains 2) (F-box/G-domain protein 2) | 0,160168764 | 0,030101 | 0,219420978 |
| P29699 | FETUA | Alpha-2-HS-glycoprotein (Countertrypin) (Fetuin-A) | -0,535887634 | 0,005013 | 0,082622968 |
| O70433 | FHL2 | Four and a half LIM domains protein 2 (FHL-2) (Skeletal muscle LIM-protein 3) (SLIM-3) | -0,965185384 | 2,28E-06 | 0,000415795 |
| Q6P9Q4 | FHOD1 | FH1/FH2 domain-containing protein 1 (Formin homolog overexpressed in spleen 1) (FHOS) (Formin homology 2 domain-containing protein 1) | 0,138074893 | 0,029413 | 0,217809595 |
| P11276 | FINC | Fibronectin (FN) [Cleaved into: Anastellin] | -0,246173924 | 0,048719 | 0,27756345 |
| Q9D824 | FIP1 | Pre-mRNA 3'-end-processing factor FIP1 (FIP1-like 1 protein) | 0,193338841 | 0,00951 | 0,120629452 |
| P30416 | FKBP4 | Peptidyl-prolyl cis-trans isomerase FKBP4 (PPIase FKBP4) (EC 5.2.1.8) (52 kDa FK506-binding protein) (52 kDa FKBP) (FKBP-52) (59 kDa immunophilin) (p59) (FK506-binding protein 4) (FKBP-4) (FKBP59) (HSP-binding immunophilin) (HBI) (Immunophilin FKBP52) (Rotamase) [Cleaved into: Peptidyl-prolyl cis-trans isomerase FKBP4, N-terminally processed] | -0,170496185 | 0,029368 | 0,217809595 |
| Q9JJ28 | FLII | Protein flightless-1 homolog | 0,153315119 | 0,016992 | 0,165769436 |
| Q60634 | FLOT2 | Flotillin-2 (Epidermal surface antigen) (ESA) (Membrane component chromosome 17 surface marker 1 homolog) | 0,198779507 | 0,006341 | 0,096338404 |
| P29391 | FRIL1 | Ferritin light chain 1 (Ferritin L subunit 1) | 0,277735156 | 0,032926 | 0,230034777 |
| P56959 | FUS | RNA-binding protein FUS (Protein pigpen) | 0,208726242 | 0,004947 | 0,082459629 |
| Q00422 | GABPA | GA-binding protein alpha chain (GABP subunit alpha) | 0,165653465 | 0,007001 | 0,103137198 |
| P62880 | GBB2 | Guanine nucleotide-binding protein G(I)/G(S)/G(T) subunit beta-2 (G protein subunit beta-2) (Transducin beta chain 2) | -0,734052133 | 1,68E-06 | 0,000356395 |
| Q91WK5 | GCSH | Glycine cleavage system H protein, mitochondrial (Lipoic acid-containing protein) | -0,348697783 | 0,000685 | 0,021939435 |
| Q9D7X8 | GGCT | Gamma-glutamylcyclotransferase (EC 4.3.2.9) | 0,201325568 | 0,009614 | 0,120876193 |
| Q80Y14 | GLRX5 | Glutaredoxin-related protein 5, mitochondrial (Monothiol glutaredoxin-5) | -0,228580795 | 0,004064 | 0,073636148 |
| Q99L27 | GMPR2 | GMP reductase 2 (GMPR 2) (EC 1.7.1.7) (Guanosine 5'-monophosphate oxidoreductase 2) (Guanosine monophosphate reductase 2) | 0,243191375 | 0,046389 | 0,270632724 |
| P27601 | GNA13 | Guanine nucleotide-binding protein subunit alpha-13 (G alpha-13) (G-protein subunit alpha-13) | 0,211029016 | 0,005323 | 0,086935286 |
| Q6R0H7 | GNAS1 | Guanine nucleotide-binding protein G(s) subunit alpha isoforms XLas (Adenylate cyclase-stimulating G alpha protein) (Extra large alphas protein) (XLalphas) | 0,139164947 | 0,035741 | 0,240254199 |
| Q921M4 | GOGA2 | Golgin subfamily A member 2 (130 kDa cis-Golgi matrix protein) (GM130) | 0,138372533 | 0,011793 | 0,133338868 |
| Q9QYE6 | GOGA5 | Golgin subfamily A member 5 (Golgin-84) (Protein Ret-II) (Protein Sumiko) | 0,292754769 | 0,020999 | 0,185617161 |
| Q9QZF2 | GPC1 | Glypican-1 [Cleaved into: Secreted glypican-1] | -0,478386881 | 4,44E-05 | 0,003014374 |
| Q3ULJ0 | GPD1L | Glycerol-3-phosphate dehydrogenase 1-like protein (EC 1.1.1.8) | 0,186751714 | 0,041794 | 0,259388001 |
| Q9CXY9 | GPI8 | GPI-anchor transamidase (GPI transamidase) (EC 3.-.-.-) (Phosphatidylinositol-glycan biosynthesis class K protein) (PIG-K) | 0,171263914 | 0,009446 | 0,120629452 |
| P46412 | GPX3 | Glutathione peroxidase 3 (GPx-3) (GSHPx-3) (EC 1.11.1.9) (Plasma glutathione peroxidase) (GPx-P) (GSHPx-P) | 0,29218738 | 0,010927 | 0,126984152 |
| O70325 | GPX4 | Phospholipid hydroperoxide glutathione peroxidase (PHGPx) (EC 1.11.1.12) (Glutathione peroxidase 4) (GPx-4) (GSHPx-4) | -0,657715489 | 2,82E-05 | 0,002164174 |
| Q91Z53 | GRHPR | Glyoxylate reductase/hydroxypyruvate reductase (EC 1.1.1.79) (EC 1.1.1.81) | -0,170633617 | 0,027534 | 0,212119635 |
| P38647 | GRP75 | Stress-70 protein, mitochondrial (75 kDa glucose-regulated protein) (GRP-75) (Heat shock 70 kDa protein 9) (Mortalin) (Peptide-binding protein 74) (PBP74) (p66 MOT) | -0,117112508 | 0,027105 | 0,210843564 |
| Q9EST1 | GSDMA | Gasdermin-A (Gasdermin-1) (Gasdermin-A1) | -1,301106837 | 0,001965 | 0,046605644 |
| Q61543 | GSLG1 | Golgi apparatus protein 1 (E-selectin ligand 1) (ESL-1) (Selel) (Golgi sialoglycoprotein MG-160) | -0,388410547 | 9,76E-06 | 0,001237666 |
| O08582 | GTPB1 | GTP-binding protein 1 (G-protein 1) (GP-1) (GP1) | 0,151021342 | 0,035023 | 0,239230719 |
| P10922 | H10 | Histone H1.0 (Histone H1') (Histone H1(0)) (MyD196) [Cleaved into: Histone H1.0, N-terminally processed] | 0,221780651 | 0,028672 | 0,216117407 |
| P43275 | H11 | Histone H1.1 (H1 VAR.3) (Histone H1a) (H1a) | 0,201249508 | 0,00787 | 0,109408805 |
| P15864 | H12 | Histone H1.2 (H1 VAR.1) (H1c) | 0,190330135 | 0,007654 | 0,108912825 |
| P43274 | H14 | Histone H1.4 (H1 VAR.2) (H1e) | 0,173572726 | 0,026949 | 0,210416436 |
| P43276 | H15 | Histone H1.5 (H1 VAR.5) (H1b) | 0,229865462 | 0,002303 | 0,049825204 |
| Q8CCK0 | H2AW | Core histone macro-H2A.2 (Histone macroH2A2) (mH2A2) | 0,159229631 | 0,041783 | 0,259388001 |
| Q9QZQ8 | H2AY | Core histone macro-H2A.1 (Histone macroH2A1) (mH2A1) (H2A.y) (H2A/y) | 0,243848686 | 0,000617 | 0,020931908 |
| P0C0S6 | H2AZ | Histone H2A.Z (H2A/z) | 0,325696476 | 0,004259 | 0,075457482 |
| P62806 | H4 | Histone H4 | 0,245362301 | 0,002266 | 0,049825204 |
| Q3UHX2 | HAP28 | 28 kDa heat- and acid-stable phosphoprotein (PDGF-associated protein) (PAP) (PDGFA-associated protein 1) (PAP1) | 0,14584623 | 0,016285 | 0,16157311 |
| O08756 | HCD2 | 3-hydroxyacyl-CoA dehydrogenase type-2 (EC 1.1.1.35) (17-beta-hydroxysteroid dehydrogenase 10) (17-beta-HSD 10) (EC 1.1.1.51) (3-hydroxy-2-methylbutyryl-CoA dehydrogenase) (EC 1.1.1.178) (3-hydroxyacyl-CoA dehydrogenase type II) (Endoplasmic reticulum-associated amyloid beta-peptide-binding protein) (Mitochondrial ribonuclease P protein 2) (Mitochondrial RNase P protein 2) (Type II HADH) | -0,22239625 | 0,001043 | 0,029817739 |
| P51859 | HDGF | Hepatoma-derived growth factor (HDGF) | 0,240402128 | 0,004902 | 0,082178485 |
| Q3UMU9 | HDGR2 | Hepatoma-derived growth factor-related protein 2 (HRP-2) | 0,181733945 | 0,034962 | 0,239230719 |
| P22315 | HEMH | Ferrochelatase, mitochondrial (EC 4.99.1.1) (Heme synthase) (Protoheme ferro-lyase) | -0,205363314 | 0,039977 | 0,254775562 |
| Q8QZS1 | HIBCH | 3-hydroxyisobutyryl-CoA hydrolase, mitochondrial (EC 3.1.2.4) (3-hydroxyisobutyryl-coenzyme A hydrolase) (HIB-CoA hydrolase) (HIBYL-CoA-H) | -0,239631142 | 0,001101 | 0,030883083 |
| P17095 | HMGA1 | High mobility group protein HMG-I/HMG-Y (HMG-I(Y)) (High mobility group AT-hook protein 1) (High mobility group protein A1) | 0,375127582 | 0,048085 | 0,275606009 |
| P63158 | HMGB1 | High mobility group protein B1 (High mobility group protein 1) (HMG-1) | 0,317566129 | 0,001273 | 0,033751711 |
| O54879 | HMGB3 | High mobility group protein B3 (High mobility group protein 2a) (HMG-2a) (High mobility group protein 4) (HMG-4) | 0,30095374 | 0,015673 | 0,15819026 |
| O35737 | HNRH1 | Heterogeneous nuclear ribonucleoprotein H (hnRNP H) [Cleaved into: Heterogeneous nuclear ribonucleoprotein H, N-terminally processed] | 0,190783498 | 0,005532 | 0,089157901 |
| P70333 | HNRH2 | Heterogeneous nuclear ribonucleoprotein H2 (hnRNP H2) (Heterogeneous nuclear ribonucleoprotein H') (hnRNP H') [Cleaved into: Heterogeneous nuclear ribonucleoprotein H2, N-terminally processed] | 0,146937593 | 0,043184 | 0,262537322 |
| Q8VDM6 | HNRL1 | Heterogeneous nuclear ribonucleoprotein U-like protein 1 | -0,490180395 | 1,42E-05 | 0,001476951 |
| Q921F4 | HNRLL | Heterogeneous nuclear ribonucleoprotein L-like | 0,206859948 | 0,046149 | 0,27031646 |
| Q60668 | HNRPD | Heterogeneous nuclear ribonucleoprotein D0 (hnRNP D0) (AU-rich element RNA-binding protein 1) | 0,148321714 | 0,023776 | 0,196218478 |
| Q8R081 | HNRPL | Heterogeneous nuclear ribonucleoprotein L (hnRNP L) | 0,240188181 | 0,007354 | 0,105149698 |
| P00493 | HPRT | Hypoxanthine-guanine phosphoribosyltransferase (HGPRT) (HGPRTase) (EC 2.4.2.8) (HPRT B) | 0,267656898 | 0,004808 | 0,081535452 |
| Q8BTX9 | HSDL1 | Inactive hydroxysteroid dehydrogenase-like protein 1 | 0,217139539 | 0,048677 | 0,27756345 |
| Q2TPA8 | HSDL2 | Hydroxysteroid dehydrogenase-like protein 2 (EC 1.-.-.-) | -0,124585551 | 0,022854 | 0,190682325 |
| Q99PR8 | HSPB2 | Heat shock protein beta-2 (HspB2) | -0,138609888 | 0,016976 | 0,165769436 |
| P35385 | HSPB7 | Heat shock protein beta-7 (HspB7) (Cardiovascular heat shock protein) (cvHsp) (Heat shock protein 25 kDa 2) (Protein p19/6.8) | -0,233213455 | 0,02048 | 0,182694444 |
| Q9D379 | HYEP | Epoxide hydrolase 1 (EC 3.3.2.9) (Epoxide hydratase) (Microsomal epoxide hydrolase) (mEH) | 0,212439472 | 0,031693 | 0,224936257 |
| Q9D6R2 | IDH3A | Isocitrate dehydrogenase [NAD] subunit alpha, mitochondrial (EC 1.1.1.41) (Isocitric dehydrogenase subunit alpha) (NAD(+)-specific ICDH subunit alpha) | -0,207751635 | 0,011702 | 0,132822946 |
| O88844 | IDHC | Isocitrate dehydrogenase [NADP] cytoplasmic (IDH) (EC 1.1.1.42) (Cytosolic NADP-isocitrate dehydrogenase) (IDP) (NADP(+)-specific ICDH) (Oxalosuccinate decarboxylase) | -0,295778252 | 0,0456 | 0,268696862 |
| Q91YJ5 | IF2M | Translation initiation factor IF-2, mitochondrial (IF-2(Mt)) (IF-2Mt) (IF2(mt)) | -0,140684955 | 0,036534 | 0,240254199 |
| Q91VC3 | IF4A3 | Eukaryotic initiation factor 4A-III (eIF-4A-III) (eIF4A-III) (EC 3.6.4.13) (ATP-dependent RNA helicase DDX48) (ATP-dependent RNA helicase eIF4A-3) (DEAD box protein 48) (Eukaryotic translation initiation factor 4A isoform 3) [Cleaved into: Eukaryotic initiation factor 4A-III, N-terminally processed] | 0,158505804 | 0,02942 | 0,217809595 |
| P59325 | IF5 | Eukaryotic translation initiation factor 5 (eIF-5) | 0,144033386 | 0,049127 | 0,278258066 |
| O55222 | ILK | Integrin-linked protein kinase (EC 2.7.11.1) | 0,144912268 | 0,045238 | 0,267667692 |
| Q91UZ5 | IMPA2 | Inositol monophosphatase 2 (IMP 2) (IMPase 2) (EC 3.1.3.25) (Inositol-1(or 4)-monophosphatase 2) (Myo-inositol monophosphatase A2) | 0,284004845 | 0,009129 | 0,118349458 |
| P85094 | ISC2A | Isochorismatase domain-containing protein 2A | -1,036886206 | 1,22E-06 | 0,000297476 |
| Q9DCB8 | ISCA2 | Iron-sulfur cluster assembly 2 homolog, mitochondrial (HESB-like domain-containing protein 1) | -0,186164846 | 0,002298 | 0,049825204 |
| Q61739 | ITA6 | Integrin alpha-6 (CD49 antigen-like family member F) (VLA-6) (CD antigen CD49f) [Cleaved into: Integrin alpha-6 heavy chain; Integrin alpha-6 light chain] | 0,206203568 | 0,035691 | 0,240254199 |
| Q9JHI5 | IVD | Isovaleryl-CoA dehydrogenase, mitochondrial (IVD) (EC 1.3.8.4) | -0,189994105 | 0,010754 | 0,125481426 |
| Q8C1D8 | IWS1 | Protein IWS1 homolog (IWS1-like protein) | 0,177894687 | 0,028089 | 0,212271439 |
| Q9WUR9 | KAD4 | Adenylate kinase 4, mitochondrial (AK 4) (EC 2.7.4.10) (EC 2.7.4.6) (Adenylate kinase 3-like) (Adenylate kinase isoenzyme 4) (GTP:AMP phosphotransferase AK4) | -0,145605343 | 0,036652 | 0,240254199 |
| O88986 | KBL | 2-amino-3-ketobutyrate coenzyme A ligase, mitochondrial (AKB ligase) (EC 2.3.1.29) (Aminoacetone synthase) (Glycine acetyltransferase) | -0,310031999 | 0,000516 | 0,018058841 |
| Q6PHZ2 | KCC2D | Calcium/calmodulin-dependent protein kinase type II subunit delta (CaM kinase II subunit delta) (CaMK-II subunit delta) (EC 2.7.11.17) | -0,291452457 | 6,64E-05 | 0,004034167 |
| Q923T9 | KCC2G | Calcium/calmodulin-dependent protein kinase type II subunit gamma (CaM kinase II subunit gamma) (CaMK-II subunit gamma) (EC 2.7.11.17) | -0,295203759 | 0,00215 | 0,048623161 |
| Q9DBP5 | KCY | UMP-CMP kinase (EC 2.7.4.14) (Deoxycytidylate kinase) (CK) (dCMP kinase) (Nucleoside-diphosphate kinase) (EC 2.7.4.6) (Uridine monophosphate/cytidine monophosphate kinase) (UMP/CMP kinase) (UMP/CMPK) | -0,190300059 | 0,022759 | 0,190682325 |
| O35071 | KIF1C | Kinesin-like protein KIF1C | 0,15790345 | 0,017863 | 0,170843438 |
| Q61768 | KINH | Kinesin-1 heavy chain (Conventional kinesin heavy chain) (Ubiquitous kinesin heavy chain) (UKHC) | -0,195684514 | 0,025232 | 0,203142138 |
| A2AUC9 | KLH41 | Kelch-like protein 41 (Kelch repeat and BTB domain-containing protein 10) | -0,55961326 | 1,15E-05 | 0,00139473 |
| P32067 | LA | Lupus La protein homolog (La autoantigen homolog) (La ribonucleoprotein) | 0,183410059 | 0,010312 | 0,123669585 |
| Q61033 | LAP2A | Lamina-associated polypeptide 2, isoforms alpha/zeta (Thymopoietin isoforms alpha/zeta) (TP alpha/zeta) | 0,172670963 | 0,009499 | 0,120629452 |
| Q61029 | LAP2B | Lamina-associated polypeptide 2, isoforms beta/delta/epsilon/gamma (Thymopoietin isoforms beta/delta/epsilon/gamma) (TP beta/delta/epsilon/gamma) | 0,313007178 | 0,007742 | 0,109408805 |
| Q8C129 | LCAP | Leucyl-cystinyl aminopeptidase (Cystinyl aminopeptidase) (EC 3.4.11.3) (Oxytocinase) (OTase) | -0,320133863 | 0,002188 | 0,048714388 |
| Q3UN02 | LCLT1 | Lysocardiolipin acyltransferase 1 (EC 2.3.1.-) (EC 2.3.1.51) (Acyl-CoA:lysocardiolipin acyltransferase 1) | 0,150711142 | 0,029368 | 0,217809595 |
| P16045 | LEG1 | Galectin-1 (Gal-1) (14 kDa lectin) (Beta-galactoside-binding lectin L-14-I) (Galaptin) (Lactose-binding lectin 1) (Lectin galactoside-binding soluble 1) (S-Lac lectin 1) | -0,197964836 | 0,006317 | 0,096338404 |
| Q6DVA0 | LEMD2 | LEM domain-containing protein 2 (Nuclear envelope transmembrane protein 25) (NET25) | 0,58515181 | 2,67E-05 | 0,002164174 |
| Q91YX5 | LGAT1 | Acyl-CoA:lysophosphatidylglycerol acyltransferase 1 (EC 2.3.1.-) | -0,14325951 | 0,040218 | 0,255035879 |
| Q9ERG0 | LIMA1 | LIM domain and actin-binding protein 1 (Epithelial protein lost in neoplasm) (mEPLIN) | 0,223288968 | 0,008187 | 0,111599897 |
| P54310 | LIPS | Hormone-sensitive lipase (HSL) (EC 3.1.1.79) | 0,291911309 | 0,008512 | 0,114948638 |
| P48678 | LMNA | Prelamin-A/C [Cleaved into: Lamin-A/C] | 0,47586813 | 6,92E-05 | 0,004034167 |
| P21619 | LMNB2 | Lamin-B2 | -0,347676151 | 0,000504 | 0,017933802 |
| P48999 | LOX5 | Arachidonate 5-lipoxygenase (5-LO) (5-lipoxygenase) (EC 1.13.11.34) | -0,232055164 | 0,007787 | 0,109408805 |
| Q6PB66 | LPPRC | Leucine-rich PPR motif-containing protein, mitochondrial (130 kDa leucine-rich protein) (LRP 130) (mLRP130) | -0,144482455 | 0,015591 | 0,157917896 |
| Q8K3W2 | LRC10 | Leucine-rich repeat-containing protein 10 (Heart-restricted leucine-rich repeat protein) | -0,687292919 | 8,96E-08 | 8,71E-05 |
| Q9QXA5 | LSM4 | U6 snRNA-associated Sm-like protein LSm4 | 0,197457249 | 0,042074 | 0,259606922 |
| Q6ZWM4 | LSM8 | U6 snRNA-associated Sm-like protein LSm8 | 0,216129649 | 0,001269 | 0,033751711 |
| Q9CQ22 | LTOR1 | Ragulator complex protein LAMTOR1 (Late endosomal/lysosomal adaptor and MAPK and MTOR activator 1) (Lipid raft adaptor protein p18) | -0,120760472 | 0,025178 | 0,203142138 |
| P70699 | LYAG | Lysosomal alpha-glucosidase (EC 3.2.1.20) (Acid maltase) | -0,432590328 | 0,000671 | 0,021753075 |
| Q9WTL7 | LYPA2 | Acyl-protein thioesterase 2 (APT-2) (EC 3.1.2.-) (Lysophospholipase 2) (Lysophospholipase II) (LPL-II) (LysoPLA II) (mLyso II) | 0,15698537 | 0,029041 | 0,217809595 |
| O35099 | M3K5 | Mitogen-activated protein kinase kinase kinase 5 (EC 2.7.11.25) (Apoptosis signal-regulating kinase 1) (ASK-1) (MAPK/ERK kinase kinase 5) (MEK kinase 5) (MEKK 5) | -0,171058199 | 0,022748 | 0,190682325 |
| A2AJI0 | MA7D1 | MAP7 domain-containing protein 1 | -0,355936236 | 7,78E-05 | 0,004127172 |
| Q922B1 | MACD1 | O-acetyl-ADP-ribose deacetylase MACROD1 (EC 3.2.2.-) (EC 3.5.1.-) (MACRO domain-containing protein 1) (Protein LRP16) ([Protein ADP-ribosylglutamate] hydrolase) | -0,299515858 | 0,001713 | 0,041634801 |
| Q9CXI5 | MANF | Mesencephalic astrocyte-derived neurotrophic factor (Arginine-rich protein) (Protein ARMET) | -0,45258147 | 0,001482 | 0,037274322 |
| Q9D1M4 | MCA3 | Eukaryotic translation elongation factor 1 epsilon-1 (Elongation factor p18) (Multisynthase complex auxiliary component p18) | -0,181564028 | 0,010003 | 0,122276261 |
| P08249 | MDHM | Malate dehydrogenase, mitochondrial (EC 1.1.1.37) | -0,145991845 | 0,015328 | 0,156339704 |
| O35678 | MGLL | Monoglyceride lipase (MGL) (EC 3.1.1.23) (Monoacylglycerol lipase) (MAGL) | -0,768847619 | 2,34E-07 | 0,000113926 |
| P61327 | MGN | Protein mago nashi homolog | 0,266318343 | 0,003353 | 0,064413722 |
| Q7TNS2 | MIC10 | MICOS complex subunit Mic10 (Mitochondrial inner membrane organizing system protein 1) | -0,435009801 | 0,000268 | 0,010545073 |
| Q8VDP3 | MICA1 | [F-actin]-monooxygenase MICAL1 (EC 1.14.13.225) (Molecule interacting with CasL protein 1) (MICAL-1) (mMical1) (NEDD9-interacting protein with calponin homology and LIM domains) | 0,184022074 | 0,040755 | 0,256765664 |
| Q9CQ86 | MIEN1 | Migration and invasion enhancer 1 | -0,16947695 | 0,019994 | 0,180820079 |
| Q8BG51 | MIRO1 | Mitochondrial Rho GTPase 1 (MIRO-1) (EC 3.6.5.-) (Ras homolog gene family member T1) | -0,414715552 | 6,00E-05 | 0,003827295 |
| Q3THE2 | ML12B | Myosin regulatory light chain 12B (Myosin regulatory light chain 2-B, smooth muscle isoform) (Myosin regulatory light chain 20 kDa) (MLC20) (Myosin regulatory light chain MRLC2) | 0,187819958 | 0,020355 | 0,182129082 |
| Q8C7H1 | MMAA | Methylmalonic aciduria type A homolog, mitochondrial (EC 3.6.-.-) | -0,249925873 | 0,007916 | 0,109439802 |
| Q9D273 | MMAB | Cob(I)yrinic acid a,c-diamide adenosyltransferase, mitochondrial (EC 2.5.1.17) (Cob(I)alamin adenosyltransferase) (Methylmalonic aciduria type B homolog) | -0,244103375 | 0,009879 | 0,122276261 |
| P26041 | MOES | Moesin (Membrane-organizing extension spike protein) | 0,160118965 | 0,021695 | 0,188260494 |
| Q80UM7 | MOGS | Mannosyl-oligosaccharide glucosidase (EC 3.2.1.106) (Glucosidase 1) (Glycoprotein-processing glucosidase I) | 0,200798767 | 0,004844 | 0,081673835 |
| Q9D023 | MPC2 | Mitochondrial pyruvate carrier 2 (Brain protein 44) | -0,169701341 | 0,011392 | 0,13031752 |
| Q920Q6 | MSI2H | RNA-binding protein Musashi homolog 2 (Musashi-2) | 0,217369631 | 0,00208 | 0,047975731 |
| Q78J03 | MSRB2 | Methionine-R-sulfoxide reductase B2, mitochondrial (MsrB2) (EC 1.8.4.12) (EC 1.8.4.14) | -0,172102941 | 0,013462 | 0,143837957 |
| Q8BKY8 | MTEF2 | Transcription termination factor 2, mitochondrial (Mitochondrial transcription termination factor 2) (mTERF2) (mTERF domain-containing protein 3, mitochondrial) | -0,167675207 | 0,018605 | 0,173938529 |
| Q9CRB8 | MTFP1 | Mitochondrial fission process protein 1 (Mitochondrial 18 kDa protein) (MTP18) | -0,122986271 | 0,039398 | 0,252578066 |
| Q9DCS2 | MTL26 | Methyltransferase-like 26 | -0,196895289 | 0,018307 | 0,17344108 |
| Q9CZU3 | MTREX | Exosome RNA helicase MTR4 (EC 3.6.4.13) (ATP-dependent helicase SKIV2L2) (Superkiller viralicidic activity 2-like 2) (TRAMP-like complex helicase) | 0,272933888 | 0,000663 | 0,021753075 |
| O88441 | MTX2 | Metaxin-2 (Mitochondrial outer membrane import complex protein 2) | -0,176920959 | 0,032583 | 0,229840195 |
| O35682 | MYADM | Myeloid-associated differentiation marker (Myeloid up-regulated protein) | -0,338015172 | 0,00035 | 0,012944646 |
| P04247 | MYG | Myoglobin | 0,33866481 | 0,010009 | 0,122276261 |
| Q9JK81 | MYG1 | UPF0160 protein MYG1, mitochondrial (Protein Gamm1) | 0,283724816 | 5,70E-05 | 0,003777201 |
| Q9DB73 | NB5R1 | NADH-cytochrome b5 reductase 1 (b5R.1) (EC 1.6.2.2) (NAD(P)H:quinone oxidoreductase type 3 polypeptide A2) | -0,473755503 | 4,09E-05 | 0,002837649 |
| Q9WV85 | NDK3 | Nucleoside diphosphate kinase 3 (NDK 3) (NDP kinase 3) (EC 2.7.4.6) (DR-nm23) (Nucleoside diphosphate kinase C) (NDPKC) (nm23-M3) | -0,15335988 | 0,010284 | 0,123669585 |
| Q9QYG0 | NDRG2 | Protein NDRG2 (N-myc downstream-regulated gene 2 protein) (Protein Ndr2) | -0,157862743 | 0,030548 | 0,220566508 |
| Q9CPP6 | NDUA5 | NADH dehydrogenase [ubiquinone] 1 alpha subcomplex subunit 5 (Complex I subunit B13) (Complex I-13kD-B) (CI-13kD-B) (NADH-ubiquinone oxidoreductase 13 kDa-B subunit) | 0,224512205 | 0,036026 | 0,240254199 |
| Q9DCJ5 | NDUA8 | NADH dehydrogenase [ubiquinone] 1 alpha subcomplex subunit 8 (Complex I-19kD) (CI-19kD) (Complex I-PGIV) (CI-PGIV) (NADH-ubiquinone oxidoreductase 19 kDa subunit) | -0,585911265 | 2,05E-07 | 0,000113926 |
| Q9DC69 | NDUA9 | NADH dehydrogenase [ubiquinone] 1 alpha subcomplex subunit 9, mitochondrial (Complex I-39kD) (CI-39kD) (NADH-ubiquinone oxidoreductase 39 kDa subunit) | -0,186503685 | 0,004646 | 0,079711554 |
| Q9D8B4 | NDUAB | NADH dehydrogenase [ubiquinone] 1 alpha subcomplex subunit 11 (Complex I-B14.7) (CI-B14.7) (NADH-ubiquinone oxidoreductase subunit B14.7) | -0,328776443 | 0,000755 | 0,022952581 |
| Q7TMF3 | NDUAC | NADH dehydrogenase [ubiquinone] 1 alpha subcomplex subunit 12 (Complex I-B17.2) (CI-B17.2) (CIB17.2) (NADH-ubiquinone oxidoreductase subunit B17.2) | -0,410664055 | 0,000237 | 0,009612549 |
| Q9DCS9 | NDUBA | NADH dehydrogenase [ubiquinone] 1 beta subcomplex subunit 10 (Complex I-PDSW) (CI-PDSW) (NADH-ubiquinone oxidoreductase PDSW subunit) | -0,227411809 | 0,003746 | 0,07005043 |
| O09111 | NDUBB | NADH dehydrogenase [ubiquinone] 1 beta subcomplex subunit 11, mitochondrial (Complex I-ESSS) (CI-ESSS) (NADH-ubiquinone oxidoreductase ESSS subunit) (Neuronal protein 15.6) (Np15.6) (p15.6) | -0,187232484 | 0,016033 | 0,160711106 |
| Q9CWG8 | NDUF7 | Protein arginine methyltransferase NDUFAF7, mitochondrial (EC 2.1.1.320) (NADH dehydrogenase [ubiquinone] complex I, assembly factor 7) (Protein midA homolog) | -0,130490678 | 0,011143 | 0,12891725 |
| Q99LY9 | NDUS5 | NADH dehydrogenase [ubiquinone] iron-sulfur protein 5 (Complex I-15 kDa) (CI-15 kDa) (NADH-ubiquinone oxidoreductase 15 kDa subunit) | -0,812441556 | 8,75E-08 | 8,71E-05 |
| P52503 | NDUS6 | NADH dehydrogenase [ubiquinone] iron-sulfur protein 6, mitochondrial (Complex I-13kD-A) (CI-13kD-A) (NADH-ubiquinone oxidoreductase 13 kDa-A subunit) | -0,139124929 | 0,031295 | 0,223194544 |
| Q91YT0 | NDUV1 | NADH dehydrogenase [ubiquinone] flavoprotein 1, mitochondrial (EC 1.6.5.3) (EC 1.6.99.3) (Complex I-51kD) (CI-51kD) (NADH-ubiquinone oxidoreductase 51 kDa subunit) | -0,426928602 | 2,59E-05 | 0,002164174 |
| Q6P5H2 | NEST | Nestin | -0,334552276 | 0,00026 | 0,010386609 |
| Q02780 | NFIA | Nuclear factor 1 A-type (NF1-A) (Nuclear factor 1/A) (CCAAT-box-binding transcription factor) (CTF) (Nuclear factor I/A) (NF-I/A) (NFI-A) (TGGCA-binding protein) | 0,21751154 | 0,009448 | 0,120629452 |
| Q9QZ23 | NFU1 | NFU1 iron-sulfur cluster scaffold homolog, mitochondrial (HIRA-interacting protein 5) (mHIRIP5) | -0,208526112 | 0,016698 | 0,164700744 |
| P10493 | NID1 | Nidogen-1 (NID-1) (Entactin) | -0,145998959 | 0,037829 | 0,245759639 |
| Q3TL44 | NLRX1 | NLR family member X1 | -0,569731306 | 3,78E-06 | 0,000648823 |
| Q99JR6 | NMNA3 | Nicotinamide/nicotinic acid mononucleotide adenylyltransferase 3 (NMN/NaMN adenylyltransferase 3) (EC 2.7.7.1) (EC 2.7.7.18) (Nicotinamide-nucleotide adenylyltransferase 3) (NMN adenylyltransferase 3) (Nicotinate-nucleotide adenylyltransferase 3) (NaMN adenylyltransferase 3) | -0,2016797 | 0,001833 | 0,04383843 |
| Q9CZ42 | NNRD | ATP-dependent (S)-NAD(P)H-hydrate dehydratase (EC 4.2.1.93) (ATP-dependent NAD(P)HX dehydratase) (Carbohydrate kinase domain-containing protein) (NAD(P)HX dehydratase) | -0,205597208 | 0,012367 | 0,138747286 |
| Q99K48 | NONO | Non-POU domain-containing octamer-binding protein (NonO protein) | 0,155780485 | 0,034488 | 0,238469099 |
| P60670 | NPL4 | Nuclear protein localization protein 4 homolog (Protein NPL4) | -0,50896936 | 8,74E-05 | 0,004554428 |
| Q64669 | NQO1 | NAD(P)H dehydrogenase [quinone] 1 (EC 1.6.5.2) (Azoreductase) (DT-diaphorase) (DTD) (Menadione reductase) (NAD(P)H:quinone oxidoreductase 1) (Phylloquinone reductase) (Quinone reductase 1) (QR1) | 0,251375857 | 0,006068 | 0,094144615 |
| O70572 | NSMA | Sphingomyelin phosphodiesterase 2 (EC 3.1.4.12) (Lyso-platelet-activating factor-phospholipase C) (Lyso-PAF-PLC) (Neutral sphingomyelinase) (N-SMase) (nSMase) | 0,124850849 | 0,044811 | 0,266221741 |
| Q1HFZ0 | NSUN2 | tRNA (cytosine(34)-C(5))-methyltransferase (EC 2.1.1.203) (Myc-induced SUN domain-containing protein) (Misu) (NOL1/NOP2/Sun domain family member 2) | 0,142833495 | 0,030488 | 0,220566508 |
| Q8VCE6 | NT5M | 5'(3')-deoxyribonucleotidase, mitochondrial (5',3'-nucleotidase, mitochondrial) (EC 3.1.3.-) (Deoxy-5'-nucleotidase 2) (dNT-2) | -0,133830907 | 0,040002 | 0,254775562 |
| Q9CWD8 | NUBPL | Iron-sulfur protein NUBPL (Nucleotide-binding protein-like) | -0,130130977 | 0,046128 | 0,27031646 |
| Q9DCN1 | NUD12 | Peroxisomal NADH pyrophosphatase NUDT12 (EC 3.6.1.22) (Nucleoside diphosphate-linked moiety X motif 12) (Nudix motif 12) | -0,164604808 | 0,043776 | 0,262985417 |
| Q9CR24 | NUDT8 | Nucleoside diphosphate-linked moiety X motif 8 (Nudix motif 8) (EC 3.6.1.-) | -0,378763511 | 0,000464 | 0,016924523 |
| P29758 | OAT | Ornithine aminotransferase, mitochondrial (EC 2.6.1.13) (Ornithine--oxo-acid aminotransferase) | -0,570029753 | 0,000351 | 0,012944646 |
| Q6P3A8 | ODBB | 2-oxoisovalerate dehydrogenase subunit beta, mitochondrial (EC 1.2.4.4) (Branched-chain alpha-keto acid dehydrogenase E1 component beta chain) (BCKDE1B) (BCKDH E1-beta) | -0,686297931 | 1,79E-05 | 0,001735881 |
| Q60597 | ODO1 | 2-oxoglutarate dehydrogenase, mitochondrial (EC 1.2.4.2) (2-oxoglutarate dehydrogenase complex component E1) (OGDC-E1) (Alpha-ketoglutarate dehydrogenase) | -0,225667924 | 0,003182 | 0,063752636 |
| P35486 | ODPA | Pyruvate dehydrogenase E1 component subunit alpha, somatic form, mitochondrial (EC 1.2.4.1) (PDHE1-A type I) | -0,145422707 | 0,043233 | 0,262537322 |
| Q9D8H7 | OMA1 | Metalloendopeptidase OMA1, mitochondrial (EC 3.4.24.-) (Overlapping with the m-AAA protease 1 homolog) | -0,136845684 | 0,016188 | 0,161166446 |
| Q8K010 | OPLA | 5-oxoprolinase (EC 3.5.2.9) (5-oxo-L-prolinase) (5-OPase) (Pyroglutamase) | -0,280657841 | 0,001648 | 0,040389028 |
| O54734 | OST48 | Dolichyl-diphosphooligosaccharide--protein glycosyltransferase 48 kDa subunit (DDOST 48 kDa subunit) (Oligosaccharyl transferase 48 kDa subunit) | 0,176669652 | 0,043996 | 0,262985417 |
| Q8VE38 | OXND1 | Oxidoreductase NAD-binding domain-containing protein 1 (EC 1.-.-.-) | -0,128611135 | 0,047453 | 0,275137781 |
| Q9D404 | OXSM | 3-oxoacyl-[acyl-carrier-protein] synthase, mitochondrial (EC 2.3.1.41) (Beta-ketoacyl-ACP synthase) | -0,251506322 | 0,018783 | 0,173938529 |
| Q60715 | P4HA1 | Prolyl 4-hydroxylase subunit alpha-1 (4-PH alpha-1) (EC 1.14.11.2) (Procollagen-proline,2-oxoglutarate-4-dioxygenase subunit alpha-1) | -0,193491768 | 0,029224 | 0,217809595 |
| Q2TBE6 | P4K2A | Phosphatidylinositol 4-kinase type 2-alpha (EC 2.7.1.67) (Phosphatidylinositol 4-kinase type II-alpha) | -0,13948912 | 0,019377 | 0,178308117 |
| Q9Z0P4 | PALM | Paralemmin-1 (Paralemmin) | 0,155953232 | 0,041464 | 0,259388001 |
| Q9ES46 | PARVB | Beta-parvin | 0,193327449 | 0,017668 | 0,169724042 |
| Q8VI36 | PAXI | Paxillin | 0,1460145 | 0,030002 | 0,219420978 |
| Q3TVI8 | PBIP1 | Pre-B-cell leukemia transcription factor-interacting protein 1 (Hematopoietic PBX-interacting protein) | -0,162059694 | 0,029161 | 0,217809595 |
| Q6W8Q3 | PC4L1 | Purkinje cell protein 4-like protein 1 (PCP4-like protein 1) | 0,237699419 | 0,002187 | 0,048714388 |
| P57724 | PCBP4 | Poly(rC)-binding protein 4 (Alpha-CP4) | 0,214302511 | 0,007877 | 0,109408805 |
| Q91ZA3 | PCCA | Propionyl-CoA carboxylase alpha chain, mitochondrial (PCCase subunit alpha) (EC 6.4.1.3) (Propanoyl-CoA:carbon dioxide ligase subunit alpha) | -0,140386751 | 0,008858 | 0,117504333 |
| Q9Z0X4 | PDE3A | cGMP-inhibited 3',5'-cyclic phosphodiesterase A (EC 3.1.4.17) (Cyclic GMP-inhibited phosphodiesterase A) (CGI-PDE A) | 0,192251513 | 0,018373 | 0,17344108 |
| P08003 | PDIA4 | Protein disulfide-isomerase A4 (EC 5.3.4.1) (Endoplasmic reticulum resident protein 72) (ER protein 72) (ERp-72) (ERp72) | -0,224886919 | 0,020735 | 0,18440493 |
| P70271 | PDLI4 | PDZ and LIM domain protein 4 (LIM protein RIL) (Reversion-induced LIM protein) | -0,290187203 | 0,03462 | 0,238469099 |
| Q4VA53 | PDS5B | Sister chromatid cohesion protein PDS5 homolog B (Androgen-induced proliferation inhibitor) (Androgen-induced prostate proliferative shutoff-associated protein AS3) | 0,199014773 | 0,010019 | 0,122276261 |
| Q08481 | PECA1 | Platelet endothelial cell adhesion molecule (PECAM-1) (CD antigen CD31) | 0,185816039 | 0,041334 | 0,259388001 |
| Q05793 | PGBM | Basement membrane-specific heparan sulfate proteoglycan core protein (HSPG) [Cleaved into: Endorepellin; LG3 peptide] | -0,121496175 | 0,026598 | 0,209125643 |
| P22437 | PGH1 | Prostaglandin G/H synthase 1 (EC 1.14.99.1) (Cyclooxygenase-1) (COX-1) (Prostaglandin H2 synthase 1) (PGH synthase 1) (PGHS-1) (PHS 1) (Prostaglandin-endoperoxide synthase 1) | 0,18958739 | 0,027793 | 0,212119635 |
| Q9DAK9 | PHP14 | 14 kDa phosphohistidine phosphatase (EC 3.9.1.3) (Phosphohistidine phosphatase 1) (PHPT1) (Protein histidine phosphatase) (PHP) | 0,250750641 | 0,010574 | 0,124983812 |
| Q9DB26 | PHYD1 | Phytanoyl-CoA dioxygenase domain-containing protein 1 (EC 1.-.-.-) | -0,314111848 | 0,000305 | 0,011542566 |
| O35691 | PININ | Pinin | 0,190575014 | 0,003356 | 0,064413722 |
| Q8BSF4 | PISD | Phosphatidylserine decarboxylase proenzyme, mitochondrial (EC 4.1.1.65) [Cleaved into: Phosphatidylserine decarboxylase beta chain; Phosphatidylserine decarboxylase alpha chain] | -0,179588455 | 0,047135 | 0,273889621 |
| P70268 | PKN1 | Serine/threonine-protein kinase N1 (EC 2.7.11.13) (Protein kinase C-like 1) (Protein kinase C-like PKN) (Protein-kinase C-related kinase 1) (Serine-threonine protein kinase N) | 0,200548048 | 0,006464 | 0,097156735 |
| P97350 | PKP1 | Plakophilin-1 | -1,037283058 | 0,010613 | 0,124983812 |
| P43883 | PLIN2 | Perilipin-2 (Adipophilin) (Adipose differentiation-related protein) (ADRP) | -0,303460849 | 0,030484 | 0,220566508 |
| Q9DBG5 | PLIN3 | Perilipin-3 (Cargo selection protein TIP47) (Mannose-6-phosphate receptor-binding protein 1) | 0,303212341 | 0,00415 | 0,074262582 |
| Q8BVZ1 | PLIN5 | Perilipin-5 (Lipid droplet-associated protein PAT-1) (Lipid storage droplet protein 5) (Myocardial LD protein) | -0,367883284 | 0,002306 | 0,049825204 |
| Q9R0M4 | PODXL | Podocalyxin (Podocalyxin-like protein 1) (PC) (PCLP-1) | 0,291146022 | 0,012639 | 0,139268033 |
| P63087 | PP1G | Serine/threonine-protein phosphatase PP1-gamma catalytic subunit (PP-1G) (EC 3.1.3.16) (Protein phosphatase 1C catalytic subunit) | -0,188590753 | 0,003302 | 0,064213044 |
| Q3UM45 | PP1R7 | Protein phosphatase 1 regulatory subunit 7 (Protein phosphatase 1 regulatory subunit 22) | 0,212014384 | 0,026322 | 0,208082265 |
| Q7TSI3 | PP6R1 | Serine/threonine-protein phosphatase 6 regulatory subunit 1 (SAPS domain family member 1) | -0,138814467 | 0,049012 | 0,278258066 |
| Q922D4 | PP6R3 | Serine/threonine-protein phosphatase 6 regulatory subunit 3 (SAPS domain family member 3) | 0,155208304 | 0,023938 | 0,196316353 |
| Q8BP40 | PPA6 | Lysophosphatidic acid phosphatase type 6 (EC 3.1.3.2) (Acid phosphatase 6, lysophosphatidic) (Acid phosphatase-like protein 1) (PACPL1) | 0,329976403 | 0,000141 | 0,00686712 |
| P09242 | PPBT | Alkaline phosphatase, tissue-nonspecific isozyme (AP-TNAP) (TNSALP) (EC 3.1.3.1) (Alkaline phosphatase 2) (Alkaline phosphatase liver/bone/kidney isozyme) | 0,258409441 | 0,019864 | 0,180820079 |
| Q9QUR6 | PPCE | Prolyl endopeptidase (PE) (EC 3.4.21.26) (Post-proline cleaving enzyme) | -0,187231276 | 0,042185 | 0,259606922 |
| Q99KR7 | PPIF | Peptidyl-prolyl cis-trans isomerase F, mitochondrial (PPIase F) (EC 5.2.1.8) (Cyclophilin D) (CyP-D) (CypD) (Cyclophilin F) (Rotamase F) | -0,226162012 | 0,001403 | 0,035899128 |
| Q9D868 | PPIH | Peptidyl-prolyl cis-trans isomerase H (PPIase H) (EC 5.2.1.8) (Rotamase H) | 0,18006699 | 0,01425 | 0,149519386 |
| Q9EPX2 | PPN | Papilin | -0,241879085 | 0,00047 | 0,016924523 |
| Q8R5J9 | PRAF3 | PRA1 family protein 3 (ADP-ribosylation factor-like protein 6-interacting protein 5) (ARL-6-interacting protein 5) (Aip-5) (Addicsin) (GTRAP3-18) (Glutamate transporter EAAC1-interacting protein) (Prenylated Rab acceptor protein 2) (Protein JWa) | -0,291215017 | 0,000736 | 0,022605699 |
| P99029 | PRDX5 | Peroxiredoxin-5, mitochondrial (EC 1.11.1.15) (Antioxidant enzyme B166) (AOEB166) (Liver tissue 2D-page spot 2D-0014IV) (PLP) (Peroxiredoxin V) (Prx-V) (Peroxisomal antioxidant enzyme) (Thioredoxin peroxidase PMP20) | -0,148756485 | 0,015417 | 0,156691327 |
| Q99KP6 | PRP19 | Pre-mRNA-processing factor 19 (EC 2.3.2.27) (Nuclear matrix protein 200) (PRP19/PSO4 homolog) (RING-type E3 ubiquitin transferase PRP19) (Senescence evasion factor) | 0,144400775 | 0,023959 | 0,196316353 |
| P62196 | PRS8 | 26S proteasome regulatory subunit 8 (26S proteasome AAA-ATPase subunit RPT6) (Proteasome 26S subunit ATPase 5) (Proteasome subunit p45) (p45/SUG) (mSUG1) | 0,183309187 | 0,011456 | 0,130534294 |
| Q52KR3 | PRUN2 | Protein prune homolog 2 (BNIP2 motif-containing molecule at the C-terminal region 1) | -0,257054261 | 0,017426 | 0,168320047 |
| Q9D820 | PRXD1 | Prolyl-tRNA synthetase associated domain-containing protein 1 (PrdX deacylase domain-containing protein 1) | -0,202878629 | 0,035293 | 0,239730685 |
| Q9WVJ2 | PSD13 | 26S proteasome non-ATPase regulatory subunit 13 (26S proteasome regulatory subunit RPN9) (26S proteasome regulatory subunit S11) (26S proteasome regulatory subunit p40.5) | 0,155628387 | 0,022496 | 0,190682325 |
| Q8BHL8 | PSMF1 | Proteasome inhibitor PI31 subunit | -0,266725502 | 0,014414 | 0,149626304 |
| P0DJE0 | PT100 | Protein PET100 homolog, mitochondrial | -0,344278201 | 0,000692 | 0,021939435 |
| Q8BGC4 | PTGR3 | Prostaglandin reductase-3 (PTGR-3) (EC 1.3.1.48) (15-oxoprostaglandin 13-reductase) (Zinc-binding alcohol dehydrogenase domain-containing protein 2) | -0,654447473 | 8,00E-07 | 0,00023329 |
| P35235 | PTN11 | Tyrosine-protein phosphatase non-receptor type 11 (EC 3.1.3.48) (Protein-tyrosine phosphatase SYP) (SH-PTP2) (SHP-2) (Shp2) | -0,191838982 | 0,044493 | 0,264869442 |
| Q66GT5 | PTPM1 | Phosphatidylglycerophosphatase and protein-tyrosine phosphatase 1 (EC 3.1.3.27) (PTEN-like phosphatase) (Phosphoinositide lipid phosphatase) (Protein-tyrosine phosphatase mitochondrial 1) (EC 3.1.3.16) (EC 3.1.3.48) | 0,131144209 | 0,029975 | 0,219420978 |
| P18052 | PTPRA | Receptor-type tyrosine-protein phosphatase alpha (Protein-tyrosine phosphatase alpha) (R-PTP-alpha) (EC 3.1.3.48) (LCA-related phosphatase) (PTPTY-28) | 0,115022636 | 0,048092 | 0,275606009 |
| P28828 | PTPRM | Receptor-type tyrosine-protein phosphatase mu (Protein-tyrosine phosphatase mu) (R-PTP-mu) (EC 3.1.3.48) | -0,24196349 | 0,011181 | 0,12891725 |
| Q64737 | PUR2 | Trifunctional purine biosynthetic protein adenosine-3 [Includes: Phosphoribosylamine--glycine ligase (EC 6.3.4.13) (Glycinamide ribonucleotide synthetase) (GARS) (Phosphoribosylglycinamide synthetase); Phosphoribosylformylglycinamidine cyclo-ligase (EC 6.3.3.1) (AIR synthase) (AIRS) (Phosphoribosyl-aminoimidazole synthetase); Phosphoribosylglycinamide formyltransferase (EC 2.1.2.2) (5'-phosphoribosylglycinamide transformylase) (GAR transformylase) (GART)] | -0,566126 | 1,44E-07 | 0,000104828 |
| Q5SUR0 | PUR4 | Phosphoribosylformylglycinamidine synthase (FGAM synthase) (FGAMS) (EC 6.3.5.3) (Formylglycinamide ribonucleotide amidotransferase) (FGAR amidotransferase) (FGAR-AT) (Formylglycinamide ribotide amidotransferase) | 0,191286141 | 0,036218 | 0,240254199 |
| P70698 | PYRG1 | CTP synthase 1 (EC 6.3.4.2) (CTP synthetase 1) (UTP--ammonia ligase 1) | -0,25696036 | 0,021158 | 0,186457169 |
| Q9CZ13 | QCR1 | Cytochrome b-c1 complex subunit 1, mitochondrial (Complex III subunit 1) (Core protein I) (Ubiquinol-cytochrome-c reductase complex core protein 1) | -0,16881565 | 0,018763 | 0,173938529 |
| P99028 | QCR6 | Cytochrome b-c1 complex subunit 6, mitochondrial (Complex III subunit 6) (Complex III subunit VIII) (Cytochrome c1 non-heme 11 kDa protein) (Mitochondrial hinge protein) (Ubiquinol-cytochrome c reductase complex 11 kDa protein) | -0,394259641 | 0,00052 | 0,018058841 |
| Q9QYS9 | QKI | Protein quaking (MqkI) (qkI) | 0,22096679 | 0,000708 | 0,022220893 |
| P35283 | RAB12 | Ras-related protein Rab-12 (Rab-13) | -0,211947913 | 0,00439 | 0,076509517 |
| P62821 | RAB1A | Ras-related protein Rab-1A (YPT1-related protein) | -0,23594093 | 0,012534 | 0,139021738 |
| Q9D1G1 | RAB1B | Ras-related protein Rab-1B | -0,146692599 | 0,030668 | 0,220885079 |
| P63001 | RAC1 | Ras-related C3 botulinum toxin substrate 1 (p21-Rac1) | -0,509919505 | 3,32E-05 | 0,002422226 |
| Q64012 | RALY | RNA-binding protein Raly (Maternally-expressed hnRNP C-related protein) (hnRNP associated with lethal yellow protein) | 0,181373292 | 0,027569 | 0,212119635 |
| Q9CT10 | RANB3 | Ran-binding protein 3 (RanBP3) | 0,20672269 | 0,006618 | 0,098499204 |
| Q99JI6 | RAP1B | Ras-related protein Rap-1b (GTP-binding protein smg p21B) | 0,146176823 | 0,02289 | 0,190682325 |
| Q8BMG7 | RBGPR | Rab3 GTPase-activating protein non-catalytic subunit (Rab3 GTPase-activating protein 150 kDa subunit) (Rab3-GAP p150) (Rab3-GAP150) (Rab3-GAP regulatory subunit) | 0,20705592 | 0,045376 | 0,267940712 |
| Q8C2Q3 | RBM14 | RNA-binding protein 14 (RNA-binding motif protein 14) | -0,655523624 | 1,26E-08 | 3,69E-05 |
| B2RY56 | RBM25 | RNA-binding protein 25 (RNA-binding motif protein 25) | 0,132666109 | 0,032923 | 0,230034777 |
| P54728 | RD23B | UV excision repair protein RAD23 homolog B (HR23B) (mHR23B) (XP-C repair-complementing complex 58 kDa protein) (p58) | -0,151585309 | 0,026978 | 0,210416436 |
| Q64374 | RGN | Regucalcin (RC) (Gluconolactonase) (GNL) (EC 3.1.1.17) (Senescence marker protein 30) (SMP-30) | -0,982692994 | 0,045689 | 0,268696862 |
| Q6ZWV3 | RL10 | 60S ribosomal protein L10 (Protein QM homolog) (Ribosomal protein L10) | -0,38221221 | 0,0014 | 0,035899128 |
| P35979 | RL12 | 60S ribosomal protein L12 | -0,592056432 | 3,20E-07 | 0,000133367 |
| P62717 | RL18A | 60S ribosomal protein L18a | -0,807447041 | 1,71E-06 | 0,000356395 |
| P62830 | RL23 | 60S ribosomal protein L23 | -0,274393872 | 0,001335 | 0,034778462 |
| Q8BP67 | RL24 | 60S ribosomal protein L24 | -0,34301203 | 0,00133 | 0,034778462 |
| P14115 | RL27A | 60S ribosomal protein L27a (L29) | -0,4296985 | 1,84E-05 | 0,001735881 |
| P62900 | RL31 | 60S ribosomal protein L31 | -0,244939003 | 0,00905 | 0,117848645 |
| P62911 | RL32 | 60S ribosomal protein L32 | 0,204077496 | 0,034807 | 0,23890147 |
| Q9D1R9 | RL34 | 60S ribosomal protein L34 | -0,443114461 | 5,73E-06 | 0,000835338 |
| Q6ZWV7 | RL35 | 60S ribosomal protein L35 | -0,176878985 | 0,016876 | 0,165749384 |
| P83882 | RL36A | 60S ribosomal protein L36a (60S ribosomal protein L44) | -0,170005193 | 0,027827 | 0,212119635 |
| P14148 | RL7 | 60S ribosomal protein L7 | -0,15627931 | 0,018988 | 0,17528149 |
| P14869 | RLA0 | 60S acidic ribosomal protein P0 (60S ribosomal protein L10E) | -0,304911824 | 0,000645 | 0,021618197 |
| Q9DCU6 | RM04 | 39S ribosomal protein L4, mitochondrial (L4mt) (MRP-L4) | 0,192157453 | 0,001992 | 0,046864871 |
| Q3TBW2 | RM10 | 39S ribosomal protein L10, mitochondrial (L10mt) (MRP-L10) | -0,473003302 | 1,83E-05 | 0,001735881 |
| Q9D1I6 | RM14 | 39S ribosomal protein L14, mitochondrial (L14mt) (MRP-L14) | 0,167609602 | 0,010013 | 0,122276261 |
| Q99N93 | RM16 | 39S ribosomal protein L16, mitochondrial (L16mt) (MRP-L16) | -0,343802828 | 0,008862 | 0,117504333 |
| Q99N92 | RM27 | 39S ribosomal protein L27, mitochondrial (L27mt) (MRP-L27) | -0,168342207 | 0,00903 | 0,117848645 |
| Q8K2M0 | RM38 | 39S ribosomal protein L38, mitochondrial (L38mt) (MRP-L38) | -0,191063483 | 0,03565 | 0,240254199 |
| Q9D1H8 | RM53 | 39S ribosomal protein L53, mitochondrial (L53mt) (MRP-L53) | -0,19588774 | 0,025379 | 0,203142138 |
| Q9CX86 | ROA0 | Heterogeneous nuclear ribonucleoprotein A0 (hnRNP A0) | 0,241836078 | 0,016157 | 0,161166446 |
| Q8BG05 | ROA3 | Heterogeneous nuclear ribonucleoprotein A3 (hnRNP A3) | 0,185233711 | 0,007205 | 0,105091796 |
| P52432 | RPAC1 | DNA-directed RNA polymerases I and III subunit RPAC1 (DNA-directed RNA polymerase I subunit C) (RNA polymerases I and III subunit AC1) (AC40) (DNA-directed RNA polymerases I and III 40 kDa polypeptide) (RPA40) (RPC40) | 0,114646063 | 0,034663 | 0,238469099 |
| P08775 | RPB1 | DNA-directed RNA polymerase II subunit RPB1 (RNA polymerase II subunit B1) (EC 2.7.7.6) (DNA-directed RNA polymerase II subunit A) (DNA-directed RNA polymerase III largest subunit) | 0,205970502 | 0,006888 | 0,101997398 |
| Q8CHG7 | RPGF2 | Rap guanine nucleotide exchange factor 2 (Cyclic nucleotide ras GEF) (CNrasGEF) (Neural RAP guanine nucleotide exchange protein) (nRap GEP) (PDZ domain-containing guanine nucleotide exchange factor 1) (PDZ-GEF1) (RA-GEF-1) (Ras/Rap1-associating GEF-1) | -0,168456936 | 0,037495 | 0,244685234 |
| P62071 | RRAS2 | Ras-related protein R-Ras2 | -0,174333963 | 0,02412 | 0,197082132 |
| Q9D6S7 | RRFM | Ribosome-recycling factor, mitochondrial (RRF) (Ribosome-releasing factor, mitochondrial) | -0,16258906 | 0,035101 | 0,239230719 |
| Q9CSH3 | RRP44 | Exosome complex exonuclease RRP44 (EC 3.1.13.-) (EC 3.1.26.-) (Protein DIS3 homolog) (Ribosomal RNA-processing protein 44) | 0,181555787 | 0,039024 | 0,250730627 |
| P62281 | RS11 | 40S ribosomal protein S11 | -0,235024364 | 0,002344 | 0,050275715 |
| P62301 | RS13 | 40S ribosomal protein S13 | 0,175232404 | 0,010678 | 0,125093016 |
| P63276 | RS17 | 40S ribosomal protein S17 | -0,400661539 | 6,04E-05 | 0,003827295 |
| P62267 | RS23 | 40S ribosomal protein S23 | 0,245683674 | 0,025909 | 0,205572428 |
| P62849 | RS24 | 40S ribosomal protein S24 | 0,187400396 | 0,020022 | 0,180820079 |
| Q99N87 | RT05 | 28S ribosomal protein S5, mitochondrial (MRP-S5) (S5mt) | -0,126020386 | 0,043372 | 0,262537322 |
| Q9D7N3 | RT09 | 28S ribosomal protein S9, mitochondrial (MRP-S9) (S9mt) | -0,276544878 | 0,001089 | 0,030854348 |
| Q9CQE3 | RT17 | 28S ribosomal protein S17, mitochondrial (MRP-S17) (S17mt) | -0,176734106 | 0,004406 | 0,076509517 |
| P57784 | RU2A | U2 small nuclear ribonucleoprotein A' (U2 snRNP A') | 0,186104699 | 0,004294 | 0,075457482 |
| P56565 | S10A1 | Protein S100-A1 (S-100 protein alpha chain) (S-100 protein subunit alpha) (S100 calcium-binding protein A1) | 1,106413073 | 0,003229 | 0,063752636 |
| P50543 | S10AB | Protein S100-A11 (Calgizzarin) (Endothelial monocyte-activating polypeptide) (EMAP) (Protein S100-C) (S100 calcium-binding protein A11) | -0,276990146 | 0,003235 | 0,063752636 |
| Q8R0Y8 | S2542 | Mitochondrial coenzyme A transporter SLC25A42 (Solute carrier family 25 member 42) | -0,133812654 | 0,047778 | 0,275264714 |
| Q60714 | S27A1 | Long-chain fatty acid transport protein 1 (FATP-1) (Fatty acid transport protein 1) (EC 6.2.1.-) (Fatty acid transport protein) (Solute carrier family 27 member 1) | 0,210247778 | 0,030029 | 0,219420978 |
| Q9D2C2 | SAAL1 | Protein SAAL1 (Synoviocyte proliferation-associated in collagen-induced arthritis protein 1) (SPACIA1) | 0,33496555 | 0,000141 | 0,00686712 |
| Q9D1J3 | SARNP | SAP domain-containing ribonucleoprotein (Nuclear protein Hcc-1) | 0,250718245 | 0,002105 | 0,047975731 |
| O35609 | SCAM3 | Secretory carrier-associated membrane protein 3 (Secretory carrier membrane protein 3) | 0,180275641 | 0,025119 | 0,203142138 |
| Q8BMD8 | SCMC1 | Calcium-binding mitochondrial carrier protein SCaMC-1 (Small calcium-binding mitochondrial carrier protein 1) (Solute carrier family 25 member 24) | 0,160028529 | 0,036568 | 0,240254199 |
| Q9D0K2 | SCOT1 | Succinyl-CoA:3-ketoacid coenzyme A transferase 1, mitochondrial (EC 2.8.3.5) (3-oxoacid CoA-transferase 1) (Somatic-type succinyl-CoA:3-oxoacid CoA-transferase) (SCOT-s) | 0,195956877 | 0,018698 | 0,173938529 |
| O35114 | SCRB2 | Lysosome membrane protein 2 (85 kDa lysosomal membrane sialoglycoprotein) (LGP85) (Lysosome membrane protein II) (LIMP II) (Scavenger receptor class B member 2) | -0,282231043 | 0,003602 | 0,068222436 |
| Q9CQA3 | SDHB | Succinate dehydrogenase [ubiquinone] iron-sulfur subunit, mitochondrial (EC 1.3.5.1) (Iron-sulfur subunit of complex II) (Ip) | -0,187651118 | 0,004027 | 0,073636148 |
| Q8C1B7 | SEP11 | Septin-11 | 0,151440668 | 0,020962 | 0,185617161 |
| Q921M3 | SF3B3 | Splicing factor 3B subunit 3 (Pre-mRNA-splicing factor SF3b 130 kDa subunit) (SF3b130) (Spliceosome-associated protein 130) (SAP 130) | 0,188801864 | 0,004294 | 0,075457482 |
| Q8VIJ6 | SFPQ | Splicing factor, proline- and glutamine-rich (DNA-binding p52/p100 complex, 100 kDa subunit) (Polypyrimidine tract-binding protein-associated-splicing factor) (PSF) (PTB-associated-splicing factor) | 0,144120957 | 0,038163 | 0,247381426 |
| P82350 | SGCA | Alpha-sarcoglycan (Alpha-SG) (50 kDa dystrophin-associated glycoprotein) (50DAG) (Adhalin) | -0,114108955 | 0,043561 | 0,262537322 |
| P82348 | SGCG | Gamma-sarcoglycan (Gamma-SG) (35 kDa dystrophin-associated glycoprotein) (35DAG) | 0,218577739 | 0,005711 | 0,090544269 |
| Q8BJU0 | SGTA | Small glutamine-rich tetratricopeptide repeat-containing protein alpha (Alpha-SGT) | -0,26679276 | 0,009586 | 0,120876193 |
| Q9JJU8 | SH3L1 | SH3 domain-binding glutamic acid-rich-like protein | -0,363963093 | 2,69E-05 | 0,002164174 |
| Q9JK48 | SHLB1 | Endophilin-B1 (SH3 domain-containing GRB2-like protein B1) | -0,168986689 | 0,006495 | 0,097156735 |
| Q9QXN0 | SHRM3 | Protein Shroom3 | 0,120113111 | 0,040705 | 0,256765664 |
| Q8K2C6 | SIR5 | NAD-dependent protein deacylase sirtuin-5, mitochondrial (EC 3.5.1.-) (Regulatory protein SIR2 homolog 5) (SIR2-like protein 5) | -0,380044248 | 7,15E-05 | 0,004034167 |
| Q7TN29 | SMAP2 | Stromal membrane-associated protein 2 (Stromal membrane-associated protein 1-like) | -0,196387085 | 0,02301 | 0,190682325 |
| Q9CW03 | SMC3 | Structural maintenance of chromosomes protein 3 (SMC protein 3) (SMC-3) (Basement membrane-associated chondroitin proteoglycan) (Bamacan) (Chondroitin sulfate proteoglycan 6) (Chromosome segregation protein SmcD) (Mad member-interacting protein 1) | 0,160522207 | 0,017131 | 0,166477456 |
| Q91ZW3 | SMCA5 | SWI/SNF-related matrix-associated actin-dependent regulator of chromatin subfamily A member 5 (EC 3.6.4.-) (Sucrose nonfermenting protein 2 homolog) (mSnf2h) | 0,282045863 | 0,005521 | 0,089157901 |
| P62317 | SMD2 | Small nuclear ribonucleoprotein Sm D2 (Sm-D2) (snRNP core protein D2) | 0,115040999 | 0,033702 | 0,234065265 |
| Q9CZ28 | SNF8 | Vacuolar-sorting protein SNF8 (ESCRT-II complex subunit VPS22) | 0,156379571 | 0,025934 | 0,205572428 |
| O09044 | SNP23 | Synaptosomal-associated protein 23 (SNAP-23) (Syndet) (Vesicle-membrane fusion protein SNAP-23) | 0,149938447 | 0,012476 | 0,138902675 |
| Q62189 | SNRPA | U1 small nuclear ribonucleoprotein A (U1 snRNP A) (U1-A) (U1A) | 0,167604779 | 0,023813 | 0,196218478 |
| Q9D8U8 | SNX5 | Sorting nexin-5 | 0,156568583 | 0,031433 | 0,22363408 |
| Q6P8X1 | SNX6 | Sorting nexin-6 [Cleaved into: Sorting nexin-6, N-terminally processed] | 0,17354225 | 0,013545 | 0,144196646 |
| Q920B9 | SP16H | FACT complex subunit SPT16 (Chromatin-specific transcription elongation factor 140 kDa subunit) (FACT 140 kDa subunit) (FACTp140) (Facilitates chromatin transcription complex subunit SPT16) | 0,236149464 | 0,008284 | 0,112396429 |
| Q7TNE3 | SPAG7 | Sperm-associated antigen 7 | 0,261233172 | 0,022437 | 0,190682325 |
| P14576 | SRP54 | Signal recognition particle 54 kDa protein (SRP54) | -0,129134029 | 0,041905 | 0,259526223 |
| Q62093 | SRSF2 | Serine/arginine-rich splicing factor 2 (Protein PR264) (Putative myelin regulatory factor 1) (MRF-1) (Splicing component, 35 kDa) (Splicing factor SC35) (SC-35) (Splicing factor, arginine/serine-rich 2) | 0,193523835 | 0,006134 | 0,094674158 |
| Q8BWF0 | SSDH | Succinate-semialdehyde dehydrogenase, mitochondrial (EC 1.2.1.24) (Aldehyde dehydrogenase family 5 member A1) (NAD(+)-dependent succinic semialdehyde dehydrogenase) | -0,162907787 | 0,045008 | 0,266843844 |
| Q62147 | SSPN | Sarcospan (K-ras oncogene-associated protein) (Kirsten-Ras-associated protein) | 0,348432959 | 0,0102 | 0,123669585 |
| Q08943 | SSRP1 | FACT complex subunit SSRP1 (Facilitates chromatin transcription complex subunit SSRP1) (Recombination signal sequence recognition protein 1) (Structure-specific recognition protein 1) (T160) | 0,21499255 | 0,0127 | 0,139268033 |
| Q9D3E6 | STAG1 | Cohesin subunit SA-1 (SCC3 homolog 1) (Stromal antigen 1) | 0,228218422 | 0,010345 | 0,123669585 |
| P42227 | STAT3 | Signal transducer and activator of transcription 3 (Acute-phase response factor) | 0,183603078 | 0,021739 | 0,188260494 |
| Q60864 | STIP1 | Stress-induced-phosphoprotein 1 (STI1) (mSTI1) (Hsc70/Hsp90-organizing protein) (Hop) | -0,479810751 | 2,27E-05 | 0,002067922 |
| Q8C9H6 | STRP2 | Striatin-interacting proteins 2 (Protein FAM40B) | 0,280451981 | 0,034557 | 0,238469099 |
| Q9Z2I9 | SUCB1 | Succinate--CoA ligase [ADP-forming] subunit beta, mitochondrial (EC 6.2.1.5) (ATP-specific succinyl-CoA synthetase subunit beta) (A-SCS) (Succinyl-CoA synthetase beta-A chain) (SCS-betaA) | -0,178152982 | 0,01022 | 0,123669585 |
| Q8BJS4 | SUN2 | SUN domain-containing protein 2 (Protein unc-84 homolog B) (Sad1/unc-84 protein-like 2) | 0,466106476 | 0,00029 | 0,011270245 |
| Q64310 | SURF4 | Surfeit locus protein 4 | -0,159609105 | 0,032964 | 0,230034777 |
| Q8BYM8 | SYCM | Probable cysteine--tRNA ligase, mitochondrial (EC 6.1.1.16) (Cysteinyl-tRNA synthetase) (CysRS) | -0,193507496 | 0,029717 | 0,218899381 |
| Q99MN1 | SYK | Lysine--tRNA ligase (EC 2.7.7.-) (EC 6.1.1.6) (Lysyl-tRNA synthetase) (LysRS) | -0,121806629 | 0,049314 | 0,278777489 |
| Q8BLY2 | SYTC2 | Probable threonine--tRNA ligase 2, cytoplasmic (EC 6.1.1.3) (Threonyl-tRNA synthetase) (ThrRS) (Threonyl-tRNA synthetase-like protein 2) | -0,126343328 | 0,043473 | 0,262537322 |
| Q9CYK1 | SYWM | Tryptophan--tRNA ligase, mitochondrial (EC 6.1.1.2) ((Mt)TrpRS) (Tryptophanyl-tRNA synthetase) (TrpRS) | 0,473149008 | 0,000216 | 0,009612549 |
| Q8K1H7 | T11L2 | T-complex protein 11-like protein 2 | 0,334453206 | 0,00307 | 0,062631645 |
| Q921F2 | TADBP | TAR DNA-binding protein 43 (TDP-43) | 0,25185597 | 0,005632 | 0,09000375 |
| P10637 | TAU | Microtubule-associated protein tau (Neurofibrillary tangle protein) (Paired helical filament-tau) (PHF-tau) | -0,264616824 | 0,006171 | 0,094744747 |
| P68368 | TBA4A | Tubulin alpha-4A chain (Alpha-tubulin 4) (Alpha-tubulin isotype M-alpha-4) (Tubulin alpha-4 chain) | -0,347654016 | 0,000295 | 0,011336551 |
| Q7TMM9 | TBB2A | Tubulin beta-2A chain | -0,186464649 | 0,019898 | 0,180820079 |
| P99024 | TBB5 | Tubulin beta-5 chain | -0,121502668 | 0,032504 | 0,229840195 |
| Q80XQ2 | TBCD5 | TBC1 domain family member 5 | 0,203129791 | 0,021249 | 0,18669793 |
| Q8CIV8 | TBCE | Tubulin-specific chaperone E (Tubulin-folding cofactor E) | 0,321292772 | 0,007346 | 0,105149698 |
| P11983 | TCPA | T-complex protein 1 subunit alpha (TCP-1-alpha) (CCT-alpha) (Tailless complex polypeptide 1A) (TCP-1-A) (Tailless complex polypeptide 1B) (TCP-1-B) | -0,407958579 | 7,48E-05 | 0,004081112 |
| Q8CGF7 | TCRG1 | Transcription elongation regulator 1 (Formin-binding protein 28) (FBP 28) (TATA box-binding protein-associated factor 2S) (Transcription factor CA150) (p144) | 0,183285447 | 0,006396 | 0,096675112 |
| Q9CY27 | TECR | Very-long-chain enoyl-CoA reductase (EC 1.3.1.93) (Synaptic glycoprotein SC2) (Trans-2,3-enoyl-CoA reductase) (TER) | 0,154498359 | 0,030841 | 0,221038409 |
| O70548 | TELT | Telethonin (Titin cap protein) | 0,328588418 | 0,027854 | 0,212119635 |
| Q80ZW2 | THEM6 | Protein THEM6 | -0,254476081 | 0,00162 | 0,040203758 |
| Q8BWT1 | THIM | 3-ketoacyl-CoA thiolase, mitochondrial (EC 2.3.1.16) (Acetyl-CoA acyltransferase) (Beta-ketothiolase) (Mitochondrial 3-oxoacyl-CoA thiolase) | -0,119521716 | 0,037815 | 0,245759639 |
| Q8BH55 | THNS1 | Threonine synthase-like 1 (TSH1) | -0,404492678 | 4,30E-06 | 0,000696938 |
| O08583 | THOC4 | THO complex subunit 4 (Tho4) (Ally of AML-1 and LEF-1) (Aly/REF export factor) (REF1-I) (RNA and export factor-binding protein 1) (Transcriptional coactivator Aly/REF) | 0,222219143 | 0,018362 | 0,17344108 |
| P62075 | TIM13 | Mitochondrial import inner membrane translocase subunit Tim13 | -0,465469357 | 0,000229 | 0,009612549 |
| Q9WTQ8 | TIM23 | Mitochondrial import inner membrane translocase subunit Tim23 | -0,278281111 | 0,003219 | 0,063752636 |
| Q3TMP8 | TM38A | Trimeric intracellular cation channel type A (TRIC-A) (TRICA) (27 kDa sarcoplasmic reticulum protein) (Mitsugumin-33A) (SPR-27) (Transmembrane protein 38A) | 0,239529824 | 0,027924 | 0,212119635 |
| Q8BK08 | TMM11 | Transmembrane protein 11, mitochondrial (Protein PM1) | 0,185083385 | 0,00875 | 0,117076745 |
| Q9CQN3 | TOM6 | Mitochondrial import receptor subunit TOM6 homolog (Overexpressed breast tumor protein homolog) (Translocase of outer membrane 6 kDa subunit homolog) | -0,181256158 | 0,042289 | 0,259696416 |
| O55013 | TPPC3 | Trafficking protein particle complex subunit 3 (BET3 homolog) | 0,154286546 | 0,00498 | 0,082544873 |
| Q921I1 | TRFE | Serotransferrin (Transferrin) (Beta-1 metal-binding globulin) (Siderophilin) | -0,740401794 | 0,001202 | 0,033085203 |
| Q38HM4 | TRI63 | E3 ubiquitin-protein ligase TRIM63 (EC 2.3.2.27) (Muscle-specific RING finger protein 1) (MuRF-1) (MuRF1) (Muscle RING finger protein 1) (RING-type E3 ubiquitin transferase TRIM63) (Tripartite motif-containing protein 63) | 0,241384497 | 0,008623 | 0,115919325 |
| Q61187 | TS101 | Tumor susceptibility gene 101 protein (ESCRT-I complex subunit TSG101) | 0,192094321 | 0,005646 | 0,09000375 |
| Q62348 | TSN | Translin (EC 3.1.-.-) (Component 3 of promoter of RISC) (C3PO) (Testis/brain RNA-binding protein) (TB-RBP) | 0,22611202 | 0,012859 | 0,140385658 |
| Q6P4T2 | U520 | U5 small nuclear ribonucleoprotein 200 kDa helicase (EC 3.6.4.13) (BRR2 homolog) (U5 snRNP-specific 200 kDa protein) (U5-200KD) | 0,127977152 | 0,021908 | 0,1888568 |
| O08810 | U5S1 | 116 kDa U5 small nuclear ribonucleoprotein component (Elongation factor Tu GTP-binding domain-containing protein 2) (U5 snRNP-specific protein, 116 kDa) (U5-116 kDa) | 0,201762477 | 0,002557 | 0,054452455 |
| P68037 | UB2L3 | Ubiquitin-conjugating enzyme E2 L3 (EC 2.3.2.23) (E2 ubiquitin-conjugating enzyme L3) (UbcM4) (Ubiquitin carrier protein L3) (Ubiquitin-protein ligase L3) | -0,177561411 | 0,021728 | 0,188260494 |
| Q8C878 | UBA3 | NEDD8-activating enzyme E1 catalytic subunit (EC 6.2.1.-) (NEDD8-activating enzyme E1C) (Ubiquitin-activating enzyme E1C) (Ubiquitin-like modifier-activating enzyme 3) (Ubiquitin-activating enzyme 3) | 0,139660168 | 0,022974 | 0,190682325 |
| P63280 | UBC9 | SUMO-conjugating enzyme UBC9 (EC 2.3.2.-) (RING-type E3 SUMO transferase UBC9) (SUMO-protein ligase) (Ubiquitin carrier protein 9) (mUBC9) (Ubiquitin carrier protein I) (Ubiquitin-conjugating enzyme E2 I) (Ubiquitin-protein ligase I) | 0,169147082 | 0,019902 | 0,180820079 |
| Q8CCJ3 | UFL1 | E3 UFM1-protein ligase 1 (EC 2.3.2.-) (E3 UFM1-protein transferase 1) | -0,156982334 | 0,036069 | 0,240254199 |
| Q8VBW6 | ULA1 | NEDD8-activating enzyme E1 regulatory subunit (Amyloid beta precursor protein-binding protein 1, 59 kDa) (APP-BP1) (Amyloid protein-binding protein 1) | -0,42270558 | 0,000235 | 0,009612549 |
| Q9CWU6 | UQCC1 | Ubiquinol-cytochrome-c reductase complex assembly factor 1 (Basic FGF-repressed Zic-binding protein) (mbFZb) (Ubiquinol-cytochrome c reductase complex chaperone CBP3 homolog) | -0,197472764 | 0,015979 | 0,160711106 |
| Q5NCI0 | URGCP | Up-regulator of cell proliferation (HBV X protein up-regulated gene 4 protein homolog) (HBxAg up-regulated gene 4 protein homolog) | 0,204849982 | 0,022372 | 0,190682325 |
| Q06318 | UTER | Uteroglobin (Clara cell 17 kDa protein) (Clara cell phospholipid-binding protein) (CCPBP) (Clara cells 10 kDa secretory protein) (CC10) (PCB-binding protein) (Secretoglobin family 1A member 1) | -0,867293514 | 0,047843 | 0,275264714 |
| P51863 | VA0D1 | V-type proton ATPase subunit d 1 (V-ATPase subunit d 1) (P39) (Physophilin) (V-ATPase 40 kDa accessory protein) (V-ATPase AC39 subunit) (Vacuolar proton pump subunit d 1) | 0,15131235 | 0,014824 | 0,152800671 |
| Q62465 | VAT1 | Synaptic vesicle membrane protein VAT-1 homolog (EC 1.-.-.-) | 0,186927026 | 0,008982 | 0,117848645 |
| Q60930 | VDAC2 | Voltage-dependent anion-selective channel protein 2 (VDAC-2) (mVDAC2) (Outer mitochondrial membrane protein porin 2) (Voltage-dependent anion-selective channel protein 6) (VDAC-6) (mVDAC6) | -0,151780293 | 0,025489 | 0,203142138 |
| Q9R1Z8 | VINEX | Vinexin (SH3 domain-containing protein SH3P3) (SH3-containing adapter molecule 1) (SCAM-1) (Sorbin and SH3 domain-containing protein 3) | -0,188629718 | 0,027645 | 0,212119635 |
| Q99KC8 | VMA5A | von Willebrand factor A domain-containing protein 5A (Loss of heterozygosity 11 chromosomal region 2 gene A protein homolog) | 0,22938597 | 0,020213 | 0,181978067 |
| Q9EQH3 | VPS35 | Vacuolar protein sorting-associated protein 35 (Maternal-embryonic 3) (Vesicle protein sorting 35) | 0,182954466 | 0,017179 | 0,166477456 |
| Q8BH43 | WASF2 | Wiskott-Aldrich syndrome protein family member 2 (WASP family protein member 2) (Protein WAVE-2) | 0,171159239 | 0,009511 | 0,120629452 |
| O88342 | WDR1 | WD repeat-containing protein 1 (Actin-interacting protein 1) (AIP1) | -0,315411053 | 0,04762 | 0,275137781 |
| Q00519 | XDH | Xanthine dehydrogenase/oxidase [Includes: Xanthine dehydrogenase (XD) (EC 1.17.1.4); Xanthine oxidase (XO) (EC 1.17.3.2) (Xanthine oxidoreductase) (XOR)] | 0,127952607 | 0,018536 | 0,173938529 |
| P27641 | XRCC5 | X-ray repair cross-complementing protein 5 (EC 3.6.4.-) (ATP-dependent DNA helicase 2 subunit 2) (ATP-dependent DNA helicase II 80 kDa subunit) (CTC box-binding factor 85 kDa subunit) (CTC85) (CTCBF) (DNA repair protein XRCC5) (Ku autoantigen protein p86 homolog) (Ku80) (Nuclear factor IV) | 0,243155707 | 0,002097 | 0,047975731 |
| Q3UPF5 | ZCCHV | Zinc finger CCCH-type antiviral protein 1 (ADP-ribosyltransferase diphtheria toxin-like 13) (ARTD13) | -0,172091259 | 0,021948 | 0,1888568 |
| Q9Z0U1 | ZO2 | Tight junction protein ZO-2 (Tight junction protein 2) (Zona occludens protein 2) (Zonula occludens protein 2) | 0,136571201 | 0,013029 | 0,141288342 |
| Q62523 | ZYX | Zyxin | -0,561557134 | 6,80E-05 | 0,004034167 |

**Supplementary Data Table 5. List of significantly altered senescence-related proteins (*p*-value<0.05) in Physiologically aged mice**

| **Uniprot ID** | **Protein Symbol** | **Protein Description** | **Average Log2 Ratio** | ***p*-value** | **Adjusted *p*-value** |
| --- | --- | --- | --- | --- | --- |
| Q8VD66 | ABHD4 | Protein ABHD4 (EC 3.-.-.-) (Alpha/beta hydrolase domain-containing protein 4) (Abhydrolase domain-containing protein 4) (Alpha/beta-hydrolase 4) (Lyso-N-acylphosphatidylethanolamine lipase) | 0,248931523 | 0,005881 | 0,091744717 |
| P28352 | APEX1 | DNA-(apurinic or apyrimidinic site) lyase (EC 3.1.-.-) (EC 4.2.99.18) (APEX nuclease) (APEN) (Apurinic-apyrimidinic endonuclease 1) (AP endonuclease 1) (REF-1) (Redox factor-1) [Cleaved into: DNA-(apurinic or apyrimidinic site) lyase, mitochondrial] | -0,280376441 | 0,00127 | 0,033751711 |
| Q9Z277 | BAZ1B | Tyrosine-protein kinase BAZ1B (EC 2.7.10.2) (Bromodomain adjacent to zinc finger domain protein 1B) (Williams syndrome transcription factor homolog) (Williams-Beuren syndrome chromosomal region 9 protein homolog) | 0,129076502 | 0,041749 | 0,259388001 |
| Q9WVC3 | CAV2 | Caveolin-2 | -0,273504107 | 0,002818 | 0,059147505 |
| P23198 | CBX3 | Chromobox protein homolog 3 (Heterochromatin protein 1 homolog gamma) (HP1 gamma) (M32) (Modifier 2 protein) | 0,191189309 | 0,005808 | 0,091573162 |
| Q8BMS4 | COQ3 | Ubiquinone biosynthesis O-methyltransferase, mitochondrial (3-demethylubiquinol 3-O-methyltransferase) (EC 2.1.1.64) (Polyprenyldihydroxybenzoate methyltransferase) (EC 2.1.1.114) | -0,238851286 | 0,003997 | 0,073636148 |
| Q9DAR7 | DCPS | m7GpppX diphosphatase (EC 3.6.1.59) (DCS-1) (Decapping scavenger enzyme) (Hint-related 7meGMP-directed hydrolase) (Histidine triad nucleotide-binding protein 5) (Histidine triad protein member 5) (HINT-5) (Scavenger mRNA-decapping enzyme DcpS) | 0,186058408 | 0,018207 | 0,17344108 |
| Q9CWS0 | DDAH1 | N(G),N(G)-dimethylarginine dimethylaminohydrolase 1 (DDAH-1) (Dimethylarginine dimethylaminohydrolase 1) (EC 3.5.3.18) (DDAHI) (Dimethylargininase-1) | -0,286442274 | 0,001165 | 0,032359119 |
| Q6Q899 | DDX58 | Probable ATP-dependent RNA helicase DDX58 (EC 3.6.4.13) (DEAD box protein 58) (RIG-I-like receptor 1) (RLR-1) (Retinoic acid-inducible gene 1 protein) (RIG-1) (Retinoic acid-inducible gene I protein) (RIG-I) | 0,250038602 | 0,04445 | 0,264869442 |
| Q7TNV0 | DEK | Protein DEK | 0,212517783 | 0,001626 | 0,040203758 |
| Q99L04 | DHRS1 | Dehydrogenase/reductase SDR family member 1 (EC 1.1.-.-) | -0,251148457 | 0,001751 | 0,042208398 |
| P84089 | ERH | Enhancer of rudimentary homolog (Mer) | 0,232110926 | 0,002898 | 0,060389411 |
| Q05816 | FABP5 | Fatty acid-binding protein 5 (Epidermal-type fatty acid-binding protein) (E-FABP) (Fatty acid-binding protein, epidermal) (Keratinocyte lipid-binding protein) (Psoriasis-associated fatty acid-binding protein homolog) (PA-FABP) | -0,498434268 | 0,000971 | 0,028033803 |
| P56959 | FUS | RNA-binding protein FUS (Protein pigpen) | 0,208726242 | 0,004947 | 0,082459629 |
| Q9D7X8 | GGCT | Gamma-glutamylcyclotransferase (EC 4.3.2.9) | 0,201325568 | 0,009614 | 0,120876193 |
| Q9QZF2 | GPC1 | Glypican-1 [Cleaved into: Secreted glypican-1] | -0,478386881 | 4,44E-05 | 0,003014374 |
| P17095 | HMGA1 | High mobility group protein HMG-I/HMG-Y (HMG-I(Y)) (High mobility group AT-hook protein 1) (High mobility group protein A1) | 0,375127582 | 0,048085 | 0,275606009 |
| P63158 | HMGB1 | High mobility group protein B1 (High mobility group protein 1) (HMG-1) | 0,317566129 | 0,001273 | 0,033751711 |
| O54879 | HMGB3 | High mobility group protein B3 (High mobility group protein 2a) (HMG-2a) (High mobility group protein 4) (HMG-4) | 0,30095374 | 0,015673 | 0,15819026 |
| Q91UZ5 | IMPA2 | Inositol monophosphatase 2 (IMP 2) (IMPase 2) (EC 3.1.3.25) (Inositol-1(or 4)-monophosphatase 2) (Myo-inositol monophosphatase A2) | 0,284004845 | 0,009129 | 0,118349458 |
| P21619 | LMNB2 | Lamin-B2 | -0,347676151 | 0,000504 | 0,017933802 |
| Q9D023 | MPC2 | Mitochondrial pyruvate carrier 2 (Brain protein 44) | -0,169701341 | 0,011392 | 0,13031752 |
| Q9R0M4 | PODXL | Podocalyxin (Podocalyxin-like protein 1) (PC) (PCLP-1) | 0,291146022 | 0,012639 | 0,139268033 |
| Q9D868 | PPIH | Peptidyl-prolyl cis-trans isomerase H (PPIase H) (EC 5.2.1.8) (Rotamase H) | 0,18006699 | 0,01425 | 0,149519386 |
| P28828 | PTPRM | Receptor-type tyrosine-protein phosphatase mu (Protein-tyrosine phosphatase mu) (R-PTP-mu) (EC 3.1.3.48) | -0,24196349 | 0,011181 | 0,12891725 |
| Q9D2C2 | SAAL1 | Protein SAAL1 (Synoviocyte proliferation-associated in collagen-induced arthritis protein 1) (SPACIA1) | 0,33496555 | 0,000141 | 0,00686712 |
| Q921M3 | SF3B3 | Splicing factor 3B subunit 3 (Pre-mRNA-splicing factor SF3b 130 kDa subunit) (SF3b130) (Spliceosome-associated protein 130) (SAP 130) | 0,188801864 | 0,004294 | 0,075457482 |
| Q8VIJ6 | SFPQ | Splicing factor, proline- and glutamine-rich (DNA-binding p52/p100 complex, 100 kDa subunit) (Polypyrimidine tract-binding protein-associated-splicing factor) (PSF) (PTB-associated-splicing factor) | 0,144120957 | 0,038163 | 0,247381426 |
| Q9CW03 | SMC3 | Structural maintenance of chromosomes protein 3 (SMC protein 3) (SMC-3) (Basement membrane-associated chondroitin proteoglycan) (Bamacan) (Chondroitin sulfate proteoglycan 6) (Chromosome segregation protein SmcD) (Mad member-interacting protein 1) | 0,160522207 | 0,017131 | 0,166477456 |
| Q62189 | SNRPA | U1 small nuclear ribonucleoprotein A (U1 snRNP A) (U1-A) (U1A) | 0,167604779 | 0,023813 | 0,196218478 |
| Q9D8U8 | SNX5 | Sorting nexin-5 | 0,156568583 | 0,031433 | 0,22363408 |
| Q08943 | SSRP1 | FACT complex subunit SSRP1 (Facilitates chromatin transcription complex subunit SSRP1) (Recombination signal sequence recognition protein 1) (Structure-specific recognition protein 1) (T160) | 0,21499255 | 0,0127 | 0,139268033 |

**Supplementary Data Table 6. List of significantly altered senescence-related proteins (p-value<0.05) in HGPS mice**

| **Uniprot ID** | **Protein Symbol** | **Protein Description** | **Average Log2 Ratio** | ***p*-value** | **Adjusted *p*-value** |
| --- | --- | --- | --- | --- | --- |
| Q8VD66 | ABHD4 | Protein ABHD4 (EC 3.-.-.-) (Alpha/beta hydrolase domain-containing protein 4) (Abhydrolase domain-containing protein 4) (Alpha/beta-hydrolase 4) (Lyso-N-acylphosphatidylethanolamine lipase) | 0,248931523 | 0,005881 | 0,091744717 |
| P28352 | APEX1 | DNA-(apurinic or apyrimidinic site) lyase (EC 3.1.-.-) (EC 4.2.99.18) (APEX nuclease) (APEN) (Apurinic-apyrimidinic endonuclease 1) (AP endonuclease 1) (REF-1) (Redox factor-1) [Cleaved into: DNA-(apurinic or apyrimidinic site) lyase, mitochondrial] | -0,280376441 | 0,00127 | 0,033751711 |
| Q9Z277 | BAZ1B | Tyrosine-protein kinase BAZ1B (EC 2.7.10.2) (Bromodomain adjacent to zinc finger domain protein 1B) (Williams syndrome transcription factor homolog) (Williams-Beuren syndrome chromosomal region 9 protein homolog) | 0,129076502 | 0,041749 | 0,259388001 |
| Q9WVC3 | CAV2 | Caveolin-2 | -0,273504107 | 0,002818 | 0,059147505 |
| P23198 | CBX3 | Chromobox protein homolog 3 (Heterochromatin protein 1 homolog gamma) (HP1 gamma) (M32) (Modifier 2 protein) | 0,191189309 | 0,005808 | 0,091573162 |
| Q8BMS4 | COQ3 | Ubiquinone biosynthesis O-methyltransferase, mitochondrial (3-demethylubiquinol 3-O-methyltransferase) (EC 2.1.1.64) (Polyprenyldihydroxybenzoate methyltransferase) (EC 2.1.1.114) | -0,238851286 | 0,003997 | 0,073636148 |
| Q9DAR7 | DCPS | m7GpppX diphosphatase (EC 3.6.1.59) (DCS-1) (Decapping scavenger enzyme) (Hint-related 7meGMP-directed hydrolase) (Histidine triad nucleotide-binding protein 5) (Histidine triad protein member 5) (HINT-5) (Scavenger mRNA-decapping enzyme DcpS) | 0,186058408 | 0,018207 | 0,17344108 |
| Q9CWS0 | DDAH1 | N(G),N(G)-dimethylarginine dimethylaminohydrolase 1 (DDAH-1) (Dimethylarginine dimethylaminohydrolase 1) (EC 3.5.3.18) (DDAHI) (Dimethylargininase-1) | -0,286442274 | 0,001165 | 0,032359119 |
| Q6Q899 | DDX58 | Probable ATP-dependent RNA helicase DDX58 (EC 3.6.4.13) (DEAD box protein 58) (RIG-I-like receptor 1) (RLR-1) (Retinoic acid-inducible gene 1 protein) (RIG-1) (Retinoic acid-inducible gene I protein) (RIG-I) | 0,250038602 | 0,04445 | 0,264869442 |
| Q7TNV0 | DEK | Protein DEK | 0,212517783 | 0,001626 | 0,040203758 |
| Q99L04 | DHRS1 | Dehydrogenase/reductase SDR family member 1 (EC 1.1.-.-) | -0,251148457 | 0,001751 | 0,042208398 |
| P84089 | ERH | Enhancer of rudimentary homolog (Mer) | 0,232110926 | 0,002898 | 0,060389411 |
| Q05816 | FABP5 | Fatty acid-binding protein 5 (Epidermal-type fatty acid-binding protein) (E-FABP) (Fatty acid-binding protein, epidermal) (Keratinocyte lipid-binding protein) (Psoriasis-associated fatty acid-binding protein homolog) (PA-FABP) | -0,498434268 | 0,000971 | 0,028033803 |
| P56959 | FUS | RNA-binding protein FUS (Protein pigpen) | 0,208726242 | 0,004947 | 0,082459629 |
| Q9D7X8 | GGCT | Gamma-glutamylcyclotransferase (EC 4.3.2.9) | 0,201325568 | 0,009614 | 0,120876193 |
| Q9QZF2 | GPC1 | Glypican-1 [Cleaved into: Secreted glypican-1] | -0,478386881 | 4,44E-05 | 0,003014374 |
| P17095 | HMGA1 | High mobility group protein HMG-I/HMG-Y (HMG-I(Y)) (High mobility group AT-hook protein 1) (High mobility group protein A1) | 0,375127582 | 0,048085 | 0,275606009 |
| P63158 | HMGB1 | High mobility group protein B1 (High mobility group protein 1) (HMG-1) | 0,317566129 | 0,001273 | 0,033751711 |
| O54879 | HMGB3 | High mobility group protein B3 (High mobility group protein 2a) (HMG-2a) (High mobility group protein 4) (HMG-4) | 0,30095374 | 0,015673 | 0,15819026 |
| Q91UZ5 | IMPA2 | Inositol monophosphatase 2 (IMP 2) (IMPase 2) (EC 3.1.3.25) (Inositol-1(or 4)-monophosphatase 2) (Myo-inositol monophosphatase A2) | 0,284004845 | 0,009129 | 0,118349458 |
| P21619 | LMNB2 | Lamin-B2 | -0,347676151 | 0,000504 | 0,017933802 |
| Q9D023 | MPC2 | Mitochondrial pyruvate carrier 2 (Brain protein 44) | -0,169701341 | 0,011392 | 0,13031752 |
| Q9R0M4 | PODXL | Podocalyxin (Podocalyxin-like protein 1) (PC) (PCLP-1) | 0,291146022 | 0,012639 | 0,139268033 |
| Q9D868 | PPIH | Peptidyl-prolyl cis-trans isomerase H (PPIase H) (EC 5.2.1.8) (Rotamase H) | 0,18006699 | 0,01425 | 0,149519386 |
| P28828 | PTPRM | Receptor-type tyrosine-protein phosphatase mu (Protein-tyrosine phosphatase mu) (R-PTP-mu) (EC 3.1.3.48) | -0,24196349 | 0,011181 | 0,12891725 |
| Q9D2C2 | SAAL1 | Protein SAAL1 (Synoviocyte proliferation-associated in collagen-induced arthritis protein 1) (SPACIA1) | 0,33496555 | 0,000141 | 0,00686712 |
| Q921M3 | SF3B3 | Splicing factor 3B subunit 3 (Pre-mRNA-splicing factor SF3b 130 kDa subunit) (SF3b130) (Spliceosome-associated protein 130) (SAP 130) | 0,188801864 | 0,004294 | 0,075457482 |
| Q8VIJ6 | SFPQ | Splicing factor, proline- and glutamine-rich (DNA-binding p52/p100 complex, 100 kDa subunit) (Polypyrimidine tract-binding protein-associated-splicing factor) (PSF) (PTB-associated-splicing factor) | 0,144120957 | 0,038163 | 0,247381426 |
| Q9CW03 | SMC3 | Structural maintenance of chromosomes protein 3 (SMC protein 3) (SMC-3) (Basement membrane-associated chondroitin proteoglycan) (Bamacan) (Chondroitin sulfate proteoglycan 6) (Chromosome segregation protein SmcD) (Mad member-interacting protein 1) | 0,160522207 | 0,017131 | 0,166477456 |
| Q62189 | SNRPA | U1 small nuclear ribonucleoprotein A (U1 snRNP A) (U1-A) (U1A) | 0,167604779 | 0,023813 | 0,196218478 |
| Q9D8U8 | SNX5 | Sorting nexin-5 | 0,156568583 | 0,031433 | 0,22363408 |
| Q08943 | SSRP1 | FACT complex subunit SSRP1 (Facilitates chromatin transcription complex subunit SSRP1) (Recombination signal sequence recognition protein 1) (Structure-specific recognition protein 1) (T160) | 0,21499255 | 0,0127 | 0,139268033 |

**Supplementary Data Table 7. List of total significantly altered proteins (p-value<0.05) in Physiologically aged humans**

| **Uniprot ID** | **Protein Symbol** | **Protein Description** | **Average Log2 Ratio** | ***p*-value** | **Adjusted *p*-value** |
| --- | --- | --- | --- | --- | --- |
| P13746 | 1A11 | HLA class I histocompatibility antigen, A-11 alpha chain (MHC class I antigen A*11) | 1,260876744 | 0,02082 | 0,234998667 |
| P49902 | 5NTC | Cytosolic purine 5'-nucleotidase (EC 3,1,3,5) (Cytosolic 5'-nucleotidase II) | -0,482197608 | 0,02833 | 0,257481581 |
| P56378 | 68MP | 6,8 kDa mitochondrial proteolipid | 0,384166739 | 0,04546 | 0,320857721 |
| P02763 | A1AG1 | Alpha-1-acid glycoprotein 1 (AGP 1) (Orosomucoid-1) (OMD 1) | 1,429845059 | 0,0127 | 0,199766061 |
| P19652 | A1AG2 | Alpha-1-acid glycoprotein 2 (AGP 2) (Orosomucoid-2) (OMD 2) | 1,247128837 | 0,0029 | 0,097030859 |
| P01009 | A1AT | Alpha-1-antitrypsin (Alpha-1 protease inhibitor) (Alpha-1-antiproteinase) (Serpin A1) [Cleaved into: Short peptide from AAT (SPAAT)] | 1,508778501 | 0,00062 | 0,043603785 |
| P04217 | A1BG | Alpha-1B-glycoprotein (Alpha-1-B glycoprotein) | 0,962183057 | 0,01479 | 0,211823261 |
| P02750 | A2GL | Leucine-rich alpha-2-glycoprotein (LRG) | 1,574039594 | 0,00114 | 0,059039382 |
| P01023 | A2MG | Alpha-2-macroglobulin (Alpha-2-M) (C3 and PZP-like alpha-2-macroglobulin domain-containing protein 5) | 0,987786924 | 0,00319 | 0,101469804 |
| Q9UDR5 | AASS | Alpha-aminoadipic semialdehyde synthase, mitochondrial (LKR/SDH) [Includes: Lysine ketoglutarate reductase (LKR) (LOR) (EC 1,5,1,8); Saccharopine dehydrogenase (SDH) (EC 1,5,1,9)] | -0,445354258 | 0,03904 | 0,299900797 |
| P33897 | ABCD1 | ATP-binding cassette sub-family D member 1 (Adrenoleukodystrophy protein) (ALDP) | -0,53973928 | 0,00267 | 0,097030859 |
| Q9NUQ8 | ABCF3 | ATP-binding cassette sub-family F member 3 | -0,560347613 | 0,0179 | 0,225054195 |
| P08910 | ABHD2 | Monoacylglycerol lipase ABHD2 (EC 3,1,1,23) (2-arachidonoylglycerol hydrolase) (Abhydrolase domain-containing protein 2) (Lung alpha/beta hydrolase 2) (Protein PHPS1-2) | -0,780055671 | 0,0236 | 0,236759134 |
| O94929 | ABLM3 | Actin-binding LIM protein 3 (abLIM-3) (Actin-binding LIM protein family member 3) | -0,50547184 | 0,04097 | 0,308530859 |
| Q15018 | ABRX2 | BRISC complex subunit Abraxas 2 (Abraxas brother protein 1) (Protein FAM175B) | 0,655674166 | 0,00135 | 0,063609836 |
| P07108 | ACBP | Acyl-CoA-binding protein (ACBP) (Diazepam-binding inhibitor) (DBI) (Endozepine) (EP) | 0,748212915 | 0,02684 | 0,251863828 |
| Q709F0 | ACD11 | Acyl-CoA dehydrogenase family member 11 (ACAD-11) (EC 1,3,99,-) | -0,370571559 | 0,02154 | 0,235752629 |
| P63261 | ACTG | Actin, cytoplasmic 2 (Gamma-actin) [Cleaved into: Actin, cytoplasmic 2, N-terminally processed] | 0,76296384 | 0,00139 | 0,064232864 |
| Q08043 | ACTN3 | Alpha-actinin-3 (Alpha-actinin skeletal muscle isoform 3) (F-actin cross-linking protein) | -0,528294414 | 0,02618 | 0,24857069 |
| P61163 | ACTZ | Alpha-centractin (Centractin) (ARP1) (Actin-RPV) (Centrosome-associated actin homolog) | 0,647385055 | 0,00641 | 0,142774153 |
| P11766 | ADHX | Alcohol dehydrogenase class-3 (EC 1,1,1,1) (Alcohol dehydrogenase 5) (Alcohol dehydrogenase class chi chain) (Alcohol dehydrogenase class-III) (Glutathione-dependent formaldehyde dehydrogenase) (FALDH) (FDH) (GSH-FDH) (EC 1,1,1,-) (S-(hydroxymethyl)glutathione dehydrogenase) (EC 1,1,1,284) | 0,340683982 | 0,02206 | 0,235814224 |
| Q15848 | ADIPO | Adiponectin (30 kDa adipocyte complement-related protein) (Adipocyte complement-related 30 kDa protein) (ACRP30) (Adipocyte, C1q and collagen domain-containing protein) (Adipose most abundant gene transcript 1 protein) (apM-1) (Gelatin-binding protein) | 0,878803239 | 0,03649 | 0,289691475 |
| Q15847 | ADIRF | Adipogenesis regulatory factor (Adipogenesis factor rich in obesity) (Adipose most abundant gene transcript 2 protein) (Adipose-specific protein 2) (apM-2) | 0,862238423 | 0,00034 | 0,030744046 |
| P12236 | ADT3 | ADP/ATP translocase 3 (ADP,ATP carrier protein 3) (ADP,ATP carrier protein, isoform T2) (ANT 2) (Adenine nucleotide translocator 3) (ANT 3) (Solute carrier family 25 member 6) [Cleaved into: ADP/ATP translocase 3, N-terminally processed] | 0,66051446 | 0,00087 | 0,048517846 |
| Q96SZ5 | AEDO | 2-aminoethanethiol dioxygenase (EC 1,13,11,19) (Cysteamine dioxygenase) | -0,511581413 | 0,03418 | 0,278696741 |
| P43652 | AFAM | Afamin (Alpha-albumin) (Alpha-Alb) | 0,903724745 | 0,0054 | 0,133124752 |
| Q09666 | AHNK | Neuroblast differentiation-associated protein AHNAK (Desmoyokin) | 0,520916815 | 0,00525 | 0,131177383 |
| Q02952 | AKA12 | A-kinase anchor protein 12 (AKAP-12) (A-kinase anchor protein 250 kDa) (AKAP 250) (Gravin) (Myasthenia gravis autoantigen) | 0,500243847 | 0,04511 | 0,320244891 |
| P02768 | ALBU | Serum albumin | 1,642716161 | 0,00037 | 0,031592167 |
| P05062 | ALDOB | Fructose-bisphosphate aldolase B (EC 4,1,2,13) (Liver-type aldolase) | -0,495392694 | 0,0109 | 0,188999161 |
| P15121 | ALDR | Aldose reductase (AR) (EC 1,1,1,21) (Aldehyde reductase) (Aldo-keto reductase family 1 member B1) | -0,536594506 | 0,00842 | 0,163422907 |
| P35858 | ALS | Insulin-like growth factor-binding protein complex acid labile subunit (ALS) | 0,482240842 | 0,04865 | 0,327613246 |
| P02760 | AMBP | Protein AMBP [Cleaved into: Alpha-1-microglobulin (Protein HC) (Alpha-1 microglycoprotein) (Complex-forming glycoprotein heterogeneous in charge); Inter-alpha-trypsin inhibitor light chain (ITI-LC) (Bikunin) (EDC1) (HI-30) (Uronic-acid-rich protein); Trypstatin] | 0,794353781 | 0,00027 | 0,025999051 |
| Q9UKU9 | ANGL2 | Angiopoietin-related protein 2 (Angiopoietin-like protein 2) | 0,823966302 | 0,02118 | 0,234998667 |
| Q12955 | ANK3 | Ankyrin-3 (ANK-3) (Ankyrin-G) | -0,256102825 | 0,04307 | 0,312814857 |
| P01008 | ANT3 | Antithrombin-III (ATIII) (Serpin C1) | 1,077267895 | 0,00185 | 0,0780729 |
| P04083 | ANXA1 | Annexin A1 (Annexin I) (Annexin-1) (Calpactin II) (Calpactin-2) (Chromobindin-9) (Lipocortin I) (Phospholipase A2 inhibitory protein) (p35) | 0,490768394 | 0,01411 | 0,207999951 |
| P08758 | ANXA5 | Annexin A5 (Anchorin CII) (Annexin V) (Annexin-5) (Calphobindin I) (CBP-I) (Endonexin II) (Lipocortin V) (Placental anticoagulant protein 4) (PP4) (Placental anticoagulant protein I) (PAP-I) (Thromboplastin inhibitor) (Vascular anticoagulant-alpha) (VAC-alpha) | 0,581033928 | 0,00127 | 0,062074957 |
| P21397 | AOFA | Amine oxidase [flavin-containing] A (EC 1,4,3,4) (Monoamine oxidase type A) (MAO-A) | 0,313509344 | 0,02302 | 0,236759134 |
| Q96CW1 | AP2M1 | AP-2 complex subunit mu (AP-2 mu chain) (Adaptin-mu2) (Adaptor protein complex AP-2 subunit mu) (Adaptor-related protein complex 2 subunit mu) (Clathrin assembly protein complex 2 mu medium chain) (Clathrin coat assembly protein AP50) (Clathrin coat-associated protein AP50) (HA2 50 kDa subunit) (Plasma membrane adaptor AP-2 50 kDa protein) | 0,373192578 | 0,02273 | 0,236759134 |
| P50583 | AP4A | Bis(5'-nucleosyl)-tetraphosphatase [asymmetrical] (EC 3,6,1,17) (Diadenosine 5',5'''-P1,P4-tetraphosphate asymmetrical hydrolase) (Ap4A hydrolase) (Ap4Aase) (Diadenosine tetraphosphatase) (Nucleoside diphosphate-linked moiety X motif 2) (Nudix motif 2) | -0,414437286 | 0,01654 | 0,215479037 |
| P02647 | APOA1 | Apolipoprotein A-I (Apo-AI) (ApoA-I) (Apolipoprotein A1) [Cleaved into: Proapolipoprotein A-I (ProapoA-I); Truncated apolipoprotein A-I (Apolipoprotein A-I(1-242))] | 1,806590499 | 0,00101 | 0,053902346 |
| P02652 | APOA2 | Apolipoprotein A-II (Apo-AII) (ApoA-II) (Apolipoprotein A2) [Cleaved into: Proapolipoprotein A-II (ProapoA-II); Truncated apolipoprotein A-II (Apolipoprotein A-II(1-76))] | 2,028411083 | 0,00014 | 0,017783943 |
| P04114 | APOB | Apolipoprotein B-100 (Apo B-100) [Cleaved into: Apolipoprotein B-48 (Apo B-48)] | 0,966778511 | 0,00733 | 0,153784738 |
| P02654 | APOC1 | Apolipoprotein C-I (Apo-CI) (ApoC-I) (Apolipoprotein C1) [Cleaved into: Truncated apolipoprotein C-I] | 1,114962027 | 0,00442 | 0,120465537 |
| P02656 | APOC3 | Apolipoprotein C-III (Apo-CIII) (ApoC-III) (Apolipoprotein C3) | 1,771271605 | 0,00381 | 0,112227954 |
| P05090 | APOD | Apolipoprotein D (Apo-D) (ApoD) | 0,784489539 | 0,00066 | 0,045425357 |
| P02649 | APOE | Apolipoprotein E (Apo-E) | 1,120302814 | 0,00061 | 0,043603785 |
| P02749 | APOH | Beta-2-glycoprotein 1 (APC inhibitor) (Activated protein C-binding protein) (Anticardiolipin cofactor) (Apolipoprotein H) (Apo-H) (Beta-2-glycoprotein I) (B2GPI) (Beta(2)GPI) | 1,065040132 | 0,00102 | 0,053902346 |
| O14791 | APOL1 | Apolipoprotein L1 (Apolipoprotein L) (Apo-L) (ApoL) (Apolipoprotein L-I) (ApoL-I) | 0,453673075 | 0,04968 | 0,329845885 |
| P10398 | ARAF | Serine/threonine-protein kinase A-Raf (EC 2,7,11,1) (Proto-oncogene A-Raf) (Proto-oncogene A-Raf-1) (Proto-oncogene Pks) | -0,490403188 | 0,00439 | 0,120430617 |
| Q92974 | ARHG2 | Rho guanine nucleotide exchange factor 2 (Guanine nucleotide exchange factor H1) (GEF-H1) (Microtubule-regulated Rho-GEF) (Proliferating cell nucleolar antigen p40) | -0,583834976 | 0,00283 | 0,097030859 |
| P04424 | ARLY | Argininosuccinate lyase (ASAL) (EC 4,3,2,1) (Arginosuccinase) | -0,362874577 | 0,02301 | 0,236759134 |
| P15848 | ARSB | Arylsulfatase B (ASB) (EC 3,1,6,12) (N-acetylgalactosamine-4-sulfatase) (G4S) | 0,69303988 | 0,01302 | 0,201422202 |
| O00192 | ARVC | Armadillo repeat protein deleted in velo-cardio-facial syndrome | -0,310152428 | 0,04774 | 0,325912134 |
| Q13510 | ASAH1 | Acid ceramidase (AC) (ACDase) (Acid CDase) (EC 3,5,1,23) (Acylsphingosine deacylase) (N-acylsphingosine amidohydrolase) (Putative 32 kDa heart protein) (PHP32) [Cleaved into: Acid ceramidase subunit alpha; Acid ceramidase subunit beta] | 1,046278962 | 0,00068 | 0,045663466 |
| O95671 | ASML | N-acetylserotonin O-methyltransferase-like protein (ASMTL) (EC 2,1,1,-) | -0,33482334 | 0,04322 | 0,313290058 |
| Q9BZE9 | ASPC1 | Tether containing UBX domain for GLUT4 (Alveolar soft part sarcoma chromosomal region candidate gene 1 protein) (Alveolar soft part sarcoma locus) (Renal papillary cell carcinoma protein 17) (UBX domain-containing protein 9) | -0,341668849 | 0,03015 | 0,263375577 |
| Q12797 | ASPH | Aspartyl/asparaginyl beta-hydroxylase (EC 1,14,11,16) (Aspartate beta-hydroxylase) (ASP beta-hydroxylase) (Peptide-aspartate beta-dioxygenase) | 0,314059673 | 0,01315 | 0,201757374 |
| P00966 | ASSY | Argininosuccinate synthase (EC 6,3,4,5) (Citrulline--aspartate ligase) | 0,510086283 | 0,0281 | 0,257481581 |
| P98196 | AT11A | Probable phospholipid-transporting ATPase IH (EC 3,6,3,1) (ATPase IS) (ATPase class VI type 11A) (P4-ATPase flippase complex alpha subunit ATP11A) | -0,328417455 | 0,03513 | 0,282929135 |
| P50993 | AT1A2 | Sodium/potassium-transporting ATPase subunit alpha-2 (Na(+)/K(+) ATPase alpha-2 subunit) (EC 3,6,3,9) (Sodium pump subunit alpha-2) | -0,43153144 | 0,0319 | 0,268193682 |
| Q7Z3C6 | ATG9A | Autophagy-related protein 9A (APG9-like 1) (mATG9) | -0,494132793 | 0,01925 | 0,230638173 |
| O00244 | ATOX1 | Copper transport protein ATOX1 (Metal transport protein ATX1) | 0,418101355 | 0,00754 | 0,156590759 |
| O75947 | ATP5H | ATP synthase subunit d, mitochondrial (ATPase subunit d) | 0,440441293 | 0,00809 | 0,163422907 |
| O75964 | ATP5L | ATP synthase subunit g, mitochondrial (ATPase subunit g) | 0,458003208 | 0,0227 | 0,236759134 |
| P30049 | ATPD | ATP synthase subunit delta, mitochondrial (ATP synthase F1 subunit delta) (F-ATPase delta subunit) | -0,281808449 | 0,04782 | 0,325912134 |
| O75173 | ATS4 | A disintegrin and metalloproteinase with thrombospondin motifs 4 (ADAM-TS 4) (ADAM-TS4) (ADAMTS-4) (EC 3,4,24,82) (ADMP-1) (Aggrecanase-1) | 1,987633652 | 1,74E-06 | 0,001705882 |
| P02730 | B3AT | Band 3 anion transport protein (Anion exchange protein 1) (AE 1) (Anion exchanger 1) (Solute carrier family 4 member 1) (CD antigen CD233) | 0,672634065 | 0,03893 | 0,299900797 |
| Q9NWV8 | BABA1 | BRISC and BRCA1-A complex member 1 (Mediator of RAP80 interactions and targeting subunit of 40 kDa) (New component of the BRCA1-A complex) | 0,333097705 | 0,01252 | 0,199766061 |
| O14874 | BCKD | [3-methyl-2-oxobutanoate dehydrogenase [lipoamide]] kinase, mitochondrial (EC 2,7,11,4) (Branched-chain alpha-ketoacid dehydrogenase kinase) (BCKD-kinase) (BCKDHKIN) | -0,391070553 | 0,01853 | 0,226221613 |
| Q10588 | BST1 | ADP-ribosyl cyclase/cyclic ADP-ribose hydrolase 2 (EC 3,2,2,6) (ADP-ribosyl cyclase 2) (Bone marrow stromal antigen 1) (BST-1) (Cyclic ADP-ribose hydrolase 2) (cADPr hydrolase 2) (CD antigen CD157) | 0,99804934 | 0,00244 | 0,094833427 |
| Q10589 | BST2 | Bone marrow stromal antigen 2 (BST-2) (HM1,24 antigen) (Tetherin) (CD antigen CD317) | -0,412220693 | 0,01227 | 0,199766061 |
| Q99622 | C10 | Protein C10 | 0,425941656 | 0,01533 | 0,212766683 |
| P02747 | C1QC | Complement C1q subcomponent subunit C | 0,527138549 | 0,03961 | 0,302225352 |
| P00736 | C1R | Complement C1r subcomponent (EC 3,4,21,41) (Complement component 1 subcomponent r) [Cleaved into: Complement C1r subcomponent heavy chain; Complement C1r subcomponent light chain] | 0,607492675 | 0,03697 | 0,29139789 |
| P04003 | C4BPA | C4b-binding protein alpha chain (C4bp) (Proline-rich protein) (PRP) | 0,692952509 | 0,04215 | 0,311743911 |
| Q99643 | C560 | Succinate dehydrogenase cytochrome b560 subunit, mitochondrial (Integral membrane protein CII-3) (QPs-1) (QPs1) (Succinate dehydrogenase complex subunit C) (Succinate-ubiquinone oxidoreductase cytochrome B large subunit) (CYBL) | 0,547856867 | 0,01263 | 0,199766061 |
| Q9BV19 | CA050 | Uncharacterized protein C1orf50 | 0,852670616 | 0,0013 | 0,062074957 |
| Q08289 | CACB2 | Voltage-dependent L-type calcium channel subunit beta-2 (CAB2) (Calcium channel voltage-dependent subunit beta 2) (Lambert-Eaton myasthenic syndrome antigen B) (MYSB) | -0,368978117 | 0,01498 | 0,212742239 |
| Q13137 | CACO2 | Calcium-binding and coiled-coil domain-containing protein 2 (Antigen nuclear dot 52 kDa protein) (Nuclear domain 10 protein NDP52) (Nuclear domain 10 protein 52) (Nuclear dot protein 52) | -0,428208627 | 0,04391 | 0,315790315 |
| Q5T440 | CAF17 | Putative transferase CAF17, mitochondrial (EC 2,1,-,-) (Iron-sulfur cluster assembly factor homolog) | -0,536603473 | 0,02352 | 0,236759134 |
| P00915 | CAH1 | Carbonic anhydrase 1 (EC 4,2,1,1) (Carbonate dehydratase I) (Carbonic anhydrase B) (CAB) (Carbonic anhydrase I) (CA-I) | 1,836704964 | 0,00083 | 0,047830967 |
| P00918 | CAH2 | Carbonic anhydrase 2 (EC 4,2,1,1) (Carbonate dehydratase II) (Carbonic anhydrase C) (CAC) (Carbonic anhydrase II) (CA-II) | 0,599801259 | 0,00755 | 0,156590759 |
| P22748 | CAH4 | Carbonic anhydrase 4 (EC 4,2,1,1) (Carbonate dehydratase IV) (Carbonic anhydrase IV) (CA-IV) | -0,395811361 | 0,0236 | 0,236759134 |
| P27797 | CALR | Calreticulin (CRP55) (Calregulin) (Endoplasmic reticulum resident protein 60) (ERp60) (HACBP) (grp60) | 0,604560606 | 0,01534 | 0,212766683 |
| P27824 | CALX | Calnexin (IP90) (Major histocompatibility complex class I antigen-binding protein p88) (p90) | 0,338086713 | 0,03727 | 0,292457436 |
| Q01518 | CAP1 | Adenylyl cyclase-associated protein 1 (CAP 1) | 0,458301108 | 0,04549 | 0,320857721 |
| Q14444 | CAPR1 | Caprin-1 (Cell cycle-associated protein 1) (Cytoplasmic activation- and proliferation-associated protein 1) (GPI-anchored membrane protein 1) (GPI-anchored protein p137) (GPI-p137) (p137GPI) (Membrane component chromosome 11 surface marker 1) (RNA granule protein 105) | 0,449056132 | 0,00836 | 0,163422907 |
| P07858 | CATB | Cathepsin B (EC 3,4,22,1) (APP secretase) (APPS) (Cathepsin B1) [Cleaved into: Cathepsin B light chain; Cathepsin B heavy chain] | 0,756734454 | 0,00671 | 0,146096973 |
| P07339 | CATD | Cathepsin D (EC 3,4,23,5) [Cleaved into: Cathepsin D light chain; Cathepsin D heavy chain] | 0,804702437 | 0,00084 | 0,047830967 |
| Q9UBR2 | CATZ | Cathepsin Z (EC 3,4,18,1) (Cathepsin P) (Cathepsin X) | 0,804368004 | 0,00292 | 0,097030859 |
| Q03135 | CAV1 | Caveolin-1 | 0,445369389 | 0,0288 | 0,257900048 |
| O95810 | CAVN2 | Caveolae-associated protein 2 (Cavin-2) (PS-p68) (Phosphatidylserine-binding protein) (Serum deprivation-response protein) | 0,412990618 | 0,01537 | 0,212766683 |
| P08185 | CBG | Corticosteroid-binding globulin (CBG) (Serpin A6) (Transcortin) | 2,050910631 | 8,32E-05 | 0,013863419 |
| Q9GZT6 | CC90B | Coiled-coil domain-containing protein 90B, mitochondrial | 0,403825675 | 0,01297 | 0,201422202 |
| Q96ER9 | CCD51 | Coiled-coil domain-containing protein 51 | -0,523703763 | 0,00729 | 0,153784738 |
| Q7Z6B0 | CCD91 | Coiled-coil domain-containing protein 91 (GGA-binding partner) (p56 accessory protein) | 0,371499141 | 0,04089 | 0,308530859 |
| P48509 | CD151 | CD151 antigen (GP27) (Membrane glycoprotein SFA-1) (Platelet-endothelial tetraspan antigen 3) (PETA-3) (Tetraspanin-24) (Tspan-24) (CD antigen CD151) | 0,403424917 | 0,01625 | 0,213209762 |
| P16671 | CD36 | Platelet glycoprotein 4 (Fatty acid translocase) (FAT) (Glycoprotein IIIb) (GPIIIB) (Leukocyte differentiation antigen CD36) (PAS IV) (PAS-4) (Platelet collagen receptor) (Platelet glycoprotein IV) (GPIV) (Thrombospondin receptor) (CD antigen CD36) | 0,755066517 | 0,00276 | 0,097030859 |
| Q08722 | CD47 | Leukocyte surface antigen CD47 (Antigenic surface determinant protein OA3) (Integrin-associated protein) (IAP) (Protein MER6) (CD antigen CD47) | 0,285602869 | 0,0439 | 0,315790315 |
| P60033 | CD81 | CD81 antigen (26 kDa cell surface protein TAPA-1) (Target of the antiproliferative antibody 1) (Tetraspanin-28) (Tspan-28) (CD antigen CD81) | 1,322755108 | 5,33E-05 | 0,01161673 |
| P21926 | CD9 | CD9 antigen (5H9 antigen) (Cell growth-inhibiting gene 2 protein) (Leukocyte antigen MIC3) (Motility-related protein) (MRP-1) (Tetraspanin-29) (Tspan-29) (p24) (CD antigen CD9) | 0,574663602 | 0,0042 | 0,115915698 |
| Q6P1J9 | CDC73 | Parafibromin (Cell division cycle protein 73 homolog) (Hyperparathyroidism 2 protein) | -0,315019618 | 0,01562 | 0,212766683 |
| Q07002 | CDK18 | Cyclin-dependent kinase 18 (EC 2,7,11,22) (Cell division protein kinase 18) (PCTAIRE-motif protein kinase 3) (Serine/threonine-protein kinase PCTAIRE-3) | -1,111997505 | 0,0001 | 0,014543495 |
| Q711Q0 | CEFIP | Cardiac-enriched FHL2-interacting protein | -0,30961485 | 0,04115 | 0,309187411 |
| Q7Z7K6 | CENPV | Centromere protein V (CENP-V) (Nuclear protein p30) (Proline-rich protein 6) | -0,456386613 | 0,01755 | 0,2249457 |
| P00450 | CERU | Ceruloplasmin (EC 1,16,3,1) (Ferroxidase) | 0,743886599 | 0,01222 | 0,199766061 |
| P00751 | CFAB | Complement factor B (EC 3,4,21,47) (C3/C5 convertase) (Glycine-rich beta glycoprotein) (GBG) (PBF2) (Properdin factor B) [Cleaved into: Complement factor B Ba fragment; Complement factor B Bb fragment] | 0,73366418 | 0,01923 | 0,230638173 |
| P08603 | CFAH | Complement factor H (H factor 1) | 0,741973499 | 0,00811 | 0,163422907 |
| P61604 | CH10 | 10 kDa heat shock protein, mitochondrial (Hsp10) (10 kDa chaperonin) (Chaperonin 10) (CPN10) (Early-pregnancy factor) (EPF) | 0,618752905 | 0,02032 | 0,234761058 |
| P10809 | CH60 | 60 kDa heat shock protein, mitochondrial (EC 3,6,4,9) (60 kDa chaperonin) (Chaperonin 60) (CPN60) (Heat shock protein 60) (HSP-60) (Hsp60) (HuCHA60) (Mitochondrial matrix protein P1) (P60 lymphocyte protein) | 0,522361025 | 0,00291 | 0,097030859 |
| Q14839 | CHD4 | Chromodomain-helicase-DNA-binding protein 4 (CHD-4) (EC 3,6,4,12) (ATP-dependent helicase CHD4) (Mi-2 autoantigen 218 kDa protein) (Mi2-beta) | -0,324095898 | 0,04099 | 0,308530859 |
| O75339 | CILP1 | Cartilage intermediate layer protein 1 (CILP-1) (Cartilage intermediate-layer protein) [Cleaved into: Cartilage intermediate layer protein 1 C1; Cartilage intermediate layer protein 1 C2] | 1,672519445 | 2,49E-07 | 0,000489228 |
| Q15642 | CIP4 | Cdc42-interacting protein 4 (Protein Felic) (Salt tolerant protein) (hSTP) (Thyroid receptor-interacting protein 10) (TR-interacting protein 10) (TRIP-10) | -0,372182653 | 0,01999 | 0,233319483 |
| Q14011 | CIRBP | Cold-inducible RNA-binding protein (A18 hnRNP) (Glycine-rich RNA-binding protein CIRP) | -0,381600104 | 0,01508 | 0,212742239 |
| Q9NZ45 | CISD1 | CDGSH iron-sulfur domain-containing protein 1 (MitoNEET) | 0,48701065 | 0,0026 | 0,097030859 |
| Q8N5K1 | CISD2 | CDGSH iron-sulfur domain-containing protein 2 (Endoplasmic reticulum intermembrane small protein) (MitoNEET-related 1 protein) (Miner1) (Nutrient-deprivation autophagy factor-1) (NAF-1) | 0,704247206 | 0,00079 | 0,047747284 |
| Q7Z460 | CLAP1 | CLIP-associating protein 1 (Cytoplasmic linker-associated protein 1) (Multiple asters homolog 1) (Protein Orbit homolog 1) (hOrbit1) | -0,355471254 | 0,02439 | 0,241532022 |
| Q86T13 | CLC14 | C-type lectin domain family 14 member A (Epidermal growth factor receptor 5) (EGFR-5) | -0,574261081 | 0,00252 | 0,096013416 |
| P51797 | CLCN6 | Chloride transport protein 6 (Chloride channel protein 6) (ClC-6) | 0,646051305 | 0,01391 | 0,20665294 |
| O00299 | CLIC1 | Chloride intracellular channel protein 1 (Chloride channel ABP) (Nuclear chloride ion channel 27) (NCC27) (Regulatory nuclear chloride ion channel protein) (hRNCC) | 0,619236674 | 0,00242 | 0,094833427 |
| P10909 | CLUS | Clusterin (Aging-associated gene 4 protein) (Apolipoprotein J) (Apo-J) (Complement cytolysis inhibitor) (CLI) (Complement-associated protein SP-40,40) (Ku70-binding protein 1) (NA1/NA2) (Testosterone-repressed prostate message 2) (TRPM-2) [Cleaved into: Clusterin beta chain (ApoJalpha) (Complement cytolysis inhibitor a chain); Clusterin alpha chain (ApoJbeta) (Complement cytolysis inhibitor b chain)] | 0,923701516 | 0,02215 | 0,235814224 |
| Q9UJS0 | CMC2 | Calcium-binding mitochondrial carrier protein Aralar2 (Citrin) (Mitochondrial aspartate glutamate carrier 2) (Solute carrier family 25 member 13) | -0,263263971 | 0,04202 | 0,311743911 |
| P01024 | CO3 | Complement C3 (C3 and PZP-like alpha-2-macroglobulin domain-containing protein 1) [Cleaved into: Complement C3 beta chain; C3-beta-c (C3bc); Complement C3 alpha chain; C3a anaphylatoxin; Acylation stimulating protein (ASP) (C3adesArg); Complement C3b alpha' chain; Complement C3c alpha' chain fragment 1; Complement C3dg fragment; Complement C3g fragment; Complement C3d fragment; Complement C3f fragment; Complement C3c alpha' chain fragment 2] | 0,651511818 | 0,01621 | 0,213209762 |
| P0C0L5 | CO4B | Complement C4-B (Basic complement C4) (C3 and PZP-like alpha-2-macroglobulin domain-containing protein 3) [Cleaved into: Complement C4 beta chain; Complement C4-B alpha chain; C4a anaphylatoxin; C4b-B; C4d-B; Complement C4 gamma chain] | 1,10770045 | 0,00483 | 0,126093358 |
| P13671 | CO6 | Complement component C6 | 0,793996826 | 0,01418 | 0,208277421 |
| P12109 | CO6A1 | Collagen alpha-1(VI) chain | 0,356741062 | 0,03173 | 0,268193682 |
| P12111 | CO6A3 | Collagen alpha-3(VI) chain | 0,258024477 | 0,03862 | 0,298767304 |
| A6NMZ7 | CO6A6 | Collagen alpha-6(VI) chain | 0,644596787 | 0,04617 | 0,32449905 |
| P10643 | CO7 | Complement component C7 | 0,78325471 | 0,02041 | 0,234761058 |
| P07357 | CO8A | Complement component C8 alpha chain (Complement component 8 subunit alpha) | 0,975578986 | 0,01518 | 0,212766683 |
| P27658 | CO8A1 | Collagen alpha-1(VIII) chain (Endothelial collagen) [Cleaved into: Vastatin] | 1,67279764 | 6,05E-05 | 0,011864953 |
| P07358 | CO8B | Complement component C8 beta chain (Complement component 8 subunit beta) | 0,953436751 | 0,02725 | 0,253562451 |
| P02748 | CO9 | Complement component C9 [Cleaved into: Complement component C9a; Complement component C9b] | 1,346748393 | 0,00178 | 0,075678131 |
| O75128 | COBL | Protein cordon-bleu | -0,363803656 | 0,0159 | 0,213209762 |
| P23528 | COF1 | Cofilin-1 (18 kDa phosphoprotein) (p18) (Cofilin, non-muscle isoform) | 0,539563628 | 0,0217 | 0,235752629 |
| P39059 | COFA1 | Collagen alpha-1(XV) chain [Cleaved into: Restin (Endostatin-XV) (Related to endostatin) (Restin-I); Restin-2 (Restin-II); Restin-3 (Restin-III); Restin-4 (Restin-IV)] | -0,26116798 | 0,049 | 0,327955691 |
| Q96P44 | COLA1 | Collagen alpha-1(XXI) chain | -0,634529931 | 0,01235 | 0,199766061 |
| Q6QEF8 | CORO6 | Coronin-6 (Coronin-like protein E) (Clipin-E) | -0,426497498 | 0,03248 | 0,270276513 |
| P10606 | COX5B | Cytochrome c oxidase subunit 5B, mitochondrial (Cytochrome c oxidase polypeptide Vb) | 0,372305995 | 0,04286 | 0,312451384 |
| P15954 | COX7C | Cytochrome c oxidase subunit 7C, mitochondrial (Cytochrome c oxidase polypeptide VIIc) | 0,613554447 | 0,02848 | 0,257481581 |
| Q7Z7K0 | COXM1 | COX assembly mitochondrial protein homolog (Cmc1p) | 0,369108558 | 0,0318 | 0,268193682 |
| P22792 | CPN2 | Carboxypeptidase N subunit 2 (Carboxypeptidase N 83 kDa chain) (Carboxypeptidase N large subunit) (Carboxypeptidase N polypeptide 2) (Carboxypeptidase N regulatory subunit) | 1,093173004 | 0,0089 | 0,169218477 |
| Q9BRF8 | CPPED | Serine/threonine-protein phosphatase CPPED1 (EC 3,1,3,16) (Calcineurin-like phosphoesterase domain-containing protein 1) (Complete S-transactivated protein 1) | 0,392100816 | 0,01796 | 0,225054195 |
| Q9H8M1 | CQ10B | Coenzyme Q-binding protein COQ10 homolog B, mitochondrial | 0,353389623 | 0,04275 | 0,312246058 |
| P23508 | CRCM | Colorectal mutant cancer protein (Protein MCC) | -0,424520602 | 0,0134 | 0,201757374 |
| P46109 | CRKL | Crk-like protein | 0,385552254 | 0,00593 | 0,138394519 |
| Q5TZA2 | CROCC | Rootletin (Ciliary rootlet coiled-coil protein) | -1,40271798 | 0,00552 | 0,134510312 |
| Q9Y534 | CSDC2 | Cold shock domain-containing protein C2 (RNA-binding protein PIPPin) | -0,434559997 | 0,02329 | 0,236759134 |
| Q4G0I0 | CSMT1 | Protein CCSMST1 | 0,531734623 | 0,0291 | 0,259726687 |
| Q99627 | CSN8 | COP9 signalosome complex subunit 8 (SGN8) (Signalosome subunit 8) (COP9 homolog) (hCOP9) (JAB1-containing signalosome subunit 8) | -0,284706717 | 0,04258 | 0,311743911 |
| P0CG12 | CTF8A | Chromosome transmission fidelity protein 8 homolog isoform 2 (Decreased expression in renal and prostate cancer protein) | -0,346404122 | 0,01541 | 0,212766683 |
| Q8IWA5 | CTL2 | Choline transporter-like protein 2 (Solute carrier family 44 member 2) | -0,392471605 | 0,01273 | 0,199766061 |
| O60716 | CTND1 | Catenin delta-1 (Cadherin-associated Src substrate) (CAS) (p120 catenin) (p120(ctn)) (p120(cas)) | -0,446617664 | 0,02131 | 0,234998667 |
| Q9P013 | CWC15 | Spliceosome-associated protein CWC15 homolog | -0,357410347 | 0,01972 | 0,23220076 |
| P99999 | CYC | Cytochrome c | 0,388652328 | 0,02458 | 0,241832734 |
| Q15438 | CYH1 | Cytohesin-1 (PH, SEC7 and coiled-coil domain-containing protein 1) (SEC7 homolog B2-1) | -0,369505283 | 0,03761 | 0,293869001 |
| P98082 | DAB2 | Disabled homolog 2 (Adaptor molecule disabled-2) (Differentially expressed in ovarian carcinoma 2) (DOC-2) (Differentially-expressed protein 2) | 0,549359821 | 0,03538 | 0,284308771 |
| Q8TEB1 | DCA11 | DDB1- and CUL4-associated factor 11 (WD repeat-containing protein 23) | -0,324604255 | 0,03364 | 0,27485911 |
| Q96C86 | DCPS | m7GpppX diphosphatase (EC 3,6,1,59) (DCS-1) (Decapping scavenger enzyme) (Hint-related 7meGMP-directed hydrolase) (Histidine triad nucleotide-binding protein 5) (Histidine triad protein member 5) (HINT-5) (Scavenger mRNA-decapping enzyme DcpS) | -0,396526265 | 0,00919 | 0,172373188 |
| Q7L014 | DDX46 | Probable ATP-dependent RNA helicase DDX46 (EC 3,6,4,13) (DEAD box protein 46) (PRP5 homolog) | -0,280566913 | 0,03024 | 0,263556732 |
| P26196 | DDX6 | Probable ATP-dependent RNA helicase DDX6 (EC 3,6,4,13) (ATP-dependent RNA helicase p54) (DEAD box protein 6) (Oncogene RCK) | -0,571344508 | 0,01366 | 0,204507527 |
| O75064 | DEN4B | DENN domain-containing protein 4B | -0,390986992 | 0,02542 | 0,243710647 |
| Q53GQ0 | DHB12 | Very-long-chain 3-oxoacyl-CoA reductase (EC 1,1,1,330) (17-beta-hydroxysteroid dehydrogenase 12) (17-beta-HSD 12) (3-ketoacyl-CoA reductase) (KAR) (Estradiol 17-beta-dehydrogenase 12) (EC 1,1,1,62) (Short chain dehydrogenase/reductase family 12C member 1) | 0,67246616 | 0,00052 | 0,040899265 |
| Q9UBM7 | DHCR7 | 7-dehydrocholesterol reductase (7-DHC reductase) (EC 1,3,1,21) (Putative sterol reductase SR-2) (Sterol Delta(7)-reductase) | 0,568032234 | 0,00521 | 0,13089915 |
| P00367 | DHE3 | Glutamate dehydrogenase 1, mitochondrial (GDH 1) (EC 1,4,1,3) | -0,404767441 | 0,00938 | 0,173604714 |
| Q9Y394 | DHRS7 | Dehydrogenase/reductase SDR family member 7 (EC 1,1,-,-) (Retinal short-chain dehydrogenase/reductase 4) (retSDR4) (Short chain dehydrogenase/reductase family 34C member 1) | 0,312856544 | 0,03734 | 0,292457436 |
| O60231 | DHX16 | Pre-mRNA-splicing factor ATP-dependent RNA helicase DHX16 (EC 3,6,4,13) (ATP-dependent RNA helicase #3) (DEAH-box protein 16) | -0,477168117 | 0,00726 | 0,153784738 |
| Q9UNQ2 | DIM1 | Probable dimethyladenosine transferase (EC 2,1,1,183) (DIM1 dimethyladenosine transferase 1 homolog) (DIM1 dimethyladenosine transferase 1-like) (Probable 18S rRNA (adenine(1779)-N(6)/adenine(1780)-N(6))-dimethyltransferase) (Probable 18S rRNA dimethylase) (Probable S-adenosylmethionine-6-N',N'-adenosyl(rRNA) dimethyltransferase) | 0,416187181 | 0,01986 | 0,232646998 |
| Q9P265 | DIP2B | Disco-interacting protein 2 homolog B (DIP2 homolog B) | -0,322335619 | 0,01858 | 0,226221613 |
| Q96LL9 | DJC30 | DnaJ homolog subfamily C member 30 (Williams-Beuren syndrome chromosomal region 18 protein) | 0,328444185 | 0,04242 | 0,311743911 |
| O60832 | DKC1 | H/ACA ribonucleoprotein complex subunit DKC1 (EC 5,4,99,-) (CBF5 homolog) (Dyskerin) (Nopp140-associated protein of 57 kDa) (Nucleolar protein NAP57) (Nucleolar protein family A member 4) (snoRNP protein DKC1) | 0,529885662 | 0,00485 | 0,126093358 |
| Q12959 | DLG1 | Disks large homolog 1 (Synapse-associated protein 97) (SAP-97) (SAP97) (hDlg) | -0,397244988 | 0,03271 | 0,270635276 |
| Q9H3Z4 | DNJC5 | DnaJ homolog subfamily C member 5 (Ceroid-lipofuscinosis neuronal protein 4) (Cysteine string protein) (CSP) | 0,401032049 | 0,01964 | 0,23220076 |
| Q9ULA0 | DNPEP | Aspartyl aminopeptidase (EC 3,4,11,21) | -0,335114261 | 0,03153 | 0,268193682 |
| Q9NY33 | DPP3 | Dipeptidyl peptidase 3 (EC 3,4,14,4) (Dipeptidyl aminopeptidase III) (Dipeptidyl arylamidase III) (Dipeptidyl peptidase III) (DPP III) (Enkephalinase B) | 0,642216101 | 0,01257 | 0,199766061 |
| Q02487 | DSC2 | Desmocollin-2 (Cadherin family member 2) (Desmocollin-3) (Desmosomal glycoprotein II) (Desmosomal glycoprotein III) | -0,405674269 | 0,00975 | 0,176207469 |
| P51452 | DUS3 | Dual specificity protein phosphatase 3 (EC 3,1,3,16) (EC 3,1,3,48) (Dual specificity protein phosphatase VHR) (Vaccinia H1-related phosphatase) (VHR) | -0,382913835 | 0,04184 | 0,311743911 |
| Q96FJ2 | DYL2 | Dynein light chain 2, cytoplasmic (8 kDa dynein light chain b) (DLC8b) (Dynein light chain LC8-type 2) | 0,663558726 | 0,00519 | 0,13089915 |
| Q13011 | ECH1 | Delta(3,5)-Delta(2,4)-dienoyl-CoA isomerase, mitochondrial (EC 5,3,3,-) | 0,35949895 | 0,02246 | 0,236759134 |
| Q9NTX5 | ECHD1 | Ethylmalonyl-CoA decarboxylase (EC 4,1,1,94) (Enoyl-CoA hydratase domain-containing protein 1) (Methylmalonyl-CoA decarboxylase) (MMCD) (EC 4,1,1,41) | 0,458557062 | 0,00393 | 0,114007879 |
| Q9BUP0 | EFHD1 | EF-hand domain-containing protein D1 (EF-hand domain-containing protein 1) (Swiprosin-2) | 0,637054308 | 0,02344 | 0,236759134 |
| Q7Z2Z2 | EFL1 | Elongation factor-like GTPase 1 (Elongation factor Tu GTP-binding domain-containing protein 1) (Elongation factor-like 1) (Protein FAM42A) | -0,389387437 | 0,0263 | 0,248649026 |
| Q9GZT9 | EGLN1 | Egl nine homolog 1 (EC 1,14,11,29) (Hypoxia-inducible factor prolyl hydroxylase 2) (HIF-PH2) (HIF-prolyl hydroxylase 2) (HPH-2) (Prolyl hydroxylase domain-containing protein 2) (PHD2) (SM-20) | -0,35167939 | 0,04822 | 0,326654608 |
| Q8N3D4 | EH1L1 | EH domain-binding protein 1-like protein 1 | -0,556334462 | 0,00626 | 0,141663963 |
| P49770 | EI2BB | Translation initiation factor eIF-2B subunit beta (S20I15) (S20III15) (eIF-2B GDP-GTP exchange factor subunit beta) | -0,278862668 | 0,02547 | 0,243710647 |
| O75822 | EIF3J | Eukaryotic translation initiation factor 3 subunit J (eIF3j) (Eukaryotic translation initiation factor 3 subunit 1) (eIF-3-alpha) (eIF3 p35) | 0,501739851 | 0,01372 | 0,204641547 |
| P15502 | ELN | Elastin (Tropoelastin) | 1,673513344 | 0,00044 | 0,036639722 |
| O94919 | ENDD1 | Endonuclease domain-containing 1 protein (EC 3,1,30,-) | 0,364804258 | 0,02811 | 0,257481581 |
| P13929 | ENOB | Beta-enolase (EC 4,2,1,11) (2-phospho-D-glycerate hydro-lyase) (Enolase 3) (Muscle-specific enolase) (MSE) (Skeletal muscle enolase) | -0,475918719 | 0,02644 | 0,24931612 |
| Q5NDL2 | EOGT | EGF domain-specific O-linked N-acetylglucosamine transferase (EC 2,4,1,255) (Extracellular O-linked N-acetylglucosamine transferase) | -0,489820753 | 0,03203 | 0,268193682 |
| Q9UM22 | EPDR1 | Mammalian ependymin-related protein 1 (MERP-1) (Upregulated in colorectal cancer gene 1 protein) | 0,743055124 | 0,00653 | 0,143094484 |
| P30040 | ERP29 | Endoplasmic reticulum resident protein 29 (ERp29) (Endoplasmic reticulum resident protein 28) (ERp28) (Endoplasmic reticulum resident protein 31) (ERp31) | 0,490600942 | 0,00201 | 0,082556156 |
| Q96AP7 | ESAM | Endothelial cell-selective adhesion molecule | -0,342675457 | 0,02474 | 0,241832734 |
| P23141 | EST1 | Liver carboxylesterase 1 (Acyl-coenzyme A:cholesterol acyltransferase) (ACAT) (Brain carboxylesterase hBr1) (Carboxylesterase 1) (CE-1) (hCE-1) (EC 3,1,1,1) (Cocaine carboxylesterase) (Egasyn) (HMSE) (Methylumbelliferyl-acetate deacetylase 1) (EC 3,1,1,56) (Monocyte/macrophage serine esterase) (Retinyl ester hydrolase) (REH) (Serine esterase 1) (Triacylglycerol hydrolase) (TGH) | 0,880335242 | 0,01122 | 0,192078217 |
| P50502 | F10A1 | Hsc70-interacting protein (Hip) (Aging-associated protein 2) (Progesterone receptor-associated p48 protein) (Protein FAM10A1) (Putative tumor suppressor ST13) (Renal carcinoma antigen NY-REN-33) (Suppression of tumorigenicity 13 protein) | 0,235010369 | 0,04866 | 0,327613246 |
| Q96ND0 | F210A | Protein FAM210A | 0,629111617 | 0,00366 | 0,110182266 |
| Q9H0Q0 | FA49A | Protein FAM49A | 0,297379403 | 0,04251 | 0,311743911 |
| Q52LJ0 | FA98B | Protein FAM98B | -0,310262485 | 0,04744 | 0,325912134 |
| P15090 | FABP4 | Fatty acid-binding protein, adipocyte (Adipocyte lipid-binding protein) (ALBP) (Adipocyte-type fatty acid-binding protein) (A-FABP) (AFABP) (Fatty acid-binding protein 4) | 0,80546444 | 0,00792 | 0,161852265 |
| Q05397 | FAK1 | Focal adhesion kinase 1 (FADK 1) (EC 2,7,10,2) (Focal adhesion kinase-related nonkinase) (FRNK) (Protein phosphatase 1 regulatory subunit 71) (PPP1R71) (Protein-tyrosine kinase 2) (p125FAK) (pp125FAK) | -0,82055621 | 0,0001 | 0,014543495 |
| Q7L8L6 | FAKD5 | FAST kinase domain-containing protein 5, mitochondrial | -0,41712406 | 0,02851 | 0,257481581 |
| P49327 | FAS | Fatty acid synthase (EC 2,3,1,85) [Includes: [Acyl-carrier-protein] S-acetyltransferase (EC 2,3,1,38); [Acyl-carrier-protein] S-malonyltransferase (EC 2,3,1,39); 3-oxoacyl-[acyl-carrier-protein] synthase (EC 2,3,1,41); 3-oxoacyl-[acyl-carrier-protein] reductase (EC 1,1,1,100); 3-hydroxyacyl-[acyl-carrier-protein] dehydratase (EC 4,2,1,59); Enoyl-[acyl-carrier-protein] reductase (EC 1,3,1,39); Oleoyl-[acyl-carrier-protein] hydrolase (EC 3,1,2,14)] | 0,812383438 | 0,03249 | 0,270276513 |
| Q9UBX5 | FBLN5 | Fibulin-5 (FIBL-5) (Developmental arteries and neural crest EGF-like protein) (Dance) (Urine p50 protein) (UP50) | -0,585395887 | 0,00306 | 0,099881772 |
| Q5T0N5 | FBP1L | Formin-binding protein 1-like (Transducer of Cdc42-dependent actin assembly protein 1) (Toca-1) | -0,326374902 | 0,04247 | 0,311743911 |
| Q9UH90 | FBX40 | F-box only protein 40 (Muscle disease-related protein) | -0,60118436 | 0,01132 | 0,193012395 |
| Q96AC1 | FERM2 | Fermitin family homolog 2 (Kindlin-2) (Mitogen-inducible gene 2 protein) (MIG-2) (Pleckstrin homology domain-containing family C member 1) (PH domain-containing family C member 1) | -0,251885497 | 0,04434 | 0,316650023 |
| P02765 | FETUA | Alpha-2-HS-glycoprotein (Alpha-2-Z-globulin) (Ba-alpha-2-glycoprotein) (Fetuin-A) [Cleaved into: Alpha-2-HS-glycoprotein chain A; Alpha-2-HS-glycoprotein chain B] | 1,250524943 | 0,00152 | 0,069315142 |
| Q96M96 | FGD4 | FYVE, RhoGEF and PH domain-containing protein 4 (Actin filament-binding protein frabin) (FGD1-related F-actin-binding protein) (Zinc finger FYVE domain-containing protein 6) | -0,417150759 | 0,01332 | 0,201757374 |
| Q2V2M9 | FHOD3 | FH1/FH2 domain-containing protein 3 (Formactin-2) (Formin homolog overexpressed in spleen 2) (hFHOS2) | -0,387814172 | 0,01063 | 0,188143812 |
| P02675 | FIBB | Fibrinogen beta chain [Cleaved into: Fibrinopeptide B; Fibrinogen beta chain] | 0,994839917 | 0,03822 | 0,29744878 |
| P20930 | FILA | Filaggrin | -0,591668771 | 0,0397 | 0,302349948 |
| Q9Y3D6 | FIS1 | Mitochondrial fission 1 protein (FIS1 homolog) (hFis1) (Tetratricopeptide repeat protein 11) (TPR repeat protein 11) | 0,364544198 | 0,01825 | 0,226221613 |
| A5D6W6 | FITM1 | Fat storage-inducing transmembrane protein 1 (Fat-inducing protein 1) | -0,320539885 | 0,02187 | 0,235814224 |
| Q8N6M3 | FITM2 | Fat storage-inducing transmembrane protein 2 (Fat-inducing protein 2) | 0,467156688 | 0,01559 | 0,212766683 |
| P21333 | FLNA | Filamin-A (FLN-A) (Actin-binding protein 280) (ABP-280) (Alpha-filamin) (Endothelial actin-binding protein) (Filamin-1) (Non-muscle filamin) | 0,592799722 | 0,03149 | 0,268193682 |
| O75955 | FLOT1 | Flotillin-1 | -0,411089803 | 0,02728 | 0,253562451 |
| Q16595 | FRDA | Frataxin, mitochondrial (EC 1,16,3,1) (Friedreich ataxia protein) (Fxn) [Cleaved into: Frataxin intermediate form (i-FXN); Frataxin(56-210) (m56-FXN); Frataxin(78-210) (d-FXN) (m78-FXN); Frataxin mature form (Frataxin(81-210)) (m81-FXN)] | 0,530129089 | 0,00948 | 0,174607283 |
| P02794 | FRIH | Ferritin heavy chain (Ferritin H subunit) (EC 1,16,3,1) (Cell proliferation-inducing gene 15 protein) [Cleaved into: Ferritin heavy chain, N-terminally processed] | 0,881109519 | 0,0426 | 0,311743911 |
| Q9BZ67 | FRMD8 | FERM domain-containing protein 8 | -0,405506422 | 0,01846 | 0,226221613 |
| A1L4K1 | FSD2 | Fibronectin type III and SPRY domain-containing protein 2 (SPRY domain-containing protein 1) | -0,41541852 | 0,01343 | 0,201757374 |
| Q9BWH2 | FUND2 | FUN14 domain-containing protein 2 (Cervical cancer proto-oncogene 3 protein) (HCC-3) (Hepatitis C virus core-binding protein 6) | 0,599998078 | 0,00923 | 0,172373188 |
| P11413 | G6PD | Glucose-6-phosphate 1-dehydrogenase (G6PD) (EC 1,1,1,49) | 0,650272556 | 0,023 | 0,236759134 |
| P50440 | GATM | Glycine amidinotransferase, mitochondrial (EC 2,1,4,1) (L-arginine:glycine amidinotransferase) (Transamidinase) | 1,088649492 | 0,00054 | 0,041879292 |
| Q9BSJ2 | GCP2 | Gamma-tubulin complex component 2 (GCP-2) (hGCP2) (Gamma-ring complex protein 103 kDa) (h103p) (hGrip103) (Spindle pole body protein Spc97 homolog) (hSpc97) | 0,493179617 | 0,00493 | 0,126430808 |
| P23434 | GCSH | Glycine cleavage system H protein, mitochondrial (Lipoic acid-containing protein) | 0,613466008 | 0,02456 | 0,241832734 |
| P52566 | GDIR2 | Rho GDP-dissociation inhibitor 2 (Rho GDI 2) (Ly-GDI) (Rho-GDI beta) | 0,498628247 | 0,02948 | 0,260978097 |
| P07093 | GDN | Glia-derived nexin (GDN) (Peptidase inhibitor 7) (PI-7) (Protease nexin 1) (PN-1) (Protease nexin I) (Serpin E2) | 1,0780396 | 0,03253 | 0,270276513 |
| O75223 | GGCT | Gamma-glutamylcyclotransferase (EC 4,3,2,9) (Cytochrome c-releasing factor 21) | 1,115494726 | 1,88E-05 | 0,007375622 |
| Q92820 | GGH | Gamma-glutamyl hydrolase (EC 3,4,19,9) (Conjugase) (GH) (Gamma-Glu-X carboxypeptidase) | 0,697630237 | 0,00603 | 0,139953969 |
| Q8N2G8 | GHDC | GH3 domain-containing protein | -0,318162645 | 0,02164 | 0,235752629 |
| Q9Y5P6 | GMPPB | Mannose-1-phosphate guanyltransferase beta (EC 2,7,7,13) (GDP-mannose pyrophosphorylase B) (GTP-mannose-1-phosphate guanylyltransferase beta) | 0,31918971 | 0,04907 | 0,327955691 |
| P46926 | GNPI1 | Glucosamine-6-phosphate isomerase 1 (EC 3,5,99,6) (Glucosamine-6-phosphate deaminase 1) (GNPDA 1) (GlcN6P deaminase 1) (Oscillin) | 0,404223103 | 0,0284 | 0,257481581 |
| Q9HCL2 | GPAT1 | Glycerol-3-phosphate acyltransferase 1, mitochondrial (GPAT-1) (EC 2,3,1,15) | 0,274781902 | 0,02415 | 0,239812457 |
| Q53EU6 | GPAT3 | Glycerol-3-phosphate acyltransferase 3 (GPAT-3) (EC 2,3,1,15) (1-acyl-sn-glycerol-3-phosphate O-acyltransferase 10) (AGPAT 10) (1-acyl-sn-glycerol-3-phosphate O-acyltransferase 9) (1-AGP acyltransferase 9) (1-AGPAT 9) (EC 2,3,1,51) (Acyl-CoA:glycerol-3-phosphate acyltransferase 3) (hGPAT3) (Lung cancer metastasis-associated protein 1) (Lysophosphatidic acid acyltransferase theta) (LPAAT-theta) (MAG-1) | -0,536022967 | 0,00169 | 0,074278921 |
| Q92917 | GPKOW | G-patch domain and KOW motifs-containing protein (G-patch domain-containing protein 5) (Protein MOS2 homolog) (Protein T54) | -0,406943508 | 0,04705 | 0,325912134 |
| Q86YR5 | GPSM1 | G-protein-signaling modulator 1 (Activator of G-protein signaling 3) | 0,371900068 | 0,03325 | 0,273349722 |
| P07203 | GPX1 | Glutathione peroxidase 1 (GPx-1) (GSHPx-1) (EC 1,11,1,9) (Cellular glutathione peroxidase) | -0,29319386 | 0,04774 | 0,325912134 |
| P36969 | GPX4 | Phospholipid hydroperoxide glutathione peroxidase (PHGPx) (EC 1,11,1,12) (Glutathione peroxidase 4) (GPx-4) (GSHPx-4) | 0,31231312 | 0,03088 | 0,266198755 |
| P48637 | GSHB | Glutathione synthetase (GSH synthetase) (GSH-S) (EC 6,3,2,3) (Glutathione synthase) | 0,494763498 | 0,0023 | 0,091530158 |
| P49840 | GSK3A | Glycogen synthase kinase-3 alpha (GSK-3 alpha) (EC 2,7,11,26) (Serine/threonine-protein kinase GSK3A) (EC 2,7,11,1) | -0,34454891 | 0,02278 | 0,236759134 |
| P78417 | GSTO1 | Glutathione S-transferase omega-1 (GSTO-1) (EC 2,5,1,18) (Glutathione S-transferase omega 1-1) (GSTO 1-1) (Glutathione-dependent dehydroascorbate reductase) (EC 1,8,5,1) (Monomethylarsonic acid reductase) (MMA(V) reductase) (EC 1,20,4,2) (S-(Phenacyl)glutathione reductase) (SPG-R) | 0,755551143 | 0,00023 | 0,022900288 |
| Q9P0M6 | H2AW | Core histone macro-H2A,2 (Histone macroH2A2) (mH2A2) | 0,516078278 | 0,02185 | 0,235814224 |
| P0C0S5 | H2AZ | Histone H2A,Z (H2A/z) | -0,394425097 | 0,02669 | 0,251064959 |
| Q99880 | H2B1L | Histone H2B type 1-L (Histone H2B,c) (H2B/c) | 0,415206841 | 0,02856 | 0,257481581 |
| Q13442 | HAP28 | 28 kDa heat- and acid-stable phosphoprotein (PDGF-associated protein) (PAP) (PDGFA-associated protein 1) (PAP1) | -0,613998031 | 0,00477 | 0,125487524 |
| P69905 | HBA | Hemoglobin subunit alpha (Alpha-globin) (Hemoglobin alpha chain) | 0,987800022 | 0,01071 | 0,188371309 |
| P68871 | HBB | Hemoglobin subunit beta (Beta-globin) (Hemoglobin beta chain) [Cleaved into: LVV-hemorphin-7; Spinorphin] | 1,114885976 | 0,00528 | 0,131177383 |
| P02042 | HBD | Hemoglobin subunit delta (Delta-globin) (Hemoglobin delta chain) | 0,817704461 | 0,01227 | 0,199766061 |
| Q16836 | HCDH | Hydroxyacyl-coenzyme A dehydrogenase, mitochondrial (HCDH) (EC 1,1,1,35) (Medium and short-chain L-3-hydroxyacyl-coenzyme A dehydrogenase) (Short-chain 3-hydroxyacyl-CoA dehydrogenase) | 0,313324858 | 0,04636 | 0,324911402 |
| Q7Z4H3 | HDDC2 | HD domain-containing protein 2 (Hepatitis C virus NS5A-transactivated protein 2) (HCV NS5A-transactivated protein 2) | 0,761450905 | 0,0003 | 0,027785329 |
| Q9Y5Z4 | HEBP2 | Heme-binding protein 2 (Placental protein 23) (PP23) (Protein SOUL) | 0,849413742 | 0,00231 | 0,091530158 |
| Q9ULT8 | HECD1 | E3 ubiquitin-protein ligase HECTD1 (EC 2,3,2,26) (E3 ligase for inhibin receptor) (EULIR) (HECT domain-containing protein 1) (HECT-type E3 ubiquitin transferase HECTD1) | -0,534360584 | 0,00564 | 0,135618752 |
| P02790 | HEMO | Hemopexin (Beta-1B-glycoprotein) | 1,671121279 | 8,48E-05 | 0,013863419 |
| P05546 | HEP2 | Heparin cofactor 2 (Heparin cofactor II) (HC-II) (Protease inhibitor leuserpin-2) (HLS2) (Serpin D1) | 0,848051339 | 0,00874 | 0,167176556 |
| P07686 | HEXB | Beta-hexosaminidase subunit beta (EC 3,2,1,52) (Beta-N-acetylhexosaminidase subunit beta) (Hexosaminidase subunit B) (Cervical cancer proto-oncogene 7 protein) (HCC-7) (N-acetyl-beta-glucosaminidase subunit beta) [Cleaved into: Beta-hexosaminidase subunit beta chain B; Beta-hexosaminidase subunit beta chain A] | 0,518884583 | 0,02479 | 0,241832734 |
| P35914 | HMGCL | Hydroxymethylglutaryl-CoA lyase, mitochondrial (HL) (HMG-CoA lyase) (EC 4,1,3,4) (3-hydroxy-3-methylglutarate-CoA lyase) | 0,469696484 | 0,01956 | 0,23220076 |
| P50135 | HNMT | Histamine N-methyltransferase (HMT) (EC 2,1,1,8) | 0,800227222 | 0,0008 | 0,047747284 |
| Q8IWW8 | HOT | Hydroxyacid-oxoacid transhydrogenase, mitochondrial (HOT) (EC 1,1,99,24) (Alcohol dehydrogenase iron-containing protein 1) (ADHFe1) (Fe-containing alcohol dehydrogenase) | -0,415805727 | 0,04698 | 0,325912134 |
| P00738 | HPT | Haptoglobin (Zonulin) [Cleaved into: Haptoglobin alpha chain; Haptoglobin beta chain] | 1,719062234 | 0,02349 | 0,236759134 |
| Q92598 | HS105 | Heat shock protein 105 kDa (Antigen NY-CO-25) (Heat shock 110 kDa protein) | 0,826863815 | 0,0041 | 0,114007879 |
| Q96MM6 | HS12B | Heat shock 70 kDa protein 12B | -0,345249081 | 0,01618 | 0,213209762 |
| O75506 | HSBP1 | Heat shock factor-binding protein 1 (Nasopharyngeal carcinoma-associated antigen 13) (NPC-A-13) | 0,325831689 | 0,03551 | 0,28484168 |
| P34932 | HSP74 | Heat shock 70 kDa protein 4 (HSP70RY) (Heat shock 70-related protein APG-2) | 0,522096226 | 0,00273 | 0,097030859 |
| Q16082 | HSPB2 | Heat shock protein beta-2 (HspB2) (DMPK-binding protein) (MKBP) | -0,459112861 | 0,01258 | 0,199766061 |
| Q12988 | HSPB3 | Heat shock protein beta-3 (HspB3) (Heat shock 17 kDa protein) (HSP 17) (Protein 3) | -0,41085277 | 0,0211 | 0,234998667 |
| P52789 | HXK2 | Hexokinase-2 (EC 2,7,1,1) (Hexokinase type II) (HK II) (Muscle form hexokinase) | -0,713734865 | 0,0184 | 0,226221613 |
| P05155 | IC1 | Plasma protease C1 inhibitor (C1 Inh) (C1Inh) (C1 esterase inhibitor) (C1-inhibiting factor) (Serpin G1) | 1,16147074 | 0,01112 | 0,191369497 |
| P13598 | ICAM2 | Intercellular adhesion molecule 2 (ICAM-2) (CD antigen CD102) | 0,38454476 | 0,02124 | 0,234998667 |
| P56537 | IF6 | Eukaryotic translation initiation factor 6 (eIF-6) (B(2)GCN homolog) (B4 integrin interactor) (CAB) (p27(BBP)) | 0,597589524 | 0,00822 | 0,163422907 |
| P78318 | IGBP1 | Immunoglobulin-binding protein 1 (B-cell signal transduction molecule alpha 4) (Protein alpha-4) (CD79a-binding protein 1) (Protein phosphatase 2/4/6 regulatory subunit) (Renal carcinoma antigen NY-REN-16) | 0,303130213 | 0,04441 | 0,316650023 |
| P01876 | IGHA1 | Immunoglobulin heavy constant alpha 1 (Ig alpha-1 chain C region) (Ig alpha-1 chain C region BUR) (Ig alpha-1 chain C region TRO) | 1,938824859 | 2,30E-05 | 0,008201724 |
| P01880 | IGHD | Immunoglobulin heavy constant delta (Ig delta chain C region) (Ig delta chain C region NIG-65) (Ig delta chain C region WAH) | 0,638255203 | 0,01559 | 0,212766683 |
| P01857 | IGHG1 | Immunoglobulin heavy constant gamma 1 (Ig gamma-1 chain C region) (Ig gamma-1 chain C region EU) (Ig gamma-1 chain C region KOL) (Ig gamma-1 chain C region NIE) | 1,47086495 | 0,00165 | 0,074278921 |
| P01859 | IGHG2 | Immunoglobulin heavy constant gamma 2 (Ig gamma-2 chain C region) (Ig gamma-2 chain C region DOT) (Ig gamma-2 chain C region TIL) (Ig gamma-2 chain C region ZIE) | 2,55459753 | 7,02E-05 | 0,013112951 |
| P01860 | IGHG3 | Immunoglobulin heavy constant gamma 3 (HDC) (Heavy chain disease protein) (Ig gamma-3 chain C region) | 0,771874184 | 0,03448 | 0,279408322 |
| P01861 | IGHG4 | Immunoglobulin heavy constant gamma 4 (Ig gamma-4 chain C region) | 0,890365012 | 0,0235 | 0,236759134 |
| P01871 | IGHM | Immunoglobulin heavy constant mu (Ig mu chain C region) (Ig mu chain C region BOT) (Ig mu chain C region GAL) (Ig mu chain C region OU) | 0,87752594 | 0,00062 | 0,043603785 |
| P01834 | IGKC | Immunoglobulin kappa constant (Ig kappa chain C region) (Ig kappa chain C region AG) (Ig kappa chain C region CUM) (Ig kappa chain C region EU) (Ig kappa chain C region OU) (Ig kappa chain C region ROY) (Ig kappa chain C region TI) | 1,740320435 | 4,75E-05 | 0,011475286 |
| B9A064 | IGLL5 | Immunoglobulin lambda-like polypeptide 5 (G lambda-1) (Germline immunoglobulin lambda 1) | 2,389119754 | 1,83E-05 | 0,007375622 |
| P30740 | ILEU | Leukocyte elastase inhibitor (LEI) (Monocyte/neutrophil elastase inhibitor) (EI) (M/NEI) (Peptidase inhibitor 2) (PI-2) (Serpin B1) | 0,571499389 | 0,00287 | 0,097030859 |
| Q9Y2B9 | IPKG | cAMP-dependent protein kinase inhibitor gamma (PKI-gamma) | 0,335784826 | 0,04257 | 0,311743911 |
| Q15181 | IPYR | Inorganic pyrophosphatase (EC 3,6,1,1) (Pyrophosphate phospho-hydrolase) (PPase) | 0,585487307 | 0,00836 | 0,163422907 |
| Q9H2U2 | IPYR2 | Inorganic pyrophosphatase 2, mitochondrial (EC 3,6,1,1) (Pyrophosphatase SID6-306) (Pyrophosphate phospho-hydrolase 2) (PPase 2) | 0,473258124 | 0,04439 | 0,316650023 |
| Q13576 | IQGA2 | Ras GTPase-activating-like protein IQGAP2 | 0,471610628 | 0,00813 | 0,163422907 |
| P56199 | ITA1 | Integrin alpha-1 (CD49 antigen-like family member A) (Laminin and collagen receptor) (VLA-1) (CD antigen CD49a) | -0,300674077 | 0,0292 | 0,259726687 |
| P18084 | ITB5 | Integrin beta-5 | 0,31598011 | 0,03469 | 0,280492718 |
| Q9BY32 | ITPA | Inosine triphosphate pyrophosphatase (ITPase) (Inosine triphosphatase) (EC 3,6,1,9) (Non-canonical purine NTP pyrophosphatase) (Non-standard purine NTP pyrophosphatase) (Nucleoside-triphosphate diphosphatase) (Nucleoside-triphosphate pyrophosphatase) (NTPase) (Putative oncogene protein hlc14-06-p) | 1,172434669 | 0,00291 | 0,097030859 |
| Q9P266 | JCAD | Junctional protein associated with coronary artery disease (JCAD) | -0,393543328 | 0,03173 | 0,268193682 |
| Q9BR39 | JPH2 | Junctophilin-2 (JP-2) (Junctophilin type 2) | -0,630262165 | 0,00052 | 0,040899265 |
| P29622 | KAIN | Kallistatin (Kallikrein inhibitor) (Peptidase inhibitor 4) (PI-4) (Serpin A4) | 0,650002823 | 0,01022 | 0,182203157 |
| Q6NY19 | KANK3 | KN motif and ankyrin repeat domain-containing protein 3 (Ankyrin repeat domain-containing protein 47) | -0,29621394 | 0,02113 | 0,234998667 |
| Q13554 | KCC2B | Calcium/calmodulin-dependent protein kinase type II subunit beta (CaM kinase II subunit beta) (CaMK-II subunit beta) (EC 2,7,11,17) | -0,407249142 | 0,00583 | 0,137765097 |
| Q53GT1 | KLH22 | Kelch-like protein 22 | -0,787863695 | 0,03172 | 0,268193682 |
| P01042 | KNG1 | Kininogen-1 (Alpha-2-thiol proteinase inhibitor) (Fitzgerald factor) (High molecular weight kininogen) (HMWK) (Williams-Fitzgerald-Flaujeac factor) [Cleaved into: Kininogen-1 heavy chain; T-kinin (Ile-Ser-Bradykinin); Bradykinin (Kallidin I); Lysyl-bradykinin (Kallidin II); Kininogen-1 light chain; Low molecular weight growth-promoting factor] | 1,230308022 | 0,00202 | 0,082556156 |
| P51812 | KS6A3 | Ribosomal protein S6 kinase alpha-3 (S6K-alpha-3) (EC 2,7,11,1) (90 kDa ribosomal protein S6 kinase 3) (p90-RSK 3) (p90RSK3) (Insulin-stimulated protein kinase 1) (ISPK-1) (MAP kinase-activated protein kinase 1b) (MAPK-activated protein kinase 1b) (MAPKAP kinase 1b) (MAPKAPK-1b) (Ribosomal S6 kinase 2) (RSK-2) (pp90RSK2) | -0,41750526 | 0,01459 | 0,211171406 |
| P04433 | KV311 | Immunoglobulin kappa variable 3-11 (Ig kappa chain V-III region VG) | 0,561963739 | 0,01404 | 0,20783937 |
| P01611 | KVD12 | Immunoglobulin kappa variable 1D-12 (Ig kappa chain V-I region Wes) | 1,393073393 | 0,00017 | 0,019060893 |
| Q53H82 | LACB2 | Endoribonuclease LACTB2 (EC 3,1,27,-) (Beta-lactamase-like protein 2) | 0,496542176 | 0,01786 | 0,225054195 |
| P11279 | LAMP1 | Lysosome-associated membrane glycoprotein 1 (LAMP-1) (Lysosome-associated membrane protein 1) (CD107 antigen-like family member A) (CD antigen CD107a) | 0,347595867 | 0,04396 | 0,315790315 |
| Q71RC2 | LARP4 | La-related protein 4 (La ribonucleoprotein domain family member 4) | -0,639575699 | 0,00959 | 0,175766361 |
| Q9UIQ6 | LCAP | Leucyl-cystinyl aminopeptidase (Cystinyl aminopeptidase) (EC 3,4,11,3) (Insulin-regulated membrane aminopeptidase) (Insulin-responsive aminopeptidase) (IRAP) (Oxytocinase) (OTase) (Placental leucine aminopeptidase) (P-LAP) [Cleaved into: Leucyl-cystinyl aminopeptidase, pregnancy serum form] | 0,363274963 | 0,02407 | 0,239646222 |
| P31025 | LCN1 | Lipocalin-1 (Tear lipocalin) (Tlc) (Tear prealbumin) (TP) (von Ebner gland protein) (VEG protein) | -1,068327079 | 0,00622 | 0,141663963 |
| P00338 | LDHA | L-lactate dehydrogenase A chain (LDH-A) (EC 1,1,1,27) (Cell proliferation-inducing gene 19 protein) (LDH muscle subunit) (LDH-M) (Renal carcinoma antigen NY-REN-59) | 0,516127388 | 0,02567 | 0,24500416 |
| P07195 | LDHB | L-lactate dehydrogenase B chain (LDH-B) (EC 1,1,1,27) (LDH heart subunit) (LDH-H) (Renal carcinoma antigen NY-REN-46) | 0,516879582 | 0,0058 | 0,137765097 |
| P09382 | LEG1 | Galectin-1 (Gal-1) (14 kDa laminin-binding protein) (HLBP14) (14 kDa lectin) (Beta-galactoside-binding lectin L-14-I) (Galaptin) (HBL) (HPL) (Lactose-binding lectin 1) (Lectin galactoside-binding soluble 1) (Putative MAPK-activating protein PM12) (S-Lac lectin 1) | 0,788996135 | 0,00174 | 0,07484788 |
| P17931 | LEG3 | Galectin-3 (Gal-3) (35 kDa lectin) (Carbohydrate-binding protein 35) (CBP 35) (Galactose-specific lectin 3) (Galactoside-binding protein) (GALBP) (IgE-binding protein) (L-31) (Laminin-binding protein) (Lectin L-29) (Mac-2 antigen) | -0,369342718 | 0,04247 | 0,311743911 |
| Q8NC56 | LEMD2 | LEM domain-containing protein 2 (hLEM2) | -0,306897753 | 0,03121 | 0,268193682 |
| P06858 | LIPL | Lipoprotein lipase (LPL) (EC 3,1,1,34) | -0,481322503 | 0,02547 | 0,243710647 |
| Q03252 | LMNB2 | Lamin-B2 | -0,372600036 | 0,02752 | 0,255168841 |
| Q08397 | LOXL1 | Lysyl oxidase homolog 1 (EC 1,4,3,-) (Lysyl oxidase-like protein 1) (LOL) | 1,197311537 | 0,037 | 0,29139789 |
| Q8N9N7 | LRC57 | Leucine-rich repeat-containing protein 57 | 0,70099018 | 8,23E-05 | 0,013863419 |
| Q14766 | LTBP1 | Latent-transforming growth factor beta-binding protein 1 (LTBP-1) (Transforming growth factor beta-1-binding protein 1) (TGF-beta1-BP-1) | 0,421247157 | 0,01866 | 0,226221613 |
| P80748 | LV321 | Immunoglobulin lambda variable 3-21 (Ig lambda chain V-III region LOI) (Ig lambda chain V-V region DEL) (Ig lambda chain V-VII region MOT) | 1,503403181 | 0,00015 | 0,017783943 |
| P10253 | LYAG | Lysosomal alpha-glucosidase (EC 3,2,1,20) (Acid maltase) (Aglucosidase alfa) [Cleaved into: 76 kDa lysosomal alpha-glucosidase; 70 kDa lysosomal alpha-glucosidase] | 0,387474736 | 0,02797 | 0,257481581 |
| Q8WZA0 | LZIC | Protein LZIC (Leucine zipper and CTNNBIP1 domain-containing protein) (Leucine zipper and ICAT homologous domain-containing protein) | 0,257894593 | 0,04241 | 0,311743911 |
| P55145 | MANF | Mesencephalic astrocyte-derived neurotrophic factor (Arginine-rich protein) (Protein ARMET) | 0,355724069 | 0,03572 | 0,285591713 |
| Q66K74 | MAP1S | Microtubule-associated protein 1S (MAP-1S) (BPY2-interacting protein 1) (Microtubule-associated protein 8) (Variable charge Y chromosome 2-interacting protein 1) (VCY2-interacting protein 1) (VCY2IP-1) [Cleaved into: MAP1S heavy chain; MAP1S light chain] | -0,526679715 | 0,02312 | 0,236759134 |
| P49137 | MAPK2 | MAP kinase-activated protein kinase 2 (MAPK-activated protein kinase 2) (MAPKAP kinase 2) (MAPKAP-K2) (MAPKAPK-2) (MK-2) (MK2) (EC 2,7,11,1) | -0,395909931 | 0,02613 | 0,24857069 |
| Q7Z434 | MAVS | Mitochondrial antiviral-signaling protein (MAVS) (CARD adapter inducing interferon beta) (Cardif) (Interferon beta promoter stimulator protein 1) (IPS-1) (Putative NF-kappa-B-activating protein 031N) (Virus-induced-signaling adapter) (VISA) | -0,428421133 | 0,00396 | 0,114007879 |
| P40925 | MDHC | Malate dehydrogenase, cytoplasmic (EC 1,1,1,37) (Cytosolic malate dehydrogenase) (Diiodophenylpyruvate reductase) (EC 1,1,1,96) | 0,58975067 | 0,00708 | 0,150838919 |
| Q14696 | MESD | LRP chaperone MESD (LDLR chaperone MESD) (Mesoderm development LRP chaperone MESD) (Mesoderm development candidate 2) (Mesoderm development protein) (Renal carcinoma antigen NY-REN-61) | 0,46196531 | 0,00828 | 0,163422907 |
| Q9H7H0 | MET17 | Methyltransferase-like protein 17, mitochondrial (EC 2,1,1,-) (False p73 target gene protein) (Methyltransferase 11 domain-containing protein 1) (Protein RSM22 homolog, mitochondrial) | -0,632431348 | 0,0012 | 0,060334988 |
| P55083 | MFAP4 | Microfibril-associated glycoprotein 4 | -1,065997783 | 0,00069 | 0,045663466 |
| Q08431 | MFGM | Lactadherin (Breast epithelial antigen BA46) (HMFG) (MFGM) (Milk fat globule-EGF factor 8) (MFG-E8) (SED1) [Cleaved into: Lactadherin short form; Medin] | 0,524167449 | 0,01268 | 0,199766061 |
| Q9H019 | MFR1L | Mitochondrial fission regulator 1-like | -0,354632455 | 0,00893 | 0,169218477 |
| Q9H3U5 | MFSD1 | Major facilitator superfamily domain-containing protein 1 (Smooth muscle cell-associated protein 4) (SMAP-4) | -0,386381836 | 0,02836 | 0,257481581 |
| Q5XKP0 | MIC13 | MICOS complex subunit MIC13 (Protein P117) | 0,381054177 | 0,01257 | 0,199766061 |
| Q9NX63 | MIC19 | MICOS complex subunit MIC19 (Coiled-coil-helix-coiled-coil-helix domain-containing protein 3) | -0,297881331 | 0,03856 | 0,298767304 |
| Q9BUR5 | MIC26 | MICOS complex subunit MIC26 (Apolipoprotein O) (MICOS complex subunit MIC23) (Protein FAM121B) | 0,362905977 | 0,02124 | 0,234998667 |
| Q6UXV4 | MIC27 | MICOS complex subunit MIC27 (Apolipoprotein O-like) (Protein FAM121A) | 0,402681283 | 0,0213 | 0,234998667 |
| P45984 | MK09 | Mitogen-activated protein kinase 9 (MAP kinase 9) (MAPK 9) (EC 2,7,11,24) (JNK-55) (Stress-activated protein kinase 1a) (SAPK1a) (Stress-activated protein kinase JNK2) (c-Jun N-terminal kinase 2) | -0,284549941 | 0,0344 | 0,279305323 |
| Q14165 | MLEC | Malectin | -0,462449802 | 0,00472 | 0,125487524 |
| Q01449 | MLRA | Myosin regulatory light chain 2, atrial isoform (MLC-2a) (MLC2a) (Myosin light chain 2a) (Myosin regulatory light chain 7) | 1,050881361 | 0,0128 | 0,200054655 |
| P10916 | MLRV | Myosin regulatory light chain 2, ventricular/cardiac muscle isoform (MLC-2) (MLC-2v) (Cardiac myosin light chain 2) (Myosin light chain 2, slow skeletal/ventricular muscle isoform) (MLC-2s/v) (Ventricular myosin light chain 2) | 0,357999023 | 0,02103 | 0,234998667 |
| Q9NZB8 | MOCS1 | Molybdenum cofactor biosynthesis protein 1 (Cell migration-inducing gene 11 protein) (Molybdenum cofactor synthesis-step 1 protein A-B) [Includes: GTP 3',8-cyclase (EC 4,1,99,22) (Molybdenum cofactor biosynthesis protein A); Cyclic pyranopterin monophosphate synthase (EC 4,6,1,17) (Molybdenum cofactor biosynthesis protein C)] | -0,573897783 | 0,00652 | 0,143094484 |
| P53985 | MOT1 | Monocarboxylate transporter 1 (MCT 1) (Solute carrier family 16 member 1) | 0,386025007 | 0,03668 | 0,290052814 |
| Q14168 | MPP2 | MAGUK p55 subfamily member 2 (Discs large homolog 2) (Protein MPP2) | -0,416430156 | 0,01853 | 0,226221613 |
| P49006 | MRP | MARCKS-related protein (MARCKS-like protein 1) (Macrophage myristoylated alanine-rich C kinase substrate) (Mac-MARCKS) (MacMARCKS) | 0,553514116 | 0,03939 | 0,301136743 |
| Q96DH6 | MSI2H | RNA-binding protein Musashi homolog 2 (Musashi-2) | -0,290909909 | 0,01836 | 0,226221613 |
| Q96S19 | MTL26 | Methyltransferase-like 26 | -0,62644985 | 0,00574 | 0,137385262 |
| P42285 | MTREX | Exosome RNA helicase MTR4 (EC 3,6,4,13) (ATP-dependent RNA helicase DOB1) (ATP-dependent RNA helicase SKIV2L2) (Superkiller viralicidic activity 2-like 2) (TRAMP-like complex helicase) | -0,774300458 | 0,02544 | 0,243710647 |
| Q5HYI7 | MTX3 | Metaxin-3 | -0,482851181 | 0,00298 | 0,098224502 |
| P20591 | MX1 | Interferon-induced GTP-binding protein Mx1 (Interferon-induced protein p78) (IFI-78K) (Interferon-regulated resistance GTP-binding protein MxA) (Myxoma resistance protein 1) (Myxovirus resistance protein 1) [Cleaved into: Interferon-induced GTP-binding protein Mx1, N-terminally processed] | -1,421960024 | 0,00402 | 0,114007879 |
| P20592 | MX2 | Interferon-induced GTP-binding protein Mx2 (Interferon-regulated resistance GTP-binding protein MxB) (Myxovirus resistance protein 2) (p78-related protein) | -0,613262334 | 0,0204 | 0,234761058 |
| Q8IUG5 | MY18B | Unconventional myosin-XVIIIb | -0,514056121 | 0,00698 | 0,149608152 |
| O75592 | MYCB2 | E3 ubiquitin-protein ligase MYCBP2 (EC 2,3,2,-) (Myc-binding protein 2) (Protein associated with Myc) | 0,250238442 | 0,04939 | 0,328959493 |
| P12882 | MYH1 | Myosin-1 (Myosin heavy chain 1) (Myosin heavy chain 2x) (MyHC-2x) (Myosin heavy chain IIx/d) (MyHC-IIx/d) (Myosin heavy chain, skeletal muscle, adult 1) | -1,707943146 | 3,80E-05 | 0,009944827 |
| Q7Z406 | MYH14 | Myosin-14 (Myosin heavy chain 14) (Myosin heavy chain, non-muscle IIc) (Non-muscle myosin heavy chain IIc) (NMHC II-C) | -0,394171574 | 0,00821 | 0,163422907 |
| Q9Y2K3 | MYH15 | Myosin-15 (Myosin heavy chain 15) | -0,456630067 | 0,00551 | 0,134510312 |
| A7E2Y1 | MYH7B | Myosin-7B (Antigen MLAA-21) (Myosin cardiac muscle beta chain) (Myosin heavy chain 7B, cardiac muscle beta isoform) (Slow A MYH14) | -0,790546394 | 0,00071 | 0,045663466 |
| Q32MK0 | MYLK3 | Myosin light chain kinase 3 (EC 2,7,11,18) (Cardiac-MyBP-C-associated Ca/CaM kinase) (Cardiac-MLCK) | -0,372278866 | 0,0162 | 0,213209762 |
| Q9NZM1 | MYOF | Myoferlin (Fer-1-like protein 3) | 0,488221313 | 0,02548 | 0,243710647 |
| A2RRP1 | NBAS | Neuroblastoma-amplified sequence (Neuroblastoma-amplified gene protein) | -0,359088896 | 0,00968 | 0,17584173 |
| P15531 | NDKA | Nucleoside diphosphate kinase A (NDK A) (NDP kinase A) (EC 2,7,4,6) (Granzyme A-activated DNase) (GAAD) (Metastasis inhibition factor nm23) (NM23-H1) (Tumor metastatic process-associated protein) | 0,485830448 | 0,00311 | 0,100882921 |
| P22392 | NDKB | Nucleoside diphosphate kinase B (NDK B) (NDP kinase B) (EC 2,7,4,6) (C-myc purine-binding transcription factor PUF) (Histidine protein kinase NDKB) (EC 2,7,13,3) (nm23-H2) | 0,554195268 | 0,00325 | 0,102119691 |
| O95182 | NDUA7 | NADH dehydrogenase [ubiquinone] 1 alpha subcomplex subunit 7 (Complex I-B14,5a) (CI-B14,5a) (NADH-ubiquinone oxidoreductase subunit B14,5a) | 0,369294626 | 0,0487 | 0,327613246 |
| O43676 | NDUB3 | NADH dehydrogenase [ubiquinone] 1 beta subcomplex subunit 3 (Complex I-B12) (CI-B12) (NADH-ubiquinone oxidoreductase B12 subunit) | 0,458355603 | 0,01691 | 0,21888084 |
| Q7Z3B1 | NEGR1 | Neuronal growth regulator 1 (IgLON family member 4) | 0,529633472 | 0,02631 | 0,248649026 |
| Q9BYT8 | NEUL | Neurolysin, mitochondrial (EC 3,4,24,16) (Angiotensin-binding protein) (Microsomal endopeptidase) (MEP) (Mitochondrial oligopeptidase M) (Neurotensin endopeptidase) | 0,561377945 | 0,04797 | 0,325912134 |
| Q15599 | NHRF2 | Na(+)/H(+) exchange regulatory cofactor NHE-RF2 (NHERF-2) (NHE3 kinase A regulatory protein E3KARP) (SRY-interacting protein 1) (SIP-1) (Sodium-hydrogen exchanger regulatory factor 2) (Solute carrier family 9 isoform A3 regulatory factor 2) (Tyrosine kinase activator protein 1) (TKA-1) | -0,31056789 | 0,0219 | 0,235814224 |
| P14543 | NID1 | Nidogen-1 (NID-1) (Entactin) | -0,367565492 | 0,01329 | 0,201757374 |
| Q96D46 | NMD3 | 60S ribosomal export protein NMD3 (hNMD3) | -0,419144411 | 0,00853 | 0,164055531 |
| Q8IVI9 | NOSTN | Nostrin (BM247 homolog) (Nitric oxide synthase traffic inducer) (Nitric oxide synthase trafficker) (eNOS-trafficking inducer) | -0,565752688 | 0,00931 | 0,173052847 |
| P61916 | NPC2 | NPC intracellular cholesterol transporter 2 (Epididymal secretory protein E1) (Human epididymis-specific protein 1) (He1) (Niemann-Pick disease type C2 protein) | 1,36282824 | 5,21E-07 | 0,000680806 |
| Q9Y639 | NPTN | Neuroplastin (Stromal cell-derived receptor 1) (SDR-1) | -0,403387594 | 0,02822 | 0,257481581 |
| Q9UNZ2 | NSF1C | NSFL1 cofactor p47 (UBX domain-containing protein 2C) (p97 cofactor p47) | -0,326022121 | 0,02967 | 0,261063402 |
| P61970 | NTF2 | Nuclear transport factor 2 (NTF-2) (Placental protein 15) (PP15) | 0,733796934 | 0,01146 | 0,193784086 |
| Q8WUM0 | NU133 | Nuclear pore complex protein Nup133 (133 kDa nucleoporin) (Nucleoporin Nup133) | -0,313576671 | 0,02813 | 0,257481581 |
| P03891 | NU2M | NADH-ubiquinone oxidoreductase chain 2 (EC 1,6,5,3) (NADH dehydrogenase subunit 2) | -0,390871734 | 0,0143 | 0,208833198 |
| Q9NRX3 | NUA4L | NADH dehydrogenase [ubiquinone] 1 alpha subcomplex subunit 4-like 2 (NADH-ubiquinone oxidoreductase MLRQ subunit homolog) (NUOMS) | 0,448145257 | 0,01304 | 0,201422202 |
| Q14249 | NUCG | Endonuclease G, mitochondrial (Endo G) (EC 3,1,30,-) | -0,267357389 | 0,04239 | 0,311743911 |
| O75147 | OBSL1 | Obscurin-like protein 1 | -0,325180448 | 0,02944 | 0,260978097 |
| Q9ULD0 | OGDHL | 2-oxoglutarate dehydrogenase-like, mitochondrial (EC 1,2,4,-) (2-oxoglutarate dehydrogenase complex component E1-like) (OGDC-E1-like) (Alpha-ketoglutarate dehydrogenase-like) | 0,729948916 | 0,03264 | 0,270603898 |
| O14841 | OPLA | 5-oxoprolinase (EC 3,5,2,9) (5-oxo-L-prolinase) (5-OPase) (Pyroglutamase) | -0,545254395 | 0,00368 | 0,110182266 |
| Q96CV9 | OPTN | Optineurin (E3-14,7K-interacting protein) (FIP-2) (Huntingtin yeast partner L) (Huntingtin-interacting protein 7) (HIP-7) (Huntingtin-interacting protein L) (NEMO-related protein) (Optic neuropathy-inducing protein) (Transcription factor IIIA-interacting protein) (TFIIIA-IntP) | 0,320744163 | 0,03075 | 0,266195983 |
| Q9Y3B8 | ORN | Oligoribonuclease, mitochondrial (EC 3,1,-,-) (RNA exonuclease 2 homolog) (Small fragment nuclease) | 0,582260108 | 0,00061 | 0,043603785 |
| Q9NPF4 | OSGEP | Probable tRNA N6-adenosine threonylcarbamoyltransferase (EC 2,3,1,234) (N6-L-threonylcarbamoyladenine synthase) (t(6)A synthase) (O-sialoglycoprotein endopeptidase) (hOSGEP) (t(6)A37 threonylcarbamoyladenosine biosynthesis protein OSGEP) (tRNA threonylcarbamoyladenosine biosynthesis protein OSGEP) | -0,257243219 | 0,03206 | 0,268193682 |
| Q06190 | P2R3A | Serine/threonine-protein phosphatase 2A regulatory subunit B'' subunit alpha (PP2A subunit B isoform PR72/PR130) (PP2A subunit B isoform R3 isoform) (PP2A subunit B isoforms B''-PR72/PR130) (PP2A subunit B isoforms B72/B130) (Serine/threonine-protein phosphatase 2A 72/130 kDa regulatory subunit B) | -0,389849473 | 0,00404 | 0,114007879 |
| P27986 | P85A | Phosphatidylinositol 3-kinase regulatory subunit alpha (PI3-kinase regulatory subunit alpha) (PI3K regulatory subunit alpha) (PtdIns-3-kinase regulatory subunit alpha) (Phosphatidylinositol 3-kinase 85 kDa regulatory subunit alpha) (PI3-kinase subunit p85-alpha) (PtdIns-3-kinase regulatory subunit p85-alpha) | -1,072093174 | 0,01467 | 0,211583932 |
| Q9UKS6 | PACN3 | Protein kinase C and casein kinase substrate in neurons protein 3 (SH3 domain-containing protein 6511) | -0,38018957 | 0,01065 | 0,188143812 |
| O14832 | PAHX | Phytanoyl-CoA dioxygenase, peroxisomal (EC 1,14,11,18) (Phytanic acid oxidase) (Phytanoyl-CoA alpha-hydroxylase) (PhyH) | -0,824978542 | 0,00046 | 0,037282544 |
| Q16822 | PCKGM | Phosphoenolpyruvate carboxykinase [GTP], mitochondrial (PEPCK-M) (EC 4,1,1,32) | 0,482058821 | 0,02211 | 0,235814224 |
| O95613 | PCNT | Pericentrin (Kendrin) (Pericentrin-B) | -0,27198731 | 0,04199 | 0,311743911 |
| P49585 | PCY1A | Choline-phosphate cytidylyltransferase A (EC 2,7,7,15) (CCT-alpha) (CTP:phosphocholine cytidylyltransferase A) (CCT A) (CT A) (Phosphorylcholine transferase A) | 0,385224512 | 0,01615 | 0,213209762 |
| Q14123 | PDE1C | Calcium/calmodulin-dependent 3',5'-cyclic nucleotide phosphodiesterase 1C (Cam-PDE 1C) (EC 3,1,4,17) (hCam-3) | 0,425259527 | 0,01719 | 0,22102165 |
| Q15120 | PDK3 | [Pyruvate dehydrogenase (acetyl-transferring)] kinase isozyme 3, mitochondrial (EC 2,7,11,2) (Pyruvate dehydrogenase kinase isoform 3) | 0,26098901 | 0,04774 | 0,325912134 |
| P36955 | PEDF | Pigment epithelium-derived factor (PEDF) (Cell proliferation-inducing gene 35 protein) (EPC-1) (Serpin F1) | 1,083730481 | 0,00492 | 0,126430808 |
| P41219 | PERI | Peripherin (Neurofilament 4) | -0,610565571 | 0,01485 | 0,211823261 |
| O15212 | PFD6 | Prefoldin subunit 6 (Protein Ke2) | -0,502775713 | 0,00792 | 0,161852265 |
| Q96PD5 | PGRP2 | N-acetylmuramoyl-L-alanine amidase (EC 3,5,1,28) (Peptidoglycan recognition protein 2) (Peptidoglycan recognition protein long) (PGRP-L) | 1,289625761 | 0,00213 | 0,086162838 |
| P21810 | PGS1 | Biglycan (Bone/cartilage proteoglycan I) (PG-S1) | 1,056607293 | 0,00076 | 0,047747284 |
| P53611 | PGTB2 | Geranylgeranyl transferase type-2 subunit beta (EC 2,5,1,60) (Geranylgeranyl transferase type II subunit beta) (GGTase-II-beta) (Rab geranyl-geranyltransferase subunit beta) (Rab GG transferase beta) (Rab GGTase beta) (Rab geranylgeranyltransferase subunit beta) (Type II protein geranyl-geranyltransferase subunit beta) | -0,302673884 | 0,03312 | 0,272868479 |
| Q96S52 | PIGS | GPI transamidase component PIG-S (Phosphatidylinositol-glycan biosynthesis class S protein) | 0,440300146 | 0,02021 | 0,234761058 |
| Q8IWE5 | PKHM2 | Pleckstrin homology domain-containing family M member 2 (PH domain-containing family M member 2) (Salmonella-induced filaments A and kinesin-interacting protein) (SifA and kinesin-interacting protein) | -0,431440177 | 0,02385 | 0,238655908 |
| Q16513 | PKN2 | Serine/threonine-protein kinase N2 (EC 2,7,11,13) (PKN gamma) (Protein kinase C-like 2) (Protein-kinase C-related kinase 2) | 0,72917283 | 0,02103 | 0,234998667 |
| Q99569 | PKP4 | Plakophilin-4 (p0071) | -0,542586542 | 0,0108 | 0,188957641 |
| Q99943 | PLCA | 1-acyl-sn-glycerol-3-phosphate acyltransferase alpha (EC 2,3,1,51) (1-acylglycerol-3-phosphate O-acyltransferase 1) (1-AGP acyltransferase 1) (1-AGPAT 1) (Lysophosphatidic acid acyltransferase alpha) (LPAAT-alpha) (Protein G15) | -0,399962663 | 0,01832 | 0,226221613 |
| O60240 | PLIN1 | Perilipin-1 (Lipid droplet-associated protein) | 2,388120822 | 3,86E-06 | 0,003030192 |
| Q99541 | PLIN2 | Perilipin-2 (Adipophilin) (Adipose differentiation-related protein) (ADRP) | 1,365582031 | 3,48E-05 | 0,009944827 |
| Q96Q06 | PLIN4 | Perilipin-4 (Adipocyte protein S3-12) | 0,641885934 | 0,00507 | 0,129189254 |
| Q9NP80 | PLPL8 | Calcium-independent phospholipase A2-gamma (EC 3,1,1,5) (Intracellular membrane-associated calcium-independent phospholipase A2 gamma) (iPLA2-gamma) (PNPLA-gamma) (Patatin-like phospholipase domain-containing protein 8) (iPLA2-2) | -0,302195228 | 0,01987 | 0,232646998 |
| P13797 | PLST | Plastin-3 (T-plastin) | 0,297138002 | 0,04654 | 0,324973393 |
| P27169 | PON1 | Serum paraoxonase/arylesterase 1 (PON 1) (EC 3,1,1,2) (EC 3,1,1,81) (EC 3,1,8,1) (Aromatic esterase 1) (A-esterase 1) (K-45) (Serum aryldialkylphosphatase 1) | 0,728043347 | 0,04498 | 0,32014742 |
| Q9HBU9 | POPD2 | Popeye domain-containing protein 2 (Popeye protein 2) | -0,516234628 | 0,00474 | 0,125487524 |
| Q08209 | PP2BA | Serine/threonine-protein phosphatase 2B catalytic subunit alpha isoform (EC 3,1,3,16) (CAM-PRP catalytic subunit) (Calmodulin-dependent calcineurin A subunit alpha isoform) | -0,343497872 | 0,02871 | 0,257625811 |
| P62937 | PPIA | Peptidyl-prolyl cis-trans isomerase A (PPIase A) (EC 5,2,1,8) (Cyclophilin A) (Cyclosporin A-binding protein) (Rotamase A) [Cleaved into: Peptidyl-prolyl cis-trans isomerase A, N-terminally processed] | 0,316415071 | 0,048 | 0,325912134 |
| P35813 | PPM1A | Protein phosphatase 1A (EC 3,1,3,16) (Protein phosphatase 2C isoform alpha) (PP2C-alpha) (Protein phosphatase IA) | -0,269558097 | 0,04423 | 0,316650023 |
| P50897 | PPT1 | Palmitoyl-protein thioesterase 1 (PPT-1) (EC 3,1,2,22) (Palmitoyl-protein hydrolase 1) | 1,533775208 | 4,80E-06 | 0,003135415 |
| Q96S44 | PRPK | EKC/KEOPS complex subunit TP53RK (EC 3,6,-,-) (Atypical serine/threonine protein kinase TP53RK) (Nori-2) (TP53-regulating kinase) (EC 2,7,11,1) (p53-related protein kinase) | -0,317958247 | 0,02162 | 0,235752629 |
| P25789 | PSA4 | Proteasome subunit alpha type-4 (EC 3,4,25,1) (Macropain subunit C9) (Multicatalytic endopeptidase complex subunit C9) (Proteasome component C9) (Proteasome subunit L) | 0,614193154 | 0,01186 | 0,197987642 |
| P28066 | PSA5 | Proteasome subunit alpha type-5 (EC 3,4,25,1) (Macropain zeta chain) (Multicatalytic endopeptidase complex zeta chain) (Proteasome zeta chain) | 0,681483004 | 0,02053 | 0,234998667 |
| O14818 | PSA7 | Proteasome subunit alpha type-7 (EC 3,4,25,1) (Proteasome subunit RC6-1) (Proteasome subunit XAPC7) | 0,848369431 | 0,0034 | 0,104947021 |
| O00487 | PSDE | 26S proteasome non-ATPase regulatory subunit 14 (EC 3,4,19,-) (26S proteasome regulatory subunit RPN11) (26S proteasome-associated PAD1 homolog 1) | 0,646237319 | 0,00683 | 0,147445448 |
| Q15008 | PSMD6 | 26S proteasome non-ATPase regulatory subunit 6 (26S proteasome regulatory subunit RPN7) (26S proteasome regulatory subunit S10) (Breast cancer-associated protein SGA-113M) (Phosphonoformate immuno-associated protein 4) (Proteasome regulatory particle subunit p44S10) (p42A) | -0,292703702 | 0,03041 | 0,263871504 |
| Q8WXF1 | PSPC1 | Paraspeckle component 1 (Paraspeckle protein 1) | -0,459139976 | 0,03988 | 0,303141461 |
| P41222 | PTGDS | Prostaglandin-H2 D-isomerase (EC 5,3,99,2) (Beta-trace protein) (Cerebrin-28) (Glutathione-independent PGD synthase) (Lipocalin-type prostaglandin-D synthase) (Prostaglandin-D2 synthase) (PGD2 synthase) (PGDS) (PGDS2) | 0,871400006 | 0,01588 | 0,213209762 |
| Q96QR8 | PURB | Transcriptional activator protein Pur-beta (Purine-rich element-binding protein B) | 0,364497864 | 0,01909 | 0,230420592 |
| Q9NRF8 | PYRG2 | CTP synthase 2 (EC 6,3,4,2) (CTP synthetase 2) (UTP--ammonia ligase 2) | -0,482480628 | 0,00321 | 0,101469804 |
| Q96PU8 | QKI | Protein quaking (Hqk) (HqkI) | -0,442149093 | 0,00249 | 0,095855316 |
| P61026 | RAB10 | Ras-related protein Rab-10 | 0,631515474 | 0,00119 | 0,060334988 |
| Q9H0U4 | RAB1B | Ras-related protein Rab-1B | 0,766823383 | 0,00088 | 0,048773128 |
| Q5U651 | RAIN | Ras-interacting protein 1 (Rain) | -0,28625453 | 0,0292 | 0,259726687 |
| O14807 | RASM | Ras-related protein M-Ras (Ras-related protein R-Ras3) | 0,362232986 | 0,03907 | 0,299900797 |
| Q9NTZ6 | RBM12 | RNA-binding protein 12 (RNA-binding motif protein 12) (SH3/WW domain anchor protein in the nucleus) (SWAN) | -0,371303675 | 0,00561 | 0,135618752 |
| P54727 | RD23B | UV excision repair protein RAD23 homolog B (HR23B) (hHR23B) (XP-C repair-complementing complex 58 kDa protein) (p58) | 0,289320663 | 0,02466 | 0,241832734 |
| P46063 | RECQ1 | ATP-dependent DNA helicase Q1 (EC 3,6,4,12) (DNA helicase, RecQ-like type 1) (RecQ1) (DNA-dependent ATPase Q1) (RecQ protein-like 1) | 0,395371345 | 0,02255 | 0,236759134 |
| Q00765 | REEP5 | Receptor expression-enhancing protein 5 (Polyposis locus protein 1) (Protein TB2) | 0,46818852 | 0,02719 | 0,253562451 |
| O15211 | RGL2 | Ral guanine nucleotide dissociation stimulator-like 2 (RalGDS-like 2) (RalGDS-like factor) (Ras-associated protein RAB2L) | -0,258807981 | 0,03207 | 0,268193682 |
| Q15382 | RHEB | GTP-binding protein Rheb (Ras homolog enriched in brain) | 0,439616664 | 0,01779 | 0,225054195 |
| P52758 | RIDA | 2-iminobutanoate/2-iminopropanoate deaminase (EC 3,5,99,10) (14,5 kDa translational inhibitor protein) (hp14,5) (p14,5) (Heat-responsive protein 12) (Reactive intermediate imine deaminase A homolog) (Translation inhibitor L-PSP ribonuclease) (EC 3,1,-,-) (UK114 antigen homolog) | 0,879621764 | 0,00036 | 0,031592167 |
| Q96NA2 | RILP | Rab-interacting lysosomal protein | -0,418803889 | 0,03853 | 0,298767304 |
| Q92901 | RL3L | 60S ribosomal protein L3-like (Large ribosomal subunit protein uL3-like) | -0,353154573 | 0,01546 | 0,212766683 |
| Q7Z2W9 | RM21 | 39S ribosomal protein L21, mitochondrial (L21mt) (MRP-L21) (Mitochondrial large ribosomal subunit protein bL21m) | -0,43736359 | 0,00285 | 0,097030859 |
| Q5XPI4 | RN123 | E3 ubiquitin-protein ligase RNF123 (EC 2,3,2,27) (Kip1 ubiquitination-promoting complex protein 1) (RING finger protein 123) (RING-type E3 ubiquitin transferase RNF123) | -0,358960667 | 0,01929 | 0,230638173 |
| Q9HB90 | RRAGC | Ras-related GTP-binding protein C (Rag C) (RagC) (GTPase-interacting protein 2) (TIB929) | 0,406820699 | 0,00759 | 0,15668516 |
| P10301 | RRAS | Ras-related protein R-Ras (p23) | -0,367146002 | 0,02149 | 0,235752629 |
| P62854 | RS26 | 40S ribosomal protein S26 (Small ribosomal subunit protein eS26) | -0,365210522 | 0,02982 | 0,261063402 |
| P62857 | RS28 | 40S ribosomal protein S28 (Small ribosomal subunit protein eS28) | -0,388200813 | 0,04768 | 0,325912134 |
| P51398 | RT29 | 28S ribosomal protein S29, mitochondrial (MRP-S29) (S29mt) (Death-associated protein 3) (DAP-3) (Ionizing radiation resistance conferring protein) (Mitochondrial small ribosomal subunit protein mS29) | 0,370726974 | 0,04097 | 0,308530859 |
| O95197 | RTN3 | Reticulon-3 (Homolog of ASY protein) (HAP) (Neuroendocrine-specific protein-like 2) (NSP-like protein 2) (Neuroendocrine-specific protein-like II) (NSP-like protein II) (NSPLII) | 0,832714319 | 0,00021 | 0,021671075 |
| Q9Y265 | RUVB1 | RuvB-like 1 (EC 3,6,4,12) (49 kDa TATA box-binding protein-interacting protein) (49 kDa TBP-interacting protein) (54 kDa erythrocyte cytosolic protein) (ECP-54) (INO80 complex subunit H) (Nuclear matrix protein 238) (NMP 238) (Pontin 52) (TIP49a) (TIP60-associated protein 54-alpha) (TAP54-alpha) | -0,367776209 | 0,01176 | 0,197123561 |
| Q9Y230 | RUVB2 | RuvB-like 2 (EC 3,6,4,12) (48 kDa TATA box-binding protein-interacting protein) (48 kDa TBP-interacting protein) (51 kDa erythrocyte cytosolic protein) (ECP-51) (INO80 complex subunit J) (Repressing pontin 52) (Reptin 52) (TIP49b) (TIP60-associated protein 54-beta) (TAP54-beta) | -0,294926811 | 0,02468 | 0,241832734 |
| P60903 | S10AA | Protein S100-A10 (Calpactin I light chain) (Calpactin-1 light chain) (Cellular ligand of annexin II) (S100 calcium-binding protein A10) (p10 protein) (p11) | 0,997247052 | 0,00613 | 0,141437168 |
| Q9Y6Y8 | S23IP | SEC23-interacting protein (p125) | 0,354762273 | 0,01577 | 0,213209762 |
| Q99808 | S29A1 | Equilibrative nucleoside transporter 1 (Equilibrative nitrobenzylmercaptopurine riboside-sensitive nucleoside transporter) (Equilibrative NBMPR-sensitive nucleoside transporter) (Nucleoside transporter, es-type) (Solute carrier family 29 member 1) | 0,39951438 | 0,01094 | 0,188999161 |
| Q8N357 | S35F6 | Solute carrier family 35 member F6 (ANT2-binding protein) (ANT2BP) (Transport and Golgi organization 9 homolog) | 0,349303964 | 0,02039 | 0,234761058 |
| P0DJI8 | SAA1 | Serum amyloid A-1 protein (SAA) [Cleaved into: Amyloid protein A (Amyloid fibril protein AA); Serum amyloid protein A(2-104); Serum amyloid protein A(3-104); Serum amyloid protein A(2-103); Serum amyloid protein A(2-102); Serum amyloid protein A(4-101)] | 1,413603647 | 0,00139 | 0,064232864 |
| P0DJI9 | SAA2 | Serum amyloid A-2 protein (SAA2) | 2,286064587 | 0,00852 | 0,164055531 |
| P35542 | SAA4 | Serum amyloid A-4 protein (Constitutively expressed serum amyloid A protein) (C-SAA) | 1,695881123 | 0,02359 | 0,236759134 |
| P02743 | SAMP | Serum amyloid P-component (SAP) (9,5S alpha-1-glycoprotein) [Cleaved into: Serum amyloid P-component(1-203)] | 2,418954644 | 5,74E-06 | 0,003214276 |
| P07602 | SAP | Prosaposin (Proactivator polypeptide) [Cleaved into: Saposin-A (Protein A); Saposin-B-Val; Saposin-B (Cerebroside sulfate activator) (CSAct) (Dispersin) (Sphingolipid activator protein 1) (SAP-1) (Sulfatide/GM1 activator); Saposin-C (A1 activator) (Co-beta-glucosidase) (Glucosylceramidase activator) (Sphingolipid activator protein 2) (SAP-2); Saposin-D (Component C) (Protein C)] | 1,02447255 | 0,00097 | 0,052931003 |
| P17900 | SAP3 | Ganglioside GM2 activator (Cerebroside sulfate activator protein) (GM2-AP) (Sphingolipid activator protein 3) (SAP-3) [Cleaved into: Ganglioside GM2 activator isoform short] | 1,108499115 | 3,56E-05 | 0,009944827 |
| Q15020 | SART3 | Squamous cell carcinoma antigen recognized by T-cells 3 (SART-3) (Tat-interacting protein of 110 kDa) (Tip110) (p110 nuclear RNA-binding protein) | -0,286043546 | 0,03808 | 0,296947114 |
| Q96F10 | SAT2 | Diamine acetyltransferase 2 (EC 2,3,1,57) (Polyamine N-acetyltransferase 2) (Spermidine/spermine N(1)-acetyltransferase 2) (Thialysine N-epsilon-acetyltransferase) | -0,420264168 | 0,03087 | 0,266198755 |
| Q13228 | SBP1 | Methanethiol oxidase (MTO) (EC 1,8,3,4) (56 kDa selenium-binding protein) (SBP56) (SP56) (Selenium-binding protein 1) | 0,366863866 | 0,04908 | 0,327955691 |
| O15027 | SC16A | Protein transport protein Sec16A (SEC16 homolog A) | -0,406280181 | 0,01967 | 0,23220076 |
| P48061 | SDF1 | Stromal cell-derived factor 1 (SDF-1) (hSDF-1) (C-X-C motif chemokine 12) (Intercrine reduced in hepatomas) (IRH) (hIRH) (Pre-B cell growth-stimulating factor) (PBSF) [Cleaved into: SDF-1-beta(3-72); SDF-1-alpha(3-67)] | 2,165496888 | 1,73E-07 | 0,000489228 |
| Q99442 | SEC62 | Translocation protein SEC62 (Translocation protein 1) (TP-1) (hTP-1) | 0,592700531 | 0,00196 | 0,081863588 |
| Q96EE3 | SEH1 | Nucleoporin SEH1 (GATOR complex protein SEH1) (Nup107-160 subcomplex subunit SEH1) (SEC13-like protein) | -0,925045292 | 0,00265 | 0,097030859 |
| Q14141 | SEPT6 | Septin-6 | 0,959614523 | 0,00011 | 0,014543495 |
| Q15637 | SF01 | Splicing factor 1 (Mammalian branch point-binding protein) (BBP) (mBBP) (Transcription factor ZFM1) (Zinc finger gene in MEN1 locus) (Zinc finger protein 162) | -0,457115162 | 0,01767 | 0,224990548 |
| Q12874 | SF3A3 | Splicing factor 3A subunit 3 (SF3a60) (Spliceosome-associated protein 61) (SAP 61) | -0,340774728 | 0,02971 | 0,261063402 |
| O75533 | SF3B1 | Splicing factor 3B subunit 1 (Pre-mRNA-splicing factor SF3b 155 kDa subunit) (SF3b155) (Spliceosome-associated protein 155) (SAP 155) | -0,499451429 | 0,00628 | 0,141663963 |
| Q13435 | SF3B2 | Splicing factor 3B subunit 2 (Pre-mRNA-splicing factor SF3b 145 kDa subunit) (SF3b145) (Spliceosome-associated protein 145) (SAP 145) | -0,409425193 | 0,01649 | 0,215479037 |
| Q9Y3B4 | SF3B6 | Splicing factor 3B subunit 6 (Pre-mRNA branch site protein p14) (SF3b 14 kDa subunit) (SF3B14a) (Spliceosome-associated protein, 14-kDa) (Splicing factor 3b, subunit 6, 14kDa) | -0,317663408 | 0,04515 | 0,320244891 |
| Q16585 | SGCB | Beta-sarcoglycan (Beta-SG) (43 kDa dystrophin-associated glycoprotein) (43DAG) (A3b) | 0,392467041 | 0,03156 | 0,268193682 |
| O43556 | SGCE | Epsilon-sarcoglycan (Epsilon-SG) | -0,480054632 | 0,04336 | 0,313753743 |
| P55822 | SH3BG | SH3 domain-binding glutamic acid-rich protein (SH3BGR protein) (21-glutamic acid-rich protein) (21-GARP) | 0,588326566 | 0,00077 | 0,047747284 |
| O75368 | SH3L1 | SH3 domain-binding glutamic acid-rich-like protein | 0,567490522 | 0,00062 | 0,043603785 |
| Q9H299 | SH3L3 | SH3 domain-binding glutamic acid-rich-like protein 3 (SH3 domain-binding protein 1) (SH3BP-1) | 0,706602943 | 0,0033 | 0,102722264 |
| Q9BYB0 | SHAN3 | SH3 and multiple ankyrin repeat domains protein 3 (Shank3) (Proline-rich synapse-associated protein 2) (ProSAP2) | -0,284342623 | 0,04222 | 0,311743911 |
| Q9HAT2 | SIAE | Sialate O-acetylesterase (EC 3,1,1,53) (H-Lse) (Sialic acid-specific 9-O-acetylesterase) | 0,540457304 | 0,02508 | 0,243520716 |
| Q9NR45 | SIAS | Sialic acid synthase (N-acetylneuraminate synthase) (EC 2,5,1,56) (N-acetylneuraminate-9-phosphate synthase) (EC 2,5,1,57) (N-acetylneuraminic acid phosphate synthase) (N-acetylneuraminic acid synthase) | 0,467438678 | 0,01084 | 0,188957641 |
| P0DJ93 | SIM13 | Small integral membrane protein 13 | -0,324653278 | 0,02282 | 0,236759134 |
| Q9P270 | SLAI2 | SLAIN motif-containing protein 2 | -1,227745258 | 0,00018 | 0,019801709 |
| Q9NWH9 | SLTM | SAFB-like transcription modulator (Modulator of estrogen-induced transcription) | -0,365377822 | 0,02197 | 0,235814224 |
| P62314 | SMD1 | Small nuclear ribonucleoprotein Sm D1 (Sm-D1) (Sm-D autoantigen) (snRNP core protein D1) | -0,907142443 | 0,03291 | 0,271696224 |
| Q99747 | SNAG | Gamma-soluble NSF attachment protein (SNAP-gamma) (N-ethylmaleimide-sensitive factor attachment protein gamma) | 0,356077545 | 0,0271 | 0,253562451 |
| P00441 | SODC | Superoxide dismutase [Cu-Zn] (EC 1,15,1,1) (Superoxide dismutase 1) (hSod1) | 0,941224557 | 0,02311 | 0,236759134 |
| P04179 | SODM | Superoxide dismutase [Mn], mitochondrial (EC 1,15,1,1) | 0,465773878 | 0,02079 | 0,234998667 |
| O75391 | SPAG7 | Sperm-associated antigen 7 | -0,322556783 | 0,01614 | 0,213209762 |
| P50453 | SPB9 | Serpin B9 (Cytoplasmic antiproteinase 3) (CAP-3) (CAP3) (Peptidase inhibitor 9) (PI-9) | -0,365625845 | 0,03493 | 0,281900627 |
| Q15005 | SPCS2 | Signal peptidase complex subunit 2 (EC 3,4,-,-) (Microsomal signal peptidase 25 kDa subunit) (SPase 25 kDa subunit) | 0,923242795 | 0,00647 | 0,143094484 |
| Q9UQ90 | SPG7 | Paraplegin (EC 3,4,24,-) (Cell matrix adhesion regulator) (Spastic paraplegia 7 protein) | -0,298030001 | 0,04657 | 0,324973393 |
| Q86TD4 | SRCA | Sarcalumenin | -0,53400295 | 0,00084 | 0,047830967 |
| P78539 | SRPX | Sushi repeat-containing protein SRPX | 1,563656662 | 1,68E-05 | 0,007375622 |
| Q9UQ35 | SRRM2 | Serine/arginine repetitive matrix protein 2 (300 kDa nuclear matrix antigen) (Serine/arginine-rich splicing factor-related nuclear matrix protein of 300 kDa) (SR-related nuclear matrix protein of 300 kDa) (Ser/Arg-related nuclear matrix protein of 300 kDa) (Splicing coactivator subunit SRm300) (Tax-responsive enhancer element-binding protein 803) (TaxREB803) | -0,265100187 | 0,04731 | 0,325912134 |
| O75494 | SRS10 | Serine/arginine-rich splicing factor 10 (40 kDa SR-repressor protein) (SRrp40) (FUS-interacting serine-arginine-rich protein 1) (Splicing factor SRp38) (Splicing factor, arginine/serine-rich 13A) (TLS-associated protein with Ser-Arg repeats) (TASR) (TLS-associated protein with SR repeats) (TLS-associated serine-arginine protein) (TLS-associated SR protein) | -0,336754812 | 0,04788 | 0,325912134 |
| Q01130 | SRSF2 | Serine/arginine-rich splicing factor 2 (Protein PR264) (Splicing component, 35 kDa) (Splicing factor SC35) (SC-35) (Splicing factor, arginine/serine-rich 2) | -0,418166176 | 0,00964 | 0,175766361 |
| P50225 | ST1A1 | Sulfotransferase 1A1 (ST1A1) (EC 2,8,2,1) (Aryl sulfotransferase 1) (HAST1/HAST2) (Phenol sulfotransferase 1) (Phenol-sulfating phenol sulfotransferase 1) (P-PST 1) (ST1A3) (Thermostable phenol sulfotransferase) (Ts-PST) | 1,174600114 | 0,01869 | 0,226221613 |
| Q92783 | STAM1 | Signal transducing adapter molecule 1 (STAM-1) | 0,480111741 | 0,02311 | 0,236759134 |
| P27105 | STOM | Erythrocyte band 7 integral membrane protein (Protein 7,2b) (Stomatin) | -0,265835898 | 0,0336 | 0,27485911 |
| P61956 | SUMO2 | Small ubiquitin-related modifier 2 (SUMO-2) (HSMT3) (SMT3 homolog 2) (SUMO-3) (Sentrin-2) (Ubiquitin-like protein SMT3B) (Smt3B) | 0,421743953 | 0,02219 | 0,235814224 |
| P51687 | SUOX | Sulfite oxidase, mitochondrial (EC 1,8,3,1) | 0,452496674 | 0,00589 | 0,138298051 |
| O15260 | SURF4 | Surfeit locus protein 4 | -0,280898776 | 0,0494 | 0,328959493 |
| Q8NHG7 | SVIP | Small VCP/p97-interacting protein | -0,457860497 | 0,01338 | 0,201757374 |
| P49590 | SYHM | Probable histidine--tRNA ligase, mitochondrial (EC 6,1,1,21) (Histidine--tRNA ligase-like) (Histidyl-tRNA synthetase) (HisRS) | -0,325783748 | 0,03036 | 0,263871504 |
| P57105 | SYJ2B | Synaptojanin-2-binding protein (Mitochondrial outer membrane protein 25) | -0,288630213 | 0,03659 | 0,289923172 |
| Q15031 | SYLM | Probable leucine--tRNA ligase, mitochondrial (EC 6,1,1,4) (Leucyl-tRNA synthetase) (LeuRS) | -0,419875577 | 0,02253 | 0,236759134 |
| Q5VXT5 | SYPL2 | Synaptophysin-like protein 2 | -0,549509248 | 0,00632 | 0,141663963 |
| Q5T160 | SYRM | Probable arginine--tRNA ligase, mitochondrial (EC 6,1,1,19) (Arginyl-tRNA synthetase) (ArgRS) | -0,353706625 | 0,02979 | 0,261063402 |
| P37840 | SYUA | Alpha-synuclein (Non-A beta component of AD amyloid) (Non-A4 component of amyloid precursor) (NACP) | 0,875506701 | 0,00272 | 0,097030859 |
| Q9Y2Z4 | SYYM | Tyrosine--tRNA ligase, mitochondrial (EC 6,1,1,1) (Tyrosyl-tRNA synthetase) (TyrRS) | -0,323433071 | 0,01432 | 0,208833198 |
| Q9H061 | T126A | Transmembrane protein 126A | 0,353467816 | 0,00995 | 0,178189563 |
| Q9C0D5 | TANC1 | Protein TANC1 (Tetratricopeptide repeat, ankyrin repeat and coiled-coil domain-containing protein 1) | -0,383565768 | 0,04864 | 0,327613246 |
| Q6NXT6 | TAPT1 | Transmembrane anterior posterior transformation protein 1 homolog (Cytomegalovirus partial fusion receptor) | -0,467133054 | 0,01602 | 0,213209762 |
| Q7Z7G0 | TARSH | Target of Nesh-SH3 (Tarsh) (ABI gene family member 3-binding protein) (Nesh-binding protein) (NeshBP) | 1,032685804 | 0,00014 | 0,017783943 |
| P10636 | TAU | Microtubule-associated protein tau (Neurofibrillary tangle protein) (Paired helical filament-tau) (PHF-tau) | -0,862610732 | 0,0039 | 0,114007879 |
| A6NHL2 | TBAL3 | Tubulin alpha chain-like 3 | -0,791775244 | 0,0017 | 0,074278921 |
| Q9HA65 | TBC17 | TBC1 domain family member 17 | -0,449746176 | 0,01935 | 0,230638173 |
| Q9ULP9 | TBC24 | TBC1 domain family member 24 | -0,286419084 | 0,04393 | 0,315790315 |
| Q92609 | TBCD5 | TBC1 domain family member 5 | -0,274128581 | 0,04303 | 0,312814857 |
| O95759 | TBCD8 | TBC1 domain family member 8 (AD 3) (Vascular Rab-GAP/TBC-containing protein) | -0,323653285 | 0,0497 | 0,329845885 |
| Q15185 | TEBP | Prostaglandin E synthase 3 (EC 5,3,99,3) (Cytosolic prostaglandin E2 synthase) (cPGES) (Hsp90 co-chaperone) (Progesterone receptor complex p23) (Telomerase-binding protein p23) | 0,56837719 | 0,0099 | 0,178117363 |
| Q9UKN8 | TF3C4 | General transcription factor 3C polypeptide 4 (EC 2,3,1,48) (TF3C-delta) (Transcription factor IIIC 90 kDa subunit) (TFIIIC 90 kDa subunit) (TFIIIC90) (Transcription factor IIIC subunit delta) | -0,564078621 | 0,00126 | 0,062074957 |
| O43493 | TGON2 | Trans-Golgi network integral membrane protein 2 (TGN38 homolog) (TGN46) (TGN48) (Trans-Golgi network protein TGN51) | 0,600828779 | 0,0032 | 0,101469804 |
| Q9BXR0 | TGT | Queuine tRNA-ribosyltransferase catalytic subunit 1 (EC 2,4,2,29) (Guanine insertion enzyme) (tRNA-guanine transglycosylase) | -0,382573702 | 0,01765 | 0,224990548 |
| P05543 | THBG | Thyroxine-binding globulin (Serpin A7) (T4-binding globulin) | 0,92910598 | 0,00272 | 0,097030859 |
| P10599 | THIO | Thioredoxin (Trx) (ATL-derived factor) (ADF) (Surface-associated sulphydryl protein) (SASP) | 0,371767394 | 0,02117 | 0,234998667 |
| Q9Y3D7 | TIM16 | Mitochondrial import inner membrane translocase subunit TIM16 (Mitochondria-associated granulocyte macrophage CSF-signaling molecule) (Presequence translocated-associated motor subunit PAM16) | -0,310016705 | 0,02486 | 0,241943883 |
| Q9Y5J7 | TIM9 | Mitochondrial import inner membrane translocase subunit Tim9 | -0,320688325 | 0,04588 | 0,323065719 |
| P35625 | TIMP3 | Metalloproteinase inhibitor 3 (Protein MIG-5) (Tissue inhibitor of metalloproteinases 3) (TIMP-3) | 3,116116312 | 4,97E-05 | 0,011475286 |
| Q9Y3B3 | TMED7 | Transmembrane emp24 domain-containing protein 7 (p24 family protein gamma-3) (p24gamma3) (p27) | 0,496322299 | 0,00922 | 0,172373188 |
| P49755 | TMEDA | Transmembrane emp24 domain-containing protein 10 (21 kDa transmembrane-trafficking protein) (S31III125) (S31I125) (Tmp-21-I) (Transmembrane protein Tmp21) (p23) (p24 family protein delta-1) (p24delta1) (p24delta) | 0,437258062 | 0,01506 | 0,212742239 |
| Q9NYL9 | TMOD3 | Tropomodulin-3 (Ubiquitous tropomodulin) (U-Tmod) | 0,498262616 | 0,00408 | 0,114007879 |
| P19237 | TNNI1 | Troponin I, slow skeletal muscle (Troponin I, slow-twitch isoform) | -1,229437232 | 0,04882 | 0,32788441 |
| P45379 | TNNT2 | Troponin T, cardiac muscle (TnTc) (Cardiac muscle troponin T) (cTnT) | -0,255910266 | 0,04726 | 0,325912134 |
| Q63HR2 | TNS2 | Tensin-2 (EC 3,1,3,-) (C1 domain-containing phosphatase and tensin homolog) (C1-TEN) (Tensin-like C1 domain-containing phosphatase) | -0,420985419 | 0,02096 | 0,234998667 |
| Q9NS69 | TOM22 | Mitochondrial import receptor subunit TOM22 homolog (hTom22) (1C9-2) (Translocase of outer membrane 22 kDa subunit homolog) | 0,280155551 | 0,02228 | 0,23612256 |
| Q15785 | TOM34 | Mitochondrial import receptor subunit TOM34 (hTom34) (Translocase of outer membrane 34 kDa subunit) | 0,589381449 | 0,00842 | 0,163422907 |
| O94826 | TOM70 | Mitochondrial import receptor subunit TOM70 (Mitochondrial precursor proteins import receptor) (Translocase of outer membrane 70 kDa subunit) (Translocase of outer mitochondrial membrane protein 70) | 0,486695062 | 0,00399 | 0,114007879 |
| Q86SZ2 | TPC6B | Trafficking protein particle complex subunit 6B | -0,326267778 | 0,03617 | 0,287727408 |
| P55327 | TPD52 | Tumor protein D52 (Protein N8) | 0,343529193 | 0,02868 | 0,257625811 |
| O43399 | TPD54 | Tumor protein D54 (hD54) (Tumor protein D52-like 2) | 0,519674182 | 0,03153 | 0,268193682 |
| P09493 | TPM1 | Tropomyosin alpha-1 chain (Alpha-tropomyosin) (Tropomyosin-1) | 0,298130157 | 0,03917 | 0,300019036 |
| P51580 | TPMT | Thiopurine S-methyltransferase (EC 2,1,1,67) (Thiopurine methyltransferase) | -0,374388641 | 0,01483 | 0,211823261 |
| Q8IUR0 | TPPC5 | Trafficking protein particle complex subunit 5 | -0,301305697 | 0,04783 | 0,325912134 |
| Q9BX59 | TPSNR | Tapasin-related protein (TAPASIN-R) (TAP-binding protein-like) (TAP-binding protein-related protein) (TAPBP-R) (Tapasin-like) | -0,583475525 | 0,03886 | 0,299900797 |
| P02787 | TRFE | Serotransferrin (Transferrin) (Beta-1 metal-binding globulin) (Siderophilin) | 0,95478388 | 0,0003 | 0,027785329 |
| Q6ZMU5 | TRI72 | Tripartite motif-containing protein 72 (Mitsugumin-53) (Mg53) | -0,617026092 | 0,01449 | 0,210422921 |
| Q96Q11 | TRNT1 | CCA tRNA nucleotidyltransferase 1, mitochondrial (EC 2,7,7,72) (Mitochondrial tRNA nucleotidyl transferase, CCA-adding) (mt CCA-adding enzyme) (mt tRNA CCA-diphosphorylase) (mt tRNA CCA-pyrophosphorylase) (mt tRNA adenylyltransferase) | 0,406305949 | 0,01596 | 0,213209762 |
| Q8WWH5 | TRUB1 | Probable tRNA pseudouridine synthase 1 (EC 5,4,99,-) | 0,394011698 | 0,0045 | 0,121657442 |
| P49815 | TSC2 | Tuberin (Tuberous sclerosis 2 protein) | -0,31724092 | 0,01833 | 0,226221613 |
| Q99598 | TSNAX | Translin-associated protein X (Translin-associated factor X) | -0,398638477 | 0,03736 | 0,292457436 |
| P35443 | TSP4 | Thrombospondin-4 | 0,636156207 | 0,01137 | 0,193012395 |
| P30536 | TSPO | Translocator protein (Mitochondrial benzodiazepine receptor) (PKBS) (Peripheral-type benzodiazepine receptor) (PBR) | 1,221782806 | 3,17E-05 | 0,009944827 |
| Q86TV6 | TTC7B | Tetratricopeptide repeat protein 7B (TPR repeat protein 7B) (Tetratricopeptide repeat protein 7-like-1) (TPR repeat protein 7-like-1) | -0,373417415 | 0,01713 | 0,220982924 |
| O14907 | TX1B3 | Tax1-binding protein 3 (Glutaminase-interacting protein 3) (Tax interaction protein 1) (TIP-1) (Tax-interacting protein 1) | 0,472402384 | 0,02957 | 0,261063402 |
| Q9H3M7 | TXNIP | Thioredoxin-interacting protein (Thioredoxin-binding protein 2) (Vitamin D3 up-regulated protein 1) | -0,984051523 | 0,00351 | 0,106830512 |
| P53007 | TXTP | Tricarboxylate transport protein, mitochondrial (Citrate transport protein) (CTP) (Solute carrier family 25 member 1) (Tricarboxylate carrier protein) | 0,509857341 | 0,04059 | 0,307888961 |
| P63313 | TYB10 | Thymosin beta-10 | 0,848477052 | 0,02397 | 0,239194989 |
| P62328 | TYB4 | Thymosin beta-4 (T beta-4) (Fx) [Cleaved into: Hematopoietic system regulatory peptide (Seraspenide)] | 0,895633135 | 0,00169 | 0,074278921 |
| P68036 | UB2L3 | Ubiquitin-conjugating enzyme E2 L3 (EC 2,3,2,23) (E2 ubiquitin-conjugating enzyme L3) (L-UBC) (UbcH7) (Ubiquitin carrier protein L3) (Ubiquitin-conjugating enzyme E2-F1) (Ubiquitin-protein ligase L3) | 0,399808188 | 0,04803 | 0,325912134 |
| Q8WVY7 | UBCP1 | Ubiquitin-like domain-containing CTD phosphatase 1 (EC 3,1,3,16) (Nuclear proteasome inhibitor UBLCP1) | 0,33425522 | 0,01211 | 0,199766061 |
| P62256 | UBE2H | Ubiquitin-conjugating enzyme E2 H (EC 2,3,2,23) ((E3-independent) E2 ubiquitin-conjugating enzyme H) (EC 2,3,2,24) (E2 ubiquitin-conjugating enzyme H) (UbcH2) (Ubiquitin carrier protein H) (Ubiquitin-conjugating enzyme E2-20K) (Ubiquitin-protein ligase H) | 0,677933751 | 0,0001 | 0,014543495 |
| O95155 | UBE4B | Ubiquitin conjugation factor E4 B (EC 2,3,2,27) (Homozygously deleted in neuroblastoma 1) (RING-type E3 ubiquitin transferase E4 B) (Ubiquitin fusion degradation protein 2) | -0,353634833 | 0,01559 | 0,212766683 |
| Q04323 | UBXN1 | UBX domain-containing protein 1 (SAPK substrate protein 1) (UBA/UBX 33,3 kDa protein) | -0,311102521 | 0,03575 | 0,285591713 |
| Q96QD9 | UIF | UAP56-interacting factor (Forty-two-three domain-containing protein 1) (Protein 40-2-3) | -0,510167703 | 0,01265 | 0,199766061 |
| Q9NVA1 | UQCC1 | Ubiquinol-cytochrome-c reductase complex assembly factor 1 (Basic FGF-repressed Zic-binding protein) (bFGF-repressed Zic-binding protein) (bFZb) (Ubiquinol-cytochrome c reductase complex chaperone CBP3 homolog) | 0,304744194 | 0,04639 | 0,324911402 |
| Q9BRT2 | UQCC2 | Ubiquinol-cytochrome-c reductase complex assembly factor 2 (Breast cancer-associated protein SGA-81M) (Mitochondrial nucleoid factor 1) (Mitochondrial protein M19) | 0,402438822 | 0,01155 | 0,194487304 |
| Q96IX5 | USMG5 | Up-regulated during skeletal muscle growth protein 5 (Diabetes-associated protein in insulin-sensitive tissues) (HCV F-transactivated protein 2) | 0,771627952 | 0,00113 | 0,059039382 |
| Q9Y5K8 | VATD | V-type proton ATPase subunit D (V-ATPase subunit D) (V-ATPase 28 kDa accessory protein) (Vacuolar proton pump subunit D) | -0,72699729 | 0,00281 | 0,097030859 |
| Q9UI12 | VATH | V-type proton ATPase subunit H (V-ATPase subunit H) (Nef-binding protein 1) (NBP1) (Protein VMA13 homolog) (V-ATPase 50/57 kDa subunits) (Vacuolar proton pump subunit H) (Vacuolar proton pump subunit SFD) | -0,371973746 | 0,01333 | 0,201757374 |
| P21796 | VDAC1 | Voltage-dependent anion-selective channel protein 1 (VDAC-1) (hVDAC1) (Outer mitochondrial membrane protein porin 1) (Plasmalemmal porin) (Porin 31HL) (Porin 31HM) | 0,584731785 | 0,0008 | 0,047747284 |
| P45880 | VDAC2 | Voltage-dependent anion-selective channel protein 2 (VDAC-2) (hVDAC2) (Outer mitochondrial membrane protein porin 2) | 0,576324125 | 0,00071 | 0,045663466 |
| Q9Y277 | VDAC3 | Voltage-dependent anion-selective channel protein 3 (VDAC-3) (hVDAC3) (Outer mitochondrial membrane protein porin 3) | 1,209162346 | 0,00011 | 0,014543495 |
| P08670 | VIME | Vimentin | 0,351722151 | 0,02133 | 0,234998667 |
| P98155 | VLDLR | Very low-density lipoprotein receptor (VLDL receptor) (VLDL-R) | 0,588996776 | 0,00468 | 0,125487524 |
| Q93050 | VPP1 | V-type proton ATPase 116 kDa subunit a isoform 1 (V-ATPase 116 kDa isoform a1) (Clathrin-coated vesicle/synaptic vesicle proton pump 116 kDa subunit) (Vacuolar adenosine triphosphatase subunit Ac116) (Vacuolar proton pump subunit 1) (Vacuolar proton translocating ATPase 116 kDa subunit a isoform 1) | -0,406377097 | 0,00629 | 0,141663963 |
| Q9NRW7 | VPS45 | Vacuolar protein sorting-associated protein 45 (h-VPS45) (hlVps45) | -0,340073446 | 0,01794 | 0,225054195 |
| O75351 | VPS4B | Vacuolar protein sorting-associated protein 4B (EC 3,6,4,6) (Cell migration-inducing gene 1 protein) (Suppressor of K(+) transport growth defect 1) (Protein SKD1) | 0,409106738 | 0,00374 | 0,111128847 |
| Q9UID3 | VPS51 | Vacuolar protein sorting-associated protein 51 homolog (Another new gene 2 protein) (Protein fat-free homolog) | -0,378283982 | 0,02517 | 0,243710647 |
| Q8N3P4 | VPS8 | Vacuolar protein sorting-associated protein 8 homolog | -0,31038252 | 0,02822 | 0,257481581 |
| P02774 | VTDB | Vitamin D-binding protein (DBP) (VDB) (Gc protein-derived macrophage activating factor) (Gc-MAF) (GcMAF) (Gc-globulin) (Group-specific component) (Gc) (Vitamin D-binding protein-macrophage activating factor) (DBP-maf) | 0,969269872 | 0,01668 | 0,216561717 |
| P04004 | VTNC | Vitronectin (VN) (S-protein) (Serum-spreading factor) (V75) [Cleaved into: Vitronectin V65 subunit; Vitronectin V10 subunit; Somatomedin-B] | 1,380444357 | 0,00129 | 0,062074957 |
| Q641Q2 | WAC2A | WASH complex subunit 2A | 0,223636646 | 0,04783 | 0,325912134 |
| Q6UXN9 | WDR82 | WD repeat-containing protein 82 (Protein TMEM113) (Swd2) | -0,388973499 | 0,03429 | 0,278977054 |
| Q96S55 | WRIP1 | ATPase WRNIP1 (EC 3,6,1,3) (Werner helicase-interacting protein 1) | -0,346464877 | 0,01262 | 0,199766061 |
| Q8N9V3 | WSDU1 | WD repeat, SAM and U-box domain-containing protein 1 | -0,363378979 | 0,0335 | 0,274849318 |
| Q9NQW7 | XPP1 | Xaa-Pro aminopeptidase 1 (EC 3,4,11,9) (Aminoacylproline aminopeptidase) (Cytosolic aminopeptidase P) (Soluble aminopeptidase P) (sAmp) (X-Pro aminopeptidase 1) (X-prolyl aminopeptidase 1, soluble) | 0,306314453 | 0,03178 | 0,268193682 |
| P25311 | ZA2G | Zinc-alpha-2-glycoprotein (Zn-alpha-2-GP) (Zn-alpha-2-glycoprotein) | 1,296882113 | 0,00351 | 0,106830512 |
| Q5BKZ1 | ZN326 | DBIRD complex subunit ZNF326 (Zinc finger protein 326) (Zinc finger protein interacting with mRNPs and DBC1) | -0,426689522 | 0,0453 | 0,320684198 |
| Q07157 | ZO1 | Tight junction protein ZO-1 (Tight junction protein 1) (Zona occludens protein 1) (Zonula occludens protein 1) | -0,292366834 | 0,03609 | 0,287688149 |
| Q9UK55 | ZPI | Protein Z-dependent protease inhibitor (PZ-dependent protease inhibitor) (PZI) (Serpin A10) | 1,096820041 | 0,00027 | 0,025999051 |

**Supplementary Data Table 8. List of total significantly altered proteins (p-value<0.05) in HGPS humans**

| **Uniprot ID** | **Protein Symbol** | **Protein Description** | **Average Log2 Ratio** | ***p*-value** | **Adjusted *p*-value** |
| --- | --- | --- | --- | --- | --- |
| P31946 | 1433B | 14-3-3 protein beta/alpha (Protein 1054) (Protein kinase C inhibitor protein 1) (KCIP-1) [Cleaved into: 14-3-3 protein beta/alpha, N-terminally processed] | -0,467086537 | 0,02468 | 0.150055430955809 |
| Q13542 | 4EBP2 | Eukaryotic translation initiation factor 4E-binding protein 2 (4E-BP2) (eIF4E-binding protein 2) | -1,028776345 | 0,04301 | 0.188876486048564 |
| O95336 | 6PGL | 6-phosphogluconolactonase (6PGL) (EC 3.1.1.31) | 0,577829573 | 0,01063 | 0.109626270622104 |
| Q8IZ83 | A16A1 | Aldehyde dehydrogenase family 16 member A1 | 0,694486598 | 0,01096 | 0.110439860103028 |
| P04217 | A1BG | Alpha-1B-glycoprotein (Alpha-1-B glycoprotein) | 2,353673143 | 0,00062 | 0.0460405438333303 |
| P08697 | A2AP | Alpha-2-antiplasmin (Alpha-2-AP) (Alpha-2-plasmin inhibitor) (Alpha-2-PI) (Serpin F2) | 1,277200906 | 0,0002 | 0.0423116847048915 |
| P01023 | A2MG | Alpha-2-macroglobulin (Alpha-2-M) (C3 and PZP-like alpha-2-macroglobulin domain-containing protein 5) | 2,864087645 | 0,00053 | 0.0460405438333303 |
| Q96GX2 | A7L3B | Ataxin-7-like protein 3B | -0,441760757 | 0,02277 | 0.146428755002108 |
| O43741 | AAKB2 | 5'-AMP-activated protein kinase subunit beta-2 (AMPK subunit beta-2) | -0,565727575 | 0,00795 | 0.102154997647559 |
| P54619 | AAKG1 | 5'-AMP-activated protein kinase subunit gamma-1 (AMPK gamma1) (AMPK subunit gamma-1) (AMPKg) | -0,57744455 | 0,00889 | 0.103882752534462 |
| Q13131 | AAPK1 | 5'-AMP-activated protein kinase catalytic subunit alpha-1 (AMPK subunit alpha-1) (EC 2.7.11.1) (Acetyl-CoA carboxylase kinase) (ACACA kinase) (EC 2.7.11.27) (Hydroxymethylglutaryl-CoA reductase kinase) (HMGCR kinase) (EC 2.7.11.31) (Tau-protein kinase PRKAA1) (EC 2.7.11.26) | 0,594928116 | 0,01546 | 0.124751718030612 |
| Q9UDR5 | AASS | Alpha-aminoadipic semialdehyde synthase, mitochondrial (LKR/SDH) [Includes: Lysine ketoglutarate reductase (LKR) (LOR) (EC 1.5.1.8); Saccharopine dehydrogenase (SDH) (EC 1.5.1.9)] | -0,715119746 | 0,00365 | 0.0767928025552317 |
| O94911 | ABCA8 | ATP-binding cassette sub-family A member 8 | -0,719187502 | 0,01743 | 0.13109533821508 |
| O60706 | ABCC9 | ATP-binding cassette sub-family C member 9 (Sulfonylurea receptor 2) | 0,600097196 | 0,0158 | 0.126187856364557 |
| P08910 | ABHD2 | Monoacylglycerol lipase ABHD2 (EC 3.1.1.23) (2-arachidonoylglycerol hydrolase) (Abhydrolase domain-containing protein 2) (Lung alpha/beta hydrolase 2) (Protein PHPS1-2) | -2,035879268 | 0,0005 | 0.0460405438333303 |
| Q9NUJ1 | ABHDA | Mycophenolic acid acyl-glucuronide esterase, mitochondrial (EC 3.1.1.93) (Alpha/beta hydrolase domain-containing protein 10) (Abhydrolase domain-containing protein 10) | -0,712861603 | 0,00308 | 0.0761554511764352 |
| O95870 | ABHGA | Protein ABHD16A (EC 3.-.-.-) (Alpha/beta hydrolase domain-containing protein 16A) (Abhydrolase domain-containing protein 16A) (HLA-B-associated transcript 5) (Protein G5) | 0,419625868 | 0,04507 | 0.194027039546718 |
| O94929 | ABLM3 | Actin-binding LIM protein 3 (abLIM-3) (Actin-binding LIM protein family member 3) | -0,501911288 | 0,01954 | 0.137096124047053 |
| Q15057 | ACAP2 | Arf-GAP with coiled-coil, ANK repeat and PH domain-containing protein 2 (Centaurin-beta-2) (Cnt-b2) | 0,555613213 | 0,00841 | 0.103396212083272 |
| P45954 | ACDSB | Short/branched chain specific acyl-CoA dehydrogenase, mitochondrial (SBCAD) (EC 1.3.8.5) (2-methyl branched chain acyl-CoA dehydrogenase) (2-MEBCAD) (2-methylbutyryl-coenzyme A dehydrogenase) (2-methylbutyryl-CoA dehydrogenase) | -0,542645974 | 0,0142 | 0.120558249672262 |
| Q9UKV3 | ACINU | Apoptotic chromatin condensation inducer in the nucleus (Acinus) | 0,440990826 | 0,02346 | 0.147127385900632 |
| O96019 | ACL6A | Actin-like protein 6A (53 kDa BRG1-associated factor A) (Actin-related protein Baf53a) (ArpNbeta) (BRG1-associated factor 53A) (BAF53A) (INO80 complex subunit K) | 0,472409518 | 0,0218 | 0.142960405215203 |
| P53396 | ACLY | ATP-citrate synthase (EC 2.3.3.8) (ATP-citrate (pro-S-)-lyase) (ACL) (Citrate cleavage enzyme) | 0,643160287 | 0,00751 | 0.101606052982629 |
| Q9NPJ3 | ACO13 | Acyl-coenzyme A thioesterase 13 (Acyl-CoA thioesterase 13) (EC 3.1.2.-) (Thioesterase superfamily member 2) [Cleaved into: Acyl-coenzyme A thioesterase 13, N-terminally processed] | -0,640789497 | 0,00664 | 0.0960549400412821 |
| O14561 | ACPM | Acyl carrier protein, mitochondrial (ACP) (CI-SDAP) (NADH-ubiquinone oxidoreductase 9.6 kDa subunit) | -0,788000912 | 0,01567 | 0.125898326767456 |
| Q96CM8 | ACSF2 | Acyl-CoA synthetase family member 2, mitochondrial (EC 6.2.1.-) | -0,937194019 | 0,00156 | 0.0666730305919975 |
| O95573 | ACSL3 | Long-chain-fatty-acid--CoA ligase 3 (EC 6.2.1.3) (Long-chain acyl-CoA synthetase 3) (LACS 3) | 0,674281576 | 0,01881 | 0.13488646867884 |
| Q08043 | ACTN3 | Alpha-actinin-3 (Alpha-actinin skeletal muscle isoform 3) (F-actin cross-linking protein) | 1,200812439 | 0,00105 | 0.0549153122487198 |
| O14672 | ADA10 | Disintegrin and metalloproteinase domain-containing protein 10 (ADAM 10) (EC 3.4.24.81) (CDw156) (Kuzbanian protein homolog) (Mammalian disintegrin-metalloprotease) (CD antigen CD156c) | 0,661690989 | 0,0124 | 0.114216522917893 |
| P78536 | ADA17 | Disintegrin and metalloproteinase domain-containing protein 17 (ADAM 17) (EC 3.4.24.86) (Snake venom-like protease) (TNF-alpha convertase) (TNF-alpha-converting enzyme) (CD antigen CD156b) | 0,867718431 | 0,0018 | 0.0697416810103427 |
| O43306 | ADCY6 | Adenylate cyclase type 6 (EC 4.6.1.1) (ATP pyrophosphate-lyase 6) (Adenylate cyclase type VI) (Adenylyl cyclase 6) (Ca(2+)-inhibitable adenylyl cyclase) | 0,655305306 | 0,01798 | 0.132658871887263 |
| Q8TED9 | AF1L1 | Actin filament-associated protein 1-like 1 (AFAP1-like protein 1) | 0,885028 | 0,00617 | 0.0954651715986055 |
| P43652 | AFAM | Afamin (Alpha-albumin) (Alpha-Alb) | 2,064988394 | 0,00322 | 0.0761554511764352 |
| Q9UKV8 | AGO2 | Protein argonaute-2 (Argonaute2) (hAgo2) (EC 3.1.26.n2) (Argonaute RISC catalytic component 2) (Eukaryotic translation initiation factor 2C 2) (eIF-2C 2) (eIF2C 2) (PAZ Piwi domain protein) (PPD) (Protein slicer) | 0,551752691 | 0,00919 | 0.104035630640557 |
| Q9H9G7 | AGO3 | Protein argonaute-3 (Argonaute3) (hAgo3) (EC 3.1.26.n2) (Argonaute RISC catalytic component 3) (Eukaryotic translation initiation factor 2C 3) (eIF-2C 3) (eIF2C 3) | 0,767211597 | 0,00644 | 0.0954651715986055 |
| O00468 | AGRIN | Agrin [Cleaved into: Agrin N-terminal 110 kDa subunit; Agrin C-terminal 110 kDa subunit; Agrin C-terminal 90 kDa fragment (C90); Agrin C-terminal 22 kDa fragment (C22)] | 1,07099125 | 0,00114 | 0.0574149968045633 |
| P14550 | AK1A1 | Alcohol dehydrogenase [NADP(+)] (EC 1.1.1.2) (Aldehyde reductase) (Aldo-keto reductase family 1 member A1) | -0,403940109 | 0,03349 | 0.171926237130089 |
| Q04828 | AK1C1 | Aldo-keto reductase family 1 member C1 (EC 1.1.1.-) (20-alpha-hydroxysteroid dehydrogenase) (20-alpha-HSD) (EC 1.1.1.149) (Chlordecone reductase homolog HAKRC) (Dihydrodiol dehydrogenase 1/2) (DD1/DD2) (High-affinity hepatic bile acid-binding protein) (HBAB) (Indanol dehydrogenase) (EC 1.1.1.112) (Trans-1,2-dihydrobenzene-1,2-diol dehydrogenase) (EC 1.3.1.20) | -0,524066238 | 0,02863 | 0.160816207866737 |
| Q99996 | AKAP9 | A-kinase anchor protein 9 (AKAP-9) (A-kinase anchor protein 350 kDa) (AKAP 350) (hgAKAP 350) (A-kinase anchor protein 450 kDa) (AKAP 450) (AKAP 120-like protein) (Centrosome- and Golgi-localized PKN-associated protein) (CG-NAP) (Protein hyperion) (Protein kinase A-anchoring protein 9) (PRKA9) (Protein yotiao) | 0,607783218 | 0,00762 | 0.101606052982629 |
| P49419 | AL7A1 | Alpha-aminoadipic semialdehyde dehydrogenase (Alpha-AASA dehydrogenase) (EC 1.2.1.31) (Aldehyde dehydrogenase family 7 member A1) (EC 1.2.1.3) (Antiquitin-1) (Betaine aldehyde dehydrogenase) (EC 1.2.1.8) (Delta1-piperideine-6-carboxylate dehydrogenase) (P6c dehydrogenase) | -0,504352305 | 0,01542 | 0.124751718030612 |
| P02768 | ALBU | Serum albumin | 1,986151096 | 0,00381 | 0.0771477016366249 |
| P05091 | ALDH2 | Aldehyde dehydrogenase, mitochondrial (EC 1.2.1.3) (ALDH class 2) (ALDH-E2) (ALDHI) | -0,435549328 | 0,0352 | 0.175403701257271 |
| P15121 | ALDR | Aldose reductase (AR) (EC 1.1.1.21) (Aldehyde reductase) (Aldo-keto reductase family 1 member B1) | -0,439440542 | 0,0396 | 0.183590812765362 |
| P35858 | ALS | Insulin-like growth factor-binding protein complex acid labile subunit (ALS) | 2,181431426 | 5,2E-05 | 0.0300468839677373 |
| P39687 | AN32A | Acidic leucine-rich nuclear phosphoprotein 32 family member A (Acidic nuclear phosphoprotein pp32) (pp32) (Leucine-rich acidic nuclear protein) (LANP) (Mapmodulin) (Potent heat-stable protein phosphatase 2A inhibitor I1PP2A) (Putative HLA-DR-associated protein I) (PHAPI) | 0,519700072 | 0,01236 | 0.114216522917893 |
| Q92688 | AN32B | Acidic leucine-rich nuclear phosphoprotein 32 family member B (Acidic protein rich in leucines) (Putative HLA-DR-associated protein I-2) (PHAPI2) (Silver-stainable protein SSP29) | 0,638784731 | 0,00764 | 0.101606052982629 |
| Q9P2R3 | ANFY1 | Rabankyrin-5 (Rank-5) (Ankyrin repeat and FYVE domain-containing protein 1) (Ankyrin repeats hooked to a zinc finger motif) | 0,511731736 | 0,03228 | 0.168698420035416 |
| Q9UKU9 | ANGL2 | Angiopoietin-related protein 2 (Angiopoietin-like protein 2) | 1,151143171 | 0,00874 | 0.103882752534462 |
| P16157 | ANK1 | Ankyrin-1 (ANK-1) (Ankyrin-R) (Erythrocyte ankyrin) | 2,323587612 | 0,0001 | 0.0300468839677373 |
| Q9GZV1 | ANKR2 | Ankyrin repeat domain-containing protein 2 (Skeletal muscle ankyrin repeat protein) (hArpp) | -0,784033618 | 0,018 | 0.132658871887263 |
| Q4KMQ2 | ANO6 | Anoctamin-6 (Small-conductance calcium-activated nonselective cation channel) (SCAN channel) (Transmembrane protein 16F) | 0,609566965 | 0,0078 | 0.102154997647559 |
| O75179 | ANR17 | Ankyrin repeat domain-containing protein 17 (Gene trap ankyrin repeat protein) (Serologically defined breast cancer antigen NY-BR-16) | 0,561554454 | 0,0304 | 0.163786460869851 |
| P01008 | ANT3 | Antithrombin-III (ATIII) (Serpin C1) | 1,473823009 | 0,00059 | 0.0460405438333303 |
| P50995 | ANX11 | Annexin A11 (56 kDa autoantigen) (Annexin XI) (Annexin-11) (Calcyclin-associated annexin 50) (CAP-50) | -0,407777729 | 0,02875 | 0.160853551860048 |
| P04083 | ANXA1 | Annexin A1 (Annexin I) (Annexin-1) (Calpactin II) (Calpactin-2) (Chromobindin-9) (Lipocortin I) (Phospholipase A2 inhibitory protein) (p35) | 0,584213233 | 0,02566 | 0.152888992161152 |
| P08758 | ANXA5 | Annexin A5 (Anchorin CII) (Annexin V) (Annexin-5) (Calphobindin I) (CBP-I) (Endonexin II) (Lipocortin V) (Placental anticoagulant protein 4) (PP4) (Placental anticoagulant protein I) (PAP-I) (Thromboplastin inhibitor) (Vascular anticoagulant-alpha) (VAC-alpha) | 0,674638453 | 0,00596 | 0.0946658042155592 |
| P21397 | AOFA | Amine oxidase [flavin-containing] A (EC 1.4.3.4) (Monoamine oxidase type A) (MAO-A) | -0,612116676 | 0,00647 | 0.0954651715986055 |
| P27338 | AOFB | Amine oxidase [flavin-containing] B (EC 1.4.3.4) (Monoamine oxidase type B) (MAO-B) | -0,460094634 | 0,03732 | 0.179750367939979 |
| Q92572 | AP3S1 | AP-3 complex subunit sigma-1 (AP-3 complex subunit sigma-3A) (Adaptor-related protein complex 3 subunit sigma-1) (Clathrin-associated/assembly/adaptor protein, small 3) (Sigma-3A-adaptin) (Sigma3A-adaptin) (Sigma-adaptin 3a) | 0,455616968 | 0,02128 | 0.141512996692124 |
| Q9BZZ5 | API5 | Apoptosis inhibitor 5 (API-5) (Antiapoptosis clone 11 protein) (AAC-11) (Cell migration-inducing gene 8 protein) (Fibroblast growth factor 2-interacting factor) (FIF) (Protein XAGL) | 0,490193125 | 0,01695 | 0.130074948166753 |
| P08519 | APOA | Apolipoprotein(a) (Apo(a)) (Lp(a)) (EC 3.4.21.-) | 1,995422714 | 0,00457 | 0.0840666208225303 |
| P02647 | APOA1 | Apolipoprotein A-I (Apo-AI) (ApoA-I) (Apolipoprotein A1) [Cleaved into: Proapolipoprotein A-I (ProapoA-I); Truncated apolipoprotein A-I (Apolipoprotein A-I(1-242))] | 1,926039339 | 0,00075 | 0.0473327203168651 |
| P02652 | APOA2 | Apolipoprotein A-II (Apo-AII) (ApoA-II) (Apolipoprotein A2) [Cleaved into: Proapolipoprotein A-II (ProapoA-II); Truncated apolipoprotein A-II (Apolipoprotein A-II(1-76))] | 1,475898948 | 0,01838 | 0.133184461911317 |
| P06727 | APOA4 | Apolipoprotein A-IV (Apo-AIV) (ApoA-IV) (Apolipoprotein A4) | 0,551390463 | 0,01802 | 0.132658871887263 |
| P04114 | APOB | Apolipoprotein B-100 (Apo B-100) [Cleaved into: Apolipoprotein B-48 (Apo B-48)] | 2,3830244 | 0,0025 | 0.073956300294973 |
| P02656 | APOC3 | Apolipoprotein C-III (Apo-CIII) (ApoC-III) (Apolipoprotein C3) | 1,061955482 | 0,03037 | 0.163786460869851 |
| P02649 | APOE | Apolipoprotein E (Apo-E) | 1,352584116 | 0,00532 | 0.0915630731121834 |
| P02749 | APOH | Beta-2-glycoprotein 1 (APC inhibitor) (Activated protein C-binding protein) (Anticardiolipin cofactor) (Apolipoprotein H) (Apo-H) (Beta-2-glycoprotein I) (B2GPI) (Beta(2)GPI) | 1,129228252 | 0,00257 | 0.073992460491634 |
| O14791 | APOL1 | Apolipoprotein L1 (Apolipoprotein L) (Apo-L) (ApoL) (Apolipoprotein L-I) (ApoL-I) | 2,333364005 | 0,00012 | 0.0300468839677373 |
| P07741 | APT | Adenine phosphoribosyltransferase (APRT) (EC 2.4.2.7) | 0,757806783 | 0,01167 | 0.112094959351594 |
| Q66PJ3 | AR6P4 | ADP-ribosylation factor-like protein 6-interacting protein 4 (ARL-6-interacting protein 4) (Aip-4) (HSP-975) (HSVI-binding protein) (SR-15) (SRp25) (SR-25) (Splicing factor SRrp37) | 1,440562685 | 0,00041 | 0.0460405438333303 |
| O15143 | ARC1B | Actin-related protein 2/3 complex subunit 1B (Arp2/3 complex 41 kDa subunit) (p41-ARC) | 0,819423777 | 0,01071 | 0.109626270622104 |
| P62330 | ARF6 | ADP-ribosylation factor 6 | 0,412972081 | 0,04437 | 0.192303711595896 |
| Q9NWB6 | ARGL1 | Arginine and glutamate-rich protein 1 | 1,218405543 | 0,00133 | 0.0605895871587186 |
| Q92888 | ARHG1 | Rho guanine nucleotide exchange factor 1 (115 kDa guanine nucleotide exchange factor) (p115-RhoGEF) (p115RhoGEF) (Sub1.5) | 0,714613069 | 0,01479 | 0.12260574009143 |
| Q92974 | ARHG2 | Rho guanine nucleotide exchange factor 2 (Guanine nucleotide exchange factor H1) (GEF-H1) (Microtubule-regulated Rho-GEF) (Proliferating cell nucleolar antigen p40) | 0,560019145 | 0,02071 | 0.139808301680561 |
| Q96PE2 | ARHGH | Rho guanine nucleotide exchange factor 17 (164 kDa Rho-specific guanine-nucleotide exchange factor) (p164-RhoGEF) (p164RhoGEF) (Tumor endothelial marker 4) | 0,398049539 | 0,03016 | 0.16369295498359 |
| Q8NDY3 | ARHL1 | [Protein ADP-ribosylarginine] hydrolase-like protein 1 (EC 3.2.-.-) (ADP-ribosylhydrolase 2) | -0,520848867 | 0,01326 | 0.11792703450348 |
| O15511 | ARPC5 | Actin-related protein 2/3 complex subunit 5 (Arp2/3 complex 16 kDa subunit) (p16-ARC) | 0,67611661 | 0,01143 | 0.111960961329472 |
| Q9ULZ3 | ASC | Apoptosis-associated speck-like protein containing a CARD (hASC) (Caspase recruitment domain-containing protein 5) (PYD and CARD domain-containing protein) (Target of methylation-induced silencing 1) | 0,775472111 | 0,00483 | 0.0864075069907863 |
| Q9BXN1 | ASPN | Asporin (Periodontal ligament-associated protein 1) (PLAP-1) | 1,42117732 | 0,00439 | 0.0823103193899197 |
| Q8IUZ5 | AT2L2 | 5-phosphohydroxy-L-lysine phospho-lyase (EC 4.2.3.134) (Alanine--glyoxylate aminotransferase 2-like 2) | -0,51653812 | 0,01168 | 0.112094959351594 |
| O95260 | ATE1 | Arginyl-tRNA--protein transferase 1 (Arginyltransferase 1) (R-transferase 1) (EC 2.3.2.8) (Arginine-tRNA--protein transferase 1) | -0,477559768 | 0,04283 | 0.188876486048564 |
| O75964 | ATP5L | ATP synthase subunit g, mitochondrial (ATPase subunit g) | -1,171150653 | 0,0162 | 0.127878060348469 |
| O75110 | ATP9A | Probable phospholipid-transporting ATPase IIA (EC 3.6.3.1) (ATPase class II type 9A) | 0,823107532 | 0,00266 | 0.0755562066413184 |
| P36542 | ATPG | ATP synthase subunit gamma, mitochondrial (ATP synthase F1 subunit gamma) (F-ATPase gamma subunit) | -0,500631595 | 0,01919 | 0.135774205040182 |
| P48047 | ATPO | ATP synthase subunit O, mitochondrial (Oligomycin sensitivity conferral protein) (OSCP) | -0,598385385 | 0,00629 | 0.0954651715986055 |
| O75882 | ATRN | Attractin (DPPT-L) (Mahogany homolog) | 0,623956827 | 0,00575 | 0.0945193206940691 |
| O75173 | ATS4 | A disintegrin and metalloproteinase with thrombospondin motifs 4 (ADAM-TS 4) (ADAM-TS4) (ADAMTS-4) (EC 3.4.24.82) (ADMP-1) (Aggrecanase-1) | 1,329822295 | 0,00146 | 0.0658126006797819 |
| P48751 | B3A3 | Anion exchange protein 3 (AE 3) (Anion exchanger 3) (CAE3/BAE3) (Cardiac/brain band 3-like protein) (Neuronal band 3-like protein) (Solute carrier family 4 member 3) | -0,532189483 | 0,00998 | 0.109336835224251 |
| P02730 | B3AT | Band 3 anion transport protein (Anion exchange protein 1) (AE 1) (Anion exchanger 1) (Solute carrier family 4 member 1) (CD antigen CD233) | 2,668036047 | 0,00051 | 0.0460405438333303 |
| Q16611 | BAK | Bcl-2 homologous antagonist/killer (Apoptosis regulator BAK) (Bcl-2-like protein 7) (Bcl2-L-7) | -0,573792936 | 0,00729 | 0.100728702589889 |
| Q9BYV7 | BCDO2 | Beta,beta-carotene 9',10'-oxygenase (EC 1.13.11.71) (B-diox-II) (Beta-carotene dioxygenase 2) | -0,779061172 | 0,03942 | 0.183590812765362 |
| Q9NYF8 | BCLF1 | Bcl-2-associated transcription factor 1 (Btf) (BCLAF1 and THRAP3 family member 1) | 0,423204047 | 0,03979 | 0.183630278609946 |
| Q15582 | BGH3 | Transforming growth factor-beta-induced protein ig-h3 (Beta ig-h3) (Kerato-epithelin) (RGD-containing collagen-associated protein) (RGD-CAP) | 1,333299802 | 0,02869 | 0.160816207866737 |
| P08236 | BGLR | Beta-glucuronidase (EC 3.2.1.31) (Beta-G1) | 0,540401419 | 0,02701 | 0.157616920811605 |
| Q8TD16 | BICD2 | Protein bicaudal D homolog 2 (Bic-D 2) | 0,493152448 | 0,02152 | 0.141824722558827 |
| P55957 | BID | BH3-interacting domain death agonist (p22 BID) (BID) [Cleaved into: BH3-interacting domain death agonist p15 (p15 BID); BH3-interacting domain death agonist p13 (p13 BID); BH3-interacting domain death agonist p11 (p11 BID)] | 0,376868974 | 0,04132 | 0.186041493203554 |
| Q9UBW5 | BIN2 | Bridging integrator 2 (Breast cancer-associated protein 1) | 0,595226666 | 0,04011 | 0.18383048391046 |
| P30043 | BLVRB | Flavin reductase (NADPH) (FR) (EC 1.5.1.30) (Biliverdin reductase B) (BVR-B) (EC 1.3.1.24) (Biliverdin-IX beta-reductase) (Green heme-binding protein) (GHBP) (NADPH-dependent diaphorase) (NADPH-flavin reductase) (FLR) | 0,822231897 | 0,00468 | 0.085284948858929 |
| O95861 | BPNT1 | 3'(2'),5'-bisphosphate nucleotidase 1 (EC 3.1.3.7) (Bisphosphate 3'-nucleotidase 1) (PAP-inositol 1,4-phosphatase) (PIP) | -0,44456082 | 0,02441 | 0.149305815497977 |
| O60885 | BRD4 | Bromodomain-containing protein 4 (Protein HUNK1) | 0,789282286 | 0,00194 | 0.0703852100453896 |
| O43684 | BUB3 | Mitotic checkpoint protein BUB3 | 0,595293436 | 0,00805 | 0.102154997647559 |
| P41223 | BUD31 | Protein BUD31 homolog (Protein EDG-2) (Protein G10 homolog) | 0,981868502 | 0,00206 | 0.0714822379670151 |
| P02745 | C1QA | Complement C1q subcomponent subunit A | 1,57671222 | 0,0037 | 0.0767928025552317 |
| P02746 | C1QB | Complement C1q subcomponent subunit B | 1,019208911 | 0,03578 | 0.176070195019166 |
| P02747 | C1QC | Complement C1q subcomponent subunit C | 1,503430214 | 0,00616 | 0.0954651715986055 |
| Q9NPY3 | C1QR1 | Complement component C1q receptor (C1q/MBL/SPA receptor) (C1qR) (C1qR(p)) (C1qRp) (CDw93) (Complement component 1 q subcomponent receptor 1) (Matrix-remodeling-associated protein 4) (CD antigen CD93) | 1,258366358 | 0,0065 | 0.0954651715986055 |
| P00736 | C1R | Complement C1r subcomponent (EC 3.4.21.41) (Complement component 1 subcomponent r) [Cleaved into: Complement C1r subcomponent heavy chain; Complement C1r subcomponent light chain] | 1,204390867 | 0,02168 | 0.142413043392443 |
| P09871 | C1S | Complement C1s subcomponent (EC 3.4.21.42) (C1 esterase) (Complement component 1 subcomponent s) [Cleaved into: Complement C1s subcomponent heavy chain; Complement C1s subcomponent light chain] | 1,25827434 | 0,00221 | 0.0729440600880325 |
| Q86YS7 | C2CD5 | C2 domain-containing protein 5 (C2 domain-containing phosphoprotein of 138 kDa) | 0,581000004 | 0,007 | 0.0985319097118505 |
| P04003 | C4BPA | C4b-binding protein alpha chain (C4bp) (Proline-rich protein) (PRP) | 2,000800211 | 0,00085 | 0.0495496681815962 |
| Q99643 | C560 | Succinate dehydrogenase cytochrome b560 subunit, mitochondrial (Integral membrane protein CII-3) (QPs-1) (QPs1) (Succinate dehydrogenase complex subunit C) (Succinate-ubiquinone oxidoreductase cytochrome B large subunit) (CYBL) | -0,843503945 | 0,04528 | 0.194611484643512 |
| Q9P1Z2 | CACO1 | Calcium-binding and coiled-coil domain-containing protein 1 (Calphoglin) (Coiled-coil coactivator protein) (Sarcoma antigen NY-SAR-3) | -0,522378705 | 0,02544 | 0.15245012576966 |
| Q13137 | CACO2 | Calcium-binding and coiled-coil domain-containing protein 2 (Antigen nuclear dot 52 kDa protein) (Nuclear domain 10 protein NDP52) (Nuclear domain 10 protein 52) (Nuclear dot protein 52) | -0,558001557 | 0,02849 | 0.160816207866737 |
| P33151 | CADH5 | Cadherin-5 (7B4 antigen) (Vascular endothelial cadherin) (VE-cadherin) (CD antigen CD144) | 0,38473113 | 0,04261 | 0.188876486048564 |
| P00915 | CAH1 | Carbonic anhydrase 1 (EC 4.2.1.1) (Carbonate dehydratase I) (Carbonic anhydrase B) (CAB) (Carbonic anhydrase I) (CA-I) | 2,930397788 | 7,5E-05 | 0.0300468839677373 |
| P00918 | CAH2 | Carbonic anhydrase 2 (EC 4.2.1.1) (Carbonate dehydratase II) (Carbonic anhydrase C) (CAC) (Carbonic anhydrase II) (CA-II) | 1,66346184 | 0,00035 | 0.0460405438333303 |
| P07451 | CAH3 | Carbonic anhydrase 3 (EC 4.2.1.1) (Carbonate dehydratase III) (Carbonic anhydrase III) (CA-III) | 0,547309781 | 0,03961 | 0.183590812765362 |
| P49913 | CAMP | Cathelicidin antimicrobial peptide (18 kDa cationic antimicrobial protein) (CAP-18) (hCAP-18) [Cleaved into: Antibacterial peptide FALL-39 (FALL-39 peptide antibiotic); Antibacterial peptide LL-37] | 2,864629741 | 0,00091 | 0.0511783336599741 |
| Q01518 | CAP1 | Adenylyl cyclase-associated protein 1 (CAP 1) | 0,527387187 | 0,01412 | 0.120558249672262 |
| P40123 | CAP2 | Adenylyl cyclase-associated protein 2 (CAP 2) | -0,377385223 | 0,03932 | 0.183376978972405 |
| P20160 | CAP7 | Azurocidin (Cationic antimicrobial protein CAP37) (Heparin-binding protein) (HBP) (hHBP) | 2,9508876 | 0,00168 | 0.0673661906302801 |
| Q96LW7 | CAR19 | Caspase recruitment domain-containing protein 19 (Bcl10-interacting CARD protein) (BinCARD) | 0,491881785 | 0,02909 | 0.161086911482475 |
| P31944 | CASPE | Caspase-14 (CASP-14) (EC 3.4.22.-) [Cleaved into: Caspase-14 subunit p17, mature form; Caspase-14 subunit p10, mature form; Caspase-14 subunit p20, intermediate form; Caspase-14 subunit p8, intermediate form] | 1,076475168 | 0,00286 | 0.0759058199506327 |
| P04040 | CATA | Catalase (EC 1.11.1.6) | 1,185731073 | 0,0003 | 0.0460405438333303 |
| P08311 | CATG | Cathepsin G (CG) (EC 3.4.21.20) | 2,111947163 | 0,00094 | 0.0512931662027789 |
| P51636 | CAV2 | Caveolin-2 | 0,482360111 | 0,03374 | 0.172516669078315 |
| P52907 | CAZA1 | F-actin-capping protein subunit alpha-1 (CapZ alpha-1) | 0,502818094 | 0,02242 | 0.14536422715807 |
| P08185 | CBG | Corticosteroid-binding globulin (CBG) (Serpin A6) (Transcortin) | 1,870892846 | 0,00047 | 0.0460405438333303 |
| P15088 | CBPA3 | Mast cell carboxypeptidase A (MC-CPA) (EC 3.4.17.1) (Carboxypeptidase A3) | -0,560958017 | 0,01849 | 0.133184461911317 |
| Q96IY4 | CBPB2 | Carboxypeptidase B2 (EC 3.4.17.20) (Carboxypeptidase U) (CPU) (Plasma carboxypeptidase B) (pCPB) (Thrombin-activable fibrinolysis inhibitor) (TAFI) | 1,006698146 | 0,03672 | 0.177966828074799 |
| Q9Y646 | CBPQ | Carboxypeptidase Q (EC 3.4.17.-) (Lysosomal dipeptidase) (Plasma glutamate carboxypeptidase) | 0,454681111 | 0,03512 | 0.175225461224328 |
| Q9GZT6 | CC90B | Coiled-coil domain-containing protein 90B, mitochondrial | -0,363602416 | 0,04119 | 0.185884549229643 |
| Q8IX12 | CCAR1 | Cell division cycle and apoptosis regulator protein 1 (Cell cycle and apoptosis regulatory protein 1) (CARP-1) (Death inducer with SAP domain) | 0,456503187 | 0,01921 | 0.135774205040182 |
| Q8N163 | CCAR2 | Cell cycle and apoptosis regulator protein 2 (Cell division cycle and apoptosis regulator protein 2) (DBIRD complex subunit KIAA1967) (Deleted in breast cancer gene 1 protein) (DBC-1) (DBC.1) (NET35) (p30 DBC) | 0,487283351 | 0,02205 | 0.144376805185818 |
| Q86WR0 | CCD25 | Coiled-coil domain-containing protein 25 | -0,395045179 | 0,0307 | 0.164699828413934 |
| P48509 | CD151 | CD151 antigen (GP27) (Membrane glycoprotein SFA-1) (Platelet-endothelial tetraspan antigen 3) (PETA-3) (Tetraspanin-24) (Tspan-24) (CD antigen CD151) | 1,008375321 | 0,0246 | 0.149799934784929 |
| P16070 | CD44 | CD44 antigen (CDw44) (Epican) (Extracellular matrix receptor III) (ECMR-III) (GP90 lymphocyte homing/adhesion receptor) (HUTCH-I) (Heparan sulfate proteoglycan) (Hermes antigen) (Hyaluronate receptor) (Phagocytic glycoprotein 1) (PGP-1) (Phagocytic glycoprotein I) (PGP-I) (CD antigen CD44) | 0,960047352 | 0,01695 | 0.130074948166753 |
| P30260 | CDC27 | Cell division cycle protein 27 homolog (Anaphase-promoting complex subunit 3) (APC3) (CDC27 homolog) (CDC27Hs) (H-NUC) | 0,357507365 | 0,0485 | 0.201503542294068 |
| Q16543 | CDC37 | Hsp90 co-chaperone Cdc37 (Hsp90 chaperone protein kinase-targeting subunit) (p50Cdc37) [Cleaved into: Hsp90 co-chaperone Cdc37, N-terminally processed] | -0,403237122 | 0,03074 | 0.164699828413934 |
| Q7Z7K6 | CENPV | Centromere protein V (CENP-V) (Nuclear protein p30) (Proline-rich protein 6) | 0,696451448 | 0,0341 | 0.173091625713912 |
| P00450 | CERU | Ceruloplasmin (EC 1.16.3.1) (Ferroxidase) | 1,741298598 | 0,01068 | 0.109626270622104 |
| P00751 | CFAB | Complement factor B (EC 3.4.21.47) (C3/C5 convertase) (Glycine-rich beta glycoprotein) (GBG) (PBF2) (Properdin factor B) [Cleaved into: Complement factor B Ba fragment; Complement factor B Bb fragment] | 1,077355173 | 0,01975 | 0.137370395567352 |
| P08603 | CFAH | Complement factor H (H factor 1) | 1,807106626 | 0,00025 | 0.0456520678124653 |
| P05156 | CFAI | Complement factor I (EC 3.4.21.45) (C3B/C4B inactivator) [Cleaved into: Complement factor I heavy chain; Complement factor I light chain] | 1,639908162 | 0,0004 | 0.0460405438333303 |
| Q14839 | CHD4 | Chromodomain-helicase-DNA-binding protein 4 (CHD-4) (EC 3.6.4.12) (ATP-dependent helicase CHD4) (Mi-2 autoantigen 218 kDa protein) (Mi2-beta) | 0,432543831 | 0,037 | 0.178718378708741 |
| Q9Y259 | CHKB | Choline/ethanolamine kinase (Choline kinase beta) (CK) (CKB) (EC 2.7.1.32) (Choline kinase-like protein) (Ethanolamine kinase) (EK) (EC 2.7.1.82) (Ethanolamine kinase beta) (EKB) (choline/ethanolamine kinase beta) (CKEKB) | 0,587963657 | 0,01814 | 0.132794115698057 |
| Q7LBR1 | CHM1B | Charged multivesicular body protein 1b (CHMP1.5) (Chromatin-modifying protein 1b) (CHMP1b) (Vacuolar protein sorting-associated protein 46-2) (Vps46-2) (hVps46-2) | 0,419606426 | 0,02678 | 0.15719965481106 |
| Q99653 | CHP1 | Calcineurin B homologous protein 1 (Calcineurin B-like protein) (Calcium-binding protein CHP) (Calcium-binding protein p22) (EF-hand calcium-binding domain-containing protein p22) | 0,398847865 | 0,02933 | 0.161086911482475 |
| Q9Y2V2 | CHSP1 | Calcium-regulated heat-stable protein 1 (Calcium-regulated heat-stable protein of 24 kDa) (CRHSP-24) | 0,906081601 | 0,00213 | 0.0725913486972866 |
| O75339 | CILP1 | Cartilage intermediate layer protein 1 (CILP-1) (Cartilage intermediate-layer protein) [Cleaved into: Cartilage intermediate layer protein 1 C1; Cartilage intermediate layer protein 1 C2] | 1,385650505 | 0,02103 | 0.140961498869228 |
| Q9NZ45 | CISD1 | CDGSH iron-sulfur domain-containing protein 1 (MitoNEET) | -0,705489373 | 0,02983 | 0.162736953292786 |
| Q86T13 | CLC14 | C-type lectin domain family 14 member A (Epidermal growth factor receptor 5) (EGFR-5) | 0,646834686 | 0,02285 | 0.146428755002108 |
| P09496 | CLCA | Clathrin light chain A (Lca) | 0,813557457 | 0,00823 | 0.102945529755352 |
| P51797 | CLCN6 | Chloride transport protein 6 (Chloride channel protein 6) (ClC-6) | 1,909808478 | 6,9E-05 | 0.0300468839677373 |
| P30622 | CLIP1 | CAP-Gly domain-containing linker protein 1 (Cytoplasmic linker protein 1) (Cytoplasmic linker protein 170 alpha-2) (CLIP-170) (Reed-Sternberg intermediate filament-associated protein) (Restin) | -0,378270517 | 0,04692 | 0.198769565715963 |
| P10909 | CLUS | Clusterin (Aging-associated gene 4 protein) (Apolipoprotein J) (Apo-J) (Complement cytolysis inhibitor) (CLI) (Complement-associated protein SP-40,40) (Ku70-binding protein 1) (NA1/NA2) (Testosterone-repressed prostate message 2) (TRPM-2) [Cleaved into: Clusterin beta chain (ApoJalpha) (Complement cytolysis inhibitor a chain); Clusterin alpha chain (ApoJbeta) (Complement cytolysis inhibitor b chain)] | 1,578711402 | 0,01203 | 0.113024663824624 |
| O75746 | CMC1 | Calcium-binding mitochondrial carrier protein Aralar1 (Mitochondrial aspartate glutamate carrier 1) (Solute carrier family 25 member 12) | -0,35554022 | 0,04299 | 0.188876486048564 |
| A5YKK6 | CNOT1 | CCR4-NOT transcription complex subunit 1 (CCR4-associated factor 1) (Negative regulator of transcription subunit 1 homolog) (NOT1H) (hNOT1) | 0,619065893 | 0,01068 | 0.109626270622104 |
| P02452 | CO1A1 | Collagen alpha-1(I) chain (Alpha-1 type I collagen) | 1,108184399 | 0,00931 | 0.10490368555784 |
| P08123 | CO1A2 | Collagen alpha-2(I) chain (Alpha-2 type I collagen) | 1,152382689 | 0,02275 | 0.146428755002108 |
| P06681 | CO2 | Complement C2 (EC 3.4.21.43) (C3/C5 convertase) [Cleaved into: Complement C2b fragment; Complement C2a fragment] | 1,434719032 | 0,00249 | 0.073956300294973 |
| P01024 | CO3 | Complement C3 (C3 and PZP-like alpha-2-macroglobulin domain-containing protein 1) [Cleaved into: Complement C3 beta chain; C3-beta-c (C3bc); Complement C3 alpha chain; C3a anaphylatoxin; Acylation stimulating protein (ASP) (C3adesArg); Complement C3b alpha' chain; Complement C3c alpha' chain fragment 1; Complement C3dg fragment; Complement C3g fragment; Complement C3d fragment; Complement C3f fragment; Complement C3c alpha' chain fragment 2] | 1,898034812 | 0,00131 | 0.0604884183584463 |
| P02462 | CO4A1 | Collagen alpha-1(IV) chain [Cleaved into: Arresten] | 1,413717011 | 0,00101 | 0.0538418641701199 |
| P0C0L5 | CO4B | Complement C4-B (Basic complement C4) (C3 and PZP-like alpha-2-macroglobulin domain-containing protein 3) [Cleaved into: Complement C4 beta chain; Complement C4-B alpha chain; C4a anaphylatoxin; C4b-B; C4d-B; Complement C4 gamma chain] | 1,803789191 | 0,00315 | 0.0761554511764352 |
| P01031 | CO5 | Complement C5 (C3 and PZP-like alpha-2-macroglobulin domain-containing protein 4) [Cleaved into: Complement C5 beta chain; Complement C5 alpha chain; C5a anaphylatoxin; Complement C5 alpha' chain] | 1,661728941 | 0,00434 | 0.0817441384449513 |
| P20908 | CO5A1 | Collagen alpha-1(V) chain | 1,075283053 | 0,01269 | 0.115380023220833 |
| P05997 | CO5A2 | Collagen alpha-2(V) chain | 1,020476832 | 0,0037 | 0.0767928025552317 |
| P13671 | CO6 | Complement component C6 | 1,517153587 | 0,00043 | 0.0460405438333303 |
| P12109 | CO6A1 | Collagen alpha-1(VI) chain | 1,06782291 | 0,03416 | 0.173091625713912 |
| A6NMZ7 | CO6A6 | Collagen alpha-6(VI) chain | 0,823750811 | 0,0031 | 0.0761554511764352 |
| P10643 | CO7 | Complement component C7 | 1,094308761 | 0,0396 | 0.183590812765362 |
| P07357 | CO8A | Complement component C8 alpha chain (Complement component 8 subunit alpha) | 1,69150398 | 0,00347 | 0.0764488368109507 |
| P27658 | CO8A1 | Collagen alpha-1(VIII) chain (Endothelial collagen) [Cleaved into: Vastatin] | 0,915435372 | 0,00331 | 0.0761554511764352 |
| P07358 | CO8B | Complement component C8 beta chain (Complement component 8 subunit beta) | 1,772948467 | 0,00344 | 0.0764488368109507 |
| P07360 | CO8G | Complement component C8 gamma chain | 2,062975473 | 0,00583 | 0.0945193206940691 |
| P02748 | CO9 | Complement component C9 [Cleaved into: Complement component C9a; Complement component C9b] | 0,837475261 | 0,03954 | 0.183590812765362 |
| Q99715 | COCA1 | Collagen alpha-1(XII) chain | 1,742733321 | 0,03222 | 0.168698420035416 |
| Q05707 | COEA1 | Collagen alpha-1(XIV) chain (Undulin) | 1,378239648 | 0,00611 | 0.0954651715986055 |
| P39059 | COFA1 | Collagen alpha-1(XV) chain [Cleaved into: Restin (Endostatin-XV) (Related to endostatin) (Restin-I); Restin-2 (Restin-II); Restin-3 (Restin-III); Restin-4 (Restin-IV)] | 0,456987986 | 0,03494 | 0.175099419027731 |
| Q9UP83 | COG5 | Conserved oligomeric Golgi complex subunit 5 (COG complex subunit 5) (13S Golgi transport complex 90 kDa subunit) (GTC-90) (Component of oligomeric Golgi complex 5) (Golgi transport complex 1) | 0,445937408 | 0,04789 | 0.200113362152407 |
| Q9GZQ3 | COMD5 | COMM domain-containing protein 5 (Hypertension-related calcium-regulated gene protein) (HCaRG) | 0,449935098 | 0,01965 | 0.137370395567352 |
| Q8NI60 | COQ8A | Atypical kinase COQ8A, mitochondrial (EC 2.7.-.-) (Chaperone activity of bc1 complex-like) (Chaperone-ABC1-like) (Coenzyme Q protein 8A) (aarF domain-containing protein kinase 3) | -0,650646361 | 0,0374 | 0.179750367939979 |
| O75208 | COQ9 | Ubiquinone biosynthesis protein COQ9, mitochondrial | -0,854846665 | 0,01742 | 0.13109533821508 |
| P31146 | COR1A | Coronin-1A (Coronin-like protein A) (Clipin-A) (Coronin-like protein p57) (Tryptophan aspartate-containing coat protein) (TACO) | 1,734944968 | 0,00802 | 0.102154997647559 |
| Q9ULV4 | COR1C | Coronin-1C (Coronin-3) (hCRNN4) | 0,641674075 | 0,0069 | 0.0978049310072672 |
| Q6QEF8 | CORO6 | Coronin-6 (Coronin-like protein E) (Clipin-E) | 0,590613487 | 0,03156 | 0.167048392255575 |
| P57737 | CORO7 | Coronin-7 (Crn7) (70 kDa WD repeat tumor rejection antigen homolog) | 1,058518548 | 0,00513 | 0.0899766492445311 |
| Q14019 | COTL1 | Coactosin-like protein | 0,974536711 | 0,00253 | 0.073956300294973 |
| Q49B96 | COX19 | Cytochrome c oxidase assembly protein COX19 (hCOX19) | -0,52683998 | 0,03217 | 0.168698420035416 |
| O14548 | COX7R | Cytochrome c oxidase subunit 7A-related protein, mitochondrial (COX7a-related protein) (Cytochrome c oxidase subunit VIIa-related protein) (EB1) | -0,648033498 | 0,02787 | 0.160530825210271 |
| P22792 | CPN2 | Carboxypeptidase N subunit 2 (Carboxypeptidase N 83 kDa chain) (Carboxypeptidase N large subunit) (Carboxypeptidase N polypeptide 2) (Carboxypeptidase N regulatory subunit) | 2,101904542 | 0,00049 | 0.0460405438333303 |
| Q8N684 | CPSF7 | Cleavage and polyadenylation specificity factor subunit 7 (Cleavage and polyadenylation specificity factor 59 kDa subunit) (CPSF 59 kDa subunit) (Cleavage factor Im complex 59 kDa subunit) (CFIm59) (Pre-mRNA cleavage factor Im 59 kDa subunit) | 0,486067242 | 0,03781 | 0.180046072726016 |
| P50416 | CPT1A | Carnitine O-palmitoyltransferase 1, liver isoform (CPT1-L) (EC 2.3.1.21) (Carnitine O-palmitoyltransferase I, liver isoform) (CPT I) (CPTI-L) (Carnitine palmitoyltransferase 1A) | 0,5553928 | 0,0089 | 0.103882752534462 |
| Q92523 | CPT1B | Carnitine O-palmitoyltransferase 1, muscle isoform (CPT1-M) (EC 2.3.1.21) (Carnitine O-palmitoyltransferase I, muscle isoform) (CPT I) (CPTI-M) (Carnitine palmitoyltransferase 1B) (Carnitine palmitoyltransferase I-like protein) | -0,51382006 | 0,01299 | 0.11709919100697 |
| P23786 | CPT2 | Carnitine O-palmitoyltransferase 2, mitochondrial (EC 2.3.1.21) (Carnitine palmitoyltransferase II) (CPT II) | -0,715002422 | 0,00762 | 0.101606052982629 |
| P52943 | CRIP2 | Cysteine-rich protein 2 (CRP-2) (Protein ESP1) | -0,369567265 | 0,04936 | 0.204011338764978 |
| P02741 | CRP | C-reactive protein [Cleaved into: C-reactive protein(1-205)] | -1,614376089 | 0,03401 | 0.173091625713912 |
| Q9Y534 | CSDC2 | Cold shock domain-containing protein C2 (RNA-binding protein PIPPin) | -0,956263088 | 0,00448 | 0.0828018336726045 |
| P13611 | CSPG2 | Versican core protein (Chondroitin sulfate proteoglycan core protein 2) (Chondroitin sulfate proteoglycan 2) (Glial hyaluronate-binding protein) (GHAP) (Large fibroblast proteoglycan) (PG-M) | 1,138752014 | 0,03189 | 0.168140913413884 |
| P33240 | CSTF2 | Cleavage stimulation factor subunit 2 (CF-1 64 kDa subunit) (Cleavage stimulation factor 64 kDa subunit) (CSTF 64 kDa subunit) (CstF-64) | 1,061920202 | 0,00056 | 0.0460405438333303 |
| Q9H0L4 | CSTFT | Cleavage stimulation factor subunit 2 tau variant (CF-1 64 kDa subunit tau variant) (Cleavage stimulation factor 64 kDa subunit tau variant) (CSTF 64 kDa subunit tau variant) (TauCstF-64) | 0,744858224 | 0,00268 | 0.0755562066413184 |
| Q8WYA6 | CTBL1 | Beta-catenin-like protein 1 (Nuclear-associated protein) (NAP) (Testis development protein NYD-SP19) | 0,523012349 | 0,02411 | 0.149130557227562 |
| Q13363 | CTBP1 | C-terminal-binding protein 1 (CtBP1) (EC 1.1.1.-) | 0,623270064 | 0,00621 | 0.0954651715986055 |
| P56545 | CTBP2 | C-terminal-binding protein 2 (CtBP2) | 0,665790386 | 0,0048 | 0.0864075069907863 |
| O60716 | CTND1 | Catenin delta-1 (Cadherin-associated Src substrate) (CAS) (p120 catenin) (p120(ctn)) (p120(cas)) | 0,419664688 | 0,04592 | 0.195567594177145 |
| P04839 | CY24B | Cytochrome b-245 heavy chain (EC 1.-.-.-) (CGD91-phox) (Cytochrome b(558) subunit beta) (Cytochrome b558 subunit beta) (Heme-binding membrane glycoprotein gp91phox) (NADPH oxidase 2) (Neutrophil cytochrome b 91 kDa polypeptide) (Superoxide-generating NADPH oxidase heavy chain subunit) (gp91-1) (gp91-phox) (p22 phagocyte B-cytochrome) | 1,371994318 | 0,00058 | 0.0460405438333303 |
| Q9NRG7 | D39U1 | Epimerase family protein SDR39U1 (EC 1.1.1.-) (Short-chain dehydrogenase/reductase family 39U member 1) | -0,592367778 | 0,00749 | 0.101606052982629 |
| Q86Y56 | DAAF5 | Dynein assembly factor 5, axonemal (HEAT repeat-containing protein 2) | 0,541217643 | 0,01626 | 0.128091561070101 |
| Q9Y4D1 | DAAM1 | Disheveled-associated activator of morphogenesis 1 | 0,862509884 | 0,00355 | 0.0767928025552317 |
| Q5VWQ8 | DAB2P | Disabled homolog 2-interacting protein (DAB2 interaction protein) (DAB2-interacting protein) (ASK-interacting protein 1) (AIP-1) (DOC-2/DAB-2 interactive protein) | 0,360156496 | 0,04578 | 0.195289951852626 |
| P08174 | DAF | Complement decay-accelerating factor (CD antigen CD55) | 0,469720155 | 0,02275 | 0.146428755002108 |
| Q8TEB1 | DCA11 | DDB1- and CUL4-associated factor 11 (WD repeat-containing protein 23) | 0,512589697 | 0,03001 | 0.163245932457869 |
| Q58WW2 | DCAF6 | DDB1- and CUL4-associated factor 6 (Androgen receptor complex-associated protein) (ARCAP) (IQ motif and WD repeat-containing protein 1) (Nuclear receptor interaction protein) (NRIP) | 0,469627094 | 0,01673 | 0.129685065944364 |
| Q96C86 | DCPS | m7GpppX diphosphatase (EC 3.6.1.59) (DCS-1) (Decapping scavenger enzyme) (Hint-related 7meGMP-directed hydrolase) (Histidine triad nucleotide-binding protein 5) (Histidine triad protein member 5) (HINT-5) (Scavenger mRNA-decapping enzyme DcpS) | 0,515388415 | 0,01748 | 0.13109533821508 |
| O95865 | DDAH2 | N(G),N(G)-dimethylarginine dimethylaminohydrolase 2 (DDAH-2) (Dimethylarginine dimethylaminohydrolase 2) (EC 3.5.3.18) (DDAHII) (Dimethylargininase-2) (Protein G6a) (S-phase protein) | 0,491888056 | 0,02614 | 0.154637297234182 |
| Q5TDH0 | DDI2 | Protein DDI1 homolog 2 (EC 3.4.23.-) | 1,319217539 | 0,00345 | 0.0764488368109507 |
| Q92841 | DDX17 | Probable ATP-dependent RNA helicase DDX17 (EC 3.6.4.13) (DEAD box protein 17) (DEAD box protein p72) (DEAD box protein p82) (RNA-dependent helicase p72) | 0,569371196 | 0,01768 | 0.131931842633018 |
| Q9NR30 | DDX21 | Nucleolar RNA helicase 2 (EC 3.6.4.13) (DEAD box protein 21) (Gu-alpha) (Nucleolar RNA helicase Gu) (Nucleolar RNA helicase II) (RH II/Gu) | 0,68672972 | 0,00898 | 0.103994216063343 |
| Q9BUQ8 | DDX23 | Probable ATP-dependent RNA helicase DDX23 (EC 3.6.4.13) (100 kDa U5 snRNP-specific protein) (DEAD box protein 23) (PRP28 homolog) (U5-100kD) | 0,659413981 | 0,00567 | 0.0945193206940691 |
| Q86XP3 | DDX42 | ATP-dependent RNA helicase DDX42 (EC 3.6.4.13) (DEAD box protein 42) (RNA helicase-like protein) (RHELP) (RNA helicase-related protein) (RNAHP) (SF3b DEAD box protein) (Splicing factor 3B-associated 125 kDa protein) (SF3b125) | 0,611279141 | 0,02922 | 0.161086911482475 |
| Q7L014 | DDX46 | Probable ATP-dependent RNA helicase DDX46 (EC 3.6.4.13) (DEAD box protein 46) (PRP5 homolog) | 0,415591272 | 0,02506 | 0.151159838745206 |
| P59665 | DEF1 | Neutrophil defensin 1 (Defensin, alpha 1) (HNP-1) (HP-1) (HP1) [Cleaved into: HP 1-56; Neutrophil defensin 2 (HNP-2) (HP-2) (HP2)] | 2,849341148 | 0,00236 | 0.073956300294973 |
| Q9HBH1 | DEFM | Peptide deformylase, mitochondrial (EC 3.5.1.88) (Polypeptide deformylase) | 0,65835921 | 0,00861 | 0.103882752534462 |
| P35659 | DEK | Protein DEK | 0,392827136 | 0,0383 | 0.180923549477534 |
| Q9Y315 | DEOC | Deoxyribose-phosphate aldolase (DERA) (EC 4.1.2.4) (2-deoxy-D-ribose 5-phosphate aldolase) (Phosphodeoxyriboaldolase) (Deoxyriboaldolase) | 0,749277141 | 0,00919 | 0.104035630640557 |
| Q07507 | DERM | Dermatopontin (Tyrosine-rich acidic matrix protein) (TRAMP) | 1,051190507 | 0,01971 | 0.137370395567352 |
| Q16854 | DGUOK | Deoxyguanosine kinase, mitochondrial (dGK) (EC 2.7.1.113) | 0,410703977 | 0,04727 | 0.198769565715963 |
| P00367 | DHE3 | Glutamate dehydrogenase 1, mitochondrial (GDH 1) (EC 1.4.1.3) | -0,396147647 | 0,03778 | 0.180046072726016 |
| Q9BTZ2 | DHRS4 | Dehydrogenase/reductase SDR family member 4 (EC 1.1.1.184) (NADPH-dependent carbonyl reductase/NADP-retinol dehydrogenase) (CR) (PHCR) (NADPH-dependent retinol dehydrogenase/reductase) (NRDR) (humNRDR) (Peroxisomal short-chain alcohol dehydrogenase) (PSCD) (SCAD-SRL) (Short chain dehydrogenase/reductase family 25C member 2) (Short-chain dehydrogenase/reductase family member 4) | 0,584478101 | 0,02416 | 0.149199506809439 |
| O43143 | DHX15 | Pre-mRNA-splicing factor ATP-dependent RNA helicase DHX15 (EC 3.6.4.13) (ATP-dependent RNA helicase #46) (DEAH box protein 15) | 0,51279182 | 0,0202 | 0.138876689931427 |
| O60231 | DHX16 | Pre-mRNA-splicing factor ATP-dependent RNA helicase DHX16 (EC 3.6.4.13) (ATP-dependent RNA helicase #3) (DEAH-box protein 16) | 0,571845557 | 0,00761 | 0.101606052982629 |
| Q7L2E3 | DHX30 | Putative ATP-dependent RNA helicase DHX30 (EC 3.6.4.13) (DEAH box protein 30) | 0,377911782 | 0,03561 | 0.176070195019166 |
| Q08211 | DHX9 | ATP-dependent RNA helicase A (EC 3.6.4.13) (DEAH box protein 9) (DExH-box helicase 9) (Leukophysin) (LKP) (Nuclear DNA helicase II) (NDH II) (RNA helicase A) | 0,647468469 | 0,00705 | 0.0987831142033136 |
| Q9UNQ2 | DIM1 | Probable dimethyladenosine transferase (EC 2.1.1.183) (DIM1 dimethyladenosine transferase 1 homolog) (DIM1 dimethyladenosine transferase 1-like) (Probable 18S rRNA (adenine(1779)-N(6)/adenine(1780)-N(6))-dimethyltransferase) (Probable 18S rRNA dimethylase) (Probable S-adenosylmethionine-6-N',N'-adenosyl(rRNA) dimethyltransferase) | 0,760085257 | 0,01714 | 0.130557270141694 |
| Q9P265 | DIP2B | Disco-interacting protein 2 homolog B (DIP2 homolog B) | 0,534148174 | 0,04832 | 0.200968705633505 |
| Q9Y2H0 | DLGP4 | Disks large-associated protein 4 (DAP-4) (PSD-95/SAP90-binding protein 4) (SAP90/PSD-95-associated protein 4) (SAPAP-4) | 0,52593318 | 0,04764 | 0.199844821555941 |
| Q9H3Z4 | DNJC5 | DnaJ homolog subfamily C member 5 (Ceroid-lipofuscinosis neuronal protein 4) (Cysteine string protein) (CSP) | -0,80319003 | 0,00193 | 0.0703852100453896 |
| Q9ULA0 | DNPEP | Aspartyl aminopeptidase (EC 3.4.11.21) | 0,588980981 | 0,00761 | 0.101606052982629 |
| Q14185 | DOCK1 | Dedicator of cytokinesis protein 1 (180 kDa protein downstream of CRK) (DOCK180) | 0,617413889 | 0,01153 | 0.111960961329472 |
| Q96HP0 | DOCK6 | Dedicator of cytokinesis protein 6 | 0,469048327 | 0,025 | 0.151098195447372 |
| Q96N67 | DOCK7 | Dedicator of cytokinesis protein 7 | 0,472683914 | 0,01889 | 0.134978321559308 |
| Q12882 | DPYD | Dihydropyrimidine dehydrogenase [NADP(+)] (DHPDHase) (DPD) (EC 1.3.1.2) (Dihydrothymine dehydrogenase) (Dihydrouracil dehydrogenase) | 0,598091809 | 0,03135 | 0.166379218606509 |
| Q14195 | DPYL3 | Dihydropyrimidinase-related protein 3 (DRP-3) (Collapsin response mediator protein 4) (CRMP-4) (Unc-33-like phosphoprotein 1) (ULIP-1) | 0,79448988 | 0,00398 | 0.0785848374826466 |
| P33316 | DUT | Deoxyuridine 5'-triphosphate nucleotidohydrolase, mitochondrial (dUTPase) (EC 3.6.1.23) (dUTP pyrophosphatase) | 0,598980378 | 0,01029 | 0.109336835224251 |
| Q9Y2J2 | E41L3 | Band 4.1-like protein 3 (4.1B) (Differentially expressed in adenocarcinoma of the lung protein 1) (DAL-1) [Cleaved into: Band 4.1-like protein 3, N-terminally processed] | 0,688008597 | 0,00836 | 0.103093659718687 |
| O75521 | ECI2 | Enoyl-CoA delta isomerase 2, mitochondrial (EC 5.3.3.8) (DRS-1) (Delta(3),delta(2)-enoyl-CoA isomerase) (D3,D2-enoyl-CoA isomerase) (Diazepam-binding inhibitor-related protein 1) (DBI-related protein 1) (Dodecenoyl-CoA isomerase) (Hepatocellular carcinoma-associated antigen 88) (Peroxisomal 3,2-trans-enoyl-CoA isomerase) (pECI) (Renal carcinoma antigen NY-REN-1) | -0,38374961 | 0,04577 | 0.195289951852626 |
| Q16610 | ECM1 | Extracellular matrix protein 1 (Secretory component p85) | 1,459173011 | 0,00307 | 0.0761554511764352 |
| Q05639 | EF1A2 | Elongation factor 1-alpha 2 (EF-1-alpha-2) (Eukaryotic elongation factor 1 A-2) (eEF1A-2) (Statin-S1) | -0,525550884 | 0,01589 | 0.126423982233753 |
| Q9BUP0 | EFHD1 | EF-hand domain-containing protein D1 (EF-hand domain-containing protein 1) (Swiprosin-2) | 0,977813651 | 0,02281 | 0.146428755002108 |
| P43897 | EFTS | Elongation factor Ts, mitochondrial (EF-Ts) (EF-TsMt) | -0,379805548 | 0,03868 | 0.181679496107777 |
| Q63HQ2 | EGFLA | Pikachurin (Agrin-like protein) (EGF-like, fibronectin type-III and laminin G-like domain-containing protein) | 0,58982561 | 0,03132 | 0.166379218606509 |
| Q9UI10 | EI2BD | Translation initiation factor eIF-2B subunit delta (eIF-2B GDP-GTP exchange factor subunit delta) | 0,369129223 | 0,03736 | 0.179750367939979 |
| Q53HC9 | EIPR1 | EARP-interacting protein (Endosome-associated recycling protein-interacting protein) (Tumor-suppressing STF cDNA 1 protein) (Tumor-suppressing subchromosomal transferable fragment candidate gene 1 protein) | 0,452323883 | 0,04399 | 0.19148637136756 |
| Q15717 | ELAV1 | ELAV-like protein 1 (Hu-antigen R) (HuR) | 0,444488786 | 0,02963 | 0.162099394483668 |
| P08246 | ELNE | Neutrophil elastase (EC 3.4.21.37) (Bone marrow serine protease) (Elastase-2) (Human leukocyte elastase) (HLE) (Medullasin) (PMN elastase) | 3,581833115 | 0,0013 | 0.0604884183584463 |
| Q00013 | EM55 | 55 kDa erythrocyte membrane protein (p55) (Membrane protein, palmitoylated 1) | 0,555492803 | 0,02657 | 0.15671727829563 |
| Q32P44 | EMAL3 | Echinoderm microtubule-associated protein-like 3 (EMAP-3) | 0,859481224 | 0,00332 | 0.0761554511764352 |
| Q9HC35 | EMAL4 | Echinoderm microtubule-associated protein-like 4 (EMAP-4) (Restrictedly overexpressed proliferation-associated protein) (Ropp 120) | 0,653749236 | 0,00656 | 0.0955763298171194 |
| Q9Y6C2 | EMIL1 | EMILIN-1 (Elastin microfibril interface-located protein 1) (Elastin microfibril interfacer 1) | 1,32395064 | 0,02067 | 0.139808301680561 |
| O94919 | ENDD1 | Endonuclease domain-containing 1 protein (EC 3.1.30.-) | 0,555787546 | 0,02352 | 0.147143576855328 |
| P14625 | ENPL | Endoplasmin (94 kDa glucose-regulated protein) (GRP-94) (Heat shock protein 90 kDa beta member 1) (Tumor rejection antigen 1) (gp96 homolog) | 0,498558566 | 0,02236 | 0.145159783023037 |
| Q9Y6X5 | ENPP4 | Bis(5'-adenosyl)-triphosphatase ENPP4 (EC 3.6.1.29) (AP3A hydrolase) (AP3Aase) (Ectonucleotide pyrophosphatase/phosphodiesterase family member 4) (E-NPP 4) (NPP-4) | -0,594187812 | 0,01838 | 0.133184461911317 |
| Q5NDL2 | EOGT | EGF domain-specific O-linked N-acetylglucosamine transferase (EC 2.4.1.255) (Extracellular O-linked N-acetylglucosamine transferase) | 0,379580179 | 0,04189 | 0.187981866319486 |
| P11171 | EPB41 | Protein 4.1 (P4.1) (4.1R) (Band 4.1) (EPB4.1) | 0,847950694 | 0,00181 | 0.0697416810103427 |
| P16452 | EPB42 | Erythrocyte membrane protein band 4.2 (Erythrocyte protein 4.2) (P4.2) | 1,429485723 | 0,0004 | 0.0460405438333303 |
| P10768 | ESTD | S-formylglutathione hydrolase (FGH) (EC 3.1.2.12) (Esterase D) (Methylumbelliferyl-acetate deacetylase) (EC 3.1.1.56) | -0,650101376 | 0,01361 | 0.11913827875583 |
| Q9BSJ8 | ESYT1 | Extended synaptotagmin-1 (E-Syt1) (Membrane-bound C2 domain-containing protein) | 0,450942228 | 0,02514 | 0.151211529477276 |
| A0FGR8 | ESYT2 | Extended synaptotagmin-2 (E-Syt2) (Chr2Syt) | 0,458623129 | 0,03757 | 0.179967517866609 |
| Q6IPR1 | ETFR1 | Electron transfer flavoprotein regulatory factor 1 (LYR motif-containing protein 5) | -0,596572795 | 0,01023 | 0.109336835224251 |
| Q01844 | EWS | RNA-binding protein EWS (EWS oncogene) (Ewing sarcoma breakpoint region 1 protein) | 0,452700943 | 0,02058 | 0.139808301680561 |
| Q15024 | EXOS7 | Exosome complex component RRP42 (Exosome component 7) (Ribosomal RNA-processing protein 42) (p8) | 0,457070062 | 0,02439 | 0.149305815497977 |
| Q06265 | EXOS9 | Exosome complex component RRP45 (Autoantigen PM/Scl 1) (Exosome component 9) (P75 polymyositis-scleroderma overlap syndrome-associated autoantigen) (Polymyositis/scleroderma autoantigen 1) (Polymyositis/scleroderma autoantigen 75 kDa) (PM/Scl-75) | 0,653020153 | 0,02894 | 0.161086911482475 |
| Q96A26 | F162A | Protein FAM162A (E2-induced gene 5 protein) (Growth and transformation-dependent protein) (HGTD-P) | -0,880839022 | 0,0059 | 0.0945193206940691 |
| Q6UWH4 | F198B | Protein FAM198B (Expressed in nerve and epithelium during development) | 1,169798891 | 0,00483 | 0.0864075069907863 |
| Q96ND0 | F210A | Protein FAM210A | -0,480303358 | 0,03318 | 0.171297602744958 |
| Q9BRX8 | F213A | Redox-regulatory protein FAM213A (Peroxiredoxin-like 2 activated in M-CSF stimulated monocytes) (Protein PAMM) | -0,620196471 | 0,01316 | 0.117794149768834 |
| P00742 | FA10 | Coagulation factor X (EC 3.4.21.6) (Stuart factor) (Stuart-Prower factor) [Cleaved into: Factor X light chain; Factor X heavy chain; Activated factor Xa heavy chain] | 1,667205866 | 0,00218 | 0.0729440600880325 |
| P00748 | FA12 | Coagulation factor XII (EC 3.4.21.38) (Hageman factor) (HAF) [Cleaved into: Coagulation factor XIIa heavy chain; Beta-factor XIIa part 1; Coagulation factor XIIa light chain (Beta-factor XIIa part 2)] | 2,158135927 | 0,0012 | 0.0586177938091161 |
| Q9H0Q0 | FA49A | Protein FAM49A | 0,779697638 | 0,00387 | 0.0773552715354599 |
| Q12805 | FBLN3 | EGF-containing fibulin-like extracellular matrix protein 1 (Extracellular protein S1-5) (Fibrillin-like protein) (Fibulin-3) (FIBL-3) | 1,429557737 | 0,04706 | 0.198769565715963 |
| P35555 | FBN1 | Fibrillin-1 [Cleaved into: Asprosin] | 0,961357789 | 0,00154 | 0.0665699421028979 |
| Q5T0N5 | FBP1L | Formin-binding protein 1-like (Transducer of Cdc42-dependent actin assembly protein 1) (Toca-1) | 0,493205326 | 0,01561 | 0.125740156273994 |
| P22087 | FBRL | rRNA 2'-O-methyltransferase fibrillarin (EC 2.1.1.-) (34 kDa nucleolar scleroderma antigen) (Histone-glutamine methyltransferase) | 0,869110237 | 0,00283 | 0.0759058199506327 |
| Q9Y3I1 | FBX7 | F-box only protein 7 | 1,388750366 | 0,00192 | 0.0703852100453896 |
| Q13630 | FCL | GDP-L-fucose synthase (EC 1.1.1.271) (GDP-4-keto-6-deoxy-D-mannose-3,5-epimerase-4-reductase) (Protein FX) (Red cell NADP(H)-binding protein) (Short-chain dehydrogenase/reductase family 4E member 1) | 0,756187536 | 0,0163 | 0.128147931661901 |
| P02765 | FETUA | Alpha-2-HS-glycoprotein (Alpha-2-Z-globulin) (Ba-alpha-2-glycoprotein) (Fetuin-A) [Cleaved into: Alpha-2-HS-glycoprotein chain A; Alpha-2-HS-glycoprotein chain B] | 2,124801841 | 0,00032 | 0.0460405438333303 |
| P05230 | FGF1 | Fibroblast growth factor 1 (FGF-1) (Acidic fibroblast growth factor) (aFGF) (Endothelial cell growth factor) (ECGF) (Heparin-binding growth factor 1) (HBGF-1) | 0,528990751 | 0,03078 | 0.164699828413934 |
| P49789 | FHIT | Bis(5'-adenosyl)-triphosphatase (EC 3.6.1.29) (AP3A hydrolase) (AP3Aase) (Diadenosine 5',5'''-P1,P3-triphosphate hydrolase) (Dinucleosidetriphosphatase) (Fragile histidine triad protein) | -0,578169423 | 0,02407 | 0.149130557227562 |
| Q13642 | FHL1 | Four and a half LIM domains protein 1 (FHL-1) (Skeletal muscle LIM-protein 1) (SLIM) (SLIM-1) | -0,595497703 | 0,01189 | 0.112411439550033 |
| Q03591 | FHR1 | Complement factor H-related protein 1 (FHR-1) (H factor-like protein 1) (H-factor-like 1) (H36) | 0,830483431 | 0,00409 | 0.0790778462733819 |
| P02671 | FIBA | Fibrinogen alpha chain [Cleaved into: Fibrinopeptide A; Fibrinogen alpha chain] | 1,758208643 | 0,0067 | 0.0963211792898087 |
| P02675 | FIBB | Fibrinogen beta chain [Cleaved into: Fibrinopeptide B; Fibrinogen beta chain] | 1,882771561 | 0,00585 | 0.0945193206940691 |
| P02679 | FIBG | Fibrinogen gamma chain | 1,966887885 | 0,00709 | 0.0988418700944849 |
| P02751 | FINC | Fibronectin (FN) (Cold-insoluble globulin) (CIG) [Cleaved into: Anastellin; Ugl-Y1; Ugl-Y2; Ugl-Y3] | 1,455780445 | 0,02225 | 0.144706839087195 |
| Q02790 | FKBP4 | Peptidyl-prolyl cis-trans isomerase FKBP4 (PPIase FKBP4) (EC 5.2.1.8) (51 kDa FK506-binding protein) (FKBP51) (52 kDa FK506-binding protein) (52 kDa FKBP) (FKBP-52) (59 kDa immunophilin) (p59) (FK506-binding protein 4) (FKBP-4) (FKBP59) (HSP-binding immunophilin) (HBI) (Immunophilin FKBP52) (Rotamase) [Cleaved into: Peptidyl-prolyl cis-trans isomerase FKBP4, N-terminally processed] | 0,43749502 | 0,02064 | 0.139808301680561 |
| Q13451 | FKBP5 | Peptidyl-prolyl cis-trans isomerase FKBP5 (PPIase FKBP5) (EC 5.2.1.8) (51 kDa FK506-binding protein) (51 kDa FKBP) (FKBP-51) (54 kDa progesterone receptor-associated immunophilin) (Androgen-regulated protein 6) (FF1 antigen) (FK506-binding protein 5) (FKBP-5) (FKBP54) (p54) (HSP90-binding immunophilin) (Rotamase) | -0,62991543 | 0,00988 | 0.109011590445672 |
| P21333 | FLNA | Filamin-A (FLN-A) (Actin-binding protein 280) (ABP-280) (Alpha-filamin) (Endothelial actin-binding protein) (Filamin-1) (Non-muscle filamin) | 0,501982685 | 0,0319 | 0.168140913413884 |
| Q8IVF7 | FMNL3 | Formin-like protein 3 (Formin homology 2 domain-containing protein 3) (WW domain-binding protein 3) (WBP-3) | 0,803474533 | 0,00275 | 0.0759058199506327 |
| Q06787 | FMR1 | Synaptic functional regulator FMR1 (Fragile X mental retardation protein 1) (FMRP) (Protein FMR-1) | 0,403795958 | 0,0285 | 0.160816207866737 |
| Q5VW36 | FOCAD | Focadhesin | 0,465395748 | 0,01664 | 0.129654315874851 |
| Q9P2B2 | FPRP | Prostaglandin F2 receptor negative regulator (CD9 partner 1) (CD9P-1) (Glu-Trp-Ile EWI motif-containing protein F) (EWI-F) (Prostaglandin F2-alpha receptor regulatory protein) (Prostaglandin F2-alpha receptor-associated protein) (CD antigen CD315) | 0,488387696 | 0,01411 | 0.120558249672262 |
| Q16595 | FRDA | Frataxin, mitochondrial (EC 1.16.3.1) (Friedreich ataxia protein) (Fxn) [Cleaved into: Frataxin intermediate form (i-FXN); Frataxin(56-210) (m56-FXN); Frataxin(78-210) (d-FXN) (m78-FXN); Frataxin mature form (Frataxin(81-210)) (m81-FXN)] | -0,853665261 | 0,03443 | 0.173957097597315 |
| Q9BZ67 | FRMD8 | FERM domain-containing protein 8 | 0,501280493 | 0,02865 | 0.160816207866737 |
| O94915 | FRYL | Protein furry homolog-like (ALL1-fused gene from chromosome 4p12 protein) | 0,413337211 | 0,043 | 0.188876486048564 |
| Q16658 | FSCN1 | Fascin (55 kDa actin-bundling protein) (Singed-like protein) (p55) | 0,885044963 | 0,00285 | 0.0759058199506327 |
| Q96AE4 | FUBP1 | Far upstream element-binding protein 1 (FBP) (FUSE-binding protein 1) (DNA helicase V) (hDH V) | 0,543785441 | 0,01063 | 0.109626270622104 |
| Q96I24 | FUBP3 | Far upstream element-binding protein 3 (FUSE-binding protein 3) | 0,581064446 | 0,00711 | 0.0988418700944849 |
| Q9BWH2 | FUND2 | FUN14 domain-containing protein 2 (Cervical cancer proto-oncogene 3 protein) (HCC-3) (Hepatitis C virus core-binding protein 6) | -0,868543392 | 0,01086 | 0.110439860103028 |
| Q495W5 | FUT11 | Alpha-(1,3)-fucosyltransferase 11 (EC 2.4.1.-) (Fucosyltransferase XI) (Fuc-TXI) (FucT-XI) (Galactoside 3-L-fucosyltransferase 11) (Fucosyltransferase 11) | 0,617491695 | 0,02849 | 0.160816207866737 |
| Q9BQS8 | FYCO1 | FYVE and coiled-coil domain-containing protein 1 (Zinc finger FYVE domain-containing protein 7) | 0,409815345 | 0,03298 | 0.171077679982414 |
| P11413 | G6PD | Glucose-6-phosphate 1-dehydrogenase (G6PD) (EC 1.1.1.49) | 0,916313543 | 0,0341 | 0.173091625713912 |
| O95479 | G6PE | GDH/6PGL endoplasmic bifunctional protein [Includes: Glucose 1-dehydrogenase (EC 1.1.1.47) (Glucose-6-phosphate dehydrogenase) (EC 1.1.1.363); 6-phosphogluconolactonase (6PGL) (EC 3.1.1.31)] | 0,664399565 | 0,02046 | 0.139808301680561 |
| Q14376 | GALE | UDP-glucose 4-epimerase (EC 5.1.3.2) (Galactowaldenase) (UDP-N-acetylgalactosamine 4-epimerase) (UDP-GalNAc 4-epimerase) (UDP-N-acetylglucosamine 4-epimerase) (UDP-GlcNAc 4-epimerase) (EC 5.1.3.7) (UDP-galactose 4-epimerase) | 0,43734849 | 0,02584 | 0.153565430017805 |
| Q10471 | GALT2 | Polypeptide N-acetylgalactosaminyltransferase 2 (EC 2.4.1.41) (Polypeptide GalNAc transferase 2) (GalNAc-T2) (pp-GaNTase 2) (Protein-UDP acetylgalactosaminyltransferase 2) (UDP-GalNAc:polypeptide N-acetylgalactosaminyltransferase 2) [Cleaved into: Polypeptide N-acetylgalactosaminyltransferase 2 soluble form] | 0,506585217 | 0,02066 | 0.139808301680561 |
| Q14697 | GANAB | Neutral alpha-glucosidase AB (EC 3.2.1.84) (Alpha-glucosidase 2) (Glucosidase II subunit alpha) | 0,515476092 | 0,03227 | 0.168698420035416 |
| Q9H4G4 | GAPR1 | Golgi-associated plant pathogenesis-related protein 1 (GAPR-1) (Golgi-associated PR-1 protein) (Glioma pathogenesis-related protein 2) (GliPR 2) | 0,615828667 | 0,01529 | 0.124447705141253 |
| Q9H0R6 | GATA | Glutamyl-tRNA(Gln) amidotransferase subunit A, mitochondrial (Glu-AdT subunit A) (EC 6.3.5.7) (Glutaminyl-tRNA synthase-like protein 1) | -0,562839402 | 0,02712 | 0.157837513271819 |
| Q9UBI6 | GBG12 | Guanine nucleotide-binding protein G(I)/G(S)/G(O) subunit gamma-12 | -0,626552295 | 0,04323 | 0.189426544291097 |
| P60520 | GBRL2 | Gamma-aminobutyric acid receptor-associated protein-like 2 (GABA(A) receptor-associated protein-like 2) (Ganglioside expression factor 2) (GEF-2) (General protein transport factor p16) (Golgi-associated ATPase enhancer of 16 kDa) (GATE-16) (MAP1 light chain 3-related protein) | -0,398377606 | 0,04913 | 0.203460874759469 |
| Q9BSJ2 | GCP2 | Gamma-tubulin complex component 2 (GCP-2) (hGCP2) (Gamma-ring complex protein 103 kDa) (h103p) (hGrip103) (Spindle pole body protein Spc97 homolog) (hSpc97) | 0,380900204 | 0,03577 | 0.176070195019166 |
| P23434 | GCSH | Glycine cleavage system H protein, mitochondrial (Lipoic acid-containing protein) | -0,662100085 | 0,04712 | 0.198769565715963 |
| Q02108 | GCYA1 | Guanylate cyclase soluble subunit alpha-1 (GCS-alpha-1) (EC 4.6.1.2) (Guanylate cyclase soluble subunit alpha-3) (GCS-alpha-3) (Soluble guanylate cyclase large subunit) | 0,37874688 | 0,0478 | 0.200071443309294 |
| P52566 | GDIR2 | Rho GDP-dissociation inhibitor 2 (Rho GDI 2) (Ly-GDI) (Rho-GDI beta) | 0,473958737 | 0,01805 | 0.132658871887263 |
| P07093 | GDN | Glia-derived nexin (GDN) (Peptidase inhibitor 7) (PI-7) (Protease nexin 1) (PN-1) (Protease nexin I) (Serpin E2) | 1,568180168 | 0,01075 | 0.109778311681309 |
| Q9NXC2 | GFOD1 | Glucose-fructose oxidoreductase domain-containing protein 1 (EC 1.-.-.-) | 0,501866412 | 0,042 | 0.188028909639907 |
| Q06210 | GFPT1 | Glutamine--fructose-6-phosphate aminotransferase [isomerizing] 1 (EC 2.6.1.16) (D-fructose-6-phosphate amidotransferase 1) (Glutamine:fructose-6-phosphate amidotransferase 1) (GFAT 1) (GFAT1) (Hexosephosphate aminotransferase 1) | 0,606380292 | 0,02376 | 0.147884574150362 |
| Q8WWP7 | GIMA1 | GTPase IMAP family member 1 (Immunity-associated protein 1) (hIMAP1) | 0,35252844 | 0,04492 | 0.193722382532051 |
| Q9NUV9 | GIMA4 | GTPase IMAP family member 4 (Immunity-associated nucleotide 1 protein) (IAN-1) (hIAN1) (Immunity-associated protein 4) | 0,854795203 | 0,002 | 0.0713274753495378 |
| Q96F15 | GIMA5 | GTPase IMAP family member 5 (Immunity-associated nucleotide 4-like 1 protein) (Immunity-associated nucleotide 5 protein) (IAN-5) (hIAN5) (Immunity-associated protein 3) | -0,611929284 | 0,04535 | 0.194611484643512 |
| Q7Z3D6 | GLUCM | D-glutamate cyclase, mitochondrial (EC 4.2.1.48) | 1,750779885 | 0,04581 | 0.195289951852626 |
| O60547 | GMDS | GDP-mannose 4,6 dehydratase (EC 4.2.1.47) (GDP-D-mannose dehydratase) (GMD) | 0,760428566 | 0,035 | 0.175099419027731 |
| O60234 | GMFG | Glia maturation factor gamma (GMF-gamma) | 0,920535567 | 0,00589 | 0.0945193206940691 |
| P36915 | GNL1 | Guanine nucleotide-binding protein-like 1 (GTP-binding protein HSR1) | 0,899220613 | 0,00253 | 0.073956300294973 |
| Q8TDQ7 | GNPI2 | Glucosamine-6-phosphate isomerase 2 (EC 3.5.99.6) (Glucosamine-6-phosphate deaminase 2) (GNPDA 2) (GlcN6P deaminase 2) (Glucosamine-6-phosphate isomerase SB52) | -0,928899612 | 0,00087 | 0.0495496681815962 |
| Q08379 | GOGA2 | Golgin subfamily A member 2 (130 kDa cis-Golgi matrix protein) (GM130) (GM130 autoantigen) (Golgin-95) | 0,583947299 | 0,01455 | 0.12221379390711 |
| O14653 | GOSR2 | Golgi SNAP receptor complex member 2 (27 kDa Golgi SNARE protein) (Membrin) | 0,580948632 | 0,01118 | 0.111041531957605 |
| Q8N335 | GPD1L | Glycerol-3-phosphate dehydrogenase 1-like protein (GPD1-L) (EC 1.1.1.8) | -0,421046496 | 0,04817 | 0.200742464336218 |
| P28676 | GRAN | Grancalcin | 1,492960761 | 0,00319 | 0.0761554511764352 |
| Q4V328 | GRAP1 | GRIP1-associated protein 1 (GRASP-1) [Cleaved into: GRASP-1 C-terminal chain (30kDa C-terminus form)] | 0,438468453 | 0,02859 | 0.160816207866737 |
| P21266 | GSTM3 | Glutathione S-transferase Mu 3 (EC 2.5.1.18) (GST class-mu 3) (GSTM3-3) (hGSTM3-3) | -1,106186203 | 0,02894 | 0.161086911482475 |
| P09211 | GSTP1 | Glutathione S-transferase P (EC 2.5.1.18) (GST class-pi) (GSTP1-1) | -0,537743559 | 0,01098 | 0.110439860103028 |
| Q8NBJ5 | GT251 | Procollagen galactosyltransferase 1 (EC 2.4.1.50) (Collagen beta(1-O)galactosyltransferase 1) (ColGalT 1) (Glycosyltransferase 25 family member 1) (Hydroxylysine galactosyltransferase 1) | 0,733961996 | 0,00642 | 0.0954651715986055 |
| P11166 | GTR1 | Solute carrier family 2, facilitated glucose transporter member 1 (Glucose transporter type 1, erythrocyte/brain) (GLUT-1) (HepG2 glucose transporter) | 1,625515607 | 0,01372 | 0.119323066804449 |
| Q8TDB8 | GTR14 | Solute carrier family 2, facilitated glucose transporter member 14 (Glucose transporter type 14) (GLUT-14) | 2,059234518 | 0,00588 | 0.0945193206940691 |
| P14672 | GTR4 | Solute carrier family 2, facilitated glucose transporter member 4 (Glucose transporter type 4, insulin-responsive) (GLUT-4) | 1,175017994 | 0,00321 | 0.0761554511764352 |
| P07305 | H10 | Histone H1.0 (Histone H1') (Histone H1(0)) [Cleaved into: Histone H1.0, N-terminally processed] | 0,67878793 | 0,01095 | 0.110439860103028 |
| P16403 | H12 | Histone H1.2 (Histone H1c) (Histone H1d) (Histone H1s-1) | 0,896389365 | 0,00282 | 0.0759058199506327 |
| P16402 | H13 | Histone H1.3 (Histone H1c) (Histone H1s-2) | 2,211393932 | 0,00045 | 0.0460405438333303 |
| P10412 | H14 | Histone H1.4 (Histone H1b) (Histone H1s-4) | 0,947798061 | 0,00107 | 0.0551770841842723 |
| P16401 | H15 | Histone H1.5 (Histone H1a) (Histone H1b) (Histone H1s-3) | 1,861457473 | 0,00502 | 0.0886056420052009 |
| Q92522 | H1X | Histone H1x | 0,70786299 | 0,02102 | 0.140961498869228 |
| O75367 | H2AY | Core histone macro-H2A.1 (Histone macroH2A1) (mH2A1) (Histone H2A.y) (H2A/y) (Medulloblastoma antigen MU-MB-50.205) | 0,671370825 | 0,02216 | 0.144403200936269 |
| P0C0S5 | H2AZ | Histone H2A.Z (H2A/z) | 0,661053645 | 0,01241 | 0.114216522917893 |
| Q99880 | H2B1L | Histone H2B type 1-L (Histone H2B.c) (H2B/c) | 0,914522528 | 0,00902 | 0.103994216063343 |
| Q8N257 | H2B3B | Histone H2B type 3-B (H2B type 12) | 0,720807484 | 0,0034 | 0.0764488368109507 |
| P62805 | H4 | Histone H4 | 0,758991401 | 0,00251 | 0.073956300294973 |
| O00165 | HAX1 | HCLS1-associated protein X-1 (HS1-associating protein X-1) (HAX-1) (HS1-binding protein 1) (HSP1BP-1) | 0,554741546 | 0,02922 | 0.161086911482475 |
| P69905 | HBA | Hemoglobin subunit alpha (Alpha-globin) (Hemoglobin alpha chain) | 2,815553651 | 0,00011 | 0.0300468839677373 |
| P68871 | HBB | Hemoglobin subunit beta (Beta-globin) (Hemoglobin beta chain) [Cleaved into: LVV-hemorphin-7; Spinorphin] | 2,775275168 | 8,8E-05 | 0.0300468839677373 |
| P02042 | HBD | Hemoglobin subunit delta (Delta-globin) (Hemoglobin delta chain) | 2,719098093 | 0,00011 | 0.0300468839677373 |
| P51610 | HCFC1 | Host cell factor 1 (HCF) (HCF-1) (C1 factor) (CFF) (VCAF) (VP16 accessory protein) [Cleaved into: HCF N-terminal chain 1; HCF N-terminal chain 2; HCF N-terminal chain 3; HCF N-terminal chain 4; HCF N-terminal chain 5; HCF N-terminal chain 6; HCF C-terminal chain 1; HCF C-terminal chain 2; HCF C-terminal chain 3; HCF C-terminal chain 4; HCF C-terminal chain 5; HCF C-terminal chain 6] | 0,3962817 | 0,0346 | 0.174037405274487 |
| Q7Z4H3 | HDDC2 | HD domain-containing protein 2 (Hepatitis C virus NS5A-transactivated protein 2) (HCV NS5A-transactivated protein 2) | 0,5288009 | 0,03787 | 0.180046072726016 |
| Q9BXW7 | HDHD5 | Haloacid dehalogenase-like hydrolase domain-containing 5 (Cat eye syndrome critical region protein 5) | 0,470611985 | 0,02315 | 0.146880219728601 |
| Q7Z4Q2 | HEAT3 | HEAT repeat-containing protein 3 | 0,892240058 | 0,0091 | 0.104035630640557 |
| P02790 | HEMO | Hemopexin (Beta-1B-glycoprotein) | 1,835599907 | 0,00355 | 0.0767928025552317 |
| P05546 | HEP2 | Heparin cofactor 2 (Heparin cofactor II) (HC-II) (Protease inhibitor leuserpin-2) (HLS2) (Serpin D1) | 2,018057337 | 8,9E-05 | 0.0300468839677373 |
| Q9NWT6 | HIF1N | Hypoxia-inducible factor 1-alpha inhibitor (EC 1.14.11.30) (EC 1.14.11.n4) (Factor inhibiting HIF-1) (FIH-1) (Hypoxia-inducible factor asparagine hydroxylase) | 0,426236855 | 0,03764 | 0.180039845845289 |
| P49773 | HINT1 | Histidine triad nucleotide-binding protein 1 (EC 3.-.-.-) (Adenosine 5'-monophosphoramidase) (Protein kinase C inhibitor 1) (Protein kinase C-interacting protein 1) (PKCI-1) | -0,609544655 | 0,00889 | 0.103882752534462 |
| O00291 | HIP1 | Huntingtin-interacting protein 1 (HIP-1) (Huntingtin-interacting protein I) (HIP-I) | 0,44683906 | 0,04298 | 0.188876486048564 |
| P17096 | HMGA1 | High mobility group protein HMG-I/HMG-Y (HMG-I(Y)) (High mobility group AT-hook protein 1) (High mobility group protein A1) (High mobility group protein R) | 0,639390288 | 0,03932 | 0.183376978972405 |
| P09429 | HMGB1 | High mobility group protein B1 (High mobility group protein 1) (HMG-1) | 0,7692113 | 0,02128 | 0.141512996692124 |
| P05114 | HMGN1 | Non-histone chromosomal protein HMG-14 (High mobility group nucleosome-binding domain-containing protein 1) | 1,293793757 | 0,01037 | 0.109336835224251 |
| P09601 | HMOX1 | Heme oxygenase 1 (HO-1) (EC 1.14.14.18) | 0,834051422 | 0,01736 | 0.13109533821508 |
| P55795 | HNRH2 | Heterogeneous nuclear ribonucleoprotein H2 (hnRNP H2) (FTP-3) (Heterogeneous nuclear ribonucleoprotein H') (hnRNP H') [Cleaved into: Heterogeneous nuclear ribonucleoprotein H2, N-terminally processed] | 0,661990608 | 0,0192 | 0.135774205040182 |
| P31942 | HNRH3 | Heterogeneous nuclear ribonucleoprotein H3 (hnRNP H3) (Heterogeneous nuclear ribonucleoprotein 2H9) (hnRNP 2H9) | 0,602107173 | 0,0201 | 0.138808588545449 |
| Q9BUJ2 | HNRL1 | Heterogeneous nuclear ribonucleoprotein U-like protein 1 (Adenovirus early region 1B-associated protein 5) (E1B-55 kDa-associated protein 5) (E1B-AP5) | 0,575259599 | 0,01236 | 0.114216522917893 |
| P07910 | HNRPC | Heterogeneous nuclear ribonucleoproteins C1/C2 (hnRNP C1/C2) | 0,45472259 | 0,02211 | 0.144403200936269 |
| Q14103 | HNRPD | Heterogeneous nuclear ribonucleoprotein D0 (hnRNP D0) (AU-rich element RNA-binding protein 1) | 0,410993144 | 0,03019 | 0.16369295498359 |
| P14866 | HNRPL | Heterogeneous nuclear ribonucleoprotein L (hnRNP L) | 0,5495426 | 0,03576 | 0.176070195019166 |
| P52272 | HNRPM | Heterogeneous nuclear ribonucleoprotein M (hnRNP M) | 0,625406466 | 0,00653 | 0.0955763298171194 |
| Q9NSC5 | HOME3 | Homer protein homolog 3 (Homer-3) | 0,664911452 | 0,01176 | 0.112208078457086 |
| Q96ED9 | HOOK2 | Protein Hook homolog 2 (h-hook2) (hHK2) | 0,605238011 | 0,01965 | 0.137370395567352 |
| Q8IWW8 | HOT | Hydroxyacid-oxoacid transhydrogenase, mitochondrial (HOT) (EC 1.1.99.24) (Alcohol dehydrogenase iron-containing protein 1) (ADHFe1) (Fe-containing alcohol dehydrogenase) | -0,368318745 | 0,04981 | 0.204754344144234 |
| Q5SSJ5 | HP1B3 | Heterochromatin protein 1-binding protein 3 (Protein HP1-BP74) | 0,602059422 | 0,00701 | 0.0985319097118505 |
| P00492 | HPRT | Hypoxanthine-guanine phosphoribosyltransferase (HGPRT) (HGPRTase) (EC 2.4.2.8) | 0,684221992 | 0,00907 | 0.104035630640557 |
| P00738 | HPT | Haptoglobin (Zonulin) [Cleaved into: Haptoglobin alpha chain; Haptoglobin beta chain] | 3,064162088 | 0,00681 | 0.0971872059109455 |
| P04196 | HRG | Histidine-rich glycoprotein (Histidine-proline-rich glycoprotein) (HPRG) | 1,669416989 | 7E-05 | 0.0300468839677373 |
| Q92598 | HS105 | Heat shock protein 105 kDa (Antigen NY-CO-25) (Heat shock 110 kDa protein) | 0,492615932 | 0,01917 | 0.135774205040182 |
| O75506 | HSBP1 | Heat shock factor-binding protein 1 (Nasopharyngeal carcinoma-associated antigen 13) (NPC-A-13) | -0,510644698 | 0,01639 | 0.128553516049245 |
| O14558 | HSPB6 | Heat shock protein beta-6 (HspB6) (Heat shock 20 kDa-like protein p20) | -0,436236203 | 0,04206 | 0.188103353984359 |
| P23083 | HV102 | Immunoglobulin heavy variable 1-2 (Ig heavy chain V-I region ND) (Ig heavy chain V-I region V35) | 1,055026378 | 0,0079 | 0.102154997647559 |
| P01768 | HV330 | Immunoglobulin heavy variable 3-30 (Ig heavy chain V-III region BUR) (Ig heavy chain V-III region CAM) (Ig heavy chain V-III region GA) (Ig heavy chain V-III region NIE) | 1,128756826 | 0,00648 | 0.0954651715986055 |
| P52790 | HXK3 | Hexokinase-3 (EC 2.7.1.1) (Hexokinase type III) (HK III) | 1,306082636 | 0,00649 | 0.0954651715986055 |
| Q9H1B7 | I2BPL | Interferon regulatory factor 2-binding protein-like (Enhanced at puberty protein 1) | 0,88564188 | 0,00525 | 0.0907497112995325 |
| P17936 | IBP3 | Insulin-like growth factor-binding protein 3 (IBP-3) (IGF-binding protein 3) (IGFBP-3) | 2,044109875 | 0,00012 | 0.0300468839677373 |
| Q16270 | IBP7 | Insulin-like growth factor-binding protein 7 (IBP-7) (IGF-binding protein 7) (IGFBP-7) (IGFBP-rP1) (MAC25 protein) (PGI2-stimulating factor) (Prostacyclin-stimulating factor) (Tumor-derived adhesion factor) (TAF) | 0,601852088 | 0,04239 | 0.188743139597737 |
| P05155 | IC1 | Plasma protease C1 inhibitor (C1 Inh) (C1Inh) (C1 esterase inhibitor) (C1-inhibiting factor) (Serpin G1) | 1,535359716 | 0,00125 | 0.0596963516380526 |
| Q14197 | ICT1 | Peptidyl-tRNA hydrolase ICT1, mitochondrial (EC 3.1.1.29) (39S ribosomal protein L58, mitochondrial) (MRP-L58) (Digestion substraction 1) (DS-1) (Immature colon carcinoma transcript 1 protein) (Mitochondrial large ribosomal subunit protein mL62) | 0,516130751 | 0,02066 | 0.139808301680561 |
| Q16666 | IF16 | Gamma-interferon-inducible protein 16 (Ifi-16) (Interferon-inducible myeloid differentiation transcriptional activator) | 0,439451732 | 0,02951 | 0.161662882550704 |
| P38919 | IF4A3 | Eukaryotic initiation factor 4A-III (eIF-4A-III) (eIF4A-III) (EC 3.6.4.13) (ATP-dependent RNA helicase DDX48) (ATP-dependent RNA helicase eIF4A-3) (DEAD box protein 48) (Eukaryotic initiation factor 4A-like NUK-34) (Eukaryotic translation initiation factor 4A isoform 3) (Nuclear matrix protein 265) (NMP 265) (hNMP 265) [Cleaved into: Eukaryotic initiation factor 4A-III, N-terminally processed] | 0,538197399 | 0,03456 | 0.174037405274487 |
| P01876 | IGHA1 | Immunoglobulin heavy constant alpha 1 (Ig alpha-1 chain C region) (Ig alpha-1 chain C region BUR) (Ig alpha-1 chain C region TRO) | 1,381608863 | 0,01851 | 0.133184461911317 |
| P01880 | IGHD | Immunoglobulin heavy constant delta (Ig delta chain C region) (Ig delta chain C region NIG-65) (Ig delta chain C region WAH) | 2,767799692 | 0,0003 | 0.0460405438333303 |
| P01857 | IGHG1 | Immunoglobulin heavy constant gamma 1 (Ig gamma-1 chain C region) (Ig gamma-1 chain C region EU) (Ig gamma-1 chain C region KOL) (Ig gamma-1 chain C region NIE) | 1,564973109 | 0,00425 | 0.0804977214407954 |
| P01859 | IGHG2 | Immunoglobulin heavy constant gamma 2 (Ig gamma-2 chain C region) (Ig gamma-2 chain C region DOT) (Ig gamma-2 chain C region TIL) (Ig gamma-2 chain C region ZIE) | 1,647278145 | 0,00129 | 0.0604884183584463 |
| P01860 | IGHG3 | Immunoglobulin heavy constant gamma 3 (HDC) (Heavy chain disease protein) (Ig gamma-3 chain C region) | 1,125762663 | 0,00192 | 0.0703852100453896 |
| P01871 | IGHM | Immunoglobulin heavy constant mu (Ig mu chain C region) (Ig mu chain C region BOT) (Ig mu chain C region GAL) (Ig mu chain C region OU) | 2,633568164 | 0,00034 | 0.0460405438333303 |
| P01834 | IGKC | Immunoglobulin kappa constant (Ig kappa chain C region) (Ig kappa chain C region AG) (Ig kappa chain C region CUM) (Ig kappa chain C region EU) (Ig kappa chain C region OU) (Ig kappa chain C region ROY) (Ig kappa chain C region TI) | 1,400530792 | 0,0006 | 0.0460405438333303 |
| B9A064 | IGLL5 | Immunoglobulin lambda-like polypeptide 5 (G lambda-1) (Germline immunoglobulin lambda 1) | 1,65670901 | 0,00751 | 0.101606052982629 |
| Q70UQ0 | IKIP | Inhibitor of nuclear factor kappa-B kinase-interacting protein (I kappa-B kinase-interacting protein) (IKBKB-interacting protein) (IKK-interacting protein) | 1,091262254 | 0,00772 | 0.102154997647559 |
| Q12906 | ILF3 | Interleukin enhancer-binding factor 3 (Double-stranded RNA-binding protein 76) (DRBP76) (M-phase phosphoprotein 4) (MPP4) (Nuclear factor associated with dsRNA) (NFAR) (Nuclear factor of activated T-cells 90 kDa) (NF-AT-90) (Translational control protein 80) (TCP80) | 0,424151196 | 0,03135 | 0.166379218606509 |
| Q9H0C8 | ILKAP | Integrin-linked kinase-associated serine/threonine phosphatase 2C (ILKAP) (EC 3.1.3.16) | 0,605944871 | 0,00878 | 0.103882752534462 |
| Q27J81 | INF2 | Inverted formin-2 (HBEBP2-binding protein C) | 0,803579648 | 0,00308 | 0.0761554511764352 |
| Q96PE3 | INP4A | Type I inositol 3,4-bisphosphate 4-phosphatase (EC 3.1.3.66) (Inositol polyphosphate 4-phosphatase type I) | 0,663881867 | 0,01134 | 0.111960961329472 |
| P06213 | INSR | Insulin receptor (IR) (EC 2.7.10.1) (CD antigen CD220) [Cleaved into: Insulin receptor subunit alpha; Insulin receptor subunit beta] | 0,584374734 | 0,01485 | 0.12260574009143 |
| Q9Y2B9 | IPKG | cAMP-dependent protein kinase inhibitor gamma (PKI-gamma) | -0,509484353 | 0,02595 | 0.153725182711907 |
| P05154 | IPSP | Plasma serine protease inhibitor (Acrosomal serine protease inhibitor) (Plasminogen activator inhibitor 3) (PAI-3) (PAI3) (Protein C inhibitor) (PCI) (Serpin A5) | 0,796782218 | 0,0106 | 0.109626270622104 |
| Q6DN90 | IQEC1 | IQ motif and SEC7 domain-containing protein 1 (ADP-ribosylation factors guanine nucleotide-exchange protein 100) (ADP-ribosylation factors guanine nucleotide-exchange protein 2) (Brefeldin-resistant Arf-GEF 2 protein) (BRAG2) | 0,438601064 | 0,02917 | 0.161086911482475 |
| Q13576 | IQGA2 | Ras GTPase-activating-like protein IQGAP2 | 0,883411125 | 0,00116 | 0.057603561428605 |
| O14896 | IRF6 | Interferon regulatory factor 6 (IRF-6) | -0,432422356 | 0,03928 | 0.183376978972405 |
| P11215 | ITAM | Integrin alpha-M (CD11 antigen-like family member B) (CR-3 alpha chain) (Cell surface glycoprotein MAC-1 subunit alpha) (Leukocyte adhesion receptor MO1) (Neutrophil adherence receptor) (CD antigen CD11b) | 1,207870901 | 0,00244 | 0.073956300294973 |
| P05107 | ITB2 | Integrin beta-2 (Cell surface adhesion glycoproteins LFA-1/CR3/p150,95 subunit beta) (Complement receptor C3 subunit beta) (CD antigen CD18) | 1,283852382 | 0,01217 | 0.113600722143195 |
| P19827 | ITIH1 | Inter-alpha-trypsin inhibitor heavy chain H1 (ITI heavy chain H1) (ITI-HC1) (Inter-alpha-inhibitor heavy chain 1) (Inter-alpha-trypsin inhibitor complex component III) (Serum-derived hyaluronan-associated protein) (SHAP) | 1,863575355 | 0,01053 | 0.109544379673903 |
| P19823 | ITIH2 | Inter-alpha-trypsin inhibitor heavy chain H2 (ITI heavy chain H2) (ITI-HC2) (Inter-alpha-inhibitor heavy chain 2) (Inter-alpha-trypsin inhibitor complex component II) (Serum-derived hyaluronan-associated protein) (SHAP) | 1,603031191 | 0,00294 | 0.0761554511764352 |
| Q14624 | ITIH4 | Inter-alpha-trypsin inhibitor heavy chain H4 (ITI heavy chain H4) (ITI-HC4) (Inter-alpha-inhibitor heavy chain 4) (Inter-alpha-trypsin inhibitor family heavy chain-related protein) (IHRP) (Plasma kallikrein sensitive glycoprotein 120) (Gp120) (PK-120) [Cleaved into: 70 kDa inter-alpha-trypsin inhibitor heavy chain H4; 35 kDa inter-alpha-trypsin inhibitor heavy chain H4] | 1,695196289 | 0,00868 | 0.103882752534462 |
| Q86UX2 | ITIH5 | Inter-alpha-trypsin inhibitor heavy chain H5 (ITI heavy chain H5) (ITI-HC5) (Inter-alpha-inhibitor heavy chain 5) | 0,693729519 | 0,03446 | 0.173957097597315 |
| Q14643 | ITPR1 | Inositol 1,4,5-trisphosphate receptor type 1 (IP3 receptor isoform 1) (IP3R 1) (InsP3R1) (Type 1 inositol 1,4,5-trisphosphate receptor) (Type 1 InsP3 receptor) | 0,459426515 | 0,01944 | 0.136861581624076 |
| P23458 | JAK1 | Tyrosine-protein kinase JAK1 (EC 2.7.10.2) (Janus kinase 1) (JAK-1) | -0,472820728 | 0,02768 | 0.160363747693279 |
| Q9P266 | JCAD | Junctional protein associated with coronary artery disease (JCAD) | 1,006739969 | 0,01482 | 0.12260574009143 |
| Q9BR39 | JPH2 | Junctophilin-2 (JP-2) (Junctophilin type 2) | 0,439056582 | 0,0307 | 0.164699828413934 |
| Q9P260 | K1468 | LisH domain and HEAT repeat-containing protein KIAA1468 | 0,608253289 | 0,02786 | 0.160530825210271 |
| P29622 | KAIN | Kallistatin (Kallikrein inhibitor) (Peptidase inhibitor 4) (PI-4) (Serpin A4) | 1,285861888 | 0,00159 | 0.0670897963950065 |
| Q16773 | KAT1 | Kynurenine--oxoglutarate transaminase 1 (EC 2.6.1.7) (Cysteine-S-conjugate beta-lyase) (EC 4.4.1.13) (Glutamine transaminase K) (GTK) (Glutamine--phenylpyruvate transaminase) (EC 2.6.1.64) (Kynurenine aminotransferase 1) (Kynurenine aminotransferase I) (KATI) (Kynurenine--oxoglutarate transaminase I) | -0,659287295 | 0,00793 | 0.102154997647559 |
| O75600 | KBL | 2-amino-3-ketobutyrate coenzyme A ligase, mitochondrial (AKB ligase) (EC 2.3.1.29) (Aminoacetone synthase) (Glycine acetyltransferase) | -1,149435561 | 0,00668 | 0.0963114437053782 |
| P48730 | KC1D | Casein kinase I isoform delta (CKI-delta) (CKId) (EC 2.7.11.1) (Tau-protein kinase CSNK1D) (EC 2.7.11.26) | 0,59414042 | 0,03809 | 0.180517494421774 |
| Q13557 | KCC2D | Calcium/calmodulin-dependent protein kinase type II subunit delta (CaM kinase II subunit delta) (CaMK-II subunit delta) (EC 2.7.11.17) | -0,363338415 | 0,04128 | 0.186041493203554 |
| Q16774 | KGUA | Guanylate kinase (EC 2.7.4.8) (GMP kinase) | 0,688817472 | 0,0075 | 0.101606052982629 |
| O00139 | KIF2A | Kinesin-like protein KIF2A (Kinesin-2) (hK2) | 0,42729273 | 0,03283 | 0.170772626139264 |
| Q9UJP4 | KLH21 | Kelch-like protein 21 | 0,522122568 | 0,01529 | 0.124447705141253 |
| Q9H511 | KLH31 | Kelch-like protein 31 (BTB and kelch domain-containing protein 6) (Kelch repeat and BTB domain-containing protein 1) (Kelch-like protein KLHL) | -0,836856934 | 0,00284 | 0.0759058199506327 |
| O60662 | KLH41 | Kelch-like protein 41 (Kel-like protein 23) (Kelch repeat and BTB domain-containing protein 10) (Kelch-related protein 1) (Sarcosin) | -0,775094308 | 0,01053 | 0.109544379673903 |
| P03952 | KLKB1 | Plasma kallikrein (EC 3.4.21.34) (Fletcher factor) (Kininogenin) (Plasma prekallikrein) (PKK) [Cleaved into: Plasma kallikrein heavy chain; Plasma kallikrein light chain] | 1,535318666 | 8,1E-05 | 0.0300468839677373 |
| P01042 | KNG1 | Kininogen-1 (Alpha-2-thiol proteinase inhibitor) (Fitzgerald factor) (High molecular weight kininogen) (HMWK) (Williams-Fitzgerald-Flaujeac factor) [Cleaved into: Kininogen-1 heavy chain; T-kinin (Ile-Ser-Bradykinin); Bradykinin (Kallidin I); Lysyl-bradykinin (Kallidin II); Kininogen-1 light chain; Low molecular weight growth-promoting factor] | 1,814492499 | 0,00053 | 0.0460405438333303 |
| P46019 | KPB2 | Phosphorylase b kinase regulatory subunit alpha, liver isoform (Phosphorylase kinase alpha L subunit) | 0,571317437 | 0,00824 | 0.102945529755352 |
| O60256 | KPRB | Phosphoribosyl pyrophosphate synthase-associated protein 2 (PRPP synthase-associated protein 2) (41 kDa phosphoribosypyrophosphate synthetase-associated protein) (PAP41) | 0,585054292 | 0,02729 | 0.158571050899145 |
| P23919 | KTHY | Thymidylate kinase (EC 2.7.4.9) (dTMP kinase) | 0,517107838 | 0,01837 | 0.133184461911317 |
| P04433 | KV311 | Immunoglobulin kappa variable 3-11 (Ig kappa chain V-III region VG) | 1,826629157 | 0,01145 | 0.111960961329472 |
| P01611 | KVD12 | Immunoglobulin kappa variable 1D-12 (Ig kappa chain V-I region Wes) | 1,285157005 | 0,00065 | 0.0460405438333303 |
| P83111 | LACTB | Serine beta-lactamase-like protein LACTB, mitochondrial (EC 3.4.-.-) | -0,705901515 | 0,00641 | 0.0954651715986055 |
| O15230 | LAMA5 | Laminin subunit alpha-5 (Laminin-10 subunit alpha) (Laminin-11 subunit alpha) (Laminin-15 subunit alpha) | 0,471531947 | 0,04281 | 0.188876486048564 |
| P13473 | LAMP2 | Lysosome-associated membrane glycoprotein 2 (LAMP-2) (Lysosome-associated membrane protein 2) (CD107 antigen-like family member B) (LGP-96) (CD antigen CD107b) | 0,604472426 | 0,02337 | 0.146880219728601 |
| O43813 | LANC1 | LanC-like protein 1 (40 kDa erythrocyte membrane protein) (p40) | -0,401078361 | 0,02854 | 0.160816207866737 |
| Q6PKG0 | LARP1 | La-related protein 1 (La ribonucleoprotein domain family member 1) | 0,395431429 | 0,04775 | 0.200071443309294 |
| Q14739 | LBR | Lamin-B receptor (Integral nuclear envelope inner membrane protein) (LMN2R) | 0,959314807 | 0,00149 | 0.0658126006797819 |
| O95232 | LC7L3 | Luc7-like protein 3 (Cisplatin resistance-associated-overexpressed protein) (Luc7A) (Okadaic acid-inducible phosphoprotein OA48-18) (cAMP regulatory element-associated protein 1) (CRE-associated protein 1) (CREAP-1) | 0,488292967 | 0,01851 | 0.133184461911317 |
| Q8NC56 | LEMD2 | LEM domain-containing protein 2 (hLEM2) | 0,732680176 | 0,00516 | 0.0899924708376971 |
| O43766 | LIAS | Lipoyl synthase, mitochondrial (EC 2.8.1.8) (Lipoate synthase) (LS) (Lip-syn) (Lipoic acid synthase) | -0,540901985 | 0,02132 | 0.141512996692124 |
| Q9UHB6 | LIMA1 | LIM domain and actin-binding protein 1 (Epithelial protein lost in neoplasm) | 0,680918325 | 0,00356 | 0.0767928025552317 |
| P06858 | LIPL | Lipoprotein lipase (LPL) (EC 3.1.1.34) | -1,009292074 | 0,01415 | 0.120558249672262 |
| P02545 | LMNA | Prelamin-A/C [Cleaved into: Lamin-A/C (70 kDa lamin) (Renal carcinoma antigen NY-REN-32)] | 0,955749287 | 0,00161 | 0.0671419509793915 |
| P20700 | LMNB1 | Lamin-B1 | 0,763876956 | 0,03026 | 0.163712810448877 |
| Q6P5Q4 | LMOD2 | Leiomodin-2 (Cardiac leiomodin) (C-LMOD) (Leiomodin) | -0,575295886 | 0,04316 | 0.189335709503698 |
| Q96JA1 | LRIG1 | Leucine-rich repeats and immunoglobulin-like domains protein 1 (LIG-1) | 0,694913191 | 0,0058 | 0.0945193206940691 |
| Q9BX40 | LS14B | Protein LSM14 homolog B (Protein FAM61B) (RNA-associated protein 55B) (hRAP55B) | -0,404864907 | 0,04254 | 0.188876486048564 |
| Q3MHD2 | LSM12 | Protein LSM12 homolog | 0,513207336 | 0,01375 | 0.119323429379425 |
| P62310 | LSM3 | U6 snRNA-associated Sm-like protein LSm3 | 0,677482033 | 0,00642 | 0.0954651715986055 |
| P33241 | LSP1 | Lymphocyte-specific protein 1 (47 kDa actin-binding protein) (52 kDa phosphoprotein) (pp52) (Lymphocyte-specific antigen WP34) | 1,857970484 | 0,0006 | 0.0460405438333303 |
| Q14766 | LTBP1 | Latent-transforming growth factor beta-binding protein 1 (LTBP-1) (Transforming growth factor beta-1-binding protein 1) (TGF-beta1-BP-1) | 1,071528013 | 0,00911 | 0.104035630640557 |
| Q8N2S1 | LTBP4 | Latent-transforming growth factor beta-binding protein 4 (LTBP-4) | 1,454464868 | 0,04993 | 0.204754344144234 |
| P51884 | LUM | Lumican (Keratan sulfate proteoglycan lumican) (KSPG lumican) | 0,563066279 | 0,01203 | 0.113024663824624 |
| P80748 | LV321 | Immunoglobulin lambda variable 3-21 (Ig lambda chain V-III region LOI) (Ig lambda chain V-V region DEL) (Ig lambda chain V-VII region MOT) | 1,34385677 | 0,00024 | 0.0456520678124653 |
| Q9BS40 | LXN | Latexin (Endogenous carboxypeptidase inhibitor) (ECI) (Protein MUM) (Tissue carboxypeptidase inhibitor) (TCI) | 1,005643912 | 0,01179 | 0.112267836107064 |
| Q9HD34 | LYRM4 | LYR motif-containing protein 4 | -0,611570751 | 0,03467 | 0.174093522433991 |
| P61626 | LYSC | Lysozyme C (EC 3.2.1.17) (1,4-beta-N-acetylmuramidase C) | 1,722528388 | 0,00069 | 0.0461437135818825 |
| O95819 | M4K4 | Mitogen-activated protein kinase kinase kinase kinase 4 (EC 2.7.11.1) (HPK/GCK-like kinase HGK) (MAPK/ERK kinase kinase kinase 4) (MEK kinase kinase 4) (MEKKK 4) (Nck-interacting kinase) | 0,805608145 | 0,00282 | 0.0759058199506327 |
| P23368 | MAOM | NAD-dependent malic enzyme, mitochondrial (NAD-ME) (EC 1.1.1.38) (Malic enzyme 2) | -0,903729797 | 0,04417 | 0.191862273940179 |
| Q16798 | MAON | NADP-dependent malic enzyme, mitochondrial (NADP-ME) (EC 1.1.1.40) (Malic enzyme 3) | -0,535875892 | 0,03543 | 0.175701151598843 |
| Q16644 | MAPK3 | MAP kinase-activated protein kinase 3 (MAPK-activated protein kinase 3) (MAPKAP kinase 3) (MAPKAP-K3) (MAPKAPK-3) (MK-3) (EC 2.7.11.1) (Chromosome 3p kinase) (3pK) | -0,769600434 | 0,01045 | 0.109336835224251 |
| Q15555 | MARE2 | Microtubule-associated protein RP/EB family member 2 (APC-binding protein EB2) (End-binding protein 2) (EB2) | -0,374242598 | 0,0399 | 0.18383048391046 |
| Q7KZI7 | MARK2 | Serine/threonine-protein kinase MARK2 (EC 2.7.11.1) (EC 2.7.11.26) (ELKL motif kinase 1) (EMK-1) (MAP/microtubule affinity-regulating kinase 2) (PAR1 homolog) (PAR1 homolog b) (Par-1b) (Par1b) | 0,565945219 | 0,01271 | 0.115380023220833 |
| P43243 | MATR3 | Matrin-3 | 0,501040342 | 0,01605 | 0.12714580081316 |
| Q7Z434 | MAVS | Mitochondrial antiviral-signaling protein (MAVS) (CARD adapter inducing interferon beta) (Cardif) (Interferon beta promoter stimulator protein 1) (IPS-1) (Putative NF-kappa-B-activating protein 031N) (Virus-induced-signaling adapter) (VISA) | -0,467757964 | 0,02077 | 0.139973423301439 |
| Q96RQ3 | MCCA | Methylcrotonoyl-CoA carboxylase subunit alpha, mitochondrial (MCCase subunit alpha) (EC 6.4.1.4) (3-methylcrotonyl-CoA carboxylase 1) (3-methylcrotonyl-CoA carboxylase biotin-containing subunit) (3-methylcrotonyl-CoA:carbon dioxide ligase subunit alpha) | -0,648992003 | 0,01661 | 0.129654315874851 |
| Q9HCC0 | MCCB | Methylcrotonoyl-CoA carboxylase beta chain, mitochondrial (MCCase subunit beta) (EC 6.4.1.4) (3-methylcrotonyl-CoA carboxylase 2) (3-methylcrotonyl-CoA carboxylase non-biotin-containing subunit) (3-methylcrotonyl-CoA:carbon dioxide ligase subunit beta) | -0,419511693 | 0,04288 | 0.188876486048564 |
| Q9Y316 | MEMO1 | Protein MEMO1 (C21orf19-like protein) (Hepatitis C virus NS5A-transactivated protein 7) (HCV NS5A-transactivated protein 7) (Mediator of ErbB2-driven cell motility 1) (Mediator of cell motility 1) (Memo-1) | 0,716362915 | 0,01322 | 0.117794149768834 |
| Q8N6R0 | MET13 | Methyltransferase-like protein 13 (EC 2.1.1.-) | 0,852880241 | 0,00286 | 0.0759058199506327 |
| Q9H7H0 | MET17 | Methyltransferase-like protein 17, mitochondrial (EC 2.1.1.-) (False p73 target gene protein) (Methyltransferase 11 domain-containing protein 1) (Protein RSM22 homolog, mitochondrial) | 0,499197609 | 0,01262 | 0.115083692479736 |
| Q8IWA4 | MFN1 | Mitofusin-1 (EC 3.6.5.-) (Fzo homolog) (Transmembrane GTPase MFN1) | -0,433858437 | 0,02437 | 0.149305815497977 |
| Q14728 | MFS10 | Major facilitator superfamily domain-containing protein 10 (Tetracycline transporter-like protein) | 0,708383013 | 0,01478 | 0.12260574009143 |
| P26572 | MGAT1 | Alpha-1,3-mannosyl-glycoprotein 2-beta-N-acetylglucosaminyltransferase (EC 2.4.1.101) (N-glycosyl-oligosaccharide-glycoprotein N-acetylglucosaminyltransferase I) (GNT-I) (GlcNAc-T I) | 0,469531693 | 0,01669 | 0.129654315874851 |
| Q8N4Q1 | MIA40 | Mitochondrial intermembrane space import and assembly protein 40 (Coiled-coil-helix-coiled-coil-helix domain-containing protein 4) | -1,019793243 | 0,00323 | 0.0761554511764352 |
| Q6UXV4 | MIC27 | MICOS complex subunit MIC27 (Apolipoprotein O-like) (Protein FAM121A) | -0,731881053 | 0,01042 | 0.109336835224251 |
| P20774 | MIME | Mimecan (Osteoglycin) (Osteoinductive factor) (OIF) | 1,241930108 | 0,0008 | 0.0484403941566856 |
| P14780 | MMP9 | Matrix metalloproteinase-9 (MMP-9) (EC 3.4.24.35) (92 kDa gelatinase) (92 kDa type IV collagenase) (Gelatinase B) (GELB) [Cleaved into: 67 kDa matrix metalloproteinase-9; 82 kDa matrix metalloproteinase-9] | 2,077434767 | 0,00487 | 0.0864075069907863 |
| Q96T76 | MMS19 | MMS19 nucleotide excision repair protein homolog (hMMS19) (MET18 homolog) (MMS19-like protein) | 0,545788007 | 0,02366 | 0.14774479275927 |
| P41218 | MNDA | Myeloid cell nuclear differentiation antigen | 2,091672035 | 0,00113 | 0.0573447930042465 |
| O96007 | MOC2B | Molybdopterin synthase catalytic subunit (EC 2.8.1.12) (MOCO1-B) (Molybdenum cofactor synthesis protein 2 large subunit) (Molybdenum cofactor synthesis protein 2B) (MOCS2B) (Molybdopterin-synthase large subunit) (MPT synthase large subunit) | -0,498783399 | 0,03391 | 0.173091625713912 |
| O15427 | MOT4 | Monocarboxylate transporter 4 (MCT 4) (Solute carrier family 16 member 3) | 1,551765801 | 0,01885 | 0.13488646867884 |
| O95563 | MPC2 | Mitochondrial pyruvate carrier 2 (Brain protein 44) | -1,222321472 | 0,00094 | 0.0512931662027789 |
| P34949 | MPI | Mannose-6-phosphate isomerase (EC 5.3.1.8) (Phosphohexomutase) (Phosphomannose isomerase) (PMI) | -0,642473577 | 0,01116 | 0.111041531957605 |
| Q9UBG0 | MRC2 | C-type mannose receptor 2 (C-type lectin domain family 13 member E) (Endocytic receptor 180) (Macrophage mannose receptor 2) (Urokinase-type plasminogen activator receptor-associated protein) (UPAR-associated protein) (Urokinase receptor-associated protein) (CD antigen CD280) | 1,459630172 | 0,00082 | 0.0485527134016778 |
| P49959 | MRE11 | Double-strand break repair protein MRE11 (Double-strand break repair protein MRE11A) (Meiotic recombination 11 homolog 1) (MRE11 homolog 1) (Meiotic recombination 11 homolog A) (MRE11 homolog A) | 0,556018075 | 0,01989 | 0.137802712336179 |
| Q6IN84 | MRM1 | rRNA methyltransferase 1, mitochondrial (EC 2.1.1.-) (16S rRNA (guanosine(1145)-2'-O)-methyltransferase) (16S rRNA [Gm1145] 2'-O-methyltransferase) | 0,608989825 | 0,0303 | 0.163712810448877 |
| P43246 | MSH2 | DNA mismatch repair protein Msh2 (hMSH2) (MutS protein homolog 2) | 1,072998427 | 0,00414 | 0.0790778462733819 |
| Q13330 | MTA1 | Metastasis-associated protein MTA1 | 0,829588351 | 0,00605 | 0.0953178672429245 |
| O94776 | MTA2 | Metastasis-associated protein MTA2 (Metastasis-associated 1-like 1) (MTA1-L1 protein) (p53 target protein in deacetylase complex) | 0,441366104 | 0,03596 | 0.176070600102387 |
| Q96QG7 | MTMR9 | Myotubularin-related protein 9 | 0,557480259 | 0,0085 | 0.103753456880926 |
| P42285 | MTREX | Exosome RNA helicase MTR4 (EC 3.6.4.13) (ATP-dependent RNA helicase DOB1) (ATP-dependent RNA helicase SKIV2L2) (Superkiller viralicidic activity 2-like 2) (TRAMP-like complex helicase) | 1,835111488 | 0,00183 | 0.0697416810103427 |
| O75648 | MTU1 | Mitochondrial tRNA-specific 2-thiouridylase 1 (EC 2.8.1.14) (MTO2 homolog) | -0,388749617 | 0,04822 | 0.200742464336218 |
| P53602 | MVD1 | Diphosphomevalonate decarboxylase (EC 4.1.1.33) (Mevalonate (diphospho)decarboxylase) (MDDase) (Mevalonate pyrophosphate decarboxylase) | 0,557459722 | 0,02162 | 0.14226454436859 |
| P20591 | MX1 | Interferon-induced GTP-binding protein Mx1 (Interferon-induced protein p78) (IFI-78K) (Interferon-regulated resistance GTP-binding protein MxA) (Myxoma resistance protein 1) (Myxovirus resistance protein 1) [Cleaved into: Interferon-induced GTP-binding protein Mx1, N-terminally processed] | -0,979202695 | 0,0427 | 0.188876486048564 |
| Q96S97 | MYADM | Myeloid-associated differentiation marker (Protein SB135) | 0,805797726 | 0,00327 | 0.0761554511764352 |
| Q99836 | MYD88 | Myeloid differentiation primary response protein MyD88 | 1,056680431 | 0,00332 | 0.0761554511764352 |
| P12882 | MYH1 | Myosin-1 (Myosin heavy chain 1) (Myosin heavy chain 2x) (MyHC-2x) (Myosin heavy chain IIx/d) (MyHC-IIx/d) (Myosin heavy chain, skeletal muscle, adult 1) | -0,757795466 | 0,01678 | 0.129807282147575 |
| P35580 | MYH10 | Myosin-10 (Cellular myosin heavy chain, type B) (Myosin heavy chain 10) (Myosin heavy chain, non-muscle IIb) (Non-muscle myosin heavy chain B) (NMMHC-B) (Non-muscle myosin heavy chain IIb) (NMMHC II-b) (NMMHC-IIB) | 0,556475134 | 0,03415 | 0.173091625713912 |
| Q9Y2K3 | MYH15 | Myosin-15 (Myosin heavy chain 15) | 1,027717279 | 0,0148 | 0.12260574009143 |
| Q9Y623 | MYH4 | Myosin-4 (Myosin heavy chain 2b) (MyHC-2b) (Myosin heavy chain 4) (Myosin heavy chain IIb) (MyHC-IIb) (Myosin heavy chain, skeletal muscle, fetal) | 1,263580842 | 0,00048 | 0.0460405438333303 |
| A7E2Y1 | MYH7B | Myosin-7B (Antigen MLAA-21) (Myosin cardiac muscle beta chain) (Myosin heavy chain 7B, cardiac muscle beta isoform) (Slow A MYH14) | -0,576764162 | 0,01369 | 0.119314186167483 |
| P13535 | MYH8 | Myosin-8 (Myosin heavy chain 8) (Myosin heavy chain, skeletal muscle, perinatal) (MyHC-perinatal) | 0,574867017 | 0,02689 | 0.15719965481106 |
| P60660 | MYL6 | Myosin light polypeptide 6 (17 kDa myosin light chain) (LC17) (Myosin light chain 3) (MLC-3) (Myosin light chain alkali 3) (Myosin light chain A3) (Smooth muscle and nonmuscle myosin light chain alkali 6) | 0,374237524 | 0,04436 | 0.192303711595896 |
| P14649 | MYL6B | Myosin light chain 6B (Myosin light chain 1 slow-twitch muscle A isoform) (MLC1sa) (Smooth muscle and nonmuscle myosin light chain alkali 6B) | 0,550203622 | 0,0255 | 0.15245012576966 |
| O43795 | MYO1B | Unconventional myosin-Ib (MYH-1c) (Myosin I alpha) (MMI-alpha) (MMIa) | 0,568486801 | 0,00965 | 0.106870225730999 |
| Q9UM54 | MYO6 | Unconventional myosin-VI (Unconventional myosin-6) | 0,429120663 | 0,02443 | 0.149305815497977 |
| Q9NZM1 | MYOF | Myoferlin (Fer-1-like protein 3) | 0,734109255 | 0,01249 | 0.114520400967055 |
| Q9BWU0 | NADAP | Kanadaptin (Human lung cancer oncogene 3 protein) (HLC-3) (Kidney anion exchanger adapter protein) (Solute carrier family 4 anion exchanger member 1 adapter protein) | 0,642105906 | 0,01169 | 0.112094959351594 |
| Q4G0N4 | NAKD2 | NAD kinase 2, mitochondrial (EC 2.7.1.23) (Mitochondrial NAD kinase) (NAD kinase domain-containing protein 1, mitochondrial) | 0,514539849 | 0,01866 | 0.134002084277297 |
| Q8NEY1 | NAV1 | Neuron navigator 1 (Pore membrane and/or filament-interacting-like protein 3) (Steerin-1) (Unc-53 homolog 1) (unc53H1) | 0,940728226 | 0,01403 | 0.120558249672262 |
| Q9NXR1 | NDE1 | Nuclear distribution protein nudE homolog 1 (NudE) | 0,648579395 | 0,01321 | 0.117794149768834 |
| P22392 | NDKB | Nucleoside diphosphate kinase B (NDK B) (NDP kinase B) (EC 2.7.4.6) (C-myc purine-binding transcription factor PUF) (Histidine protein kinase NDKB) (EC 2.7.13.3) (nm23-H2) | -0,4519499 | 0,04112 | 0.185884549229643 |
| Q9ULP0 | NDRG4 | Protein NDRG4 (Brain development-related molecule 1) (N-myc downstream-regulated gene 4 protein) (Vascular smooth muscle cell-associated protein 8) (SMAP-8) | -0,858632476 | 0,0287 | 0.160816207866737 |
| P56556 | NDUA6 | NADH dehydrogenase [ubiquinone] 1 alpha subcomplex subunit 6 (Complex I-B14) (CI-B14) (LYR motif-containing protein 6) (NADH-ubiquinone oxidoreductase B14 subunit) | 0,597197579 | 0,0125 | 0.114520400967055 |
| O43676 | NDUB3 | NADH dehydrogenase [ubiquinone] 1 beta subcomplex subunit 3 (Complex I-B12) (CI-B12) (NADH-ubiquinone oxidoreductase B12 subunit) | -0,872753733 | 0,02775 | 0.160514915195513 |
| O95169 | NDUB8 | NADH dehydrogenase [ubiquinone] 1 beta subcomplex subunit 8, mitochondrial (Complex I-ASHI) (CI-ASHI) (NADH-ubiquinone oxidoreductase ASHI subunit) | -0,416421281 | 0,03068 | 0.164699828413934 |
| Q9BU61 | NDUF3 | NADH dehydrogenase [ubiquinone] 1 alpha subcomplex assembly factor 3 | 0,572713686 | 0,01544 | 0.124751718030612 |
| Q96SB3 | NEB2 | Neurabin-2 (Neurabin-II) (Protein phosphatase 1 regulatory subunit 9B) (Spinophilin) | 0,39707622 | 0,04412 | 0.191839076219091 |
| Q9NVZ3 | NECP2 | Adaptin ear-binding coat-associated protein 2 (NECAP endocytosis-associated protein 2) (NECAP-2) | 0,507983398 | 0,02851 | 0.160816207866737 |
| Q92692 | NECT2 | Nectin-2 (Herpes virus entry mediator B) (Herpesvirus entry mediator B) (HveB) (Nectin cell adhesion molecule 2) (Poliovirus receptor-related protein 2) (CD antigen CD112) | 0,814578626 | 0,00399 | 0.0785848374826466 |
| Q96PU5 | NED4L | E3 ubiquitin-protein ligase NEDD4-like (EC 2.3.2.26) (HECT-type E3 ubiquitin transferase NED4L) (NEDD4.2) (Nedd4-2) | 1,002504804 | 0,00955 | 0.106645552538302 |
| O60524 | NEMF | Nuclear export mediator factor NEMF (Antigen NY-CO-1) (Serologically defined colon cancer antigen 1) | 0,414919863 | 0,02642 | 0.156041415189838 |
| Q8NFW8 | NEUA | N-acylneuraminate cytidylyltransferase (EC 2.7.7.43) (CMP-N-acetylneuraminic acid synthase) (CMP-NeuNAc synthase) | 0,652096839 | 0,01803 | 0.132658871887263 |
| P19838 | NFKB1 | Nuclear factor NF-kappa-B p105 subunit (DNA-binding factor KBF1) (EBP-1) (Nuclear factor of kappa light polypeptide gene enhancer in B-cells 1) [Cleaved into: Nuclear factor NF-kappa-B p50 subunit] | 0,777773197 | 0,00867 | 0.103882752534462 |
| P07196 | NFL | Neurofilament light polypeptide (NF-L) (68 kDa neurofilament protein) (Neurofilament triplet L protein) | 0,573140755 | 0,03629 | 0.177468177682117 |
| Q9BZQ8 | NIBAN | Protein Niban (Cell growth-inhibiting gene 39 protein) (Protein FAM129A) | -0,521253147 | 0,01106 | 0.110505462518784 |
| Q96TA1 | NIBL1 | Niban-like protein 1 (Meg-3) (Melanoma invasion by ERK) (MINERVA) (Protein FAM129B) | 0,466699175 | 0,02444 | 0.149305815497977 |
| Q8IW45 | NNRD | ATP-dependent (S)-NAD(P)H-hydrate dehydratase (EC 4.2.1.93) (ATP-dependent NAD(P)HX dehydratase) (Carbohydrate kinase domain-containing protein) (NAD(P)HX dehydratase) | -0,356011992 | 0,04715 | 0.198769565715963 |
| Q9BZE4 | NOG1 | Nucleolar GTP-binding protein 1 (Chronic renal failure gene protein) (GTP-binding protein NGB) | 0,853512755 | 0,00794 | 0.102154997647559 |
| O60936 | NOL3 | Nucleolar protein 3 (Apoptosis repressor with CARD) (Muscle-enriched cytoplasmic protein) (Myp) (Nucleolar protein of 30 kDa) (Nop30) | -0,389078484 | 0,03965 | 0.183590812765362 |
| Q14978 | NOLC1 | Nucleolar and coiled-body phosphoprotein 1 (140 kDa nucleolar phosphoprotein) (Nopp140) (Hepatitis C virus NS5A-transactivated protein 13) (HCV NS5A-transactivated protein 13) (Nucleolar 130 kDa protein) (Nucleolar phosphoprotein p130) | 0,618665679 | 0,01033 | 0.109336835224251 |
| Q15233 | NONO | Non-POU domain-containing octamer-binding protein (NonO protein) (54 kDa nuclear RNA- and DNA-binding protein) (55 kDa nuclear protein) (DNA-binding p52/p100 complex, 52 kDa subunit) (NMT55) (p54(nrb)) (p54nrb) | 0,407687501 | 0,03501 | 0.175099419027731 |
| P29474 | NOS3 | Nitric oxide synthase, endothelial (EC 1.14.13.39) (Constitutive NOS) (cNOS) (EC-NOS) (Endothelial NOS) (eNOS) (NOS type III) (NOSIII) | 0,555848915 | 0,01034 | 0.109336835224251 |
| Q8IVI9 | NOSTN | Nostrin (BM247 homolog) (Nitric oxide synthase traffic inducer) (Nitric oxide synthase trafficker) (eNOS-trafficking inducer) | -0,641448878 | 0,00554 | 0.0940424135447919 |
| P06748 | NPM | Nucleophosmin (NPM) (Nucleolar phosphoprotein B23) (Nucleolar protein NO38) (Numatrin) | 0,524828021 | 0,01399 | 0.120558249672262 |
| Q9BS92 | NPS3B | Protein NipSnap homolog 3B (NipSnap3B) (SNAP1) | -0,809789009 | 0,00447 | 0.0828018336726045 |
| Q8IXM6 | NRM | Nurim (Nuclear envelope membrane protein) (Nuclear rim protein) | 0,633363011 | 0,01141 | 0.111960961329472 |
| O14786 | NRP1 | Neuropilin-1 (Vascular endothelial cell growth factor 165 receptor) (CD antigen CD304) | 0,432570027 | 0,03543 | 0.175701151598843 |
| P03886 | NU1M | NADH-ubiquinone oxidoreductase chain 1 (EC 1.6.5.3) (NADH dehydrogenase subunit 1) | -0,68110841 | 0,01105 | 0.110505462518784 |
| Q9Y5Y2 | NUBP2 | Cytosolic Fe-S cluster assembly factor NUBP2 (Nucleotide-binding protein 2) (NBP 2) | 0,500845698 | 0,01666 | 0.129654315874851 |
| Q9H1E3 | NUCKS | Nuclear ubiquitous casein and cyclin-dependent kinase substrate 1 (P1) | 0,555791451 | 0,02327 | 0.146880219728601 |
| P19338 | NUCL | Nucleolin (Protein C23) | 0,404405737 | 0,04998 | 0.204754344144234 |
| Q9BW91 | NUDT9 | ADP-ribose pyrophosphatase, mitochondrial (EC 3.6.1.13) (ADP-ribose diphosphatase) (ADP-ribose phosphohydrolase) (Adenosine diphosphoribose pyrophosphatase) (ADPR-PPase) (Nucleoside diphosphate-linked moiety X motif 9) (Nudix motif 9) | -0,408448432 | 0,04019 | 0.18383048391046 |
| Q14980 | NUMA1 | Nuclear mitotic apparatus protein 1 (Nuclear matrix protein-22) (NMP-22) (Nuclear mitotic apparatus protein) (NuMA protein) (SP-H antigen) | 0,424234968 | 0,03811 | 0.180517494421774 |
| Q9UBU9 | NXF1 | Nuclear RNA export factor 1 (Tip-associated protein) (Tip-associating protein) (mRNA export factor TAP) | 0,457859782 | 0,01973 | 0.137370395567352 |
| Q6DKJ4 | NXN | Nucleoredoxin (EC 1.8.1.8) | 0,738708607 | 0,00317 | 0.0761554511764352 |
| Q5SWX8 | ODR4 | Protein odr-4 homolog (hODR-4) (LAG1-interacting protein) (Transactivated by transforming growth factor beta protein 1) | 0,429767788 | 0,04881 | 0.202356616939561 |
| Q9NTK5 | OLA1 | Obg-like ATPase 1 (DNA damage-regulated overexpressed in cancer 45) (DOC45) (GTP-binding protein 9) | -0,478593558 | 0,04532 | 0.194611484643512 |
| Q6UWY5 | OLFL1 | Olfactomedin-like protein 1 | 0,943817995 | 0,03758 | 0.179967517866609 |
| Q9NRN5 | OLFL3 | Olfactomedin-like protein 3 (HNOEL-iso) (hOLF44) | 0,676006972 | 0,03585 | 0.176070600102387 |
| Q96E52 | OMA1 | Metalloendopeptidase OMA1, mitochondrial (EC 3.4.24.-) (Metalloprotease-related protein 1) (MPRP-1) (Overlapping with the m-AAA protease 1 homolog) | -0,922441922 | 0,00382 | 0.0771477016366249 |
| Q9Y3B8 | ORN | Oligoribonuclease, mitochondrial (EC 3.1.-.-) (RNA exonuclease 2 homolog) (Small fragment nuclease) | -0,409127592 | 0,04864 | 0.201882368160302 |
| Q96SU4 | OSBL9 | Oxysterol-binding protein-related protein 9 (ORP-9) (OSBP-related protein 9) | 0,928320224 | 0,00181 | 0.0697416810103427 |
| Q32P28 | P3H1 | Prolyl 3-hydroxylase 1 (EC 1.14.11.7) (Growth suppressor 1) (Leucine- and proline-enriched proteoglycan 1) (Leprecan-1) | 0,984799141 | 0,0081 | 0.102455695466046 |
| O15460 | P4HA2 | Prolyl 4-hydroxylase subunit alpha-2 (4-PH alpha-2) (EC 1.14.11.2) (Procollagen-proline,2-oxoglutarate-4-dioxygenase subunit alpha-2) | 0,715608747 | 0,01318 | 0.117794149768834 |
| P54886 | P5CS | Delta-1-pyrroline-5-carboxylate synthase (P5CS) (Aldehyde dehydrogenase family 18 member A1) [Includes: Glutamate 5-kinase (GK) (EC 2.7.2.11) (Gamma-glutamyl kinase); Gamma-glutamyl phosphate reductase (GPR) (EC 1.2.1.41) (Glutamate-5-semialdehyde dehydrogenase) (Glutamyl-gamma-semialdehyde dehydrogenase)] | 0,927744005 | 0,00215 | 0.0727236209046166 |
| O14683 | P5I11 | Tumor protein p53-inducible protein 11 (p53-induced gene 11 protein) | 0,694910698 | 0,00801 | 0.102154997647559 |
| Q15102 | PA1B3 | Platelet-activating factor acetylhydrolase IB subunit gamma (EC 3.1.1.47) (PAF acetylhydrolase 29 kDa subunit) (PAF-AH 29 kDa subunit) (PAF-AH subunit gamma) (PAFAH subunit gamma) | 0,767649735 | 0,00225 | 0.0735786562712696 |
| Q9UP65 | PA24C | Cytosolic phospholipase A2 gamma (cPLA2-gamma) (EC 3.1.1.4) (Phospholipase A2 group IVC) | 0,436198559 | 0,02331 | 0.146880219728601 |
| P05121 | PAI1 | Plasminogen activator inhibitor 1 (PAI) (PAI-1) (Endothelial plasminogen activator inhibitor) (Serpin E1) | 1,858333535 | 0,00256 | 0.073992460491634 |
| Q460N5 | PAR14 | Poly [ADP-ribose] polymerase 14 (PARP-14) (EC 2.4.2.30) (ADP-ribosyltransferase diphtheria toxin-like 8) (ARTD8) (B aggressive lymphoma protein 2) | 0,460080976 | 0,02331 | 0.146880219728601 |
| P09874 | PARP1 | Poly [ADP-ribose] polymerase 1 (PARP-1) (EC 2.4.2.30) (ADP-ribosyltransferase diphtheria toxin-like 1) (ARTD1) (NAD(+) ADP-ribosyltransferase 1) (ADPRT 1) (Poly[ADP-ribose] synthase 1) | 0,547534342 | 0,03848 | 0.1811476445859 |
| Q9HBI1 | PARVB | Beta-parvin (Affixin) | -0,611844524 | 0,00798 | 0.102154997647559 |
| Q9BUH6 | PAXX | Protein PAXX (Paralog of XRCC4 and XLF) (XRCC4-like small protein) | 0,485959807 | 0,02394 | 0.148779323593954 |
| Q15154 | PCM1 | Pericentriolar material 1 protein (PCM-1) (hPCM-1) | 0,408597875 | 0,03708 | 0.178871835755882 |
| O75340 | PDCD6 | Programmed cell death protein 6 (Apoptosis-linked gene 2 protein homolog) (ALG-2) | 0,391185851 | 0,03199 | 0.168322690360001 |
| Q6L8Q7 | PDE12 | 2',5'-phosphodiesterase 12 (2'-PDE) (2-PDE) (EC 3.1.4.-) (Mitochondrial deadenylase) (EC 3.1.13.4) | 0,419820167 | 0,04347 | 0.190053906769213 |
| P13667 | PDIA4 | Protein disulfide-isomerase A4 (EC 5.3.4.1) (Endoplasmic reticulum resident protein 70) (ER protein 70) (ERp70) (Endoplasmic reticulum resident protein 72) (ER protein 72) (ERp-72) (ERp72) | 0,438625289 | 0,04225 | 0.188734490032972 |
| P50479 | PDLI4 | PDZ and LIM domain protein 4 (LIM protein RIL) (Reversion-induced LIM protein) | 1,377173804 | 0,02149 | 0.141824722558827 |
| Q29RF7 | PDS5A | Sister chromatid cohesion protein PDS5 homolog A (Cell proliferation-inducing gene 54 protein) (Sister chromatid cohesion protein 112) (SCC-112) | 0,565585863 | 0,00884 | 0.103882752534462 |
| Q15121 | PEA15 | Astrocytic phosphoprotein PEA-15 (15 kDa phosphoprotein enriched in astrocytes) (Phosphoprotein enriched in diabetes) (PED) | 0,396565936 | 0,03651 | 0.177612813348651 |
| P30086 | PEBP1 | Phosphatidylethanolamine-binding protein 1 (PEBP-1) (HCNPpp) (Neuropolypeptide h3) (Prostatic-binding protein) (Raf kinase inhibitor protein) (RKIP) [Cleaved into: Hippocampal cholinergic neurostimulating peptide (HCNP)] | -0,570884453 | 0,01258 | 0.115044256595559 |
| P05164 | PERM | Myeloperoxidase (MPO) (EC 1.11.2.2) [Cleaved into: Myeloperoxidase; 89 kDa myeloperoxidase; 84 kDa myeloperoxidase; Myeloperoxidase light chain; Myeloperoxidase heavy chain] | 2,521979383 | 0,00634 | 0.0954651715986055 |
| O00541 | PESC | Pescadillo homolog | 0,775496098 | 0,02679 | 0.15719965481106 |
| Q99471 | PFD5 | Prefoldin subunit 5 (Myc modulator 1) (c-Myc-binding protein Mm-1) | -0,521539638 | 0,01731 | 0.13109533821508 |
| P17858 | PFKAL | ATP-dependent 6-phosphofructokinase, liver type (ATP-PFK) (PFK-L) (EC 2.7.1.11) (6-phosphofructokinase type B) (Phosphofructo-1-kinase isozyme B) (PFK-B) (Phosphohexokinase) | 0,669259216 | 0,01341 | 0.118216489113017 |
| P18669 | PGAM1 | Phosphoglycerate mutase 1 (EC 5.4.2.11) (EC 5.4.2.4) (BPG-dependent PGAM 1) (Phosphoglycerate mutase isozyme B) (PGAM-B) | 0,53034659 | 0,02903 | 0.161086911482475 |
| P98160 | PGBM | Basement membrane-specific heparan sulfate proteoglycan core protein (HSPG) (Perlecan) (PLC) [Cleaved into: Endorepellin; LG3 peptide] | 0,713403484 | 0,02688 | 0.15719965481106 |
| Q9H7Z7 | PGES2 | Prostaglandin E synthase 2 (Membrane-associated prostaglandin E synthase-2) (mPGE synthase-2) (Microsomal prostaglandin E synthase 2) (mPGES-2) (Prostaglandin-H(2) E-isomerase) (EC 5.3.99.3) [Cleaved into: Prostaglandin E synthase 2 truncated form] | -0,412042033 | 0,03533 | 0.175701151598843 |
| P09619 | PGFRB | Platelet-derived growth factor receptor beta (PDGF-R-beta) (PDGFR-beta) (EC 2.7.10.1) (Beta platelet-derived growth factor receptor) (Beta-type platelet-derived growth factor receptor) (CD140 antigen-like family member B) (Platelet-derived growth factor receptor 1) (PDGFR-1) (CD antigen CD140b) | 0,702238795 | 0,00404 | 0.0788100563354666 |
| Q96PD5 | PGRP2 | N-acetylmuramoyl-L-alanine amidase (EC 3.5.1.28) (Peptidoglycan recognition protein 2) (Peptidoglycan recognition protein long) (PGRP-L) | 1,108079388 | 0,00044 | 0.0460405438333303 |
| P21810 | PGS1 | Biglycan (Bone/cartilage proteoglycan I) (PG-S1) | 1,176533021 | 0,00313 | 0.0761554511764352 |
| Q92696 | PGTA | Geranylgeranyl transferase type-2 subunit alpha (EC 2.5.1.60) (Geranylgeranyl transferase type II subunit alpha) (Rab geranyl-geranyltransferase subunit alpha) (Rab GG transferase alpha) (Rab GGTase alpha) (Rab geranylgeranyltransferase subunit alpha) | -0,425743433 | 0,04053 | 0.184276431928827 |
| Q86SQ0 | PHLB2 | Pleckstrin homology-like domain family B member 2 (Protein LL5-beta) | 0,575062149 | 0,02711 | 0.157837513271819 |
| P61457 | PHS | Pterin-4-alpha-carbinolamine dehydratase (PHS) (EC 4.2.1.96) (4-alpha-hydroxy-tetrahydropterin dehydratase) (Dimerization cofactor of hepatocyte nuclear factor 1-alpha) (DCoH) (Dimerization cofactor of HNF1) (Phenylalanine hydroxylase-stimulating protein) (Pterin carbinolamine dehydratase) (PCD) | 0,551152182 | 0,01152 | 0.111960961329472 |
| Q969N2 | PIGT | GPI transamidase component PIG-T (Phosphatidylinositol-glycan biosynthesis class T protein) | 0,613661963 | 0,00747 | 0.101606052982629 |
| Q00169 | PIPNA | Phosphatidylinositol transfer protein alpha isoform (PI-TP-alpha) (PtdIns transfer protein alpha) (PtdInsTP alpha) | -0,409717933 | 0,03397 | 0.173091625713912 |
| Q9GZP4 | PITH1 | PITH domain-containing protein 1 | 0,435167504 | 0,02129 | 0.141512996692124 |
| Q8IWE5 | PKHM2 | Pleckstrin homology domain-containing family M member 2 (PH domain-containing family M member 2) (Salmonella-induced filaments A and kinesin-interacting protein) (SifA and kinesin-interacting protein) | 0,549583283 | 0,01406 | 0.120558249672262 |
| Q8TD55 | PKHO2 | Pleckstrin homology domain-containing family O member 2 (PH domain-containing family O member 2) (Pleckstrin homology domain-containing family Q member 1) (PH domain-containing family Q member 1) | 0,683406491 | 0,00541 | 0.0923083755221002 |
| Q99943 | PLCA | 1-acyl-sn-glycerol-3-phosphate acyltransferase alpha (EC 2.3.1.51) (1-acylglycerol-3-phosphate O-acyltransferase 1) (1-AGP acyltransferase 1) (1-AGPAT 1) (Lysophosphatidic acid acyltransferase alpha) (LPAAT-alpha) (Protein G15) | 0,517220819 | 0,04461 | 0.192680841415088 |
| O60240 | PLIN1 | Perilipin-1 (Lipid droplet-associated protein) | 1,466687762 | 0,01035 | 0.109336835224251 |
| Q99541 | PLIN2 | Perilipin-2 (Adipophilin) (Adipose differentiation-related protein) (ADRP) | 0,911021125 | 0,00166 | 0.0673661906302801 |
| P00747 | PLMN | Plasminogen (EC 3.4.21.7) [Cleaved into: Plasmin heavy chain A; Activation peptide; Angiostatin; Plasmin heavy chain A, short form; Plasmin light chain B] | 2,084292368 | 0,00029 | 0.0460405438333303 |
| Q02809 | PLOD1 | Procollagen-lysine,2-oxoglutarate 5-dioxygenase 1 (EC 1.14.11.4) (Lysyl hydroxylase 1) (LH1) | 0,877732187 | 0,02472 | 0.150084617127734 |
| O60568 | PLOD3 | Procollagen-lysine,2-oxoglutarate 5-dioxygenase 3 (EC 1.14.11.4) (Lysyl hydroxylase 3) (LH3) | 1,06323661 | 0,0038 | 0.0771477016366249 |
| P13796 | PLSL | Plastin-2 (L-plastin) (LC64P) (Lymphocyte cytosolic protein 1) (LCP-1) | 1,11884096 | 0,01107 | 0.110505462518784 |
| O43157 | PLXB1 | Plexin-B1 (Semaphorin receptor SEP) | 0,539587311 | 0,02995 | 0.163163481826501 |
| Q9Y4D7 | PLXD1 | Plexin-D1 | 0,503114157 | 0,01521 | 0.124447705141253 |
| P07738 | PMGE | Bisphosphoglycerate mutase (BPGM) (EC 5.4.2.4) (2,3-bisphosphoglycerate mutase, erythrocyte) (2,3-bisphosphoglycerate synthase) (EC 5.4.2.11) (2,3-diphosphoglycerate mutase) (DPGM) (BPG-dependent PGAM) | 1,407310484 | 0,00034 | 0.0460405438333303 |
| Q6XQN6 | PNCB | Nicotinate phosphoribosyltransferase (NAPRTase) (EC 6.3.4.21) (FHA-HIT-interacting protein) (Nicotinate phosphoribosyltransferase domain-containing protein 1) | 0,666891026 | 0,00588 | 0.0945193206940691 |
| Q96T60 | PNKP | Bifunctional polynucleotide phosphatase/kinase (DNA 5'-kinase/3'-phosphatase) (Polynucleotide kinase-3'-phosphatase) [Includes: Polynucleotide 3'-phosphatase (EC 3.1.3.32) (2'(3')-polynucleotidase); Polynucleotide 5'-hydroxyl-kinase (EC 2.7.1.78)] | 0,737648445 | 0,03296 | 0.171077679982414 |
| P00491 | PNPH | Purine nucleoside phosphorylase (PNP) (EC 2.4.2.1) (Inosine phosphorylase) (Inosine-guanosine phosphorylase) | 1,071456202 | 0,00046 | 0.0460405438333303 |
| Q8WVV4 | POF1B | Protein POF1B (Premature ovarian failure protein 1B) | 0,45062035 | 0,02438 | 0.149305815497977 |
| P27169 | PON1 | Serum paraoxonase/arylesterase 1 (PON 1) (EC 3.1.1.2) (EC 3.1.1.81) (EC 3.1.8.1) (Aromatic esterase 1) (A-esterase 1) (K-45) (Serum aryldialkylphosphatase 1) | 1,857126431 | 0,00076 | 0.0473327203168651 |
| B7ZBB8 | PP13G | Protein phosphatase 1 regulatory subunit 3G | 1,176041423 | 0,00087 | 0.0495496681815962 |
| P05186 | PPBT | Alkaline phosphatase, tissue-nonspecific isozyme (AP-TNAP) (TNSALP) (EC 3.1.3.1) (Alkaline phosphatase liver/bone/kidney isozyme) | -0,722897415 | 0,01842 | 0.133184461911317 |
| P23284 | PPIB | Peptidyl-prolyl cis-trans isomerase B (PPIase B) (EC 5.2.1.8) (CYP-S1) (Cyclophilin B) (Rotamase B) (S-cyclophilin) (SCYLP) | 0,540355556 | 0,01276 | 0.115590326418118 |
| Q9Y3C6 | PPIL1 | Peptidyl-prolyl cis-trans isomerase-like 1 (PPIase) (EC 5.2.1.8) (Rotamase PPIL1) | -0,558854719 | 0,03784 | 0.180046072726016 |
| P26678 | PPLA | Cardiac phospholamban (PLB) | 1,91408599 | 0,00016 | 0.0377697242370153 |
| P49593 | PPM1F | Protein phosphatase 1F (EC 3.1.3.16) (Ca(2+)/calmodulin-dependent protein kinase phosphatase) (CaM-kinase phosphatase) (CaMKPase) (Partner of PIX 2) (Protein fem-2 homolog) (hFem-2) | 0,46568194 | 0,03544 | 0.175701151598843 |
| Q8N3J5 | PPM1K | Protein phosphatase 1K, mitochondrial (EC 3.1.3.16) (PP2C domain-containing protein phosphatase 1K) (PP2C-like mitochondrial protein) (PP2C-type mitochondrial phosphoprotein phosphatase) (PTMP) (Protein phosphatase 2C isoform kappa) (PP2C-kappa) | -0,844947659 | 0,00399 | 0.0785848374826466 |
| P50897 | PPT1 | Palmitoyl-protein thioesterase 1 (PPT-1) (EC 3.1.2.22) (Palmitoyl-protein hydrolase 1) | -0,678043451 | 0,02458 | 0.149799934784929 |
| O75400 | PR40A | Pre-mRNA-processing factor 40 homolog A (Fas ligand-associated factor 1) (Formin-binding protein 11) (Formin-binding protein 3) (Huntingtin yeast partner A) (Huntingtin-interacting protein 10) (HIP-10) (Huntingtin-interacting protein A) (Renal carcinoma antigen NY-REN-6) | 0,397134266 | 0,03188 | 0.168140913413884 |
| Q06830 | PRDX1 | Peroxiredoxin-1 (EC 1.11.1.15) (Natural killer cell-enhancing factor A) (NKEF-A) (Proliferation-associated gene protein) (PAG) (Thioredoxin peroxidase 2) (Thioredoxin-dependent peroxide reductase 2) | -0,636001159 | 0,00775 | 0.102154997647559 |
| P30041 | PRDX6 | Peroxiredoxin-6 (EC 1.11.1.15) (1-Cys peroxiredoxin) (1-Cys PRX) (24 kDa protein) (Acidic calcium-independent phospholipase A2) (aiPLA2) (EC 3.1.1.4) (Antioxidant protein 2) (Liver 2D page spot 40) (Non-selenium glutathione peroxidase) (NSGPx) (Red blood cells page spot 12) | -0,514512925 | 0,03972 | 0.183630278609946 |
| P51888 | PRELP | Prolargin (Proline-arginine-rich end leucine-rich repeat protein) | 0,773144005 | 0,01295 | 0.116996060183678 |
| P78527 | PRKDC | DNA-dependent protein kinase catalytic subunit (DNA-PK catalytic subunit) (DNA-PKcs) (EC 2.7.11.1) (DNPK1) (p460) | 0,45474839 | 0,02251 | 0.145713783110726 |
| P07737 | PROF1 | Profilin-1 (Epididymis tissue protein Li 184a) (Profilin I) | 0,468134697 | 0,02926 | 0.161086911482475 |
| P35080 | PROF2 | Profilin-2 (Profilin II) | -0,45297354 | 0,04005 | 0.18383048391046 |
| P07225 | PROS | Vitamin K-dependent protein S | 0,775303589 | 0,0047 | 0.085284948858929 |
| O43172 | PRP4 | U4/U6 small nuclear ribonucleoprotein Prp4 (PRP4 homolog) (hPrp4) (U4/U6 snRNP 60 kDa protein) (WD splicing factor Prp4) | 0,883970132 | 0,00618 | 0.0954651715986055 |
| Q6P2Q9 | PRP8 | Pre-mRNA-processing-splicing factor 8 (220 kDa U5 snRNP-specific protein) (PRP8 homolog) (Splicing factor Prp8) (p220) | 0,585801536 | 0,01003 | 0.109336835224251 |
| O43395 | PRPF3 | U4/U6 small nuclear ribonucleoprotein Prp3 (Pre-mRNA-splicing factor 3) (hPrp3) (U4/U6 snRNP 90 kDa protein) | 0,54635278 | 0,01386 | 0.119960852796423 |
| P60891 | PRPS1 | Ribose-phosphate pyrophosphokinase 1 (EC 2.7.6.1) (PPRibP) (Phosphoribosyl pyrophosphate synthase I) (PRS-I) | 0,545547223 | 0,01909 | 0.135774205040182 |
| P24158 | PRTN3 | Myeloblastin (EC 3.4.21.76) (AGP7) (C-ANCA antigen) (Leukocyte proteinase 3) (PR-3) (PR3) (Neutrophil proteinase 4) (NP-4) (P29) (Wegener autoantigen) | 3,472428242 | 0,00102 | 0.0538418641701199 |
| Q99436 | PSB7 | Proteasome subunit beta type-7 (EC 3.4.25.1) (Macropain chain Z) (Multicatalytic endopeptidase complex chain Z) (Proteasome subunit Z) | -0,541857009 | 0,01997 | 0.138103098631934 |
| O75475 | PSIP1 | PC4 and SFRS1-interacting protein (CLL-associated antigen KW-7) (Dense fine speckles 70 kDa protein) (DFS 70) (Lens epithelium-derived growth factor) (Transcriptional coactivator p75/p52) | 0,57032053 | 0,01786 | 0.132658871887263 |
| Q16401 | PSMD5 | 26S proteasome non-ATPase regulatory subunit 5 (26S protease subunit S5 basic) (26S proteasome subunit S5B) | 0,661549762 | 0,00734 | 0.100950143902858 |
| Q9UL46 | PSME2 | Proteasome activator complex subunit 2 (11S regulator complex subunit beta) (REG-beta) (Activator of multicatalytic protease subunit 2) (Proteasome activator 28 subunit beta) (PA28b) (PA28beta) | -0,721369886 | 0,01445 | 0.121895247571125 |
| Q8WXF1 | PSPC1 | Paraspeckle component 1 (Paraspeckle protein 1) | 0,472446223 | 0,03324 | 0.171297602744958 |
| P26599 | PTBP1 | Polypyrimidine tract-binding protein 1 (PTB) (57 kDa RNA-binding protein PPTB-1) (Heterogeneous nuclear ribonucleoprotein I) (hnRNP I) | 0,649572181 | 0,01188 | 0.112411439550033 |
| Q9UKA9 | PTBP2 | Polypyrimidine tract-binding protein 2 (Neural polypyrimidine tract-binding protein) (Neurally-enriched homolog of PTB) (PTB-like protein) | 0,590248936 | 0,01449 | 0.121933727203822 |
| P29350 | PTN6 | Tyrosine-protein phosphatase non-receptor type 6 (EC 3.1.3.48) (Hematopoietic cell protein-tyrosine phosphatase) (Protein-tyrosine phosphatase 1C) (PTP-1C) (Protein-tyrosine phosphatase SHP-1) (SH-PTP1) | 1,081625233 | 0,01032 | 0.109336835224251 |
| P08575 | PTPRC | Receptor-type tyrosine-protein phosphatase C (EC 3.1.3.48) (Leukocyte common antigen) (L-CA) (T200) (CD antigen CD45) | 0,944983721 | 0,00167 | 0.0673661906302801 |
| Q6GMV3 | PTRD1 | Putative peptidyl-tRNA hydrolase PTRHD1 (EC 3.1.1.29) (Peptidyl-tRNA hydrolase domain-containing protein 1) | -1,023330303 | 0,02025 | 0.138876689931427 |
| O15067 | PUR4 | Phosphoribosylformylglycinamidine synthase (FGAM synthase) (FGAMS) (EC 6.3.5.3) (Formylglycinamide ribonucleotide amidotransferase) (FGAR amidotransferase) (FGAR-AT) (Formylglycinamide ribotide amidotransferase) | 0,65489904 | 0,00538 | 0.0921423483083817 |
| Q8N142 | PURA1 | Adenylosuccinate synthetase isozyme 1 (AMPSase 1) (AdSS 1) (EC 6.3.4.4) (Adenylosuccinate synthetase, basic isozyme) (Adenylosuccinate synthetase, muscle isozyme) (M-type adenylosuccinate synthetase) (IMP--aspartate ligase 1) | 0,702785606 | 0,00311 | 0.0761554511764352 |
| Q96QR8 | PURB | Transcriptional activator protein Pur-beta (Purine-rich element-binding protein B) | 0,507066597 | 0,01694 | 0.130074948166753 |
| P15151 | PVR | Poliovirus receptor (Nectin-like protein 5) (NECL-5) (CD antigen CD155) | 0,563832135 | 0,00997 | 0.109336835224251 |
| O96011 | PX11B | Peroxisomal membrane protein 11B (Peroxin-11B) (Peroxisomal biogenesis factor 11B) (Protein PEX11 homolog beta) (PEX11-beta) | 0,405463886 | 0,03337 | 0.171528636510207 |
| Q92626 | PXDN | Peroxidasin homolog (EC 1.11.1.7) (Melanoma-associated antigen MG50) (Vascular peroxidase 1) (p53-responsive gene 2 protein) | 0,757961116 | 0,00625 | 0.0954651715986055 |
| Q9NR77 | PXMP2 | Peroxisomal membrane protein 2 (22 kDa peroxisomal membrane protein) | 0,722974719 | 0,02025 | 0.138876689931427 |
| P06737 | PYGL | Glycogen phosphorylase, liver form (EC 2.4.1.1) | 0,686939471 | 0,04337 | 0.189843976360776 |
| P27708 | PYR1 | CAD protein [Includes: Glutamine-dependent carbamoyl-phosphate synthase (EC 6.3.5.5); Aspartate carbamoyltransferase (EC 2.1.3.2); Dihydroorotase (EC 3.5.2.3)] | 1,009468435 | 0,00188 | 0.0703852100453896 |
| P17812 | PYRG1 | CTP synthase 1 (EC 6.3.4.2) (CTP synthetase 1) (UTP--ammonia ligase 1) | 0,500202265 | 0,03461 | 0.174037405274487 |
| Q08257 | QOR | Quinone oxidoreductase (EC 1.6.5.5) (NADPH:quinone reductase) (Zeta-crystallin) | -0,440260933 | 0,03505 | 0.175099419027731 |
| O95825 | QORL1 | Quinone oxidoreductase-like protein 1 (EC 1.-.-.-) (Protein 4P11) (Quinone oxidoreductase homolog 1) (QOH-1) (Zeta-crystallin homolog) | 0,427029731 | 0,04004 | 0.18383048391046 |
| Q9BZG1 | RAB34 | Ras-related protein Rab-34 (Ras-related protein Rab-39) (Ras-related protein Rah) | 0,5336014 | 0,01362 | 0.11913827875583 |
| P20338 | RAB4A | Ras-related protein Rab-4A | -0,529588277 | 0,04747 | 0.199326629362477 |
| P61018 | RAB4B | Ras-related protein Rab-4B | 0,811872074 | 0,00345 | 0.0764488368109507 |
| P20340 | RAB6A | Ras-related protein Rab-6A (Rab-6) | 0,437537166 | 0,03431 | 0.173610022145665 |
| P51151 | RAB9A | Ras-related protein Rab-9A | -0,425991994 | 0,02897 | 0.161086911482475 |
| P63000 | RAC1 | Ras-related C3 botulinum toxin substrate 1 (Cell migration-inducing gene 5 protein) (Ras-like protein TC25) (p21-Rac1) | -0,664821261 | 0,01459 | 0.122253320238255 |
| P55042 | RAD | GTP-binding protein RAD (RAD1) (Ras associated with diabetes) | 1,074553904 | 0,00073 | 0.0473327203168651 |
| Q92878 | RAD50 | DNA repair protein RAD50 (hRAD50) (EC 3.6.-.-) | 0,530784138 | 0,01025 | 0.109336835224251 |
| Q9P0K7 | RAI14 | Ankycorbin (Ankyrin repeat and coiled-coil structure-containing protein) (Novel retinal pigment epithelial cell protein) (Retinoic acid-induced protein 14) | 0,472950142 | 0,01829 | 0.133184461911317 |
| Q5U651 | RAIN | Ras-interacting protein 1 (Rain) | 0,557913013 | 0,00834 | 0.103093659718687 |
| P62834 | RAP1A | Ras-related protein Rap-1A (C21KG) (G-22K) (GTP-binding protein smg p21A) (Ras-related protein Krev-1) | -0,664541988 | 0,00593 | 0.0945792628211344 |
| Q99969 | RARR2 | Retinoic acid receptor responder protein 2 (Chemerin) (RAR-responsive protein TIG2) (Tazarotene-induced gene 2 protein) | 0,682192675 | 0,04007 | 0.18383048391046 |
| Q8IXT5 | RB12B | RNA-binding protein 12B (RNA-binding motif protein 12B) | 1,084754011 | 0,00441 | 0.0823103193899197 |
| P51159 | RB27A | Ras-related protein Rab-27A (Rab-27) (GTP-binding protein Ram) | 0,712929398 | 0,01045 | 0.109336835224251 |
| Q9H082 | RB33B | Ras-related protein Rab-33B | 0,631001623 | 0,03022 | 0.16369295498359 |
| Q9NTZ6 | RBM12 | RNA-binding protein 12 (RNA-binding motif protein 12) (SH3/WW domain anchor protein in the nucleus) (SWAN) | 0,563026495 | 0,00884 | 0.103882752534462 |
| Q96PK6 | RBM14 | RNA-binding protein 14 (Paraspeckle protein 2) (PSP2) (RNA-binding motif protein 14) (RRM-containing coactivator activator/modulator) (Synaptotagmin-interacting protein) (SYT-interacting protein) | 1,012431479 | 0,0059 | 0.0945193206940691 |
| Q96T37 | RBM15 | RNA-binding protein 15 (One-twenty two protein 1) (RNA-binding motif protein 15) | 0,590327771 | 0,02021 | 0.138876689931427 |
| P98179 | RBM3 | RNA-binding protein 3 (RNA-binding motif protein 3) (RNPL) | -0,717424348 | 0,01319 | 0.117794149768834 |
| Q14498 | RBM39 | RNA-binding protein 39 (CAPER alpha) (Hepatocellular carcinoma protein 1) (RNA-binding motif protein 39) (RNA-binding region-containing protein 2) (Splicing factor HCC1) | 0,558892754 | 0,01667 | 0.129654315874851 |
| P38159 | RBMX | RNA-binding motif protein, X chromosome (Glycoprotein p43) (Heterogeneous nuclear ribonucleoprotein G) (hnRNP G) [Cleaved into: RNA-binding motif protein, X chromosome, N-terminally processed] | 0,97386887 | 0,00154 | 0.0665699421028979 |
| Q9P258 | RCC2 | Protein RCC2 (RCC1-like protein TD-60) (Telophase disk protein of 60 kDa) | 1,120902198 | 0,00037 | 0.0460405438333303 |
| Q96D15 | RCN3 | Reticulocalbin-3 (EF-hand calcium-binding protein RLP49) | 1,555170204 | 0,00203 | 0.0713274753495378 |
| P46063 | RECQ1 | ATP-dependent DNA helicase Q1 (EC 3.6.4.12) (DNA helicase, RecQ-like type 1) (RecQ1) (DNA-dependent ATPase Q1) (RecQ protein-like 1) | 0,513989728 | 0,04199 | 0.188028909639907 |
| Q13123 | RED | Protein Red (Cytokine IK) (IK factor) (Protein RER) | 0,428830656 | 0,04055 | 0.184276431928827 |
| Q00765 | REEP5 | Receptor expression-enhancing protein 5 (Polyposis locus protein 1) (Protein TB2) | -0,822528333 | 0,00243 | 0.073956300294973 |
| P02753 | RET4 | Retinol-binding protein 4 (Plasma retinol-binding protein) (PRBP) (RBP) [Cleaved into: Plasma retinol-binding protein(1-182); Plasma retinol-binding protein(1-181); Plasma retinol-binding protein(1-179); Plasma retinol-binding protein(1-176)] | 1,095539815 | 0,0498 | 0.204754344144234 |
| Q9HD89 | RETN | Resistin (Adipose tissue-specific secretory factor) (ADSF) (C/EBP-epsilon-regulated myeloid-specific secreted cysteine-rich protein) (Cysteine-rich secreted protein A12-alpha-like 2) (Cysteine-rich secreted protein FIZZ3) | 0,77152667 | 0,02826 | 0.160816207866737 |
| Q6NUM9 | RETST | All-trans-retinol 13,14-reductase (EC 1.3.99.23) (All-trans-13,14-dihydroretinol saturase) (RetSat) (PPAR-alpha-regulated and starvation-induced gene protein) | -0,377999914 | 0,03749 | 0.179949680895175 |
| Q9H4X1 | RGCC | Regulator of cell cycle RGCC (Response gene to complement 32 protein) (RGC-32) | -0,872529581 | 0,0142 | 0.120558249672262 |
| O15211 | RGL2 | Ral guanine nucleotide dissociation stimulator-like 2 (RalGDS-like 2) (RalGDS-like factor) (Ras-associated protein RAB2L) | 0,516790397 | 0,01163 | 0.112094959351594 |
| Q96CC6 | RHDF1 | Inactive rhomboid protein 1 (iRhom1) (Epidermal growth factor receptor-related protein) (Rhomboid 5 homolog 1) (Rhomboid family member 1) (p100hRho) | 0,533074885 | 0,0398 | 0.183630278609946 |
| P42331 | RHG25 | Rho GTPase-activating protein 25 (Rho-type GTPase-activating protein 25) | 0,783855456 | 0,00346 | 0.0764488368109507 |
| Q9NRY4 | RHG35 | Rho GTPase-activating protein 35 (Glucocorticoid receptor DNA-binding factor 1) (Glucocorticoid receptor repression factor 1) (GRF-1) (Rho GAP p190A) (p190-A) | 1,296850568 | 0,00018 | 0.039418750910064 |
| P61586 | RHOA | Transforming protein RhoA (Rho cDNA clone 12) (h12) | 0,978942188 | 0,0038 | 0.0771477016366249 |
| Q6R327 | RICTR | Rapamycin-insensitive companion of mTOR (AVO3 homolog) (hAVO3) | 0,46890143 | 0,02215 | 0.144403200936269 |
| P52758 | RIDA | 2-iminobutanoate/2-iminopropanoate deaminase (EC 3.5.99.10) (14.5 kDa translational inhibitor protein) (hp14.5) (p14.5) (Heat-responsive protein 12) (Reactive intermediate imine deaminase A homolog) (Translation inhibitor L-PSP ribonuclease) (EC 3.1.-.-) (UK114 antigen homolog) | -0,709645338 | 0,01511 | 0.123957180633967 |
| Q6ZS17 | RIPR1 | Rho family-interacting cell polarization regulator 1 | 0,414542618 | 0,04729 | 0.198769565715963 |
| P27635 | RL10 | 60S ribosomal protein L10 (Laminin receptor homolog) (Large ribosomal subunit protein uL16) (Protein QM) (Ribosomal protein L10) (Tumor suppressor QM) | 0,354099163 | 0,04928 | 0.203866488649309 |
| P26373 | RL13 | 60S ribosomal protein L13 (Breast basic conserved protein 1) (Large ribosomal subunit protein eL13) | 0,609000008 | 0,00917 | 0.104035630640557 |
| P40429 | RL13A | 60S ribosomal protein L13a (23 kDa highly basic protein) (Large ribosomal subunit protein uL13) | 0,977363902 | 0,00411 | 0.0790778462733819 |
| P50914 | RL14 | 60S ribosomal protein L14 (CAG-ISL 7) (Large ribosomal subunit protein eL14) | 0,426304626 | 0,02319 | 0.146880219728601 |
| P61313 | RL15 | 60S ribosomal protein L15 (Large ribosomal subunit protein eL15) | 1,021849462 | 0,02126 | 0.141512996692124 |
| P18621 | RL17 | 60S ribosomal protein L17 (60S ribosomal protein L23) (Large ribosomal subunit protein uL22) (PD-1) | 0,526784787 | 0,01041 | 0.109336835224251 |
| Q07020 | RL18 | 60S ribosomal protein L18 (Large ribosomal subunit protein eL18) | 0,898977889 | 0,00204 | 0.0713274753495378 |
| P84098 | RL19 | 60S ribosomal protein L19 (Large ribosomal subunit protein eL19) | 0,671583525 | 0,00521 | 0.090331683207465 |
| O76021 | RL1D1 | Ribosomal L1 domain-containing protein 1 (CATX-11) (Cellular senescence-inhibited gene protein) (Protein PBK1) | 0,822824481 | 0,00514 | 0.0899766492445311 |
| P83731 | RL24 | 60S ribosomal protein L24 (60S ribosomal protein L30) (Large ribosomal subunit protein eL24) | 0,595705068 | 0,02542 | 0.15245012576966 |
| P46776 | RL27A | 60S ribosomal protein L27a (Large ribosomal subunit protein uL15) | 0,553210134 | 0,01032 | 0.109336835224251 |
| P39023 | RL3 | 60S ribosomal protein L3 (HIV-1 TAR RNA-binding protein B) (TARBP-B) (Large ribosomal subunit protein uL3) | 0,853836594 | 0,0036 | 0.0767928025552317 |
| P62899 | RL31 | 60S ribosomal protein L31 (Large ribosomal subunit protein eL31) | 0,490751729 | 0,01701 | 0.130074948166753 |
| P62910 | RL32 | 60S ribosomal protein L32 (Large ribosomal subunit protein eL32) | 0,741524241 | 0,00274 | 0.0759058199506327 |
| P42766 | RL35 | 60S ribosomal protein L35 (Large ribosomal subunit protein uL29) | 0,878318068 | 0,00264 | 0.0755323007278025 |
| Q9Y3U8 | RL36 | 60S ribosomal protein L36 (Large ribosomal subunit protein eL36) | 0,486552779 | 0,04233 | 0.188743139597737 |
| Q969Q0 | RL36L | 60S ribosomal protein L36a-like (Large ribosomal subunit protein eL42-like) | 0,438977236 | 0,03645 | 0.177612813348651 |
| P61927 | RL37 | 60S ribosomal protein L37 (G1.16) (Large ribosomal subunit protein eL37) | 0,667762973 | 0,01815 | 0.132794115698057 |
| P36578 | RL4 | 60S ribosomal protein L4 (60S ribosomal protein L1) (Large ribosomal subunit protein uL4) | 0,430824275 | 0,02672 | 0.15719965481106 |
| Q02878 | RL6 | 60S ribosomal protein L6 (Large ribosomal subunit protein eL6) (Neoplasm-related protein C140) (Tax-responsive enhancer element-binding protein 107) (TaxREB107) | 0,471877746 | 0,04118 | 0.185884549229643 |
| P18124 | RL7 | 60S ribosomal protein L7 (Large ribosomal subunit protein uL30) | 0,495040421 | 0,02403 | 0.149130557227562 |
| P62917 | RL8 | 60S ribosomal protein L8 (Large ribosomal subunit protein uL2) | 0,662307312 | 0,01586 | 0.126423982233753 |
| Q96DB5 | RMD1 | Regulator of microtubule dynamics protein 1 (RMD-1) (hRMD-1) (Protein FAM82B) | -0,460793666 | 0,02842 | 0.160816207866737 |
| Q96LZ7 | RMD2 | Regulator of microtubule dynamics protein 2 (RMD-2) (hRMD-2) (Protein FAM82A1) | -0,732817451 | 0,04449 | 0.192406103238607 |
| Q9UBS8 | RNF14 | E3 ubiquitin-protein ligase RNF14 (EC 2.3.2.31) (Androgen receptor-associated protein 54) (HFB30) (RING finger protein 14) (Triad2 protein) | -0,564754212 | 0,00826 | 0.102945529755352 |
| Q13151 | ROA0 | Heterogeneous nuclear ribonucleoprotein A0 (hnRNP A0) | 0,516425332 | 0,04042 | 0.184122814311903 |
| P22626 | ROA2 | Heterogeneous nuclear ribonucleoproteins A2/B1 (hnRNP A2/B1) | 0,541411977 | 0,01341 | 0.118216489113017 |
| P51991 | ROA3 | Heterogeneous nuclear ribonucleoprotein A3 (hnRNP A3) | 0,836776547 | 0,00802 | 0.102154997647559 |
| P24928 | RPB1 | DNA-directed RNA polymerase II subunit RPB1 (RNA polymerase II subunit B1) (EC 2.7.7.6) (DNA-directed RNA polymerase II subunit A) (DNA-directed RNA polymerase III largest subunit) (RNA-directed RNA polymerase II subunit RPB1) (EC 2.7.7.48) | 0,704435179 | 0,00312 | 0.0761554511764352 |
| P30876 | RPB2 | DNA-directed RNA polymerase II subunit RPB2 (EC 2.7.7.6) (DNA-directed RNA polymerase II 140 kDa polypeptide) (DNA-directed RNA polymerase II subunit B) (RNA polymerase II subunit 2) (RNA polymerase II subunit B2) | 0,590048392 | 0,02141 | 0.141803063485846 |
| Q9NQG5 | RPR1B | Regulation of nuclear pre-mRNA domain-containing protein 1B (Cell cycle-related and expression-elevated protein in tumor) | 0,611902146 | 0,00954 | 0.106645552538302 |
| Q5VT52 | RPRD2 | Regulation of nuclear pre-mRNA domain-containing protein 2 | 0,565155443 | 0,04375 | 0.190659129510615 |
| Q9P2E9 | RRBP1 | Ribosome-binding protein 1 (180 kDa ribosome receptor homolog) (RRp) (ES/130-related protein) (Ribosome receptor protein) | 0,728464137 | 0,00404 | 0.0788100563354666 |
| Q15050 | RRS1 | Ribosome biogenesis regulatory protein homolog | 0,556098481 | 0,03818 | 0.180630335358898 |
| P62277 | RS13 | 40S ribosomal protein S13 (Small ribosomal subunit protein uS15) | 0,587988575 | 0,01699 | 0.130074948166753 |
| P62244 | RS15A | 40S ribosomal protein S15a (Small ribosomal subunit protein uS8) | 0,544362562 | 0,01235 | 0.114216522917893 |
| P60866 | RS20 | 40S ribosomal protein S20 (Small ribosomal subunit protein uS10) | 0,441385733 | 0,04014 | 0.18383048391046 |
| P62266 | RS23 | 40S ribosomal protein S23 (Small ribosomal subunit protein uS12) | 0,620090489 | 0,00609 | 0.0954651715986055 |
| P62847 | RS24 | 40S ribosomal protein S24 (Small ribosomal subunit protein eS24) | 0,521253066 | 0,03363 | 0.172214098674328 |
| P62857 | RS28 | 40S ribosomal protein S28 (Small ribosomal subunit protein eS28) | 0,612849727 | 0,03927 | 0.183376978972405 |
| P62701 | RS4X | 40S ribosomal protein S4, X isoform (SCR10) (Single copy abundant mRNA protein) (Small ribosomal subunit protein eS4) | 0,51775751 | 0,01769 | 0.131931842633018 |
| P62753 | RS6 | 40S ribosomal protein S6 (Phosphoprotein NP33) (Small ribosomal subunit protein eS6) | 0,457468647 | 0,02859 | 0.160816207866737 |
| P46781 | RS9 | 40S ribosomal protein S9 (Small ribosomal subunit protein uS4) | 0,808867991 | 0,00365 | 0.0767928025552317 |
| P08865 | RSSA | 40S ribosomal protein SA (37 kDa laminin receptor precursor) (37LRP) (37/67 kDa laminin receptor) (LRP/LR) (67 kDa laminin receptor) (67LR) (Colon carcinoma laminin-binding protein) (Laminin receptor 1) (LamR) (Laminin-binding protein precursor p40) (LBP/p40) (Multidrug resistance-associated protein MGr1-Ag) (NEM/1CHD4) (Small ribosomal subunit protein uS2) | 0,439598031 | 0,03104 | 0.165624853763808 |
| P82912 | RT11 | 28S ribosomal protein S11, mitochondrial (MRP-S11) (S11mt) (Cervical cancer proto-oncogene 2 protein) (HCC-2) (Mitochondrial small ribosomal subunit protein uS11m) | 0,447104848 | 0,02348 | 0.147127385900632 |
| Q92541 | RTF1 | RNA polymerase-associated protein RTF1 homolog | 1,251747765 | 0,00379 | 0.0771477016366249 |
| P08621 | RU17 | U1 small nuclear ribonucleoprotein 70 kDa (U1 snRNP 70 kDa) (U1-70K) (snRNP70) | 0,643835115 | 0,03595 | 0.176070600102387 |
| P09661 | RU2A | U2 small nuclear ribonucleoprotein A' (U2 snRNP A') | 0,415325408 | 0,02818 | 0.160816207866737 |
| P08579 | RU2B | U2 small nuclear ribonucleoprotein B'' (U2 snRNP B'') | 0,492106415 | 0,01578 | 0.126187856364557 |
| Q96GQ5 | RUS1 | RUS1 family protein C16orf58 | 1,038680505 | 0,00312 | 0.0761554511764352 |
| P62304 | RUXE | Small nuclear ribonucleoprotein E (snRNP-E) (Sm protein E) (Sm-E) (SmE) | 0,657578567 | 0,01205 | 0.113024663824624 |
| A8MWD9 | RUXGL | Putative small nuclear ribonucleoprotein G-like protein 15 | 0,467200625 | 0,02594 | 0.153725182711907 |
| P28702 | RXRB | Retinoic acid receptor RXR-beta (Nuclear receptor subfamily 2 group B member 2) (Retinoid X receptor beta) | 0,729361518 | 0,00296 | 0.0761554511764352 |
| P25815 | S100P | Protein S100-P (Migration-inducing gene 9 protein) (MIG9) (Protein S100-E) (S100 calcium-binding protein P) | 1,562549892 | 0,00067 | 0.0461437135818825 |
| P23297 | S10A1 | Protein S100-A1 (S-100 protein alpha chain) (S-100 protein subunit alpha) (S100 calcium-binding protein A1) | -0,768744217 | 0,01933 | 0.136364775277862 |
| P05109 | S10A8 | Protein S100-A8 (Calgranulin-A) (Calprotectin L1L subunit) (Cystic fibrosis antigen) (CFAG) (Leukocyte L1 complex light chain) (Migration inhibitory factor-related protein 8) (MRP-8) (p8) (S100 calcium-binding protein A8) (Urinary stone protein band A) | 1,963365782 | 0,00011 | 0.0300468839677373 |
| P06702 | S10A9 | Protein S100-A9 (Calgranulin-B) (Calprotectin L1H subunit) (Leukocyte L1 complex heavy chain) (Migration inhibitory factor-related protein 14) (MRP-14) (p14) (S100 calcium-binding protein A9) | 2,141461182 | 0,00066 | 0.0460405438333303 |
| P31949 | S10AB | Protein S100-A11 (Calgizzarin) (Metastatic lymph node gene 70 protein) (MLN 70) (Protein S100-C) (S100 calcium-binding protein A11) [Cleaved into: Protein S100-A11, N-terminally processed] | 0,567397381 | 0,00858 | 0.103855107408487 |
| P80511 | S10AC | Protein S100-A12 (CGRP) (Calcium-binding protein in amniotic fluid 1) (CAAF1) (Calgranulin-C) (CAGC) (Extracellular newly identified RAGE-binding protein) (EN-RAGE) (Migration inhibitory factor-related protein 6) (MRP-6) (p6) (Neutrophil S100 protein) (S100 calcium-binding protein A12) [Cleaved into: Calcitermin] | 1,913068473 | 0,00037 | 0.0460405438333303 |
| Q5K4L6 | S27A3 | Long-chain fatty acid transport protein 3 (FATP-3) (Fatty acid transport protein 3) (EC 6.2.1.-) (Solute carrier family 27 member 3) (Very long-chain acyl-CoA synthetase homolog 3) (VLCS-3) | 0,594042049 | 0,02686 | 0.15719965481106 |
| P61619 | S61A1 | Protein transport protein Sec61 subunit alpha isoform 1 (Sec61 alpha-1) | 0,457319061 | 0,02144 | 0.141803063485846 |
| P35542 | SAA4 | Serum amyloid A-4 protein (Constitutively expressed serum amyloid A protein) (C-SAA) | 1,553867798 | 0,009 | 0.103994216063343 |
| Q9UBT2 | SAE2 | SUMO-activating enzyme subunit 2 (EC 2.3.2.-) (Anthracycline-associated resistance ARX) (Ubiquitin-like 1-activating enzyme E1B) (Ubiquitin-like modifier-activating enzyme 2) | 0,499432379 | 0,01464 | 0.12245128905555 |
| Q15424 | SAFB1 | Scaffold attachment factor B1 (SAF-B) (SAF-B1) (HSP27 estrogen response element-TATA box-binding protein) (HSP27 ERE-TATA-binding protein) | 0,432062812 | 0,02546 | 0.15245012576966 |
| Q9Y512 | SAM50 | Sorting and assembly machinery component 50 homolog (Transformation-related gene 3 protein) (TRG-3) | -0,47426118 | 0,0207 | 0.139808301680561 |
| Q9Y6B6 | SAR1B | GTP-binding protein SAR1b (GTP-binding protein B) (GTBPB) | -0,523682459 | 0,02334 | 0.146880219728601 |
| Q6SZW1 | SARM1 | Sterile alpha and TIR motif-containing protein 1 (Sterile alpha and Armadillo repeat protein) (Sterile alpha motif domain-containing protein 2) (MyD88-5) (SAM domain-containing protein 2) (Tir-1 homolog) | 0,734213922 | 0,04944 | 0.204121656015476 |
| Q15020 | SART3 | Squamous cell carcinoma antigen recognized by T-cells 3 (SART-3) (Tat-interacting protein of 110 kDa) (Tip110) (p110 nuclear RNA-binding protein) | 0,471225154 | 0,01735 | 0.13109533821508 |
| P67812 | SC11A | Signal peptidase complex catalytic subunit SEC11A (EC 3.4.21.89) (Endopeptidase SP18) (Microsomal signal peptidase 18 kDa subunit) (SPase 18 kDa subunit) (SEC11 homolog A) (SEC11-like protein 1) (SPC18) | 0,619072428 | 0,01152 | 0.111960961329472 |
| O15027 | SC16A | Protein transport protein Sec16A (SEC16 homolog A) | 0,4838866 | 0,03332 | 0.171517898123243 |
| P53992 | SC24C | Protein transport protein Sec24C (SEC24-related protein C) | 0,565254683 | 0,01184 | 0.112411439550033 |
| O94979 | SC31A | Protein transport protein Sec31A (ABP125) (ABP130) (SEC31-like protein 1) (SEC31-related protein A) (Web1-like protein) | 0,800264219 | 0,00325 | 0.0761554511764352 |
| Q14108 | SCRB2 | Lysosome membrane protein 2 (85 kDa lysosomal membrane sialoglycoprotein) (LGP85) (CD36 antigen-like 2) (Lysosome membrane protein II) (LIMP II) (Scavenger receptor class B member 2) (CD antigen CD36) | -0,584004519 | 0,00957 | 0.106645552538302 |
| Q96GA7 | SDSL | Serine dehydratase-like (L-serine deaminase) (L-serine dehydratase/L-threonine deaminase) (L-threonine dehydratase) (TDH) (EC 4.3.1.19) (Serine dehydratase 2) (SDH 2) (EC 4.3.1.17) | -0,716889063 | 0,01156 | 0.111960961329472 |
| P55735 | SEC13 | Protein SEC13 homolog (GATOR complex protein SEC13) (SEC13-like protein 1) (SEC13-related protein) | 0,42660791 | 0,02931 | 0.161086911482475 |
| Q9UGP8 | SEC63 | Translocation protein SEC63 homolog | 0,398122247 | 0,04977 | 0.204754344144234 |
| Q8IZQ5 | SELH | Selenoprotein H (SelH) | 0,86277784 | 0,0022 | 0.0729440600880325 |
| Q99985 | SEM3C | Semaphorin-3C (Semaphorin-E) (Sema E) | 0,899988924 | 0,00897 | 0.103994216063343 |
| Q96LD8 | SENP8 | Sentrin-specific protease 8 (EC 3.4.22.-) (Deneddylase-1) (NEDD8-specific protease 1) (Protease, cysteine 2) (Sentrin/SUMO-specific protease SENP8) | -0,571998068 | 0,0326 | 0.169804057839623 |
| P49908 | SEPP1 | Selenoprotein P (SeP) | 0,865525713 | 0,01096 | 0.110439860103028 |
| O43175 | SERA | D-3-phosphoglycerate dehydrogenase (3-PGDH) (EC 1.1.1.95) (2-oxoglutarate reductase) (EC 1.1.1.399) (Malate dehydrogenase) (EC 1.1.1.37) | 0,967287049 | 0,03092 | 0.165214630545001 |
| P50454 | SERPH | Serpin H1 (47 kDa heat shock protein) (Arsenic-transactivated protein 3) (AsTP3) (Cell proliferation-inducing gene 14 protein) (Collagen-binding protein) (Colligin) (Rheumatoid arthritis-related antigen RA-A47) | 1,202870175 | 0,00069 | 0.0461437135818825 |
| Q01105 | SET | Protein SET (HLA-DR-associated protein II) (Inhibitor of granzyme A-activated DNase) (IGAAD) (PHAPII) (Phosphatase 2A inhibitor I2PP2A) (I-2PP2A) (Template-activating factor I) (TAF-I) | 0,416542918 | 0,04366 | 0.190482327798759 |
| Q13435 | SF3B2 | Splicing factor 3B subunit 2 (Pre-mRNA-splicing factor SF3b 145 kDa subunit) (SF3b145) (Spliceosome-associated protein 145) (SAP 145) | 0,448231434 | 0,01898 | 0.135332764765101 |
| Q15393 | SF3B3 | Splicing factor 3B subunit 3 (Pre-mRNA-splicing factor SF3b 130 kDa subunit) (SF3b130) (STAF130) (Spliceosome-associated protein 130) (SAP 130) | 0,476447141 | 0,02374 | 0.147884574150362 |
| Q9Y3B4 | SF3B6 | Splicing factor 3B subunit 6 (Pre-mRNA branch site protein p14) (SF3b 14 kDa subunit) (SF3B14a) (Spliceosome-associated protein, 14-kDa) (Splicing factor 3b, subunit 6, 14kDa) | 0,642226826 | 0,02568 | 0.152888992161152 |
| P23246 | SFPQ | Splicing factor, proline- and glutamine-rich (100 kDa DNA-pairing protein) (hPOMp100) (DNA-binding p52/p100 complex, 100 kDa subunit) (Polypyrimidine tract-binding protein-associated-splicing factor) (PSF) (PTB-associated-splicing factor) | 0,517931704 | 0,01746 | 0.13109533821508 |
| Q16586 | SGCA | Alpha-sarcoglycan (Alpha-SG) (50 kDa dystrophin-associated glycoprotein) (50DAG) (Adhalin) (Dystroglycan-2) | -0,349538919 | 0,04542 | 0.194688736896108 |
| Q99720 | SGMR1 | Sigma non-opioid intracellular receptor 1 (Aging-associated gene 8 protein) (SR31747-binding protein) (SR-BP) (Sigma 1-type opioid receptor) (SIG-1R) (Sigma1-receptor) (Sigma1R) (hSigmaR1) | 0,613271356 | 0,04357 | 0.190307153040718 |
| Q9H299 | SH3L3 | SH3 domain-binding glutamic acid-rich-like protein 3 (SH3 domain-binding protein 1) (SH3BP-1) | 0,839076025 | 0,0025 | 0.073956300294973 |
| Q9UQ13 | SHOC2 | Leucine-rich repeat protein SHOC-2 (Protein soc-2 homolog) (Protein sur-8 homolog) | 0,524795132 | 0,0323 | 0.168698420035416 |
| Q96FS4 | SIPA1 | Signal-induced proliferation-associated protein 1 (Sipa-1) (GTPase-activating protein Spa-1) (p130 SPA-1) | 0,754490199 | 0,00325 | 0.0761554511764352 |
| Q15796 | SMAD2 | Mothers against decapentaplegic homolog 2 (MAD homolog 2) (Mothers against DPP homolog 2) (JV18-1) (Mad-related protein 2) (hMAD-2) (SMAD family member 2) (SMAD 2) (Smad2) (hSMAD2) | 0,462024154 | 0,02302 | 0.146880219728601 |
| Q14683 | SMC1A | Structural maintenance of chromosomes protein 1A (SMC protein 1A) (SMC-1-alpha) (SMC-1A) (Sb1.8) | 0,494284682 | 0,01364 | 0.11913827875583 |
| Q9UQE7 | SMC3 | Structural maintenance of chromosomes protein 3 (SMC protein 3) (SMC-3) (Basement membrane-associated chondroitin proteoglycan) (Bamacan) (Chondroitin sulfate proteoglycan 6) (Chromosome-associated polypeptide) (hCAP) | 0,349882296 | 0,04718 | 0.198769565715963 |
| P51532 | SMCA4 | Transcription activator BRG1 (EC 3.6.4.-) (ATP-dependent helicase SMARCA4) (BRG1-associated factor 190A) (BAF190A) (Mitotic growth and transcription activator) (Protein BRG-1) (Protein brahma homolog 1) (SNF2-beta) (SWI/SNF-related matrix-associated actin-dependent regulator of chromatin subfamily A member 4) | 0,533846738 | 0,02797 | 0.160816207866737 |
| P62314 | SMD1 | Small nuclear ribonucleoprotein Sm D1 (Sm-D1) (Sm-D autoantigen) (snRNP core protein D1) | 0,808653633 | 0,01527 | 0.124447705141253 |
| P62318 | SMD3 | Small nuclear ribonucleoprotein Sm D3 (Sm-D3) (snRNP core protein D3) | 0,730659282 | 0,00563 | 0.0943336318850801 |
| A6NHR9 | SMHD1 | Structural maintenance of chromosomes flexible hinge domain-containing protein 1 (SMC hinge domain-containing protein 1) | 1,279178065 | 0,00851 | 0.103753456880926 |
| Q8TAQ2 | SMRC2 | SWI/SNF complex subunit SMARCC2 (BRG1-associated factor 170) (BAF170) (SWI/SNF complex 170 kDa subunit) (SWI/SNF-related matrix-associated actin-dependent regulator of chromatin subfamily C member 2) | 0,481895968 | 0,03151 | 0.166991253015236 |
| Q8NB12 | SMYD1 | Histone-lysine N-methyltransferase SMYD1 (EC 2.1.1.43) (SET and MYND domain-containing protein 1) | -0,543288623 | 0,01803 | 0.132658871887263 |
| Q6GMV2 | SMYD5 | SET and MYND domain-containing protein 5 (EC 2.1.1.-) (Protein NN8-4AG) (Retinoic acid-induced protein 15) | -0,533941331 | 0,01425 | 0.120674495674537 |
| Q53GS9 | SNUT2 | U4/U6.U5 tri-snRNP-associated protein 2 (Inactive ubiquitin-specific peptidase 39) (SAD1 homolog) (U4/U6.U5 tri-snRNP-associated 65 kDa protein) (65K) | 0,788440659 | 0,00803 | 0.102154997647559 |
| Q13573 | SNW1 | SNW domain-containing protein 1 (Nuclear protein SkiP) (Nuclear receptor coactivator NCoA-62) (Ski-interacting protein) | 0,43053501 | 0,03244 | 0.169193971121812 |
| Q9UMY4 | SNX12 | Sorting nexin-12 | -0,445833569 | 0,0275 | 0.159552134451148 |
| Q969T3 | SNX21 | Sorting nexin-21 (Sorting nexin L) (SNX-L) | 0,613560441 | 0,00691 | 0.0978049310072672 |
| Q96L92 | SNX27 | Sorting nexin-27 | 0,386668455 | 0,03787 | 0.180046072726016 |
| P08294 | SODE | Extracellular superoxide dismutase [Cu-Zn] (EC-SOD) (EC 1.15.1.1) | 0,95940971 | 0,00123 | 0.0596963516380526 |
| P18583 | SON | Protein SON (Bax antagonist selected in saccharomyces 1) (BASS1) (Negative regulatory element-binding protein) (NRE-binding protein) (Protein DBP-5) (SON3) | 0,461800211 | 0,01708 | 0.13029397549501 |
| Q9Y5B9 | SP16H | FACT complex subunit SPT16 (Chromatin-specific transcription elongation factor 140 kDa subunit) (FACT 140 kDa subunit) (FACTp140) (Facilitates chromatin transcription complex subunit SPT16) (hSPT16) | 0,712799071 | 0,00415 | 0.0790778462733819 |
| Q96P63 | SPB12 | Serpin B12 | 0,926394778 | 0,04684 | 0.198769565715963 |
| P19623 | SPEE | Spermidine synthase (SPDSY) (EC 2.5.1.16) (Putrescine aminopropyltransferase) | 0,692092038 | 0,03852 | 0.1811476445859 |
| O75934 | SPF27 | Pre-mRNA-splicing factor SPF27 (Breast carcinoma-amplified sequence 2) (DNA amplified in mammary carcinoma 1 protein) (Spliceosome-associated protein SPF 27) | 0,387376489 | 0,03306 | 0.171282532397201 |
| Q9HCB6 | SPON1 | Spondin-1 (F-spondin) (Vascular smooth muscle cell growth-promoting factor) | 0,69533837 | 0,0445 | 0.192406103238607 |
| Q8WW59 | SPRY4 | SPRY domain-containing protein 4 | -0,56104242 | 0,00947 | 0.106455395758856 |
| P02549 | SPTA1 | Spectrin alpha chain, erythrocytic 1 (Erythroid alpha-spectrin) | 2,168136697 | 3,8E-05 | 0.0300468839677373 |
| P11277 | SPTB1 | Spectrin beta chain, erythrocytic (Beta-I spectrin) | 0,484935044 | 0,02129 | 0.141512996692124 |
| O15269 | SPTC1 | Serine palmitoyltransferase 1 (EC 2.3.1.50) (Long chain base biosynthesis protein 1) (LCB 1) (Serine-palmitoyl-CoA transferase 1) (SPT 1) (SPT1) | 0,482005146 | 0,02898 | 0.161086911482475 |
| O15042 | SR140 | U2 snRNP-associated SURP motif-containing protein (140 kDa Ser/Arg-rich domain protein) (U2-associated protein SR140) | 0,923368215 | 0,0021 | 0.0720954147183514 |
| Q14247 | SRC8 | Src substrate cortactin (Amplaxin) (Oncogene EMS1) | -0,384582291 | 0,03848 | 0.1811476445859 |
| P08240 | SRPRA | Signal recognition particle receptor subunit alpha (SR-alpha) (Docking protein alpha) (DP-alpha) | 0,428868891 | 0,02314 | 0.146880219728601 |
| P78539 | SRPX | Sushi repeat-containing protein SRPX | 0,880494483 | 0,00852 | 0.103753456880926 |
| Q9UQ35 | SRRM2 | Serine/arginine repetitive matrix protein 2 (300 kDa nuclear matrix antigen) (Serine/arginine-rich splicing factor-related nuclear matrix protein of 300 kDa) (SR-related nuclear matrix protein of 300 kDa) (Ser/Arg-related nuclear matrix protein of 300 kDa) (Splicing coactivator subunit SRm300) (Tax-responsive enhancer element-binding protein 803) (TaxREB803) | 0,598730527 | 0,01334 | 0.118076797909896 |
| Q9BXP5 | SRRT | Serrate RNA effector molecule homolog (Arsenite-resistance protein 2) | 0,45134231 | 0,02809 | 0.160816207866737 |
| O75494 | SRS10 | Serine/arginine-rich splicing factor 10 (40 kDa SR-repressor protein) (SRrp40) (FUS-interacting serine-arginine-rich protein 1) (Splicing factor SRp38) (Splicing factor, arginine/serine-rich 13A) (TLS-associated protein with Ser-Arg repeats) (TASR) (TLS-associated protein with SR repeats) (TLS-associated serine-arginine protein) (TLS-associated SR protein) | 0,754492686 | 0,03648 | 0.177612813348651 |
| Q16629 | SRSF7 | Serine/arginine-rich splicing factor 7 (Splicing factor 9G8) (Splicing factor, arginine/serine-rich 7) | 0,568247729 | 0,04032 | 0.183872362438851 |
| Q08945 | SSRP1 | FACT complex subunit SSRP1 (Chromatin-specific transcription elongation factor 80 kDa subunit) (Facilitates chromatin transcription complex 80 kDa subunit) (FACT 80 kDa subunit) (FACTp80) (Facilitates chromatin transcription complex subunit SSRP1) (Recombination signal sequence recognition protein 1) (Structure-specific recognition protein 1) (hSSRP1) (T160) | 0,588415314 | 0,01753 | 0.131227758234791 |
| Q9NY15 | STAB1 | Stabilin-1 (Fasciclin, EGF-like, laminin-type EGF-like and link domain-containing scavenger receptor 1) (FEEL-1) (MS-1 antigen) | 0,530450252 | 0,04618 | 0.196421194110723 |
| P42226 | STAT6 | Signal transducer and activator of transcription 6 (IL-4 Stat) | 0,555432428 | 0,04105 | 0.185884549229643 |
| O94804 | STK10 | Serine/threonine-protein kinase 10 (EC 2.7.11.1) (Lymphocyte-oriented kinase) | 0,556914428 | 0,01154 | 0.111960961329472 |
| Q9UJZ1 | STML2 | Stomatin-like protein 2, mitochondrial (SLP-2) (EPB72-like protein 2) (Paraprotein target 7) (Paratarg-7) | -0,59462769 | 0,00644 | 0.0954651715986055 |
| Q86Y82 | STX12 | Syntaxin-12 | 0,468116109 | 0,01829 | 0.133184461911317 |
| O14662 | STX16 | Syntaxin-16 (Syn16) | 0,684552153 | 0,03833 | 0.180923549477534 |
| Q8IX01 | SUGP2 | SURP and G-patch domain-containing protein 2 (Arginine/serine-rich-splicing factor 14) (Splicing factor, arginine/serine-rich 14) | 0,639371316 | 0,00877 | 0.103882752534462 |
| Q9UH99 | SUN2 | SUN domain-containing protein 2 (Protein unc-84 homolog B) (Rab5-interacting protein) (Rab5IP) (Sad1/unc-84 protein-like 2) | 1,160414849 | 0,00076 | 0.0473327203168651 |
| Q5JTZ9 | SYAM | Alanine--tRNA ligase, mitochondrial (EC 6.1.1.7) (Alanyl-tRNA synthetase) (AlaRS) | 0,480370945 | 0,02259 | 0.145950138465754 |
| Q96A49 | SYAP1 | Synapse-associated protein 1 (BSD domain-containing signal transducer and Akt interactor protein) (BSTA) | -0,53384609 | 0,0248 | 0.150350167155027 |
| P14868 | SYDC | Aspartate--tRNA ligase, cytoplasmic (EC 6.1.1.12) (Aspartyl-tRNA synthetase) (AspRS) (Cell proliferation-inducing gene 40 protein) | 0,432944087 | 0,04953 | 0.204280163930216 |
| Q9UMS6 | SYNP2 | Synaptopodin-2 (Genethonin-2) (Myopodin) | -0,535789601 | 0,01045 | 0.109336835224251 |
| Q9BW92 | SYTM | Threonine--tRNA ligase, mitochondrial (EC 6.1.1.3) (Threonyl-tRNA synthetase) (ThrRS) (Threonyl-tRNA synthetase-like 1) | 0,358754258 | 0,04252 | 0.188876486048564 |
| P37840 | SYUA | Alpha-synuclein (Non-A beta component of AD amyloid) (Non-A4 component of amyloid precursor) (NACP) | 0,923272175 | 0,00781 | 0.102154997647559 |
| Q9H061 | T126A | Transmembrane protein 126A | -0,555308229 | 0,03311 | 0.171292325765253 |
| Q96EM0 | T3HPD | Trans-3-hydroxy-L-proline dehydratase (EC 4.2.1.77) (Trans-L-3-hydroxyproline dehydratase) | 0,851369811 | 0,02822 | 0.160816207866737 |
| Q15750 | TAB1 | TGF-beta-activated kinase 1 and MAP3K7-binding protein 1 (Mitogen-activated protein kinase kinase kinase 7-interacting protein 1) (TGF-beta-activated kinase 1-binding protein 1) (TAK1-binding protein 1) | 0,782069879 | 0,00352 | 0.0767928025552317 |
| P37802 | TAGL2 | Transgelin-2 (Epididymis tissue protein Li 7e) (SM22-alpha homolog) | 0,638919813 | 0,01806 | 0.132658871887263 |
| Q9H2K8 | TAOK3 | Serine/threonine-protein kinase TAO3 (EC 2.7.11.1) (Cutaneous T-cell lymphoma-associated antigen HD-CL-09) (CTCL-associated antigen HD-CL-09) (Dendritic cell-derived protein kinase) (JNK/SAPK-inhibitory kinase) (Jun kinase-inhibitory kinase) (Kinase from chicken homolog A) (hKFC-A) (Thousand and one amino acid protein 3) | 0,865857691 | 0,00338 | 0.0764488368109507 |
| Q7Z7G0 | TARSH | Target of Nesh-SH3 (Tarsh) (ABI gene family member 3-binding protein) (Nesh-binding protein) (NeshBP) | 1,108382931 | 0,00364 | 0.0767928025552317 |
| Q6P1N9 | TATD1 | Putative deoxyribonuclease TATDN1 (EC 3.1.21.-) (Hepatocarcinoma high expression protein) | -0,602756347 | 0,04031 | 0.183872362438851 |
| P07437 | TBB5 | Tubulin beta chain (Tubulin beta-5 chain) | 0,48193474 | 0,04571 | 0.195289951852626 |
| Q9BUF5 | TBB6 | Tubulin beta-6 chain (Tubulin beta class V) | 0,689840307 | 0,04812 | 0.200742464336218 |
| Q9NVG8 | TBC13 | TBC1 domain family member 13 | 0,448156264 | 0,03651 | 0.177612813348651 |
| Q66K14 | TBC9B | TBC1 domain family member 9B | 0,430194324 | 0,02801 | 0.160816207866737 |
| Q9BTW9 | TBCD | Tubulin-specific chaperone D (Beta-tubulin cofactor D) (tfcD) (SSD-1) (Tubulin-folding cofactor D) | 0,419184477 | 0,03696 | 0.178718378708741 |
| Q9Y4P3 | TBL2 | Transducin beta-like protein 2 (WS beta-transducin repeats protein) (WS-betaTRP) (Williams-Beuren syndrome chromosomal region 13 protein) | 0,578748136 | 0,02289 | 0.146471517358329 |
| Q13428 | TCOF | Treacle protein (Treacher Collins syndrome protein) | 0,930049416 | 0,00678 | 0.096987147592169 |
| O14776 | TCRG1 | Transcription elongation regulator 1 (TATA box-binding protein-associated factor 2S) (Transcription factor CA150) | 0,761764298 | 0,00234 | 0.073956300294973 |
| Q9NZ01 | TECR | Very-long-chain enoyl-CoA reductase (EC 1.3.1.93) (Synaptic glycoprotein SC2) (Trans-2,3-enoyl-CoA reductase) (TER) | 0,529290125 | 0,02064 | 0.139808301680561 |
| O15273 | TELT | Telethonin (Titin cap protein) | -0,776732572 | 0,00871 | 0.103882752534462 |
| P22105 | TENX | Tenascin-X (TN-X) (Hexabrachion-like protein) | 1,091784954 | 0,00486 | 0.0864075069907863 |
| P05452 | TETN | Tetranectin (TN) (C-type lectin domain family 3 member B) (Plasminogen kringle 4-binding protein) | 1,457869254 | 0,00827 | 0.102945529755352 |
| P52655 | TF2AA | Transcription initiation factor IIA subunit 1 (General transcription factor IIA subunit 1) (TFIIAL) (Transcription initiation factor TFIIA 42 kDa subunit) (TFIIA-42) [Cleaved into: Transcription initiation factor IIA alpha chain (TFIIA p35 subunit); Transcription initiation factor IIA beta chain (TFIIA p19 subunit)] | 0,43378734 | 0,02434 | 0.149305815497977 |
| Q9UKN8 | TF3C4 | General transcription factor 3C polypeptide 4 (EC 2.3.1.48) (TF3C-delta) (Transcription factor IIIC 90 kDa subunit) (TFIIIC 90 kDa subunit) (TFIIIC90) (Transcription factor IIIC subunit delta) | 0,661624529 | 0,00562 | 0.0943336318850801 |
| P42765 | THIM | 3-ketoacyl-CoA thiolase, mitochondrial (EC 2.3.1.16) (Acetyl-CoA acyltransferase) (Beta-ketothiolase) (Mitochondrial 3-oxoacyl-CoA thiolase) (T1) | -0,629440259 | 0,01416 | 0.120558249672262 |
| P00734 | THRB | Prothrombin (EC 3.4.21.5) (Coagulation factor II) [Cleaved into: Activation peptide fragment 1; Activation peptide fragment 2; Thrombin light chain; Thrombin heavy chain] | 2,056988092 | 0,00064 | 0.0460405438333303 |
| Q6ZMP0 | THSD4 | Thrombospondin type-1 domain-containing protein 4 (A disintegrin and metalloproteinase with thrombospondin motifs-like protein 6) (ADAMTS-like protein 6) (ADAMTSL-6) | 0,852524109 | 0,04662 | 0.198114702396706 |
| P25325 | THTM | 3-mercaptopyruvate sulfurtransferase (MST) (EC 2.8.1.2) | 0,586036756 | 0,03324 | 0.171297602744958 |
| Q13263 | TIF1B | Transcription intermediary factor 1-beta (TIF1-beta) (E3 SUMO-protein ligase TRIM28) (EC 2.3.2.27) (KRAB-associated protein 1) (KAP-1) (KRAB-interacting protein 1) (KRIP-1) (Nuclear corepressor KAP-1) (RING finger protein 96) (RING-type E3 ubiquitin transferase TIF1-beta) (Tripartite motif-containing protein 28) | 0,465285566 | 0,03202 | 0.168322690360001 |
| Q9Y5J7 | TIM9 | Mitochondrial import inner membrane translocase subunit Tim9 | -0,523360688 | 0,01331 | 0.11806561221821 |
| O75663 | TIPRL | TIP41-like protein (Putative MAPK-activating protein PM10) (Type 2A-interacting protein) (TIP) | -0,589643777 | 0,01499 | 0.123290954242387 |
| Q8WZ42 | TITIN | Titin (EC 2.7.11.1) (Connectin) (Rhabdomyosarcoma antigen MU-RMS-40.14) | -0,395456955 | 0,03666 | 0.177966828074799 |
| Q3LXA3 | TKFC | Triokinase/FMN cyclase (Bifunctional ATP-dependent dihydroxyacetone kinase/FAD-AMP lyase (cyclizing)) [Includes: ATP-dependent dihydroxyacetone kinase (DHA kinase) (EC 2.7.1.28) (EC 2.7.1.29) (Glycerone kinase) (Triokinase) (Triose kinase); FAD-AMP lyase (cyclizing) (EC 4.6.1.15) (FAD-AMP lyase (cyclic FMN forming)) (FMN cyclase)] | 0,523783527 | 0,04073 | 0.184903170886882 |
| Q9BVC6 | TM109 | Transmembrane protein 109 (Mitsugumin-23) (Mg23) | 0,621869613 | 0,00855 | 0.103835198154595 |
| Q4V9L6 | TM119 | Transmembrane protein 119 (Osteoblast induction factor) (OBIF) | 1,368202838 | 0,02509 | 0.151159838745206 |
| Q96I45 | TM141 | Transmembrane protein 141 | 0,369670507 | 0,04791 | 0.200113362152407 |
| A0PJW6 | TM223 | Transmembrane protein 223 | -0,652033748 | 0,01174 | 0.112208078457086 |
| Q8WUH6 | TM263 | Transmembrane protein 263 | 0,456046254 | 0,02853 | 0.160816207866737 |
| Q969M1 | TM40L | Mitochondrial import receptor subunit TOM40B (Protein TOMM40-like) | -0,809884176 | 0,00717 | 0.0993450268446226 |
| Q96HV5 | TM41A | Transmembrane protein 41A | 0,576936899 | 0,01089 | 0.110439860103028 |
| Q13445 | TMED1 | Transmembrane emp24 domain-containing protein 1 (Interleukin-1 receptor-like 1 ligand) (Putative T1/ST2 receptor-binding protein) (p24 family protein gamma-1) (Tp24) (p24gamma1) | 0,508262762 | 0,02487 | 0.150535290153693 |
| Q15363 | TMED2 | Transmembrane emp24 domain-containing protein 2 (Membrane protein p24A) (p24) (p24 family protein beta-1) (p24beta1) | 0,577356562 | 0,01729 | 0.13109533821508 |
| Q96MV1 | TMM56 | Transmembrane protein 56 | -0,484034698 | 0,02925 | 0.161086911482475 |
| Q9BUB7 | TMM70 | Transmembrane protein 70, mitochondrial | -0,372796341 | 0,0417 | 0.187538691516333 |
| Q9H3N1 | TMX1 | Thioredoxin-related transmembrane protein 1 (Thioredoxin domain-containing protein 1) (Transmembrane Trx-related protein) | 0,880153583 | 0,00989 | 0.109011590445672 |
| P63316 | TNNC1 | Troponin C, slow skeletal and cardiac muscles (TN-C) | -0,510336126 | 0,04495 | 0.193722382532051 |
| Q15785 | TOM34 | Mitochondrial import receptor subunit TOM34 (hTom34) (Translocase of outer membrane 34 kDa subunit) | 0,779522119 | 0,00463 | 0.0848101324768568 |
| O96008 | TOM40 | Mitochondrial import receptor subunit TOM40 homolog (Protein Haymaker) (Translocase of outer membrane 40 kDa subunit homolog) (p38.5) | -0,584414019 | 0,03676 | 0.177966828074799 |
| P11387 | TOP1 | DNA topoisomerase 1 (EC 5.99.1.2) (DNA topoisomerase I) | 0,588928324 | 0,01539 | 0.124751718030612 |
| Q02880 | TOP2B | DNA topoisomerase 2-beta (EC 5.99.1.3) (DNA topoisomerase II, beta isozyme) | 0,892028798 | 0,00148 | 0.0658126006797819 |
| A5PLN9 | TPC13 | Trafficking protein particle complex subunit 13 | 0,513862001 | 0,01689 | 0.130074948166753 |
| P55327 | TPD52 | Tumor protein D52 (Protein N8) | -0,561449071 | 0,03119 | 0.166181718381279 |
| P07951 | TPM2 | Tropomyosin beta chain (Beta-tropomyosin) (Tropomyosin-2) | -0,610559414 | 0,03578 | 0.176070195019166 |
| P67936 | TPM4 | Tropomyosin alpha-4 chain (TM30p1) (Tropomyosin-4) | 0,643620873 | 0,01032 | 0.109336835224251 |
| Q8IUR0 | TPPC5 | Trafficking protein particle complex subunit 5 | 0,559908116 | 0,01122 | 0.111167464928122 |
| Q9Y2L5 | TPPC8 | Trafficking protein particle complex subunit 8 (Protein TRS85 homolog) | 0,451041181 | 0,02098 | 0.140961498869228 |
| O94811 | TPPP | Tubulin polymerization-promoting protein (TPPP) (25 kDa brain-specific protein) (TPPP/p25) (p24) (p25-alpha) | 0,43662302 | 0,041 | 0.185884549229643 |
| Q9BW30 | TPPP3 | Tubulin polymerization-promoting protein family member 3 (TPPP/p20) | 0,767340532 | 0,00241 | 0.073956300294973 |
| O15533 | TPSN | Tapasin (TPN) (TPSN) (NGS-17) (TAP-associated protein) (TAP-binding protein) | 0,538310623 | 0,02569 | 0.152888992161152 |
| P62995 | TRA2B | Transformer-2 protein homolog beta (TRA-2 beta) (TRA2-beta) (hTRA2-beta) (Splicing factor, arginine/serine-rich 10) (Transformer-2 protein homolog B) | 0,775951241 | 0,01603 | 0.12714580081316 |
| Q9UPV9 | TRAK1 | Trafficking kinesin-binding protein 1 (106 kDa O-GlcNAc transferase-interacting protein) | 0,511383791 | 0,01614 | 0.12764681059851 |
| P02787 | TRFE | Serotransferrin (Transferrin) (Beta-1 metal-binding globulin) (Siderophilin) | 2,158823126 | 1,8E-05 | 0.0300468839677373 |
| P02788 | TRFL | Lactotransferrin (Lactoferrin) (EC 3.4.21.-) (Growth-inhibiting protein 12) (Talalactoferrin) [Cleaved into: Lactoferricin-H (Lfcin-H); Kaliocin-1; Lactoferroxin-A; Lactoferroxin-B; Lactoferroxin-C] | 2,383947222 | 0,00077 | 0.0474502355471915 |
| O75382 | TRIM3 | Tripartite motif-containing protein 3 (Brain-expressed RING finger protein) (RING finger protein 22) (RING finger protein 97) | 0,673715632 | 0,00415 | 0.0790778462733819 |
| Q15643 | TRIPB | Thyroid receptor-interacting protein 11 (TR-interacting protein 11) (TRIP-11) (Clonal evolution-related gene on chromosome 14 protein) (Golgi-associated microtubule-binding protein 210) (GMAP-210) (Trip230) | 0,796087867 | 0,00252 | 0.073956300294973 |
| Q9Y5S1 | TRPV2 | Transient receptor potential cation channel subfamily V member 2 (TrpV2) (Osm-9-like TRP channel 2) (OTRPC2) (Vanilloid receptor-like protein 1) (VRL-1) | 0,628519757 | 0,0066 | 0.0958235018254441 |
| Q8WWH5 | TRUB1 | Probable tRNA pseudouridine synthase 1 (EC 5.4.99.-) | 0,772166365 | 0,00313 | 0.0761554511764352 |
| P07996 | TSP1 | Thrombospondin-1 (Glycoprotein G) | 1,43610081 | 0,00168 | 0.0673661906302801 |
| P35443 | TSP4 | Thrombospondin-4 | 1,722336259 | 0,00183 | 0.0697416810103427 |
| Q5SRH9 | TT39A | Tetratricopeptide repeat protein 39A (TPR repeat protein 39A) (Differentially expressed in MCF7 with estradiol protein 6) (DEME-6) | -0,380102713 | 0,0424 | 0.188743139597737 |
| Q6DKK2 | TTC19 | Tetratricopeptide repeat protein 19, mitochondrial (TPR repeat protein 19) | -0,433656929 | 0,02783 | 0.160530825210271 |
| Q86TV6 | TTC7B | Tetratricopeptide repeat protein 7B (TPR repeat protein 7B) (Tetratricopeptide repeat protein 7-like-1) (TPR repeat protein 7-like-1) | 0,394208773 | 0,04288 | 0.188876486048564 |
| Q12792 | TWF1 | Twinfilin-1 (Protein A6) (Protein tyrosine kinase 9) | 0,541408548 | 0,01017 | 0.109336835224251 |
| Q8NBS9 | TXND5 | Thioredoxin domain-containing protein 5 (Endoplasmic reticulum resident protein 46) (ER protein 46) (ERp46) (Thioredoxin-like protein p46) | 0,441413026 | 0,04022 | 0.18383048391046 |
| Q9H3M7 | TXNIP | Thioredoxin-interacting protein (Thioredoxin-binding protein 2) (Vitamin D3 up-regulated protein 1) | -0,586161819 | 0,04182 | 0.187870082281016 |
| P26368 | U2AF2 | Splicing factor U2AF 65 kDa subunit (U2 auxiliary factor 65 kDa subunit) (hU2AF(65)) (hU2AF65) (U2 snRNP auxiliary factor large subunit) | 0,485725179 | 0,02317 | 0.146880219728601 |
| P0DN76 | U2AF5 | Splicing factor U2AF 35 kDa subunit-like protein (U2 small nuclear RNA auxiliary factor 1-like protein 5) | 0,776783384 | 0,0092 | 0.104035630640557 |
| O75643 | U520 | U5 small nuclear ribonucleoprotein 200 kDa helicase (EC 3.6.4.13) (Activating signal cointegrator 1 complex subunit 3-like 1) (BRR2 homolog) (U5 snRNP-specific 200 kDa protein) (U5-200KD) | 0,505751314 | 0,01441 | 0.121811791788254 |
| P61088 | UBE2N | Ubiquitin-conjugating enzyme E2 N (EC 2.3.2.23) (Bendless-like ubiquitin-conjugating enzyme) (E2 ubiquitin-conjugating enzyme N) (Ubc13) (UbcH13) (Ubiquitin carrier protein N) (Ubiquitin-protein ligase N) | -0,466689329 | 0,03887 | 0.182368104742193 |
| P17480 | UBF1 | Nucleolar transcription factor 1 (Autoantigen NOR-90) (Upstream-binding factor 1) (UBF-1) | 0,628872788 | 0,00562 | 0.0943336318850801 |
| P68543 | UBX2A | UBX domain-containing protein 2A (UBX domain-containing protein 4) | 0,955516587 | 0,01207 | 0.113024663824624 |
| P15374 | UCHL3 | Ubiquitin carboxyl-terminal hydrolase isozyme L3 (UCH-L3) (EC 3.4.19.12) (Ubiquitin thioesterase L3) | -0,346975214 | 0,0472 | 0.198769565715963 |
| Q8IX04 | UEVLD | Ubiquitin-conjugating enzyme E2 variant 3 (UEV-3) (EV and lactate/malate dehydrogenase domain-containing protein) | 0,445356212 | 0,01986 | 0.137802712336179 |
| O60701 | UGDH | UDP-glucose 6-dehydrogenase (UDP-Glc dehydrogenase) (UDP-GlcDH) (UDPGDH) (EC 1.1.1.22) | 1,053589163 | 0,00962 | 0.106870225730999 |
| Q96QD9 | UIF | UAP56-interacting factor (Forty-two-three domain-containing protein 1) (Protein 40-2-3) | 0,405228478 | 0,03573 | 0.176070195019166 |
| P11172 | UMPS | Uridine 5'-monophosphate synthase (UMP synthase) [Includes: Orotate phosphoribosyltransferase (OPRT) (OPRTase) (EC 2.4.2.10); Orotidine 5'-phosphate decarboxylase (ODC) (EC 4.1.1.23) (OMPdecase)] | 0,745003579 | 0,006 | 0.0948090470651261 |
| Q9NVA1 | UQCC1 | Ubiquinol-cytochrome-c reductase complex assembly factor 1 (Basic FGF-repressed Zic-binding protein) (bFGF-repressed Zic-binding protein) (bFZb) (Ubiquinol-cytochrome c reductase complex chaperone CBP3 homolog) | -0,636792481 | 0,02335 | 0.146880219728601 |
| Q9BRT2 | UQCC2 | Ubiquinol-cytochrome-c reductase complex assembly factor 2 (Breast cancer-associated protein SGA-81M) (Mitochondrial nucleoid factor 1) (Mitochondrial protein M19) | -0,500269467 | 0,01498 | 0.123290954242387 |
| Q86UX7 | URP2 | Fermitin family homolog 3 (Kindlin-3) (MIG2-like protein) (Unc-112-related protein 2) | 1,325487791 | 0,0295 | 0.161662882550704 |
| Q08AM6 | VAC14 | Protein VAC14 homolog (Tax1-binding protein 2) | 0,352014713 | 0,04716 | 0.198769565715963 |
| P51809 | VAMP7 | Vesicle-associated membrane protein 7 (VAMP-7) (Synaptobrevin-like protein 1) (Tetanus-insensitive VAMP) (Ti-VAMP) | 0,751263528 | 0,00231 | 0.073956300294973 |
| Q99536 | VAT1 | Synaptic vesicle membrane protein VAT-1 homolog (EC 1.-.-.-) | -0,462179891 | 0,03361 | 0.172214098674328 |
| P17948 | VGFR1 | Vascular endothelial growth factor receptor 1 (VEGFR-1) (EC 2.7.10.1) (Fms-like tyrosine kinase 1) (FLT-1) (Tyrosine-protein kinase FRT) (Tyrosine-protein kinase receptor FLT) (FLT) (Vascular permeability factor receptor) | 1,670935944 | 0,04565 | 0.195289951852626 |
| P08670 | VIME | Vimentin | 0,894254304 | 0,00824 | 0.102945529755352 |
| P38435 | VKGC | Vitamin K-dependent gamma-carboxylase (EC 4.1.1.90) (Gamma-glutamyl carboxylase) (Peptidyl-glutamate 4-carboxylase) (Vitamin K gamma glutamyl carboxylase) | 0,590887585 | 0,01239 | 0.114216522917893 |
| Q86VN1 | VPS36 | Vacuolar protein-sorting-associated protein 36 (ELL-associated protein of 45 kDa) (ESCRT-II complex subunit VPS36) | -0,411749985 | 0,03594 | 0.176070600102387 |
| O75351 | VPS4B | Vacuolar protein sorting-associated protein 4B (EC 3.6.4.6) (Cell migration-inducing gene 1 protein) (Suppressor of K(+) transport growth defect 1) (Protein SKD1) | 0,508934943 | 0,01483 | 0.12260574009143 |
| Q8N3P4 | VPS8 | Vacuolar protein sorting-associated protein 8 homolog | 0,829121498 | 0,01345 | 0.118307495449371 |
| P02774 | VTDB | Vitamin D-binding protein (DBP) (VDB) (Gc protein-derived macrophage activating factor) (Gc-MAF) (GcMAF) (Gc-globulin) (Group-specific component) (Gc) (Vitamin D-binding protein-macrophage activating factor) (DBP-maf) | 1,167808263 | 0,00199 | 0.0713274753495378 |
| P04004 | VTNC | Vitronectin (VN) (S-protein) (Serum-spreading factor) (V75) [Cleaved into: Vitronectin V65 subunit; Vitronectin V10 subunit; Somatomedin-B] | 1,551106717 | 0,01573 | 0.126145843622979 |
| Q9BV38 | WDR18 | WD repeat-containing protein 18 | 0,672427258 | 0,00368 | 0.0767928025552317 |
| Q9H7D7 | WDR26 | WD repeat-containing protein 26 (CUL4- and DDB1-associated WDR protein 2) (Myocardial ischemic preconditioning up-regulated protein 2) | 0,690796828 | 0,00384 | 0.0771477016366249 |
| Q9GZS3 | WDR61 | WD repeat-containing protein 61 (Meiotic recombination REC14 protein homolog) (SKI8 homolog) (Ski8) [Cleaved into: WD repeat-containing protein 61, N-terminally processed] | -0,416187134 | 0,038 | 0.180446151432194 |
| Q96S55 | WRIP1 | ATPase WRNIP1 (EC 3.6.1.3) (Werner helicase-interacting protein 1) | 1,158282187 | 0,00064 | 0.0460405438333303 |
| Q9UIA9 | XPO7 | Exportin-7 (Exp7) (Ran-binding protein 16) | 0,619652101 | 0,00829 | 0.102945529755352 |
| P13010 | XRCC5 | X-ray repair cross-complementing protein 5 (EC 3.6.4.-) (86 kDa subunit of Ku antigen) (ATP-dependent DNA helicase 2 subunit 2) (ATP-dependent DNA helicase II 80 kDa subunit) (CTC box-binding factor 85 kDa subunit) (CTC85) (CTCBF) (DNA repair protein XRCC5) (Ku80) (Ku86) (Lupus Ku autoantigen protein p86) (Nuclear factor IV) (Thyroid-lupus autoantigen) (TLAA) (X-ray repair complementing defective repair in Chinese hamster cells 5 (double-strand-break rejoining)) | 0,421526314 | 0,02976 | 0.162543965335756 |
| P12956 | XRCC6 | X-ray repair cross-complementing protein 6 (EC 3.6.4.-) (EC 4.2.99.-) (5'-deoxyribose-5-phosphate lyase Ku70) (5'-dRP lyase Ku70) (70 kDa subunit of Ku antigen) (ATP-dependent DNA helicase 2 subunit 1) (ATP-dependent DNA helicase II 70 kDa subunit) (CTC box-binding factor 75 kDa subunit) (CTC75) (CTCBF) (DNA repair protein XRCC6) (Lupus Ku autoantigen protein p70) (Ku70) (Thyroid-lupus autoantigen) (TLAA) (X-ray repair complementing defective repair in Chinese hamster cells 6) | 0,3949786 | 0,03655 | 0.177612813348651 |
| Q7Z2W4 | ZCCHV | Zinc finger CCCH-type antiviral protein 1 (ADP-ribosyltransferase diphtheria toxin-like 13) (ARTD13) (Zinc finger CCCH domain-containing protein 2) (Zinc finger antiviral protein) (ZAP) | 0,507874385 | 0,03906 | 0.183043764764278 |
| Q9UK55 | ZPI | Protein Z-dependent protease inhibitor (PZ-dependent protease inhibitor) (PZI) (Serpin A10) | 1,480536421 | 0,00055 | 0.0460405438333303 |

**Supplementary Data Table 9. List of significantly altered senescence-related proteins (p-value<0.05) in physiologically aged humans**

| **Uniprot ID** | **Protein Symbol** | **Protein Description** | **Average Log2 Ratio** | ***p*-value** | **Adjusted *p*-value** |
| --- | --- | --- | --- | --- | --- |
| Q9UDR5 | AASS | Alpha-aminoadipic semialdehyde synthase, mitochondrial (LKR/SDH) [Includes: Lysine ketoglutarate reductase (LKR) (LOR) (EC 1,5,1,8); Saccharopine dehydrogenase (SDH) (EC 1,5,1,9)] | -0,445354258 | 0,03904 | 0,299900797 |
| P02654 | APOC1 | Apolipoprotein C-I (Apo-CI) (ApoC-I) (Apolipoprotein C1) [Cleaved into: Truncated apolipoprotein C-I] | 1,114962027 | 0,00442 | 0,120465537 |
| Q13510 | ASAH1 | Acid ceramidase (AC) (ACDase) (Acid CDase) (EC 3,5,1,23) (Acylsphingosine deacylase) (N-acylsphingosine amidohydrolase) (Putative 32 kDa heart protein) (PHP32) [Cleaved into: Acid ceramidase subunit alpha; Acid ceramidase subunit beta] | 1,046278962 | 0,00068 | 0,045663466 |
| P16671 | CD36 | Platelet glycoprotein 4 (Fatty acid translocase) (FAT) (Glycoprotein IIIb) (GPIIIB) (Leukocyte differentiation antigen CD36) (PAS IV) (PAS-4) (Platelet collagen receptor) (Platelet glycoprotein IV) (GPIV) (Thrombospondin receptor) (CD antigen CD36) | 0,755066517 | 0,00276 | 0,097030859 |
| P60033 | CD81 | CD81 antigen (26 kDa cell surface protein TAPA-1) (Target of the antiproliferative antibody 1) (Tetraspanin-28) (Tspan-28) (CD antigen CD81) | 1,322755108 | 5,33E-05 | 0,01161673 |
| Q9UJS0 | CMC2 | Calcium-binding mitochondrial carrier protein Aralar2 (Citrin) (Mitochondrial aspartate glutamate carrier 2) (Solute carrier family 25 member 13) | -0,263263971 | 0,04202 | 0,311743911 |
| P98082 | DAB2 | Disabled homolog 2 (Adaptor molecule disabled-2) (Differentially expressed in ovarian carcinoma 2) (DOC-2) (Differentially-expressed protein 2) | 0,549359821 | 0,03538 | 0,284308771 |
| Q96C86 | DCPS | m7GpppX diphosphatase (EC 3,6,1,59) (DCS-1) (Decapping scavenger enzyme) (Hint-related 7meGMP-directed hydrolase) (Histidine triad nucleotide-binding protein 5) (Histidine triad protein member 5) (HINT-5) (Scavenger mRNA-decapping enzyme DcpS) | -0,396526265 | 0,00919 | 0,172373188 |
| Q9Y394 | DHRS7 | Dehydrogenase/reductase SDR family member 7 (EC 1,1,-,-) (Retinal short-chain dehydrogenase/reductase 4) (retSDR4) (Short chain dehydrogenase/reductase family 34C member 1) | 0,312856544 | 0,03734 | 0,292457436 |
| O60832 | DKC1 | H/ACA ribonucleoprotein complex subunit DKC1 (EC 5,4,99,-) (CBF5 homolog) (Dyskerin) (Nopp140-associated protein of 57 kDa) (Nucleolar protein NAP57) (Nucleolar protein family A member 4) (snoRNP protein DKC1) | 0,529885662 | 0,00485 | 0,126093358 |
| P49327 | FAS | Fatty acid synthase (EC 2,3,1,85) [Includes: [Acyl-carrier-protein] S-acetyltransferase (EC 2,3,1,38); [Acyl-carrier-protein] S-malonyltransferase (EC 2,3,1,39); 3-oxoacyl-[acyl-carrier-protein] synthase (EC 2,3,1,41); 3-oxoacyl-[acyl-carrier-protein] reductase (EC 1,1,1,100); 3-hydroxyacyl-[acyl-carrier-protein] dehydratase (EC 4,2,1,59); Enoyl-[acyl-carrier-protein] reductase (EC 1,3,1,39); Oleoyl-[acyl-carrier-protein] hydrolase (EC 3,1,2,14)] | 0,812383438 | 0,03249 | 0,270276513 |
| O75223 | GGCT | Gamma-glutamylcyclotransferase (EC 4,3,2,9) (Cytochrome c-releasing factor 21) | 1,115494726 | 1,88E-05 | 0,007375622 |
| Q92820 | GGH | Gamma-glutamyl hydrolase (EC 3,4,19,9) (Conjugase) (GH) (Gamma-Glu-X carboxypeptidase) | 0,697630237 | 0,00603 | 0,139953969 |
| P07686 | HEXB | Beta-hexosaminidase subunit beta (EC 3,2,1,52) (Beta-N-acetylhexosaminidase subunit beta) (Hexosaminidase subunit B) (Cervical cancer proto-oncogene 7 protein) (HCC-7) (N-acetyl-beta-glucosaminidase subunit beta) [Cleaved into: Beta-hexosaminidase subunit beta chain B; Beta-hexosaminidase subunit beta chain A] | 0,518884583 | 0,02479 | 0,241832734 |
| Q03252 | LMNB2 | Lamin-B2 | -0,372600036 | 0,02752 | 0,255168841 |
| Q9H3U5 | MFSD1 | Major facilitator superfamily domain-containing protein 1 (Smooth muscle cell-associated protein 4) (SMAP-4) | -0,386381836 | 0,02836 | 0,257481581 |
| P20591 | MX1 | Interferon-induced GTP-binding protein Mx1 (Interferon-induced protein p78) (IFI-78K) (Interferon-regulated resistance GTP-binding protein MxA) (Myxoma resistance protein 1) (Myxovirus resistance protein 1) [Cleaved into: Interferon-induced GTP-binding protein Mx1, N-terminally processed] | -1,421960024 | 0,00402 | 0,114007879 |
| P61916 | NPC2 | NPC intracellular cholesterol transporter 2 (Epididymal secretory protein E1) (Human epididymis-specific protein 1) (He1) (Niemann-Pick disease type C2 protein) | 1,36282824 | 5,21E-07 | 0,000680806 |
| Q96CV9 | OPTN | Optineurin (E3-14,7K-interacting protein) (FIP-2) (Huntingtin yeast partner L) (Huntingtin-interacting protein 7) (HIP-7) (Huntingtin-interacting protein L) (NEMO-related protein) (Optic neuropathy-inducing protein) (Transcription factor IIIA-interacting protein) (TFIIIA-IntP) | 0,320744163 | 0,03075 | 0,266195983 |
| O95613 | PCNT | Pericentrin (Kendrin) (Pericentrin-B) | -0,27198731 | 0,04199 | 0,311743911 |
| Q00765 | REEP5 | Receptor expression-enhancing protein 5 (Polyposis locus protein 1) (Protein TB2) | 0,46818852 | 0,02719 | 0,253562451 |
| P10301 | RRAS | Ras-related protein R-Ras (p23) | -0,367146002 | 0,02149 | 0,235752629 |
| Q9C0D5 | TANC1 | Protein TANC1 (Tetratricopeptide repeat, ankyrin repeat and coiled-coil domain-containing protein 1) | -0,383565768 | 0,04864 | 0,327613246 |
| P35625 | TIMP3 | Metalloproteinase inhibitor 3 (Protein MIG-5) (Tissue inhibitor of metalloproteinases 3) (TIMP-3) | 3,116116312 | 4,97E-05 | 0,011475286 |
| P09493 | TPM1 | Tropomyosin alpha-1 chain (Alpha-tropomyosin) (Tropomyosin-1) | 0,298130157 | 0,03917 | 0,300019036 |
| Q9H3M7 | TXNIP | Thioredoxin-interacting protein (Thioredoxin-binding protein 2) (Vitamin D3 up-regulated protein 1) | -0,984051523 | 0,00351 | 0,106830512 |
| P62256 | UBE2H | Ubiquitin-conjugating enzyme E2 H (EC 2,3,2,23) ((E3-independent) E2 ubiquitin-conjugating enzyme H) (EC 2,3,2,24) (E2 ubiquitin-conjugating enzyme H) (UbcH2) (Ubiquitin carrier protein H) (Ubiquitin-conjugating enzyme E2-20K) (Ubiquitin-protein ligase H) | 0,677933751 | 0,0001 | 0,014543495 |

**Supplementary Data Table 10. List of significantly altered senescence-related proteins (p-value<0.05) in HGPS human individuals**

| **Uniprot ID** | **Protein Symbol** | **Protein Description** | **Average Log2 Ratio** | ***p*-value** | **Adjusted *p*-value** |
| --- | --- | --- | --- | --- | --- |
| Q9UDR5 | AASS | Alpha-aminoadipic semialdehyde synthase, mitochondrial (LKR/SDH) [Includes: Lysine ketoglutarate reductase (LKR) (LOR) (EC 1.5.1.8); Saccharopine dehydrogenase (SDH) (EC 1.5.1.9)] | -0,715119746 | 0,00365 | 0.0767928025552317 |
| O43684 | BUB3 | Mitotic checkpoint protein BUB3 | 0,595293436 | 0,00805 | 0.102154997647559 |
| P40123 | CAP2 | Adenylyl cyclase-associated protein 2 (CAP 2) | -0,377385223 | 0,03932 | 0.183376978972405 |
| P51636 | CAV2 | Caveolin-2 | 0,482360111 | 0,03374 | 0.172516669078315 |
| Q96C86 | DCPS | m7GpppX diphosphatase (EC 3.6.1.59) (DCS-1) (Decapping scavenger enzyme) (Hint-related 7meGMP-directed hydrolase) (Histidine triad nucleotide-binding protein 5) (Histidine triad protein member 5) (HINT-5) (Scavenger mRNA-decapping enzyme DcpS) | 0,515388415 | 0,01748 | 0.13109533821508 |
| P35659 | DEK | Protein DEK | 0,392827136 | 0,0383 | 0.180923549477534 |
| O43143 | DHX15 | Pre-mRNA-splicing factor ATP-dependent RNA helicase DHX15 (EC 3.6.4.13) (ATP-dependent RNA helicase #46) (DEAH box protein 15) | 0,51279182 | 0,0202 | 0.138876689931427 |
| P33316 | DUT | Deoxyuridine 5'-triphosphate nucleotidohydrolase, mitochondrial (dUTPase) (EC 3.6.1.23) (dUTP pyrophosphatase) | 0,598980378 | 0,01029 | 0.109336835224251 |
| Q13642 | FHL1 | Four and a half LIM domains protein 1 (FHL-1) (Skeletal muscle LIM-protein 1) (SLIM) (SLIM-1) | -0,595497703 | 0,01189 | 0.112411439550033 |
| Q13451 | FKBP5 | Peptidyl-prolyl cis-trans isomerase FKBP5 (PPIase FKBP5) (EC 5.2.1.8) (51 kDa FK506-binding protein) (51 kDa FKBP) (FKBP-51) (54 kDa progesterone receptor-associated immunophilin) (Androgen-regulated protein 6) (FF1 antigen) (FK506-binding protein 5) (FKBP-5) (FKBP54) (p54) (HSP90-binding immunophilin) (Rotamase) | -0,62991543 | 0,00988 | 0.109011590445672 |
| Q96AE4 | FUBP1 | Far upstream element-binding protein 1 (FBP) (FUSE-binding protein 1) (DNA helicase V) (hDH V) | 0,543785441 | 0,01063 | 0.109626270622104 |
| Q06210 | GFPT1 | Glutamine--fructose-6-phosphate aminotransferase [isomerizing] 1 (EC 2.6.1.16) (D-fructose-6-phosphate amidotransferase 1) (Glutamine:fructose-6-phosphate amidotransferase 1) (GFAT 1) (GFAT1) (Hexosephosphate aminotransferase 1) | 0,606380292 | 0,02376 | 0.147884574150362 |
| P17096 | HMGA1 | High mobility group protein HMG-I/HMG-Y (HMG-I(Y)) (High mobility group AT-hook protein 1) (High mobility group protein A1) (High mobility group protein R) | 0,639390288 | 0,03932 | 0.183376978972405 |
| P09429 | HMGB1 | High mobility group protein B1 (High mobility group protein 1) (HMG-1) | 0,7692113 | 0,02128 | 0.141512996692124 |
| Q12906 | ILF3 | Interleukin enhancer-binding factor 3 (Double-stranded RNA-binding protein 76) (DRBP76) (M-phase phosphoprotein 4) (MPP4) (Nuclear factor associated with dsRNA) (NFAR) (Nuclear factor of activated T-cells 90 kDa) (NF-AT-90) (Translational control protein 80) (TCP80) | 0,424151196 | 0,03135 | 0.166379218606509 |
| Q14739 | LBR | Lamin-B receptor (Integral nuclear envelope inner membrane protein) (LMN2R) | 0,959314807 | 0,00149 | 0.0658126006797819 |
| P20700 | LMNB1 | Lamin-B1 | 0,763876956 | 0,03026 | 0.163712810448877 |
| P62310 | LSM3 | U6 snRNA-associated Sm-like protein LSm3 | 0,677482033 | 0,00642 | 0.0954651715986055 |
| P51884 | LUM | Lumican (Keratan sulfate proteoglycan lumican) (KSPG lumican) | 0,563066279 | 0,01203 | 0.113024663824624 |
| Q9BS40 | LXN | Latexin (Endogenous carboxypeptidase inhibitor) (ECI) (Protein MUM) (Tissue carboxypeptidase inhibitor) (TCI) | 1,005643912 | 0,01179 | 0.112267836107064 |
| P20591 | MX1 | Interferon-induced GTP-binding protein Mx1 (Interferon-induced protein p78) (IFI-78K) (Interferon-regulated resistance GTP-binding protein MxA) (Myxoma resistance protein 1) (Myxovirus resistance protein 1) [Cleaved into: Interferon-induced GTP-binding protein Mx1, N-terminally processed] | -0,979202695 | 0,0427 | 0.188876486048564 |
| O95563 | MPC2 | Mitochondrial pyruvate carrier 2 (Brain protein 44) | -1,222321472 | 0,00094 | 0.0512931662027789 |
| P49959 | MRE11 | Double-strand break repair protein MRE11 (Double-strand break repair protein MRE11A) (Meiotic recombination 11 homolog 1) (MRE11 homolog 1) (Meiotic recombination 11 homolog A) (MRE11 homolog A) | 0,556018075 | 0,01989 | 0.137802712336179 |
| P43246 | MSH2 | DNA mismatch repair protein Msh2 (hMSH2) (MutS protein homolog 2) | 1,072998427 | 0,00414 | 0.0790778462733819 |
| Q8IXM6 | NRM | Nurim (Nuclear envelope membrane protein) (Nuclear rim protein) | 0,633363011 | 0,01141 | 0.111960961329472 |
| O15460 | P4HA2 | Prolyl 4-hydroxylase subunit alpha-2 (4-PH alpha-2) (EC 1.14.11.2) (Procollagen-proline,2-oxoglutarate-4-dioxygenase subunit alpha-2) | 0,715608747 | 0,01318 | 0.117794149768834 |
| P09874 | PARP1 | Poly [ADP-ribose] polymerase 1 (PARP-1) (EC 2.4.2.30) (ADP-ribosyltransferase diphtheria toxin-like 1) (ARTD1) (NAD(+) ADP-ribosyltransferase 1) (ADPRT 1) (Poly[ADP-ribose] synthase 1) | 0,547534342 | 0,03848 | 0.1811476445859 |
| Q15121 | PEA15 | Astrocytic phosphoprotein PEA-15 (15 kDa phosphoprotein enriched in astrocytes) (Phosphoprotein enriched in diabetes) (PED) | 0,396565936 | 0,03651 | 0.177612813348651 |
| Q02809 | PLOD1 | Procollagen-lysine,2-oxoglutarate 5-dioxygenase 1 (EC 1.14.11.4) (Lysyl hydroxylase 1) (LH1) | 0,877732187 | 0,02472 | 0.150084617127734 |
| P60891 | PRPS1 | Ribose-phosphate pyrophosphokinase 1 (EC 2.7.6.1) (PPRibP) (Phosphoribosyl pyrophosphate synthase I) (PRS-I) | 0,545547223 | 0,01909 | 0.135774205040182 |
| P26599 | PTBP1 | Polypyrimidine tract-binding protein 1 (PTB) (57 kDa RNA-binding protein PPTB-1) (Heterogeneous nuclear ribonucleoprotein I) (hnRNP I) | 0,649572181 | 0,01188 | 0.112411439550033 |
| Q9NR77 | PXMP2 | Peroxisomal membrane protein 2 (22 kDa peroxisomal membrane protein) | 0,722974719 | 0,02025 | 0.138876689931427 |
| Q9P0K7 | RAI14 | Ankycorbin (Ankyrin repeat and coiled-coil structure-containing protein) (Novel retinal pigment epithelial cell protein) (Retinoic acid-induced protein 14) | 0,472950142 | 0,01829 | 0.133184461911317 |
| P38159 | RBMX | RNA-binding motif protein, X chromosome (Glycoprotein p43) (Heterogeneous nuclear ribonucleoprotein G) (hnRNP G) [Cleaved into: RNA-binding motif protein, X chromosome, N-terminally processed] | 0,97386887 | 0,00154 | 0.0665699421028979 |
| Q9P258 | RCC2 | Protein RCC2 (RCC1-like protein TD-60) (Telophase disk protein of 60 kDa) | 1,120902198 | 0,00037 | 0.0460405438333303 |
| Q00765 | REEP5 | Receptor expression-enhancing protein 5 (Polyposis locus protein 1) (Protein TB2) | -0,822528333 | 0,00243 | 0.073956300294973 |
| Q01105 | SET | Protein SET (HLA-DR-associated protein II) (Inhibitor of granzyme A-activated DNase) (IGAAD) (PHAPII) (Phosphatase 2A inhibitor I2PP2A) (I-2PP2A) (Template-activating factor I) (TAF-I) | 0,416542918 | 0,04366 | 0.190482327798759 |
| Q15393 | SF3B3 | Splicing factor 3B subunit 3 (Pre-mRNA-splicing factor SF3b 130 kDa subunit) (SF3b130) (STAF130) (Spliceosome-associated protein 130) (SAP 130) | 0,476447141 | 0,02374 | 0.147884574150362 |
| P23246 | SFPQ | Splicing factor, proline- and glutamine-rich (100 kDa DNA-pairing protein) (hPOMp100) (DNA-binding p52/p100 complex, 100 kDa subunit) (Polypyrimidine tract-binding protein-associated-splicing factor) (PSF) (PTB-associated-splicing factor) | 0,517931704 | 0,01746 | 0.13109533821508 |
| Q9UQE7 | SMC3 | Structural maintenance of chromosomes protein 3 (SMC protein 3) (SMC-3) (Basement membrane-associated chondroitin proteoglycan) (Bamacan) (Chondroitin sulfate proteoglycan 6) (Chromosome-associated polypeptide) (hCAP) | 0,349882296 | 0,04718 | 0.198769565715963 |
| Q16629 | SRSF7 | Serine/arginine-rich splicing factor 7 (Splicing factor 9G8) (Splicing factor, arginine/serine-rich 7) | 0,568247729 | 0,04032 | 0.183872362438851 |
| Q08945 | SSRP1 | FACT complex subunit SSRP1 (Chromatin-specific transcription elongation factor 80 kDa subunit) (Facilitates chromatin transcription complex 80 kDa subunit) (FACT 80 kDa subunit) (FACTp80) (Facilitates chromatin transcription complex subunit SSRP1) (Recombination signal sequence recognition protein 1) (Structure-specific recognition protein 1) (hSSRP1) (T160) | 0,588415314 | 0,01753 | 0.131227758234791 |
| Q86Y82 | STX12 | Syntaxin-12 | 0,468116109 | 0,01829 | 0.133184461911317 |
| P07951 | TPM2 | Tropomyosin beta chain (Beta-tropomyosin) (Tropomyosin-2) | -0,610559414 | 0,03578 | 0.176070195019166 |
| P62995 | TRA2B | Transformer-2 protein homolog beta (TRA-2 beta) (TRA2-beta) (hTRA2-beta) (Splicing factor, arginine/serine-rich 10) (Transformer-2 protein homolog B) | 0,775951241 | 0,01603 | 0.12714580081316 |
| Q9Y5S1 | TRPV2 | Transient receptor potential cation channel subfamily V member 2 (TrpV2) (Osm-9-like TRP channel 2) (OTRPC2) (Vanilloid receptor-like protein 1) (VRL-1) | 0,628519757 | 0,0066 | 0.0958235018254441 |
| Q9H3M7 | TXNIP | Thioredoxin-interacting protein (Thioredoxin-binding protein 2) (Vitamin D3 up-regulated protein 1) | -0,586161819 | 0,04182 | 0.187870082281016 |
| P11172 | UMPS | Uridine 5'-monophosphate synthase (UMP synthase) [Includes: Orotate phosphoribosyltransferase (OPRT) (OPRTase) (EC 2.4.2.10); Orotidine 5'-phosphate decarboxylase (ODC) (EC 4.1.1.23) (OMPdecase)] | 0,745003579 | 0,006 | 0.0948090470651261 |
| Q9H7D7 | WDR26 | WD repeat-containing protein 26 (CUL4- and DDB1-associated WDR protein 2) (Myocardial ischemic preconditioning up-regulated protein 2) | 0,690796828 | 0,00384 | 0.0771477016366249 |

**Supporting table 11: List of all drug concentrations used for the short-term and long-term cytotoxicity tests in ISO- and HGPS-iPSC-CMs.**

| **Compounds** | **Tested Concentrations**  **(48h)** | **Tested Concentrations (10 days)** | **Most effective Acute concentrations used in literature** | **Cell types** | **Treatment**  **duration** | **Decrease of progerin**  **?** | **Ref.** |
| --- | --- | --- | --- | --- | --- | --- | --- |
| Rapa | 1 nM 4 nM | 1 nM 4 nM | 10 nM | Rat CMs | 24h | N.A. | [8] |
| Lona | 2 µM 8 µM | 2 µM 8 µM | 20 μM | Human iPSC SMCs | 48h | N.T. | [9] |
| MG-132 | 500 nM  2 µM | 125 nM  250 nM | 5 μM | HGPS FBs | 24h and 48h | Yes | [10] |
| Fisetin | 2 µM 20 µM | 2 µM 10 µM | 50 μM | Senescent mouse and human fibroblasts | 48h | N.A. | [11] |
| D+Q | 8 nM + 1.6 µM  32 nM + 6.4 µM | 8 nM + 1.6 µM  16 nM + 3.2 µM | 80 nM + 16 μM | Senescent human fat cell progenitors, human ECs, mouse BM-MSCs, MEFs | 48h | N.A. | [12] |
| NPY | 10 nM  40 nM | 10 nM  40 nM | 100 nΜ | Human HGPS FBs | 6h  1 week | Yes | [5] |

CM – cardiomyocyte; N.A. – not applicable; N.T. – not tested

**Supplementary Data Table 12 List of relevant antioxidant proteins with altered expression upon treating HGPS-iPSC-CMs with Lona**

| **Uniprot ID** | **Protein Symbol** | **Protein Description** | **Average Log2 Ratio** | ***p*-value** | **Adjusted *p*-value** |
| --- | --- | --- | --- | --- | --- |
| P04179 | SOD2 | Superoxide dismutase [Mn], mitochondrial | 0.56222036 | 3.34E-15 | 1.2741635E-13 |
| P22352 | GPX3 | Glutathione peroxidase 3 | 0.47674486 | 0.607205 | 0.3831745 |
| P36969 | GPX4 | Phospholipid hydroperoxide glutathione peroxidase, mitochondrial | 0.25801198 | 0.0073357 | 0.0177590 |
| Q96SL4 | GPX7 | Glutathione peroxidase 7 | 0.22463707 | 0.0731816 | 0.09933406 |

**Supplementary Data Table 13. List of total significantly altered proteins (p-value<0.05) in Lona-treated LAKI mice, when compared with Untreated LAKI controls.**

| **Uniprot ID** | **Protein Symbol** | **Protein Description** | **Average Log2 Ratio** | ***p*-value** | **Adjusted *p*-value** |
| --- | --- | --- | --- | --- | --- |
| A3KFX0 | 5NT1A | Cytosolic 5'-nucleotidase 1A | 0,136070656 | 0,025368 | 0,815053346 |
| Q6P542 | ABCF1 | ATP-binding cassette sub-family F member 1 | -0,113156739 | 0,013002 | 0,687122582 |
| Q8BL65 | ABLM2 | Actin-binding LIM protein 2 | -0,076522742 | 0,013222 | 0,687122582 |
| Q8VCW8 | ACSF2 | Acyl-CoA synthetase family member 2, mitochondrial | 0,268306435 | 0,036757 | 0,839048388 |
| P00329 | ADH1 | Alcohol dehydrogenase 1 | -0,412575975 | 2,43E-05 | 0,026052545 |
| P47740 | AL3A2 | Fatty aldehyde dehydrogenase | -0,088081254 | 0,01883 | 0,739546641 |
| P06728 | APOA4 | Apolipoprotein A-IV | -0,139385279 | 0,010826 | 0,633030862 |
| P08226 | APOE | Apolipoprotein E | -0,299471184 | 0,002054 | 0,337384746 |
| Q8K4I3 | ARHG6 | Rho guanine nucleotide exchange factor 6 | 0,152441265 | 0,023116 | 0,774263036 |
| A6H630 | ARMT1 | Protein-glutamate O-methyltransferase | -0,199691314 | 7,81E-05 | 0,044919352 |
| P98203 | ARVC | Armadillo repeat protein deleted in velo-cardio-facial syndrome homolog | -0,214379064 | 0,0008 | 0,211561006 |
| Q9EPB4 | ASC | Apoptosis-associated speck-like protein containing a CARD | -0,208095192 | 0,022056 | 0,76860298 |
| Q04519 | ASM | Sphingomyelin phosphodiesterase | -0,200496039 | 0,0217 | 0,763918354 |
| Q9EPE9 | AT131 | Manganese-transporting ATPase 13A1 | -0,209980975 | 0,038091 | 0,850221017 |
| O54962 | BAF | Barrier-to-autointegration factor | -0,282305019 | 0,038198 | 0,850221017 |
| Q60739 | BAG1 | BAG family molecular chaperone regulator 1 | -0,048066839 | 0,009932 | 0,622754493 |
| Q91YN9 | BAG2 | BAG family molecular chaperone regulator 2 | -0,005911084 | 0,025515 | 0,815053346 |
| Q80XN0 | BDH | D-beta-hydroxybutyrate dehydrogenase, mitochondrial | 0,281857901 | 0,01127 | 0,637391741 |
| P82198 | BGH3 | Transforming growth factor-beta-induced protein ig-h3 | -0,226220819 | 0,017702 | 0,739546641 |
| Q64152 | BTF3 | Transcription factor BTF3 | -0,066864165 | 0,002828 | 0,394149379 |
| Q8BIF0 | C99L2 | CD99 antigen-like protein 2 | -0,195497179 | 0,026773 | 0,819215298 |
| P83917 | CBX1 | Chromobox protein homolog 1 | -0,172007317 | 0,048258 | 0,923571273 |
| Q61147 | CERU | Ceruloplasmin | -0,22476408 | 0,020255 | 0,741287169 |
| P03953 | CFAD | Complement factor D | -0,259196045 | 0,000247 | 0,114165097 |
| P06909 | CFAH | Complement factor H | -0,12726177 | 0,029841 | 0,819215298 |
| Q9D1P4 | CHRD1 | Cysteine and histidine-rich domain-containing protein 1 | -0,745218881 | 0,015586 | 0,693519632 |
| Q9CQE8 | CN166 | UPF0568 protein C14orf166 homolog | -0,068840439 | 0,002856 | 0,394149379 |
| P06684 | CO5 | Complement C5 | 0,265600755 | 0,000884 | 0,211561006 |
| Q8C6K9 | CO6A6 | Collagen alpha-6(VI) chain | -0,127909044 | 0,012068 | 0,650543919 |
| Q8K0C8 | COX19 | Cytochrome c oxidase assembly protein COX19 | -0,29212698 | 0,003572 | 0,456376156 |
| Q8VHY0 | CSPG4 | Chondroitin sulfate proteoglycan 4 | -0,036705489 | 0,033368 | 0,835940968 |
| P26231 | CTNA1 | Catenin alpha-1 | -0,09926097 | 0,021326 | 0,758490128 |
| Q9WTX6 | CUL1 | Cullin-1 | -0,187992458 | 0,030172 | 0,819215298 |
| Q9CXW3 | CYBP | Calcyclin-binding protein | -0,206033967 | 0,01115 | 0,637391741 |
| Q9Z0Y1 | DCTN3 | Dynactin subunit 3 | -0,137268456 | 0,007364 | 0,599413668 |
| E9Q557 | DESP | Desmoplakin | -0,056523111 | 0,036581 | 0,839048388 |
| Q9QYI4 | DJB12 | DnaJ homolog subfamily B member 12 | -0,384915596 | 0,048236 | 0,923571273 |
| P63037 | DNJA1 | DnaJ homolog subfamily A member 1 | -0,188652882 | 0,025211 | 0,815053346 |
| O55111 | DSG2 | Desmoglein-2 | -0,132254902 | 0,019659 | 0,741287169 |
| Q8BJW6 | EIF2A | Eukaryotic translation initiation factor 2A | -0,095098431 | 0,043275 | 0,885856051 |
| Q9EP72 | EMC7 | ER membrane protein complex subunit 7 | -0,072447469 | 0,027156 | 0,819215298 |
| P48193 | EPB41 | Protein 4.1 | -0,162363765 | 0,018635 | 0,739546641 |
| P49222 | EPB42 | Erythrocyte membrane protein band 4.2 | -0,140618042 | 0,0056 | 0,53664797 |
| Q9DC16 | ERGI1 | Endoplasmic reticulum-Golgi intermediate compartment protein 1 | -1,744838209 | 0,042352 | 0,885537151 |
| Q922E6 | FAKD2 | FAST kinase domain-containing protein 2 | 0,114887468 | 0,022733 | 0,76890649 |
| Q61554 | FBN1 | Fibrillin-1 | -0,17183109 | 0,002412 | 0,378253577 |
| E9PV24 | FIBA | Fibrinogen alpha chain | 0,201707153 | 0,041564 | 0,879733084 |
| Q8K0E8 | FIBB | Fibrinogen beta chain | 0,231723575 | 0,045209 | 0,914033092 |
| Q8VCM7 | FIBG | Fibrinogen gamma chain | 0,215969723 | 0,000978 | 0,211561006 |
| P11276 | FINC | Fibronectin | 0,203684916 | 0,000981 | 0,211561006 |
| Q9JJ28 | FLII | Protein flightless-1 homolog | -0,153507083 | 0,020476 | 0,741287169 |
| Q9CS72 | FLIP1 | Filamin-A-interacting protein 1 | -0,090305473 | 0,015557 | 0,693519632 |
| Q8K2I3 | FMO2 | Dimethylaniline monooxygenase [N-oxide-forming] 2 | -0,286126781 | 0,049008 | 0,923571273 |
| P09528 | FRIH | Ferritin heavy chain | -0,408289162 | 0,015356 | 0,693519632 |
| Q3U0V1 | FUBP2 | Far upstream element-binding protein 2 | -0,086099216 | 0,04895 | 0,923571273 |
| Q3TQB2 | FXRD1 | FAD-dependent oxidoreductase domain-containing protein 1 | 0,19165909 | 0,011949 | 0,650543919 |
| Q9D6M3 | GHC1 | Mitochondrial glutamate carrier 1 | 0,102689592 | 0,039967 | 0,863507749 |
| Q922P9 | GLYR1 | Putative oxidoreductase GLYR1 | -0,119933414 | 0,008179 | 0,599413668 |
| Q9CRC9 | GNPI2 | Glucosamine-6-phosphate isomerase 2 | -0,050485688 | 0,033625 | 0,835940968 |
| Q8BH60 | GOPC | Golgi-associated PDZ and coiled-coil motif-containing protein | 1,775256526 | 0,000344 | 0,131681665 |
| Q8C0L9 | GPCP1 | Glycerophosphocholine phosphodiesterase GPCPD1 | 0,07182072 | 0,001695 | 0,320112 |
| P11352 | GPX1 | Glutathione peroxidase 1 | 0,493188182 | 6,25E-07 | 0,002155468 |
| Q8CCK0 | H2AW | Core histone macro-H2A.2 | -0,238914319 | 0,034208 | 0,835940968 |
| P0C0S6;Q3THW5 | H2AZ;H2AV | Histone H2A.Z;Histone H2A.V | -0,216081374 | 0,03802 | 0,850221017 |
| Q8VC19 | HEM1 | 5-aminolevulinate synthase, nonspecific, mitochondrial | -3,241716701 | 0,035014 | 0,838882162 |
| Q6PAV2 | HERC4 | Probable E3 ubiquitin-protein ligase HERC4 | 0,226973688 | 0,010357 | 0,622754493 |
| Q8VD75 | HIP1 | Huntingtin-interacting protein 1 | 0,310788639 | 0,046539 | 0,920851405 |
| P70333 | HNRH2 | Heterogeneous nuclear ribonucleoprotein H2 | -0,053481361 | 0,039719 | 0,863507749 |
| Q9ESB3 | HRG | Histidine-rich glycoprotein | 0,136339627 | 0,048258 | 0,923571273 |
| Q9CZJ2 | HS12B | Heat shock 70 kDa protein 12B | -0,123261377 | 0,028393 | 0,819215298 |
| P51125 | ICAL | Calpastatin | -0,004914849 | 0,019293 | 0,739546641 |
| P35330 | ICAM2 | Intercellular adhesion molecule 2 | -0,106062139 | 0,009245 | 0,613394518 |
| Q9JHR7 | IDE | Insulin-degrading enzyme | -0,361331237 | 0,000682 | 0,211561006 |
| P60843 | IF4A1 | Eukaryotic initiation factor 4A-I | -0,030505578 | 0,013777 | 0,687122582 |
| P59325 | IF5 | Eukaryotic translation initiation factor 5 | -0,036127825 | 0,029352 | 0,819215298 |
| P01867 | IGG2B | Ig gamma-2B chain C region | 0,355462295 | 0,010469 | 0,622754493 |
| P01878 | IGHA | Ig alpha chain C region | -0,558429093 | 0,010202 | 0,622754493 |
| Q8R0F6 | ILKAP | Integrin-linked kinase-associated serine/threonine phosphatase 2C | -0,461137502 | 0,001962 | 0,337384746 |
| Q9JHU9 | INO1 | Inositol-3-phosphate synthase 1 | -0,214738913 | 0,008861 | 0,599413668 |
| P63248 | IPKA | cAMP-dependent protein kinase inhibitor alpha | -0,19381533 | 0,049033 | 0,923571273 |
| O70139 | IPKG | cAMP-dependent protein kinase inhibitor gamma | -0,201342394 | 0,03113 | 0,82000378 |
| Q9JHP7 | KDEL1 | KDEL motif-containing protein 1 | -0,235733755 | 0,036064 | 0,839048388 |
| Q60675 | LAMA2 | Laminin subunit alpha-2 | -0,064607611 | 0,022656 | 0,76890649 |
| P02469 | LAMB1 | Laminin subunit beta-1 | -0,09960076 | 0,013496 | 0,687122582 |
| P02468 | LAMC1 | Laminin subunit gamma-1 | -0,079407717 | 0,032508 | 0,835940968 |
| Q3U9G9 | LBR | Lamin-B receptor | -0,102996174 | 0,014181 | 0,687122582 |
| Q5SUF2 | LC7L3 | Luc7-like protein 3 | -0,150142068 | 0,031136 | 0,82000378 |
| Q99JW4 | LIMS1 | LIM and senescent cell antigen-like-containing domain protein 1 | -0,148753216 | 0,027954 | 0,819215298 |
| Q8VCM4 | LIPT | Lipoyltransferase 1, mitochondrial | -0,128453841 | 0,030459 | 0,819215298 |
| Q9D009 | LIPT2 | Putative lipoyltransferase 2, mitochondrial | 0,423842157 | 0,024231 | 0,801950884 |
| P48678 | LMNA | Prelamin-A/C | -0,12822893 | 0,043017 | 0,885856051 |
| P14733 | LMNB1 | Lamin-B1 | -0,109849964 | 0,033147 | 0,835940968 |
| P21619 | LMNB2 | Lamin-B2 | -0,211346555 | 0,01907 | 0,739546641 |
| O35900 | LSM2 | U6 snRNA-associated Sm-like protein LSm2 | -0,100724975 | 0,030364 | 0,819215298 |
| Q91V16 | LYRM5 | LYR motif-containing protein 5 | -0,14663621 | 0,014078 | 0,687122582 |
| Q8K2I4 | MANBA | Beta-mannosidase | -0,338529905 | 0,034024 | 0,835940968 |
| Q3KNM2 | MARH5 | E3 ubiquitin-protein ligase MARCH5 | 0,108129348 | 0,01434 | 0,687122582 |
| Q8R3C0 | MCMBP | Mini-chromosome maintenance complex-binding protein | -0,571984168 | 0,020627 | 0,741287169 |
| Q9Z2D6 | MECP2 | Methyl-CpG-binding protein 2 | -0,158082425 | 0,049525 | 0,923571273 |
| Q8BI84 | MIA3 | Melanoma inhibitory activity protein 3 | -0,182924005 | 0,016109 | 0,693519632 |
| Q9QVP4 | MLRA | Myosin regulatory light chain 2, atrial isoform | -1,44892047 | 0,005039 | 0,514788593 |
| Q07113 | MPRI | Cation-independent mannose-6-phosphate receptor | -0,146648677 | 0,032382 | 0,835940968 |
| Q8BKY8 | MTEF2 | Transcription termination factor 2, mitochondrial | 0,214763688 | 0,000857 | 0,211561006 |
| Q8VCM5 | MUL1 | Mitochondrial ubiquitin ligase activator of NFKB 1 | -0,227257704 | 0,018128 | 0,739546641 |
| Q91Z83 | MYH7 | Myosin-7 | -0,760929394 | 3,03E-05 | 0,026052545 |
| P09541 | MYL4 | Myosin light chain 4 | -1,645105487 | 0,049283 | 0,923571273 |
| Q6PGB6 | NAA50 | N-alpha-acetyltransferase 50 | -0,029507467 | 0,016283 | 0,693519632 |
| TRYP_PIG | NaN | Trypsin - Sus scrofa (Pig). | -0,453776196 | 0,015576 | 0,693519632 |
| O70310 | NMT1 | Glycylpeptide N-tetradecanoyltransferase 1 | -0,028280114 | 0,047594 | 0,923571273 |
| Q99ME9 | NOG1 | Nucleolar GTP-binding protein 1 | -0,195475732 | 0,016242 | 0,693519632 |
| P60670 | NPL4 | Nuclear protein localization protein 4 homolog | -0,040838134 | 0,017894 | 0,739546641 |
| Q80XB4 | NRAP | Nebulin-related-anchoring protein | -0,140135761 | 0,000265 | 0,114165097 |
| Q8JZU0 | NUD13 | Nucleoside diphosphate-linked moiety X motif 13 | 0,344386644 | 3,78E-05 | 0,026052545 |
| Q61206 | PA1B2 | Platelet-activating factor acetylhydrolase IB subunit beta | -0,16829193 | 0,018526 | 0,739546641 |
| O35386 | PAHX | Phytanoyl-CoA dioxygenase, peroxisomal | 0,196626215 | 0,008854 | 0,599413668 |
| P11103 | PARP1 | Poly [ADP-ribose] polymerase 1 | -0,122698313 | 0,028505 | 0,819215298 |
| P48725 | PCNT | Pericentrin | 0,271817321 | 0,028597 | 0,819215298 |
| Q8BG81 | PDIP3 | Polymerase delta-interacting protein 3 | -0,505468312 | 0,028576 | 0,819215298 |
| P15331 | PERI | Peripherin | -0,239806882 | 0,007609 | 0,599413668 |
| Q68FH0 | PKP4 | Plakophilin-4 | -0,149059091 | 0,002809 | 0,394149379 |
| Q9D3P8 | PLRKT | Plasminogen receptor (KT) | 0,281433483 | 0,01914 | 0,739546641 |
| Q62009 | POSTN | Periostin | 0,373264603 | 0,041904 | 0,881512077 |
| Q9CR16 | PPID | Peptidyl-prolyl cis-trans isomerase D | -0,110154654 | 0,028683 | 0,819215298 |
| Q99MR9 | PPR3A | Protein phosphatase 1 regulatory subunit 3A | -0,099458339 | 0,040297 | 0,863507749 |
| Q9Z0S9 | PRAF1 | Prenylated Rab acceptor protein 1 | -0,32464305 | 0,003998 | 0,492590026 |
| O35593 | PSDE | 26S proteasome non-ATPase regulatory subunit 14 | -0,009561707 | 0,026373 | 0,819215298 |
| Q9CX56 | PSMD8 | 26S proteasome non-ATPase regulatory subunit 8 | -0,104630218 | 0,034778 | 0,838882162 |
| P70698 | PYRG1 | CTP synthase 1 | -0,153332946 | 0,004438 | 0,497797202 |
| Q8CEE7 | RDH13 | Retinol dehydrogenase 13 | 0,049115525 | 0,00979 | 0,622754493 |
| Q9D7N6 | RM30 | 39S ribosomal protein L30, mitochondrial | -0,098032981 | 0,001763 | 0,320112 |
| Q9CZ83 | RM55 | 39S ribosomal protein L55, mitochondrial | 0,268854359 | 0,004473 | 0,497797202 |
| P70336 | ROCK2 | Rho-associated protein kinase 2 | 0,032736952 | 0,042739 | 0,885856051 |
| P62858 | RS28 | 40S ribosomal protein S28 | 0,006906739 | 0,008196 | 0,599413668 |
| Q9CQV5 | RT24 | 28S ribosomal protein S24, mitochondrial | -0,117587874 | 0,039315 | 0,863507749 |
| P55012 | S12A2 | Solute carrier family 12 member 2 | 0,184973388 | 0,040552 | 0,863600186 |
| Q9DCP2 | S38A3 | Sodium-coupled neutral amino acid transporter 3 | -0,176234321 | 0,034407 | 0,835940968 |
| Q8VCA8 | SCRN2 | Secernin-2 | -0,115746226 | 0,045304 | 0,914033092 |
| Q8BJU0 | SGTA | Small glutamine-rich tetratricopeptide repeat-containing protein alpha | -0,134144439 | 0,043394 | 0,885856051 |
| Q6P549 | SHIP2 | Phosphatidylinositol 3,4,5-trisphosphate 5-phosphatase 2 | 0,137731902 | 0,005073 | 0,514788593 |
| Q91ZW3 | SMCA5 | SWI/SNF-related matrix-associated actin-dependent regulator of chromatin subfamily A member 5 | -0,144650765 | 0,008694 | 0,599413668 |
| P62315 | SMD1 | Small nuclear ribonucleoprotein Sm D1 | -0,198896287 | 0,009583 | 0,622754493 |
| Q6PDG5 | SMRC2 | SWI/SNF complex subunit SMARCC2 | -0,205954614 | 0,048447 | 0,923571273 |
| O09164 | SODE | Extracellular superoxide dismutase [Cu-Zn] | -0,108921732 | 0,036852 | 0,839048388 |
| Q9QX47 | SON | Protein SON | -0,327991973 | 0,046348 | 0,920851405 |
| Q03734 | SPA3M | Serine protease inhibitor A3M | 0,208645287 | 0,036942 | 0,839048388 |
| Q3ULF4 | SPG7 | Paraplegin | 0,107369562 | 0,006906 | 0,581132057 |
| P97364 | SPS2 | Selenide, water dikinase 2 | 0,340171706 | 0,029223 | 0,819215298 |
| P08032 | SPTA1 | Spectrin alpha chain, erythrocytic 1 | -0,114552652 | 0,006252 | 0,541857659 |
| P15508 | SPTB1 | Spectrin beta chain, erythrocytic | -0,062351833 | 0,040125 | 0,863507749 |
| Q8R1R3 | STAR7 | StAR-related lipid transfer protein 7, mitochondrial | -0,02352597 | 0,04671 | 0,920851405 |
| P54227 | STMN1 | Stathmin | -0,373305119 | 0,007971 | 0,599413668 |
| Q8K4T3 | STRAB | STE20-related kinase adapter protein beta | 0,82789726 | 0,00612 | 0,541857659 |
| P58404 | STRN4 | Striatin-4 | -0,103756449 | 0,03543 | 0,839048388 |
| Q9ER00 | STX12 | Syntaxin-12 | -0,064702371 | 0,004638 | 0,499983833 |
| O70439 | STX7 | Syntaxin-7 | -0,264634089 | 0,001275 | 0,258741822 |
| Q9D8Y1 | T126A | Transmembrane protein 126A | 0,138233501 | 0,022471 | 0,76890649 |
| Q9D1R1 | T126B | Complex I assembly factor TMEM126B, mitochondrial | 0,11366506 | 0,016 | 0,693519632 |
| P37804 | TAGL | Transgelin | 0,301869187 | 0,004459 | 0,497797202 |
| P48428 | TBCA | Tubulin-specific chaperone A | -0,058677659 | 0,030632 | 0,819215298 |
| Q3TL26 | TFB2M | Dimethyladenosine transferase 2, mitochondrial | 0,336739887 | 0,000667 | 0,211561006 |
| Q62351 | TFR1 | Transferrin receptor protein 1 | 0,140307268 | 0,024407 | 0,801950884 |
| P61939 | THBG | Thyroxine-binding globulin | -0,369959939 | 0,003378 | 0,448179287 |
| Q99J36 | THUM1 | THUMP domain-containing protein 1 | -0,122868564 | 0,032175 | 0,835940968 |
| Q9CQV1 | TIM16 | Mitochondrial import inner membrane translocase subunit TIM16 | -0,124534814 | 0,026773 | 0,819215298 |
| Q71LX4 | TLN2 | Talin-2 | -0,100176595 | 0,0288 | 0,819215298 |
| Q9R0Q3 | TMED2 | Transmembrane emp24 domain-containing protein 2 | -0,16963504 | 0,046646 | 0,920851405 |
| Q8CGB6 | TNS2 | Tensin-2 | -0,248630565 | 0,005331 | 0,525529664 |
| Q62393 | TPD52 | Tumor protein D52 | -0,142692503 | 0,03941 | 0,863507749 |
| Q9CRB6 | TPPP3 | Tubulin polymerization-promoting protein family member 3 | -0,319184324 | 0,026546 | 0,819215298 |
| Q9ERP3 | TRI54 | Tripartite motif-containing protein 54 | -0,060923046 | 0,00876 | 0,599413668 |
| Q9D8Z2 | TRIA1 | TP53-regulated inhibitor of apoptosis 1 | 0,104606246 | 0,036967 | 0,839048388 |
| Q91W90 | TXND5 | Thioredoxin domain-containing protein 5 | 0,086711762 | 0,020358 | 0,741287169 |
| Q6P4T2 | U520 | U5 small nuclear ribonucleoprotein 200 kDa helicase | -0,117576038 | 0,028592 | 0,819215298 |
| O08810 | U5S1 | 116 kDa U5 small nuclear ribonucleoprotein component | -0,133921632 | 0,034075 | 0,835940968 |
| O08759 | UBE3A | Ubiquitin-protein ligase E3A | -0,038444049 | 0,008783 | 0,599413668 |
| P35123 | UBP4 | Ubiquitin carboxyl-terminal hydrolase 4 | -0,340968331 | 0,014308 | 0,687122582 |
| Q922Y1 | UBXN1 | UBX domain-containing protein 1 | -0,161912454 | 0,006282 | 0,541857659 |
| Q99PL6 | UBXN6 | UBX domain-containing protein 6 | 0,090182712 | 0,030598 | 0,819215298 |
| Q9EQH3 | VPS35 | Vacuolar protein sorting-associated protein 35 | -0,127555266 | 0,015034 | 0,693519632 |
| P97390 | VPS45 | Vacuolar protein sorting-associated protein 45 | 0,170909074 | 0,019834 | 0,741287169 |
| Q9ERF3 | WDR61 | WD repeat-containing protein 61 | -0,120291857 | 0,011679 | 0,649888728 |
| O70373 | XIRP1 | Xin actin-binding repeat-containing protein 1 | -0,058535212 | 0,036027 | 0,839048388 |
| Q9JMD0 | ZN207 | BUB3-interacting and GLEBS motif-containing protein ZNF207 | -0,289234353 | 0,005911 | 0,541857659 |
| P39447 | ZO1 | Tight junction protein ZO-1 | -0,126424402 | 0,034325 | 0,835940968 |

**Supplementary Data Table 14. List of significantly altered senescence-related proteins (p-value<0.05) in Lona-treated LAKI mice, when compared with Untreated LAKI controls**

| **Uniprot ID** | **Protein Symbol** | **Protein Description** | **Average Log2 Ratio** | ***p*-value** | **Adjusted *p*-value** |
| --- | --- | --- | --- | --- | --- |
| Q9EPB4 | ASC | Apoptosis-associated speck-like protein containing a CARD | -0,208095192 | 0,022056 | 0,76860298 |
| Q04519 | ASM | Sphingomyelin phosphodiesterase | -0,200496039 | 0,0217 | 0,763918354 |
| O54962 | BAF | Barrier-to-autointegration factor | -0,282305019 | 0,038198 | 0,850221017 |
| Q9EP72 | EMC7 | ER membrane protein complex subunit 7 | -0,072447469 | 0,027156 | 0,819215298 |
| P11276 | FINC | Fibronectin | 0,203684916 | 0,000981 | 0,211561006 |
| P09528 | FRIH | Ferritin heavy chain | -0,408289162 | 0,015356 | 0,693519632 |
| Q3U0V1 | FUBP2 | Far upstream element-binding protein 2 | -0,086099216 | 0,04895 | 0,923571273 |
| P02468 | LAMC1 | Laminin subunit gamma-1 | -0,079407717 | 0,032508 | 0,835940968 |
| Q3U9G9 | LBR | Lamin-B receptor | -0,102996174 | 0,014181 | 0,687122582 |
| P14733 | LMNB1 | Lamin-B1 | -0,109849964 | 0,033147 | 0,835940968 |
| P21619 | LMNB2 | Lamin-B2 | -0,211346555 | 0,01907 | 0,739546641 |
| O35900 | LSM2 | U6 snRNA-associated Sm-like protein LSm2 | -0,100724975 | 0,030364 | 0,819215298 |
| Q8K2I4 | MANBA | Beta-mannosidase | -0,338529905 | 0,034024 | 0,835940968 |
| Q8R3C0 | MCMBP | Mini-chromosome maintenance complex-binding protein | -0,571984168 | 0,020627 | 0,741287169 |
| Q07113 | MPRI | Cation-independent mannose-6-phosphate receptor | -0,146648677 | 0,032382 | 0,835940968 |
| P11103 | PARP1 | Poly [ADP-ribose] polymerase 1 | -0,122698313 | 0,028505 | 0,819215298 |
| P48725 | PCNT | Pericentrin | 0,271817321 | 0,028597 | 0,819215298 |
| Q62009 | POSTN | Periostin | 0,373264603 | 0,041904 | 0,881512077 |
| P62315 | SMD1 | Small nuclear ribonucleoprotein Sm D1 | -0,198896287 | 0,009583 | 0,622754493 |
| P54227 | STMN1 | Stathmin | -0,373305119 | 0,007971 | 0,599413668 |
| Q9ER00 | STX12 | Syntaxin-12 | -0,064702371 | 0,004638 | 0,499983833 |

**Supplementary Table 15. Specific sets of primers and Probe (FAM) for Progerin using Taqman technology.**

| **Gene** | **Forward primer sequence**  **(5′-3′)** | **Reverse primer sequence**  **(5′-3′)** |
| --- | --- | --- |
| *EDN3* | GTCAGCAGTAATTCAAGGAC | CTTGACTTCAACCTCCTTTG |
| *GAPDH* | AGCCACATCGCTCAGACACC | GTACTCAGCGCCAGCATCG |
| *GATA4* | TCCCTCTTCCCTCCTCAAAT | TCAGCGTGTAAAGGCATCTG |
| *GDF15* | CGAAGACTCCAGATTCCG | ACTTCTGGCGTGAGTATC |
| *LMNA* | TCAGGAGCCCAGGTGGGC | TGACCAGATTGTCCCCGAAGC |
| *MYL2* | AGGCTGATTACGTTCGGGAA | TCTCTTCTCCGTGGGTGATG |
| *TBX5* | GCTGGAAGGCGGATGTTT | GATCGTCGGCAGGTACAATG |
| *TBXT* | TGCTTCCCTGAGACCCAGTT | GATCACTTCTTTCCTTTGCATCAAG |
| *TGFB2* | AGATTTGCAGGTATTGATGG | ATTTCTAAAGCAATAGGCCG |
| *TNNT2* | TCCAGAAGACAGAGCGGAAA | CTTCATTCAGGTGGTCAATGG |

**Supplementary Table 16. Specific sets of primers and Probe (FAM) for Progerin using Taqman technology.**

| **Gene** | **Forward primer sequence**  **(5′-3′)** | **Reverse primer sequence**  **(5′-3′)** | **Probe sequence**  **(5′-3′)** |
| --- | --- | --- | --- |
| Progerin | TGAGTACAACCTGCGCTCAC | TGGCAGGTCCCAGATTACAT | CGGGAGCCCAGAGCTCCCAGAA |

**References**

1.	Nissan, X., et al., *Unique preservation of neural cells in Hutchinson- Gilford progeria syndrome is due to the expression of the neural-specific miR-9 microRNA.* Cell Rep, 2012. **2**(1): p. 1-9.

2.	Hummon, A.B., et al., *Isolation and solubilization of proteins after TRIzol extraction of RNA and DNA from patient material following prolonged storage.* Biotechniques, 2007. **42**(4): p. 467-70, 472.

3.	Holden, P. and W.A. Horton, *Crude subcellular fractionation of cultured mammalian cell lines.* BMC Res Notes, 2009. **2**: p. 243.

4.	Ferreira, L.L., et al., *Doxorubicin persistently rewires cardiac circadian homeostasis in mice.* Arch Toxicol, 2020. **94**(1): p. 257-271.

5.	Aveleira, C.A., et al., *Neuropeptide Y Enhances Progerin Clearance and Ameliorates the Senescent Phenotype of Human Hutchinson-Gilford Progeria Syndrome Cells.* J Gerontol A Biol Sci Med Sci, 2020. **75**(6): p. 1073-1078.

6.	Santinha, D., et al., *Remodeling of the Cardiac Extracellular Matrix Proteome During Chronological and Pathological Aging.* Mol Cell Proteomics, 2024. **23**(1): p. 100706.

7.	Perez-Riverol, Y., et al., *The PRIDE database resources in 2022: a hub for mass spectrometry-based proteomics evidences.* Nucleic Acids Res, 2022. **50**(D1): p. D543-D552.

8.	Gao, G., et al., *Rapamycin regulates the balance between cardiomyocyte apoptosis and autophagy in chronic heart failure by inhibiting mTOR signaling.* Int J Mol Med, 2020. **45**(1): p. 195-209.

9.	Pitrez, P.R., et al., *Vulnerability of progeroid smooth muscle cells to biomechanical forces is mediated by MMP13.* Nat Commun, 2020. **11**(1): p. 4110.

10.	Harhouri, K., et al., *MG132-induced progerin clearance is mediated by autophagy activation and splicing regulation.* EMBO Mol Med, 2017. **9**(9): p. 1294-1313.

11.	Yousefzadeh, M.J., et al., *Fisetin is a senotherapeutic that extends health and lifespan.* EBioMedicine, 2018. **36**: p. 18-28.

12.	Zhu, Y., et al., *The Achilles' heel of senescent cells: from transcriptome to senolytic drugs.* Aging Cell, 2015. **14**(4): p. 644-58.
